# Transcriptomic insights into the tritrophic plant–pathogen–mycoparasite interaction reveal coordinated reprogramming fungal secretomes and plant amino acid metabolism

**DOI:** 10.1101/2025.11.24.690098

**Authors:** Kazuya Maeda, Mariko Kouda, Mai Ohara, Takumi Kawase, Koki Saito, Eishin Iwao, Hirotoshi Sushida, Tomoko Suzuki, Takuya Sumita, Yuichiro Iida

**Author notes:** Correspondence to Y. Iida.

## Abstract

- The tomato–*Cladosporium fulvum* (syn. *Fulvia fulva*) pathosystem has served as a model for the gene-for-gene concept of effectors and resistance proteins, but this binary framework does not include the potential influence of other microbial participants. Here we describe the dramatic changes in gene expression of all members of the tritrophic interaction among tomato, *C. fulvum*, and mycoparasitic fungus *Hansfordia pulvinata*.
- Transcriptomic analyses of the mycoparasite *H. pulvinata* during parasitism of *C. fulvum* on tomato *on planta* and *in vitro* revealed a dramatic upregulation of genes encoding small secreted proteins during mycoparasitism, notably, a Nep1-like protein (HpNlp1) lacked typical necrosis-inducing activity but induced the accumulation of antifungal compounds inhibiting spore germination of *C. fulvum*.
- Similarly, in *C. fulvum* parasitized by *H. pulvinata*, effector genes were highly expressed. Strikingly, effector protein Ecp2 was found to share structural similarity with pathogen killer toxin 4 proteins and had broad-spectrum antifungal activity, indicating a dual function in fungal competition and *Cf-ECP2*-mediated plant resistance.
- In tomato plants infected by *C. fulvum* parasitized by *H. pulvinata*, primary metabolism and defense-related genes were exclusively activated. These results suggest that in the tritrophic interaction, the mycoparasite simultaneously suppressed the pathogen and induced plant resistance. This study uncovers a multilayered molecular network in which the mycoparasite coordinates pathogen suppression and plant defense within the tritrophic interaction.

## INTRODUCTION

The biotrophic filamentous fungus *Cladosporium fulvum* Cooke [syn. *Fulvia fulva* (Cooke) Cif.] causes tomato leaf mold, a serious economic threat due to reducing fruit yield and quality and occasionally killing tomato plants (*Solanum lycopersicum* L.) (Thomma *et al*., 2005; de Wit *et al*., 2009). The fungus enters tomato leaves through open stomata in high humidity, colonizes the apoplast, and secretes numerous effector proteins to suppress plant immunity and facilitate infection (Mesarich *et al*., 2023). Commercial tomato cultivars have thus been bred with *Cf* resistance genes encoding receptor-like proteins (RLPs) with extracytoplasmic leucine-rich repeats from wild relatives (Mesarich *et al*., 2023). These RLPs recognize the corresponding *C. fulvum* effector proteins as virulence factors via a gene-for-gene relationship and trigger a hypersensitive response to restrict the growth of *C. fulvum* (Mesarich *et al*., 2023). However, *C. fulvum* rapidly evolves new races through mutation or loss of effector genes, thereby evading plant immunity (Iida *et al*., 2015; de la Rosa *et al*., 2024). The extensive use of resistant cultivars carrying only one or a few *Cf* genes has consequently led to the emergence of *C. fulvum* races that can overcome the resistance of the tomato plants (Luderer *et al*., 2002; Mesarich *et al*., 2014).

To date, over 100 effector genes have been identified in *C. fulvum* (Mesarich *et al*., 2018; Zaccaron & Stergiopoulos, 2024). Most encode small cysteine-rich proteins that contain an N-terminal signal peptide for secretion. The identified avirulence (Avr) and extracellular proteins (Ecp) effectors—such as Avr2, Avr4, Avr4E, Avr5, Avr9B, Avr9C, Ecp1, Ecp2, Ecp4, Ecp5, and Ecp6—exemplify the complex coevolution between fungal virulence factors and their cognate *Cf* immune receptors (Wulff *et al*., 2009; de la Rosa *et al*., 2024). Several effectors have been functionally characterized. Avr2 acts as a cysteine protease inhibitor that targets the apoplastic cysteine proteases Rcr3 and Pip1, thereby interfering with basal defense mechanisms conserved across solanaceous plants (Rooney *et al*., 2005; Shabab *et al*., 2008; Kourelis *et al*., 2020). Ecp6 suppresses chitin-triggered immunity by sequestering chitin oligomers released from the fungal cell wall, thus evading recognition by host pattern-recognition receptors (de Jonge *et al*., 2010). In contrast, Ecp2 induces cell death in both tomato and nonhost plants, suggesting a dual role as a potential virulence and avirulence factor (de Kock *et al*., 2004; de Wit *et al*., 2012). Notably, the Ecp2 effector contains an Hce2 (homolog of *Cladosporium fulvum Ecp2*) domain that is widely conserved across diverse fungal and bacterial taxa, implying a fundamental role for this domain in microbe–plant interactions (Stergiopoulos *et al*., 2012). However, the biochemical function of Ecp2 remains unclear. Of additional concern is that *C. fulvum* has developed resistance to fungicides (Yan *et al*., 2008; Watanabe *et al*., 2017). The extensive breakdown of the resistance conferred by single *Cf* resistance genes and the emergence of fungicide-resistant strains emphasize the need for sustainable disease management strategies beyond conventional breeding and chemical approaches. To better understand the molecular basis of disease suppression, previous studies have primarily focused on the *C. fulvum*–tomato interaction within the framework of the gene-for-gene model, but tritrophic interactions involving additional microorganisms remain largely unexplored.

*Hansfordia pulvinata* (Berk. & M.A. Curtis) S. Hughes (syn. *Dicyma pulvinata*) is a mycoparasitic fungus serendipitously discovered on tomato leaves infected with *C. fulvum* in a greenhouse (Iida *et al*., 2018). It parasitizes *C. fulvum* and overgrows lesions, effectively suppressing disease progression, and has broad mycoparasitic activity against all known *C. fulvum* races (Iida *et al*., 2018). *H. pulvinata* recognizes *C. fulvum* hyphae, coils around them, and directly penetrates the hyphae, then forms intercellular structures within the host fungus (Peresse & Le Picard, 1980; Maeda *et al*., 2025). It also produces an eremophilane-type sesquiterpenoid, 13-deoxyphomenone (also known as sporogen-AO1), which has antifungal activity against *C. fulvum* (Tirilly *et al*., 1983; Maeda *et al*., 2025). The compound was initially identified as a sporulation-inducing factor in the koji mold *Aspergillus oryzae* (Tanaka *et al*., 1984a, 1984b). Comparative genomics recently revealed that the biosynthetic gene cluster for 13-deoxyphomenone (*DPH*) in *H. pulvinata* shares high synteny with that in *Aspergillus* species, suggesting horizontal gene transfer from an ancestral *Aspergillus* lineage (Maeda *et al*., 2025). Subsequently, *H. pulvinata* appears to have retained this gene cluster to support its mycoparasitic lifestyle, repurposing the endogeni c sporogenesis function in *Aspergillus* species into exogenic antifungal activity that inhibits spore germination and hyphal elongation of *C. fulvum*. However, the molecular mechanisms underlying its mycoparasitism remain largely unknown. Since its mycoparasitism is not restricted to the phyllosphere (Maeda *et al*., 2025), comparative analyses of fungal–fungal interactions under different mycoparasitic conditions could provide new insights on the molecular basis of fungal host recognition and leaf-surface biocontrol mechanisms.

The *C. fulvum*–tomato interaction has long served as a valuable model for dissecting gene-for-gene dynamics in plant–pathogen systems (Mesarich et al. 2023.). Recent chromosome-scale genome assemblies of multiple *C. fulvum* races have substantially advanced our understanding of its genome structure and effector repertoire (Zaccaron *et al*., 2022; Zaccaron & Stergiopoulos, 2024). Nevertheless, the molecular basis governing fungal–fungal recognition and the tritrophic interactions with tomato plants remain largely uncharacterized. Here, we present transcriptomic evidence revealing how the mycoparasitic fungus *H. pulvinata* modulates both *C. fulvum* and tomato responses, providing molecular insights into the interplay among plant, pathogen and mycoparasite. These findings underscore the potential of the foliar mycoparasitic fungus as a promising biocontrol agent within complex plant–microbe ecosystems.

## MATERIALS AND METHODS

### Culture conditions for fungal strains and plants, treatments and assays

Fungal strains *H. pulvinata* 414-3 and *C. fulvum* CF301 were cultured on half-strength potato dextrose agar (PDA; BD Difco, Sparks, NJ, USA) or minimal medium (MM) agar (15 g sucrose, 5 g ammonium tartrate, 1 g NH_4_NO_3_, 1 g KH_2_PO_4_, 0.5 g MgSO_4_×7H_2_O, 0.1 g NaCl, 0.1 g CaCl_2_×H_2_O, 25 μL 0.2 mg mL^−1^ biotin, 15 g agar and 1 mL trace elements per liter) at 25 °C in the dark for 1 (*H. pulvinata*) and 2 weeks (*C. fulvum*) (Maeda *et al*., 2025). Spores were collected in distilled water and adjusted to 1 × 10^6^ spores mL^−1^ using a hemocytometer. For *in vitro* assays, 100 μL of each spore suspension was spread onto nylon membranes on PDA and incubated for 1 (*H. pulvinata*) or 2 weeks (*C. fulvum*) (hereafter “*in vitro Hp*” and “*in vitro Cf*” conditions; Fig. **S1a**). For establishing the mycoparasitism *in vitro*, membranes bearing *C. fulvum* colonies were transferred to water agar (nutrient-poor) and sprayed with 1 mL of *H. pulvinata* spore suspension (1 × 10^6^ spores mL^−1^); plates were then incubated at 25 °C in the dark for 2 weeks (*in vitro Cf/Hp*; Fig. **S1a**).

Seedlings of tomato (*Solanum lycopersicum*) cv. Moneymaker (lacking all known *Cf* resistance genes) were grown at 25 °C with 16 h light/8 h dark. Four-week-old plants were either grown in high humidity as healthy controls for another 2 weeks (Healthy leaf; Fig. **S1a**) or sprayed with approximately 3 mL of *C. fulvum* spore suspension (1 × 10^6^ spores mL^−1^), then incubated in high humidity for 2 weeks until lesions developed (“*on planta Cf*” interaction; Fig. **S1a**). For establishing the mycoparasitism on plants, plants with leaf mold lesions were sprayed with approximately 3 mL of *H. pulvinata* spore suspension (1 × 10⁶ conidia mL⁻¹) and incubated under high humidity at 25 °C for 2 weeks until lesions were overgrown by white mycelia of *H. pulvinata* (“*on planta Cf/Hp*”; Fig. **S1a**).

*N. benthamiana* plants were grown for 4 weeks at 25 °C with a 16 h light/8 h dark. *Arabidopsis thaliana* ecotype Col-0 seeds were surface-sterilized in 70% (v/v) ethanol for 1 min, rinsed three times with sterile distilled water, plated on Murashige and Skoog (MS) agar (4.33 g MS basal medium (Sigma-Aldrich, St. Louis, MO, USA) and 20 g sucrose per liter, pH 5.7), stratified at 4 °C in the dark for 1 month, then transferred to soil and grown for 5 weeks at 25 °C with 16 h light/8 h dark.

### Scanning electron microscopy (SEM)

Details of the sample preparation and imaging procedures for SEM are described previously (Maeda *et al*., 2025). Nylon membranes with fungal cultures were cut into about 5-mm squares and fixed in 2.5% (v/v) glutaraldehyde in 100 mM cacodylate buffer (pH 7.4) overnight at 4 °C, then 1% (w/v) osmium tetroxide in 100 mM cacodylate buffer (pH 7.4) for 1 h at room temperature. Samples were then dehydrated with a graded ethanol series, critical-point dried in absolute ethanol using Leica EM CPD300 critical point dryer (Leica Microsystems, Wetzlar, Germany), then mounted on stubs, coated with platinum-palladium, and observed with a Hitachi SU-8220 scanning electron microscope (Tokyo, Japan).

### RNAseq and transcriptome analysis

Three biological replicates were collected for each condition described above. Total RNA was extracted using the RNeasy Plant Mini Kit (Qiagen, Hilden, Germany) according to the manufacturer’s instructions. Poly(A) mRNA was enriched with the NEBNext Poly(A) mRNA Magnetic Isolation Module (New England Biolabs, Ipswich, MA, USA), and libraries were prepared using the NEBNext Ultra II Directional RNA Library Prep Kit (NEB). Libraries were sequenced on an Illumina NovaSeq 6000 platform (Illumina, San Diego, CA, USA) with the PE150 kit (150 bp × 2 paired-end), producing 2 or 4 Gb of data per sample.

Raw read quality was assessed using FastQC Ver. 16.0.1 (bioinformatics.babraham.ac.uk/projects/fastqc). Low-quality bases and adapter sequences were removed using Trimmomatic Ver. 0.39 (Bolger *et al*., 2014). Clean reads were aligned to the respective reference genomes using HISAT2 Ver. 2.2.1; SAM files were converted to BAM files with SamTools Ver. 1.18 (Li *et al*., 2009; Kim *et al*., 2019). Read counts were generated using Rsubread Ver. 2.20.0. Differentially expressed genes (DEGs) were identified with DESeq2 using thresholds of |log₂(Fold-change)| ≥ 1 and adjusted *P*-value (padj) ≤ 0.05 (Love *et al*., 2014). Gene Ontology (GO) and KEGG (Kyoto Encyclopedia of Genes and Genomes) enrichment analyses were performed using TBtools-II Ver. 2.119 (Aleksander *et al*., 2023; Chen *et al*., 2023; Kanehisa *et al*., 2025). Data were visualized using SRplot (Tang *et al*., 2023).

### Bioinformatic analysis

Orthologous coding sequences were identified using OrthoFinder Ver. 2.5.5 (Emms & Kelly, 2019); 100 single-copy orthologs were selected for multiple sequence alignment. Sequence alignments were generated using MAFFT Ver. 7.526 and subsequently trimmed with TrimAl Ver. 1.4 (Katoh and Standley, 2013; Capella-Gutiérrez *et al*., 2009). Maximum-likelihood phylogenetic trees were constructed with RAxML Ver. 8.2.12 using 1,000 bootstrap replicates and the GTRCAT or PROTCATAUTO model (Stamatakis, 2014). The consistency of tree topology was assessed using approximately unbiased (AU) tests (Shimodaira, 2002) with 10,000 bootstrap replicates as previously described (Maeda *et al*., 2025). Signal peptides were predicted with SignalP-6.0 (Teufel *et al*., 2022), candidate effectors with EffectorP-fungi 3.0 (Sperschneider & Dodds, 2022), and subcellular localization with LOCALIZER (Sperschneider *et al*., 2017). Secondary metabolite biosynthetic gene clusters were identified using the fungal version of antiSMASH (Blin *et al*., 2023). GPI-anchored proteins were predicted with NetGPI 1.1 (Gíslason *et al*., 2021), and carbohydrate-active enzymes (CAZymes) were annotated using DIAMOND (Zhang *et al*., 2018). Default parameters were used for all analyses unless otherwise specified.

For sequence similarity comparison, the EMBOSS Needle tool was used for pairwise alignments (Madeira *et al*., 2024). Protein domains and motifs were predicted using the Pfam database and the MEME Suite, respectively (Bailey *et al*., 2015; Mistry *et al*., 2021). The 3D structures of proteins were predicted using ColabFold Ver. 1.5.5 (Mirdita *et al*., 2022). Five models were generated with three iterative recycles each, and the models were ranked according to their predicted local distance difference test (pLDDT) scores. The top-ranked models were refined by AMBER energy minimization to resolve steric clashes and improve stereochemical geometry. The resulting relaxed structures were used for subsequent structural analyses. The structure of Ecp2 was analyzed using the Dali server to retrieve structural homologs based on a non-redundant dataset clustered at 25% sequence identity (PDB25), and structures were compared with TM-align (Zhang & Skolnick, 2005; Holm, 2022).

### Plasmid construction and protein expression

Necrosis-inducing protein 1-Like (NLP) protein coding sequences were synthesized (VectorBuilder, IL, USA), cloned into the pSfinx vector using a primer set (Table **S1**) and NEBuilder HiFi assembly kit (NEB). The NLP genes were digested with EcoRI and subcloned into the pPIC9K vector (Thermo Fisher Scientific, Waltham, MA, USA) for expression in *Pichia pastoris* GS115 strain according to the manufacturer’s instructions. Transformants were selected on YPD medium (Fujifilm Wako, Osaka, Japan) containing G418 (Fujifilm Wako) and confirmed by PCR using the primer set in Table **S1**. Expression was induced by adding 5 mL of the pre-culture grown in BMGY (buffered minimal medium supplemented with 1% glycerol, 2% peptone, and 1% yeast extract, 1.34% amino acid–free yeast nitrogen base, 100 mM potassium phosphate buffer [pH 6.0], and 4 × 10⁻⁵% D-biotin) broth to 100 mL of BMMY (1.5 % [w/v] methanol instead of glycerol in BMGY), and incubating the culture at 30 °C and 300 rpm for 96 h. Supernatants were collected by centrifugation at 12,000 rpm for 10 min.

The gene *Ecp2* was PCR-amplified from genomic DNA of *C. fulvum* using a primer set (Table **S1**) and cloned into pET-28b vector (Merck, Darmstadt, Germany) to express an N-terminal His_6_-tagged protein in *Escherichia coli* strain BL21(DE3). Expression was induced in LB medium with 1 mM IPTG at 16 °C for 16 h. Cells were resuspended with 50 mM KPi buffer (pH 7.4) and disrupted by sonication on ice (10 min: 2 s on–4 s off). Insoluble fractions were solubilized in 10 mL of denaturing buffer (6 M guanidine hydrochloride, 10 mM Tris, 10 mM β-mercaptoethanol, pH 8.0) for 1 h at room temperature. Proteins were purified in a Ni-affinity column HisTrap HP (Cytiva, Medemblik, The Netherlands), concentrated using Vivaspin 6 (GE Healthcare, IL, USA) and quantified with a Qubit fluorometer (Thermo Fisher Scientific). BL21(DE3) carrying empty pET-28b served as a negative control.

### Western blot analysis

All protein samples were heated at 95 °C for 10 min in sample buffer (1% SDS, 1% 2-mercaptoethanol, 10 mM Tris–HCl, 20% (v/v) glycerol, pH 6.8) for denaturation. The proteins were separated in 12% SDS-polyacrylamide gel and transferred onto PVDF membrane (Bio-Rad, Hercules, CA, USA). Membranes were blocked with 5% skim milk in TBS-T for 1 h before antibody incubation. Anti-His and anti-HA monoclonal antibodies (Cell Signaling Technology, Danvers, MA, USA) were diluted in Can Get Signal solution 1 (1:10,000) (TOYOBO, Osaka, Japan) and used as primary antibodies to detect Ecp2 and NLPs, respectively. Horseradish peroxidase-conjugated anti-rabbit antibody was diluted with Can Get Signal solution 2 (1:2,000) (TOYOBO) for use as the secondary antibody. Signals were visualized using an ECL western detection kit (GE Healthcare, Chicago, IL, USA) and captured using a ChemiDoc imaging system (Bio-Rad).

### Functional assay of NLPs

pSfinx plasmids were introduced into *Agrobacterium tumefaciens* strain GV3101 via electroporation (Ma *et al*., 2012). Agrobacterium suspensions were adjusted to OD_600_ = 0.3 in infiltration buffer (2.03 g MgCl_2_×6H_2_O, 1.95 g MES, 500 μL of 200 mM acetosyringone per liter, pH 5.6) and used to infiltrate leaves on 4-week-old plants of *N. benthamiana* by syringe pressure. Leaves were monitored for necrosis 14 days after infiltration. Necrosis was visualized by trypan blue staining (Ma *et al*., 2012) and quantified using ilastik (Ver. 1.4.1) (Berg *et al*., 2019).

*P. pastoris* culture supernatant containing NLP proteins or BM medium (BMGY without glycerol) was mixed at a 1:1 ratio with *C. fulvum* spore suspension (1 × 10^6^ spores mL^−1^). A 40 µL drop of each mixture was placed on a hydrophobic glass slide and incubated at room temperature for 24 h. After incubation, germination of 200 spores per treatment was assessed using a Nikon E600 light microscope (Tokyo, Japan). Culture supernatants containing NLP proteins were collected from *P. pastoris* and used to syringe-infiltrate *N. benthamiana* leaves. Twenty-four hours after infiltration, apoplastic fluid was collected by vacuum infiltration with distilled water, followed by centrifugation (1,000 rpm, 10 min) (Joosten, 2012). Equal volumes (10 µL) of *C. fulvum* spore suspension and apoplast fluid were then incubated at room temperature for 24 h. A light microscope (Nikon) was then used to assess 200 spores for germination.

Synthetic peptides of nlp24 (GenScript Biotech, Nanjing, China) were prepared in 500 nM stock solutions in dimethyl sulfoxide. Reactive oxygen species (ROS) were detected as previously described with slight modifications (Yang *et al*., 2022). Leaf disks (0.125 cm²) of *N. benthamiana* were floated on sterile distilled water for 24 h, then treated with 10 μL of a substrate mixture (5 μL of 10 mM L-012 [Fujifilm Wako] and 5 μL of protein or peptide). Luminescence was recorded every 2 min for 40 min using a Tristar3 Multimode Reader (Berthold Technologies, Bad Wildbad, Germany).

### Functional assays of Ecp2

The abaxial side of *Cf-ECP2* tomato leaves was infiltrated with purified Ecp2 protein (final concentration 1 μM) using a syringe. Leaves were incubated at 25 °C with 16 h light/8 h dark for 6 d and checked for a hypersensitive response (HR). *In vitro* antifungal activity was assayed as previously described (Chavarro-Carrero *et al*., 2024). Briefly, fungal spores were suspended in 5% PDB containing 15 μM Ecp2 or empty vector control (1 × 10⁴ spores mL^−1^), then 100 μL of the samples (three wells per sample) was incubated in wells of 96-well plates at 25 °C for 4 days. For each well, five fields of view were photographed (*N* = 15). Fungal growth was quantified and compared with the control using the Hybrid Cell Count module of the BZ-800X (KEYENCE, Osaka, Japan).

### Statistical analyses

R version 4.4.2 (R Core Team, 2024) was used for all analyses. The proportion of necrosis relative to total leaf area in *N. benthamiana* was assessed using Dunnett’s test, with the empty vector as the control (*N* = 3). Spore germination rate of *C. fulvum* treated with NLPs was analyzed by one-way ANOVA followed by Tukey’s HSD test (*N* = 3). The spore germination rate of *C. fulvum* in apoplastic fluid from *N. benthamiana* leaves treated with NLPs was compared using the Mann–Whitney *U* test, with the buffer as the control (*N* = 12). The relative growth rate of fungi and yeast treated with Ecp2 was compared using Student’s *t*-test, with the empty vector as the control (*N* = 15).

## RESULTS

### Overview of RNA-seq datasets

RNA was extracted from healthy control leaves, leaves infected with *C. fulvum* (*on planta Cf*), and leaves with *C. fulvum* lesions overgrown by *H. pulvinata* (*on planta Cf/Hp*) (Fig. **1a**; Fig. **S1a**). Because *H. pulvinata* parasitizes *C. fulvum* on leaf surfaces and on nutrient-poor water agar (Iida *et al*., 2018; Maeda *et al*., 2025), fungal RNA was also extracted from *in vitro* cultures of *H. pulvinata* alone (*in vitro Hp*), *C. fulvum* alone (*in vitro Cf*), and dual cultures of *C. fulvum* and *H. pulvinata* (*in vitro Cf/Hp*) on water agar (Fig. **1a**; Fig. **S1a**). Typical mycoparasitic growth characterized by white mycelium of *H. pulvinata* was observed at 14 days after inoculation for the *on planta Cf/Hp* and *in vitro Cf/Hp* conditions (Fig. **1a**). Microscopic observation confirmed that *H. pulvinata* penetrated *C. fulvum* hyphae (Fig. **1b**), and that *C. fulvum* entered tomato leaves through stomata (Fig. **S1b**) as previously described (Maeda *et al*., 2025). Also, hyphae of *H. pulvinata* elongated on the leaf surface and entered stomata in the presence of *C. fulvum* (Fig. **S1b**). Raw reads were aligned to the reference genome sequences of *H. pulvinata*, *C. fulvum* and tomato plants using HISAT2. The proportion of uniquely mapped reads varied across samples, ranging from 10.3 % and 88.1 % (Table **S2**), reflecting the multi organismal nature of several samples. Principal component analysis (PCA) of normalized expression values revealed clear clustering of biological replicates by treatment for each organism, indicating high within-treatment reproducibility (Fig. **S1c**). In the transcriptome analysis, we identified DEGs using adjusted *p*-value (padj) ≤ 0.05 and |log_2_ Fold-change| ≥ 1.0. Functional enrichment of DEGs was performed using KEGG pathway annotation to reveal biological processes that are positively and negatively regulated during mycoparasitism.

**Fig. 1.**
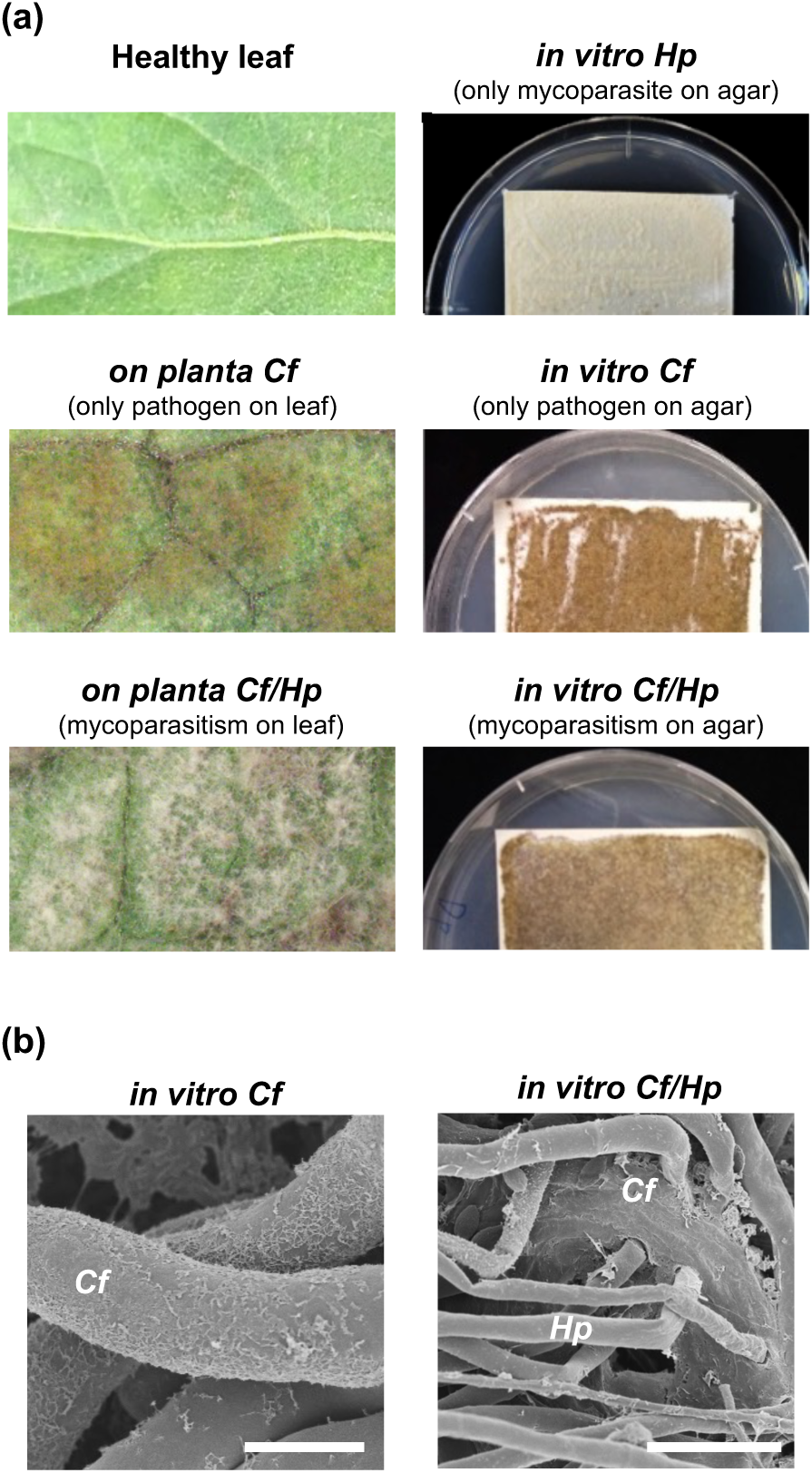
*Cladosporium fulvum* parasitized by mycoparasitic fungus *Hansfordia pulvinata*. (a) Representative images of a healthy tomato leaf, lesions caused by *C. fulvum* (*on planta Cf*), and lesions covered by white mycelium of *H. pulvinata* (*on planta Cf/Hp*). Tomato plants were inoculated with *C. fulvum*, and lesions developed after 2 weeks. *H. pulvinata* was then inoculated onto the diseased leaves, and white mycelia appeared on the lesions a week after the second inoculation. *In vitro* cultures of *H. pulvinata* (*in vitro Hp*) and *C. fulvum* (*in vitro Cf*) on nylon membranes on water agar after transfer from PDA. *H. pulvinata* parasitized *C. fulvum* colonies on water agar (*in vitro Cf/Hp*). (b) Scanning electron micrographs of hyphae of *C. fulvum* and *H. pulvinata* in *in vitro Cf* and *Cf/Hp* conditions. *H. pulvinata* hyphae have penetrated *C. fulvum* hyphae in the *in vitro Cf/Hp* condition. Bar indicates 10 μm.

### Transcriptomic reprogramming in *H. pulvinata* during parasitisation of *C. fulvum*

Transcriptome analysis of *H. pulvinata* during parasitisation of *C. fulvum in vitro* (*in vitro Cf/Hp*) and *on planta* (*on planta Cf/Hp*) and the control (*in vitro Hp*) was used to identify DEGs involved in mycoparasitism based on gene annotation (Table **S3**). During parasitisation of *C. fulvum in vitro* (*in vitro Cf/Hp*), 1,336 genes were significantly upregulated and 738 genes were downregulated (Fig. **2a**), whereas 1,793 were upregulated and 2,086 were downregulated during parasitisation of *C. fulvum* grown on tomato leaves (*on planta Cf/Hp*). A core set of 891 upregulated DEGs was shared between the *in vitro Cf/Hp* and the *on planta Cf/Hp* conditions and a core set of 557 genes was downregulated and were mapped to 21 upregulated and 25 downregulated KEGG pathways (Fig. **2b**), suggesting in *H. pulvinata* a conserved transcriptional program was activated during parasitisation of *C. fulvum* both *in vitro* and *on planta*.

**Fig. 2.**
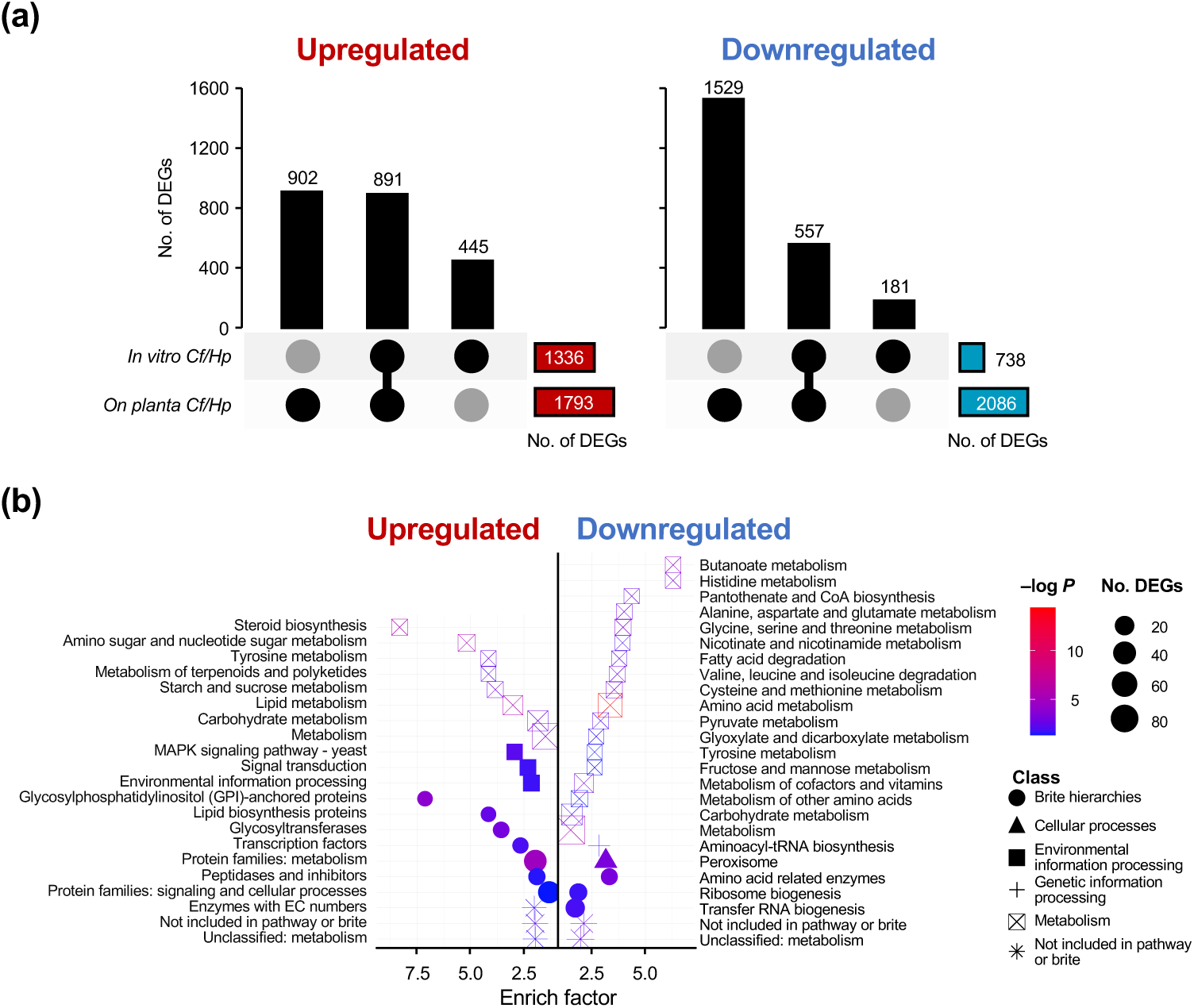
Differentially expressed genes (DEGs) and enriched KEGG pathways in mycoparasite *Hansfordia pulvinata*. (a) UpSet plot of DEGs in the *in vitro* and *on planta* mycoparasitic conditions (*in vitro Cf/Hp* and *on planta Cf/Hp*; Fig. 1a). Connected dots indicate shared DEGs; Black single dots represent condition-specific DEGs. Vertical bars show the number of up- and downregulated genes; colored horizontal bars indicate the total number of DEGs detected in each interaction. (b) Bubble plot of KEGG pathway enrichment analysis based on the 891 and 557 shared DEGs shown in (a). Keys: color of symbols indicates significance level of enrichment (–log adjusted *P*); size of symbol indicates the number of enriched DEGs in each pathway; the symbol indicates the functional class of the enriched DEGs.

KEGG enrichment indicated reprogramming of primary and secondary metabolism in *H. pulvinata* during parasitisation. KEGG pathways involved in secondary metabolism, including steroids, terpenoids, and polyketides, were strongly upregulated. Genome mining prediction revealed 33 putative secondary metabolite biosynthetic gene clusters in *H. pulvinata*—12 non-ribosomal peptide synthetases (NRPSs), 10 type I polyketide synthases (T1PKSs), 7 terpene synthetases, 3 fungal ribosomally synthesized and post-translationally modified peptides (fungal-RiPPs), and an isocyanide—all of which underwent a marked transcriptional change during parasitisation of *C. fulvum* (Table **S4**). Deoxyphomenone biosynthesis (*DPH*) genes were also upregulated, consistent with *H. pulvinata* producing the antifungal sesquiterpenoid 13-deoxyphomenone during parasitisation (Table **S4**) (Tirilly *et al*., 1983; Maeda *et al*., 2025). Interestingly, multiple amino acid metabolism pathways were consistently downregulated, despite *H. pulvinata* growing vegetatively during parasitisation (Fig. **2b**). Given the central role of amino acids for protein synthesis and vegetative growth, this KEGG pattern suggested that *H. pulvinata* may acquire amino acids derived from *C. fulvum* or tomato plants to supplement its amino acid pool during mycoparasitism.

### Expression profile and functional analysis of small secreted proteins in *H. pulvinata*

A comparison of the genome structures of *H. pulvinata* with those of fungi and the mycoparasite *Trichoderma atroviride* revealed that *H. pulvinata* had the smallest number of genes encoding carbohydrate-active enzymes (CAZymes), and the highest number of genes encoding secreted proteins (Fig. **S3**; Table **S3**). To analyze the secretome of *H. pulvinata*, we filtered 891 DEGs commonly upregulated in both mycoparasitic interactions (*in vitro Cf/Hp* and *on planta Cf/Hp*) and identified 165 genes encoding small secreted proteins (Fig. **3a**). Although most of them could not be annotated for function, 27 were annotated as genes previously reported to be associated with pathogenicity in plant pathogenic fungi, including glycoside hydrolase, T2 family ribonuclease, EPL1-like protein (orthologous to Ecp45 effector protein in *C. fulvum*), hydrophobin, necrosis-inducing protein, KP4-type killer toxin proteins, cutinase-like protein, superoxide dismutase protein, endoglucanase, and metalloprotease (Table **S5**). We focused on a necrosis inducing protein (DIP_10004627), designated *HpNLP1*, which is an Nep1-like protein (NLP) homologous to PsojNIP, because *HpNLP1* showed a particularly high –log_10_ padj, indicating a high level of significance (Fig. **3b**; Fig. **S4**). NLPs represent a widespread family of virulence-associated proteins in plant pathogens that trigger necrosis and generation of ROS in host plants (Seidl & Van den Ackerveken, 2019). Expression of *HpNLP1* increased 34-fold and 14-fold during parasitisation of *C. fulvum in vitro* and *on planta*, respectively (Table **S5**). The NLP gene *HpNLP2* (DIP_10008029) is also present in *H. pulvinata*, but its expression was unchanged during parasitisation of *C. fulvum*.

**Fig. 3.**
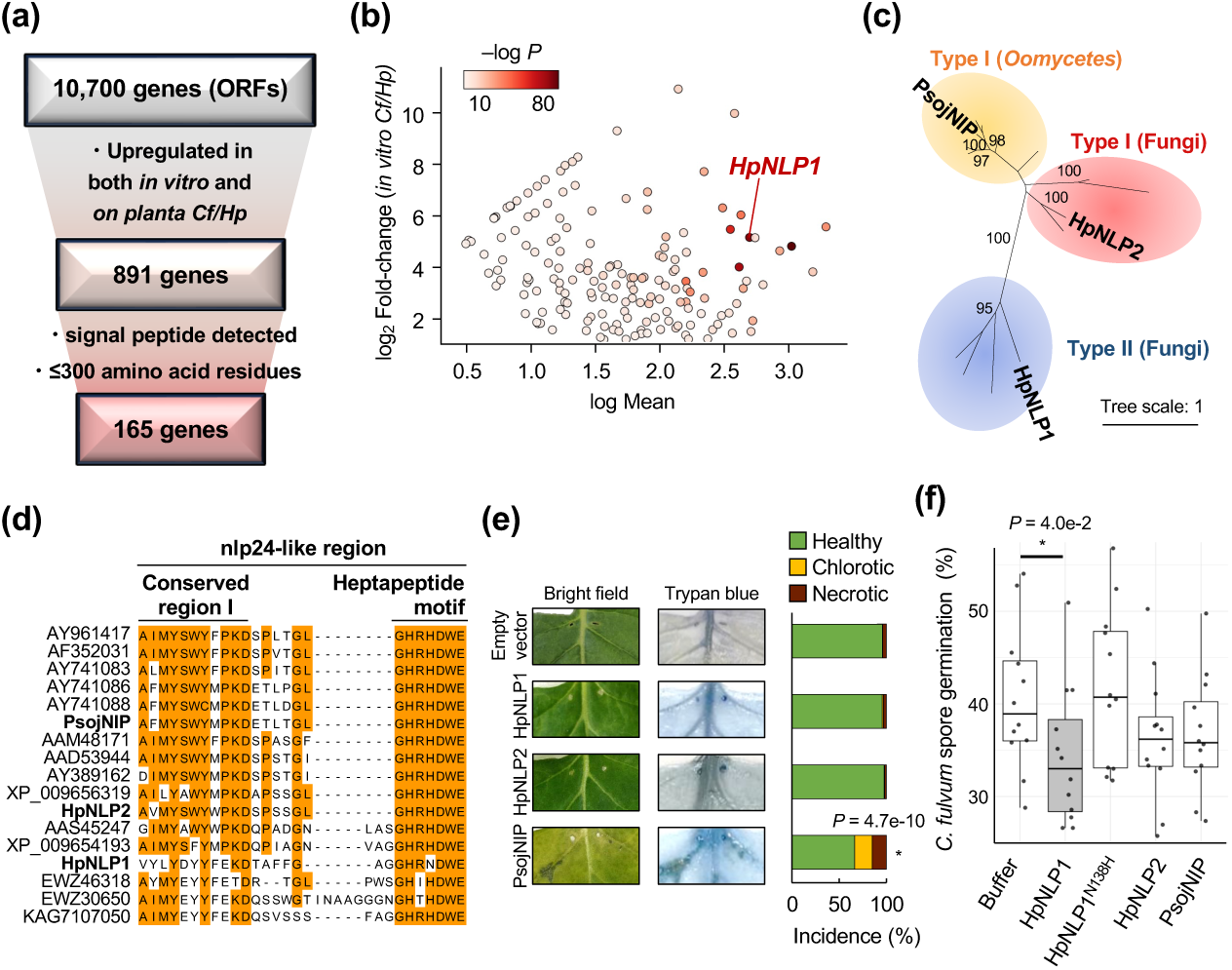
A Nep1-like protein (NLP) secreted from the mycoparasite *Hansfordia pulvinata* induces a plant-mediated toxicity against spores of *Cladosporium fulvum*. (a) Workflow for identifying small secreted proteins synthesized in *H. pulvinata* during mycoparasitism. (b) MA plot for expression levels of 165 selected genes. Cutoff thresholds are log_2_ Fold-change (±1.0) for *in vitro Cf/Hp* interaction and log Mean (0.3). (c) Phylogenetic tree constructed based on NLP amino acid sequences using the maximum likelihood method with 1,000 bootstrap replicates. Type I and Type II have two and four cysteine residues, respectively. Proteins analyzed in this study are in bold. (d) Multiple sequence alignment of NLP using MAFFT. Conserved residues are highlighted in orange. (e) Transient expression assays of NLPs in *Nicotiana benthamiana* via agroinfiltration. Chlorosis and necrosis were assessed 14 days after infiltration by trypan blue staining. Similar results were obtained in three independent experiments; representative results are in Supplementary Fig. S5c. (f) Percentage germination of conidia of *C. fulvum* after 24-h treatment with apoplastic fluid from *N. benthamiana* leaves that had been infiltrated with different NLPs or buffer. Asterisks indicate significant differences between treatments determined using Dunnett’s or the Mann–Whitney *U* test.

Phylogenetic analysis placed HpNlp1 and HpNlp2 within the type II and fungal type I clades, respectively, in NLPs (Fig. **3c**). HpNlp1 contained four cysteine residues predicted to form two disulfide bonds between positions 39–65 and 82–87 (Fig. **S4a** and **b**). Multiple sequence alignment showed that both HpNlp1 and HpNlp2 contained the conserved region I and the C-terminal heptapeptide GHRHDWE motif; this conserved 24-amino-acid peptide (nlp24) is recognized as a microbe- or pathogen-associated molecular pattern (MAMP/PAMP) (Oome *et al*., 2014). The tryptophan (W) and proline (P) residues essential for necrosis-inducing activity in the conserved region I (Zhou *et al*., 2012) were conserved in HpNlp2 but replaced by tyrosine (Y) and glutamic acid (E) in HpNlp1 (Fig. **3d**), which was the only NLP strongly expressed during mycoparasitism. In the heptapeptide motif of HpNlp1, the second histidine (H) was replaced by asparagine (N) (Fig. **3d**). Structural superposition of PsojNIP and HpNlp1 yielded a TM-score of 0.86 and a root mean square deviation (RMSD) of 1.93 Å, demonstrating that despite mutations in the nlp24-like regions of HpNlp1, the global protein structure was largely preserved, as indicated by these similar values compared to the PsojNIP (Fig. **S4c**).

Recombinant proteins PsojNIP, HpNlp1 and HpNlp2 were produced in *P. pastoris* and used to infiltrate *Nicotiana benthamiana* leaves. Western blot analysis of HpNlp1 demonstrated an additional band larger than the predicted molecular weight, suggesting dimer formation (Fig. **S5a** and **b**). As previously reported, PsojNIP induced necrosis and chlorosis (Qutob *et al*., 2002), whereas HpNlp1 and HpNlp2 did not trigger visible necrosis or ROS accumulation (Fig. **3e**; Fig. **S5c** and **d**). The high expression of *HpNLP1* gene under *in vitro Cf/Hp* condition, in the absence of the plant, suggested that HpNlp1 might directly suppress *C. fulvum* growth. Contrary to this expectation, no such direct antifungal activity was detected against *C. fulvum* spore germination and hyphal elongation (Fig. **S5e**). Remarkably, however, apoplastic fluid—where *C. fulvum* develops—collected from *N. benthamiana* leaves that had been infiltrated with HpNlp1 strongly suppressed *C. fulvum* spore germination, whereas the protein with a point mutantation (HpNlp1^N138H^) failed to elicit antifungal activity (Fig. **3f**). Neither of the nlp24 peptides of HpNlp1 and HpNlp1^N138H^ induced RLP23-mediated resistance (Fig. **S5f**). These findings together demonstrated that HpNlp1 diverges from typical necrosis-inducing NLPs in plant pathogens, inducing the accumulation of apoplastic antifungal substances that suppress *C. fulvum* spore germination instead of promoting host cell death.

### Transcriptomic responses of pathogen *C. fulvum* parasitized by *H. pulvinata*

Next, we investigated the DEGs of *C. fulvum* through KEGG pathway analysis after tomato was infected by *C. fulvum* (*on planta Cf*), dual infection on tomato leaves (*on planta Cf/Hp*), and dual culture on agar medium (*in vitro Cf/Hp*) (Fig. **1a**). DEGs in *C. fulvum* were identified during parasitisation relative to the *in vitro Cf* control, in which the fungus was cultured on water agar. A total of 1,902 genes were upregulated and 1,783 genes were downregulated when *C. fulvum* infected tomato leaves (*on planta Cf*) (Fig. **4a**). Under the two mycoparasitic conditions, 2,195 upregulated and 2,435 downregulated genes were detected *on planta Cf/Hp*, and 1,479 upregulated and 845 downregulated were detected *in vitro Cf/Hp*. We identified 363 upregulated and 158 downregulated genes shared between the two mycoparasitic conditions (*in vitro Cf/Hp* and *on planta Cf/Hp*), indicating a conserved transcriptional response to *H. pulvinata* attack. KEGG enrichment analysis assigned these shared DEGs to seven upregulated and three downregulated pathways, revealing strong activation of protein folding and processing pathways in *C. fulvum* during parasitisation by *H. pulvinata* (Fig. **4b**). Conversely, genes involved in amino acid metabolism were consistently downregulated, as was also observed in *H. pulvinata*, suggesting coordinated suppression of primary metabolism in this fungus-fungus interaction.

**Fig. 4.**
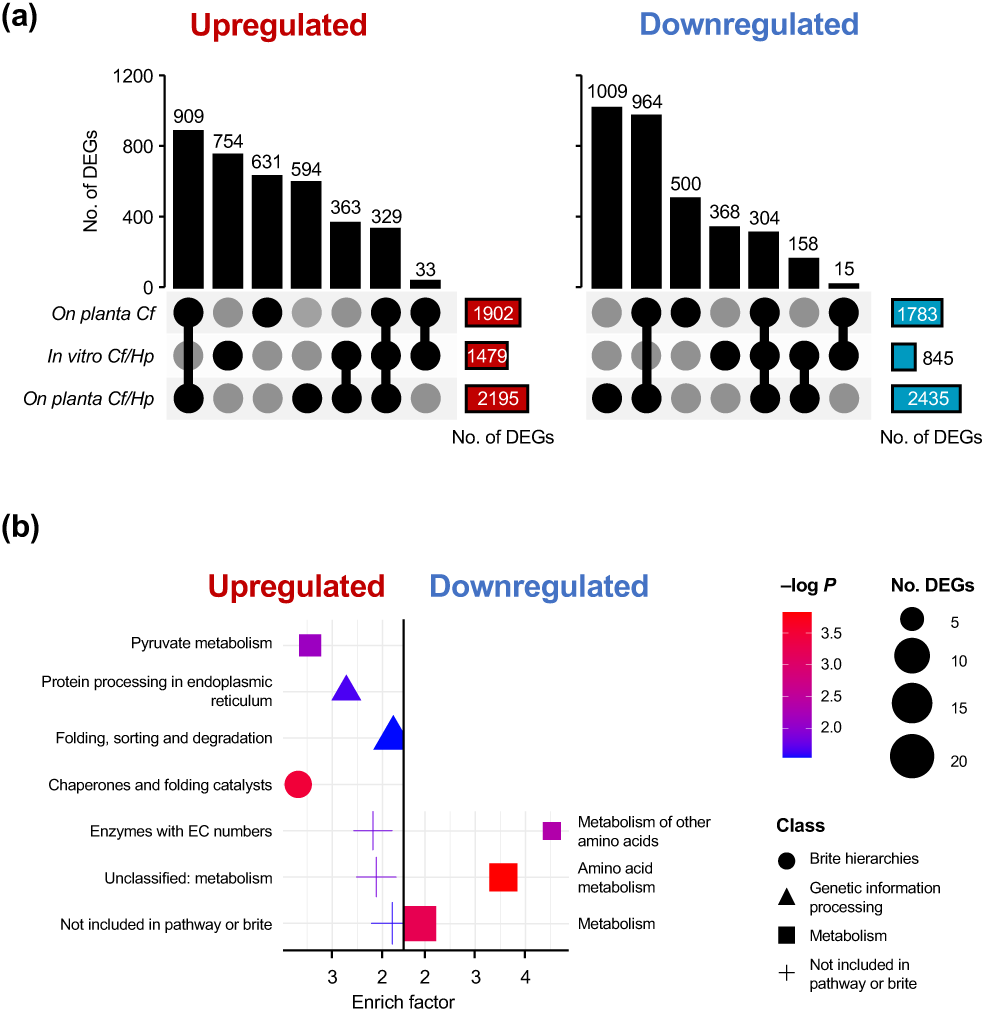
Differentially expressed genes (DEGs) and enriched KEGG pathways in the pathogen *Cladosporium fulvum*. (a) UpSet plot showing DEGs identified in the *on planta Cf, in vitro Cf/Hp,* and *on planta Cf/HP* conditions (see Fig. 1A). Connected dots indicate shared DEGs; Black single dots represent condition-specific DEGs. Vertical bars show the numbers of up- and downregulated genes; colored horizontal bars indicate the total number of DEGs detected in each condition. (b) Bubble plot of KEGG pathway enrichment analysis based on the 363 and 158 shared DEGs shown in (a). Keys: color of symbols indicates significant level of enrichment (–log adjusted *P*); size of symbol indicates the number of enriched DEGs in each pathway; the symbol indicates the functional class of the enriched DEGs.

In *C. fulvum*, protein folding and processing pathways were strongly activated when parasitized by *H. pulvinata* (Fig. **4a**). These responses are indicative of endoplasmic reticulum (ER) stress and accumulation of misfolded proteins, phenomena often triggered by exposure to antifungal compounds (Chaillot *et al*., 2015). Similar to the results in the mycoparasite *H. pulvinata*, the downregulation of secondary metabolism-related genes suggested that *C. fulvum* reallocated resources from the production of secondary metabolites to survive and mitigate stress. The expression profiles of DEGs detected in *C. fulvum* were further analyzed based on functional classification as previously reported by Zaccaron & Stergiopoulos (2024). Among the 42 genes associated with secondary metabolite biosynthesis, 15 (35.7 %) and 18 (42.8 %) were downregulated in the *on planta Cf* and *on planta Cf/Hp* interaction, respectively (Fig. **5a**; Table **S6**). Thirteen genes, including *PKS6* (CLAFUR5_12905), a key enzyme in cladofulvin biosynthesis (Griffiths *et al*., 2018), were consistently downregulated in both interactions.

**Fig. 5.**
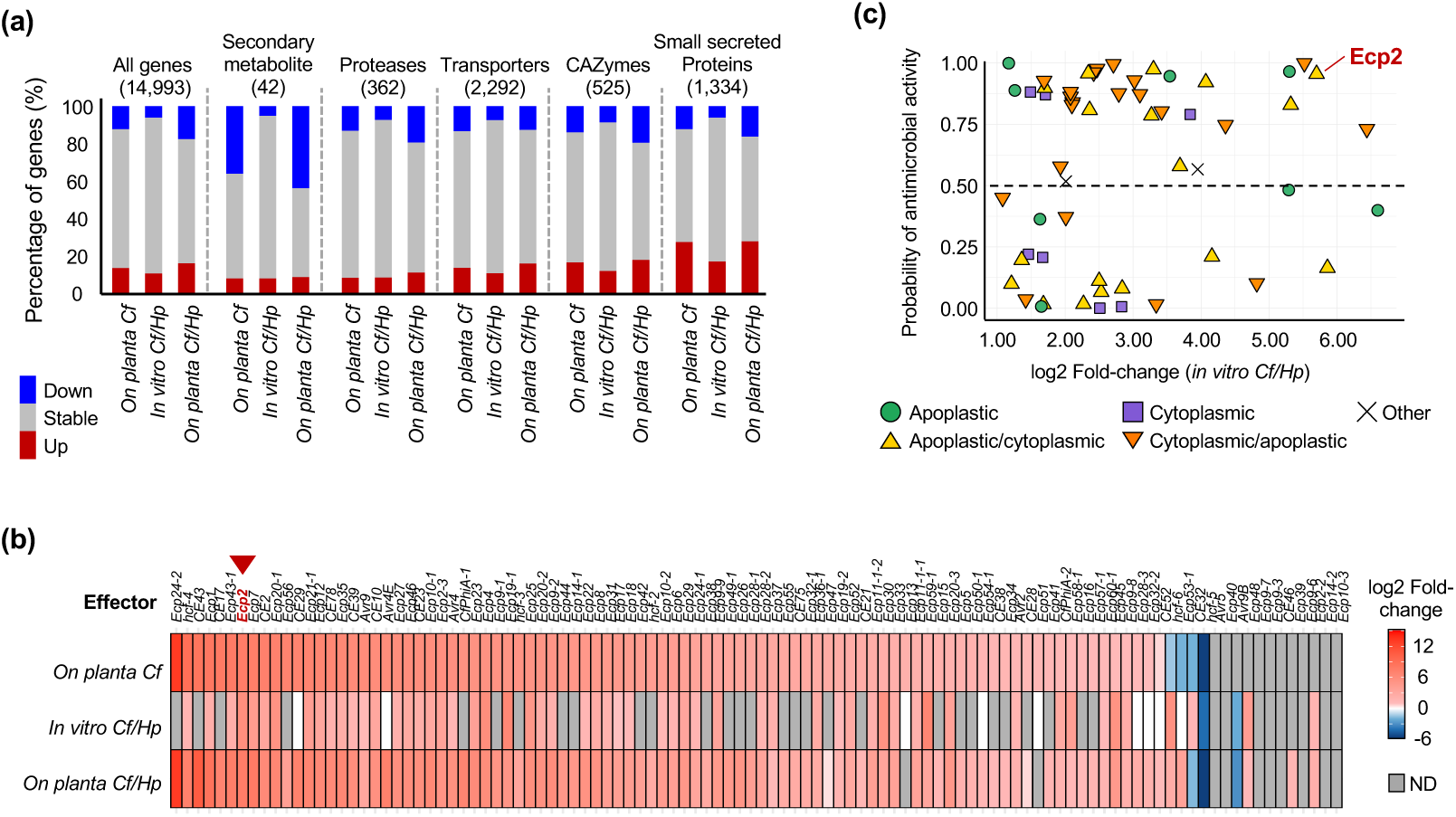
Effector genes in the pathogen *Cladosporium fulvum* were highly expressed during mycoparasitism by *Hansfordia pulvinata.* (a) Functional classification of DEGs identified for the *on planta Cf*, *in vitro Cf/Hp* and *on planta Cf/Hp* interactions. Red: upregulated, stable: grey, blue: downregulated. The number of genes assigned to each category is in parentheses. (b) Heatmap showing expression profiles of effector genes. The red-to-blue gradient represents up- and downregulated expression, respectively. Genes with an adjusted *P*-value ≥0.05 are indicated in gray as not detected (ND). Red arrowhead indicates *Ecp2*. (c) Antimicrobial activity prediction for 56 effector genes upregulated during the *in vitro Cf/Hp* interaction (determined using the AMAPEC program). Symbols denote protein location predicted by EffectorP-fungi 3.0.

In contrast, the category of small secreted proteins had the highest number of upregulated DEGs, including numerous known or predicted effectors (Fig. **5a**). An expression heatmap of effector genes showed that, as expected, the majority of the 106 effector genes were highly expressed during infection of tomato by *C. fulvum* (*on planta Cf*) (Fig. **5b**). Expression of effector genes are generally low on rich synthetic media (Mesarich *et al*., 2014); however surprisingly, 56 effector genes were upregulated in the *in vitro* coculture of *C. fulvum* and *H. pulvinata* (*in vitro Cf/Hp*) (Fig. **5b**). On tomato leaves, effector expression in *C. fulvum* remained high regardless of the presence of *H. pulvinata* (Fig. **S6**). These results suggested that *C. fulvum* effectors may play a defensive role, acting as countermeasures against mycoparasitic attack by *H. pulvinata* or are upregulated due to nitrogen limitation caused by parasitisation by *H. pulvinata.* To investigate this possibility, we conducted an AMAPEC analysis, which predicts antimicrobial proteins based on 3D-structural signatures (Mesny *et al*., 2024). Of the 56 effector genes that were upregulated under the *in vitro Cf/Hp* condition, 34 were predicted to possess antimicrobial activity, suggesting that many effectors may act as antifungal proteins (Fig. **5c**; Table **S7**). Among these, we focused on the effector Ecp2, which is known to trigger a hypersensitive response (HR) in tomato cultivars carrying the *Cf-ECP2* resistance gene.

### Structural and functional similarity between *C. fulvum* Ecp2 effector and antifungal killer toxin 4 proteins

The Alphafold2-predicted structure of Ecp2 yielded high confidence scores, with a pLDDT of 91.3 and a pTM of 0.872, indicating reliable local and global structures (Fig. **6a**). Structural similarity search using the DALI server identified two killer toxin 4 (KP4)-like proteins, ZtKP4 in *Zymoseptoria tritici* and UmVKP4 in *Ustilago maydis*, despite their low amino acid sequence similarity (2.9 to 36.9%) (Table **S8**). Comparative structural analysis showed that the overall arrangement of the α-helices and multiple β-sheets were conserved among these proteins (Fig. **6a** and **S7a**). The TM-scores for Ecp2 relative to ZtKP4 and UmVKP4 were 0.81 and 0.59, respectively, supporting that Ecp2 had the same structural fold as the KP4 protein family according to the SCOP and CATH classification criteria (Fig. **6b**).

**Fig. 6.**
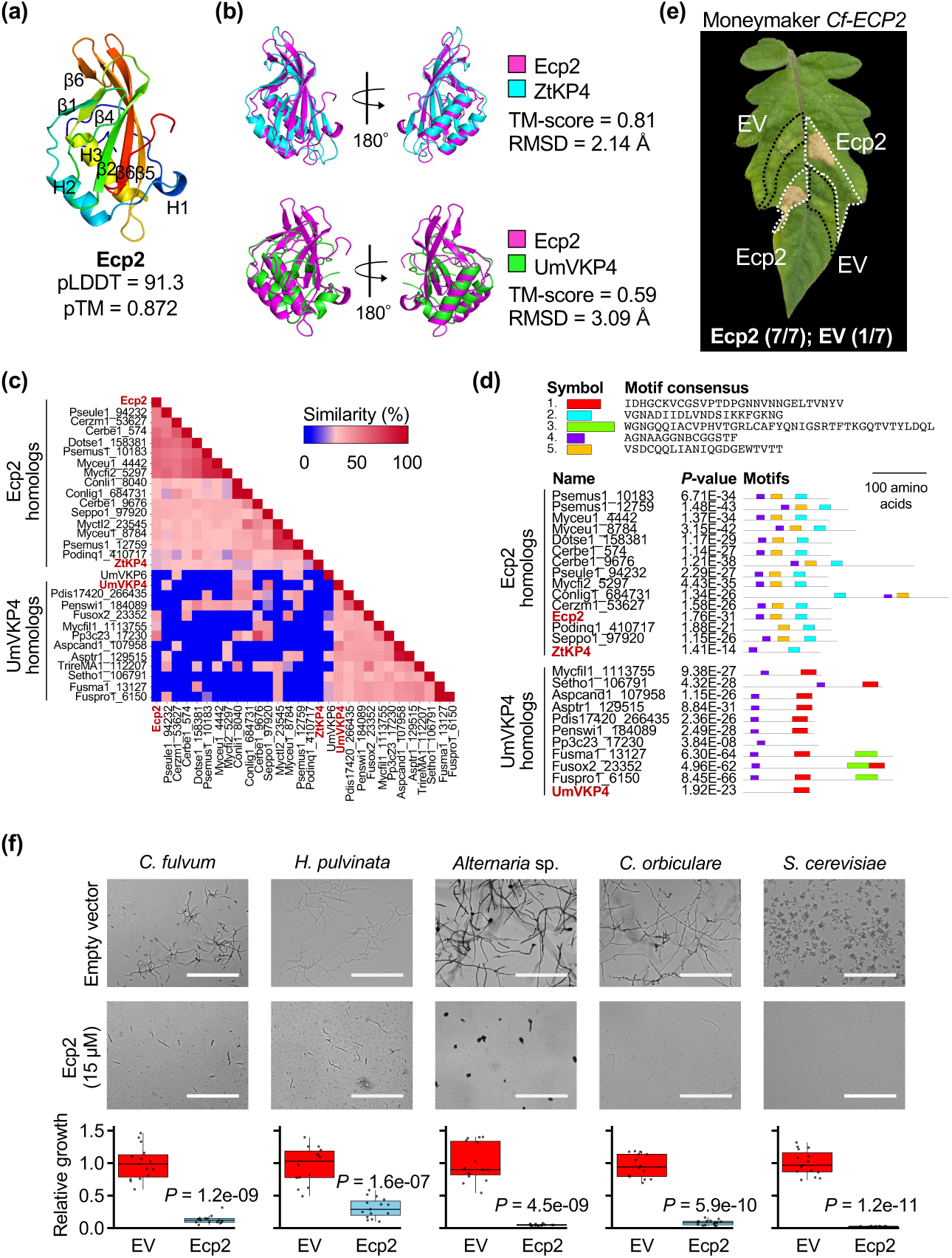
Sequence dissimilarity and structural similarity between *Cladosporium fulvum* effector Ecp2 and killer toxin KP4 proteins. (a) Predicted 3D structure of Ecp2 generated using AlphaFold2. The color spectrum from blue to red indicates the N- to C-terminal positions of residues. (b) Structural comparison of Ecp2 with ZtKP4 and UmVKP4 proteins. The superposition of the aligned proteins was evaluated using TM-align, and the TM-score and RMSD values are shown. (c) Heatmap showing sequence similarity among Ecp2, ZtKP4, and UmVKP4 proteins as identified by BLASTp analysis. (d) Conserved motifs of the proteins identified by MEME motif analysis. (e) Representative tomato leaf from 4-week-old tomato plant carrying the *Cf-ECP2* resistance gene at 6 days after infiltration with purified Ecp2 protein produced by *Escherichia coli*. White dotted outline: area infiltrated with infiltrated with 1 μM Ecp2; black dotted outline: infiltrated with control protein purified from *E. coli* harboring an empty vector (EV). Numbers in parentheses indicate the number of leaves with a hypersensitive response (HR) relative to the number of total leaves. (f) *In vitro* antifungal activity of Ecp2 against *C. fulvum*, *H. pulvinata*, *Alternaria* sp., *Colletotrichum orbiculare*, and *Saccharomyces cerevisiae*. Spores were cultured on 5% potato dextrose agar with or without Ecp2. Relative hyphal length was determined 4 days after treatment. Bars = 250 μm. Significant differences between EV and Ecp2 treatments were assessed using Student’s t-test, and *p*-values are indicated.

To explore the evolutionary relationships among these KP4-like proteins, we searched for homologs of Ecp2, ZtKP4, and UmVKP4. Nineteen Ecp2 homologs were identified across three fungal classes, while only one ZtKP4 homolog was detected in *Zymoseptoria tritici* (syn. *Mycosphaerella graminicola*) (Table **S9**). A total of 188 UmVKP4 homologs were found in 10 classes. Based on sequence similarity, these proteins clustered into two groups; Ecp2 and ZtKP4 were placed in the same clade (Fig. **6c**). Motif analysis revealed distinct conserved amino acid motifs in each group, and UmVKP4 group lacked an Hce2 domain (Fig. **6d**). While Ecp2 and ZtKP4 shared two conserved disulfide bonds, UmVKP4 had five, suggesting differences in structural stability and folding despite their shared overall architecture (Fig. **S7b**). These results indicated that Ecp2 and UmVKP4 protein groups likely originated from two distinct evolutionary lineages.

Given that Ecp2 is structurally and phylogenetically close to ZtKP4, which has antifungal activity (de Guillen *et al*., 2024), we investigated the antifungal potential of Ecp2. Infiltration of *Cf-ECP2* tomato leaves with recombinant Ecp2 (Fig. **S7c**) triggered a specific HR (Fig. **6e**). At the high concentrations used, the active Ecp2 protein significantly inhibited hyphal growth in diverse fungi: *C. fulvum*, *H. pulvinata*, *Alternaria* sp. isolated from tomato fruits, the plant pathogen *Colletotrichum orbiculare* and the yeast *Saccharomyces cerevisiae* (Fig. **6f**). Together, these results demonstrated that Ecp2 has a dual role: inducing *Cf-ECP2*-mediated immunity in plant while inhibiting fungal growth. *H. pulvinata* exhibited marginally higher resistance than the other tested fungi and yeast to Ecp2.

### Enrichment analysis of DEGs and KEGG pathways in tomato plants

We also analyzed differential gene expression in tomato leaves inoculated with *C. fulvum* and inoculated with *C. fulvum* followed by *H. pulvinata*, each compared with uninoculated control leaves (Fig. **1a**). Infection by *C. fulvum* alone led to 3,231 upregulated and 4,980 downregulated genes, whereas in the presence of *H. pulvinata* (*on planta Cf/Hp*), 3,465 genes were upregulated and 5,679 downregulated (Fig. **7a**). Among these, 1,619 up- and 2,153 downregulated genes were uniquely responsive to *H. pulvinata*, mapping to 35 and 33 enriched KEGG pathways, respectively (Fig. **7b**). Pathways associated with plant–pathogen interaction, phytoalexin biosynthesis (stilbenoid, diarylheptanoid and gingerol), biosynthesis of unsaturated fatty acids, other secondary metabolites, phenylpropanoid biosynthesis and plant hormone signal transduction were induced when *H. pulvinata* parasitized *C. fulvum* on leaves (Fig. **7b**). Genes encoding calcium-dependent protein kinases, PR1, RIN4, MAP kinase, WRKY22 and defensins were highly expressed (Fig. **S8a**; Table **S10**). These results indicated that *H. pulvinata* triggers plant immunity during parasitisation of *C. fulvum* on tomato leaves. In contrast, pathways associated with photosynthesis antenna proteins and porphyrin metabolism, which is involved in chlorophyll biosynthesis, were upregulated during infection by *C. fulvum* alone, but significantly downregulated when parasitized by *H. pulvinata* (Fig. **7b**; Fig. **S8b** and **c**), indicating that *H. pulvinata* triggered pronounced plant resistance responses during parasitisation of *C. fulvum*.

**Fig. 7.**
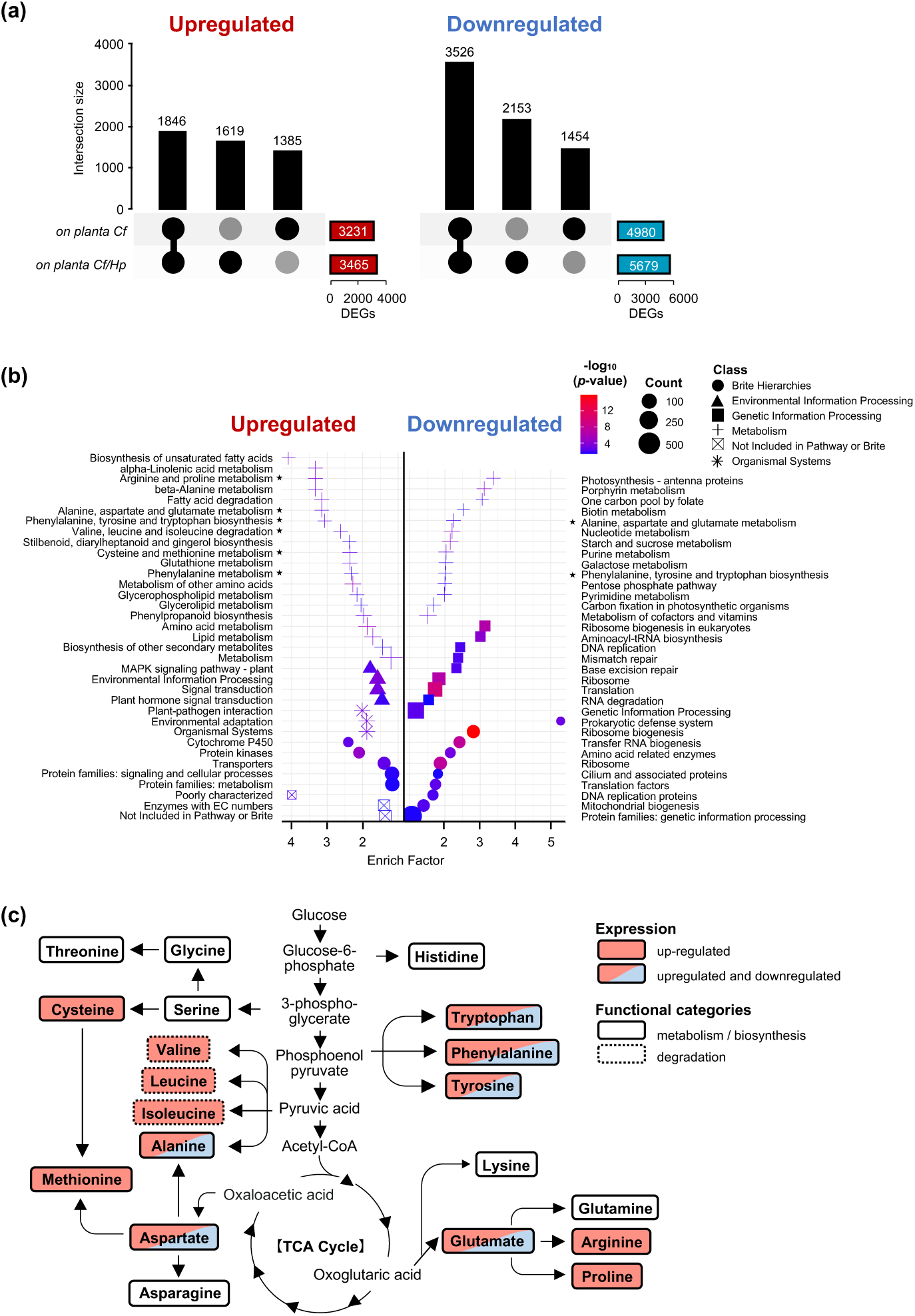
Differentially expressed genes (DEGs) and enriched KEGG pathways in tomato plants. (a) UpSet plot of DEGs in the *on planta Cf* and *on planta Cf/Hp* conditions (see Fig. 1a). Healthy leaves were used as a control. Connected dots indicate shared DEGs; Black single dots represent condition-specific DEGs. Vertical bars show the number of up- and downregulated genes; colored horizontal bars indicate the total number of DEGs detected in each interaction. (b) Bubble plot of KEGG pathway enrichment analysis based on the 1,619 and 2,153 DEGs shown in (a). Keys: color of symbols indicates significance level of enrichment (–log adjusted *P*); size of symbol indicates the number of enriched DEGs in each pathway; the symbol indicates the functional class of the enriched DEGs. Stars indicate pathways highlighted in (c). (c) KEGG pathways involved in amino acid biosynthesis in tomato.

Strikingly, 13 amino acid biosynthesis and metabolism pathways were strongly activated in the *on planta Cf/Hp* condition (Fig. 7b and c). Although amino acid metabolism was broadly repressed in both *C. fulvum* and *H. pulvinata* during mycoparasitism (Fig. 2b; Fig. 4b), it was upregulated in tomato (Fig. 7b and c). The pronounced activation of amino acid biosynthesis pathways in tomato suggests that the plant supplies amino acids to both the pathogen *C. fulvum* and its mycoparasite *H. pulvinata* during the tritrophic interaction.

## DISCUSSION

In this study, we analyzed gene expression in tomato plants, the phytopathogenic fungus *C. fulvum*, and its mycoparasite *H. pulvinata* alone or during their bitrophic and tritrophic interaction to capture coordinated and distinct responses. We thus discovered previously unrecognized aspects of molecular crosstalk and revealed how the transcriptome of each organism was dynamically reprogrammed during these interactions.

### Evolutionary and functional insights into an atypical NLP in the biocontrol mycoparasite *H. pulvinata*

The identification of 165 secreted small proteins upregulated during mycoparasitism highlights the central role of the secretome of *H. pulvinata*. Among them, HpNlp1, an atypical NLP, lacked both necrosis-inducing activity or RLP23 receptor-mediated response in plants, unlike canonical NLPs such as PsojNIP (Qutob *et al*., 2006; Böhm *et al*., 2014; Oome *et al*., 2014). Instead, HpNlp1 accumulated antifungal substances in the apoplast and inhibited spore germination of *C. fulvum*. Although *H. pulvinata* cannot grow independently on tomato leaves (Iida *et al*., 2018), it follows the hyphae of *C. fulvum* that enter the stomata of tomato leaves (Fig. **S1b**), indicating that the two fungi co-occur in the tomato apoplast. Furthermore, the functional divergence of the NLPs between the pathogen and mycoparasite indicates the evolutionary plasticity of NLP family. Despite its structural similarity to PsojNIP, HpNlp1 has distinct biological functions, suggesting that subtle amino acid modifications can drive functional divergence within the NLP family. If *H. pulvinata* induced a strong necrotic or immune response in tomato, the host fungus *C. fulvum* would decline, limiting its own nutrient supply. Therefore, the high upregulation of a noncytotoxic NLP such as HpNlp1 during mycoparasitism is evolutionarily reasonable. Notably, the *HpNLP1* gene was expressed in *H. pulvinata* when parasiting its host *on planta* and *in vitro*, suggesting that it recognizes *C. fulvum* itself rather than plant-derived signals. Although HpNlp1 did not directly inhibit *C. fulvum* growth, it indirectly promoted the accumulation of antifungal metabolites, suggesting coevolution of *H. pulvinata* with not only its fungal host *C. fulvum* but also tomato plants.

KEGG enrichment analysis indicated that *H. pulvinata* activated multiple secondary metabolic pathways, particularly those related to terpenoids, polyketides, steroids and *DPH* biosynthetic gene cluster, during mycoparasitism. These findings agree with previous findings that *H. pulvinata* produces 13-deoxyformenone, an eremophilane-type sesquiterpenoid, which is fungistatic to *C. fulvum* (Maeda *et al*., 2025). The production of antifungal substances triggered by HpNlp1 in the tomato apoplast was correlated with the induction of genes encoding defensins and phytoalexins, which were expressed only when *C. fulvum* was parasitized by *H. pulvinata* on tomato leaves. Given that similar noncytotoxic NLPs from the biocontrol oomycete *Pythium oligandrum* also induce plant defensins (Yang *et al*., 2022), HpNlp1 may represent a conserved mechanism among biocontrol agents that modulate plant immunity without causing cell death. *H. pulvinata* may employ a two-tiered strategy to preserve the spores of its host fungus *C. fulvum*, combining a direct antifungal sesquiterpenoid with an indirect, noncytotoxic protein, HpNlp1.

### Ecp2 effector as a novel antifungal protein in the pathogen *C. fulvum*

KP4 family proteins, broadly conserved in phytopathogenic fungi, are antifungal but do not interact with the plant. For instance, *U. maydis* UmVKP4 suppresses the growth of susceptible strains of the same species by inhibiting calcium-dependent pathways that regulate the cell cycle and morphogenesis (Gu *et al*., 1995; Gage *et al*., 2002). *Z. tritici* ZtKP4 has fungicidal activity even against itself (de Guillen *et al*., 2024). Our structural modeling showed that *C. fulvum* Ecp2 shared similarity with KP4 proteins and had broad-spectrum antifungal activity ranging from filamentous fungi to yeast, including self-inhibition similar to that of ZtKP4. The β1–β2 loop in Ecp2 should functionally correspond to the β3–β4 loop of UmVKP4, which mediates interactions with intracellular proteins (Gu *et al*., 1995). However, Ecp2 lacked the key lysine residue (Lys42) that UmVKP4 requires for full antifungal activity (Gage *et al*., 2001), suggesting a similar mechanism underlying antifungal action but involving an alternative molecular interface. During the arms race between tomato and *C. fulvum*, a KP4-like antifungal protein may have been diverted to Ecp2 effector. However, the structural basis underlying *CfECP2*-mediated immune recognition and how Ecp2 acquired the dual ability to interact with both fungal and plant targets remain unresolved.

Ecp2 was strongly expressed even during *in vitro* mycoparasitism, suggesting a dual role as both a virulence factor against plants and an antifungal protein in microbial competition. Similar dual functionality is seen in the root endophyte *Serendipita* spp., where a chitinase encoding effector is induced during both plant colonization and microbial interaction (Eichfeld *et al*., 2024). The vascular wilt fungus *Verticillium dahliae* secretes antimicrobial effectors such as VdAve1 and VdAMP2 to gain an ecological advantage in the soil biome (Snelders *et al*., 2020). The soil-borne white root rot pathogen *Rosellinia necatrix*, which is phylogenetically related to *V. dahliae*, secretes antimicrobial effector proteins during host plant colonization (Chavarro-Carrero *et al*., 2024). While the rhizosphere harbors highly diverse and competitive microbial communities, the diversity of the microbial community in the endosphere such as the apoplastic space is generally relatively low (Snelders *et al*., 2022) because colonization of the apoplast is typically restricted to specialized microbes that can secrete effectors to suppress host immunity (Rocafort *et al*., 2020). In fact, despite previous predictions that some apoplastic effectors produced by *C. fulvum* may have antimicrobial activity, no direct evidence had been found so far (Mesarich *et al*., 2018). Our finding indicates the potential for *C. fulvum* not only to utilize effectors for efficient colonization of tomato leaves but also to compete with/inhibit other microbes in the endosphere and phyllosphere. Although *H. pulvinata* was moderately tolerant of Ecp2, the molecular basis for this tolerance during mycoparasitism remains unclear.

The Hce2 (Homologs of *C. fulvum* Ecp2) superfamily is widely conserved across the *Ascomycota* and *Basidiomycota*, suggesting that its origin predates the divergence of the *Dikarya* lineage (Stergiopoulos *et al*., 2012). In contrast, *U. maydis* UmVKP4, which shares structural similarity with Ecp2, lacks the Hce2 domain and shared sequence motifs, and differs in the number of disulfide bonds (Fig. **S7b**). These findings suggest that UmVKP4 represents a case of convergent evolution, resulting in structural resemblance to Ecp2 and ZtKP4. Molecular evolutionary analyses suggest that the amino acid substitution underlying Hce2 function diversification was fixed early in evolution (Stergiopoulos *et al*., 2012). Ancestral Hce2 proteins may initially act as antimicrobial factors to provide competitive advantages in microbe-rich habitats, and have been co-opted as an effector to inhibit plant immunity. Mutations in effector genes that allow *C. fulvum* to evade *Cf* resistance in plants may increase its susceptibility to antagonistic microbes such as *H. pulvinata*. This hypothesis suggests that mycoparasitic biocontrol agents can circumvent the classical arms race between plant and pathogen, providing a sustainable disease management.

### Potential metabolic trade-offs in amino acid pathways between fungi and tomato plants

The physiological changes found in tomato plants highlight the dynamic interplay between microbial interplay and plant defense responses. The expression of genes in multiple amino acid biosynthetic pathways were enhanced only when *H. pulvinata* parasitized *C. fulvum* on tomato leaves. Amino acids are key mediators of plant–microbe interactions, functioning as signaling molecules that activate defense and as nutritional sources that are exploited by pathogens (Moormann *et al*., 2022; Tünnermann *et al*., 2022). Indeed, infection by *C. fulvum* leads to the accumulation of free amino acids in the apoplast (Solomon & Oliver, 2001). Because the acquisition of plant-derived amino acids is energetically advantageous to microbes, many plant pathogens modulate expression of genes encoding amino acid transporters to optimize uptake (Sonawala *et al*., 2018). Thus, the regulation of amino acid assimilation and transport in plants can be a decisive factor shaping the outcome of plant–pathogen interactions.

In contrast, amino acid biosynthetic pathways in *H. pulvinata* and *C. fulvum* were downregulated under mycoparasitic conditions, suggesting that tomato-derived amino acids could be shared or sequentially utilized by these fungi (Fig. **8**). In this metabolic trade-off, tomato plants might invest in amino acids that benefit *H. pulvinata*, while receiving biocontrol protection in return. Although amino acids are fundamental primary metabolites during fungal vegetative growth, the metabolic profile of *H. pulvinata*, which exhibited vigorous hyphal growth during mycoparasitism, deviated from this conventional paradigm. Because *H. pulvinata* parasitizes *C. fulvum*, it likely becomes the recipient of these plant-derived nutrients via its host *C. fulvum*, forming a potential metabolic trade-off in the tritrophic interaction. In this context, the activation of amino acid biosynthesis in tomato may provide a nutritional foundation that ultimately supports colonization and biocontrol activity by *H. pulvinata*. Although *C. fulvum* may also acquire amino acids from plants, these resources are likely reallocated to *H. pulvinata*, the ultimate beneficiary of the tritrophic interaction. Similar strategies have been reported for the biocontrol fungus *Pseudozyma flocculosa,* active against powdery mildews, which transiently suppresses photosynthesis to promote amino acid release (Laur *et al*., 2018). The mycoparasitic fungus *Trichoderma erinaceum* exploits environmental amino acids to enhance the synthesis of antifungal secondary metabolites derived from amino acids (Guo *et al*., 2020). Together, these results suggested that shifts in amino acid metabolism enable *H. pulvinata* to harness pathogen-induced plant responses to its own benefit.

**Fig. 8.**
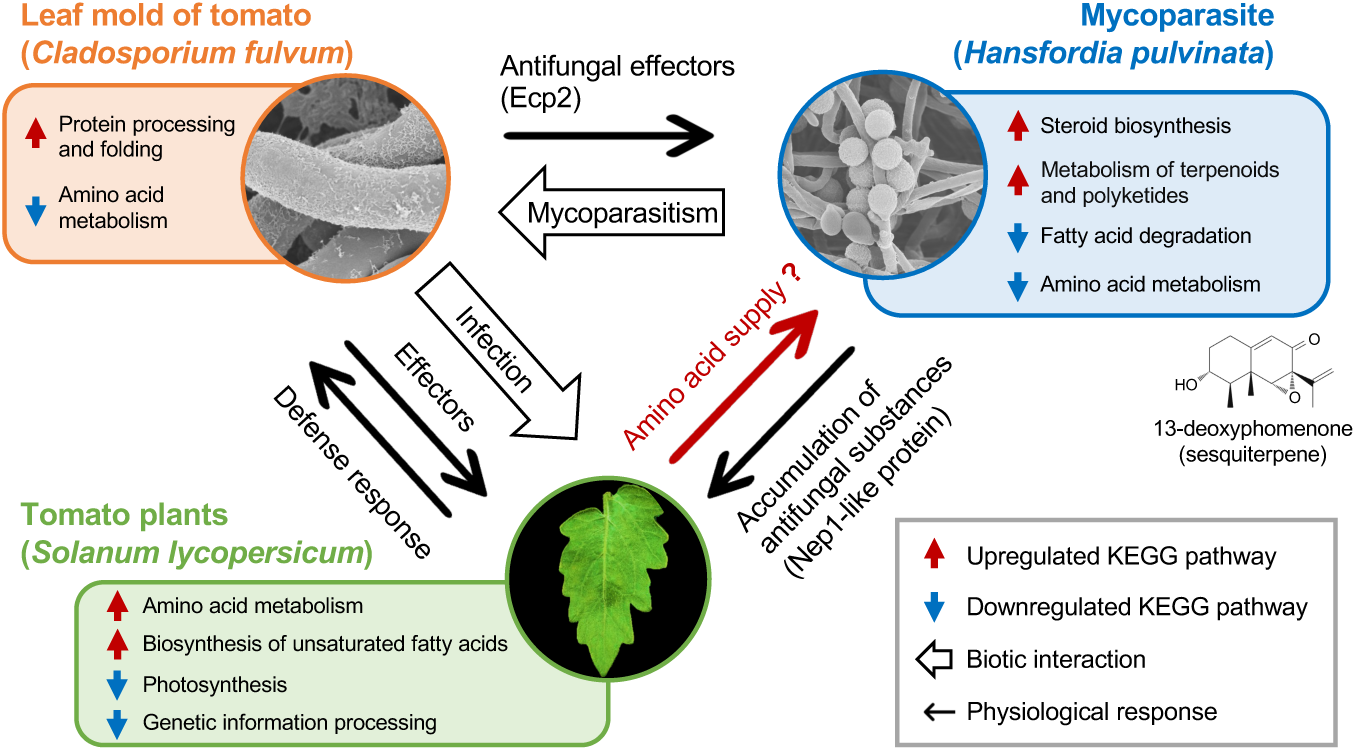
Overview of the tritrophic interaction among tomato plants, pathogen *Cladosporium fulvum*, and mycoparasite *Hansfordia pulvinata*. *C. fulvum* secretes effector proteins that function as virulence factors in susceptible tomato cultivars and antifungal proteins active against other fungi, but act as avirulence factors upon recognition by *Cf*-resistant cultivars. *H. pulvinata* parasitizes *C. fulvum* and induces the production of apoplastic antifungal substances in tomato plants via the Nep1-like protein HpNlp1. *H. pulvinata* also activates the biosynthesis of the antifungal sesquiterpene 13-deoxyphomenone during mycoparasitism. Other plant defense responses support this biocontrol effect. Pathways related to amino acid biosynthesis in the three organisms suggest that nutrients from the plant are supplied to *H. pulvinata*.

### Mycoparasitism by *H. pulvinata* modulates plant immunity and pathogen growth

In conclusion, the mycoparasite *H. pulvinata* actively reprogrammes gene expression and physiological responses in the pathogen *C. fulvum* and its tomato host to establish a stable tritrophic interaction (Fig. **8**). *H. pulvinata* induced expression of the 13-deoxyphomenone biosynthetic gene cluster in tomato plants; antimicrobial substances accumulated in the apoplast and directly suppressed *C. fulvum*, thus enhancing plant resistance. In turn, *C. fulvum* secretes effectors such as Ecp2 that exhibites roles in both fungal competition and plant immunity. During the tritrophic interaction in tomato, amino acid metabolism and defense response pathways are reprogrammed, orchestrating distinct strategies against beneficial and pathogenic microbes. Collectively, these findings underscore the dynamic interplay in plant–pathogen–mycoparasite interactions, demonstrating that *H. pulvinata* simultaneously strengthens tomato immunity and represses *C. fulvum* virulence via direct antagonism and modulation of plant metabolic pathways.

## ACKNOWLEDGEMENTS

This work was supported by a Grant-in-Aid for Scientific Research from JSPS (20H02993 and 24K08919) and the Research and Implementation Promotion Program through Open Innovation Grants (JPJ011937) from the Project of the Bio-oriented Technology Research Advancement Institution (BRAIN) (Y. Iida). We are grateful to K. Ikeda for providing fungal strains, M.H.A.J. Joosten for *Cf-ECP2* tomato seeds, D. Takemoto for the *psojNIP* gene, F. Takken for pSfinx, and Y. Kubo for the *C. orbiculare* strain. We thank Y. Kubo, M.Z. Fanani and S. Kodama for valuable suggestions, C. Tanaka for laboratory management, P.J.G.M. de Wit for critical reading of the manuscript.

## AUTHOR CONTRIBUTIONS

YI designed the study and managed the research funds. KM, HS and TaS performed RNA-seq and analyzed the data. KM, MK, MO, and TK analyzed NLPs. ToS conducted electron microscopy and EI developed a microscopic observation protocol. KM and KS studied the Ecp2 protein. KM and YI wrote the manuscript. All authors reviewed and approved the final version.

## DATA AVAILABILITY

Genomic sequences for *C. fulvum* strain Race5_Kim and *H. pulvinata* strain 414-3 are available as accessions PRJNA565804 and PRJDB8178 at NCBI BioProject, respectively (Sushida *et al*., 2019; Zaccaron & Stergiopoulos, 2024). Tomato genome data were obtained from the Solanaceae Genomics Network (Hosmani *et al*., 2019). Additional genomes are accessible via JGI MycoCosm (mycocosm.jgi.doe.gov) and the DDBJ/EMBL/GenBank repository (www.ncbi.nlm.nih.gov). Custom scripts and supporting datasets were deposited at Figshare (doi.org/10.6084/m9.figshare.30265009).

## SUPPORTING INFORMATION

**Fig. S1** Experimental treatments and principal component analysis (PCA) of transcripts from the mycoparasite *Hansfordia pulvinata* (Hp), the pathogen *Cladosporium fulvum* (Cf), and tomato plant, *Solanum lycopersicum*.

**Fig. S2** Secondary metabolite biosynthesis gene clusters in the mycoparasite *Hansfordia pulvinata*.

**Fig. S3** Phylogenetic analysis and functional characterization of proteins in fungal species.

**Fig. S4** Structure of HpNlp1.

**Fig. S5** Responses in plant leaves triggered by HpNlp1 protein.

**Fig. S6** Heat map and hierarchical clustering of effector genes expressed in *Cladosporium fulvum*.

**Fig. S7** Structural comparative analysis of Ecp2 with ZtKP4 and UmVKP4.

**Fig. S8** Differentially expressed genes (DEGs) in KEGG pathways detected in tomato plants.

**Table S1** Primer sequences used in this study.

**Table S2** Summary statistics of RNA-seq data and mapping results.

**Table S3** Functional annotation of all genes in *Hansfordia pulvinata* genome.

**Table S4** Predicted gene clusters involved in secondary metabolite biosynthesis in *Hansfordia pulvinata*.

**Table S5** Small secreted protein-coding genes (≤300 aa) in *Hansfordia pulvinata*.

**Table S6** Key genes encoding secondary metabolism enzymes in *Cladosporium fulvum*.

**Table S7** Effector genes identified in *Cladosporium fulvum*.

**Table S8** Structural homologs of Ecp2 protein identified from PDB25 using the Dali server.

**Table S9** Species possessing homologs of Ecp2 and KP4 proteins.

**Table S10** Log_2_ fold-change for defensin-like genes in tomato plants.

