## Supplementary material for "Transcriptomic insights into the tritrophic plant–pathogen–mycoparasite interaction reveal coordinated reprogramming fungal secretomes and plant amino acid metabolism": Figure S1-8

***New Phytologist* Supporting Information**

Article title: Tripartite plant–pathogen–mycoparasite interactions uncover novel mechanisms involving fungal secretomes and plant amino acid metabolisms

Authors: Kazuya Maeda, Mariko Kouda, Mai Ohara, Takumi Kawase, Koki Saito, Eishin Iwao, Hirotoshi Sushida, Tomoko Suzuki, Takuya Sumita, Yuichiro Iida

The following Supporting Information is available for this article:

**Fig. S1** Experimental treatments and principal component analysis (PCA) of transcripts from the mycoparasite *Hansfordia pulvinata* (Hp), the pathogen *Cladosporium fulvum* (Cf), and tomato plant, *Solanum lycopersicum*.

**Fig. S2** Secondary metabolite biosynthesis gene clusters in the mycoparasite *Hansfordia pulvinata*.

**Fig. S3** Phylogenetic analysis and functional characterization of proteins in fungal species.

**Fig. S4** Structure of HpNlp1.

**Fig. S5** Responses in plant leaves triggered by HpNlp1 protein.

**Fig. S6** Heat map and hierarchical clustering of effector genes expressed in *Cladosporium fulvum*.

**Fig. S7** Comparative structural analysis of Ecp2 with ZtKP4 and UmVKP4 proteins.

**Fig. S8** KEGG pathway annotations for differentially expressed genes (DEGs) detected in tomato plants.

**Table S1** Primer sequences used in this study.

**Table S2** Summary statistics of RNA-seq data and mapping results.

**Table S3** Functional annotation of all genes in *Hansfordia pulvinata* genome.

**Table S4** Predicted gene clusters involved in secondary metabolite biosynthesis in *Hansfordia pulvinata*.

**Table S5** Small secreted protein-coding genes (≤ 300 aa) in *Hansfordia pulvinata*.

**Table S6** Key genes encoding secondary metabolism enzymes in *Cladosporium fulvum*.

**Table S7** Effector genes identified in *Cladosporium fulvum*.

**Table S8** Structural homologs of Ecp2 protein identified from PDB25 using the Dali server.

**Table S9** Species with homologs of Ecp2 and KP4 proteins.

**Table S10** $Log_2$ fold change of defensin-like genes in tomato plants.

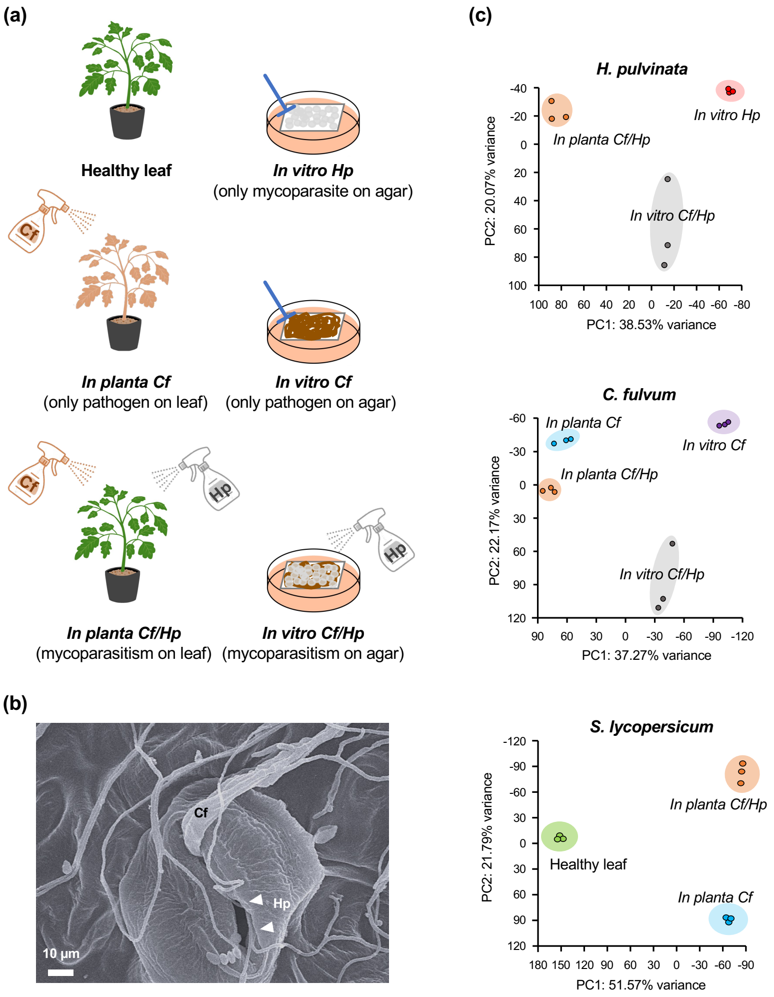


**Fig. S1** Experimental treatments and principal component analysis (PCA) of transcripts from the mycoparasite *Hansfordia pulvinata* (Hp), the pathogen *Cladosporium fulvum* (Cf), and tomato plant, *Solanum lycopersicum*. (a) Tomato leaves were sprayed with a spore suspension of *C. fulvum* (*on planta Cf*). Lesions caused by *C. fulvum* were subsequently sprayed with a spore suspension of *H. pulvinata* (*on planta Cf/Hp*). *H. pulvinata* (*in vitro Hp*) and *C. fulvum* (*in vitro Cf*) were cultured on nylon membranes on PDA, then transferred to water agar. *H. pulvinata* also parasitized *C. fulvum* on water agar (*in vitro Cf/Hp*). (b) Thick hypha of *C. fulvum* and thin hyphae of *H. pulvinata* (white arrows) entering stoma on tomato leaf in the *on planta Cf/Hp* treatment. (c) PCA plot of three biological replicates for each treatment. Axis labels indicate the percentage of total variance explained by the corresponding principal components.

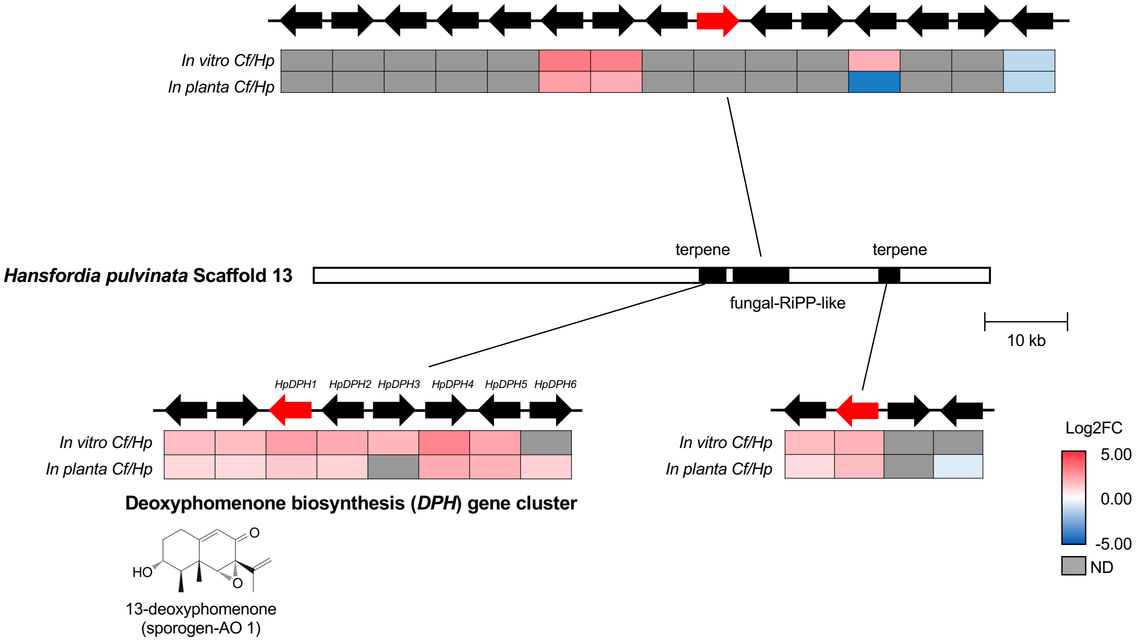


**Fig. S2** Secondary metabolite biosynthesis gene clusters in the mycoparasite *Hansfordia pulvinata*. Scaffold 13 contains three terpenoid biosynthesis gene clusters. Potential secondary metabolite biosynthetic gene clusters were predicted using AntiSMASH 7.1.0. Terpenoid backbone biosynthesis genes are highlighted with red arrows. The biosynthesis of 13-deoxyphomenone (sporogen-AO 1) has been reported by Maeda *et al*. (2025). log2FC = $\log_2$ Fold-change.

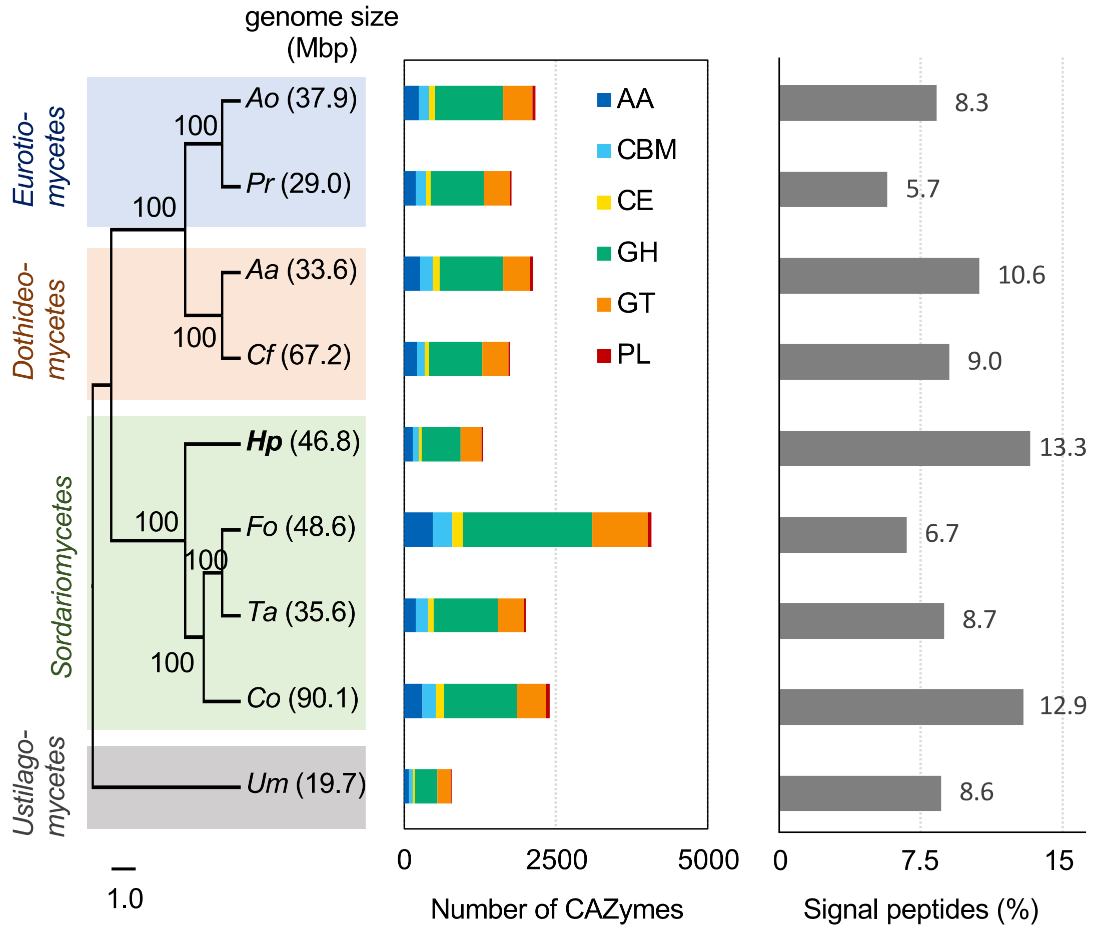


**Fig. S3** Phylogenetic analysis and functional characterization of proteins in fungal species. The phylogenetic species tree was constructed using RAxML based on 100 monocore genes (single-copy homologs in each species) with 1,000 bootstrap replicates. Species abbreviations: Ao, *Aspergillus oryzae*; Pr, *Penicillium roqueforti*; Aa, *Alternaria alternata*; Cf, *Cladosporium fulvum*; Hp, *Hansfordia pulvinata*; Fo, *Fusarium oxysporum*; Ta, *Trichoderma atroviride*; Co, *Colletotrichum orbiculare*; Um, *Ustilago maydis.* The activity of the CAZymes in each species belong to six categories: AA, auxiliary activities; CBM, carbohydrate-binding modules; CE, carbohydrate esterases; GH, glycoside hydrolases; GT, glycosyltransferases; PL, polysaccharide lyases. Signal peptides were predicted using SignalP 6.0 and are shown as the percentage of total genes to the right of each gray bar.

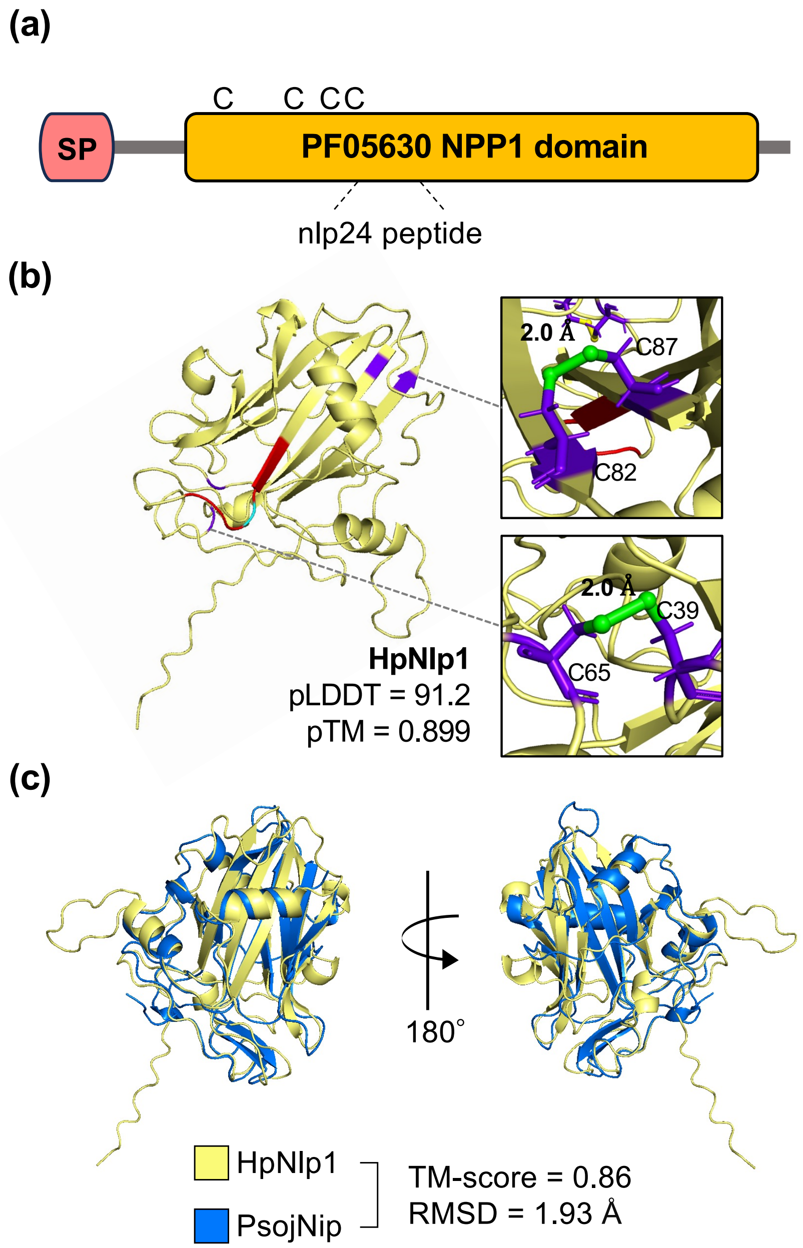


**Fig. S4** Structure of HpNlp1. (a) Protein domain organization analyzed using the InterPro database. SP, signal peptide; C, cysteine residues. (b) Predicted 3D structure of HpNlp1 generated by AlphaFold2. The conserved GHRHDWE motif is highlighted in red; nonconserved regions are in light blue. Cysteine residues are in purple, disulfide bonds in green, and numbers between adjacent cysteines are sulfur atom distances. Predicted pLDDT and pTM values are shown. (c) Structural comparison between HpNlp1 and PsojNip proteins. TM-score and RMSD were calculated using the TM-align program.

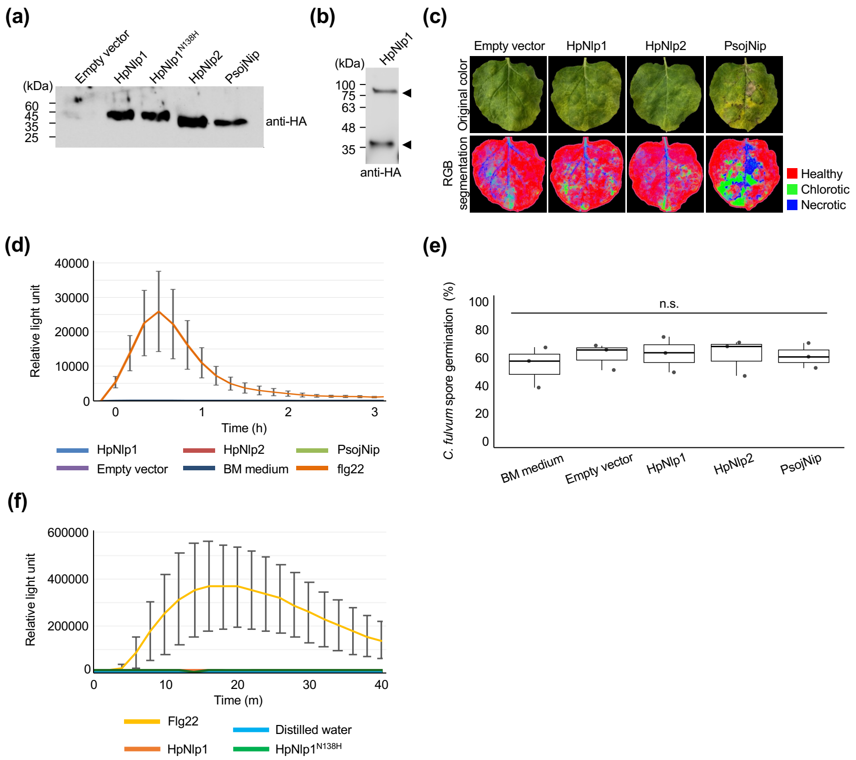


**Fig. S5** Responses in plant leaves triggered by HpNlp1 protein. (a) NLP proteins secreted from *Pichia pastoris* were detected using an anti-HA antibody and visualized by enhanced chemiluminescence. (b) Monomer and dimer bands are indicated by arrowheads. (c) Pixel classification to quantify surface area for leaf colors using the machine-learning-based software Ilastik. The program was trained interactively by manually labeling healthy (red), chlorotic (green), and necrotic (blue) regions on representative images of leaves infiltrated with PsojNip. Classification was performed using three biological replicates per treatment. Color ratios were calculated based on the number of RGB pixels and are represented in the bar graph in Fig. 3e. (d) Reactive oxygen species (ROS) measured as relative luminescence units in leaf disks of *Nicotiana benthamiana* after 3 h treatment with 500 nM flg22 or NLP proteins. All experiments were done three times. (e) Percentage of spore germination of *C. fulvum* after 24-h treatment with NLP proteins. For each of three independent replicate, 200 spores were assessed. None of the comparisons were statistically significant (n.s.) in a one-way ANOVA with post hoc Tukey's HSD test. (f) ROS burst assay in *Arabidopsis thaliana*. Leaf disks were treated with 500 nM flg22 or nlp24 peptides (HpNlp1: GHRNDWEHIAVWTRDGAAVAVA; HpNlp1$^{N138H}$: GHRHDWEHIAVWTRDGAAVAVA) for 40 min, and relative luminescence units were measured. All experiments were done three times.

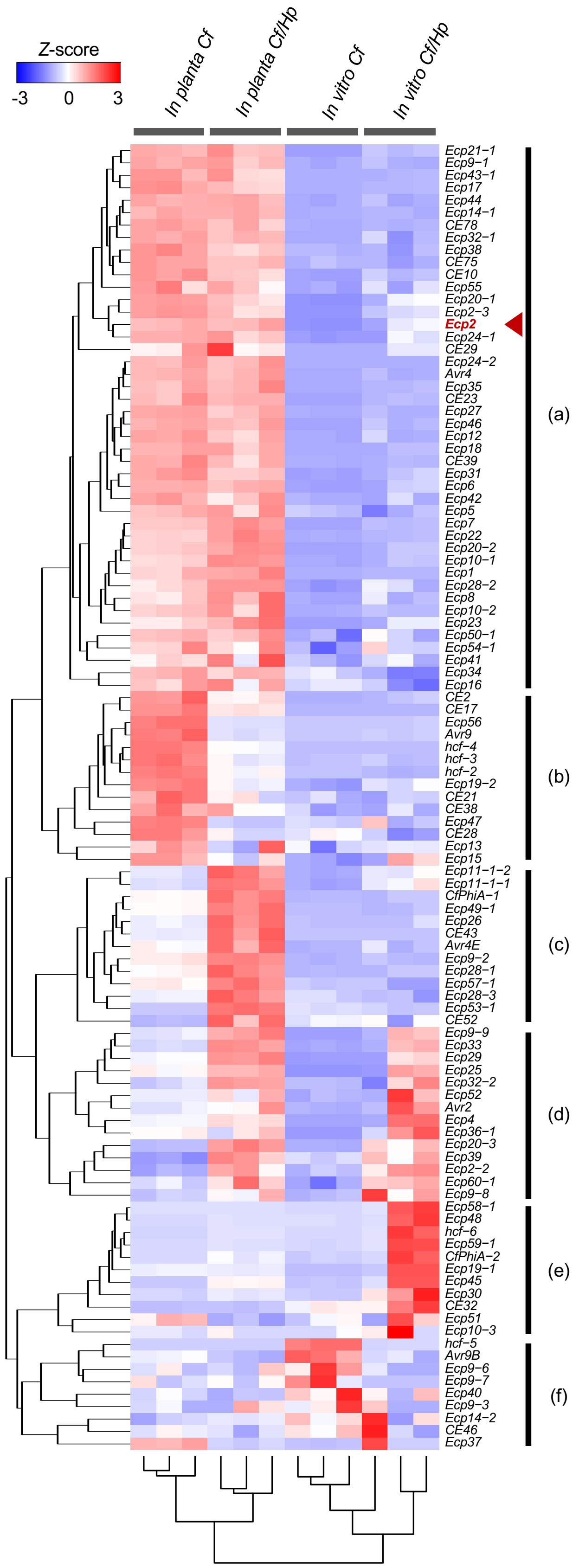


**Fig. S6** Heat map and hierarchical clustering of effector genes expressed in *Cladosporium fulvum*. Expression levels of 105 effector genes were $\log_2$ (TPM + 1)-normalized, converted to raw *Z*-scores for each gene and divided into six subsets (a–f). Genes in subsets (a) and (b) were highly expressed in tomato leaves regardless of the presence of the mycoparasite *H. pulvinata.* Expression of genes in subsets (c), (d), and (e) was induced in the presence of *H. pulvinata.* The red arrowhead marks *Ecp2*.

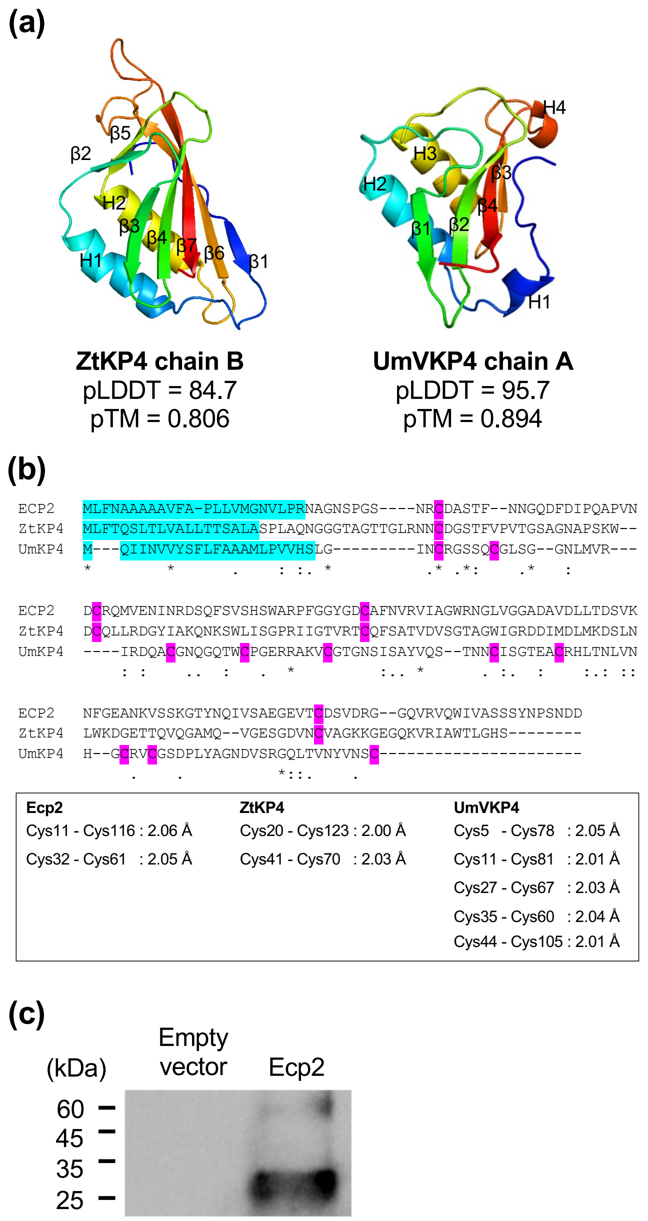


**Fig. S7** Comparative structural analysis of Ecp2 with ZtKP4 and UmVKP4 proteins. (a) Predicted structures of Ecp2, ZtKP4, and UmVKP4 generated by AlphaFold2. Structural similarity of Ecp2 relative to KP4 was evaluated using TM-score and RMSD calculated with the TM-align program. (b) Amino acid sequence alignment of Ecp2 and KP4. Signal peptides are highlighted in blue, cysteine residues in magenta. Symbols indicate conserved residues: (*) fully conserved, (:) strongly similar, (.) weakly similar. Predicted disulfide bonds in the three-dimensional structures are indicated within boxes. (c) Western blot of Ecp2 expressed in *Escherichia coli* BL21 strain. The protein was detected using an anti-His antibody.

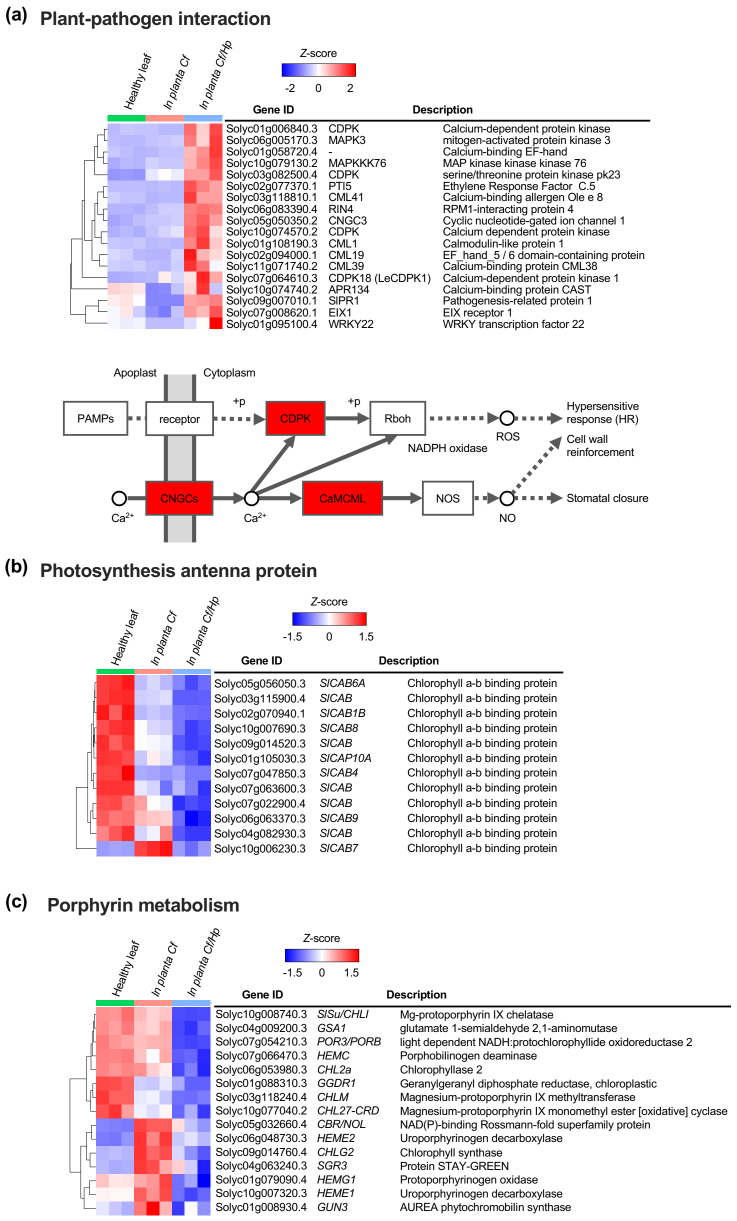


**Fig. S8** KEGG pathway annotations for differentially expressed genes (DEGs) detected in tomato plants. (a) *Z*-score heat maps and schematic of enriched DEGs in the plant–pathogen interaction in the presence of *Hansfordia pulvinata*, based on normalized TPM values. The pathway schematic was adapted from KEGG pathway model images (www.kegg.jp/kegg/pathway.html). Upregulated DEGs in tomato are in red boxes. (b, c) *Z*-score heat maps of enriched DEGs in the photosynthesis antenna protein pathway (b) and porphyrin metabolism pathway (c) in the presence of the mycoparasite *H. pulvinata*, based on normalized TPM values.
