## Supplementary material for "Transcriptomic insights into the tritrophic plant–pathogen–mycoparasite interaction reveal coordinated reprogramming fungal secretomes and plant amino acid metabolism": Table S1-10

Supplementary Table S1. Primer sequences used in this study.

| **Primer name** | **Primer sequence (5’-3’)** | **Purpose** |
|---|---|---|
| pET28b_F | GCCGCTGCTGTGATGATGATGATGATG | Linearization of pET28b+ |
| pET28b_R | GATCCGGCTGCTAACAAAGCCCGAAAG | |
| ECP2_F | GCTTTGTTAGCAGCCGGATCCTAGTCATCGTTGGACGG | Amplification of *ECP2* gene |
| ECP2_R | ATCATCATCACAGCAGCGGCAACGCTGGCAACTC | |
| pet_chk_F | CCGGATATAGTTCCTCCTTTCAGC | Detection of insertion sequence on pET28b+ |
| pet_chk_R | CACTATAGGGGAATTGTGAGCG | |
| pPIC9_AOX1_F1 | GAAGCTGTCATCGGTTACTCAGA | Detection of insertion sequence on pPIC9K |
| pPIC9_AOX1_R1 | CTCGTAAGTGCCCAACTTGAAC | |
| smaI_NLPF | GCCGGCCATTATGGCCCATGGGATTTGTTCTCTTTTCA | Amplification of NLP genes |
| smaI_NLPR | CGAGGCGGCCGATATCCCTTATTTTTCAAATTGAGGATG | |
| OX10_F | CAATCACAGTGTTGGCTTGC | Detection of insertion sequence on psfinx |
| N31_R | GACCCTATGGGCTGTGTTG | |

Supplementary Table S2. Summary statistics of RNA-seq data and mapping results.

| Sample name | Organisms | Total no. reads | Uniquely mapped no. reads | Uniquely mapped (%) |
|---|---|---|---|---|
| Healthy leaf_1 | *Solanum lycopersicum* | 35650034 | 31195431 | 87.505 |
| Healthy leaf_2 | *Solanum lycopersicum* | 34139852 | 29556835 | 86.576 |
| Healthy leaf_3 | *Solanum lycopersicum* | 31265696 | 27020308 | 86.422 |
| *in planta Cf_1* | *Solanum lycopersicum* | 27786553 | 5704640 | 20.530 |
| *in planta Cf_2* | *Solanum lycopersicum* | 31230663 | 6331914 | 20.275 |
| *in planta Cf_3* | *Solanum lycopersicum* | 26555549 | 6129447 | 23.082 |
| *in planta Cf/Hp_1* | *Solanum lycopersicum* | 30034779 | 3098914 | 10.318 |
| *in planta Cf/Hp_2* | *Solanum lycopersicum* | 30646255 | 4217836 | 13.763 |
| *in planta Cf/Hp_3* | *Solanum lycopersicum* | 31922549 | 3795064 | 11.888 |
| *in vitro Cf* _1 | *Cladosporium fulvum* | 8050429 | 6977797 | 86.676 |
| *in vitro Cf* _2 | *Cladosporium fulvum* | 9682814 | 8417685 | 86.934 |
| *in vitro Cf* _3 | *Cladosporium fulvum* | 5559785 | 4710943 | 84.732 |
| *in vitro Cf/Hp* _1 | *Cladosporium fulvum* | 5981499 | 1015894 | 16.984 |
| *in vitro Cf/Hp* _2 | *Cladosporium fulvum* | 6882711 | 1052615 | 15.294 |
| *in vitro Cf/Hp* _3 | *Cladosporium fulvum* | 38102224 | 18580666 | 48.765 |
| *in planta Cf_1* | *Cladosporium fulvum* | 28442850 | 20350042 | 71.547 |
| *in planta Cf_2* | *Cladosporium fulvum* | 31896295 | 23073556 | 72.339 |
| *in planta Cf_3* | *Cladosporium fulvum* | 27084934 | 18793425 | 69.387 |
| *in planta Cf/Hp_1* | *Cladosporium fulvum* | 30619379 | 19009433 | 62.083 |
| *in planta Cf/Hp_2* | *Cladosporium fulvum* | 31245982 | 18929296 | 60.582 |
| *in planta Cf/Hp_3* | *Cladosporium fulvum* | 32527338 | 20365382 | 62.610 |
| *in vitro Hp* _1 | *Hansfordia pulvinata* | 8923362 | 7386504 | 82.777 |
| *in vitro Hp* _2 | *Hansfordia pulvinata* | 6911253 | 5726956 | 82.864 |
| *in vitro Hp* _3 | *Hansfordia pulvinata* | 7203849 | 6348700 | 88.129 |
| *in vitro Cf/Hp* _1 | *Hansfordia pulvinata* | 8923362 | 2134533 | 23.921 |
| *in vitro Cf/Hp* _2 | *Hansfordia pulvinata* | 6911253 | 3772118 | 54.579 |
| *in vitro Cf/Hp* _3 | *Hansfordia pulvinata* | 36448769 | 18580666 | 50.977 |
| *in planta Cf/Hp_1* | *Hansfordia pulvinata* | 29928210 | 6131174 | 20.486 |
| *in planta Cf/Hp_2* | *Hansfordia pulvinata* | 30491584 | 5380508 | 17.646 |
| *in planta Cf/Hp_3* | *Hansfordia pulvinata* | 31817738 | 5671251 | 17.824 |

Supplementary Table S3. Functional annotation of all genes in *Hansfordia pulvinata* genome.

| GeneID | Description [a] | Identity (%) | E_value |
|---|---|---|---|
| DIP_10000001 | hypothetical protein TCE0_060f19332 [Talaromyces cellulolyticus] | 58.7 | 6.0E-36 |
| DIP_10000002 | hypothetical protein H633G_11182 [Metarhizium anisopliae BRIP 53284] | 53.1 | 1.0E-26 |
| DIP_10000003 | NA | | |
| DIP_10000004 | NA | | |
| DIP_10000005 | NA | | |
| DIP_10000006 | hypothetical protein CSUB01_04592 [Colletotrichum sublineola] | 30.5 | 6.0E-10 |
| DIP_10000007 | PKS1, putative Phthiocerol synthesis polyketide synthase type I PpsA [Lepidopterella palustris CBS 459.81] (Dothideomycetes) | 47.3 | 2.0E-107 |
| DIP_10000008 | hypothetical protein MSSIT_3509 [Methanosarcina siciliae T4/M] | 50.0 | 2.0E-11 |
| DIP_10000009 | hypothetical protein TCE0_060f19332 [Talaromyces cellulolyticus] | 58.7 | 6.0E-36 |
| DIP_10000010 | hypothetical protein, partial [bacterium UASB270] | 50.5 | 2.0E-18 |
| DIP_10000011 | PKS2<br>beta-ketoacyl synthase [Trichoderma harzianum] | 48.9 | 3.0E-89 |
| DIP_10000012 | NA | | |
| DIP_10000013 | NA | | |
| DIP_10000014 | NA | | |
| DIP_10000015 | NA | | |
| DIP_10000016 | NA | | |
| DIP_10000017 | NA | | |
| DIP_10000018 | NA | | |
| DIP_10000019 | NA | | |
| DIP_10000020 | NA | | |
| DIP_10000021 | NA | | |
| DIP_10000022 | NA | | |
| DIP_10000023 | NA | | |
| DIP_10000024 | NA | | |
| DIP_10000025 | NA | | |
| DIP_10000026 | hypothetical protein NECHADRAFT_55847 [Nectria haematococca mpVI 77-13-4] | 55.1 | 1.0E-76 |
| DIP_10000027 | hypothetical protein PTT_13697 [Pyrenophora teres f. teres 0-1] | 34.5 | 4.0E-18 |
| DIP_10000028 | NA | | |
| DIP_10000029 | NA | | |
| DIP_10000030 | predicted protein [Histoplasma capsulatum NAm1] | 39.8 | 1.0E-17 |
| DIP_10000031 | NA | | |
| DIP_10000032 | NA | | |
| DIP_10000033 | NA | | |
| DIP_10000034 | NA | | |
| DIP_10000035 | NA | | |
| DIP_10000036 | NA | | |
| DIP_10000037 | NA | | |
| DIP_10000038 | NA | | |
| DIP_10000039 | NA | | |
| DIP_10000040 | NA | | |
| DIP_10000041 | NA | | |
| DIP_10000042 | NA | | |
| DIP_10000043 | hypothetical protein A1O5_10006 [Cladophialophora psammophila CBS 110553] | 43.9 | 3.0E-44 |
| DIP_10000044 | putative -like protein [Neofusicoccum parvum UCRNP2] | 37.3 | 2.0E-19 |
| DIP_10000045 | nitroreductase protein [Metarhizium robertsii ARSEF 23] | 60.8 | 8.0E-83 |
| DIP_10000046 | NA | | |
| DIP_10000047 | PKS3<br>beta-ketoacyl synthase [Trichoderma harzianum] | 48.9 | 2.0E-89 |
| DIP_10000048 | putative reverse transcriptase [Macrophomina phaseolina MS6] | 35.3 | 2.0E-22 |
| DIP_10000049 | NA | | |
| DIP_10000050 | hypothetical protein PFICI_06087 [Pestalotiopsis fici W106-1] | 40 | 1.0E-64 |
| DIP_10000051 | group 1 glycosyl transferase [cyanobacterium PCC 7702] | 32.5 | 5.0E-13 |
| DIP_10000052 | NA | | |
| DIP_10000053 | NA | | |
| DIP_10000054 | NA | | |
| DIP_10000055 | hypothetical protein CSUB01_10637 [Colletotrichum sublineola] | 36.7 | 5.0E-39 |
| DIP_10000056 | NA | | |
| DIP_10000057 | hypothetical protein HMPREF1624_08178 [Sporothrix schenckii ATCC 58251] | 47.8 | 0.0 |
| DIP_10000058 | hypothetical protein ACRE_030110 [Acremonium chrysogenum ATCC 11550] | 31.8 | 1.0E-62 |
| DIP_10000059 | putative Mpv17/PMP22 family protein [Colletotrichum sublineola] | 41.6 | 3.0E-60 |
| DIP_10000060 | putative hrf1 domain-containing protein [Togninia minima UCRPA7] | 73.9 | 8.0E-171 |
| DIP_10000061 | putative signal recognition particle receptor subunit alpha protein [Eutypa lata UCREL1] | 68 | 0.0 |
| DIP_10000062 | hypothetical protein PFICI_03240 [Pestalotiopsis fici W106-1] | 57.9 | 6.0E-132 |
| DIP_10000063 | Pre-mRNA-splicing factor slt-11 [Pestalotiopsis fici W106-1] | 72.6 | 0.0 |
| DIP_10000064 | Mitochondrial zinc maintenance protein 1 [Pestalotiopsis fici W106-1] | 65.5 | 4.0E-41 |
| DIP_10000065 | putative zuotin protein [Eutypa lata UCREL1] | 82.5 | 0.0 |
| DIP_10000066 | NA | | |
| DIP_10000067 | hypothetical protein DOTSEDRAFT_75436 [Dothistroma septosporum NZE10] | 30.1 | 4.0E-25 |
| DIP_10000068 | putative endo- -beta-xylanase protein [Eutypa lata UCREL1] | 61.3 | 9.0E-116 |
| DIP_10000069 | hypothetical protein PFICI_13019 [Pestalotiopsis fici W106-1] | 67.7 | 1.0E-171 |
| DIP_10000070 | NA | | |
| DIP_10000071 | NA | | |
| DIP_10000072 | hypothetical protein Z517_09311 [Fonsecaea pedrosoi CBS 271.37] | 37.7 | 4.0E-26 |
| DIP_10000073 | NA | | |
| DIP_10000074 | hypothetical protein TD95_002244 [Thielaviopsis punctulata] | 42.3 | 5.0E-15 |
| DIP_10000075 | NA | | |
| DIP_10000076 | NA | | |
| DIP_10000077 | NA | | |
| DIP_10000078 | NA | | |
| DIP_10000079 | NA | | |
| DIP_10000080 | NA | | |
| DIP_10000081 | NA | | |
| DIP_10000082 | NA | | |
| DIP_10000083 | hypothetical protein AUEXF2481DRAFT_29316 [Aureobasidium subglaciale EXF-2481] | 33.9 | 3.0E-51 |
| DIP_10000084 | NA | | |
| DIP_10000085 | hypothetical protein CGGC5_14879 [Colletotrichum gloeosporioides Nara gc5] | 32.1 | 2.0E-09 |
| DIP_10000086 | NA | | |
| DIP_10000087 | NA | | |
| DIP_10000088 | NA | | |
| DIP_10000089 | NA | | |
| DIP_10000090 | NA | | |
| DIP_10000091 | NA | | |
| DIP_10000092 | hypothetical protein GGTG_06027 [Gaeumannomyces graminis var. tritici R3-111a-1] | 31 | 3.0E-44 |
| DIP_10000093 | NA | | |
| DIP_10000094 | hypothetical protein A1O3_10456 [Capronia epimyces CBS 606.96] | 64.2 | 2.0E-137 |
| DIP_10000095 | hypothetical protein PFICI_10677 [Pestalotiopsis fici W106-1] | 70.9 | 1.0E-156 |
| DIP_10000096 | hypothetical protein PFICI_10676 [Pestalotiopsis fici W106-1] | 67.2 | 4.0E-126 |
| DIP_10000097 | hexose transporter protein [Colletotrichum gloeosporioides Nara gc5] | 63.6 | 2.0E-143 |
| DIP_10000098 | hypothetical protein FOCG_05198 [Fusarium oxysporum f. sp. radicis-lycopersici 26381] | 45.2 | 7.0E-40 |
| DIP_10000099 | hypothetical protein PFICI_10673 [Pestalotiopsis fici W106-1] | 54.3 | 2.0E-126 |
| DIP_10000100 | hypothetical protein A1O3_10461 [Capronia epimyces CBS 606.96] | 60.1 | 5.0E-116 |
| DIP_10000101 | NA | | |
| DIP_10000102 | NA | | |
| DIP_10000103 | hypothetical protein PV07_12505 [Cladophialophora immunda] | 70.5 | 4.0E-92 |
| DIP_10000104 | kinase-related protein [Colletotrichum gloeosporioides Nara gc5] | 58.3 | 1.0E-71 |
| DIP_10000105 | NA | | |
| DIP_10000106 | NA | | |
| DIP_10000107 | NA | | |
| DIP_10000108 | NA | | |
| DIP_10000109 | delta-aminolevulinic acid dehydratase [Emmonsia crescens UAMH 3008] | 60.5 | 1.0E-12 |
| DIP_10000110 | NA | | |
| DIP_10000111 | NA | | |
| DIP_10000112 | NA | | |
| DIP_10000113 | NA | | |
| DIP_10000114 | NA | | |
| DIP_10000115 | NA | | |
| DIP_10000116 | hypothetical protein TCE0_060f19332 [Talaromyces cellulolyticus] | 58.7 | 6.0E-36 |
| DIP_10000117 | hypothetical protein H633G_11182 [Metarhizium anisopliae BRIP 53284] | 53.1 | 1.0E-26 |
| DIP_10000118 | NA | | |
| DIP_10000119 | hypothetical protein CGGC5_13047 [Colletotrichum gloeosporioides Nara gc5] | 63.1 | 2.0E-152 |
| DIP_10000120 | hypothetical protein FOXB_06689 [Fusarium oxysporum Fo5176] | 44.6 | 5.0E-27 |
| DIP_10000121 | hypothetical protein TRIATDRAFT_52247 [Trichoderma atroviride IMI 206040] | 59.7 | 5.0E-57 |
| DIP_10000122 | hypothetical protein HIM_10180 [Hirsutella minnesotensis 3608] | 44.6 | 5.0E-31 |
| DIP_10000123 | NA | | |
| DIP_10000124 | PKS4<br>beta-ketoacyl synthase [Trichoderma harzianum] | 48.9 | 1.0E-89 |
| DIP_10000125 | NA | | |
| DIP_10000126 | NA | | |
| DIP_10000127 | NA | | |
| DIP_10000128 | NA | | |
| DIP_10000129 | NA | | |
| DIP_10000130 | NA | | |
| DIP_10000131 | NA | | |
| DIP_10000132 | NA | | |
| DIP_10000133 | NA | | |
| DIP_10000134 | NA | | |
| DIP_10000135 | NA | | |
| DIP_10000136 | NA | | |
| DIP_10000137 | NA | | |
| DIP_10000138 | NA | | |
| DIP_10000139 | hypothetical protein GGTG_06027 [Gaeumannomyces graminis var. tritici R3-111a-1] | 41.3 | 2.0E-87 |
| DIP_10000140 | NA | | |
| DIP_10000141 | NA | | |
| DIP_10000142 | hypothetical protein VHEMI02445 [Torrubiella hemipterigena] | 64.2 | 0.0 |
| DIP_10000143 | NA | | |
| DIP_10000144 | NA | | |
| DIP_10000145 | hypothetical protein CGGC5_13047 [Colletotrichum gloeosporioides Nara gc5] | 62.2 | 5.0E-144 |
| DIP_10000146 | NA | | |
| DIP_10000147 | NA | | |
| DIP_10000148 | reverse transcriptase, RNaseH [Metarhizium guizhouense ARSEF 977] | 46.6 | 4.0E-28 |
| DIP_10000149 | NA | | |
| DIP_10000150 | NA | | |
| DIP_10000151 | NA | | |
| DIP_10000152 | NA | | |
| DIP_10000153 | putative frag1 dram sfk1 family protein [Eutypa lata UCREL1] | 62.2 | 2.0E-109 |
| DIP_10000154 | hypothetical protein PFICI_07893 [Pestalotiopsis fici W106-1] | 51.1 | 0.0 |
| DIP_10000155 | putative tyrosine serine protein phosphatase protein [Eutypa lata UCREL1] | 69.4 | 1.0E-150 |
| DIP_10000156 | putative actin-binding protein [Eutypa lata UCREL1] | 71.2 | 0.0 |
| DIP_10000157 | putative cyclase dehydrase family protein [Eutypa lata UCREL1] | 49.3 | 3.0E-58 |
| DIP_10000158 | NA | | |
| DIP_10000159 | putative kinetoplast-associated protein kap-like protein [Eutypa lata UCREL1] | 55.3 | 1.0E-103 |
| DIP_10000160 | putative tetratricopeptide repeat domain-containing protein [Eutypa lata UCREL1] | 64.9 | 4.0E-152 |
| DIP_10000161 | ap-3 complex subunit delta [Colletotrichum orbiculare MAFF 240422] | 62.2 | 0.0 |
| DIP_10000162 | NA | | |
| DIP_10000163 | NA | | |
| DIP_10000164 | putative rho-gtpase-activating protein 8 protein [Eutypa lata UCREL1] | 75.2 | 0.0 |
| DIP_10000165 | putative dipeptidase 1 protein [Eutypa lata UCREL1] | 57.1 | 2.0E-174 |
| DIP_10000166 | putative mitochondrial 37s ribosomal protein sws2 protein [Eutypa lata UCREL1] | 79 | 9.0E-68 |
| DIP_10000167 | putative peptidyl-prolyl cis-trans isomerase b protein [Eutypa lata UCREL1] | 62.6 | 6.0E-119 |
| DIP_10000168 | predicted protein [Nectria haematococca mpVI 77-13-4] | 82 | 1.0E-74 |
| DIP_10000169 | putative er protein big1 protein [Eutypa lata UCREL1] | 57.5 | 8.0E-97 |
| DIP_10000170 | Ribosomal RNA-processing protein-like protein [Acremonium chrysogenum ATCC 11550] | 61 | 9.0E-83 |
| DIP_10000171 | putative actin lateral binding protein [Eutypa lata UCREL1] | 88.2 | 3.0E-91 |
| DIP_10000172 | hypothetical protein UCREL1_3106 [Eutypa lata UCREL1] | 64.4 | 3.0E-103 |
| DIP_10000173 | hypothetical protein M419DRAFT_90299 [Trichoderma reesei RUT C-30] | 46.2 | 0.0 |
| DIP_10000174 | putative rrna-processing protein fcf1 protein [Eutypa lata UCREL1] | 72.9 | 6.0E-117 |
| DIP_10000175 | putative csi2 protein [Eutypa lata UCREL1] | 68 | 4.0E-110 |
| DIP_10000176 | putative abc transporter family protein [Eutypa lata UCREL1] | 75.4 | 0.0 |
| DIP_10000177 | putative ino80 chromatin remodeling complex ies1 protein [Eutypa lata UCREL1] | 66.8 | 0.0 |
| DIP_10000178 | hypothetical protein V502_02473 [Pseudogymnoascus pannorum VKM F-4520 (FW-2644)] | 76.9 | 0.0 |
| DIP_10000179 | probable multicatalytic endopeptidase complex chain PRE1 [Neurospora crassa] | 89.7 | 5.0E-137 |
| DIP_10000180 | WD domain, G-beta repeat containing protein [Metarhizium album ARSEF 1941] | 55.9 | 0.0 |
| DIP_10000181 | hypothetical protein PFICI_12101 [Pestalotiopsis fici W106-1] | 72.6 | 0.0 |
| DIP_10000182 | putative nucleosome assembly protein [Colletotrichum sublineola] | 82.5 | 0.0 |
| DIP_10000183 | putative d-3-phosphoglycerate dehydrogenase 1 protein [Eutypa lata UCREL1] | 79.5 | 0.0 |
| DIP_10000184 | putative myosin myo2 protein [Eutypa lata UCREL1] | 87.9 | 0.0 |
| DIP_10000185 | putative pos9-activating factor fap7 protein [Togninia minima UCRPA7] | 81.8 | 5.0E-105 |
| DIP_10000186 | putative inosine triphosphate pyrophosphatase protein [Eutypa lata UCREL1] | 78.1 | 7.0E-107 |
| DIP_10000187 | Serine--tRNA ligase [Pestalotiopsis fici W106-1] | 81.1 | 0.0 |
| DIP_10000188 | GTP-binding protein rhb1 [Pestalotiopsis fici W106-1] | 85.1 | 2.0E-93 |
| DIP_10000189 | hypothetical protein PFICI_12102 [Pestalotiopsis fici W106-1] | 40.5 | 1.0E-131 |
| DIP_10000190 | NA | | |
| DIP_10000191 | putative uracil phosphoribosyltransferase protein [Eutypa lata UCREL1] | 93.4 | 1.0E-169 |
| DIP_10000192 | putative transcription elongation protein [Eutypa lata UCREL1] | 73.6 | 2.0E-58 |
| DIP_10000193 | hypothetical protein FPSE_09441 [Fusarium pseudograminearum CS3096] | 71.2 | 0.0 |
| DIP_10000194 | putative 40s ribosomal protein s3ae protein [Eutypa lata UCREL1] | 95.7 | 4.0E-175 |
| DIP_10000195 | Pc22g00200 [Penicillium rubens Wisconsin 54-1255] | 63.9 | 4.0E-171 |
| DIP_10000196 | NA | | |
| DIP_10000197 | NA | | |
| DIP_10000198 | putative glycosyl hydrolase family protein [Eutypa lata UCREL1] | 66.7 | 0.0 |
| DIP_10000199 | hypothetical protein A1O9_06711 [Exophiala aquamarina CBS 119918] | 64.9 | 0.0 |
| DIP_10000200 | hypothetical protein GGTG_05621 [Gaeumannomyces graminis var. tritici R3-111a-1] | 31.2 | 2.0E-10 |
| DIP_10000201 | hypothetical protein H634G_08057 [Metarhizium anisopliae BRIP 53293] | 43.4 | 4.0E-24 |
| DIP_10000202 | NA | | |
| DIP_10000203 | pyoverdine/dityrosine biosynthesis protein, putative [Aspergillus clavatus NRRL 1] | 48.3 | 0.0 |
| DIP_10000204 | hypothetical protein SETTUDRAFT_42150 [Setosphaeria turcica Et28A] | 41.4 | 2.0E-74 |
| DIP_10000205 | hypothetical protein SNOG_14726 [Phaeosphaeria nodorum SN15] | 56.5 | 0.0 |
| DIP_10000206 | hypothetical protein FOQG_02657 [Fusarium oxysporum f. sp. raphani 54005] | 50.5 | 8.0E-73 |
| DIP_10000207 | putative autophagic vesicle breakdown protein atg22 [Ustilaginoidea virens] | 49.7 | 2.0E-163 |
| DIP_10000208 | polysaccharide lyase family 7 protein [Trichoderma atroviride IMI 206040] | 54.2 | 8.0E-81 |
| DIP_10000209 | hypothetical protein HIM_04083 [Hirsutella minnesotensis 3608] | 36 | 8.0E-53 |
| DIP_10000210 | NA | | |
| DIP_10000211 | NA | | |
| DIP_10000212 | hypothetical protein CGLO_11873 [Colletotrichum gloeosporioides Cg-14] | 78.8 | 4.0E-161 |
| DIP_10000213 | hypothetical protein PFICI_06717 [Pestalotiopsis fici W106-1] | 79.2 | 0.0 |
| DIP_10000214 | hypothetical protein MAN_05469, partial [Metarhizium anisopliae ARSEF 549] | 64.2 | 4.0E-143 |
| DIP_10000215 | hypothetical protein TRIVIDRAFT_67765 [Trichoderma virens Gv29-8] | 60.2 | 3.0E-179 |
| DIP_10000216 | protein RDR1 [Fusarium verticillioides 7600] | 44.4 | 7.0E-166 |
| DIP_10000217 | hypothetical protein PFICI_00205 [Pestalotiopsis fici W106-1] | 54.2 | 2.0E-98 |
| DIP_10000218 | putative cytochrome p450 4v3 protein [Eutypa lata UCREL1] | 52.5 | 2.0E-178 |
| DIP_10000219 | hypothetical protein PFICI_00128 [Pestalotiopsis fici W106-1] | 54.9 | 0.0 |
| DIP_10000220 | hypothetical protein PFICI_00194 [Pestalotiopsis fici W106-1] | 73.4 | 0.0 |
| DIP_10000221 | hypothetical protein PFICI_13223 [Pestalotiopsis fici W106-1] | 36.3 | 6.0E-80 |
| DIP_10000222 | hypothetical protein MYCGRDRAFT_67922 [Zymoseptoria tritici IPO323] | 43 | 8.0E-87 |
| DIP_10000223 | hypothetical protein OIDMADRAFT_121419 [Oidiodendron maius Zn] | 47.8 | 9.0E-122 |
| DIP_10000224 | hypothetical protein NECHADRAFT_44930 [Nectria haematococca mpVI 77-13-4] | 67.9 | 0.0 |
| DIP_10000225 | putative beta-glucosidase 2 protein [Eutypa lata UCREL1] | 67.3 | 0.0 |
| DIP_10000226 | putative vacuolar protein sorting-associated protein 35 protein [Eutypa lata UCREL1] | 78.6 | 0.0 |
| DIP_10000227 | hypothetical protein PFICI_00216 [Pestalotiopsis fici W106-1] | 54.3 | 0.0 |
| DIP_10000228 | NA | | |
| DIP_10000229 | hypothetical protein PFICI_07271 [Pestalotiopsis fici W106-1] | 50.7 | 5.0E-133 |
| DIP_10000230 | NA | | |
| DIP_10000231 | hypothetical protein PFICI_00663 [Pestalotiopsis fici W106-1] | 56.3 | 0.0 |
| DIP_10000232 | putative delta-1-pyrroline-5-carboxylate dehydrogenase [Diaporthe ampelina] | 72 | 0.0 |
| DIP_10000233 | putative transcription factor protein [Eutypa lata UCREL1] | 68.4 | 0.0 |
| DIP_10000234 | hypothetical protein PFICI_00239 [Pestalotiopsis fici W106-1] | 42.6 | 2.0E-154 |
| DIP_10000235 | hypothetical protein PFICI_05426 [Pestalotiopsis fici W106-1] | 42.7 | 8.0E-48 |
| DIP_10000236 | vacuolar carboxypeptidase [Colletotrichum orbiculare MAFF 240422] | 61.1 | 0.0 |
| DIP_10000237 | hypothetical protein PFICI_00247 [Pestalotiopsis fici W106-1] | 53.5 | 2.0E-65 |
| DIP_10000238 | NA | | |
| DIP_10000239 | hypothetical protein PFICI_15338 [Pestalotiopsis fici W106-1] | 77.1 | 3.0E-161 |
| DIP_10000240 | Phosphoglycerate kinase [Pestalotiopsis fici W106-1] | 89.2 | 0.0 |
| DIP_10000241 | hypothetical protein TD95_002745 [Thielaviopsis punctulata] | 88.6 | 4.0E-79 |
| DIP_10000242 | hypothetical protein UCREL1_3152 [Eutypa lata UCREL1] | 63.3 | 0.0 |
| DIP_10000243 | NA | | |
| DIP_10000244 | NA | | |
| DIP_10000245 | 3-isopropylmalate dehydratase [Pestalotiopsis fici W106-1] | 79.7 | 0.0 |
| DIP_10000246 | hypothetical protein FOIG_08060 [Fusarium oxysporum f. sp. cubense tropical race 4 54006] | 34 | 1.0E-100 |
| DIP_10000247 | NA | | |
| DIP_10000248 | NA | | |
| DIP_10000249 | NA | | |
| DIP_10000250 | NA | | |
| DIP_10000251 | NA | | |
| DIP_10000252 | NA | | |
| DIP_10000253 | NA | | |
| DIP_10000254 | NA | | |
| DIP_10000255 | hypothetical protein CGGC5_10992 [Colletotrichum gloeosporioides Nara gc5] | 36.2 | 2.0E-25 |
| DIP_10000256 | NA | | |
| DIP_10000257 | glycosyl transferase [Burkholderia sp. WSM2232] | 31.5 | 3.0E-15 |
| DIP_10000258 | NA | | |
| DIP_10000259 | NA | | |
| DIP_10000260 | pisatin demethylase [Gaeumannomyces graminis var. tritici R3-111a-1] | 54.6 | 0.0 |
| DIP_10000261 | hypothetical protein GLRG_11756 [Colletotrichum graminicola M1.001] | 32.7 | 3.0E-53 |
| DIP_10000262 | hypothetical protein NECHADRAFT_18425 [Nectria haematococca mpVI 77-13-4] | 35.4 | 9.0E-27 |
| DIP_10000263 | hypothetical protein VHEMI04368 [Torrubiella hemipterigena] | 32.8 | 2.0E-06 |
| DIP_10000264 | MFS transporter [Colletotrichum orbiculare MAFF 240422] | 67.9 | 0.0 |
| DIP_10000265 | hypothetical protein EURHEDRAFT_379064 [Aspergillus ruber CBS 135680] | 43.9 | 2.0E-155 |
| DIP_10000266 | NA | | |
| DIP_10000267 | hypothetical protein FOVG_13373 [Fusarium oxysporum f. sp. pisi HDV247] | 55.2 | 1.0E-164 |
| DIP_10000268 | NA | | |
| DIP_10000269 | hypothetical protein OIDMADRAFT_102054 [Oidiodendron maius Zn] | 76.3 | 0.0 |
| DIP_10000270 | Non-catalytic module family EXPN protein [Heterobasidion irregulare TC 32-1] | 37.2 | 4.0E-26 |
| DIP_10000271 | hypothetical protein S40293_07373 [Stachybotrys chartarum IBT 40293] | 52 | 0.0 |
| DIP_10000272 | pfs domain-containing protein, partial [Metarhizium majus ARSEF 297] | 44.4 | 7.0E-41 |
| DIP_10000273 | NA | | |
| DIP_10000274 | NA | | |
| DIP_10000275 | NA | | |
| DIP_10000276 | NA | | |
| DIP_10000277 | hypothetical protein VHEMI01022 [Torrubiella hemipterigena] | 47.8 | 1.0E-155 |
| DIP_10000278 | hypothetical protein UCDDA912_g02783 [Diaporthe ampelina] | 36.6 | 3.0E-17 |
| DIP_10000279 | integral membrane protein [Cordyceps militaris CM01] | 52.9 | 5.0E-100 |
| DIP_10000280 | NA | | |
| DIP_10000281 | hypothetical protein PFICI_11515 [Pestalotiopsis fici W106-1] | 37 | 1.0E-38 |
| DIP_10000282 | hypothetical protein FOZG_11962 [Fusarium oxysporum Fo47] | 33.6 | 6.0E-50 |
| DIP_10000283 | NA | | |
| DIP_10000284 | alpha-L-arabinofuranosidase (Arabinosidase) [Micromonospora sp. M42] | 72 | 3.0E-160 |
| DIP_10000285 | Endo-1,4-beta-xylanase C-like protein [Acremonium chrysogenum ATCC 11550] | 73.8 | 9.0E-165 |
| DIP_10000286 | hypothetical protein ASPNIDRAFT_56022 [Aspergillus niger ATCC 1015] | 34.3 | 2.0E-85 |
| DIP_10000287 | putative matrix metalloproteinase-11 protein [Togninia minima UCRPA7] | 78.7 | 9.0E-159 |
| DIP_10000288 | hypothetical protein UCRPA7_7161 [Togninia minima UCRPA7] | 69 | 0.0 |
| DIP_10000289 | NA | | |
| DIP_10000290 | NA | | |
| DIP_10000291 | NA | | |
| DIP_10000292 | hypothetical protein THITE_2109392 [Thielavia terrestris NRRL 8126] | 42.4 | 7.0E-53 |
| DIP_10000293 | hypothetical protein CGGC5_13047 [Colletotrichum gloeosporioides Nara gc5] | 60 | 3.0E-163 |
| DIP_10000294 | NA | | |
| DIP_10000295 | PKS5<br>Thiolase-like protein [Glarea lozoyensis ATCC 20868] | 50.7 | 0.0 |
| DIP_10000296 | NA | | |
| DIP_10000297 | hypothetical protein TRIVIDRAFT_160306, partial [Trichoderma virens Gv29-8] | 49.5 | 3.0E-23 |
| DIP_10000298 | hypothetical protein THAR02_11481 [Trichoderma harzianum] | 31.4 | 3.0E-22 |
| DIP_10000299 | hypothetical protein H633G_11182 [Metarhizium anisopliae BRIP 53284] | 49 | 2.0E-26 |
| DIP_10000300 | hypothetical protein FOYG_03509 [Fusarium oxysporum FOSC 3-a] | 58.1 | 1.0E-150 |
| DIP_10000301 | NA | | |
| DIP_10000302 | hypothetical protein M437DRAFT_54988, partial [Aureobasidium melanogenum CBS 110374] | 31.1 | 1.0E-24 |
| DIP_10000303 | NA | | |
| DIP_10000304 | hypothetical protein PFICI_13569 [Pestalotiopsis fici W106-1] | 46.9 | 8.0E-56 |
| DIP_10000305 | NA | | |
| DIP_10000306 | metalloprotease-like protein [Beauveria bassiana ARSEF 2860] | 61.2 | 3.0E-174 |
| DIP_10000307 | NA | | |
| DIP_10000308 | hypothetical protein M437DRAFT_54988, partial [Aureobasidium melanogenum CBS 110374] | 35.7 | 5.0E-38 |
| DIP_10000309 | NA | | |
| DIP_10000310 | hypothetical protein FOYG_03509 [Fusarium oxysporum FOSC 3-a] | 60.3 | 3.0E-166 |
| DIP_10000311 | NA | | |
| DIP_10000312 | hypothetical protein TCE0_060f19332 [Talaromyces cellulolyticus] | 58.7 | 6.0E-36 |
| DIP_10000313 | hypothetical protein CGGC5_13047 [Colletotrichum gloeosporioides Nara gc5] | 61 | 6.0E-167 |
| DIP_10000314 | squalene synthetase-like protein [Dothistroma septosporum NZE10] | 85.7 | 0.0 |
| DIP_10000315 | putative glycerophosphoryl diester phosphodiesterase family protein [Eutypa lata UCREL1] | 36.3 | 5.0E-82 |
| DIP_10000316 | NA | | |
| DIP_10000317 | hypothetical protein FOZG_17943 [Fusarium oxysporum Fo47] | 39.9 | 2.0E-22 |
| DIP_10000318 | hypothetical protein FOXB_06689 [Fusarium oxysporum Fo5176] | 44.6 | 5.0E-27 |
| DIP_10000319 | galactoside O-acetyltransferase [Trichoderma harzianum] | 74.1 | 4.0E-52 |
| DIP_10000320 | NA | | |
| DIP_10000321 | hypothetical protein PFICI_07330 [Pestalotiopsis fici W106-1] | 46.2 | 7.0E-75 |
| DIP_10000322 | NA | | |
| DIP_10000323 | hypothetical protein OOU_Y34scaffold00824g9 [Magnaporthe oryzae Y34] | 36.2 | 5.0E-16 |
| DIP_10000324 | NA | | |
| DIP_10000325 | hypothetical protein CGGC5_13047 [Colletotrichum gloeosporioides Nara gc5] | 59.1 | 2.0E-146 |
| DIP_10000326 | hypothetical protein FOC1_g10001881 [Fusarium oxysporum f. sp. cubense race 1] | 46.3 | 2.0E-135 |
| DIP_10000327 | predicted protein [Histoplasma capsulatum NAm1] | 33.8 | 8.0E-37 |
| DIP_10000328 | NA | | |
| DIP_10000329 | NA | | |
| DIP_10000330 | hypothetical protein M437DRAFT_54988, partial [Aureobasidium melanogenum CBS 110374] | 34.3 | 1.0E-36 |
| DIP_10000331 | NA | | |
| DIP_10000332 | NA | | |
| DIP_10000333 | hypothetical protein HIM_11311 [Hirsutella minnesotensis 3608] | 64 | 6.0E-148 |
| DIP_10000334 | hypothetical protein THITE_2123114 [Thielavia terrestris NRRL 8126] | 33.2 | 2.0E-37 |
| DIP_10000335 | hypothetical protein [Pasteurella multocida] | 33.8 | 3.0E-12 |
| DIP_10000336 | hypothetical protein CGGC5_5462 [Colletotrichum gloeosporioides Nara gc5] | 47.2 | 4.0E-121 |
| DIP_10000337 | hypothetical protein PFICI_06950 [Pestalotiopsis fici W106-1] | 51.8 | 0.0 |
| DIP_10000338 | NA | | |
| DIP_10000339 | hypothetical protein PFICI_14308 [Pestalotiopsis fici W106-1] | 56.9 | 0.0 |
| DIP_10000340 | hypothetical protein PFICI_14309 [Pestalotiopsis fici W106-1] | 39.2 | 5.0E-112 |
| DIP_10000341 | eukaryotic peptide chain release factor GTP-binding subunit [Gaeumannomyces graminis var. tritici R3-111a-1] | 72.9 | 0.0 |
| DIP_10000342 | putative afln vera monooxygenase protein [Eutypa lata UCREL1] | 52.3 | 0.0 |
| DIP_10000343 | NA | | |
| DIP_10000344 | predicted protein [Leptosphaeria maculans JN3] | 39.1 | 4.0E-101 |
| DIP_10000345 | hypothetical protein PFICI_09929 [Pestalotiopsis fici W106-1] | 56.5 | 7.0E-85 |
| DIP_10000346 | hypothetical protein FOIG_02217 [Fusarium oxysporum f. sp. cubense tropical race 4 54006] | 69.2 | 2.0E-33 |
| DIP_10000347 | NA | | |
| DIP_10000348 | hypothetical protein PFICI_10333 [Pestalotiopsis fici W106-1] | 85.2 | 0.0 |
| DIP_10000349 | hypothetical protein W97_07304 [Coniosporium apollinis CBS 100218] | 61.3 | 0.0 |
| DIP_10000350 | hypothetical protein PFICI_11244 [Pestalotiopsis fici W106-1] | 48.3 | 0.0 |
| DIP_10000351 | putative glycosyltransferase family 4 protein [Eutypa lata UCREL1] | 84.3 | 0.0 |
| DIP_10000352 | NA | | |
| DIP_10000353 | putative diphthamide biosynthesis protein 3 protein [Eutypa lata UCREL1] | 69.2 | 4.0E-39 |
| DIP_10000354 | ATP-binding cassette protein [Neurospora crassa OR74A] | 84.6 | 0.0 |
| DIP_10000355 | NA | | |
| DIP_10000356 | putative sugar transporter stl1 protein [Eutypa lata UCREL1] | 74.9 | 0.0 |
| DIP_10000357 | hypothetical protein GGTG_12056 [Gaeumannomyces graminis var. tritici R3-111a-1] | 30.4 | 2.0E-18 |
| DIP_10000358 | hypothetical protein ACRE_008280 [Acremonium chrysogenum ATCC 11550] | 33.3 | 2.0E-14 |
| DIP_10000359 | hypothetical protein O988_08390 [Pseudogymnoascus pannorum VKM F-3808] | 30.6 | 5.0E-83 |
| DIP_10000360 | hypothetical protein PFICI_11167 [Pestalotiopsis fici W106-1] | 43.6 | 5.0E-92 |
| DIP_10000361 | hypothetical protein UCREL1_104 [Eutypa lata UCREL1] | 48 | 4.0E-40 |
| DIP_10000362 | NA | | |
| DIP_10000363 | putative coproporphyrinogen iii oxidase protein [Eutypa lata UCREL1] | 73 | 0.0 |

| | | | |
|---|---|---|---|
| DIP_10000364 | putative bromodomain-containing factor 1 protein [Eutypa lata UCREL1] | 49.4 | 0.0 |
| DIP_10000365 | putative small nucleolar ribonucleoprotein complex component protein [Eutypa lata UCREL1] | 65.4 | 9.0E-146 |
| DIP_10000366 | putative aaa family atpase protein [Eutypa lata UCREL1] | 66.2 | 0.0 |
| DIP_10000367 | NA | | |
| DIP_10000368 | hypothetical protein PFICI_04943 [Pestalotiopsis fici W106-1] | 34.6 | 1.0E-32 |
| DIP_10000369 | NA | | |
| DIP_10000370 | hypothetical protein GGTG_05621 [Gaeumannomyces graminis var. tritici R3-111a-1] | 30.5 | 1.0E-17 |
| DIP_10000371 | hypothetical protein TRIVIDRAFT_149245 [Trichoderma virens Gv29-8] | 41.9 | 3.0E-107 |
| DIP_10000372 | hypothetical protein FGSG_11248 [Fusarium graminearum PH-1] | 57.4 | 5.0E-123 |
| DIP_10000373 | NA | | |
| DIP_10000374 | NA | | |
| DIP_10000375 | NA | | |
| DIP_10000376 | hypothetical protein THITE_42560 [Thielavia terrestris NRRL 8126] | 32.9 | 2.0E-21 |
| DIP_10000377 | putative heat shock protein sti1 protein [Eutypa lata UCREL1] | 57.5 | 0.0 |
| DIP_10000378 | hypothetical protein PFICI_13498 [Pestalotiopsis fici W106-1] | 37.1 | 3.0E-146 |
| DIP_10000379 | high affinity copper transporter [Verticillium dahliae VdLs.17] | 48.5 | 5.0E-63 |
| DIP_10000380 | hypothetical protein CGLO_02455 [Colletotrichum gloeosporioides Cg-14] | 33.2 | 2.0E-87 |
| DIP_10000381 | hypothetical protein PFICI_04080 [Pestalotiopsis fici W106-1] | 48.6 | 2.0E-157 |
| DIP_10000382 | Formaldehyde transketolase [Rasamsonia emersonii CBS 393.64] | 69.8 | 0.0 |
| DIP_10000383 | predicted protein [Trichoderma reesei QM6a] | 70.7 | 0.0 |
| DIP_10000384 | NA | | |
| DIP_10000385 | hypothetical protein THITE_2071706 [Thielavia terrestris NRRL 8126] | 49.7 | 7.0E-138 |
| DIP_10000386 | hypothetical protein V493_05328 [Pseudogymnoascus pannorum VKM F-4281 (FW-2241)] | 53.4 | 0.0 |
| DIP_10000387 | putative 3-hydroxy-3-methylglutaryl-coenzyme a reductase protein [Eutypa lata UCREL1] HMG-CoA reductase catalyzes the conversion of HMG-CoA to mevalonate | 74.9 | 0.0 |
| DIP_10000388 | hypothetical protein PFICI_[illegible] [Pestalotiopsis fici W106-1] | 50.4 | 2.0E-124 |
| DIP_10000389 | putative heterokaryon incompatibility protein [Eutypa lata UCREL1] | 34.9 | 1.0E-120 |
| DIP_10000390 | hypothetical protein A1O1_08839 [Capronia coronata CBS 617.96] | 52.6 | 6.0E-46 |
| DIP_10000391 | hypothetical protein FPSE_11641 [Fusarium pseudograminearum CS3096] | 46 | 7.0E-67 |
| DIP_10000392 | Histidine biosynthesis trifunctional protein [Pestalotiopsis fici W106-1] | 51.6 | 0.0 |
| DIP_10000393 | hypothetical protein NECHADRAFT_89622 [Nectria haematococca mpVI 77-13-4] | 57.6 | 0.0 |
| DIP_10000394 | NA | | |
| DIP_10000395 | putative cupin domain containing protein [Togninia minima UCRPA7] | 42.5 | 4.0E-46 |
| DIP_10000396 | putative membrane transporter protein [Eutypa lata UCREL1] | 83 | 0.0 |
| DIP_10000397 | hypothetical protein GALMADRAFT_32921, partial [Galerina marginata CBS 339.88] | 39.8 | 8.0E-49 |
| DIP_10000398 | putative fad binding domain-containing protein [Eutypa lata UCREL1] | 46.7 | 0.0 |
| DIP_10000399 | hypothetical protein V491_06313 [Pseudogymnoascus pannorum VKM F-3775] | 97.5 | 3.0E-48 |
| DIP_10000400 | hypothetical protein UCREL1_2027 [Eutypa lata UCREL1] | 45.3 | 1.0E-104 |
| DIP_10000401 | putative arsenate reductase protein [Eutypa lata UCREL1] | 47.3 | 3.0E-104 |
| DIP_10000402 | hypothetical protein CGLO_11902 [Colletotrichum gloeosporioides Cg-14] | 33.6 | 2.0E-58 |
| DIP_10000403 | putative nuclear elongation and deformation protein 1 protein [Eutypa lata UCREL1] | 62.7 | 0.0 |
| DIP_10000404 | NA | | |
| DIP_10000405 | hypothetical protein GGTG_07793 [Gaeumannomyces graminis var. tritici R3-111a-1] | 68 | 4.0E-65 |
| DIP_10000406 | hypothetical protein PFICI_06938 [Pestalotiopsis fici W106-1] | 60.4 | 0.0 |
| DIP_10000407 | putative nicotinate phosphoribosyltransferase protein [Eutypa lata UCREL1] | 75.9 | 0.0 |
| DIP_10000408 | hypothetical protein FOYG_11739 [Fusarium oxysporum FOSC 3-a] | 61.5 | 1.0E-171 |
| DIP_10000409 | putative alpha beta hydrolase protein [Togninia minima UCRPA7] | 44.6 | 1.0E-85 |
| DIP_10000410 | hypothetical protein TRIATDRAFT_48897 [Trichoderma atroviride IMI 206040] | 54.6 | 4.0E-128 |
| DIP_10000411 | hypothetical protein NECHADRAFT_75779 [Nectria haematococca mpVI 77-13-4] | 35.4 | 8.0E-109 |
| DIP_10000412 | hypothetical protein UCREL1_2325 [Eutypa lata UCREL1] | 35.9 | 1.0E-12 |
| DIP_10000413 | aryl-alcohol dehydrogenase Aad14, putative [Aspergillus flavus NRRL3357] | 69.5 | 0.0 |
| DIP_10000414 | conserved hypothetical protein [Paracoccidioides sp. 'lutzii' Pb01] | 39.5 | 2.0E-34 |
| DIP_10000415 | putative alpha beta hydrolase fold protein [Eutypa lata UCREL1] | 59.3 | 8.0E-117 |
| DIP_10000416 | hypothetical protein ANO11243_050190 [fungal sp. No.11243] | 48.7 | 9.0E-106 |
| DIP_10000417 | NA | | |
| DIP_10000418 | hypothetical protein PVAR5_3618 [Byssochlamys spectabilis No. 5] | 49.5 | 4.0E-27 |
| DIP_10000419 | NA | | |
| DIP_10000420 | putative short-chain dehydrogenase reductase family protein [Eutypa lata UCREL1] | 63.8 | 1.0E-127 |
| DIP_10000421 | NA | | |
| DIP_10000422 | acyl carrier protein, putative [Metarhizium acridum CQMa 102] | 73.3 | 1.0E-62 |
| DIP_10000423 | putative nadh-ubiquinone oxidoreductase 23 kda subunit protein [Eutypa lata UCREL1] | 78.8 | 6.0E-133 |
| DIP_10000424 | NA | | |
| DIP_10000425 | C6 zinc finger domain protein [Rasamsonia emersonii CBS 393.64] | 42 | 0.0 |
| DIP_10000426 | putative dna-binding protein [Eutypa lata UCREL1] | 34.3 | 7.0E-78 |
| DIP_10000427 | NA | | |
| DIP_10000428 | putative c6 transcription factor [Diaporthe ampelina] | 48.5 | 0.0 |
| DIP_10000429 | putative amidohydrolase family protein [Eutypa lata UCREL1] | 54 | 0.0 |
| DIP_10000430 | NA | | |
| DIP_10000431 | NA | | |
| DIP_10000432 | hypothetical protein PFICI_03412 [Pestalotiopsis fici W106-1] | 71.9 | 0.0 |
| DIP_10000433 | short-chain dehydrogenase [Colletotrichum higginsianum] | 58.8 | 2.0E-111 |
| DIP_10000434 | putative vacuolar import and degradation protein 27 protein [Eutypa lata UCREL1] | 83.6 | 0.0 |
| DIP_10000435 | putative -trehalose-phosphate synthase protein [Eutypa lata UCREL1] | 82.6 | 0.0 |
| DIP_10000436 | putative uracil phosphoribosyltransferase [Diaporthe ampelina] | 48.4 | 6.0E-52 |
| DIP_10000437 | predicted protein [Trichoderma reesei QM6a] | 44 | 4.0E-68 |
| DIP_10000438 | hypothetical protein THAR02_08480 [Trichoderma harzianum] | 54.3 | 0.0 |
| DIP_10000439 | hypothetical protein PFICI_11659 [Pestalotiopsis fici W106-1] | 41.8 | 4.0E-98 |
| DIP_10000440 | NA | | |
| DIP_10000441 | putative beta- -galactanase protein [Togninia minima UCRPA7] | 51.5 | 6.0E-153 |
| DIP_10000442 | hypothetical protein PFICI_13156 [Pestalotiopsis fici W106-1] | 31.1 | 4.0E-57 |
| DIP_10000443 | hypothetical protein FPSE_01176 [Fusarium pseudograminearum CS3096] | 85 | 0.0 |
| DIP_10000444 | hypothetical protein GLRG_01990, partial [Colletotrichum graminicola M1.001] | 84 | 1.0E-24 |
| DIP_10000445 | putative transcriptional regulator protein [Eutypa lata UCREL1] | 51.3 | 8.0E-64 |
| DIP_10000446 | putative bret domain protein [Eutypa lata UCREL1] | 34.3 | 1.0E-90 |
| DIP_10000447 | hypothetical protein PFICI_05361 [Pestalotiopsis fici W106-1] | 59 | 0.0 |
| DIP_10000448 | hypothetical protein PFICI_05360 [Pestalotiopsis fici W106-1] | 69.8 | 0.0 |
| DIP_10000449 | hypothetical protein TD95_000259 [Thielaviopsis punctulata] | 44.4 | 0.0 |
| DIP_10000450 | hypothetical protein NECHADRAFT_82569 [Nectria haematococca mpVI 77-13-4] | 43.8 | 8.0E-47 |
| DIP_10000451 | hypothetical protein PFICI_10079 [Pestalotiopsis fici W106-1] | 43.2 | 2.0E-145 |
| DIP_10000452 | NA | | |
| DIP_10000453 | NA | | |
| DIP_10000454 | putative immunoreactive protein [Eutypa lata UCREL1] | 43.3 | 2.0E-40 |
| DIP_10000455 | NA | | |
| DIP_10000456 | putative sulfate permease 2 protein [Eutypa lata UCREL1] | 64.1 | 0.0 |
| DIP_10000457 | hypothetical protein PFICI_12570 [Pestalotiopsis fici W106-1] | 50.2 | 0.0 |
| DIP_10000458 | hypothetical protein UCREL1_5854 [Eutypa lata UCREL1] | 40.6 | 6.0E-26 |
| DIP_10000459 | hypothetical protein FGSG_11438 [Fusarium graminearum PH-1] | 37.8 | 3.0E-80 |
| DIP_10000460 | putative dna replication regulator sld2 protein [Togninia minima UCRPA7] | 45.9 | 3.0E-120 |
| DIP_10000461 | hypothetical protein CH063_08254 [Colletotrichum higginsianum] | 51.1 | 1.0E-59 |
| DIP_10000462 | hypothetical protein PFICI_12542 [Pestalotiopsis fici W106-1] | 67.3 | 0.0 |
| DIP_10000463 | hypothetical protein PFICI_13240 [Pestalotiopsis fici W106-1] | 66.4 | 1.0E-156 |
| DIP_10000464 | hypothetical protein CPC735_066940 [Coccidioides posadasii C735 delta SOWgp] | 63.5 | 5.0E-117 |
| DIP_10000465 | NA | | |
| DIP_10000466 | hypothetical protein PFICI_02553 [Pestalotiopsis fici W106-1] | 56.2 | 0.0 |
| DIP_10000467 | putative general amino acid permease agp2 protein [Eutypa lata UCREL1] | 74.5 | 0.0 |
| DIP_10000468 | NA | | |
| DIP_10000469 | putative dna mismatch repair protein msh-2 protein [Eutypa lata UCREL1] | 82.3 | 0.0 |
| DIP_10000470 | hypothetical protein PFICI_12230 [Pestalotiopsis fici W106-1] | 54 | 1.0E-126 |
| DIP_10000471 | putative eif4-gamma eif5 eif2-epsilon protein [Eutypa lata UCREL1] | 64.4 | 0.0 |
| DIP_10000472 | hypothetical protein PFICI_12578 [Pestalotiopsis fici W106-1] | 61.8 | 0.0 |
| DIP_10000473 | NA | | |
| DIP_10000474 | hypothetical protein FOPG_18856 [Fusarium oxysporum f. sp. conglutinans race 2 54008] | 34.7 | 3.0E-11 |
| DIP_10000475 | NA | | |
| DIP_10000476 | putative glyoxylate reductase protein [Eutypa lata UCREL1] | 79.6 | 0.0 |
| DIP_10000477 | ATPase [Fusarium oxysporum f. sp. lycopersici MN25] | 74.2 | 0.0 |
| DIP_10000478 | putative plasma-membrane proton-efflux P-type ATPase [Colletotrichum sublineola] | 42.7 | 6.0E-20 |
| DIP_10000479 | putative plasma membrane atpase protein [Eutypa lata UCREL1] | 84.1 | 0.0 |
| DIP_10000480 | hypothetical protein PFICI_10580 [Pestalotiopsis fici W106-1] | 37.2 | 2.0E-110 |
| DIP_10000481 | NA | | |
| DIP_10000482 | putative phd-finger domain-containing protein [Eutypa lata UCREL1] | 57.1 | 1.0E-170 |
| DIP_10000483 | putative mitochondrial gtpase 1 protein [Eutypa lata UCREL1] | 62.8 | 9.0E-163 |
| DIP_10000484 | hypothetical protein ACRE_061050 [Acremonium chrysogenum ATCC 11550] | 61.1 | 4.0E-133 |
| DIP_10000485 | hypothetical protein Z518_05318 [Rhinocladiella mackenziei CBS 650.93] | 32.9 | 2.0E-52 |
| DIP_10000486 | putative d-arabinitol dehydrogenase protein [Eutypa lata UCREL1] | 66.3 | 1.0E-163 |
| DIP_10000487 | Peroxidase family 2 [Aspergillus parasiticus SU-1] | 61.2 | 9.0E-111 |
| DIP_10000488 | hypothetical protein MAC_09758 [Metarhizium acridum CQMa 102] | 51 | 3.0E-88 |
| DIP_10000489 | hypothetical protein V492_02219 [Pseudogymnoascus pannorum VKM F-4246] | 31 | 8.0E-14 |
| DIP_10000490 | hypothetical protein CGLO_01082 [Colletotrichum gloeosporioides Cg-14] | 67.6 | 0.0 |
| DIP_10000491 | putative 37s ribosomal protein rsm22 protein [Eutypa lata UCREL1] | 49.5 | 0.0 |
| DIP_10000492 | putative ydg sra domain-containing protein [Eutypa lata UCREL1] | 42.9 | 1.0E-80 |
| DIP_10000493 | NA | | |
| DIP_10000494 | putative mannose-6-phosphate isomerase protein [Eutypa lata UCREL1] | 88.2 | 9.0E-101 |
| DIP_10000495 | hypothetical protein PFICI_13817 [Pestalotiopsis fici W106-1] | 57.3 | 2.0E-147 |
| DIP_10000496 | hypothetical protein PFICI_13818 [Pestalotiopsis fici W106-1] | 51.9 | 3.0E-65 |
| DIP_10000497 | hypothetical protein PFICI_02255 [Pestalotiopsis fici W106-1] | 76.2 | 0.0 |
| DIP_10000498 | hypothetical protein VDAG_05358 [Verticillium dahliae VdLs.17] | 48.2 | 3.0E-122 |
| DIP_10000499 | putative fad binding domain-containing protein [Eutypa lata UCREL1] | 66.7 | 0.0 |
| DIP_10000500 | hypothetical protein HIM_05809 [Hirsutella minnesotensis 3608] | 30.7 | 1.0E-16 |
| DIP_10000501 | hypothetical protein CGLO_07212 [Colletotrichum gloeosporioides Cg-14] | 66.6 | 0.0 |
| DIP_10000502 | hypothetical protein PFICI_13953 [Pestalotiopsis fici W106-1] | 47.8 | 2.0E-68 |
| DIP_10000503 | NAD-dependent protein deacetylase hst2-like protein [Acremonium chrysogenum ATCC 11550] | 67 | 7.0E-148 |
| DIP_10000504 | putative auxin efflux carrier superfamily protein [Eutypa lata UCREL1] | 67.3 | 0.0 |
| DIP_10000505 | hypothetical protein PFICI_13952 [Pestalotiopsis fici W106-1] | 67 | 0.0 |
| DIP_10000506 | NA | | |
| DIP_10000507 | NA | | |
| DIP_10000508 | calmodulin A [Blastomyces dermatitidis SLH14081] | 50 | 4.0E-48 |
| DIP_10000509 | NA | | |
| DIP_10000510 | NA | | |
| DIP_10000511 | NA | | |
| DIP_10000512 | hypothetical protein VHEMI01823 [Torrubiella hemipterigena] | 46.5 | 3.0E-79 |
| DIP_10000513 | hypothetical protein CFIO01_11083 [Colletotrichum fioriniae PJ7] | 31.7 | 7.0E-12 |
| DIP_10000514 | hypothetical protein OOW_P131scaffold01655g1 [Magnaporthe oryzae P131] | 64.2 | 1.0E-44 |
| DIP_10000515 | hypothetical protein UCREL1_7482 [Eutypa lata UCREL1] | 72.3 | 0.0 |
| DIP_10000516 | Putative solute carrier [Podospora anserina S mat+] | 58.8 | 2.0E-170 |
| DIP_10000517 | NA | | |
| DIP_10000518 | NA | | |
| DIP_10000519 | hypothetical protein PFICI_13962 [Pestalotiopsis fici W106-1] | 61.3 | 0.0 |
| DIP_10000520 | hypothetical protein CH063_06875, partial [Colletotrichum higginsianum] | 56.5 | 0.0 |
| DIP_10000521 | putative molybdopterin biosynthesis protein moeb protein [Eutypa lata UCREL1] | 73.3 | 0.0 |
| DIP_10000522 | Similar to 5' acc. no. Q61503 [Pyronema omphalodes CBS 100304] | 30.1 | 6.0E-52 |
| DIP_10000523 | hypothetical protein THITE_2112754 [Thielavia terrestris NRRL 8126] | 44.3 | 3.0E-113 |
| DIP_10000524 | hypothetical protein PFICI_14039 [Pestalotiopsis fici W106-1] | 45.9 | 3.0E-62 |
| DIP_10000525 | NA | | |
| DIP_10000526 | hypothetical protein FGSG_09439 [Fusarium graminearum PH-1] | 77.1 | 5.0E-135 |
| DIP_10000527 | putative mitochondrial 37s ribosomal protein mrps16 protein [Eutypa lata UCREL1] | 81.1 | 2.0E-61 |
| DIP_10000528 | hypothetical protein PFICI_13828 [Pestalotiopsis fici W106-1] | 76.9 | 0.0 |
| DIP_10000529 | putative pyridoxamine 5 -phosphate oxidase protein [Eutypa lata UCREL1] | 75.7 | 8.0E-119 |
| DIP_10000530 | putative monocarboxylate permease protein [Eutypa lata UCREL1] | 60.1 | 0.0 |
| DIP_10000531 | NA | | |
| DIP_10000532 | NA | | |
| DIP_10000533 | putative malate synthase [Diaporthe ampelina] | 90.8 | 0.0 |
| DIP_10000534 | hypothetical protein THITE_2111697 [Thielavia terrestris NRRL 8126] | 82 | 0.0 |
| DIP_10000535 | predicted protein [Trichoderma reesei QM6a] | 43.6 | 0.0 |
| DIP_10000536 | putative structural maintenance of chromosome complex subunit protein [Eutypa lata UCREL1] | 49 | 0.0 |
| DIP_10000537 | conserved hypothetical protein [Talaromyces stipitatus ATCC 10500] | 43.8 | 7.0E-24 |
| DIP_10000538 | ankyrin [Sulfolobus islandicus] | 46 | 2.0E-41 |
| DIP_10000539 | hypothetical protein THAR02_11041 [Trichoderma harzianum] | 44.3 | 3.0E-152 |
| DIP_10000540 | NA | | |
| DIP_10000541 | hypothetical protein PFICI_10229 [Pestalotiopsis fici W106-1] | 57.3 | 1.0E-83 |
| DIP_10000542 | NA | | |
| DIP_10000543 | hypothetical protein THITE_2057225 [Thielavia terrestris NRRL 8126] | 49.2 | 5.0E-90 |
| DIP_10000544 | hypothetical protein A1O5_12212 [Cladophialophora psammophila CBS 110553] | 57 | 0.0 |
| DIP_10000545 | hypothetical protein PFICI_02833 [Pestalotiopsis fici W106-1] | 46 | 3.0E-109 |
| DIP_10000546 | NA | | |
| DIP_10000547 | hypothetical protein PFICI_09871 [Pestalotiopsis fici W106-1] | 54 | 6.0E-117 |
| DIP_10000548 | NA | | |
| DIP_10000549 | hypothetical protein FAVG1_03128 [Fusarium avenaceum] | 41.6 | 3.0E-86 |
| DIP_10000550 | Hsp70-like protein [Scedosporium apiospermum] | 90.3 | 0.0 |
| DIP_10000551 | NA | | |
| DIP_10000552 | putative ribosome biogenesis protein sqt1 protein [Eutypa lata UCREL1] | 68.1 | 0.0 |
| DIP_10000553 | hypothetical protein CH063_00325 [Colletotrichum higginsianum] | 42.5 | 3.0E-47 |
| DIP_10000554 | NA | | |
| DIP_10000555 | NA | | |
| DIP_10000556 | hypothetical protein UCREL1_648 [Eutypa lata UCREL1] | 65.7 | 2.0E-72 |
| DIP_10000557 | mycocerosic acid synthase [Colletotrichum fioriniae PJ7] | 47.1 | 4.0E-15 |
| DIP_10000558 | hypothetical protein CGLO_14150 [Colletotrichum gloeosporioides Cg-14] | 38.5 | 0.0 |
| DIP_10000559 | NA | | |
| DIP_10000560 | NA | | |
| DIP_10000561 | NA | | |
| DIP_10000562 | putative amp-binding enzyme protein [Eutypa lata UCREL1] | 74.4 | 0.0 |
| DIP_10000563 | hypothetical protein PFICI_11408 [Pestalotiopsis fici W106-1] | 44.9 | 0.0 |
| DIP_10000564 | hypothetical protein PFICI_11409 [Pestalotiopsis fici W106-1] | 57.2 | 0.0 |
| DIP_10000565 | putative citrate synthase protein [Eutypa lata UCREL1] | 91.2 | 0.0 |
| DIP_10000566 | Mitochondrial DNA replication protein [Pestalotiopsis fici W106-1] | 87 | 0.0 |
| DIP_10000567 | putative plasma membrane atpase protein [Eutypa lata UCREL1] | 89.2 | 0.0 |
| DIP_10000568 | hypothetical protein CH063_02578 [Colletotrichum higginsianum] | 30.6 | 5.0E-52 |
| DIP_10000569 | putative bzip transcription protein [Eutypa lata UCREL1] | 42.1 | 6.0E-123 |
| DIP_10000570 | putative nucleoporin nic96 protein [Eutypa lata UCREL1] | 74.1 | 0.0 |
| DIP_10000571 | sugar kinase [Pestalotiopsis fici W106-1] | 81.3 | 0.0 |
| DIP_10000572 | hypothetical protein [Podospora anserina S mat+] | 49.5 | 0.0 |
| DIP_10000573 | putative arp2 3 complex subunit protein [Eutypa lata UCREL1] | 80.5 | 1.0E-115 |
| DIP_10000574 | TATA-binding protein [Trichoplax adhaerens] | 59 | 1.0E-77 |
| DIP_10000575 | NA | | |
| DIP_10000576 | NA | | |
| DIP_10000577 | serine/threonine protein kinase [Mycobacterium sp. UM_Kg1] | 34.7 | 6.0E-12 |
| DIP_10000578 | putative nulp1-pending protein [Eutypa lata UCREL1] | 53.5 | 0.0 |
| DIP_10000579 | hypothetical protein V493_07758 [Pseudogymnoascus pannorum VKM F-4281 (FW-2241)] | 81 | 8.0E-44 |
| DIP_10000580 | putative alpha-l-arabinofuranosidase protein [Eutypa lata UCREL1] | 63.3 | 0.0 |
| DIP_10000581 | NA | | |
| DIP_10000582 | related to TMA46-Protein putative involved in cytoplasmic ribosome function [Fusarium fujikuroi IMI 58289] | 78 | 0.0 |
| DIP_10000583 | hypothetical protein MYCTH_2316572 [Myceliophthora thermophila ATCC 42464] | 54.2 | 4.0E-16 |
| DIP_10000584 | hypothetical protein PFICI_09430 [Pestalotiopsis fici W106-1] | 57.6 | 3.0E-81 |
| DIP_10000585 | hypothetical protein PFICI_09432 [Pestalotiopsis fici W106-1] | 56.3 | 0.0 |
| DIP_10000586 | putative acetyl-coenzyme a synthetase protein [Eutypa lata UCREL1] | 77.2 | 0.0 |
| DIP_10000587 | conserved hypothetical protein [Histoplasma capsulatum G186AR] | 78 | 0.0 |
| DIP_10000588 | hypothetical protein CH063_01310 [Colletotrichum higginsianum] | 64.3 | 2.0E-52 |
| DIP_10000589 | extracellular solute-binding protein family 5 [Colletotrichum orbiculare MAFF 240422] | 46.4 | 6.0E-123 |
| DIP_10000590 | pantothenate transporter liz1 [Colletotrichum higginsianum] | 65.7 | 0.0 |
| DIP_10000591 | hypothetical protein PFICI_11214 [Pestalotiopsis fici W106-1] | 90.9 | 0.0 |
| DIP_10000592 | hypothetical protein PFICI_09431 [Pestalotiopsis fici W106-1] | 54.7 | 4.0E-22 |
| DIP_10000593 | Serine/threonine-protein kinase nrc-2 [Neurospora tetrasperma FGSC 2508] | 84.1 | 0.0 |
| DIP_10000594 | hypothetical protein TRIVIDRAFT_129239, partial [Trichoderma virens Gv29-8] | 44.4 | 9.0E-44 |
| DIP_10000595 | probable short chain dehydrogenase [Fusarium fujikuroi IMI 58289] | 62.7 | 4.0E-106 |
| DIP_10000596 | putative proton myo-inositol cotransporter protein [Eutypa lata UCREL1] | 54.9 | 0.0 |
| DIP_10000597 | hypothetical protein HMPREF1624_08401 [Sporothrix schenckii ATCC 58251] | 43.1 | 2.0E-24 |
| DIP_10000598 | hypothetical protein ACRE_043800 [Acremonium chrysogenum ATCC 11550] | 35.7 | 3.0E-89 |
| DIP_10000599 | NA | | |
| DIP_10000600 | NA | | |
| DIP_10000601 | NA | | |
| DIP_10000602 | NA | | |
| DIP_10000603 | NA | | |
| DIP_10000604 | NA | | |
| DIP_10000605 | NA | | |
| DIP_10000606 | NA | | |
| DIP_10000607 | NA | | |
| DIP_10000608 | NA | | |
| DIP_10000609 | NA | | |
| DIP_10000610 | NA | | |
| DIP_10000611 | NA | | |
| DIP_10000612 | 26S proteasome regulatory subunit rpn6 [Pestalotiopsis fici W106-1] | 83.2 | 0.0 |
| DIP_10000613 | putative u3 small nucleolar rna associated protein [Eutypa lata UCREL1] | 83 | 0.0 |
| DIP_10000614 | hypothetical protein PFICI_08961 [Pestalotiopsis fici W106-1] | 55.5 | 6.0E-121 |
| DIP_10000615 | putative protein kinase domain-containing protein [Eutypa lata UCREL1] | 43.3 | 5.0E-128 |
| DIP_10000616 | putative glycolipid-anchored surface protein 5 protein [Eutypa lata UCREL1] | 67.3 | 0.0 |
| DIP_10000617 | NA | | |
| DIP_10000618 | hypothetical protein TRIVIDRAFT_209784 [Trichoderma virens Gv29-8] | 52.1 | 0.0 |
| DIP_10000619 | hypothetical protein PFICI_15258 [Pestalotiopsis fici W106-1] | 54.5 | 0.0 |
| DIP_10000620 | putative ubiquitin fusion degradation protein [Eutypa lata UCREL1] | 41.5 | 1.0E-113 |
| DIP_10000621 | putative hsp70 protein [Togninia minima UCRPA7] | 86.7 | 0.0 |
| DIP_10000622 | putative serine threonine-protein kinase mak protein [Togninia minima UCRPA7] | 71.4 | 0.0 |
| DIP_10000623 | hypothetical protein UCREL1_11267 [Eutypa lata UCREL1] | 48.4 | 3.0E-37 |
| DIP_10000624 | putative hydroxyproline-rich glycoprotein dz-hrgp protein [Eutypa lata UCREL1] | 34.9 | 2.0E-44 |
| DIP_10000625 | NA | | |
| DIP_10000626 | Pyoverdine biosynthesis [Metarhizium album ARSEF 1941] | 35.6 | 2.0E-57 |
| DIP_10000627 | putative ribosomal protein yme-31 protein [Eutypa lata UCREL1] | 68.2 | 3.0E-55 |
| DIP_10000628 | hypothetical protein MGG_09177 [Magnaporthe oryzae 70-15] | 43.2 | 2.0E-72 |
| DIP_10000629 | flavin-binding monooxygenase-like protein [Ophiocordyceps sinensis CO18] | 52.9 | 0.0 |
| DIP_10000630 | hypothetical protein SNOG_02027 [Phaeosphaeria nodorum SN15] | 37.1 | 3.0E-23 |
| DIP_10000631 | NA | | |
| DIP_10000632 | hypothetical protein PFICI_08850 [Pestalotiopsis fici W106-1] | 65.2 | 0.0 |
| DIP_10000633 | NA | | |
| DIP_10000634 | putative exoglucanase 3 precursor protein [Togninia minima UCRPA7] | 51.2 | 1.0E-119 |
| DIP_10000635 | hypothetical protein PFICI_07076 [Pestalotiopsis fici W106-1] | 65.8 | 0.0 |
| DIP_10000636 | NA | | |
| DIP_10000637 | RNA-dependent RNA polymerase [Rosellinia necatrix] | 50.3 | 0.0 |
| DIP_10000638 | NA | | |
| DIP_10000639 | hypothetical protein PFICI_09061 [Pestalotiopsis fici W106-1] | 62.3 | 7.0E-97 |
| DIP_10000640 | 40S ribosomal protein S23 [Pestalotiopsis fici W106-1] | 99.3 | 2.0E-98 |
| DIP_10000641 | hypothetical protein BC1G_12033 [Botrytis cinerea B05.10] | 73.5 | 8.0E-168 |
| DIP_10000642 | hypothetical protein PFICI_09058 [Pestalotiopsis fici W106-1] | 70.1 | 0.0 |
| DIP_10000643 | NA | | |
| DIP_10000644 | putative golgi matrix protein [Eutypa lata UCREL1] | 72.5 | 0.0 |
| DIP_10000645 | putative ribosomal biogenesis protein gar2 protein [Eutypa lata UCREL1] | 52.4 | 3.0E-94 |
| DIP_10000646 | hypothetical protein FVEG_11272 [Fusarium verticillioides 7600] | 43.8 | 4.0E-37 |
| DIP_10000647 | NA | | |
| DIP_10000648 | putative serine-threonine rich protein [Eutypa lata UCREL1] | 54.2 | 3.0E-82 |
| DIP_10000649 | hypothetical protein TERG_07684 [Trichophyton rubrum CBS 118892] | 65.6 | 1.0E-17 |
| DIP_10000650 | hypothetical protein UCREL1_6499 [Eutypa lata UCREL1] | 46.8 | 4.0E-80 |
| DIP_10000651 | hypothetical protein PFICI_15021 [Pestalotiopsis fici W106-1] | 49.2 | 0.0 |
| DIP_10000652 | hypothetical protein PFICI_15020 [Pestalotiopsis fici W106-1] | 44.4 | 1.0E-91 |
| DIP_10000653 | hypothetical protein THITE_2124422 [Thielavia terrestris NRRL 8126] | 65.5 | 0.0 |
| DIP_10000654 | hypothetical protein [Podospora anserina S mat+] | 50.3 | 3.0E-95 |
| DIP_10000655 | MYB DNA-binding domain-containing protein [Colletotrichum higginsianum] | 57.2 | 1.0E-103 |
| DIP_10000656 | putative cap-gly domain-containing protein [Eutypa lata UCREL1] | 73.3 | 3.0E-132 |
| DIP_10000657 | hypothetical protein FVEG_00763 [Fusarium verticillioides 7600] | 38.5 | 2.0E-100 |
| DIP_10000658 | putative vacuolar amino acid transporter 2 protein [Togninia minima UCRPA7] | 64.9 | 0.0 |
| DIP_10000659 | putative aspartic-type endopeptidase protein [Eutypa lata UCREL1] | 50.7 | 9.0E-151 |
| DIP_10000660 | hypothetical protein PFICI_15029 [Pestalotiopsis fici W106-1] | 49 | 7.0E-107 |
| DIP_10000661 | Protein get1 [Pestalotiopsis fici W106-1] | 65.7 | 3.0E-98 |
| DIP_10000662 | hypothetical protein PFICI_15027 [Pestalotiopsis fici W106-1] | 43.4 | 3.0E-87 |
| DIP_10000663 | hypothetical protein BAUCODRAFT_33279 [Baudoinia compniacensis UAMH 10762] | 92.8 | 2.0E-132 |
| DIP_10000664 | putative kow motif containing protein [Eutypa lata UCREL1] | 63.6 | 7.0E-152 |
| DIP_10000665 | NA | | |
| DIP_10000666 | NA | | |
| DIP_10000667 | hypothetical protein NECHADRAFT_39197 [Nectria haematococca mpVI 77-13-4] | 34.3 | 6.0E-127 |
| DIP_10000668 | NA | | |
| DIP_10000669 | hypothetical protein OCS_06701 [Ophiocordyceps sinensis CO18] | 37.8 | 2.0E-71 |
| DIP_10000670 | hypothetical protein PFICI_11324 [Pestalotiopsis fici W106-1] | 44.5 | 4.0E-70 |
| DIP_10000671 | putative histidine phosphotransferase hpt1p protein [Eutypa lata UCREL1] | 55.3 | 2.0E-48 |
| DIP_10000672 | hypothetical protein PFICI_11326 [Pestalotiopsis fici W106-1] | 57.8 | 1.0E-129 |
| DIP_10000673 | hypothetical protein PFICI_11327 [Pestalotiopsis fici W106-1] | 61.1 | 0.0 |
| DIP_10000674 | hypothetical protein PFICI_11327 [Pestalotiopsis fici W106-1] | 65.6 | 0.0 |
| DIP_10000675 | hypothetical protein TRIATDRAFT_236441, partial [Trichoderma atroviride IMI 206040] | 33.5 | 2.0E-111 |
| DIP_10000676 | hypothetical protein GLRG_07364 [Colletotrichum graminicola M1.001] | 47.1 | 2.0E-60 |
| DIP_10000677 | hypothetical protein PFICI_11363 [Pestalotiopsis fici W106-1] | 54.1 | 4.0E-123 |
| DIP_10000678 | armadillo repeat protein [Colletotrichum graminicola M1.001] | 42.8 | 2.0E-135 |
| DIP_10000679 | hypothetical protein PFICI_11368 [Pestalotiopsis fici W106-1] | 62.4 | 1.0E-146 |
| DIP_10000680 | hypothetical protein ACRE_045100 [Acremonium chrysogenum ATCC 11550] | 36.1 | 3.0E-29 |
| DIP_10000681 | hypothetical protein VDAG_06788 [Verticillium dahliae VdLs.17] | 48.7 | 7.0E-16 |
| DIP_10000682 | hypothetical protein DOTSEDRAFT_67492 [Dothistroma septosporum NZE10] | 53.8 | 3.0E-29 |
| DIP_10000683 | putative mfs sugar transporter protein [Eutypa lata UCREL1] | 70.2 | 0.0 |
| DIP_10000684 | hypothetical protein UCREL1_11270 [Eutypa lata UCREL1] | 53.5 | 8.0E-23 |
| DIP_10000685 | hypothetical protein PFICI_11255 [Pestalotiopsis fici W106-1] | 86.6 | 0.0 |
| DIP_10000686 | hypothetical protein CSUB01_06094 [Colletotrichum sublineola] | 44.4 | 0.0 |
| DIP_10000687 | putative sphingosine-1-phosphate lyase protein [Eutypa lata UCREL1] | 79.2 | 0.0 |
| DIP_10000688 | NA | | |
| DIP_10000689 | hypothetical protein PFICI_08856 [Pestalotiopsis fici W106-1] | 44.4 | 2.0E-112 |
| DIP_10000690 | hypothetical protein GLAREA_04354 [Glarea lozoyensis ATCC 20868] | 37.6 | 3.0E-48 |
| DIP_10000691 | NA | | |
| DIP_10000692 | NA | | |
| DIP_10000693 | hypothetical protein PFICI_09062 [Pestalotiopsis fici W106-1] | 57.5 | 4.0E-23 |
| DIP_10000694 | putative microtubule binding protein [Eutypa lata UCREL1] | 55 | 0.0 |
| DIP_10000695 | putative adenylate kinase protein [Eutypa lata UCREL1] | 74 | 4.0E-145 |
| DIP_10000696 | 26S proteasome regulatory subunit rpn-8 [Pestalotiopsis fici W106-1] | 85.3 | 0.0 |
| DIP_10000697 | hypothetical protein BAUCODRAFT_131212 [Baudoinia compniacensis UAMH 10762] | 32.8 | 1.0E-19 |
| DIP_10000698 | NA | | |
| DIP_10000699 | NA | | |
| DIP_10000700 | hypothetical protein NECHADRAFT_91510 [Nectria haematococca mpVI 77-13-4] | 39.5 | 4.0E-69 |
| DIP_10000701 | saponin hydrolase [Colletotrichum fioriniae PJ7] | 61.3 | 0.0 |
| DIP_10000702 | hypothetical protein CFIO01_10236 [Colletotrichum fioriniae PJ7] | 67.1 | 7.0E-130 |
| DIP_10000703 | NA | | |
| DIP_10000704 | ABA 3 protein [Aspergillus kawachii IFO 4308] | 64.7 | 0.0 |
| DIP_10000705 | putative cytochrome p450 [Diaporthe ampelina] | 49.2 | 3.0E-54 |
| DIP_10000706 | hypothetical protein PFICI_12381 [Pestalotiopsis fici W106-1] | 51.6 | 8.0E-120 |
| DIP_10000707 | hypothetical protein GGTG_05621 [Gaeumannomyces graminis var. tritici R3-111a-1] | 41 | 3.0E-31 |
| DIP_10000708 | NA | | |
| DIP_10000709 | hypothetical protein CGLO_12084 [Colletotrichum gloeosporioides Cg-14] | 71.2 | 0.0 |
| DIP_10000710 | NA | | |
| DIP_10000711 | phospholipase/carboxylesterase [Beauveria bassiana ARSEF 2860] | 45.2 | 3.0E-56 |
| DIP_10000712 | putative ubiquitin carboxyl-terminal hydrolase 14 protein [Eutypa lata UCREL1] | 77.7 | 0.0 |
| DIP_10000713 | putative iron sulfur cluster assembly protein 1 protein [Eutypa lata UCREL1] | 81.7 | 8.0E-110 |
| DIP_10000714 | NA | | |
| DIP_10000715 | hypothetical protein AURDEDRAFT_125994 [Auricularia delicata TFB-10046 SS5] | 52 | 3.0E-24 |
| DIP_10000716 | hypothetical protein PFICI_13175 [Pestalotiopsis fici W106-1] | 43.1 | 5.0E-129 |
| DIP_10000717 | hypothetical protein PFICI_13441 [Pestalotiopsis fici W106-1] | 61.8 | 0.0 |
| DIP_10000718 | ATP-dependent RNA helicase dbp3 [Pestalotiopsis fici W106-1] | 79.3 | 0.0 |
| DIP_10000719 | NA | | |
| DIP_10000720 | NA | | |
| DIP_10000721 | hypothetical protein UCREL1_4085 [Eutypa lata UCREL1] | 85.5 | 0.0 |
| DIP_10000722 | Mitotic spindle checkpoint component mad2 [Pestalotiopsis fici W106-1] | 80.3 | 4.0E-126 |
| DIP_10000723 | hypothetical protein PFICI_13151 [Pestalotiopsis fici W106-1] | 49.8 | 0.0 |
| DIP_10000724 | putative dna replication licensing factor mcm2 protein [Eutypa lata UCREL1] | 85.7 | 0.0 |
| DIP_10000725 | hypothetical protein PFICI_02690 [Pestalotiopsis fici W106-1] | 57.5 | 0.0 |
| DIP_10000726 | hypothetical protein PFICI_02688 [Pestalotiopsis fici W106-1] | 52.9 | 1.0E-150 |
| DIP_10000727 | putative fructose-bisphosphate aldolase protein [Togninia minima UCRPA7] | 76.2 | 6.0E-163 |
| DIP_10000728 | putative dihydroxyacetone kinase protein [Eutypa lata UCREL1] | 64.2 | 0.0 |
| DIP_10000729 | putative ribose 5-phosphate isomerase protein [Eutypa lata UCREL1] | 87.7 | 4.0E-95 |

| | | | |
|---|---|---|---|
| DIP_10000730 | triosephosphate isomerase [Magnaporthe oryzae 70-15] | 66.2 | 6.0E-109 |
| DIP_10000731 | NA | | |
| DIP_10000732 | hypothetical protein PFICI_13155 [Pestalotiopsis fici W106-1] | 55.4 | 0.0 |
| DIP_10000733 | NA | | |
| DIP_10000734 | hypothetical protein PFICI_02195 [Pestalotiopsis fici W106-1] | 64.4 | 3.0E-132 |
| DIP_10000735 | hypothetical protein GGTG_13115 [Gaeumannomyces graminis var. tritici R3-111a-1] | 34 | 2.0E-20 |
| DIP_10000736 | hypothetical protein MYCTH_2068129 [Myceliophthora thermophila ATCC 42464] | 59.2 | 0.0 |
| DIP_10000737 | hypothetical protein GGTG_02532 [Gaeumannomyces graminis var. tritici R3-111a-1] | 58.7 | 0.0 |
| DIP_10000738 | hypothetical protein PFICI_13920 [Pestalotiopsis fici W106-1] | 71.2 | 0.0 |
| DIP_10000739 | NA | | |
| DIP_10000740 | NA | | |
| DIP_10000741 | NA | | |
| DIP_10000742 | hypothetical protein PFICI_02133 [Pestalotiopsis fici W106-1] | 77.8 | 0.0 |
| DIP_10000743 | Trafficking protein particle complex subunit BET3 [Pestalotiopsis fici W106-1] | 92.2 | 1.0E-134 |
| DIP_10000744 | SOM1 protein [Colletotrichum fioriniae PJ7] | 51 | 0.0 |
| DIP_10000745 | NA | | |
| DIP_10000746 | zinc knuckle [Colletotrichum fioriniae PJ7] | 88.8 | 2.0E-126 |
| DIP_10000747 | hypothetical protein PFICI_02130 [Pestalotiopsis fici W106-1] | 41.3 | 3.0E-78 |
| DIP_10000748 | TOM core complex subunit [Ustilaginoidea virens] | 61.7 | 3.0E-18 |
| DIP_10000749 | hypothetical protein CGLO_06036 [Colletotrichum gloeosporioides Cg-14] | 81.9 | 0.0 |
| DIP_10000750 | NA | | |
| DIP_10000751 | putative short chain dehydrogenase family protein [Eutypa lata UCREL1] | 44.9 | 7.0E-64 |
| DIP_10000752 | putative major facilitator superfamily transporter protein [Eutypa lata UCREL1] | 51.8 | 0.0 |
| DIP_10000753 | hypothetical protein PFICI_02678 [Pestalotiopsis fici W106-1] | 57.8 | 0.0 |
| DIP_10000754 | putative glycoside hydrolase family 16 protein [Eutypa lata UCREL1] | 67.8 | 4.0E-130 |
| DIP_10000755 | putative short-chain dehydrogenase reductase sdr protein [Eutypa lata UCREL1] | 60.6 | 3.0E-115 |
| DIP_10000756 | putative calpain family cysteine protease protein [Eutypa lata UCREL1] | 55.4 | 0.0 |
| DIP_10000757 | putative kinetochore protein [Eutypa lata UCREL1] | 48.1 | 1.0E-65 |
| DIP_10000758 | NA | | |
| DIP_10000759 | putative mfs transporter protein [Eutypa lata UCREL1] | 79.4 | 0.0 |
| DIP_10000760 | hypothetical protein UCREL1_7756 [Eutypa lata UCREL1] | 54.4 | 0.0 |
| DIP_10000761 | hypothetical protein PFICI_10284 [Pestalotiopsis fici W106-1] | 79.3 | 1.0E-161 |
| DIP_10000762 | NA | | |
| DIP_10000763 | ESCRT-I component [Verticillium alfalfae VaMs.102] | 56.2 | 9.0E-171 |
| DIP_10000764 | putative indoleamine -dioxygenase protein [Eutypa lata UCREL1] | 52.8 | 0.0 |
| DIP_10000765 | NA | | |
| DIP_10000766 | myo-inositol 2-dehydrogenase [Cryptococcus gattii CA1873] | 69 | 0.0 |
| DIP_10000767 | hypothetical protein TREMEDRAFT_36755 [Tremella mesenterica DSM 1558] | 72.5 | 0.0 |
| DIP_10000768 | hypothetical protein CNBI0040 [Cryptococcus neoformans var. neoformans B-3501A] | 50 | 3.0E-126 |
| DIP_10000769 | putative sugar transporter stl1 [Phaeomoniella chlamydospora] | 50.7 | 0.0 |
| DIP_10000770 | dihydrodipicolinate synthase [Trichosporon asahii var. asahii CBS 8904] | 57.4 | 4.0E-132 |
| DIP_10000771 | hypothetical protein PDE_07957 [Penicillium oxalicum 114-2] | 66.9 | 0.0 |
| DIP_10000772 | L-carnitine dehydrogenase-like protein [Acremonium chrysogenum ATCC 11550] | 56.3 | 1.0E-116 |
| DIP_10000773 | hypothetical protein AN2721.2 [Aspergillus nidulans FGSC A4] | 66.8 | 2.0E-127 |
| DIP_10000774 | hypothetical protein AN2723.2 [Aspergillus nidulans FGSC A4] | 76.4 | 0.0 |
| DIP_10000775 | hypothetical protein [Cryptococcus neoformans var. neoformans JEC21] | 34.6 | 2.0E-64 |
| DIP_10000776 | hypothetical protein PV11_00953 [Exophiala sideris] | 70.2 | 2.0E-160 |
| DIP_10000777 | hypothetical protein PV05_01751 [Exophiala xenobiotica] | 66.8 | 0.0 |
| DIP_10000778 | NA | | |
| DIP_10000779 | NA | | |
| DIP_10000780 | NA | | |
| DIP_10000781 | putative inositol oxygenase 1 protein [Eutypa lata UCREL1] | 81.2 | 0.0 |
| DIP_10000782 | hypothetical protein PFICI_02676 [Pestalotiopsis fici W106-1] | 66 | 0.0 |
| DIP_10000783 | putative maltose permease mal61 protein [Eutypa lata UCREL1] | 77.9 | 0.0 |
| DIP_10000784 | NA | | |
| DIP_10000785 | NA | | |
| DIP_10000786 | hypothetical protein GLRG_01904 [Colletotrichum graminicola M1.001] | 64.3 | 0.0 |
| DIP_10000787 | pfs domain-containing protein [Colletotrichum higginsianum] | 44.5 | 2.0E-101 |
| DIP_10000788 | hypothetical protein TRIVIDRAFT_136184, partial [Trichoderma virens Gv29-8] | 50 | 2.0E-86 |
| DIP_10000789 | NA | | |
| DIP_10000790 | NA | | |
| DIP_10000791 | NA | | |
| DIP_10000792 | NA | | |
| DIP_10000793 | hypothetical protein V495_07393 [Pseudogymnoascus pannorum VKM F-4514 (FW-929)] | 31.9 | 1.0E-22 |
| DIP_10000794 | putative u4 u6 snrna-associated-splicing factor prp24 protein [Eutypa lata UCREL1] | 62.1 | 0.0 |
| DIP_10000795 | NA | | |
| DIP_10000796 | hypothetical protein UV8b_3126 [Ustilaginoidea virens] | 30.7 | 1.0E-11 |
| DIP_10000797 | NA | | |
| DIP_10000798 | NA | | |
| DIP_10000799 | hypothetical protein FOXB_09165 [Fusarium oxysporum Fo5176] | 96.5 | 0.0 |
| DIP_10000800 | putative bromodomain associated domain-containing protein [Eutypa lata UCREL1] | 64 | 1.0E-119 |
| DIP_10000801 | hypothetical protein PFICI_13191 [Pestalotiopsis fici W106-1] | 45.4 | 2.0E-141 |
| DIP_10000802 | putative isochorismatase family protein family protein [Eutypa lata UCREL1] | 45.6 | 1.0E-70 |
| DIP_10000803 | putative isochorismatase family protein family protein [Eutypa lata UCREL1] | 41.9 | 1.0E-106 |
| DIP_10000804 | hypothetical protein VDAG_06697 [Verticillium dahliae VdLs.17] | 35.6 | 2.0E-28 |
| DIP_10000805 | uncharacterized protein CPUR_05599 [Claviceps purpurea 20.1] | 56.1 | 2.0E-32 |
| DIP_10000806 | hypothetical protein NCU05816 [Neurospora crassa OR74A] | 77.2 | 6.0E-29 |
| DIP_10000807 | putative mitochondrial large ribosomal subunit protein [Eutypa lata UCREL1] | 56.9 | 6.0E-123 |
| DIP_10000808 | putative pheromone receptor 2 protein [Eutypa lata UCREL1] | 54.7 | 1.0E-104 |
| DIP_10000809 | hypothetical protein PFICI_14276 [Pestalotiopsis fici W106-1] | 82.8 | 1.0E-113 |
| DIP_10000810 | putative cytochrome p450 oxidoreductase [Diaporthe ampelina] | 53.4 | 0.0 |
| DIP_10000811 | ATP dependent RNA helicase [Sporothrix brasiliensis 5110] | 61.5 | 0.0 |
| DIP_10000812 | translational activator gcn1 [Colletotrichum gloeosporioides Nara gc5] | 75.5 | 0.0 |
| DIP_10000813 | NAD dependent epimerase/dehydratase family protein [Zymoseptoria brevis] | 56.4 | 1.0E-139 |
| DIP_10000814 | hypothetical protein MYCFIDRAFT_86556 [Pseudocercospora fijiensis CIRAD86] | 59.2 | 0.0 |
| DIP_10000815 | serine hydroxymethyltransferase [Pestalotiopsis fici W106-1] | 81.1 | 0.0 |
| DIP_10000816 | NA | | |
| DIP_10000817 | putative phosphatidyl synthase [Erysiphe necator] | 44.5 | 2.0E-22 |
| DIP_10000818 | hypothetical protein PFICI_03100 [Pestalotiopsis fici W106-1] | 77.6 | 5.0E-140 |
| DIP_10000819 | hypothetical protein PFICI_03101 [Pestalotiopsis fici W106-1] | 86.8 | 0.0 |
| DIP_10000820 | C6 transcription factor [Colletotrichum gloeosporioides Nara gc5] | 63.4 | 0.0 |
| DIP_10000821 | putative s- glutathione dehydrogenase protein [Eutypa lata UCREL1] | 78.6 | 0.0 |
| DIP_10000822 | hypothetical protein V490_05231 [Pseudogymnoascus pannorum VKM F-3557] | 73.9 | 6.0E-40 |
| DIP_10000823 | vacuolar ATPase [Colletotrichum gloeosporioides Nara gc5] | 85 | 8.0E-62 |
| DIP_10000824 | putative duf1716 domain-containing protein [Togninia minima UCRPA7] | 52.5 | 0.0 |
| DIP_10000825 | putative phosphoglycerate mutase protein [Eutypa lata UCREL1] | 63.9 | 1.0E-104 |
| DIP_10000826 | unnamed protein product [Aspergillus oryzae RIB40] | 60.2 | 6.0E-43 |
| DIP_10000827 | putative dna-directed rna polymerase ii subunit rpb2 protein [Eutypa lata UCREL1] | 88.4 | 0.0 |
| DIP_10000828 | putative glycoside hydrolase family 2 protein [Eutypa lata UCREL1] | 69 | 0.0 |
| DIP_10000829 | putative barwin-like endoglucanase protein [Eutypa lata UCREL1] | 53.9 | 4.0E-32 |
| DIP_10000830 | putative duf250 domain membrane protein [Eutypa lata UCREL1] | 82.3 | 0.0 |
| DIP_10000831 | UDP-galactopyranose mutase [Magnaporthe oryzae Y34] | 84.3 | 0.0 |
| DIP_10000832 | putative hsp90-like protein [Colletotrichum sublineola] | 59 | 0.0 |
| DIP_10000833 | putative 60s ribosomal protein l27 protein [Eutypa lata UCREL1] | 72.9 | 1.0E-82 |
| DIP_10000834 | hypothetical protein NCU01828 [Neurospora crassa OR74A] | 64.4 | 5.0E-22 |
| DIP_10000835 | hypothetical protein VDAG_03920 [Verticillium dahliae VdLs.17] | 54.2 | 5.0E-92 |
| DIP_10000836 | 4-hydroxyphenylpyruvate dioxygenase [Pestalotiopsis fici W106-1] | 67.4 | 0.0 |
| DIP_10000837 | NA | | |
| DIP_10000838 | hypothetical protein TRIATDRAFT_88643 [Trichoderma atroviride IMI 206040] | 41.6 | 4.0E-100 |
| DIP_10000839 | putative tfiih basal transcription factor complex p47 subunit protein [Eutypa lata UCREL1] | 73.8 | 0.0 |
| DIP_10000840 | putative o-acetylhomoserine -lyase protein [Togninia minima UCRPA7] | 77.4 | 0.0 |
| DIP_10000841 | hypothetical protein PFICI_11406 [Pestalotiopsis fici W106-1] | 59.6 | 2.0E-58 |
| DIP_10000842 | hypothetical protein THITE_2085426 [Thielavia terrestris NRRL 8126] | 50.8 | 1.0E-102 |
| DIP_10000843 | NA | | |
| DIP_10000844 | C6 zinc finger protein [Colletotrichum higginsianum] | 44.6 | 3.0E-129 |
| DIP_10000845 | hypothetical protein S40285_03372 [Stachybotrys chlorohalonata IBT 40285] | 36.7 | 4.0E-39 |
| DIP_10000846 | hypothetical protein PFICI_11377 [Pestalotiopsis fici W106-1] | 73.4 | 0.0 |
| DIP_10000847 | NA | | |
| DIP_10000848 | NA | | |
| DIP_10000849 | putative hydroxyproline-rich glyco protein [Eutypa lata UCREL1] | 51.2 | 4.0E-105 |
| DIP_10000850 | putative charged multivesicular body protein 3 [Diaporthe ampelina] | 30.8 | 5.0E-10 |
| DIP_10000851 | putative protein wos2 [Ustilaginoidea virens] | 32.3 | 5.0E-21 |
| DIP_10000852 | SNF7 family protein [Neurospora crassa OR74A] | 52.5 | 5.0E-78 |
| DIP_10000853 | hypothetical protein UCREL1_4982 [Eutypa lata UCREL1] | 60.6 | 3.0E-147 |
| DIP_10000854 | hypothetical protein PFICI_08971 [Pestalotiopsis fici W106-1] | 65.1 | 8.0E-63 |
| DIP_10000855 | hypothetical protein PFICI_08972 [Pestalotiopsis fici W106-1] | 63.6 | 2.0E-80 |
| DIP_10000856 | putative pseudouridine synthase 3 protein [Eutypa lata UCREL1] | 46.2 | 7.0E-157 |
| DIP_10000857 | GPI-GlcNAc transferase complex [Colletotrichum fioriniae PJ7] | 44.6 | 3.0E-52 |
| DIP_10000858 | hypothetical protein PFICI_08975 [Pestalotiopsis fici W106-1] | 60.1 | 0.0 |
| DIP_10000859 | hypothetical protein CH063_08146 [Colletotrichum higginsianum] | 52.5 | 2.0E-35 |
| DIP_10000860 | NA | | |
| DIP_10000861 | hypothetical protein H072_11364 [Dactylellina haptotyla CBS 200.50] | 31.5 | 2.0E-25 |
| DIP_10000862 | NA | | |
| DIP_10000863 | putative f-box wd-40 repeat-containing protein [Eutypa lata UCREL1] | 66.8 | 0.0 |
| DIP_10000864 | putative rna recognition motif containing protein [Eutypa lata UCREL1] | 36.5 | 2.0E-54 |
| DIP_10000865 | putative isoflavone reductase family protein [Eutypa lata UCREL1] | 49.6 | 2.0E-102 |
| DIP_10000866 | hypothetical protein NECHADRAFT_50811 [Nectria haematococca mpVI 77-13-4] | 41.3 | 4.0E-36 |
| DIP_10000867 | NA | | |
| DIP_10000868 | NA | | |
| DIP_10000869 | ribosomal protein L16p/L10e [Colletotrichum gloeosporioides Cg-14] | 68.3 | 5.0E-112 |
| DIP_10000870 | putative replication factor rfc1 c terminal domain-containing protein [Eutypa lata UCREL1] | 72.1 | 0.0 |
| DIP_10000871 | NA | | |
| DIP_10000872 | NA | | |
| DIP_10000873 | putative vacuolar transporter chaperone 2 protein [Eutypa lata UCREL1] | 68.1 | 0.0 |
| DIP_10000874 | hypothetical protein CH063_13662 [Colletotrichum higginsianum] | 61.1 | 1.0E-25 |
| DIP_10000875 | CDC24 Calponin [Colletotrichum higginsianum] | 58.2 | 0.0 |
| DIP_10000876 | related to tol protein [Fusarium fujikuroi IMI 58289] | 30.2 | 7.0E-50 |
| DIP_10000877 | predicted protein [Trichoderma reesei QM6a] | 44.3 | 3.0E-75 |
| DIP_10000878 | NA | | |
| DIP_10000879 | NA | | |
| DIP_10000880 | NA | | |
| DIP_10000881 | Major Facilitator Superfamily protein [Coccidioides posadasii C735 delta SOWgp] | 62.8 | 0.0 |
| DIP_10000882 | hypothetical protein PFICI_11397 [Pestalotiopsis fici W106-1] | 57 | 0.0 |
| DIP_10000883 | predicted protein [Nectria haematococca mpVI 77-13-4] | 68.2 | 1.0E-126 |
| DIP_10000884 | putative ankyrin repeat protein [Togninia minima UCRPA7] | 40.9 | 0.0 |
| DIP_10000885 | NA | | |
| DIP_10000886 | hypothetical protein FGSG_06003 [Fusarium graminearum PH-1] | 67 | 9.0E-144 |
| DIP_10000887 | predicted protein [Nectria haematococca mpVI 77-13-4] | 74.4 | 0.0 |
| DIP_10000888 | putative dolichyl-phosphate-mannose-protein mannosyltransferase 2 protein [Eutypa lata UCREL1] | 86.9 | 0.0 |
| DIP_10000889 | hypothetical protein PFICI_11400 [Pestalotiopsis fici W106-1] | 69.4 | 0.0 |
| DIP_10000890 | hypothetical protein THITE_2071706 [Thielavia terrestris NRRL 8126] | 55.1 | 0.0 |
| DIP_10000891 | putative ribonuclease h protein [Eutypa lata UCREL1] | 67 | 2.0E-144 |
| DIP_10000892 | putative cytidine deaminase-like protein [Eutypa lata UCREL1] | 64.1 | 0.0 |
| DIP_10000893 | hypothetical protein SPBR_01899 [Sporothrix brasiliensis 5110] | 46.9 | 2.0E-26 |
| DIP_10000894 | hypothetical protein UCREL1_9799 [Eutypa lata UCREL1] | 55.5 | 4.0E-70 |
| DIP_10000895 | NA | | |
| DIP_10000896 | NA | | |
| DIP_10000897 | hypothetical protein ANO11243_014040 [fungal sp. No.11243] | 55.9 | 4.0E-82 |
| DIP_10000898 | NA | | |
| DIP_10000899 | NA | | |
| DIP_10000900 | Leucine--tRNA ligase [Pestalotiopsis fici W106-1] | 80.1 | 0.0 |
| DIP_10000901 | hypothetical protein PFICI_09210 [Pestalotiopsis fici W106-1] | 65.1 | 1.0E-52 |
| DIP_10000902 | hypothetical protein MAA_11079 [Metarhizium robertsii ARSEF 23] | 85.9 | 2.0E-119 |
| DIP_10000903 | bZIP transcription factor, bZIP-1 [Beauveria bassiana ARSEF 2860] | 39.6 | 4.0E-19 |
| DIP_10000904 | NA | | |
| DIP_10000905 | hypothetical protein PFICI_09311 [Pestalotiopsis fici W106-1] | 68.6 | 0.0 |
| DIP_10000906 | putative uba domain-containing protein ucp14 protein [Eutypa lata UCREL1] | 46.3 | 7.0E-87 |
| DIP_10000907 | NA | | |
| DIP_10000908 | putative deacetylase complex subunit protein [Eutypa lata UCREL1] | 55.9 | 0.0 |
| DIP_10000909 | hypothetical protein PFICI_08947 [Pestalotiopsis fici W106-1] | 33.5 | 1.0E-73 |
| DIP_10000910 | hypothetical protein A1O5_00732 [Cladophialophora psammophila CBS 110553] | 73.2 | 0.0 |
| DIP_10000911 | hypothetical protein PFICI_10890 [Pestalotiopsis fici W106-1] | 67.6 | 0.0 |
| DIP_10000912 | hypothetical protein PFICI_10892 [Pestalotiopsis fici W106-1] | 54.1 | 0.0 |
| DIP_10000913 | hypothetical protein PFICI_10893 [Pestalotiopsis fici W106-1] | 66.1 | 0.0 |
| DIP_10000914 | hypothetical protein PFICI_09310 [Pestalotiopsis fici W106-1] | 48.4 | 2.0E-59 |
| DIP_10000915 | hypothetical protein PFICI_09309 [Pestalotiopsis fici W106-1] | 45.6 | 2.0E-92 |
| DIP_10000916 | putative sugar transporter stl1 protein [Eutypa lata UCREL1] | 63 | 0.0 |
| DIP_10000917 | hypothetical protein PV10_00718 [Exophiala mesophila] | 34.5 | 2.0E-92 |
| DIP_10000918 | 3-carboxy-cis,cis-muconate cycloisomerase [Exophiala sideris] | 71.3 | 0.0 |
| DIP_10000919 | NA | | |
| DIP_10000920 | putative plasma membrane fusion protein prm1 protein [Eutypa lata UCREL1] | 56.4 | 0.0 |
| DIP_10000921 | NA | | |
| DIP_10000922 | NA | | |
| DIP_10000923 | putative kh domain-containing protein [Eutypa lata UCREL1] | 65.7 | 0.0 |
| DIP_10000924 | hypothetical protein PFICI_09306 [Pestalotiopsis fici W106-1] | 71.8 | 0.0 |
| DIP_10000925 | hypothetical protein PFICI_09305 [Pestalotiopsis fici W106-1] | 64.2 | 3.0E-113 |
| DIP_10000926 | putative mitochondrial carrier protein pet8 protein [Eutypa lata UCREL1] | 59.5 | 2.0E-40 |
| DIP_10000927 | NA | | |
| DIP_10000928 | predicted protein [Trichoderma reesei QM6a] | 36.6 | 2.0E-32 |
| DIP_10000929 | uncharacterized protein FFUJ_09819 [Fusarium fujikuroi IMI 58289] | 70.2 | 3.0E-80 |
| DIP_10000930 | NA | | |
| DIP_10000931 | cyclin-C [Verticillium alfalfae VaMs.102] | 64.7 | 7.0E-155 |
| DIP_10000932 | putative calcium dependent mitochondrial carrier protein [Eutypa lata UCREL1] | 66.5 | 0.0 |
| DIP_10000933 | hypothetical protein FGSG_04300 [Fusarium graminearum PH-1] | 39.5 | 4.0E-52 |
| DIP_10000934 | predicted protein [Nectria haematococca mpVI 77-13-4] | 71.9 | 0.0 |
| DIP_10000935 | hypothetical protein V492_01959 [Pseudogymnoascus pannorum VKM F-4246] | 42.8 | 1.0E-101 |
| DIP_10000936 | NA | | |
| DIP_10000937 | NA | | |
| DIP_10000938 | hypothetical protein PFICI_09001 [Pestalotiopsis fici W106-1] | 48.1 | 2.0E-107 |
| DIP_10000939 | hypothetical protein CH063_08767 [Colletotrichum higginsianum] | 36.2 | 1.0E-63 |
| DIP_10000940 | hypothetical protein UCREL1_6517 [Eutypa lata UCREL1] | 32.4 | 9.0E-19 |
| DIP_10000941 | hypothetical protein PFICI_15190 [Pestalotiopsis fici W106-1] | 47.7 | 5.0E-105 |
| DIP_10000942 | hypothetical protein PFICI_14235 [Pestalotiopsis fici W106-1] | 31.4 | 3.0E-33 |
| DIP_10000943 | NA | | |
| DIP_10000944 | putative serine arginine repetitive matrix protein 1 protein [Eutypa lata UCREL1] | 57.7 | 2.0E-41 |
| DIP_10000945 | NA | | |
| DIP_10000946 | putative histone deacetylase hda1 protein [Eutypa lata UCREL1] | 71.9 | 0.0 |
| DIP_10000947 | putative f-box domain-containing protein [Eutypa lata UCREL1] | 31.7 | 2.0E-55 |
| DIP_10000948 | putative ribosome biogenesis protein brx1 protein [Togninia minima UCRPA7] | 80.3 | 0.0 |
| DIP_10000949 | Ras family protein [Beauveria bassiana ARSEF 2860] | 85.1 | 1.0E-141 |
| DIP_10000950 | putative pp-loop family protein [Togninia minima UCRPA7] | 48 | 1.0E-76 |
| DIP_10000951 | hypothetical protein THITE_2060163 [Thielavia terrestris NRRL 8126] | 68.9 | 2.0E-139 |
| DIP_10000952 | hypothetical protein BC1G_12823 [Botrytis cinerea B05.10] | 52.5 | 0.0 |
| DIP_10000953 | NA | | |
| DIP_10000954 | NA | | |
| DIP_10000955 | hypothetical protein PFICI_08966 [Pestalotiopsis fici W106-1] | 35.8 | 5.0E-86 |
| DIP_10000956 | NA | | |
| DIP_10000957 | NA | | |
| DIP_10000958 | hypothetical protein PFICI_08967 [Pestalotiopsis fici W106-1] | 51 | 9.0E-81 |
| DIP_10000959 | putative glycerol-3-phosphate dehydrogenase protein [Eutypa lata UCREL1] | 78 | 0.0 |
| DIP_10000960 | hypothetical protein PFICI_11459 [Pestalotiopsis fici W106-1] | 45.6 | 0.0 |
| DIP_10000961 | hypothetical protein FOVG_11731 [Fusarium oxysporum f. sp. pisi HDV247] | 65.5 | 8.0E-160 |
| DIP_10000962 | NA | | |
| DIP_10000963 | putative tfiih p62 subunit domain-containing protein [Eutypa lata UCREL1] | 66.4 | 0.0 |
| DIP_10000964 | putative oxoglutarate dehydrogenase protein [Eutypa lata UCREL1] | 85.8 | 0.0 |
| DIP_10000965 | leucine carboxyl methyltransferase, partial [Colletotrichum higginsianum] | 58.3 | 4.0E-151 |
| DIP_10000966 | Autophagy-related protein 18 [Pestalotiopsis fici W106-1] | 73 | 0.0 |
| DIP_10000967 | Pps1 dual specificity phosphatase [Trichophyton tonsurans CBS 112818] | 63.6 | 4.0E-87 |
| DIP_10000968 | hypothetical protein H634G_01513 [Metarhizium anisopliae BRIP 53293] | 42 | 2.0E-140 |
| DIP_10000969 | epithelial zinc-finger erf protein [Ophiostoma piceae UAMH 11346] | 48.4 | 2.0E-50 |
| DIP_10000970 | putative dna repair helicase rad15 protein [Eutypa lata UCREL1] | 89.6 | 0.0 |
| DIP_10000971 | putative u4 u6 small nuclear ribonucleoprotein prp3 protein [Eutypa lata UCREL1] | 71.8 | 0.0 |
| DIP_10000972 | putative dna-directed rna polymerase i subunit rpa1 protein [Eutypa lata UCREL1] | 73.8 | 0.0 |
| DIP_10000973 | conserved hypothetical protein [Aspergillus terreus NIH2624] | 49.8 | 2.0E-68 |
| DIP_10000974 | histone H3 [Neurospora crassa OR74A] | 100 | 7.0E-96 |
| DIP_10000975 | histone H4 [Botrytis cinerea B05.10] | 100 | 2.0E-67 |
| DIP_10000976 | hypothetical protein [Podospora anserina S mat+] | 44.6 | 1.0E-116 |
| DIP_10000977 | putative pentafunctional arom polypeptide protein [Eutypa lata UCREL1] | 75.3 | 0.0 |
| DIP_10000978 | hypothetical protein PFICI_07308 [Pestalotiopsis fici W106-1] | 52.5 | 5.0E-75 |
| DIP_10000979 | NA | | |
| DIP_10000980 | hypothetical protein UCREL1_2595 [Eutypa lata UCREL1] | 35 | 8.0E-62 |
| DIP_10000981 | NA | | |
| DIP_10000982 | hypothetical protein PFICI_07015 [Pestalotiopsis fici W106-1] | 68 | 0.0 |
| DIP_10000983 | putative threonine dehydratase protein [Eutypa lata UCREL1] | 85.2 | 0.0 |
| DIP_10000984 | hypothetical protein [Podospora anserina S mat+] | 52.6 | 4.0E-61 |
| DIP_10000985 | putative tldc domain containing protein 2 protein [Eutypa lata UCREL1] | 71.5 | 2.0E-120 |
| DIP_10000986 | putative rpel repeat protein [Eutypa lata UCREL1] | 80.2 | 5.0E-59 |
| DIP_10000987 | putative alpha-n-arabinofuranosidase a protein [Eutypa lata UCREL1] | 64.5 | 0.0 |
| DIP_10000988 | putative impb mucb samb family protein [Eutypa lata UCREL1] | 72.3 | 0.0 |
| DIP_10000989 | putative alanine racemase domain protein [Togninia minima UCRPA7] | 59.5 | 6.0E-136 |
| DIP_10000990 | lipase 2 [Magnaporthe oryzae 70-15] | 55.8 | 3.0E-138 |
| DIP_10000991 | hypothetical protein TRIVIDRAFT_74115 [Trichoderma virens Gv29-8] | 60.7 | 7.0E-56 |
| DIP_10000992 | putative mfs transporter protein [Eutypa lata UCREL1] | 68.7 | 0.0 |
| DIP_10000993 | fkbp-type peptidyl-prolyl [Colletotrichum orbiculare MAFF 240422] | 60.4 | 2.0E-79 |
| DIP_10000994 | NA | | |
| DIP_10000995 | hypothetical protein PV04_03465 [Capronia semiimmersa] | 44.6 | 2.0E-72 |
| DIP_10000996 | hypothetical protein PFICI_06810 [Pestalotiopsis fici W106-1] | 36.3 | 7.0E-21 |
| DIP_10000997 | hypothetical protein FVEG_04050 [Fusarium verticillioides 7600] | 70 | 1.0E-86 |
| DIP_10000998 | hypothetical protein GLRG_05471 [Colletotrichum graminicola M1.001] | 62.3 | 0.0 |
| DIP_10000999 | putative snare complex subunit protein [Eutypa lata UCREL1] | 85.2 | 4.0E-58 |
| DIP_10001000 | putative nonribosomal peptide synthetase 2 protein [Neofusicoccum parvum UCRNP2] | 40.8 | 0.0 |
| DIP_10001001 | putative l-ornithine 5-monooxygenase (l-ornithine n -oxygenase) protein [Eutypa lata UCREL1] | 69.8 | 0.0 |
| DIP_10001002 | NA | | |
| DIP_10001003 | putative udp-glucose dehydrogenase protein [Eutypa lata UCREL1] | 77.5 | 0.0 |
| DIP_10001004 | hypothetical protein HMPREF1541_11131 [Cyphellophora europaea CBS 101466] | 63.5 | 2.0E-158 |
| DIP_10001005 | hypothetical protein HMPREF1541_11132 [Cyphellophora europaea CBS 101466] | 54.7 | 0.0 |
| DIP_10001006 | hypothetical protein HMPREF1541_11133 [Cyphellophora europaea CBS 101466] | 39.4 | 1.0E-123 |
| DIP_10001007 | hypothetical protein UCREL1_2595 [Eutypa lata UCREL1] | 37.1 | 1.0E-69 |
| DIP_10001008 | hypothetical protein PFICI_09984 [Pestalotiopsis fici W106-1] | 71.2 | 0.0 |
| DIP_10001009 | Cytochrome c heme lyase [Pestalotiopsis fici W106-1] | 68.3 | 1.0E-163 |
| DIP_10001010 | NA | | |
| DIP_10001011 | hypothetical protein PFICI_07287 [Pestalotiopsis fici W106-1] | 48.6 | 2.0E-131 |
| DIP_10001012 | hypothetical protein A1O1_01660 [Capronia coronata CBS 617.96] | 35.8 | 1.0E-53 |
| DIP_10001013 | putative carbohydrate binding family 6 protein [Eutypa lata UCREL1] | 52.8 | 6.0E-99 |
| DIP_10001014 | NA | | |
| DIP_10001015 | vegetative cell wall protein gp1 [Colletotrichum higginsianum] | 56.2 | 5.0E-141 |
| DIP_10001016 | hypothetical protein PFICI_10066 [Pestalotiopsis fici W106-1] | 55.1 | 6.0E-94 |
| DIP_10001017 | hypothetical protein PFICI_05503 [Pestalotiopsis fici W106-1] | 45.2 | 4.0E-137 |
| DIP_10001018 | hypothetical protein COCHEDRAFT_1185799 [Bipolaris maydis C5] | 37.9 | 6.0E-23 |
| DIP_10001019 | hypothetical protein MGG_00443 [Magnaporthe oryzae 70-15] | 69.7 | 7.0E-24 |
| DIP_10001020 | putative rna recognition domain-containing protein [Eutypa lata UCREL1] | 62.3 | 3.0E-120 |
| DIP_10001021 | YOR378W-like protein [Saccharomyces cerevisiae x Saccharomyces kudriavzevii VIN7] | 41.1 | 4.0E-106 |
| DIP_10001022 | NA | | |
| DIP_10001023 | hypothetical protein SAPIO_CDS10918 [Scedosporium apiospermum] | 45.8 | 5.0E-88 |
| DIP_10001024 | hypothetical protein UCREL1_10528 [Eutypa lata UCREL1] | 50.3 | 2.0E-46 |
| DIP_10001025 | hypothetical protein FGSG_08082 [Fusarium graminearum PH-1] | 42.3 | 3.0E-43 |
| DIP_10001026 | NA | | |
| DIP_10001027 | hypothetical protein PFICI_10395 [Pestalotiopsis fici W106-1] | 82.2 | 2.0E-173 |
| DIP_10001028 | putative atp-dependent rna helicase mak5 protein [Eutypa lata UCREL1] | 70.1 | 0.0 |
| DIP_10001029 | NA | | |
| DIP_10001030 | hippurate hydrolase [Cordyceps militaris CM01] | 55.2 | 7.0E-163 |
| DIP_10001031 | putative chitinase protein [Eutypa lata UCREL1] | 66.4 | 0.0 |
| DIP_10001032 | hypothetical protein GGTG_00157 [Gaeumannomyces graminis var. tritici R3-111a-1] | 57.9 | 0.0 |
| DIP_10001033 | hypothetical protein PFICI_04524 [Pestalotiopsis fici W106-1] | 52.1 | 0.0 |
| DIP_10001034 | hypothetical protein PFICI_04523 [Pestalotiopsis fici W106-1] | 69.8 | 0.0 |
| DIP_10001035 | d-alanine ligase [Grosmannia clavigera kw1407] | 44.6 | 1.0E-97 |
| DIP_10001036 | hypothetical protein PFICI_04519 [Pestalotiopsis fici W106-1] | 76.4 | 5.0E-161 |
| DIP_10001037 | hypothetical protein A1O7_09106 [Cladophialophora yegresii CBS 114405] | 42.8 | 8.0E-103 |
| DIP_10001038 | putative mediator of rna polymerase ii transcription subunit 7 protein [Eutypa lata UCREL1] | 65.9 | 5.0E-101 |
| DIP_10001039 | hypothetical protein PFICI_15276 [Pestalotiopsis fici W106-1] | 40.2 | 3.0E-51 |
| DIP_10001040 | Protein hir-1 [Pestalotiopsis fici W106-1] | 75.8 | 0.0 |
| DIP_10001041 | hypothetical protein GLRG_06526 [Colletotrichum graminicola M1.001] | 54 | 9.0E-50 |
| DIP_10001042 | hypothetical protein FOC1_g10001821 [Fusarium oxysporum f. sp. cubense race 1] | 60 | 2.0E-43 |
| DIP_10001043 | NA | | |
| DIP_10001044 | hypothetical protein PFICI_10380 [Pestalotiopsis fici W106-1] | 56.8 | 0.0 |
| DIP_10001045 | NA | | |
| DIP_10001046 | NA | | |
| DIP_10001047 | C6 finger domain-containing protein [Colletotrichum gloeosporioides Nara gc5] | 42.2 | 5.0E-111 |
| DIP_10001048 | hypothetical protein PFICI_12860 [Pestalotiopsis fici W106-1] | 67.5 | 0.0 |
| DIP_10001049 | NA | | |
| DIP_10001050 | hypothetical protein A1O7_03339 [Cladophialophora yegresii CBS 114405] | 50.5 | 6.0E-139 |
| DIP_10001051 | hypothetical protein FOWG_14007 [Fusarium oxysporum f. sp. lycopersici MN25] | 52.6 | 3.0E-94 |
| DIP_10001052 | n-carbamoyl-l-amino acid hydrolase [Ophiostoma piceae UAMH 11346] | 57.5 | 0.0 |
| DIP_10001053 | putative diaminopropionate ammonia-lyase [Colletotrichum sublineola] | 49.6 | 4.0E-113 |
| DIP_10001054 | putative mms4 protein [Eutypa lata UCREL1] | 52.5 | 3.0E-76 |
| DIP_10001055 | putative adenosine amp deaminase protein [Togninia minima UCRPA7] | 40.7 | 2.0E-148 |
| DIP_10001056 | hypothetical protein UCREL1_1522 [Eutypa lata UCREL1] | 51.1 | 0.0 |
| DIP_10001057 | putative fad dependent oxidoreductase [Diaporthe ampelina] | 70.7 | 0.0 |
| DIP_10001058 | putative pho85 cyclin-1 protein [Eutypa lata UCREL1] | 74.4 | 0.0 |
| DIP_10001059 | putative cell division control protein 2 protein [Eutypa lata UCREL1] | 91.4 | 0.0 |
| DIP_10001060 | hypothetical protein PFICI_10250 [Pestalotiopsis fici W106-1] | 56.1 | 3.0E-156 |
| DIP_10001061 | putative filament-forming protein [Eutypa lata UCREL1] | 61.8 | 0.0 |
| DIP_10001062 | putative trimeric -like protein [Eutypa lata UCREL1] | 47.4 | 3.0E-45 |
| DIP_10001063 | putative nadh-ubiquinone oxidoreductase 51 kda subunit protein [Eutypa lata UCREL1] | 92.8 | 0.0 |
| DIP_10001064 | putative ex11b-like protein [Eutypa lata UCREL1] | 42.9 | 2.0E-75 |
| DIP_10001065 | NA | | |
| DIP_10001066 | NA | | |
| DIP_10001067 | NA | | |
| DIP_10001068 | putative hexaprenyldihydroxybenzoate methyltransferase protein [Eutypa lata UCREL1] | 62 | 1.0E-119 |
| DIP_10001069 | putative -dihydroxy-2-butanone 4-phosphate synthase protein [Eutypa lata UCREL1] | 78.8 | 2.0E-133 |
| DIP_10001070 | conserved hypothetical protein [Arthroderma otae CBS 113480] | 32.5 | 9.0E-72 |
| DIP_10001071 | NA | | |
| DIP_10001072 | NA | | |
| DIP_10001073 | C6 zinc finger domain containing protein [Metarhizium acridum CQMa 102] | 43.6 | 0.0 |
| DIP_10001074 | putative f-box domain-containing protein [Eutypa lata UCREL1] | 46.9 | 1.0E-112 |
| DIP_10001075 | hypothetical protein PFICI_15098 [Pestalotiopsis fici W106-1] | 66.2 | 5.0E-163 |
| DIP_10001076 | hypothetical protein FAVG1_04628 [Fusarium avenaceum] | 40.4 | 5.0E-66 |
| DIP_10001077 | NA | | |
| DIP_10001078 | NA | | |
| DIP_10001079 | putative glycoside hydrolase family 18 protein [Eutypa lata UCREL1] | 64.7 | 7.0E-153 |
| DIP_10001080 | Elongation factor G [Pestalotiopsis fici W106-1] | 80.9 | 0.0 |
| DIP_10001081 | NA | | |
| DIP_10001082 | hypothetical protein S40293_01100 [Stachybotrys chartarum IBT 40293] | 49.4 | 5.0E-74 |
| DIP_10001083 | hypothetical protein NEUTE1DRAFT_150702 [Neurospora tetrasperma FGSC 2508] | 42.9 | 5.0E-95 |
| DIP_10001084 | putative aminotransferase class i and ii protein [Eutypa lata UCREL1] | 57 | 1.0E-167 |
| DIP_10001085 | PREDICTED: lipase-like [Pyrus x bretschneideri] | 30.2 | 6.0E-12 |
| DIP_10001086 | hypothetical protein UCREL1_11462 [Eutypa lata UCREL1] | 49.7 | 4.0E-50 |
| DIP_10001087 | putative atp-binding cassette sub-family g member 2 protein [Eutypa lata UCREL1] | 82.9 | 0.0 |
| DIP_10001088 | hypothetical protein UCREL1_9941 [Eutypa lata UCREL1] | 46.9 | 3.0E-58 |
| DIP_10001089 | Nucleolar GTP-binding protein 2 [Pestalotiopsis fici W106-1] | 73.3 | 0.0 |
| DIP_10001090 | acid phosphatase [Microsporum gypseum CBS 118893] | 33.1 | 5.0E-29 |
| DIP_10001091 | putative homoisocitrate dehydrogenase [Pestalotiopsis fici W106-1] | 81.3 | 0.0 |
| DIP_10001092 | hypothetical protein PFICI_15113 [Pestalotiopsis fici W106-1] | 37.7 | 2.0E-73 |
| DIP_10001093 | putative mitochondrial 37s ribosomal protein s8 protein [Eutypa lata UCREL1] | 63.1 | 1.0E-59 |
| DIP_10001094 | putative rna recognition motif-containing protein [Eutypa lata UCREL1] | 57.3 | 6.0E-147 |
| DIP_10001095 | NA | | |

| | | | |
|---|---|---|---|
| DIP_10001096 | hypothetical protein UCREL1_9934 [Eutypa lata UCREL1] | 46.5 | 2.0E-97 |
| DIP_10001097 | putative mfs monocarboxylate transporter protein [Eutypa lata UCREL1] | 65.4 | 1.0E-179 |
| DIP_10001098 | hypothetical protein FOZG_17149 [Fusarium oxysporum Fo47] | 38.5 | 2.0E-27 |
| DIP_10001099 | hypothetical protein PFICI_07013 [Pestalotiopsis fici W106-1] | 33.3 | 1.0E-18 |
| DIP_10001100 | arylsulfatase [Capronia epimyces CBS 606.96] | 69.2 | 3.0E-178 |
| DIP_10001101 | putative 26s proteasome regulatory subunit rpn2 protein [Eutypa lata UCREL1] | 81.2 | 0.0 |
| DIP_10001102 | putative mitochondrial peroxiredoxin prx1 protein [Eutypa lata UCREL1] | 87.3 | 1.0E-147 |
| DIP_10001103 | ral2 [Verticillium dahliae VdLs.17] | 62.7 | 0.0 |
| DIP_10001104 | NA | | |
| DIP_10001105 | hypothetical protein CSUB01_00502 [Colletotrichum sublineola] | 45.3 | 7.0E-119 |
| DIP_10001106 | hypothetical protein PFICI_15151 [Pestalotiopsis fici W106-1] | 68.4 | 0.0 |
| DIP_10001107 | putative glycylpeptide n-tetradecanoyltransferase protein [Eutypa lata UCREL1] | 81 | 0.0 |
| DIP_10001108 | hypothetical protein BGHDH14_bgh01817 [Blumeria graminis f. sp. hordei DH14] | 33 | 3.0E-54 |
| DIP_10001109 | hypothetical protein PFICI_13517 [Pestalotiopsis fici W106-1] | 56.3 | 4.0E-145 |
| DIP_10001110 | putative nucleolar atpase protein [Eutypa lata UCREL1] | 78.9 | 0.0 |
| DIP_10001111 | hypothetical protein HK57_00464 [Aspergillus ustus] | 34 | 4.0E-15 |
| DIP_10001112 | putative ef hand domain-containing protein [Eutypa lata UCREL1] | 64.4 | 0.0 |
| DIP_10001113 | hypothetical protein PFICI_04753 [Pestalotiopsis fici W106-1] | 63.1 | 7.0E-135 |
| DIP_10001114 | hypothetical protein UCREL1_10921 [Eutypa lata UCREL1] | 76.3 | 6.0E-158 |
| DIP_10001115 | hypothetical protein PFICI_15149 [Pestalotiopsis fici W106-1] | 35.3 | 4.0E-130 |
| DIP_10001116 | MFS sugar transporter [Colletotrichum orbiculare MAFF 240422] | 66.1 | 0.0 |
| DIP_10001117 | hypothetical protein PFICI_13724 [Pestalotiopsis fici W106-1] | 64.7 | 9.0E-117 |
| DIP_10001118 | hypothetical protein PFICI_13732 [Pestalotiopsis fici W106-1] | 72.1 | 5.0E-143 |
| DIP_10001119 | hypothetical protein FPSE_05315 [Fusarium pseudograminearum CS3096] | 31.5 | 4.0E-29 |
| DIP_10001120 | Kynurenine 3-monooxygenase [Pestalotiopsis fici W106-1] | 75.7 | 0.0 |
| DIP_10001121 | putative methionine aminopeptidase protein [Eutypa lata UCREL1] | 83.2 | 0.0 |
| DIP_10001122 | ER-derived vesicles protein ERV14 [Pestalotiopsis fici W106-1] | 94.2 | 6.0E-92 |
| DIP_10001123 | putative pre-mrna-processing factor 17 protein [Eutypa lata UCREL1] | 78.8 | 0.0 |
| DIP_10001124 | putative tartrate dehydrogenase protein [Eutypa lata UCREL1] | 83.8 | 0.0 |
| DIP_10001125 | fungal specific transcription factor domain-containing protein [Trichoderma reesei QM6a] | 46.2 | 2.0E-115 |
| DIP_10001126 | hypothetical protein PFICI_13557 [Pestalotiopsis fici W106-1] | 52 | 3.0E-65 |
| DIP_10001127 | hypothetical protein PV11_06153 [Exophiala sideris] | 47.7 | 7.0E-55 |
| DIP_10001128 | NA | | |
| DIP_10001129 | NA | | |
| DIP_10001130 | hypothetical protein MYCFIDRAFT_44275 [Pseudocercospora fijiensis CIRAD86] | 61.1 | 6.0E-132 |
| DIP_10001131 | hypothetical protein FAVG1_04489 [Fusarium avenaceum] | 38.5 | 2.0E-07 |
| DIP_10001132 | hypothetical protein HIM_00158 [Hirsutella minnesotensis 3608] | 38.2 | 2.0E-109 |
| DIP_10001133 | putative gnat family acetyltransferase protein [Eutypa lata UCREL1] | 54.1 | 3.0E-87 |
| DIP_10001134 | putative complex 1 protein [Colletotrichum sublineola] | 49.5 | 3.0E-22 |
| DIP_10001135 | hypothetical protein [Podospora anserina S mat+] | 50.7 | 9.0E-30 |
| DIP_10001136 | hypothetical protein PFICI_07025 [Pestalotiopsis fici W106-1] | 52.8 | 2.0E-122 |
| DIP_10001137 | hypothetical protein COCHEDRAFT_1077932, partial [Bipolaris maydis C5] | 33.3 | 3.0E-44 |
| DIP_10001138 | hypothetical protein PFICI_07026 [Pestalotiopsis fici W106-1] | 78.3 | 0.0 |
| DIP_10001139 | putative metallo-beta-lactamase family protein [Eutypa lata UCREL1] | 60.6 | 3.0E-138 |
| DIP_10001140 | hypothetical protein PFICI_07023 [Pestalotiopsis fici W106-1] | 73.1 | 0.0 |
| DIP_10001141 | methyltransferase [Colletotrichum fioriniae PJ7] | 57.6 | 2.0E-102 |
| DIP_10001142 | NA | | |
| DIP_10001143 | NA | | |
| DIP_10001144 | NA | | |
| DIP_10001145 | predicted protein [Nectria haematococca mpVI 77-13-4] | 33.9 | 5.0E-85 |
| DIP_10001146 | NA | | |
| DIP_10001147 | Sulfite oxidase [Beauveria bassiana D1-5] | 42.3 | 1.0E-40 |
| DIP_10001148 | NA | | |
| DIP_10001149 | NA | | |
| DIP_10001150 | putative mrna capping nucleoside- protein [Eutypa lata UCREL1] | 48 | 2.0E-152 |
| DIP_10001151 | NA | | |
| DIP_10001152 | hypothetical protein PFICI_15092 [Pestalotiopsis fici W106-1] | 65.6 | 1.0E-56 |
| DIP_10001153 | hypothetical protein [Podospora anserina S mat+] | 51.2 | 2.0E-147 |
| DIP_10001154 | hypothetical protein TRIVIDRAFT_134921, partial [Trichoderma virens Gv29-8] | 32 | 2.0E-49 |
| DIP_10001155 | putative glycerophosphoryl diester phosphodiesterase family protein [Eutypa lata UCREL1] | 58.1 | 3.0E-162 |
| DIP_10001156 | putative purine permease protein [Eutypa lata UCREL1] | 67.5 | 0.0 |
| DIP_10001157 | NA | | |
| DIP_10001158 | putative quinic acid utilization activator [Diaporthe ampelina] | 34.5 | 6.0E-96 |
| DIP_10001159 | Quinate repressor protein [Pestalotiopsis fici W106-1] | 52.3 | 0.0 |
| DIP_10001160 | hypothetical protein MYCTH_2309056 [Myceliophthora thermophila ATCC 42464] | 54.9 | 1.0E-149 |
| DIP_10001161 | Quinate permease [Pestalotiopsis fici W106-1] | 69 | 0.0 |
| DIP_10001162 | putative quinate dehydrogenase protein [Eutypa lata UCREL1] | 52.4 | 1.0E-121 |
| DIP_10001163 | phosphotransferase enzyme family protein [Mycobacterium lentiflavum] | 32.1 | 5.0E-93 |
| DIP_10001164 | NA | | |
| DIP_10001165 | QA-X protein [Verticillium dahliae VdLs.17] | 71.7 | 8.0E-159 |
| DIP_10001166 | NA | | |
| DIP_10001167 | hypothetical protein UCDDA912_g01384 [Diaporthe ampelina] | 30.8 | 1.0E-14 |
| DIP_10001168 | hypothetical protein MYCTH_2119925 [Myceliophthora thermophila ATCC 42464] | 34.6 | 3.0E-59 |
| DIP_10001169 | putative wd repeat-containing protein jip5 protein [Eutypa lata UCREL1] | 70.6 | 0.0 |
| DIP_10001170 | putative peroxisomal catalase protein [Togninia minima UCRPA7] | 79.1 | 0.0 |
| DIP_10001171 | hypothetical protein PFICI_15079 [Pestalotiopsis fici W106-1] | 68 | 0.0 |
| DIP_10001172 | NA | | |
| DIP_10001173 | NA | | |
| DIP_10001174 | hypothetical protein H072_174 [Dactylellina haptotyla CBS 200.50] | 49.7 | 2.0E-38 |
| DIP_10001175 | NA | | |
| DIP_10001176 | NA | | |
| DIP_10001177 | NA | | |
| DIP_10001178 | NA | | |
| DIP_10001179 | hypothitical protein [Pestalotiopsis fici W106-1] | 79.4 | 0.0 |
| DIP_10001180 | NA | | |
| DIP_10001181 | endosomal peripheral membrane protein, partial [Colletotrichum higginsianum] | 72 | 0.0 |
| DIP_10001182 | Ubiquitin-conjugating enzyme spm2 [Pestalotiopsis fici W106-1] | 87.9 | 3.0E-91 |
| DIP_10001183 | hypothetical protein PFICI_10652 [Pestalotiopsis fici W106-1] | 33.4 | 3.0E-43 |
| DIP_10001184 | hypothetical protein [Podospora anserina S mat+] | 31.1 | 4.0E-56 |
| DIP_10001185 | NA | | |
| DIP_10001186 | hypothetical protein UCREL1_6845 [Eutypa lata UCREL1] | 31.6 | 1.0E-18 |
| DIP_10001187 | NA | | |
| DIP_10001188 | putative ligninase h2 precursor protein [Eutypa lata UCREL1] | 58.1 | 3.0E-131 |
| DIP_10001189 | putative dienelactone hydrolase family protein [Eutypa lata UCREL1] | 57.8 | 2.0E-118 |
| DIP_10001190 | hypothetical protein CGLO_02026 [Colletotrichum gloeosporioides Cg-14] | 55.5 | 7.0E-132 |
| DIP_10001191 | hypothetical protein UCRNP2_6846 [Neofusicoccum parvum UCRNP2] | 41 | 6.0E-20 |
| DIP_10001192 | hypothetical protein OOU_Y34scaffold00881g1 [Magnaporthe oryzae Y34] | 36.5 | 6.0E-10 |
| DIP_10001193 | hypothetical protein GGTG_05049 [Gaeumannomyces graminis var. tritici R3-111a-1] | 33.3 | 3.0E-31 |
| DIP_10001194 | hypothetical protein EURHEDRAFT_530994 [Aspergillus ruber CBS 135680] | 43.3 | 5.0E-108 |
| DIP_10001195 | putative atp-dependent dna helicase recq protein [Eutypa lata UCREL1] | 60.2 | 0.0 |
| DIP_10001196 | putative porphobilinogen deaminase protein [Eutypa lata UCREL1] | 57.5 | 2.0E-64 |
| DIP_10001197 | putative integral membrane protein [Eutypa lata UCREL1] | 42.8 | 2.0E-138 |
| DIP_10001198 | putative cell division control protein [Eutypa lata UCREL1] | 63.8 | 0.0 |
| DIP_10001199 | putative phosphoglycerate mutase family protein [Eutypa lata UCREL1] | 56.9 | 2.0E-122 |
| DIP_10001200 | DyP [Metarhizium guizhouense ARSEF 977] | 35.7 | 4.0E-15 |
| DIP_10001201 | DSBA-like thioredoxin domain-containing protein [Colletotrichum graminicola M1.001] | 54.6 | 3.0E-84 |
| DIP_10001202 | putative beta- -glucanase protein [Eutypa lata UCREL1] | 68.3 | 0.0 |
| DIP_10001203 | putative serine threonine-protein kinase protein [Eutypa lata UCREL1] | 82.7 | 0.0 |
| DIP_10001204 | putative igr protein domain-containing protein [Eutypa lata UCREL1] | 55.5 | 3.0E-71 |
| DIP_10001205 | hypothetical protein FVEG_07844 [Fusarium verticillioides 7600] | 39.6 | 0.0 |
| DIP_10001206 | NA | | |
| DIP_10001207 | putative secretion pathway protein sls2 rcy1 protein [Eutypa lata UCREL1] | 74.8 | 0.0 |
| DIP_10001208 | hypothetical protein CCM_03890 [Cordyceps militaris CM01] | 34 | 3.0E-06 |
| DIP_10001209 | putative secretory pathway protein ysy6-like protein [Eutypa lata UCREL1] | 76.5 | 3.0E-33 |
| DIP_10001210 | hypothetical protein M436DRAFT_52589 [Aureobasidium namibiae CBS 147.97] | 37.4 | 2.0E-15 |
| DIP_10001211 | putative calcium influx-promoting protein ehs1 protein [Eutypa lata UCREL1] | 51.9 | 4.0E-173 |
| DIP_10001212 | NA | | |
| DIP_10001213 | putative ef-hand superfamily ca2+-modulated protein [Eutypa lata UCREL1] | 61.5 | 2.0E-73 |
| DIP_10001214 | putative translation initiation factor protein [Eutypa lata UCREL1] | 78.7 | 1.0E-147 |
| DIP_10001215 | hypothetical protein M419DRAFT_85741 [Trichoderma reesei RUT C-30] | 67.1 | 0.0 |
| DIP_10001216 | hypothetical protein PFICI_12085 [Pestalotiopsis fici W106-1] | 80.9 | 0.0 |
| DIP_10001217 | hypothetical protein PFICI_12084 [Pestalotiopsis fici W106-1] | 44.8 | 1.0E-85 |
| DIP_10001218 | S-adenosylmethionine-dependent methyltransferase-like protein [Chaetomium thermophilum var. thermophilum DSM 1495] | 47.8 | 8.0E-66 |
| DIP_10001219 | NA | | |
| DIP_10001220 | hypothetical protein CGGC5_3682 [Colletotrichum gloeosporioides Nara gc5] | 39.6 | 7.0E-41 |
| DIP_10001221 | hypothetical protein PFICI_12080 [Pestalotiopsis fici W106-1] | 50.4 | 2.0E-78 |
| DIP_10001222 | saccharopine dehydrogenase [Colletotrichum orbiculare MAFF 240422] | 79.9 | 0.0 |
| DIP_10001223 | hypothetical protein PFICI_12082 [Pestalotiopsis fici W106-1] | 49.5 | 2.0E-83 |
| DIP_10001224 | hypothetical protein UCREL1_11137 [Eutypa lata UCREL1] | 41.6 | 2.0E-07 |
| DIP_10001225 | hypothetical protein PFICI_12083 [Pestalotiopsis fici W106-1] | 77.6 | 2.0E-154 |
| DIP_10001226 | hypothetical protein PFICI_12078 [Pestalotiopsis fici W106-1] | 77.1 | 0.0 |
| DIP_10001227 | glycerophosphoryl diester phosphodiesterase [Beauveria bassiana ARSEF 2860] | 43.6 | 1.0E-106 |
| DIP_10001228 | hypothetical protein PFICI_12073 [Pestalotiopsis fici W106-1] | 48.2 | 1.0E-176 |
| DIP_10001229 | hypothetical protein V496_00682 [Pseudogymnoascus pannorum VKM F-4515 (FW-2607)] NADPH-dependent 1-acyldihydroxyacetone phosphate reductase [Aspergillus lentulus] | 34.3 | 6.0E-36 |
| DIP_10001230 | NA | | |
| DIP_10001231 | hypothetical protein PFICI_07768 [Pestalotiopsis fici W106-1] | 56.5 | 2.0E-106 |
| DIP_10001232 | NA | | |
| DIP_10001233 | hypothetical protein PFICI_08069 [Pestalotiopsis fici W106-1] | 63.1 | 4.0E-120 |
| DIP_10001234 | hypothetical protein [Neurospora crassa] | 42.4 | 5.0E-34 |
| DIP_10001235 | hypothetical protein PFICI_07751 [Pestalotiopsis fici W106-1] | 52.1 | 0.0 |
| DIP_10001236 | putative duf410 domain-containing protein [Eutypa lata UCREL1] | 70.9 | 1.0E-160 |
| DIP_10001237 | hypothetical protein FAVG1_05774 [Fusarium avenaceum] | 41.3 | 4.0E-29 |
| DIP_10001238 | putative protein rai1 protein [Eutypa lata UCREL1] | 78.5 | 0.0 |
| DIP_10001239 | putative alpha beta fold family protein [Eutypa lata UCREL1] | 69.5 | 0.0 |
| DIP_10001240 | putative 60s ribosomal protein l4-a protein [Eutypa lata UCREL1] | 70.4 | 6.0E-170 |
| DIP_10001241 | hypothetical protein PFICI_08067 [Pestalotiopsis fici W106-1] | 47.9 | 0.0 |
| DIP_10001242 | hypothetical protein PFICI_08066 [Pestalotiopsis fici W106-1] | 77.7 | 0.0 |
| DIP_10001243 | NA | | |
| DIP_10001244 | NA | | |
| DIP_10001245 | putative ulp1 protease family protein [Eutypa lata UCREL1] | 48 | 4.0E-88 |
| DIP_10001246 | hypothetical protein PFICI_08063 [Pestalotiopsis fici W106-1] | 68.7 | 2.0E-111 |
| DIP_10001247 | putative mitochondrial import receptor subunit tom-40 protein [Eutypa lata UCREL1] | 70.9 | 0.0 |
| DIP_10001248 | hypothetical protein PFICI_07731 [Pestalotiopsis fici W106-1] | 71.2 | 0.0 |
| DIP_10001249 | ubiquitin carboxyl-terminal hydrolase family protein [Colletotrichum orbiculare MAFF 240422] | 37.8 | 1.0E-160 |
| DIP_10001250 | hypothetical protein PFICI_07740 [Pestalotiopsis fici W106-1] | 53.4 | 1.0E-135 |
| DIP_10001251 | Hsp70 nucleotide exchange factor [Pestalotiopsis fici W106-1] | 56 | 6.0E-82 |
| DIP_10001252 | hypothetical protein SMAC_04416 [Sordaria macrospora k-hell] | 100 | 0.0 |
| DIP_10001253 | peptidyl-tRNA hydrolase domain-containing protein [Magnaporthe oryzae 70-15] | 69 | 3.0E-52 |
| DIP_10001254 | hypothetical protein PFICI_06250 [Pestalotiopsis fici W106-1] | 67 | 2.0E-44 |
| DIP_10001255 | WD repeat domain-containing protein [Colletotrichum fioriniae PJ7] | 47.9 | 0.0 |
| DIP_10001256 | putative cap binding protein [Eutypa lata UCREL1] | 78.2 | 0.0 |
| DIP_10001257 | Protein kinase gsk3 [Pestalotiopsis fici W106-1] | 92.2 | 0.0 |
| DIP_10001258 | NA | | |
| DIP_10001259 | hypothetical protein MGG_12124 [Magnaporthe oryzae 70-15] | 89.4 | 2.0E-24 |
| DIP_10001260 | hypothetical protein PFICI_08210 [Pestalotiopsis fici W106-1] | 80.2 | 3.0E-174 |
| DIP_10001261 | NA | | |
| DIP_10001262 | hypothetical protein FAVG1_05753 [Fusarium avenaceum] | 39.8 | 1.0E-96 |
| DIP_10001263 | ubiquitin related modifier 1 [Colletotrichum orbiculare MAFF 240422] | 80.2 | 8.0E-62 |
| DIP_10001264 | hypothetical protein GLRG_03067 [Colletotrichum graminicola M1.001] | 55.9 | 6.0E-129 |
| DIP_10001265 | hypothetical protein MGG_03971 [Magnaporthe oryzae 70-15] | 80.6 | 2.0E-95 |
| DIP_10001266 | hypothetical protein PFICI_07726 [Pestalotiopsis fici W106-1] | 61.9 | 0.0 |
| DIP_10001267 | NA | | |
| DIP_10001268 | putative sec8 exocyst complex component specific domain-containing protein [Eutypa lata UCREL1] | 78.2 | 0.0 |
| DIP_10001269 | putative phospholipase carboxylesterase superfamily protein [Eutypa lata UCREL1] | 71.3 | 5.0E-128 |
| DIP_10001270 | putative mfs monosaccharide protein [Eutypa lata UCREL1] | 77.6 | 0.0 |
| DIP_10001271 | variant SH3 domain-containing protein [Colletotrichum gloeosporioides Cg-14] | 80.2 | 0.0 |
| DIP_10001272 | hypothetical protein V491_03052, partial [Pseudogymnoascus pannorum VKM F-3775] | 61 | 3.0E-43 |
| DIP_10001273 | putative met3-like protein [Eutypa lata UCREL1] | 70.8 | 0.0 |
| DIP_10001274 | NA | | |
| DIP_10001275 | hypothetical protein PFICI_08095 [Pestalotiopsis fici W106-1] | 81.3 | 0.0 |
| DIP_10001276 | hypothetical protein PFICI_08089 [Pestalotiopsis fici W106-1] | 50.1 | 0.0 |
| DIP_10001277 | hypothetical protein UCREL1_9500 [Eutypa lata UCREL1] | 36.9 | 1.0E-84 |
| DIP_10001278 | NA | | |
| DIP_10001279 | tuberin [Colletotrichum higginsianum] | 63.3 | 0.0 |
| DIP_10001280 | putative gtpase activating protein [Togninia minima UCRPA7] | 47.3 | 0.0 |
| DIP_10001281 | putative chromosome segregation protein [Eutypa lata UCREL1] | 70.4 | 0.0 |
| DIP_10001282 | hypothetical protein PFICI_08094 [Pestalotiopsis fici W106-1] | 46.7 | 1.0E-139 |
| DIP_10001283 | putative ccr4-not complex subunit caf16 protein [Eutypa lata UCREL1] | 79.4 | 1.0E-161 |
| DIP_10001284 | hypothetical protein NCU01391 [Neurospora crassa OR74A] | 53 | 6.0E-12 |
| DIP_10001285 | putative cell cycle control protein cwf16 protein [Togninia minima UCRPA7] | 64.2 | 4.0E-115 |
| DIP_10001286 | putative spc97 spc98 family protein [Eutypa lata UCREL1] | 46.7 | 3.0E-88 |
| DIP_10001287 | putative glycolipid 2-alpha-mannosyltransferase protein [Eutypa lata UCREL1] | 80.4 | 0.0 |
| DIP_10001288 | hypothetical protein PFICI_08099 [Pestalotiopsis fici W106-1] | 49.5 | 2.0E-129 |
| DIP_10001289 | hypothetical protein GLRG_03070 [Colletotrichum graminicola M1.001] | 47.2 | 0.0 |
| DIP_10001290 | NA | | |
| DIP_10001291 | NA | | |
| DIP_10001292 | hypothetical protein THITE_2107667 [Thielavia terrestris NRRL 8126] | 49.5 | 0.0 |
| DIP_10001293 | putative urease accessory protein [Eutypa lata UCREL1] | 50.6 | 1.0E-170 |
| DIP_10001294 | hypothetical protein SMAC_07555 [Sordaria macrospora k-hell] | 44.5 | 1.0E-80 |
| DIP_10001295 | putative pre-mrna splicing factor cwc2 protein [Eutypa lata UCREL1] | 82 | 0.0 |
| DIP_10001296 | putative acyl- dehydrogenase family member 11 protein [Eutypa lata UCREL1] | 86.7 | 0.0 |
| DIP_10001297 | hypothetical protein NECHADRAFT_101680 [Nectria haematococca mpVI 77-13-4] | 63.8 | 3.0E-37 |
| DIP_10001298 | GTP-binding protein [Colletotrichum gloeosporioides Nara gc5] | 89.5 | 0.0 |
| DIP_10001299 | hypothetical protein PFICI_07419 [Pestalotiopsis fici W106-1] | 74.4 | 0.0 |
| DIP_10001300 | hypothetical protein NECHADRAFT_106298 [Nectria haematococca mpVI 77-13-4] | 33.5 | 3.0E-16 |
| DIP_10001301 | putative multidrug resistance protein fnx1 protein [Eutypa lata UCREL1] | 74.8 | 0.0 |
| DIP_10001302 | putative chaperone-binding protein [Eutypa lata UCREL1] | 56.4 | 2.0E-116 |
| DIP_10001303 | hypothetical protein PFICI_00640 [Pestalotiopsis fici W106-1] | 36.2 | 3.0E-59 |
| DIP_10001304 | glycoside hydrolase family 63 protein [Thielavia terrestris NRRL 8126] | 51.9 | 0.0 |
| DIP_10001305 | putative dnase1 protein [Eutypa lata UCREL1] | 57.7 | 2.0E-65 |
| DIP_10001306 | oxidoreductase, FAD-binding [Aspergillus kawachii IFO 4308] | 34.8 | 1.0E-80 |
| DIP_10001307 | hypothetical protein NECHADRAFT_84947 [Nectria haematococca mpVI 77-13-4] | 42.9 | 0.0 |
| DIP_10001308 | hypothetical protein PFICI_08204 [Pestalotiopsis fici W106-1] | 44.6 | 3.0E-92 |
| DIP_10001309 | NA | | |
| DIP_10001310 | hypothetical protein PFICI_08065 [Pestalotiopsis fici W106-1] | 63 | 0.0 |
| DIP_10001311 | putative dna replication licensing factor mcm5 protein [Eutypa lata UCREL1] | 90 | 0.0 |
| DIP_10001312 | hypothetical protein PFICI_08190 [Pestalotiopsis fici W106-1] | 48.2 | 1.0E-97 |
| DIP_10001313 | tfiiic transcription initiation factor complex subunits tfc3 [Colletotrichum gloeosporioides Nara gc5] | 37.1 | 3.0E-96 |
| DIP_10001314 | putative tfiiic transcription initiation factor complex subunits tfc3 protein [Eutypa lata UCREL1] | 42.8 | 6.0E-139 |
| DIP_10001315 | NA | | |
| DIP_10001316 | NA | | |
| DIP_10001317 | hypothetical protein FOMG_03160 [Fusarium oxysporum f. sp. melonis 26406] | 56.3 | 1.0E-175 |
| DIP_10001318 | hypothetical protein PFICI_08192 [Pestalotiopsis fici W106-1] | 53.7 | 2.0E-120 |
| DIP_10001319 | putative mitochondrial chaperone bcs1 protein [Eutypa lata UCREL1] | 57.9 | 0.0 |
| DIP_10001320 | putative mechanosensitive ion channel family protein [Eutypa lata UCREL1] | 67.1 | 0.0 |
| DIP_10001321 | SAC3 family protein 1 [Ceratocystis platani] | 30.9 | 2.0E-85 |
| DIP_10001322 | hypothetical protein UCREL1_8658 [Eutypa lata UCREL1] | 46.6 | 4.0E-109 |
| DIP_10001323 | predicted protein [Nectria haematococca mpVI 77-13-4] | 77.8 | 0.0 |
| DIP_10001324 | hypothetical protein DOTSEDRAFT_159290 [Dothistroma septosporum NZE10] | 32 | 6.0E-20 |
| DIP_10001325 | hypothetical protein PFICI_07625 [Pestalotiopsis fici W106-1] | 33.7 | 5.0E-27 |
| DIP_10001326 | hypothetical protein PFICI_00383 [Pestalotiopsis fici W106-1] | 66.5 | 0.0 |
| DIP_10001327 | putative meiotically up-regulated gene 190 protein [Eutypa lata UCREL1] | 59.9 | 0.0 |
| DIP_10001328 | hypothetical protein FGSG_04160 [Fusarium graminearum PH-1] | 68.5 | 7.0E-97 |
| DIP_10001329 | putative cyanamide hydratase protein [Eutypa lata UCREL1] | 61.6 | 1.0E-110 |
| DIP_10001330 | AP-2 complex subunit mu [Pestalotiopsis fici W106-1] | 95.7 | 0.0 |
| DIP_10001331 | hypothetical protein V490_02903 [Pseudogymnoascus pannorum VKM F-3557] | 34.2 | 2.0E-06 |
| DIP_10001332 | hypothetical protein Cob_10335 [Colletotrichum orbiculare MAFF 240422] | 57.3 | 0.0 |
| DIP_10001333 | hypothetical protein PFICI_09480 [Pestalotiopsis fici W106-1] | 40.3 | 3.0E-136 |
| DIP_10001334 | putative bar domain-containing protein [Eutypa lata UCREL1] | 64.1 | 0.0 |
| DIP_10001335 | hypothetical protein MYCTH_2315449 [Myceliophthora thermophila ATCC 42464] | 69.2 | 0.0 |
| DIP_10001336 | putative ring finger ubiquitin ligase protein [Eutypa lata UCREL1] | 58.1 | 0.0 |
| DIP_10001337 | hypothetical protein GGTG_00502 [Gaeumannomyces graminis var. tritici R3-111a-1] | 58.4 | 2.0E-19 |
| DIP_10001338 | Protein disulfide-isomerase erp38 [Pestalotiopsis fici W106-1] | 69.8 | 0.0 |
| DIP_10001339 | 30S ribosomal protein S14p/S29e [Gaeumannomyces graminis var. tritici R3-111a-1] | 89.3 | 5.0E-32 |
| DIP_10001340 | putative elongation factor tu protein [Eutypa lata UCREL1] | 84.3 | 0.0 |
| DIP_10001341 | hypothetical protein PFICI_07867 [Pestalotiopsis fici W106-1] | 68.7 | 0.0 |
| DIP_10001342 | Cyanate hydratase [Pestalotiopsis fici W106-1] | 80 | 3.0E-94 |
| DIP_10001343 | NA | | |
| DIP_10001344 | NA | | |
| DIP_10001345 | putative cullin-associated nedd8-dissociated protein 2 protein [Eutypa lata UCREL1] | 67.6 | 0.0 |
| DIP_10001346 | hypothetical protein UCREL1_11122 [Eutypa lata UCREL1] | 41 | 3.0E-30 |
| DIP_10001347 | NA | | |
| DIP_10001348 | transporter [Pestalotiopsis fici W106-1] | 75.5 | 0.0 |
| DIP_10001349 | NA | | |
| DIP_10001350 | putative signal recognition particle 54 kda protein [Eutypa lata UCREL1] | 91.9 | 0.0 |
| DIP_10001351 | putative alpha-glucan synthase [Diaporthe ampelina] | 54.1 | 0.0 |
| DIP_10001352 | High-affinity methionine permease [Talaromyces islandicus] | 75.4 | 0.0 |
| DIP_10001353 | hypothetical protein VDAG_05066 [Verticillium dahliae VdLs.17] | 32.2 | 5.0E-21 |
| DIP_10001354 | putative polyamine transporter tpo5 protein [Eutypa lata UCREL1] | 69.6 | 0.0 |
| DIP_10001355 | hypothetical protein PFICI_04173 [Pestalotiopsis fici W106-1] | 59.4 | 2.0E-102 |
| DIP_10001356 | putative methyltransferase protein [Eutypa lata UCREL1] | 44.1 | 2.0E-85 |
| DIP_10001357 | NA | | |
| DIP_10001358 | hypothetical protein OIDMADRAFT_141746 [Oidiodendron maius Zn] | 79.5 | 1.0E-79 |
| DIP_10001359 | putative cupin domain-containing protein [Eutypa lata UCREL1] | 76.5 | 7.0E-73 |
| DIP_10001360 | hypothetical protein PFICI_09610 [Pestalotiopsis fici W106-1] | 56.9 | 1.0E-124 |
| DIP_10001361 | homeobox domain-containing protein [Colletotrichum higginsianum] | 66.4 | 0.0 |
| DIP_10001362 | putative p-type atpase protein [Eutypa lata UCREL1] | 61.5 | 0.0 |
| DIP_10001363 | Alcohol oxidase [Pestalotiopsis fici W106-1] | 87.8 | 0.0 |
| DIP_10001364 | NA | | |
| DIP_10001365 | NA | | |
| DIP_10001366 | NA | | |
| DIP_10001367 | NA | | |
| DIP_10001368 | putative MFS transporter [fungal sp. NRRL 50135] | 66.6 | 0.0 |
| DIP_10001369 | hypothetical protein AUEXF2481DRAFT_1954 [Aureobasidium subglaciale EXF-2481] | 57.2 | 0.0 |
| DIP_10001370 | hypothetical protein Z518_10159 [Rhinocladiella mackenziei CBS 650.93] | 71.3 | 0.0 |
| DIP_10001371 | putative 4-hydroxythreonine-4-phosphate dehydrogenase protein [Eutypa lata UCREL1] | 75.5 | 0.0 |
| DIP_10001372 | putative peptide transporter ptr2 protein [Eutypa lata UCREL1] | 73.5 | 0.0 |
| DIP_10001373 | putative rnase iii domain-containing protein [Eutypa lata UCREL1] | 51.5 | 1.0E-68 |
| DIP_10001374 | hypothetical protein PFICI_10719 [Pestalotiopsis fici W106-1] | 45.9 | 9.0E-158 |
| DIP_10001375 | putative d-isomer specific 2-hydroxyacid dehydrogenase protein [Eutypa lata UCREL1] | 57.5 | 5.0E-169 |
| DIP_10001376 | hypothetical protein MYCTH_2301726 [Myceliophthora thermophila ATCC 42464] | 34.7 | 4.0E-14 |
| DIP_10001377 | hypothetical protein PFICI_07330 [Pestalotiopsis fici W106-1] | 44.2 | 2.0E-71 |
| DIP_10001378 | hypothetical protein TCE0_013r00864 [Talaromyces cellulolyticus] | 36.2 | 4.0E-12 |
| DIP_10001379 | NA | | |
| DIP_10001380 | hypothetical protein CGLO_07477 [Colletotrichum gloeosporioides Cg-14] | 53.7 | 0.0 |
| DIP_10001381 | hypothetical protein S40293_04651 [Stachybotrys chartarum IBT 40293] | 67.3 | 0.0 |
| DIP_10001382 | hypothetical protein GLRG_03300 [Colletotrichum graminicola M1.001] | 88.9 | 1.0E-23 |
| DIP_10001383 | hypothetical protein NECHADRAFT_63362 [Nectria haematococca mpVI 77-13-4] | 63.7 | 3.0E-153 |
| DIP_10001384 | NA | | |
| DIP_10001385 | lysR family regulatory protein [Colletotrichum gloeosporioides Nara gc5] | 38.6 | 8.0E-115 |
| DIP_10001386 | hypothetical protein PFICI_10928 [Pestalotiopsis fici W106-1] | 59.3 | 0.0 |
| DIP_10001387 | putative rrna processing protein [Eutypa lata UCREL1] | 47.6 | 6.0E-75 |
| DIP_10001388 | putative phosphoprotein phosphatase protein [Eutypa lata UCREL1] | 62.6 | 1.0E-95 |
| DIP_10001389 | putative d-lactate dehydrogenase protein [Eutypa lata UCREL1] | 40.9 | 8.0E-112 |
| DIP_10001390 | NA | | |
| DIP_10001391 | putative nadp-dependent malic enzyme protein [Eutypa lata UCREL1] | 79.3 | 0.0 |
| DIP_10001392 | hypothetical protein PFICI_09536 [Pestalotiopsis fici W106-1] | 44 | 3.0E-100 |
| DIP_10001393 | NA | | |
| DIP_10001394 | Cutinase transcription factor 1 beta [Pestalotiopsis fici W106-1] | 52.4 | 0.0 |
| DIP_10001395 | NA | | |
| DIP_10001396 | hypothetical protein UCREL1_8592 [Eutypa lata UCREL1] | 50.5 | 1.0E-131 |
| DIP_10001397 | hypothetical protein PFICI_10915 [Pestalotiopsis fici W106-1] | 69.9 | 0.0 |
| DIP_10001398 | hypothetical protein CH063_08636 [Colletotrichum higginsianum] | 64.3 | 1.0E-126 |
| DIP_10001399 | NA | | |
| DIP_10001400 | short chain dehydrogenase/reductase, putative [Talaromyces stipitatus ATCC 10500] | 42.1 | 8.0E-58 |
| DIP_10001401 | predicted protein [Nectria haematococca mpVI 77-13-4] | 92.5 | 3.0E-115 |
| DIP_10001402 | putative epl1-like protein [Eutypa lata UCREL1] | 68.3 | 0.0 |
| DIP_10001403 | O-sialoglycoprotein endopeptidase [Verticillium dahliae VdLs.17] | 57.8 | 5.0E-174 |
| DIP_10001404 | putative carbohydrate-binding module family 48 protein [Eutypa lata UCREL1] | 63.4 | 0.0 |
| DIP_10001405 | NA | | |
| DIP_10001406 | NA | | |
| DIP_10001407 | putative trafficking pga2 protein [Eutypa lata UCREL1] | 63.8 | 1.0E-61 |
| DIP_10001408 | unnamed protein product [Sordaria macrospora k-hell] | 74.3 | 2.0E-56 |
| DIP_10001409 | hypothetical protein UCREL1_170 [Eutypa lata UCREL1] | 61.9 | 1.0E-107 |
| DIP_10001410 | hypothetical protein OIDMADRAFT_51371 [Oidiodendron maius Zn] | 34.5 | 2.0E-31 |
| DIP_10001411 | hypothetical protein PFICI_10867 [Pestalotiopsis fici W106-1] | 66.3 | 0.0 |
| DIP_10001412 | hypothetical protein PFICI_10868 [Pestalotiopsis fici W106-1] | 38.1 | 4.0E-25 |
| DIP_10001413 | 60S ribosomal subunit assembly/export protein LOC1 [Pestalotiopsis fici W106-1] | 67.3 | 1.0E-70 |
| DIP_10001414 | hypothetical protein THITE_2106862 [Thielavia terrestris NRRL 8126] | 88.6 | 4.0E-117 |
| DIP_10001415 | NA | | |
| DIP_10001416 | hypothetical protein PFICI_10934 [Pestalotiopsis fici W106-1] | 60.3 | 1.0E-147 |
| DIP_10001417 | putative 40s ribosomal protein s9 protein [Eutypa lata UCREL1] | 61.3 | 1.0E-109 |
| DIP_10001418 | hypothetical protein V491_04953, partial [Pseudogymnoascus pannorum VKM F-3775] | 60.2 | 7.0E-30 |
| DIP_10001419 | putative elongation factor 1 gamma domain-containing protein [Diaporthe ampelina] | 66 | 3.0E-136 |
| DIP_10001420 | hypothetical protein PV06_05052 [Exophiala oligosperma] | 79.7 | 0.0 |
| DIP_10001421 | NA | | |
| DIP_10001422 | hypothetical protein PV09_04978 [Verruconis gallopava] | 49.2 | 0.0 |
| DIP_10001423 | hypothetical protein ANO11243_042620 [fungal sp. No.11243] | 67.5 | 0.0 |
| DIP_10001424 | hypothetical protein PV05_11763 [Exophiala xenobiotica] | 57 | 3.0E-108 |
| DIP_10001425 | hypothetical protein PV06_05050 [Exophiala oligosperma] | 41.3 | 5.0E-141 |
| DIP_10001426 | hypothetical protein PV06_05049 [Exophiala oligosperma] | 48.3 | 0.0 |
| DIP_10001427 | hypothetical protein PFICI_10724 [Pestalotiopsis fici W106-1] | 55.6 | 0.0 |
| DIP_10001428 | putative 20s-pre-rrna d-site endonuclease nob1 protein [Eutypa lata UCREL1] | 55.5 | 0.0 |
| DIP_10001429 | putative h aca ribonucleoprotein complex subunit 3 protein [Eutypa lata UCREL1] | 85.1 | 3.0E-35 |
| DIP_10001430 | hypothetical protein FOXB_05504 [Fusarium oxysporum Fo5176] NAD(P)-binding protein [illegible] | 58.5 | 2.0E-178 |
| DIP_10001431 | hypothetical protein FOXB_05503 [Fusarium oxysporum Fo5176] aldehyde dehydrogenase [illegible] | 73.4 | 0.0 |
| DIP_10001432 | hypothetical protein FOVG_07054 [Fusarium oxysporum f. sp. pisi HDV247] | 61.2 | 0.0 |
| DIP_10001433 | hypothetical protein FOVG_07055 [Fusarium oxysporum f. sp. pisi HDV247] | 75.5 | 4.0E-137 |
| DIP_10001434 | polyketide synthase [Fusarium fujikuroi] Sequence ID: [illegible] | 46.6 | 0.0 |
| DIP_10001435 | hypothetical protein UCREL1_3716 [Eutypa lata UCREL1] | 55.5 | 7.0E-153 |
| DIP_10001436 | related to Tri201-trichothecene 3-O-acetyltransferase [Fusarium fujikuroi IMI 58289] | 58.6 | 0.0 |
| DIP_10001437 | hypothetical protein PFICI_09644 [Pestalotiopsis fici W106-1] | 37 | 2.0E-63 |
| DIP_10001438 | Cytosolic Fe-S cluster assembly factor CFD1 [Pestalotiopsis fici W106-1] | 82.7 | 0.0 |
| DIP_10001439 | putative s-adenosylmethionine decarboxylase proenzyme protein [Eutypa lata UCREL1] | 70.2 | 0.0 |
| DIP_10001440 | predicted protein [Trichoderma reesei QM6a] | 45 | 6.0E-65 |
| DIP_10001441 | NA | | |
| DIP_10001442 | NA | | |
| DIP_10001443 | NA | | |
| DIP_10001444 | Na(+)/H(+) antiporter [Gaeumannomyces graminis var. tritici R3-111a-1] | 56.7 | 0.0 |
| DIP_10001445 | NA | | |
| DIP_10001446 | hypothetical protein CSUB01_04821 [Colletotrichum sublineola] | 49.7 | 1.0E-84 |
| DIP_10001447 | hypothetical protein PFICI_04146 [Pestalotiopsis fici W106-1] | 57.4 | 4.0E-128 |
| DIP_10001448 | alpha/beta-Hydrolase [Glarea lozoyensis ATCC 20868] | 39.5 | 4.0E-66 |
| DIP_10001449 | putative heterokaryon incompatibility protein [Neofusicoccum parvum UCRNP2] | 34.4 | 2.0E-54 |
| DIP_10001450 | hypothetical protein PFICI_09516 [Pestalotiopsis fici W106-1] | 49.1 | 2.0E-163 |
| DIP_10001451 | putative capsule polysaccharide biosynthesis protein [Eutypa lata UCREL1] | 51.1 | 3.0E-143 |
| DIP_10001452 | GTP cyclohydrolase 1 [Pestalotiopsis fici W106-1] | 77.4 | 9.0E-170 |
| DIP_10001453 | putative transcription initiation factor tfiid subunit 11 protein [Eutypa lata UCREL1] | 65.9 | 5.0E-149 |
| DIP_10001454 | short-chain dehydrogenase [Sinorhizobium medicae] | 45.2 | 3.0E-61 |
| DIP_10001455 | NA | | |
| DIP_10001456 | putative 3-hydroxyphenylacetate 6 hydroxylase protein [Eutypa lata UCREL1] | 63.2 | 0.0 |
| DIP_10001457 | NA | | |
| DIP_10001458 | Major facilitator superfamily [Macrophomina phaseolina MS6] | 46.7 | 1.0E-140 |
| DIP_10001459 | putative bzip transcription factor protein [Eutypa lata UCREL1] | 64.9 | 7.0E-139 |
| DIP_10001460 | putative oligomeric golgi complex subunit 2 protein [Eutypa lata UCREL1] | 59.3 | 6.0E-87 |
| DIP_10001461 | hypothetical protein PFICI_11612 [Pestalotiopsis fici W106-1] | 63.9 | 1.0E-140 |

| | | | |
|---|---|---|---|
| DIP_10001462 | putative glutamyl-trna synthetase protein [Eutypa lata UCREL1] | 86.8 | 0.0 |
| DIP_10001463 | hypothetical protein AOR_1_322054 [Aspergillus oryzae RIB40] | 35.9 | 8.0E-09 |
| DIP_10001464 | Chitinase-like protein [Acremonium chrysogenum ATCC 11550] | 55.9 | 1.0E-127 |
| DIP_10001465 | hypothetical protein COCMIDRAFT_98269 [Bipolaris oryzae ATCC 44560] | 63.5 | 0.0 |
| DIP_10001466 | hypothetical protein MYCGRDRAFT_51574 [Byssochlamys spectabilis No. 5] | 70.1 | 0.0 |
| DIP_10001467 | glycoside hydrolase family 16 protein [Sclerotinia borealis F-4157] | 42.3 | 5.0E-95 |
| DIP_10001468 | hypothetical protein TRIVIDRAFT_55144 [Trichoderma virens Gv29-8] | 76.9 | 8.0E-80 |
| DIP_10001469 | hypothetical protein TRIVIDRAFT_27992 [Trichoderma virens Gv29-8] | 41.3 | 1.0E-176 |
| DIP_10001470 | hypothetical protein FOVG_00032 [Fusarium oxysporum f. sp. pisi HDV247] | 36.5 | 7.0E-105 |
| DIP_10001471 | hypothetical protein CFIO01_13418 [Colletotrichum fioriniae PJ7] | 54.2 | 0.0 |
| DIP_10001472 | putative ceramide glucosyltransferase protein [Eutypa lata UCREL1] | 60.8 | 0.0 |
| DIP_10001473 | putative tRNA-splicing endonuclease subunit sen-2 [Beauveria bassiana D1-5] | 42.4 | 2.0E-49 |
| DIP_10001474 | putative trna-splicing endonuclease subunit sen2 [Diplodia seriata] | 58.4 | 2.0E-52 |
| DIP_10001475 | putative major facilitator superfamily transporter protein [Eutypa lata UCREL1] | 59.7 | 0.0 |
| DIP_10001476 | putative clathrin assembly protein AP47 [Neurospora tetrasperma FGSC 2509] | 94.8 | 0.0 |
| DIP_10001477 | hypothetical protein CSUB01_04772 [Colletotrichum sublineola] | 46.7 | 2.0E-137 |
| DIP_10001478 | NA | | |
| DIP_10001479 | putative gdsl-like lipase acylhydrolase protein [Eutypa lata UCREL1] | 58.2 | 3.0E-80 |
| DIP_10001480 | hypothetical protein PP112 [Penicillium paxilli] | 49.7 | 1.0E-96 |
| DIP_10001481 | putative gtp-binding protein gtr1 protein [Eutypa lata UCREL1] | 92 | 0.0 |
| DIP_10001482 | putative secreted protein [Phaeomoniella chlamydospora] | 65 | 2.0E-98 |
| DIP_10001483 | putative ubx domain-containing protein [Eutypa lata UCREL1] | 68.5 | 0.0 |
| DIP_10001484 | putative mitochondrial import protein 1 protein [Eutypa lata UCREL1] | 79 | 2.0E-61 |
| DIP_10001485 | putative mitochondrial export translocase oxa1 protein [Eutypa lata UCREL1] | 48.9 | 3.0E-104 |
| DIP_10001486 | duf1688 domain-containing protein [Colletotrichum orbiculare MAFF 240422] | 69.5 | 0.0 |
| DIP_10001487 | fungal specific transcription factor domain-containing protein [Colletotrichum gloeosporioides Cg-14] | 49.6 | 1.0E-157 |
| DIP_10001488 | hypothetical protein CGLO_01171 [Colletotrichum gloeosporioides Cg-14] | 79.9 | 0.0 |
| DIP_10001489 | putative aminotransferase [Colletotrichum sublineola] | 70.6 | 2.0E-161 |
| DIP_10001490 | hypothetical protein PFICI_06341 [Pestalotiopsis fici W106-1] | 75.1 | 0.0 |
| DIP_10001491 | NA | | |
| DIP_10001492 | NA | | |
| DIP_10001493 | hypothetical protein BGT96224_2331B [Blumeria graminis f. sp. tritici 96224] | 74.6 | 5.0E-27 |
| DIP_10001494 | hypothetical protein CSUB01_07754 [Colletotrichum sublineola] | 39 | 8.0E-135 |
| DIP_10001495 | conserved hypothetical protein [Verticillium alfalfae VaMs.102] | 31 | 1.0E-60 |
| DIP_10001496 | hypothetical protein PFICI_00230 [Pestalotiopsis fici W106-1] | 60.2 | 1.0E-162 |
| DIP_10001497 | hypothetical protein PFICI_00231 [Pestalotiopsis fici W106-1] | 59.3 | 8.0E-157 |
| DIP_10001498 | putative u4 u6 small nuclear ribonucleoprotein prp4 protein [Eutypa lata UCREL1] | 72.8 | 0.0 |
| DIP_10001499 | NA | | |
| DIP_10001500 | Carbohydrate-binding domain family 9-like protein [Ophiocordyceps sinensis CO18] | 45.6 | 1.0E-107 |
| DIP_10001501 | putative serine threonine-protein kinase psk1 protein [Eutypa lata UCREL1] | 81.2 | 0.0 |
| DIP_10001502 | SH3 domain-containing protein [Magnaporthe oryzae P131] | 55.5 | 0.0 |
| DIP_10001503 | putative duf408 domain-containing protein [Eutypa lata UCREL1] | 40.5 | 1.0E-67 |
| DIP_10001504 | putative 40s ribosomal protein s17 protein [Eutypa lata UCREL1] | 95.8 | 5.0E-97 |
| DIP_10001505 | hypothetical protein UCREL1_3229 [Eutypa lata UCREL1] | 41.6 | 1.0E-117 |
| DIP_10001506 | hypothetical protein PFICI_05197 [Pestalotiopsis fici W106-1] | 70.5 | 2.0E-126 |
| DIP_10001507 | queuine tRNA-ribosyltransferase-like protein [Chaetomium thermophilum var. thermophilum DSM 1495] | 70.4 | 0.0 |
| DIP_10001508 | hypothetical protein CH063_02192 [Colletotrichum higginsianum] | 58 | 2.0E-30 |
| DIP_10001509 | hypothetical protein PFICI_05607 [Pestalotiopsis fici W106-1] | 42.5 | 3.0E-69 |
| DIP_10001510 | putative mediator of rna polymerase ii transcription subunit 21 protein [Eutypa lata UCREL1] | 73 | 1.0E-66 |
| DIP_10001511 | hypothetical protein C24B11.05 [Pestalotiopsis fici W106-1] | 77.3 | 6.0E-124 |
| DIP_10001512 | hypothetical protein GGTG_02931 [Gaeumannomyces graminis var. tritici R3-111a-1] | 37.4 | 1.0E-40 |
| DIP_10001513 | hypothetical protein PFICI_05611 [Pestalotiopsis fici W106-1] | 76.4 | 0.0 |
| DIP_10001514 | putative polyamine acetyltransferase protein [Eutypa lata UCREL1] | 60.4 | 6.0E-110 |
| DIP_10001515 | putative orotate phosphoribosyltransferase protein [Eutypa lata UCREL1] | 66.5 | 9.0E-105 |
| DIP_10001516 | putative nucleolar gtp-binding protein [Eutypa lata UCREL1] | 86.8 | 0.0 |
| DIP_10001517 | NA | | |
| DIP_10001518 | hypothetical protein UCDDS831_g02070 [Diplodia seriata] | 31.7 | 1.0E-28 |
| DIP_10001519 | hypothetical protein PFICI_05631 [Pestalotiopsis fici W106-1] | 65.6 | 4.0E-27 |
| DIP_10001520 | hypothetical protein PFICI_06816 [Pestalotiopsis fici W106-1] | 40.1 | 1.0E-77 |
| DIP_10001521 | hypothetical protein NECHADRAFT_84500 [Nectria haematococca mpVI 77-13-4] | 31.9 | 5.0E-13 |
| DIP_10001522 | kelch repeat protein [Verticillium alfalfae VaMs.102] | 58.2 | 9.0E-138 |
| DIP_10001523 | NA | | |
| DIP_10001524 | NA | | |
| DIP_10001525 | putative duf1014 domain protein [Eutypa lata UCREL1] | 68.2 | 5.0E-90 |
| DIP_10001526 | hypothetical protein PFICI_04614 [Pestalotiopsis fici W106-1] | 66.7 | 1.0E-147 |
| DIP_10001527 | NA | | |
| DIP_10001528 | hypothetical protein SMAC_01807 [Sordaria macrospora k-hell] | 57.1 | 7.0E-124 |
| DIP_10001529 | hypothetical protein THITE_2115312 [Thielavia terrestris NRRL 8126] | 60.5 | 0.0 |
| DIP_10001530 | NA | | |
| DIP_10001531 | hypothetical protein PFICI_04609 [Pestalotiopsis fici W106-1] | 65.2 | 2.0E-49 |
| DIP_10001532 | putative major facilitator superfamily transporter protein [Eutypa lata UCREL1] | 76.4 | 0.0 |
| DIP_10001533 | hypothetical protein HMPREF1541_09109 [Cyphellophora europaea CBS 101466] | 52.2 | 8.0E-99 |
| DIP_10001534 | hypothetical protein UCREL1_6589 [Eutypa lata UCREL1] | 30.4 | 4.0E-16 |
| DIP_10001535 | Centromere protein V-like protein [Acremonium chrysogenum ATCC 11550] | 37.7 | 9.0E-58 |
| DIP_10001536 | hypothetical protein PFICI_04601 [Pestalotiopsis fici W106-1] | 67.5 | 0.0 |
| DIP_10001537 | hypothetical protein PFICI_04602 [Pestalotiopsis fici W106-1] | 65.3 | 3.0E-170 |
| DIP_10001538 | hypothetical protein PFICI_04670 [Pestalotiopsis fici W106-1] | 58.2 | 0.0 |
| DIP_10001539 | hypothetical protein UCREL1_6149 [Eutypa lata UCREL1] | 33.5 | 7.0E-19 |
| DIP_10001540 | putative cue domain-containing protein [Eutypa lata UCREL1] | 63.2 | 2.0E-72 |
| DIP_10001541 | hypothetical protein S7711_06500 [Stachybotrys chartarum IBT 7711] | 52.3 | 9.0E-128 |
| DIP_10001542 | hypothetical protein S7711_06499 [Stachybotrys chartarum IBT 7711] | 48.4 | 0.0 |
| DIP_10001543 | putative eukaryotic translation initiation factor 5b protein [Eutypa lata UCREL1] | 75.8 | 0.0 |
| DIP_10001544 | putative thermotolerance protein [Eutypa lata UCREL1] | 38.1 | 3.0E-49 |
| DIP_10001545 | putative gtpase-activating protein gyp10 protein [Eutypa lata UCREL1] | 57.8 | 1.0E-143 |
| DIP_10001546 | hypothetical protein PFICI_04412 [Pestalotiopsis fici W106-1] | 71.7 | 0.0 |
| DIP_10001547 | eukaryotic translation initiation factor 5A-2 [Exophiala spinifera] | 89.4 | 2.0E-106 |
| DIP_10001548 | putative rna polymerase i specific transcription initiation factor protein [Eutypa lata UCREL1] | 45.5 | 4.0E-150 |
| DIP_10001549 | probable ubiquitin fusion protein (ubiquitin / ribosomal protein) [Claviceps purpurea 20.1] | 93.1 | 9.0E-43 |
| DIP_10001550 | hypothetical protein ACRE_046540 [Acremonium chrysogenum ATCC 11550] | 77.6 | 2.0E-153 |
| DIP_10001551 | hypothetical protein PFICI_04406 [Pestalotiopsis fici W106-1] | 54 | 3.0E-49 |
| DIP_10001552 | putative cytochrome p450 61 protein [Eutypa lata UCREL1] | 79.6 | 0.0 |
| DIP_10001553 | NA | | |
| DIP_10001554 | Glutathione S-transferase omega-like 2 [Pestalotiopsis fici W106-1] | 78.8 | 0.0 |
| DIP_10001555 | Ubiquitin-activating enzyme [Pestalotiopsis fici W106-1] | 87.2 | 0.0 |
| DIP_10001556 | putative ubiquitin-activating enzyme e1 1 protein [Eutypa lata UCREL1] | 86.4 | 0.0 |
| DIP_10001557 | hypothetical protein FVEG_06203 [Fusarium verticillioides 7600] | 30.4 | 5.0E-53 |
| DIP_10001558 | hypothetical protein PFICI_04389 [Pestalotiopsis fici W106-1] | 48.5 | 0.0 |
| DIP_10001559 | hypothetical protein PFICI_04399 [Pestalotiopsis fici W106-1] | 49.1 | 2.0E-65 |
| DIP_10001560 | putative Gamma-tubulin complex component 6 [Glarea lozoyensis 74030] | 34.7 | 5.0E-104 |
| DIP_10001561 | putative dash complex subunit ask1 protein [Eutypa lata UCREL1] | 52.9 | 1.0E-147 |
| DIP_10001562 | hypothetical protein PFICI_00591 [Pestalotiopsis fici W106-1] | 47 | 0.0 |
| DIP_10001563 | putative dna repair protein rad51 protein [Eutypa lata UCREL1] | 94.8 | 0.0 |
| DIP_10001564 | hypothetical protein PFICI_00589 [Pestalotiopsis fici W106-1] | 38.7 | 7.0E-56 |
| DIP_10001565 | NA | | |
| DIP_10001566 | hypothetical protein PFICI_04366 [Pestalotiopsis fici W106-1] | 77.3 | 0.0 |
| DIP_10001567 | hypothetical protein FPSE_09182 [Fusarium pseudograminearum CS3096] | 71.8 | 1.0E-31 |
| DIP_10001568 | putative mitochondrial pyruvate dehydrogenase kinase protein [Eutypa lata UCREL1] | 75.9 | 0.0 |
| DIP_10001569 | putative chitin synthase protein [Eutypa lata UCREL1] | 76.5 | 0.0 |
| DIP_10001570 | putative chitin synthase 8 protein [Eutypa lata UCREL1] | 83.7 | 0.0 |
| DIP_10001571 | Putative Peptidyl-prolyl cis-trans isomerase [Torrubiella hemipterigena] | 77 | 4.0E-59 |
| DIP_10001572 | NA | | |
| DIP_10001573 | NA | | |
| DIP_10001574 | NA | | |
| DIP_10001575 | NA | | |
| DIP_10001576 | putative mucin family signaling protein [Eutypa lata UCREL1] | 66.1 | 5.0E-86 |
| DIP_10001577 | putative MOSC domain-containing protein [Colletotrichum sublineola] | 48.1 | 1.0E-149 |
| DIP_10001578 | NA | | |
| DIP_10001579 | NA | | |
| DIP_10001580 | NA | | |
| DIP_10001581 | hypothetical protein PFICI_04667 [Pestalotiopsis fici W106-1] | 85.3 | 0.0 |
| DIP_10001582 | putative heat shock protein hsp88 protein [Eutypa lata UCREL1] | 80.7 | 0.0 |
| DIP_10001583 | putative acetylglutamate synthase protein [Neofusicoccum parvum UCRNP2] | 37.9 | 4.0E-87 |
| DIP_10001584 | L-xylulose reductase-like protein [Acremonium chrysogenum ATCC 11550] | 68.4 | 4.0E-115 |
| DIP_10001585 | phd finger and set domain-containing protein [Colletotrichum gloeosporioides Nara gc5] | 51 | 0.0 |
| DIP_10001586 | NA | | |
| DIP_10001587 | NA | | |
| DIP_10001588 | hypothetical protein PV04_08898 [Capronia semiimmersa] | 57.9 | 2.0E-145 |
| DIP_10001589 | hypothetical protein SAPIO_CDS0745 [Scedosporium apiospermum] | 32.7 | 2.0E-31 |
| DIP_10001590 | NA | | |
| DIP_10001591 | NA | | |
| DIP_10001592 | NA | | |
| DIP_10001593 | oxidoreductase [Magnaporthe oryzae 70-15] | 60.6 | 0.0 |
| DIP_10001594 | hypothetical protein PFICI_12207 [Pestalotiopsis fici W106-1] | 67.2 | 1.0E-166 |
| DIP_10001595 | hypothetical protein PFICI_01479 [Pestalotiopsis fici W106-1] | 59.4 | 6.0E-122 |
| DIP_10001596 | NA | | |
| DIP_10001597 | hypothetical protein [Podospora anserina S mat+] | 38.7 | 6.0E-43 |
| DIP_10001598 | hypothetical protein PFICI_01311 [Pestalotiopsis fici W106-1] | 58.1 | 3.0E-62 |
| DIP_10001599 | NA | | |
| DIP_10001600 | NA | | |
| DIP_10001601 | hypothetical protein PFICI_01474 [Pestalotiopsis fici W106-1] | 77.8 | 0.0 |
| DIP_10001602 | hypothetical protein PFICI_01473 [Pestalotiopsis fici W106-1] | 42 | 0.0 |
| DIP_10001603 | hypothetical protein PFICI_01478 [Pestalotiopsis fici W106-1] | 52 | 0.0 |
| DIP_10001604 | hypothetical protein MYCTH_42359 [Myceliophthora thermophila ATCC 42464] | 60 | 0.0 |
| DIP_10001605 | NA | | |
| DIP_10001606 | Exosome complex exonuclease dis3 [Pestalotiopsis fici W106-1] | 83 | 0.0 |
| DIP_10001607 | hypothetical protein PFICI_01477 [Pestalotiopsis fici W106-1] | 42.1 | 1.0E-123 |
| DIP_10001608 | NA | | |
| DIP_10001609 | NA | | |
| DIP_10001610 | NA | | |
| DIP_10001611 | hypothetical protein PFICI_01476 [Pestalotiopsis fici W106-1] | 70.8 | 1.0E-119 |
| DIP_10001612 | hypothetical protein PFICI_01475 [Pestalotiopsis fici W106-1] | 69.2 | 0.0 |
| DIP_10001613 | hypothetical protein VHEMI08083 [Torrubiella hemipterigena] | 52.8 | 3.0E-156 |
| DIP_10001614 | NA | | |
| DIP_10001615 | NA | | |
| DIP_10001616 | hypothetical protein CSUB01_01382 [Colletotrichum sublineola] | 45.8 | 7.0E-171 |
| DIP_10001617 | putative pre-mrna-processing protein prp40 protein [Eutypa lata UCREL1] | 68.8 | 0.0 |
| DIP_10001618 | Translation initiation factor RLI1 [Ceratocystis platani] | 86.6 | 0.0 |
| DIP_10001619 | putative chromodomain helicase hrp3 protein [Eutypa lata UCREL1] | 78.7 | 0.0 |
| DIP_10001620 | putative nad-dependent histone deacetylase sir2 protein [Eutypa lata UCREL1] | 64.4 | 0.0 |
| DIP_10001621 | hypothetical protein PFICI_01482 [Pestalotiopsis fici W106-1] | 44.5 | 0.0 |
| DIP_10001622 | hypothetical protein UCREL1_4432 [Eutypa lata UCREL1] | 39.3 | 2.0E-10 |
| DIP_10001623 | putative mfs transporter protein [Eutypa lata UCREL1] | 58.6 | 0.0 |
| DIP_10001624 | hypothetical protein M419DRAFT_77103 [Trichoderma reesei RUT C-30] | 38.7 | 2.0E-53 |
| DIP_10001625 | putative intracellular protein transport-like protein [Eutypa lata UCREL1] | 69.5 | 0.0 |
| DIP_10001626 | NA | | |
| DIP_10001627 | D-3-phosphoglycerate dehydrogenase [Exophiala aquamarina CBS 119918] | 62.3 | 2.0E-140 |
| DIP_10001628 | putative alkaline serine protease alp1 protein [Eutypa lata UCREL1] | 55.9 | 2.0E-156 |
| DIP_10001629 | hypothetical protein PV09_02278 [Verruconis gallopava] | 56.5 | 0.0 |
| DIP_10001630 | hypothetical protein CFIO01_00223 [Colletotrichum fioriniae PJ7] | 47.4 | 8.0E-170 |
| DIP_10001631 | hypothetical protein V494_03701 [Pseudogymnoascus pannorum VKM F-4513 (FW-928)] | 74.1 | 0.0 |
| DIP_10001632 | hypothetical protein PFICI_01055 [Pestalotiopsis fici W106-1] | 65.3 | 0.0 |
| DIP_10001633 | NA | | |
| DIP_10001634 | hypothetical protein PFICI_01056 [Pestalotiopsis fici W106-1] | 61.5 | 0.0 |
| DIP_10001635 | hypothetical protein VDAG_00490 [Verticillium dahliae VdLs.17] | 36.9 | 2.0E-41 |
| DIP_10001636 | putative rsc complex subunit protein [Eutypa lata UCREL1] | 67.5 | 0.0 |
| DIP_10001637 | hypothetical protein GLRG_01648 [Colletotrichum graminicola M1.001] | 49.7 | 0.0 |
| DIP_10001638 | J-type co-chaperone-like protein [Acremonium chrysogenum ATCC 11550] | 51.1 | 2.0E-75 |
| DIP_10001639 | hypothetical protein PFICI_00514 [Pestalotiopsis fici W106-1] | 71.9 | 2.0E-174 |
| DIP_10001640 | NA | | |
| DIP_10001641 | hypothetical protein PFICI_00513 [Pestalotiopsis fici W106-1] | 67.6 | 0.0 |
| DIP_10001642 | hypothetical protein, partial [Haliangium ochraceum] | 32.2 | 9.0E-07 |
| DIP_10001643 | putative wyosine base formation protein [Eutypa lata UCREL1] | 67.6 | 0.0 |
| DIP_10001644 | putative rrna-processing protein utp23 protein [Eutypa lata UCREL1] | 60.5 | 2.0E-104 |
| DIP_10001645 | putative membrane protein C6F6.13c [Ceratocystis platani] | 56.3 | 0.0 |
| DIP_10001646 | putative aspartyl protease protein [Eutypa lata UCREL1] | 76.8 | 0.0 |
| DIP_10001647 | putative transcription termination factor 2 protein [Eutypa lata UCREL1] | 75.8 | 0.0 |
| DIP_10001648 | NA | | |
| DIP_10001649 | hypothetical protein PV09_07794 [Verruconis gallopava] | 31.9 | 4.0E-07 |
| DIP_10001650 | putative phosphatidylinositol 4-kinase type ii subunit alpha protein [Eutypa lata UCREL1] | 76.1 | 0.0 |
| DIP_10001651 | hypothetical protein UCRPA7_7961 [Togninia minima UCRPA7] | 38.7 | 2.0E-25 |
| DIP_10001652 | hypothetical protein NECHADRAFT_105487 [Nectria haematococca mpVI 77-13-4] | 54.1 | 0.0 |
| DIP_10001653 | hypothetical protein PFICI_05647 [Pestalotiopsis fici W106-1] | 82.6 | 0.0 |
| DIP_10001654 | putative dna mismatch repair protein pms1 protein [Togninia minima UCRPA7] | 77.9 | 0.0 |
| DIP_10001655 | similar to GPI inositol-deacylase [Botrytis cinerea T4] | 57.3 | 0.0 |
| DIP_10001656 | NA | | |
| DIP_10001657 | hypothetical protein PFICI_14598 [Pestalotiopsis fici W106-1] | 62.4 | 0.0 |
| DIP_10001658 | hypothetical protein MGG_01552 [Magnaporthe oryzae 70-15] | 57.5 | 4.0E-106 |
| DIP_10001659 | putative long-chain-fatty-acid- ligase 1 protein [Eutypa lata UCREL1] | 81.1 | 0.0 |
| DIP_10001660 | NA | | |
| DIP_10001661 | hypothetical protein PFICI_04490 [Pestalotiopsis fici W106-1] | 46.1 | 4.0E-163 |
| DIP_10001662 | hypothetical protein PFICI_04497 [Pestalotiopsis fici W106-1] | 65.5 | 0.0 |
| DIP_10001663 | hypothetical protein PFICI_04496 [Pestalotiopsis fici W106-1] | 65.5 | 3.0E-42 |
| DIP_10001664 | oxidoreductase, short chain dehydrogenase/reductase family [Blumeria graminis f. sp. hordei DH14] | 52.2 | 5.0E-53 |
| DIP_10001665 | putative retinol dehydrogenase 13 protein [Togninia minima UCRPA7] | 66.4 | 3.0E-118 |
| DIP_10001666 | NA | | |
| DIP_10001667 | synaptobrevin [Colletotrichum gloeosporioides Cg-14] | 81.3 | 3.0E-132 |
| DIP_10001668 | NA | | |
| DIP_10001669 | hypothetical protein PFICI_07977 [Pestalotiopsis fici W106-1] | 48.5 | 0.0 |
| DIP_10001670 | short-chain dehydrogenase [Colletotrichum higginsianum] | 72.1 | 2.0E-147 |
| DIP_10001671 | Dolichol-phosphate mannosyltransferase subunit-like protein [Acremonium chrysogenum ATCC 11550] | 89.3 | 2.0E-162 |
| DIP_10001672 | hypothetical protein PFICI_15308 [Pestalotiopsis fici W106-1] | 65.7 | 3.0E-146 |
| DIP_10001673 | NA | | |
| DIP_10001674 | NA | | |
| DIP_10001675 | cation transporting ATPase [Trichoderma virens Gv29-8] | 58.5 | 0.0 |
| DIP_10001676 | NA | | |
| DIP_10001677 | RecName: Full=Endo-1,4-beta-xylanase F3; Short=Xylanase F3; AltName: Full=1,4-beta-D-xylan xylanohydrolase F3; Flags: Precursor [Aspergillus oryzae RIB40] | 60.1 | 7.0E-131 |
| DIP_10001678 | aminoglycoside 3'-phosphotransferase/choline kinase domain protein [Metarhizium robertsii] | 44.6 | 5.0E-119 |
| DIP_10001679 | NA | | |
| DIP_10001680 | hypothetical protein PFICI_01471 [Pestalotiopsis fici W106-1] | 67.3 | 0.0 |
| DIP_10001681 | Chloroperoxidase, partial [Metarhizium brunneum ARSEF 3297] | 44.7 | 9.0E-53 |
| DIP_10001682 | Putative protein of unknown function [Podospora anserina S mat+] | 31.6 | 3.0E-84 |
| DIP_10001683 | hypothetical protein UCREL1_3703 [Eutypa lata UCREL1] | 45.8 | 2.0E-109 |
| DIP_10001684 | NA | | |
| DIP_10001685 | putative short-chain dehydrogenase protein [Eutypa lata UCREL1] | 78.1 | 2.0E-124 |
| DIP_10001686 | hypothetical protein CH063_10637 [Colletotrichum higginsianum] | 52.8 | 2.0E-154 |
| DIP_10001687 | hypothetical protein PFICI_15065 [Pestalotiopsis fici W106-1] | 54 | 4.0E-78 |
| DIP_10001688 | hypothetical protein PFICI_00384 [Pestalotiopsis fici W106-1] | 41.4 | 4.0E-49 |
| DIP_10001689 | NA | | |
| DIP_10001690 | putative mapii degradation factor def1 protein [Eutypa lata UCREL1] | 62.2 | 0.0 |
| DIP_10001691 | hypothetical protein PFICI_00386 [Pestalotiopsis fici W106-1] | 39.7 | 5.0E-83 |
| DIP_10001692 | NA | | |
| DIP_10001693 | hypothetical protein EURHEDRAFT_390042 [Aspergillus ruber CBS 135680] | 47.3 | 8.0E-79 |
| DIP_10001694 | hypothetical protein CH063_00504 [Colletotrichum higginsianum] | 30.5 | 9.0E-21 |
| DIP_10001695 | putative major facilitator superfamily protein [Diplodia seriata] | 63.6 | 0.0 |
| DIP_10001696 | oxidoreductase [Azospirillum brasilense] | 41.6 | 2.0E-54 |
| DIP_10001697 | hypothetical protein PFICI_14670 [Pestalotiopsis fici W106-1] | 73.6 | 0.0 |
| DIP_10001698 | hypothetical protein PFICI_14669 [Pestalotiopsis fici W106-1] | 59.5 | 0.0 |
| DIP_10001699 | NA | | |
| DIP_10001700 | NA | | |
| DIP_10001701 | putative kelch domain-containing protein [Eutypa lata UCREL1] | 53.9 | 0.0 |
| DIP_10001702 | NA | | |
| DIP_10001703 | transcription factor cmr1 [Ophiostoma piceae UAMH 11346] | 53.3 | 6.0E-53 |
| DIP_10001704 | hypothetical protein PFICI_15323 [Pestalotiopsis fici W106-1] | 58 | 0.0 |
| DIP_10001705 | NA | | |
| DIP_10001706 | NA | | |
| DIP_10001707 | Fructose-bisphosphate aldolase [Beauveria bassiana D1-5] | 81 | 0.0 |
| DIP_10001708 | NA | | |
| DIP_10001709 | Protein kinase-like (PK-like) [Glarea lozoyensis ATCC 20868] | 38.8 | 3.0E-114 |
| DIP_10001710 | NA | | |
| DIP_10001711 | NA | | |
| DIP_10001712 | stress responsive a b barrel domain protein [Colletotrichum gloeosporioides Nara gc5] | 65.3 | 4.0E-44 |
| DIP_10001713 | hypothetical protein HMPREF1541_08530 [Cyphellophora europaea CBS 101466] | 63.4 | 4.0E-108 |
| DIP_10001714 | hypothetical protein A1O9_04222 [Exophiala aquamarina CBS 119918] | 41.6 | 7.0E-180 |
| DIP_10001715 | putative succinate-semialdehyde dehydrogenase protein [Eutypa lata UCREL1] | 74.4 | 0.0 |
| DIP_10001716 | putative 3-isopropylmalate dehydrogenase protein [Togninia minima UCRPA7] | 75.1 | 0.0 |
| DIP_10001717 | hypothetical protein PV05_08683 [Exophiala xenobiotica] | 70.4 | 0.0 |
| DIP_10001718 | NA | | |
| DIP_10001719 | hypothetical protein M438DRAFT_215983 [Aureobasidium pullulans EXF-150] | 31.7 | 1.0E-14 |
| DIP_10001720 | hypothetical protein NECHADRAFT_61453 [Nectria haematococca mpVI 77-13-4] | 62.8 | 6.0E-105 |
| DIP_10001721 | putative fructose-bisphosphate aldolase protein [Eutypa lata UCREL1] | 51.8 | 3.0E-78 |
| DIP_10001722 | putative glycoside hydrolase family 128 protein [Diaporthe ampelina] | 54 | 1.0E-93 |
| DIP_10001723 | hypothetical protein [Podospora anserina S mat+] | 49.2 | 8.0E-146 |
| DIP_10001724 | nucleosome binding protein [Colletotrichum gloeosporioides Nara gc5] | 87 | 2.0E-56 |
| DIP_10001725 | hypothetical protein PFICI_00742 [Pestalotiopsis fici W106-1] | 46.1 | 4.0E-57 |
| DIP_10001726 | putative autophagy-related protein 11 protein [Eutypa lata UCREL1] | 64.9 | 0.0 |
| DIP_10001727 | hypothetical protein UCREL1_9555 [Eutypa lata UCREL1] | 53.2 | 3.0E-136 |
| DIP_10001728 | putative 5'/3'-nucleotidase SurE [Colletotrichum sublineola] | 38.5 | 2.0E-46 |
| DIP_10001729 | NA | | |
| DIP_10001730 | putative mitochondrial inner membrane translocase subunit tim44 protein [Eutypa lata UCREL1] | 86.1 | 0.0 |
| DIP_10001731 | putative mitochondrial ribosomal protein subunit s18 protein [Eutypa lata UCREL1] | 74.6 | 5.0E-70 |
| DIP_10001732 | hypothetical protein GLRG_06400 [Colletotrichum graminicola M1.001] | 71.4 | 2.0E-112 |
| DIP_10001733 | hypothetical protein MGG_10957 [Magnaporthe oryzae 70-15] | 83 | 0.0 |
| DIP_10001734 | hypothetical protein V499_02740 [Pseudogymnoascus pannorum VKM F-103] | 37 | 1.0E-30 |
| DIP_10001735 | hypothetical protein PFICI_00751 [Pestalotiopsis fici W106-1] | 58.4 | 7.0E-167 |
| DIP_10001736 | hypothetical protein PFICI_02386 [Pestalotiopsis fici W106-1] | 39.3 | 4.0E-95 |
| DIP_10001737 | predicted protein [Chaetomium globosum CBS 148.51] | 37.5 | 2.0E-18 |
| DIP_10001738 | serine threonine-protein kinase [Ustilaginoidea virens] | 49.7 | 3.0E-91 |
| DIP_10001739 | NA | | |
| DIP_10001740 | NA | | |
| DIP_10001741 | NA | | |
| DIP_10001742 | hypothetical protein PFICI_15328 [Pestalotiopsis fici W106-1] | 78.9 | 4.0E-145 |
| DIP_10001743 | putative serine threonine-protein kinase cot-1 protein [Eutypa lata UCREL1] | 80.7 | 0.0 |
| DIP_10001744 | putative cytochrome p450 monooxygenase protein [Togninia minima UCRPA7] | 60.7 | 0.0 |
| DIP_10001745 | hypothetical protein PFICI_05501 [Pestalotiopsis fici W106-1] | 37.1 | 5.0E-44 |
| DIP_10001746 | NA | | |
| DIP_10001747 | NA | | |
| DIP_10001748 | putative tsa family thioredoxin [Phaeomoniella chlamydospora] | 70.9 | 4.0E-123 |
| DIP_10001749 | NA | | |
| DIP_10001750 | putative transcription initiation factor iib protein [Eutypa lata UCREL1] | 77.5 | 0.0 |
| DIP_10001751 | predicted protein [Chaetomium globosum CBS 148.51] | 31.7 | 2.0E-12 |
| DIP_10001752 | hypothetical protein PFICI_00057 [Pestalotiopsis fici W106-1] | 78 | 0.0 |
| DIP_10001753 | exonuclease [Colletotrichum higginsianum] | 70.7 | 6.0E-71 |
| DIP_10001754 | putative phosphoglucomutase 2 protein [Eutypa lata UCREL1] | 89.9 | 0.0 |
| DIP_10001755 | putative vps9 domain protein [Eutypa lata UCREL1] | 62.2 | 0.0 |
| DIP_10001756 | hypothetical protein PFICI_00171 [Pestalotiopsis fici W106-1] | 59 | 0.0 |
| DIP_10001757 | hypothetical protein PFICI_00135 [Pestalotiopsis fici W106-1] | 50.2 | 2.0E-59 |
| DIP_10001758 | putative dna-directed rna polymerases ii 24 kda polypeptide protein [Eutypa lata UCREL1] | 78.7 | 2.0E-143 |
| DIP_10001759 | hypothetical protein CSUB01_07330 [Colletotrichum sublineola] | 55.4 | 5.0E-179 |
| DIP_10001760 | NA | | |
| DIP_10001761 | hypothetical protein PFICI_04477 [Pestalotiopsis fici W106-1] | 49.9 | 0.0 |
| DIP_10001762 | putative glycosyl hydrolase protein [Eutypa lata UCREL1] | 70.1 | 0.0 |
| DIP_10001763 | NA | | |
| DIP_10001764 | hypothetical protein BC1G_02335 [Botrytis cinerea B05.10] | 58.6 | 0.0 |
| DIP_10001765 | putative wd repeat containing protein 36 protein [Eutypa lata UCREL1] | 73.3 | 0.0 |
| DIP_10001766 | hypothetical protein FOC4_g10012001 [Fusarium oxysporum f. sp. cubense race 4] | 66.8 | 0.0 |
| DIP_10001767 | hypothetical protein G7K_2489-t1 [Saitoella complicata NRRL Y-17804] | 31.6 | 3.0E-58 |
| DIP_10001768 | putative d-glycerate 3-kinase protein [Togninia minima UCRPA7] | 57.3 | 7.0E-120 |
| DIP_10001769 | hypothetical protein UCREL1_8632 [Eutypa lata UCREL1] | 48.5 | 6.0E-103 |
| DIP_10001770 | putative ring finger protein [Eutypa lata UCREL1] | 59.8 | 0.0 |
| DIP_10001771 | putative peroxisomal hydratase-dehydrogenase-epimerase protein [Eutypa lata UCREL1] putative peroxisomal hydratase-dehydrogenase-epimerase [Rosellinia necatrix] Sequence ID: | 80.6 | 0.0 |
| DIP_10001772 | NA | | |
| DIP_10001773 | putative vacuolar protein sorting-associated protein 26 [Colletotrichum sublineola] | 91.3 | 0.0 |
| DIP_10001774 | putative 60s ribosomal protein l9 protein [Togninia minima UCRPA7] | 86.5 | 5.0E-124 |
| DIP_10001775 | putative low affinity copper transporter protein [Eutypa lata UCREL1] | 49.5 | 7.0E-48 |
| DIP_10001776 | hypothetical protein PFICI_01333 [Pestalotiopsis fici W106-1] | 51.4 | 0.0 |
| DIP_10001777 | hypothetical protein UCREL1_8816 [Eutypa lata UCREL1] | 59 | 3.0E-88 |
| DIP_10001778 | putative vacuolar atp synthase subunit h protein [Eutypa lata UCREL1] | 73 | 0.0 |
| DIP_10001779 | methylmalonate-semialdehyde dehydrogenase [Magnaporthe oryzae 70-15] | 79.6 | 0.0 |
| DIP_10001780 | hypothetical protein PFICI_00597 [Pestalotiopsis fici W106-1] | 48.6 | 2.0E-133 |
| DIP_10001781 | putative histidine acid phosphatase protein [Eutypa lata UCREL1] | 69.4 | 0.0 |
| DIP_10001782 | hypothetical protein FGSG_00404 [Fusarium graminearum PH-1] | 47.6 | 3.0E-89 |
| DIP_10001783 | putative ubiquitin carboxyl-terminal hydrolase isozyme l3 protein [Eutypa lata UCREL1] | 65.8 | 5.0E-104 |
| DIP_10001784 | hypothetical protein CGLO_10533 [Colletotrichum gloeosporioides Cg-14] | 70.7 | 3.0E-94 |
| DIP_10001785 | DNA-directed RNA polymerase II subunit rpb7 [Pestalotiopsis fici W106-1] | 88.2 | 7.0E-111 |
| DIP_10001786 | mucin [Colletotrichum higginsianum] | 45.9 | 3.0E-77 |
| DIP_10001787 | putative vacuolar protein sorting targeting protein pep1 protein [Eutypa lata UCREL1] | 70.3 | 0.0 |
| DIP_10001788 | hypothetical protein UCREL1_113 [Eutypa lata UCREL1] | 55.6 | 0.0 |
| DIP_10001789 | General amino-acid permease GAP1 [Pestalotiopsis fici W106-1] | 71.3 | 0.0 |
| DIP_10001790 | putative mannosyl-oligosaccharide -alpha-mannosidase protein [Eutypa lata UCREL1] | 73.2 | 0.0 |
| DIP_10001791 | hypothetical protein PFICI_00614 [Pestalotiopsis fici W106-1] | 47.4 | 1.0E-42 |
| DIP_10001792 | hypothetical protein NECHADRAFT_90073 [Nectria haematococca mpVI 77-13-4] | 67.5 | 1.0E-90 |
| DIP_10001793 | Actin-related protein 2/3 complex subunit 1 [Scedosporium apiospermum] | 85 | 0.0 |
| DIP_10001794 | Ran GTPase-activating protein-like protein [Acremonium chrysogenum ATCC 11550] | 78.4 | 0.0 |
| DIP_10001795 | putative dynamin-related protein [Eutypa lata UCREL1] | 86.8 | 0.0 |
| DIP_10001796 | protein phosphatase PP2A regulatory subunit B [Fusarium verticillioides 7600] | 81.7 | 0.0 |
| DIP_10001797 | hypothetical protein PFICI_00622 [Pestalotiopsis fici W106-1] | 77.3 | 0.0 |
| DIP_10001798 | putative amp deaminase protein [Eutypa lata UCREL1] | 73.3 | 0.0 |
| DIP_10001799 | putative cytoskeleton assembly control protein sla1 protein [Eutypa lata UCREL1] | 60.3 | 0.0 |
| DIP_10001800 | putative tlc domain-containing protein [Eutypa lata UCREL1] | 69.3 | 0.0 |
| DIP_10001801 | hypothetical protein PFICI_00627 [Pestalotiopsis fici W106-1] | 60.8 | 9.0E-152 |
| DIP_10001802 | putative solute carrier family 25 member 38 protein [Eutypa lata UCREL1] | 75.2 | 3.0E-169 |
| DIP_10001803 | hypothetical protein UCREL1_3791 [Eutypa lata UCREL1] | 43.4 | 4.0E-84 |
| DIP_10001804 | putative prefoldin subunit protein [Eutypa lata UCREL1] | 75.4 | 1.0E-60 |
| DIP_10001805 | alkaline dihydroceramidase [Grosmannia clavigera kw1407] | 36.2 | 4.0E-59 |
| DIP_10001806 | hypothetical protein PFICI_00632 [Pestalotiopsis fici W106-1] | 55.3 | 0.0 |
| DIP_10001807 | hypothetical protein PFICI_00633 [Pestalotiopsis fici W106-1] | 78 | 6.0E-132 |
| DIP_10001808 | putative pre-mrna splicing protein [Eutypa lata UCREL1] | 79.9 | 0.0 |
| DIP_10001809 | hypothetical protein V496_08533 [Pseudogymnoascus pannorum VKM F-4515 (FW-2607)] | 50.8 | 6.0E-164 |
| DIP_10001810 | hypothetical protein PFICI_00637 [Pestalotiopsis fici W106-1] | 71.9 | 1.0E-79 |
| DIP_10001811 | putative adventurous gliding protein z protein [Eutypa lata UCREL1] | 40.7 | 2.0E-80 |
| DIP_10001812 | hypothetical protein PFICI_14029 [Pestalotiopsis fici W106-1] | 75.8 | 0.0 |
| DIP_10001813 | hypothetical protein PFICI_14032 [Pestalotiopsis fici W106-1] | 44.2 | 6.0E-147 |
| DIP_10001814 | hypothetical protein BGT96224_1593 [Blumeria graminis f. sp. tritici 96224] | 51.5 | 2.0E-32 |
| DIP_10001815 | putative allantoate permease protein [Eutypa lata UCREL1] | 73 | 0.0 |
| DIP_10001816 | putative glycoside hydrolase family 105 protein [Eutypa lata UCREL1] | 67.7 | 0.0 |
| DIP_10001817 | NA | | |
| DIP_10001818 | putative nicotinate-nucleotide diphosphorylase protein [Eutypa lata UCREL1] | 89.3 | 0.0 |
| DIP_10001819 | putative dimethylaniline monooxygenase [Diaporthe ampelina] | 54.8 | 0.0 |
| DIP_10001820 | putative umta methyltransferase family protein [Eutypa lata UCREL1] | 46.7 | 2.0E-83 |
| DIP_10001821 | NA | | |
| DIP_10001822 | hypothetical protein P875_00075871 [Aspergillus parasiticus SU-1] | 54.2 | 2.0E-129 |
| DIP_10001823 | putative acyltransferase protein [Eutypa lata UCREL1] | 40.2 | 4.0E-114 |
| DIP_10001824 | short-chain dehydrogenase reductase family protein [Colletotrichum orbiculare MAFF 240422] | 83.5 | 5.0E-176 |
| DIP_10001825 | hypothetical protein PFICI_13776 [Pestalotiopsis fici W106-1] | 51.2 | 9.0E-144 |
| DIP_10001826 | 2-methylcitrate synthase [Pestalotiopsis fici W106-1] | 87.9 | 0.0 |
| DIP_10001827 | putative isocitrate lyase protein [Eutypa lata UCREL1] | 84.6 | 0.0 |

| | | | |
|---|---|---|---|
| DIP_10001828 | glutamine synthetase [Colletotrichum higginsianum] | 80.7 | 0.0 |
| DIP_10001829 | hypothetical protein VITISV_010526 [Vitis vinifera] | 45.1 | 8.0E-09 |
| DIP_10001830 | NA | | |
| DIP_10001831 | NA | | |
| DIP_10001832 | hypothetical protein UCREL1_2150 [Eutypa lata UCREL1] | 75.2 | 0.0 |
| DIP_10001833 | hypothetical protein UCREL1_1195 [Eutypa lata UCREL1] | 63.2 | 3.0E-77 |
| DIP_10001834 | hypothetical protein PFICI_02420 [Pestalotiopsis fici W106-1] | 66.7 | 0.0 |
| DIP_10001835 | hypothetical protein S40285_05932 [Stachybotrys chlorohalonata IBT 40285] | 52.3 | 0.0 |
| DIP_10001836 | putative glycoside hydrolase family 3 protein [Eutypa lata UCREL1] | 71.7 | 0.0 |
| DIP_10001837 | hypothetical protein PFICI_09800 [Pestalotiopsis fici W106-1] | 64.1 | 0.0 |
| DIP_10001838 | putative acyl-protein thioesterase 1 protein [Eutypa lata UCREL1] | 64.6 | 3.0E-103 |
| DIP_10001839 | putative sedoheptulose- -bisphosphatase protein [Eutypa lata UCREL1] | 66.4 | 3.0E-156 |
| DIP_10001840 | putative 3-dehydroshikimate dehydratase protein [Eutypa lata UCREL1] | 53.9 | 1.0E-130 |
| DIP_10001841 | hypothetical protein PFICI_13805 [Pestalotiopsis fici W106-1] | 57.1 | 5.0E-172 |
| DIP_10001842 | hypothetical protein PFICI_13804 [Pestalotiopsis fici W106-1] | 54 | 8.0E-138 |
| DIP_10001843 | hypothetical protein THITE_2106964 [Thielavia terrestris NRRL 8126] | 60 | 6.0E-80 |
| DIP_10001844 | glycoside hydrolase family 16 protein [Thielavia terrestris NRRL 8126] | 54.7 | 0.0 |
| DIP_10001845 | hypothetical protein CTHT_0010870 [Chaetomium thermophilum var. thermophilum DSM 1495] | 61.6 | 0.0 |
| DIP_10001846 | hypothetical protein FOZG_12971 [Fusarium oxysporum Fo47] | 57.6 | 0.0 |
| DIP_10001847 | hypothetical protein NECHADRAFT_97495 [Nectria haematococca mpVI 77-13-4] | 53.5 | 1.0E-168 |
| DIP_10001848 | putative cop9 signalosome complex subunit 2 protein [Eutypa lata UCREL1] | 77.5 | 0.0 |
| DIP_10001849 | putative pre-mrna-splicing factor cwc25 protein [Togninia minima UCRPA7] | 56.5 | 6.0E-82 |
| DIP_10001850 | GTP-binding protein RHO3 [Pestalotiopsis fici W106-1] | 88.9 | 3.0E-133 |
| DIP_10001851 | hypothetical protein A1O5_09496 [Cladophialophora psammophila CBS 110553] | 43.2 | 7.0E-29 |
| DIP_10001852 | NA | | |
| DIP_10001853 | hypothetical protein TRIVIDRAFT_142895, partial [Trichoderma virens Gv29-8] | 36 | 9.0E-34 |
| DIP_10001854 | hypothetical protein UCREL1_7895 [Eutypa lata UCREL1] | 50.2 | 5.0E-129 |
| DIP_10001855 | hypothetical protein [Podospora anserina S mat+] | 42.1 | 0.0 |
| DIP_10001856 | putative adp-ribose pyrophosphatase protein [Eutypa lata UCREL1] | 55.1 | 8.0E-91 |
| DIP_10001857 | hypothetical protein PFICI_14051 [Pestalotiopsis fici W106-1] | 59.9 | 0.0 |
| DIP_10001858 | mitochondrial genome maintenance protein Mgr2 [Neurospora crassa OR74A] | 81.8 | 1.0E-51 |
| DIP_10001859 | putative ascorbase and cu-oxidase protein [Eutypa lata UCREL1] | 57 | 1.0E-141 |
| DIP_10001860 | putative 6-phosphofructokinase protein [Eutypa lata UCREL1] | 81.3 | 0.0 |
| DIP_10001861 | hypothetical protein PFICI_02383 [Pestalotiopsis fici W106-1] | 42 | 2.0E-146 |
| DIP_10001862 | NA | | |
| DIP_10001863 | hypothetical protein S7711_10443 [Stachybotrys chartarum IBT 7711] | 32.5 | 5.0E-24 |
| DIP_10001864 | putative glutamic acid rich protein [Eutypa lata UCREL1] | 46.5 | 3.0E-103 |
| DIP_10001865 | PARP-type zinc finger-containing protein C2A9.07c [Ceratocystis platani] | 44.7 | 3.0E-68 |
| DIP_10001866 | MFS general substrate transporter [Glarea lozoyensis ATCC 20868] | 53 | 0.0 |
| DIP_10001867 | NA | | |
| DIP_10001868 | NA | | |
| DIP_10001869 | conserved hypothetical protein [Byssochlamys spectabilis No. 5] | 42.7 | 1.0E-17 |
| DIP_10001870 | RNase [Colletotrichum gloeosporioides Nara gc5] | 46.2 | 1.0E-31 |
| DIP_10001871 | hypothetical protein THITE_2116227 [Thielavia terrestris NRRL 8126] | 49.4 | 1.0E-98 |
| DIP_10001872 | hypothetical protein LEMA_P082120.1 [Leptosphaeria maculans JN3] | 34.4 | 2.0E-14 |
| DIP_10001873 | hypothetical protein PV07_05581 [Cladophialophora immunda] | 41 | 0.0 |
| DIP_10001874 | putative negative cofactor 2 complex subunit beta protein [Eutypa lata UCREL1] | 80 | 1.0E-70 |
| DIP_10001875 | hypothetical protein PFICI_02335 [Pestalotiopsis fici W106-1] | 58.4 | 4.0E-107 |
| DIP_10001876 | NA | | |
| DIP_10001877 | hypothetical protein UCREL1_6921 [Eutypa lata UCREL1] | 45.1 | 7.0E-50 |
| DIP_10001878 | Cytochrome c oxidase assembly protein cox-16 [Pestalotiopsis fici W106-1] | 86.1 | 1.0E-69 |
| DIP_10001879 | hypothetical protein PFICI_02339 [Pestalotiopsis fici W106-1] | 48.7 | 0.0 |
| DIP_10001880 | hypothetical protein PFICI_02261 [Pestalotiopsis fici W106-1] | 55 | 0.0 |
| DIP_10001881 | NA | | |
| DIP_10001882 | hypothetical protein PFICI_02288 [Pestalotiopsis fici W106-1] | 33.6 | 2.0E-21 |
| DIP_10001883 | hypothetical protein PFICI_02290 [Pestalotiopsis fici W106-1] | 68 | 1.0E-128 |
| DIP_10001884 | aspartyl-tRNA synthetase [Fusarium oxysporum f. sp. cubense tropical race 4 54006] | 79.1 | 0.0 |
| DIP_10001885 | hypothetical protein FG05_10664 [Fusarium graminearum] | 32.6 | 1.0E-73 |
| DIP_10001886 | probable programed cell death protein [Claviceps purpurea 20.1] | 61 | 1.0E-49 |
| DIP_10001887 | RecName: Full=Ubiquinone biosynthesis protein COQ4, mitochondrial; AltName: Full=Coenzyme Q biosynthesis protein 4; Flags: Precursor [Phaeosphaeria nodorum SN15] | 56.2 | 2.0E-113 |
| DIP_10001888 | hypothetical protein PFICI_02270 [Pestalotiopsis fici W106-1] | 71.8 | 1.0E-50 |
| DIP_10001889 | hypothetical protein PFICI_02271 [Pestalotiopsis fici W106-1] | 37.9 | 4.0E-63 |
| DIP_10001890 | hypothetical protein PFICI_02287 [Pestalotiopsis fici W106-1] | 64.4 | 1.0E-148 |
| DIP_10001891 | hypothetical protein PFICI_02286 [Pestalotiopsis fici W106-1] | 48.4 | 1.0E-101 |
| DIP_10001892 | Rho GTPase [Grosmannia clavigera kw1407] | 91.5 | 8.0E-136 |
| DIP_10001893 | putative rna binding protein [Eutypa lata UCREL1] | 48.4 | 2.0E-55 |
| DIP_10001894 | putative 2-dehydropantoate 2-reductase protein [Eutypa lata UCREL1] | 39.4 | 2.0E-80 |
| DIP_10001895 | hypothetical protein PFICI_02266 [Pestalotiopsis fici W106-1] | 68.9 | 7.0E-70 |
| DIP_10001896 | NA | | |
| DIP_10001897 | NA | | |
| DIP_10001898 | hypothetical protein UCREL1_2333 [Eutypa lata UCREL1] | 63.8 | 1.0E-48 |
| DIP_10001899 | NA | | |
| DIP_10001900 | putative l-ascorbic acid binding protein [Eutypa lata UCREL1] | 58.2 | 1.0E-49 |
| DIP_10001901 | hypothetical protein PFICI_02394 [Pestalotiopsis fici W106-1] | 59.5 | 0.0 |
| DIP_10001902 | hypothetical protein VHEMI06451 [Torrubiella hemipterigena] | 43.5 | 1.0E-164 |
| DIP_10001903 | NA | | |
| DIP_10001904 | putative cbp4 domain-containing protein [Eutypa lata UCREL1] | 73.9 | 1.0E-57 |
| DIP_10001905 | putative set domain-containing protein [Eutypa lata UCREL1] | 45.6 | 2.0E-100 |
| DIP_10001906 | putative wd repeat protein [Eutypa lata UCREL1] | 68.2 | 0.0 |
| DIP_10001907 | protein transporter Sec61 subunit gamma [Magnaporthe oryzae 70-15] | 80 | 1.0E-38 |
| DIP_10001908 | putative rrm domain-containing protein [Eutypa lata UCREL1] | 63.7 | 1.0E-87 |
| DIP_10001909 | hypothetical protein Cob_05000 [Colletotrichum orbiculare MAFF 240422] | 48.6 | 1.0E-50 |
| DIP_10001910 | putative myb dna-binding domain containing protein [Eutypa lata UCREL1] | 52.3 | 2.0E-163 |
| DIP_10001911 | putative 40s ribosomal protein s4 protein [Eutypa lata UCREL1] | 90.5 | 0.0 |
| DIP_10001912 | hypothetical protein PFICI_02796 [Pestalotiopsis fici W106-1] | 73.2 | 0.0 |
| DIP_10001913 | hypothetical protein LEMA_P117350.1 [Leptosphaeria maculans JN3] | 39.2 | 6.0E-86 |
| DIP_10001914 | Nascent polypeptide-associated complex subunit beta [Pestalotiopsis fici W106-1] | 91 | 7.0E-99 |
| DIP_10001915 | hypothetical protein PFICI_02793 [Pestalotiopsis fici W106-1] | 40.8 | 4.0E-111 |
| DIP_10001916 | hypothetical protein FPSE_11700 [Fusarium pseudograminearum CS3096] | 82.5 | 3.0E-89 |
| DIP_10001917 | peroxiredoxin type-2 [Verticillium dahliae VdLs.17] | 67.1 | 1.0E-75 |
| DIP_10001918 | putative duf1348-domain-containing protein [Eutypa lata UCREL1] | 71 | 1.0E-81 |
| DIP_10001919 | hypothetical protein ANO11243_024890 [fungal sp. No.11243] | 35.7 | 2.0E-26 |
| DIP_10001920 | hypothetical protein PFICI_02297 [Pestalotiopsis fici W106-1] | 76.4 | 0.0 |
| DIP_10001921 | putative upf0041 domain protein [Eutypa lata UCREL1] | 61.8 | 1.0E-70 |
| DIP_10001922 | hypothetical protein H634G_01755 [Metarhizium anisopliae BRIP 53293] | 80.9 | 1.0E-50 |
| DIP_10001923 | duf1687 domain containing protein [Colletotrichum gloeosporioides Nara gc5] | 72.7 | 8.0E-74 |
| DIP_10001924 | NA | | |
| DIP_10001925 | hypothetical protein PFICI_02712 [Pestalotiopsis fici W106-1] | 49 | 0.0 |
| DIP_10001926 | NA | | |
| DIP_10001927 | hypothetical protein NECHADRAFT_94695 [Nectria haematococca mpVI 77-13-4] | 53 | 1.0E-114 |
| DIP_10001928 | NA | | |
| DIP_10001929 | NA | | |
| DIP_10001930 | zn 2cys6 transcription factor [Colletotrichum gloeosporioides Nara gc5] | 41 | 9.0E-155 |
| DIP_10001931 | Lipoyl synthase [Pestalotiopsis fici W106-1] | 79.7 | 0.0 |
| DIP_10001932 | NA | | |
| DIP_10001933 | hypothetical protein CSUB01_11738 [Colletotrichum sublineola] | 36.4 | 3.0E-09 |
| DIP_10001934 | hypothetical protein M422DRAFT_66939 [Sphaerobolus stellatus SS14] | 35.3 | 1.0E-21 |
| DIP_10001935 | hypothetical protein UCREL1_7374 [Eutypa lata UCREL1] | 40.4 | 2.0E-48 |
| DIP_10001936 | putative 26s proteasome regulatory subunit rpn5 protein [Eutypa lata UCREL1] | 84.5 | 0.0 |
| DIP_10001937 | putative dihydrofolate reductase protein [Eutypa lata UCREL1] | 42.5 | 9.0E-32 |
| DIP_10001938 | dihydrofolate reductase [Sporothrix brasiliensis 5110] | 50.9 | 1.0E-32 |
| DIP_10001939 | unnamed protein product [Podospora anserina S mat+] | 79 | 5.0E-28 |
| DIP_10001940 | HypE protein [Colletotrichum graminicola M1.001] | 35.5 | 5.0E-16 |
| DIP_10001941 | putative nipsnap family protein [Eutypa lata UCREL1] | 77.6 | 0.0 |
| DIP_10001942 | NA | | |
| DIP_10001943 | hypothetical protein CHGG_06752 [Chaetomium globosum CBS 148.51] | 44.5 | 7.0E-98 |
| DIP_10001944 | putative ribonuclease p subunit pop7 protein [Eutypa lata UCREL1] | 54.3 | 1.0E-59 |
| DIP_10001945 | isotrichodermin C-15 hydroxylase [Magnaporthe oryzae 70-15] | 47 | 7.0E-159 |
| DIP_10001946 | Rhamnogalacturonan acetylesterase [Fusarium oxysporum f. sp. cubense race 1] | 36.8 | 8.0E-41 |
| DIP_10001947 | hypothetical protein CGLO_02459 [Colletotrichum gloeosporioides Cg-14] | 55.6 | 0.0 |
| DIP_10001948 | NA | | |
| DIP_10001949 | purine and uridine phosphorylase [Aureobasidium melanogenum CBS 110374] | 50.7 | 2.0E-105 |
| DIP_10001950 | NA | | |
| DIP_10001951 | NA | | |
| DIP_10001952 | Ae [Aspergillus parasiticus SU-1] | 44.2 | 8.0E-100 |
| DIP_10001953 | hypothetical protein AOR_1_184054 [Aspergillus oryzae RIB40] | 59.4 | 2.0E-145 |
| DIP_10001954 | aristolochene synthase [Aspergillus oryzae RIB40] | 58.7 | 2.0E-137 |
| DIP_10001955 | short chain dehydrogenase/reductase family oxidoreductase [Aspergillus oryzae RIB40] | 56.3 | 3.0E-109 |
| DIP_10001956 | hypothetical protein Ao3042_09586 [Aspergillus oryzae 3.042] | 58.5 | 0.0 |
| DIP_10001957 | cytochrome protein [Aspergillus oryzae 3.042] | 71 | 0.0 |
| DIP_10001958 | cytochrome P450 monooxygenase [Aspergillus oryzae] | 59.9 | 0.0 |
| DIP_10001959 | efflux pump antibiotic resistance protein [Aspergillus oryzae RIB40] | 65.6 | 0.0 |
| DIP_10001960 | NA | | |
| DIP_10001961 | hypothetical protein PFICI_09869 [Pestalotiopsis fici W106-1] | 42.7 | 5.0E-50 |
| DIP_10001962 | hypothetical protein HMPREF1624_03662 [Sporothrix schenckii ATCC 58251] | 82.9 | 2.0E-65 |
| DIP_10001963 | hypothetical protein FGSG_08552 [Fusarium graminearum PH-1] | 74.6 | 2.0E-62 |
| DIP_10001964 | hypothetical protein PFICI_10558 [Pestalotiopsis fici W106-1] | 81.3 | 2.0E-43 |
| DIP_10001965 | hypothetical protein PFICI_10557 [Pestalotiopsis fici W106-1] | 74.7 | 0.0 |
| DIP_10001966 | hypothetical protein FOMG_04595 [Fusarium oxysporum f. sp. melonis 26406] | 32 | 5.0E-85 |
| DIP_10001967 | hypothetical protein FVEG_12415 [Fusarium verticillioides 7600] | 47.9 | 2.0E-153 |
| DIP_10001968 | NA | | |
| DIP_10001969 | hypothetical protein H634G_10395 [Metarhizium anisopliae BRIP 53293] | 37.5 | 4.0E-56 |
| DIP_10001970 | hypothetical protein PFICI_10563 [Pestalotiopsis fici W106-1] | 77.7 | 0.0 |
| DIP_10001971 | hypothetical protein CHGG_06497 [Chaetomium globosum CBS 148.51] | 47.2 | 7.0E-88 |
| DIP_10001972 | NA | | |
| DIP_10001973 | NA | | |
| DIP_10001974 | hypothetical protein PFICI_14107 [Pestalotiopsis fici W106-1] | 49.8 | 0.0 |
| DIP_10001975 | hypothetical protein PFICI_10569 [Pestalotiopsis fici W106-1] | 72.3 | 0.0 |
| DIP_10001976 | major facilitator superfamily transporter [Colletotrichum fioriniae PJ7] | 63.7 | 0.0 |
| DIP_10001977 | putative WW domain binding protein 11 [Colletotrichum sublineola] | 68.2 | 3.0E-106 |
| DIP_10001978 | NA | | |
| DIP_10001979 | putative serine threonine-protein kinase dbf20 protein [Eutypa lata UCREL1] | 80.4 | 0.0 |
| DIP_10001980 | hypothetical protein PFICI_10570 [Pestalotiopsis fici W106-1] | 45.3 | 2.0E-55 |
| DIP_10001981 | hypothetical protein NEUTE1DRAFT_118436 [Neurospora tetrasperma FGSC 2508] | 93.6 | 5.0E-36 |
| DIP_10001982 | hypothetical protein SAPIO_CDS6490 [Scedosporium apiospermum] | 41.5 | 2.0E-79 |
| DIP_10001983 | hypothetical protein UCREL1_8781 [Eutypa lata UCREL1] | 59.5 | 1.0E-113 |
| DIP_10001984 | NA | | |
| DIP_10001985 | NA | | |
| DIP_10001986 | putative short-chain dehydrogenase reductase sdr protein [Eutypa lata UCREL1] | 50.9 | 3.0E-96 |
| DIP_10001987 | NA | | |
| DIP_10001988 | putative chitin biosynthesis protein chs5 protein [Eutypa lata UCREL1] | 67.1 | 0.0 |
| DIP_10001989 | peptidase [Streptomyces sp. CNQ431] | 32.9 | 5.0E-12 |
| DIP_10001990 | hypothetical protein V499_09695 [Pseudogymnoascus pannorum VKM F-103] | 87.3 | 1.0E-93 |
| DIP_10001991 | hypothetical protein PFICI_14130 [Pestalotiopsis fici W106-1] | 63.3 | 6.0E-125 |
| DIP_10001992 | hypothetical protein UCREL1_8942 [Eutypa lata UCREL1] | 60.9 | 4.0E-15 |
| DIP_10001993 | putative dna repair protein [Eutypa lata UCREL1] | 82.6 | 0.0 |
| DIP_10001994 | hypothetical protein UCREL1_8940 [Eutypa lata UCREL1] | 92.5 | 6.0E-25 |
| DIP_10001995 | NA | | |
| DIP_10001996 | putative mads box protein [Eutypa lata UCREL1] | 81.9 | 5.0E-116 |
| DIP_10001997 | putative tetraspanin protein [Eutypa lata UCREL1] | 63.1 | 9.0E-98 |
| DIP_10001998 | hypothetical protein BAUCODRAFT_25915 [Baudoinia compniacensis UAMH 10762] | 42.3 | 3.0E-45 |
| DIP_10001999 | NA | | |
| DIP_10002000 | hypothetical protein TRIVIDRAFT_44566 [Trichoderma virens Gv29-8] | 44.5 | 5.0E-175 |
| DIP_10002001 | putative phosphatidylinositol:udp- c transferase pig-c protein [Eutypa lata UCREL1] | 74.3 | 0.0 |
| DIP_10002002 | NA | | |
| DIP_10002003 | NA | | |
| DIP_10002004 | putative annexin anxc4 protein [Eutypa lata UCREL1] | 58.1 | 0.0 |
| DIP_10002005 | Phytoene dehydrogenase [Pestalotiopsis fici W106-1] | 72.9 | 0.0 |
| DIP_10002006 | putative phytoene synthase protein [Eutypa lata UCREL1] | 66.7 | 0.0 |
| DIP_10002007 | putative thioesterase family protein [Eutypa lata UCREL1] | 54.3 | 2.0E-66 |
| DIP_10002008 | cryptochrome DASH [Pestalotiopsis fici W106-1] | 62.9 | 0.0 |
| DIP_10002009 | hypothetical protein S40288_05957 [Stachybotrys chartarum IBT 40288] | 32.9 | 1.0E-07 |
| DIP_10002010 | putative zinc finger protein 2 protein [Eutypa lata UCREL1] | 60.4 | 7.0E-87 |
| DIP_10002011 | hypothetical protein GLAREA_02484 [Glarea lozoyensis ATCC 20868] | 36.9 | 7.0E-36 |
| DIP_10002012 | hypothetical protein PFICI_07499 [Pestalotiopsis fici W106-1] | 45.8 | 4.0E-151 |
| DIP_10002013 | 2-methylcitrate dehydratase [Pestalotiopsis fici W106-1] | 83.1 | 0.0 |
| DIP_10002014 | hypothetical protein PFICI_13968 [Pestalotiopsis fici W106-1] | 60.5 | 4.0E-103 |
| DIP_10002015 | NA | | |
| DIP_10002016 | Endonuclease/exonuclease/phosphatase [Ophiocordyceps sinensis CO18] | 35.6 | 1.0E-87 |
| DIP_10002017 | putative casein kinase i protein [Eutypa lata UCREL1] | 93.6 | 0.0 |
| DIP_10002018 | hypothetical protein GGTG_11878 [Gaeumannomyces graminis var. tritici R3-111a-1] | 43.8 | 4.0E-145 |
| DIP_10002019 | hypothetical protein PFICI_02821 [Pestalotiopsis fici W106-1] | 65.5 | 0.0 |
| DIP_10002020 | hypothetical protein MGG_03054 [Magnaporthe oryzae 70-15] | 63.6 | 2.0E-141 |
| DIP_10002021 | ankyrin-1 [Colletotrichum higginsianum] | 39.1 | 9.0E-43 |
| DIP_10002022 | PREDICTED: serine/threonine-protein phosphatase 6 regulatory ankyrin repeat subunit A-like, partial [Stegastes partitus] | 39.5 | 3.0E-95 |
| DIP_10002023 | NA | | |
| DIP_10002024 | hypothetical protein PFICI_03035 [Pestalotiopsis fici W106-1] | 36.8 | 2.0E-124 |
| DIP_10002025 | NA | | |
| DIP_10002026 | hypothetical protein PFICI_03041 [Pestalotiopsis fici W106-1] | 49 | 6.0E-119 |
| DIP_10002027 | NAD(P)-binding domain protein, partial [Metarhizium brunneum ARSEF 3297] | 33.7 | 9.0E-54 |
| DIP_10002028 | NA | | |
| DIP_10002029 | hypothetical protein UCREL1_6379 [Eutypa lata UCREL1] | 39.1 | 4.0E-71 |
| DIP_10002030 | purine and uridine phosphorylase [Aureobasidium melanogenum CBS 110374] | 47.2 | 6.0E-91 |
| DIP_10002031 | phd transcription factor [Ophiostoma piceae UAMH 11346] | 54.6 | 1.0E-111 |
| DIP_10002032 | putative synaptobrevin protein [Togninia minima UCRPA7] | 88.7 | 1.0E-56 |
| DIP_10002033 | hypothetical protein SAPIO_CDS6398 [Scedosporium apiospermum] | 61.3 | 1.0E-156 |
| DIP_10002034 | hypothetical protein CGLO_05672 [Colletotrichum gloeosporioides Cg-14] | 80.5 | 1.0E-169 |
| DIP_10002035 | hypothetical protein PFICI_10133 [Pestalotiopsis fici W106-1] | 65.5 | 0.0 |
| DIP_10002036 | NA | | |
| DIP_10002037 | NA | | |
| DIP_10002038 | NA | | |
| DIP_10002039 | hypothetical protein NECHADRAFT_52478 [Nectria haematococca mpVI 77-13-4] | 41.7 | 2.0E-157 |
| DIP_10002040 | NA | | |
| DIP_10002041 | NA | | |
| DIP_10002042 | hypothetical protein FVEG_08414 [Fusarium verticillioides 7600] | 60.3 | 8.0E-25 |
| DIP_10002043 | hypothetical protein PV06_00121 [Exophiala oligosperma] | 77.4 | 2.0E-60 |
| DIP_10002044 | NA | | |
| DIP_10002045 | hypothetical protein CFIO01_00117 [Colletotrichum fioriniae PJ7] | 38.8 | 3.0E-93 |
| DIP_10002046 | hypothetical protein CH063_10067, partial [Colletotrichum higginsianum] | 49 | 1.0E-179 |
| DIP_10002047 | hypothetical protein A1O9_04232 [Exophiala aquamarina CBS 119918] | 41.2 | 8.0E-40 |
| DIP_10002048 | putative siderochrome-iron transporter protein [Eutypa lata UCREL1] | 51.8 | 4.0E-114 |
| DIP_10002049 | hypothetical protein UCRPA7_2912 [Togninia minima UCRPA7] | 54.6 | 4.0E-70 |
| DIP_10002050 | hypothetical protein VHEMI01022 [Torrubiella hemipterigena] | 43.1 | 1.0E-145 |
| DIP_10002051 | hypothetical protein THITE_2082502 [Thielavia terrestris NRRL 8126] | 56 | 3.0E-166 |
| DIP_10002052 | nadh oxidase [Colletotrichum gloeosporioides Nara gc5] | 68.9 | 0.0 |
| DIP_10002053 | hypothetical protein PFICI_06686 [Pestalotiopsis fici W106-1] | 59.6 | 4.0E-100 |
| DIP_10002054 | NA | | |
| DIP_10002055 | putative integral membrane protein [Eutypa lata UCREL1] | 58.7 | 4.0E-177 |
| DIP_10002056 | hypothetical protein UCREL1_10788 [Eutypa lata UCREL1] | 46.3 | 1.0E-60 |
| DIP_10002057 | hypothetical protein PFICI_06676 [Pestalotiopsis fici W106-1] | 72.6 | 4.0E-122 |
| DIP_10002058 | putative ubiquitin homeostasis protein lub1 protein [Eutypa lata UCREL1] | 73.5 | 0.0 |
| DIP_10002059 | putative wd domain-containing protein [Eutypa lata UCREL1] | 81.7 | 0.0 |
| DIP_10002060 | Putative Ribonuclease Trv [Torrubiella hemipterigena] | 63.9 | 3.0E-113 |
| DIP_10002061 | Aminoglycoside phosphotransferase, partial [Metarhizium brunneum ARSEF 3297] | 62.3 | 0.0 |
| DIP_10002062 | putative viral a-type inclusion protein repeat protein [Eutypa lata UCREL1] | 54.7 | 0.0 |
| DIP_10002063 | hypothetical protein UCREL1_11332 [Eutypa lata UCREL1] | 57.3 | 9.0E-123 |
| DIP_10002064 | putative swim zinc finger protein [Eutypa lata UCREL1] | 58.2 | 1.0E-101 |
| DIP_10002065 | hypothetical protein PV07_10054 [Cladophialophora immunda] | 36 | 3.0E-81 |
| DIP_10002066 | aflatoxin biosynthesis ketoreductase nor-1 [Colletotrichum fioriniae PJ7] | 55.2 | 2.0E-99 |
| DIP_10002067 | putative protein rds1 protein [Eutypa lata UCREL1] | 72.4 | 0.0 |
| DIP_10002068 | hypothetical protein PFICI_06688 [Pestalotiopsis fici W106-1] | 60.3 | 0.0 |
| DIP_10002069 | hypothetical protein PFICI_10231 [Pestalotiopsis fici W106-1] | 53 | 0.0 |
| DIP_10002070 | hypothetical protein PFICI_10230 [Pestalotiopsis fici W106-1] | 50.7 | 3.0E-170 |
| DIP_10002071 | NA | | |
| DIP_10002072 | hypothetical protein PFICI_04115 [Pestalotiopsis fici W106-1] | 50.9 | 3.0E-126 |
| DIP_10002073 | NA | | |
| DIP_10002074 | NA | | |
| DIP_10002075 | hypothetical protein PFICI_13020 [Pestalotiopsis fici W106-1] | 40.4 | 1.0E-50 |
| DIP_10002076 | putative swirm domain-containing protein [Togninia minima UCRPA7] | 43.4 | 1.0E-88 |
| DIP_10002077 | hypothetical protein PFICI_13022 [Pestalotiopsis fici W106-1] | 74.7 | 2.0E-161 |
| DIP_10002078 | NA | | |
| DIP_10002079 | putative g2-specific protein kinase nim-1 protein [Eutypa lata UCREL1] | 64.7 | 0.0 |
| DIP_10002080 | putative btb poz domain-containing protein [Eutypa lata UCREL1] | 67.3 | 0.0 |
| DIP_10002081 | hypothetical protein UCREL1_8587 [Eutypa lata UCREL1] | 68.7 | 5.0E-73 |
| DIP_10002082 | hypothetical protein CGLO_15139 [Colletotrichum gloeosporioides Cg-14] | 60.1 | 2.0E-97 |
| DIP_10002083 | putative phd-finger domain-containing protein [Eutypa lata UCREL1] | 65.5 | 4.0E-166 |
| DIP_10002084 | hypothetical protein UCDDA912_g01654 [Diaporthe ampelina] | 63.5 | 2.0E-61 |
| DIP_10002085 | Pc22g19620 [Penicillium rubens Wisconsin 54-1255] | 67 | 0.0 |
| DIP_10002086 | hypothetical protein PFICI_02991 [Pestalotiopsis fici W106-1] | 75.7 | 0.0 |
| DIP_10002087 | hypothetical protein PFICI_02990 [Pestalotiopsis fici W106-1] | 66 | 5.0E-154 |
| DIP_10002088 | putative rec8 protein [Eutypa lata UCREL1] | 50.7 | 2.0E-69 |
| DIP_10002089 | NA | | |
| DIP_10002090 | putative rab protein geranylgeranyltransferase component a protein [Eutypa lata UCREL1] | 57.4 | 0.0 |
| DIP_10002091 | hypothetical protein W97_04094 [Coniosporium apollinis CBS 100218] | 46.3 | 6.0E-46 |
| DIP_10002092 | putative d-arabinono- -lactone oxidase protein [Eutypa lata UCREL1] | 70.7 | 0.0 |
| DIP_10002093 | Putative phosphoribosylaminoimidazole-succinocarboxamide synthase [Podospora anserina S mat+] | 65.1 | 2.0E-141 |
| DIP_10002094 | NA | | |
| DIP_10002095 | hypothetical protein V500_01066 [Pseudogymnoascus pannorum VKM F-4518 (FW-2643)] | 35 | 2.0E-30 |
| DIP_10002096 | putative phosphoribosylaminoimidazole-succinocarboxamide synthase protein [Eutypa lata UCREL1] | 49.5 | 0.0 |
| DIP_10002097 | hypothetical protein PFICI_02960 [Pestalotiopsis fici W106-1] | 33.7 | 9.0E-26 |
| DIP_10002098 | aminotriazole resistance protein [Aspergillus oryzae RIB40] | 46.9 | 1.0E-34 |
| DIP_10002099 | hypothetical protein CCM_03245 [Cordyceps militaris CM01] | 33.1 | 3.0E-40 |
| DIP_10002100 | NA | | |
| DIP_10002101 | hypothetical protein ACRE_065570 [Acremonium chrysogenum ATCC 11550] | 49.8 | 4.0E-161 |
| DIP_10002102 | Protein SCO1 [Pestalotiopsis fici W106-1] | 72.4 | 7.0E-154 |
| DIP_10002103 | hypothetical protein PFICI_02993 [Pestalotiopsis fici W106-1] | 59.7 | 1.0E-155 |
| DIP_10002104 | hypothetical protein PFICI_02994 [Pestalotiopsis fici W106-1] | 59.4 | 0.0 |
| DIP_10002105 | putative gdp gtp exchange factor protein [Togninia minima UCRPA7] | 58.5 | 0.0 |
| DIP_10002106 | NA | | |
| DIP_10002107 | NA | | |
| DIP_10002108 | putative importin subunit beta-1 protein [Eutypa lata UCREL1] | 81.9 | 0.0 |
| DIP_10002109 | 2-isopropylmalate synthase [Pestalotiopsis fici W106-1] | 79 | 0.0 |
| DIP_10002110 | putative ferric reductase protein [Eutypa lata UCREL1] | 68.2 | 0.0 |
| DIP_10002111 | NA | | |
| DIP_10002112 | integral membrane protein [Colletotrichum gloeosporioides Cg-14] | 48.9 | 3.0E-127 |
| DIP_10002113 | hypothetical protein SAPIO_CDS4725 [Scedosporium apiospermum] | 66 | 0.0 |
| DIP_10002114 | putative anaphase-promoting complex protein [Eutypa lata UCREL1] | 72.1 | 0.0 |
| DIP_10002115 | putative complex i intermediate-associated protein 30 protein [Eutypa lata UCREL1] | 67.8 | 4.0E-136 |
| DIP_10002116 | ANTH domain-containing protein [Beauveria bassiana ARSEF 2860] | 81.8 | 0.0 |
| DIP_10002117 | NA | | |
| DIP_10002118 | putative Mating-type protein a-1 [Glarea lozoyensis 74030] | 35 | 4.0E-19 |
| DIP_10002119 | hypothetical protein PFICI_02897 [Pestalotiopsis fici W106-1] | 79.1 | 0.0 |
| DIP_10002120 | putative cytochrome c oxidase subunit protein [Eutypa lata UCREL1] | 78.5 | 2.0E-71 |
| DIP_10002121 | putative ph domain-containing protein [Eutypa lata UCREL1] | 59.3 | 0.0 |
| DIP_10002122 | NA | | |
| DIP_10002123 | putative guanylate kinase protein [Eutypa lata UCREL1] | 64.1 | 2.0E-87 |
| DIP_10002124 | DNA polymerase zeta catalytic subunit [Colletotrichum orbiculare MAFF 240422] | 62.1 | 0.0 |
| DIP_10002125 | Pre-mRNA splicing Prp18-interacting factor [Metarhizium robertsii ARSEF 23] | 66.7 | 0.0 |
| DIP_10002126 | 40S ribosomal protein S9 [Ceratocystis platani] | 92.7 | 2.0E-125 |
| DIP_10002127 | putative 60s ribosomal protein l21 protein [Eutypa lata UCREL1] | 84.4 | 7.0E-97 |
| DIP_10002128 | pH-response regulator protein palI/rim-9 [Verticillium dahliae VdLs.17] | 38.1 | 6.0E-35 |
| DIP_10002129 | hypothetical protein PFICI_02889 [Pestalotiopsis fici W106-1] | 73.7 | 0.0 |
| DIP_10002130 | putative septin [Diaporthe ampelina] | 63.2 | 0.0 |
| DIP_10002131 | hypothetical protein PFICI_02800 [Pestalotiopsis fici W106-1] | 56.3 | 1.0E-107 |
| DIP_10002132 | NA | | |
| DIP_10002133 | putative cdp-alcohol phosphatidyltransferase protein [Eutypa lata UCREL1] | 82.8 | 0.0 |
| DIP_10002134 | hypothetical protein PFICI_02924 [Pestalotiopsis fici W106-1] | 63.5 | 0.0 |
| DIP_10002135 | hypothetical protein PFICI_02923 [Pestalotiopsis fici W106-1] | 57.3 | 3.0E-108 |
| DIP_10002136 | hypothetical protein PFICI_02922 [Pestalotiopsis fici W106-1] | 63.9 | 3.0E-65 |
| DIP_10002137 | putative rna recognition domain-containing protein [Eutypa lata UCREL1] | 42.8 | 1.0E-124 |
| DIP_10002138 | hypothetical protein CTHT_0011960 [Chaetomium thermophilum var. thermophilum DSM 1495] | 60.4 | 0.0 |
| DIP_10002139 | hypothetical protein UCREL1_11339 [Eutypa lata UCREL1] | 47.8 | 2.0E-61 |
| DIP_10002140 | hypothetical protein UCREL1_11340 [Eutypa lata UCREL1] | 62.1 | 3.0E-109 |
| DIP_10002141 | putative translation initiation factor protein [Eutypa lata UCREL1] | 87.7 | 4.0E-69 |
| DIP_10002142 | hypothetical protein PFICI_02883 [Pestalotiopsis fici W106-1] | 61.5 | 0.0 |
| DIP_10002143 | putative coatomer subunit gamma protein [Eutypa lata UCREL1] | 91.6 | 0.0 |
| DIP_10002144 | hypothetical protein PFICI_02925 [Pestalotiopsis fici W106-1] | 57.2 | 0.0 |
| DIP_10002145 | hypothetical protein UCREL1_11343 [Eutypa lata UCREL1] | 64.9 | 8.0E-52 |
| DIP_10002146 | putative phosphoserine phosphatase protein [Eutypa lata UCREL1] | 73.1 | 0.0 |
| DIP_10002147 | Phosphoadenosine phosphosulfate reductase [Pestalotiopsis fici W106-1] | 66.9 | 1.0E-165 |
| DIP_10002148 | NA | | |
| DIP_10002149 | putative iron-sulfur cluster biosynthesis protein isd11 [Diaporthe ampelina] | 76.1 | 4.0E-59 |
| DIP_10002150 | hypothetical protein UCREL1_11347 [Eutypa lata UCREL1] | 37.1 | 2.0E-64 |
| DIP_10002151 | putative splicing factor 3b subunit 5 protein [Eutypa lata UCREL1] | 83.9 | 7.0E-54 |
| DIP_10002152 | putative mage family protein [Eutypa lata UCREL1] | 61.5 | 1.0E-126 |
| DIP_10002153 | hypothetical protein PFICI_02950 [Pestalotiopsis fici W106-1] | 30.1 | 2.0E-83 |
| DIP_10002154 | putative elongation factor 1-alpha protein [Eutypa lata UCREL1] | 93.1 | 0.0 |
| DIP_10002155 | NA | | |
| DIP_10002156 | hypothetical protein UCDDS831_g00870 [Diplodia seriata] | 47.9 | 6.0E-111 |
| DIP_10002157 | putative duf718 domain-containing protein [Eutypa lata UCREL1] | 69.7 | 5.0E-72 |
| DIP_10002158 | Major facilitator superfamily domain, general substrate transporter [Metarhizium album ARSEF 1941] | 43.3 | 3.0E-52 |
| DIP_10002159 | hypothetical protein PFICI_02904 [Pestalotiopsis fici W106-1] | 53 | 4.0E-101 |
| DIP_10002160 | 60S ribosomal protein L37 [Colletotrichum higginsianum] | 89.4 | 2.0E-56 |
| DIP_10002161 | putative autophagocytosis protein [Eutypa lata UCREL1] | 79.4 | 0.0 |
| DIP_10002162 | putative inner membrane protein yico protein [Eutypa lata UCREL1] | 73.6 | 0.0 |
| DIP_10002163 | hypothetical protein CFIO01_04766 [Colletotrichum fioriniae PJ7] | 31.2 | 3.0E-36 |
| DIP_10002164 | hypothetical protein V496_01753 [Pseudogymnoascus pannorum VKM F-4515 (FW-2607)] | 67.7 | 0.0 |
| DIP_10002165 | hypothetical protein SETTUDRAFT_152856 [Setosphaeria turcica Et28A] | 65.9 | 6.0E-134 |
| DIP_10002166 | hypothetical protein S7711_06147 [Stachybotrys chartarum IBT 7711] | 62.1 | 0.0 |
| DIP_10002167 | NA | | |
| DIP_10002168 | putative hexose transporter-like protein [Eutypa lata UCREL1] | 76.4 | 0.0 |
| DIP_10002169 | hypothetical protein MYCFIDRAFT_78988 [Pseudocercospora fijiensis CIRAD86] | 42 | 2.0E-74 |
| DIP_10002170 | putative fmp21-like protein [Eutypa lata UCREL1] | 65.6 | 4.0E-44 |
| DIP_10002171 | hypothetical protein CGLO_07087 [Colletotrichum gloeosporioides Cg-14] | 44 | 1.0E-37 |
| DIP_10002172 | hypothetical protein PFICI_08418 [Pestalotiopsis fici W106-1] | 51.2 | 3.0E-77 |
| DIP_10002173 | NA | | |
| DIP_10002174 | NA | | |
| DIP_10002175 | putative spx domain-containing protein [Eutypa lata UCREL1] | 78 | 0.0 |
| DIP_10002176 | NA | | |
| DIP_10002177 | putative vacuolar sorting protein 39 domain 1 protein [Eutypa lata UCREL1] | 60.4 | 0.0 |
| DIP_10002178 | hypothetical protein V499_03368 [Pseudogymnoascus pannorum VKM F-103] | 53.4 | 1.0E-15 |
| DIP_10002179 | Calcium/calmodulin-dependent protein kinase type 1G, partial [Pelecanus crispus] | 30.2 | 5.0E-10 |
| DIP_10002180 | putative glycoside hydrolase family 76 protein [Phaeomoniella chlamydospora] | 49.1 | 9.0E-121 |
| DIP_10002181 | putative tryptophan synthase protein [Eutypa lata UCREL1] | 83.8 | 0.0 |
| DIP_10002182 | hypothetical protein PFICI_08396 [Pestalotiopsis fici W106-1] | 56.2 | 4.0E-69 |
| DIP_10002183 | hypothetical protein MYCTH_2313216 [Myceliophthora thermophila ATCC 42464] | 34.5 | 2.0E-40 |
| DIP_10002184 | putative aspartate aminotransferase protein [Eutypa lata UCREL1] | 85.5 | 0.0 |
| DIP_10002185 | hypothetical protein TSTA_095930 [Talaromyces stipitatus ATCC 10500] | 33.3 | 5.0E-17 |
| DIP_10002186 | hypothetical protein TRIVIDRAFT_122772, partial [Trichoderma virens Gv29-8] | 33.3 | 6.0E-153 |
| DIP_10002187 | hypothetical protein SMAC_08329 [Sordaria macrospora k-hell] | 55.9 | 0.0 |
| DIP_10002188 | hypothetical protein PISL3812_06440 [Talaromyces islandicus] | 31.5 | 4.0E-47 |
| DIP_10002189 | alcohol dehydrogenase, putative [Byssochlamys spectabilis No. 5] | 46.7 | 4.0E-108 |
| DIP_10002190 | ZIP4 protein [Sordaria macrospora k-hell] | 40.5 | 1.0E-122 |
| DIP_10002191 | zn cluster transcription factor [Colletotrichum gloeosporioides Nara gc5] | 60.4 | 8.0E-156 |
| DIP_10002192 | putative exocyst complex component sec6 protein [Eutypa lata UCREL1] | 80.5 | 0.0 |
| DIP_10002193 | putative initiation-specific alpha-1,6-mannosyltransferase [Colletotrichum sublineola] | 44.4 | 1.0E-69 |

| | | | |
|---|---|---|---|
| DIP_10002194 | putative taurine catabolism dioxygenase protein [Eutypa lata UCREL1] | 75.7 | 0.0 |
| DIP_10002195 | NA | | |
| DIP_10002196 | hypothetical protein PFICI_09325 [Pestalotiopsis fici W106-1] | 59.7 | 0.0 |
| DIP_10002197 | hypothetical protein PFICI_09326 [Pestalotiopsis fici W106-1] | 77.6 | 0.0 |
| DIP_10002198 | hypothetical protein CGLO_16490 [Colletotrichum gloeosporioides Cg-14] | 74.9 | 0.0 |
| DIP_10002199 | hypothetical protein PFICI_11219 [Pestalotiopsis fici W106-1] | 57 | 3.0E-60 |
| DIP_10002200 | putative 60s ribosomal protein l3 protein [Eutypa lata UCREL1] | 91.1 | 0.0 |
| DIP_10002201 | hypothetical protein PFICI_11217 [Pestalotiopsis fici W106-1] | 66.7 | 0.0 |
| DIP_10002202 | NA | | |
| DIP_10002203 | hypothetical protein PFICI_11216 [Pestalotiopsis fici W106-1] | 73 | 0.0 |
| DIP_10002204 | hypothetical protein PFICI_11215 [Pestalotiopsis fici W106-1] | 46.5 | 0.0 |
| DIP_10002205 | hypothetical protein G647_06580 [Cladophialophora carrionii CBS 160.54] | 42.1 | 3.0E-151 |
| DIP_10002206 | putative zn2cys6 transcription factor protein [Eutypa lata UCREL1] | 60.3 | 0.0 |
| DIP_10002207 | NA | | |
| DIP_10002208 | putative nad-dependent epimerase dehydratase protein [Eutypa lata UCREL1] | 47.5 | 9.0E-67 |
| DIP_10002209 | NA | | |
| DIP_10002210 | putative kynurenine 3-monooxygenase protein [Eutypa lata UCREL1] | 62.6 | 0.0 |
| DIP_10002211 | NA | | |
| DIP_10002212 | NA | | |
| DIP_10002213 | Non-specific lipid-transfer protein [Pestalotiopsis fici W106-1] | 86.5 | 0.0 |
| DIP_10002214 | putative maltose permease protein [Eutypa lata UCREL1] | 78.2 | 0.0 |
| DIP_10002215 | hypothetical protein PFICI_13749 [Pestalotiopsis fici W106-1] | 59.8 | 0.0 |
| DIP_10002216 | putative integral membrane protein [Eutypa lata UCREL1] | 51.4 | 2.0E-79 |
| DIP_10002217 | hypothetical protein UCREL1_311 [Eutypa lata UCREL1] | 64.1 | 5.0E-33 |
| DIP_10002218 | hypothetical protein PFICI_09083 [Pestalotiopsis fici W106-1] | 60.9 | 1.0E-163 |
| DIP_10002219 | hypothetical protein PFICI_09084 [Pestalotiopsis fici W106-1] | 59.6 | 0.0 |
| DIP_10002220 | putative secreted aspartic proteinase protein [Eutypa lata UCREL1] | 42 | 1.0E-99 |
| DIP_10002221 | putative sterol esterase precursor protein [Eutypa lata UCREL1] | 62.6 | 0.0 |
| DIP_10002222 | NA | | |
| DIP_10002223 | putative mfs monocarboxylic acid transporter protein [Eutypa lata UCREL1] | 66 | 0.0 |
| DIP_10002224 | hypothetical protein PFICI_09070 [Pestalotiopsis fici W106-1] | 48.1 | 2.0E-30 |
| DIP_10002225 | Threonine synthase [Pestalotiopsis fici W106-1] | 83.8 | 0.0 |
| DIP_10002226 | mip family channel protein [Colletotrichum orbiculare MAFF 240422] | 77.6 | 8.0E-159 |
| DIP_10002227 | NA | | |
| DIP_10002228 | hypothetical protein GLRG_06266 [Colletotrichum graminicola M1.001] | 44.3 | 2.0E-91 |
| DIP_10002229 | serine/threonine protein kinase, partial [Streptomyces sp. R1-NS-10] | 31.3 | 3.0E-08 |
| DIP_10002230 | putative mitochondrial inner membrane protein 1 protein [Eutypa lata UCREL1] | 60.8 | 4.0E-140 |
| DIP_10002231 | putative vacuolar protein sorting-associated protein 13 protein [Eutypa lata UCREL1] | 80.5 | 0.0 |
| DIP_10002232 | putative cytochrome p450 52c1 protein [Eutypa lata UCREL1] | 42 | 5.0E-155 |
| DIP_10002233 | hypothetical protein PFICI_08465 [Pestalotiopsis fici W106-1] | 53 | 2.0E-19 |
| DIP_10002234 | hypothetical protein UCREL1_8330 [Eutypa lata UCREL1] | 48.6 | 3.0E-153 |
| DIP_10002235 | fungal specific transcription factor domain-containing protein [Colletotrichum gloeosporioides Nara gc5] | 38.6 | 9.0E-112 |
| DIP_10002236 | thiol-specific monooxygenase [Colletotrichum fioriniae PJ7] | 38.6 | 5.0E-120 |
| DIP_10002237 | hypothetical protein PFICI_13564 [Pestalotiopsis fici W106-1] | 52.9 | 4.0E-161 |
| DIP_10002238 | hypothetical protein PFICI_07012 [Pestalotiopsis fici W106-1] | 47.1 | 5.0E-75 |
| DIP_10002239 | putative set domain-containing protein [Eutypa lata UCREL1] | 39.8 | 9.0E-105 |
| DIP_10002240 | hypothetical protein UCREL1_9231 [Eutypa lata UCREL1] | 46.4 | 9.0E-39 |
| DIP_10002241 | hypothetical protein CTHT_0019830 [Chaetomium thermophilum var. thermophilum DSM 1495] | 50.7 | 8.0E-20 |
| DIP_10002242 | putative flavodoxin reductase family protein [Eutypa lata UCREL1] | 50.1 | 0.0 |
| DIP_10002243 | NA | | |
| DIP_10002244 | putative riboflavin aldehyde-forming enzyme protein [Eutypa lata UCREL1] | 38.8 | 2.0E-12 |
| DIP_10002245 | NA | | |
| DIP_10002246 | hypothetical protein EMCG_00061 [Emmonsia crescens UAMH 3008] | 37.4 | 4.0E-99 |
| DIP_10002247 | NA | | |
| DIP_10002248 | tpr domain protein [Colletotrichum orbiculare MAFF 240422] | 67.5 | 0.0 |
| DIP_10002249 | putative key lime pathogenicity protein [Eutypa lata UCREL1] | 51.7 | 6.0E-142 |
| DIP_10002250 | misfolded glycoproteins degradation protein [Colletotrichum orbiculare MAFF 240422] | 45 | 7.0E-133 |
| DIP_10002251 | fungal specific transcription factor domain-containing protein [Colletotrichum fioriniae PJ7] | 48.3 | 6.0E-180 |
| DIP_10002252 | hypothetical protein ACRE_075350 [Acremonium chrysogenum ATCC 11550] | 30.5 | 3.0E-80 |
| DIP_10002253 | hypothetical protein HIM_08874 [Hirsutella minnesotensis 3608] | 47.6 | 1.0E-152 |
| DIP_10002254 | hypothetical protein HIM_08875 [Hirsutella minnesotensis 3608] | 43.4 | 0.0 |
| DIP_10002255 | O-methyl transferase B [Metarhizium robertsii ARSEF 23] | 51.9 | 1.0E-140 |
| DIP_10002256 | hypothetical protein S40293_05700 [Stachybotrys chartarum IBT 40293] | 42.2 | 1.0E-138 |
| DIP_10002257 | hypothetical protein [Podospora anserina S mat+] | 56.4 | 0.0 |
| DIP_10002258 | Cytochrome P450 [Glarea lozoyensis ATCC 20868] | 42.2 | 5.0E-159 |
| DIP_10002259 | NAD(P)-binding domain protein [Ophiocordyceps sinensis CO18] | 31.5 | 8.0E-43 |
| DIP_10002260 | NA | | |
| DIP_10002261 | NA | | |
| DIP_10002262 | putative btn-1 protein [Eutypa lata UCREL1] | 65.7 | 0.0 |
| DIP_10002263 | autophagic death protein Aut7/IDI-7 [Sphaerulina musiva SO2202] | 100 | 1.0E-82 |
| DIP_10002264 | hypothetical protein V496_09036 [Pseudogymnoascus pannorum VKM F-4515 (FW-2607)] | 54.3 | 2.0E-144 |
| DIP_10002265 | putative geranylgeranyl transferase type-2 subunit alpha protein [Eutypa lata UCREL1] | 51.9 | 3.0E-93 |
| DIP_10002266 | putative lipase 2 protein [Eutypa lata UCREL1] | 67.5 | 2.0E-169 |
| DIP_10002267 | NA | | |
| DIP_10002268 | Topoisomerase II-associated protein PAT1 [Metarhizium album ARSEF 1941] | 54.3 | 4.0E-119 |
| DIP_10002269 | NA | | |
| DIP_10002270 | NA | | |
| DIP_10002271 | putative -domain-containing protein [Togninia minima UCRPA7] | 58.9 | 0.0 |
| DIP_10002272 | hypothetical protein UCREL1_4341 [Eutypa lata UCREL1] | 55.1 | 2.0E-18 |
| DIP_10002273 | putative cytochrome b5 [Acremonium chrysogenum ATCC 11550] | 52.8 | 2.0E-33 |
| DIP_10002274 | NA | | |
| DIP_10002275 | hypothetical protein PFICI_12330 [Pestalotiopsis fici W106-1] | 56 | 5.0E-73 |
| DIP_10002276 | hypothetical protein ACRE_017630 [Acremonium chrysogenum ATCC 11550] | 63.2 | 6.0E-142 |
| DIP_10002277 | hypothetical protein PFICI_12325 [Pestalotiopsis fici W106-1] | 49.3 | 7.0E-50 |
| DIP_10002278 | hypothetical protein PFICI_12324 [Pestalotiopsis fici W106-1] | 48.5 | 3.0E-34 |
| DIP_10002279 | f-box domain-containing protein [Colletotrichum orbiculare MAFF 240422] | 66.4 | 0.0 |
| DIP_10002280 | related to peptidylprolyl isomerase (cyclophilin)-like protein [Fusarium fujikuroi IMI 58289] | 63.2 | 0.0 |
| DIP_10002281 | NA | | |
| DIP_10002282 | NA | | |
| DIP_10002283 | NA | | |
| DIP_10002284 | hypothetical protein FAVG1_02342 [Fusarium avenaceum] | 34.3 | 0.0 |
| DIP_10002285 | NA | | |
| DIP_10002286 | NA | | |
| DIP_10002287 | putative heat shock factor protein hsf24 protein [Eutypa lata UCREL1] | 71.3 | 0.0 |
| DIP_10002288 | hypothetical protein FOPG_19181 [Fusarium oxysporum f. sp. conglutinans race 2 54008] | 34.5 | 1.0E-20 |
| DIP_10002289 | NA | | |
| DIP_10002290 | NA | | |
| DIP_10002291 | NA | | |
| DIP_10002292 | NA | | |
| DIP_10002293 | hypothetical protein VHEMI06405 [Torrubiella hemipterigena] | 35.6 | 3.0E-107 |
| DIP_10002294 | NA | | |
| DIP_10002295 | putative deoxyhypusine synthase protein [Eutypa lata UCREL1] | 86.7 | 0.0 |
| DIP_10002296 | hypothetical protein CH063_15220 [Colletotrichum higginsianum] | 61.5 | 2.0E-19 |
| DIP_10002297 | hypothetical protein PFICI_09277 [Pestalotiopsis fici W106-1] | 45.1 | 2.0E-178 |
| DIP_10002298 | putative inositol polyphosphate kinase [Ceratocystis platani] | 55.7 | 2.0E-133 |
| DIP_10002299 | hypothetical protein AOL_s00080g120 [Arthrobotrys oligospora ATCC 24927] | 38 | 5.0E-81 |
| DIP_10002300 | predicted protein [Aspergillus terreus NIH2624] | 42.2 | 7.0E-68 |
| DIP_10002301 | Alpha-1,3/1,6-mannosyltransferase alg-2 [Pestalotiopsis fici W106-1] | 61.7 | 0.0 |
| DIP_10002302 | NA | | |
| DIP_10002303 | xanthine dehydrogenase [Colletotrichum gloeosporioides Cg-14] | 83.2 | 0.0 |
| DIP_10002304 | putative xanthine dehydrogenase protein [Eutypa lata UCREL1] | 80.7 | 0.0 |
| DIP_10002305 | putative 37s ribosomal protein rsm25 protein [Eutypa lata UCREL1] | 56.6 | 1.0E-100 |
| DIP_10002306 | putative pentatricopeptide repeat protein [Eutypa lata UCREL1] | 49 | 7.0E-166 |
| DIP_10002307 | putative cytochrome-c oxidase chain viic protein [Eutypa lata UCREL1] | 74.3 | 4.0E-34 |
| DIP_10002308 | putative atp-dependent rna helicase dob1 protein [Eutypa lata UCREL1] | 82.5 | 0.0 |
| DIP_10002309 | ornithine cyclodeaminase [Metarhizium guizhouense ARSEF 977] | 41 | 1.0E-94 |
| DIP_10002310 | hypothetical protein NECHADRAFT_34486 [Nectria haematococca mpVI 77-13-4] | 56.9 | 0.0 |
| DIP_10002311 | duf895 domain membrane protein [Colletotrichum gloeosporioides Nara gc5] | 65.9 | 0.0 |
| DIP_10002312 | hypothetical protein SAPIO_CDS6575 [Scedosporium apiospermum] | 64.3 | 8.0E-178 |
| DIP_10002313 | NA | | |
| DIP_10002314 | NA | | |
| DIP_10002315 | putative membrane protein [Eutypa lata UCREL1] | 50.7 | 1.0E-88 |
| DIP_10002316 | aminotransferase C6B12.04c [Pestalotiopsis fici W106-1] | 85.5 | 0.0 |
| DIP_10002317 | UDP-N-acetylglucosamine transferase subunit ALG14 [Verticillium dahliae VdLs.17] | 34.1 | 5.0E-33 |
| DIP_10002318 | hypothetical protein PFICI_08815 [Pestalotiopsis fici W106-1] | 39.6 | 8.0E-174 |
| DIP_10002319 | putative inositolphosphorylceramide-b c-26 hydroxylase protein [Eutypa lata UCREL1] | 77.8 | 0.0 |
| DIP_10002320 | hypothetical protein PFICI_09408 [Pestalotiopsis fici W106-1] | 78.4 | 0.0 |
| DIP_10002321 | hypothetical protein ACRE_074890 [Acremonium chrysogenum ATCC 11550] | 65.1 | 3.0E-65 |
| DIP_10002322 | hypothetical protein FGSG_04731 [Fusarium graminearum PH-1] | 43.3 | 1.0E-129 |
| DIP_10002323 | putative fad binding domain containing protein [Eutypa lata UCREL1] | 58.1 | 0.0 |
| DIP_10002324 | hypothetical protein PFICI_08994 [Pestalotiopsis fici W106-1] | 49.6 | 1.0E-104 |
| DIP_10002325 | hypothetical protein HK57_00666 [Aspergillus ustus] | 39.1 | 4.0E-65 |
| DIP_10002326 | hypothetical protein UCREL1_3221 [Eutypa lata UCREL1] | 41.3 | 5.0E-36 |
| DIP_10002327 | hypothetical protein PFICI_09260 [Pestalotiopsis fici W106-1] | 41.3 | 1.0E-141 |
| DIP_10002328 | PREDICTED: probable pathogenesis-related protein CaO19.6200 [Fragaria vesca subsp. vesca] | 30.5 | 8.0E-09 |
| DIP_10002329 | hypothetical protein PFICI_09330 [Pestalotiopsis fici W106-1] | 50 | 7.0E-149 |
| DIP_10002330 | hypothetical protein PFICI_09331 [Pestalotiopsis fici W106-1] | 43.7 | 6.0E-137 |
| DIP_10002331 | glycoside hydrolase family 13 [Nectria haematococca mpVI 77-13-4] | 83.4 | 0.0 |
| DIP_10002332 | ATP-dependent Clp protease ATP-binding subunit ClpX [Fusarium oxysporum FOSC 3-a] | 59.7 | 0.0 |
| DIP_10002333 | GTP cyclohydrolase I [Physarum polycephalum] | 53.9 | 3.0E-65 |
| DIP_10002334 | conserved hypothetical protein [Aspergillus terreus NIH2624] | 38.6 | 3.0E-148 |
| DIP_10002335 | Cytochrome c oxidase polypeptide 5 [Pestalotiopsis fici W106-1] | 74.9 | 2.0E-83 |
| DIP_10002336 | hypothetical protein PFICI_09004 [Pestalotiopsis fici W106-1] | 47.8 | 0.0 |
| DIP_10002337 | hypothetical protein T310_0088 [Rasamsonia emersonii CBS 393.64] | 35.3 | 3.0E-58 |
| DIP_10002338 | NA | | |
| DIP_10002339 | hypothetical protein PFICI_11456 [Pestalotiopsis fici W106-1] | 46 | 6.0E-142 |
| DIP_10002340 | putative cdp-diacylglycerol-inositol 3-phosphatidyltransferase pis protein [Eutypa lata UCREL1] | 71.2 | 5.0E-159 |
| DIP_10002341 | orotidine 5'-phosphate decarboxylase [Pestalotiopsis microspora] | 75.2 | 0.0 |
| DIP_10002342 | putative fungal specific transcription factor domain-containing protein [Eutypa lata UCREL1] | 60.7 | 0.0 |
| DIP_10002343 | hypothetical protein SPBR_07740 [Sporothrix brasiliensis 5110] | 46.2 | 6.0E-56 |
| DIP_10002344 | NA | | |
| DIP_10002345 | putative flagellar calcium-binding protein [Eutypa lata UCREL1] | 50 | 6.0E-49 |
| DIP_10002346 | NA | | |
| DIP_10002347 | putative serine threonine-protein kinase ssp1 protein [Togninia minima UCRPA7] | 51.9 | 0.0 |
| DIP_10002348 | NA | | |
| DIP_10002349 | putative dehydrodolichyl diphosphate synthase protein [Eutypa lata UCREL1] | 72.9 | 2.0E-169 |
| DIP_10002350 | hypothetical protein PFICI_09321 [Pestalotiopsis fici W106-1] | 50.7 | 0.0 |
| DIP_10002351 | hypothetical protein PFICI_09322 [Pestalotiopsis fici W106-1] | 60.9 | 0.0 |
| DIP_10002352 | hypothetical protein NECHADRAFT_57703 [Nectria haematococca mpVI 77-13-4] | 75.3 | 0.0 |
| DIP_10002353 | putative wsc domain-containing protein [Eutypa lata UCREL1] | 57.6 | 4.0E-132 |
| DIP_10002354 | NA | | |
| DIP_10002355 | hypothetical protein PFICI_09282 [Pestalotiopsis fici W106-1] | 51.1 | 0.0 |
| DIP_10002356 | NA | | |
| DIP_10002357 | NA | | |
| DIP_10002358 | hypothetical protein PFICI_08655 [Pestalotiopsis fici W106-1] | 62.3 | 0.0 |
| DIP_10002359 | hypothetical protein CH063_07354 [Colletotrichum higginsianum] | 35.4 | 7.0E-48 |
| DIP_10002360 | putative duf895 domain membrane protein [Eutypa lata UCREL1] | 71 | 0.0 |
| DIP_10002361 | NA | | |
| DIP_10002362 | hypothetical protein PFICI_00279 [Pestalotiopsis fici W106-1] | 39.7 | 9.0E-172 |
| DIP_10002363 | hypothetical protein OIDMADRAFT_16886 [Oidiodendron maius Zn] | 92.1 | 0.0 |
| DIP_10002364 | hypothetical protein SPSK_01825 [Sporothrix schenckii 1099-18] | 43.2 | 7.0E-59 |
| DIP_10002365 | hypothetical protein SMAC_06757 [Sordaria macrospora k-hell] | 35.4 | 0.0 |
| DIP_10002366 | putative trfa protein [Eutypa lata UCREL1] | 49 | 4.0E-180 |
| DIP_10002367 | putative regulatory protein cys-3 protein [Eutypa lata UCREL1] | 69.2 | 5.0E-70 |
| DIP_10002368 | hypothetical protein PFICI_09265 [Pestalotiopsis fici W106-1] | 40.6 | 2.0E-138 |
| DIP_10002369 | hypothetical protein NFIA_094280 [Neosartorya fischeri NRRL 181] | 31.3 | 1.0E-17 |
| DIP_10002370 | hypothetical protein V492_07800 [Pseudogymnoascus pannorum VKM F-4246] | 39.1 | 2.0E-171 |
| DIP_10002371 | NA | | |
| DIP_10002372 | NA | | |
| DIP_10002373 | hypothetical protein CHGG_08223 [Chaetomium globosum CBS 148.51] | 43.3 | 2.0E-56 |
| DIP_10002374 | hypothetical protein H634G_00767 [Metarhizium anisopliae BRIP 53293] | 30.3 | 9.0E-22 |
| DIP_10002375 | 2OG-Fe(II) oxygenase [Colletotrichum higginsianum] | 59.5 | 2.0E-110 |
| DIP_10002376 | hypothetical protein PFICI_09099 [Pestalotiopsis fici W106-1] | 62.8 | 2.0E-135 |
| DIP_10002377 | putative tpa inducible protein [Eutypa lata UCREL1] | 58.1 | 5.0E-174 |
| DIP_10002378 | predicted protein [Nectria haematococca mpVI 77-13-4] | 76.7 | 0.0 |
| DIP_10002379 | putative l-aminoadipate-semialdehyde dehydrogenase large subunit protein [Botrytis cinerea BcDW1] | 64.5 | 0.0 |
| DIP_10002380 | hypothetical protein PFICI_09076 [Pestalotiopsis fici W106-1] | 48.5 | 5.0E-99 |
| DIP_10002381 | putative g2 m phase checkpoint control protein [Togninia minima UCRPA7] | 53.4 | 4.0E-111 |
| DIP_10002382 | NADH dehydrogenase (ubiquinone) 1 alpha subcomplex assembly factor 5 [Scedosporium apiospermum] | 60.2 | 7.0E-139 |
| DIP_10002383 | putative ap-2 complex subunit alpha protein [Eutypa lata UCREL1] | 87.4 | 0.0 |
| DIP_10002384 | NA | | |
| DIP_10002385 | putative family pyridoxal phosphate enzyme protein [Eutypa lata UCREL1] | 69.8 | 2.0E-126 |
| DIP_10002386 | adenylosuccinate synthetase [Magnaporthe oryzae 70-15] | 79.7 | 0.0 |
| DIP_10002387 | glycosyl hydrolase family 16 [Colletotrichum fioriniae PJ7] | 59.5 | 2.0E-122 |
| DIP_10002388 | CTP synthase [Blastomyces dermatitidis SLH14081] | 75.5 | 0.0 |
| DIP_10002389 | NA | | |
| DIP_10002390 | hypothetical protein FGSG_01435 [Fusarium graminearum PH-1] | 75.1 | 2.0E-84 |
| DIP_10002391 | hypothetical protein UCREL1_1171 [Eutypa lata UCREL1] | 52 | 1.0E-81 |
| DIP_10002392 | glutamate synthase [Pestalotiopsis fici W106-1] | 87.6 | 0.0 |
| DIP_10002393 | hypothetical protein PFICI_08391 [Pestalotiopsis fici W106-1] | 63.4 | 0.0 |
| DIP_10002394 | putative g-patch domain-containing protein [Eutypa lata UCREL1] | 59.7 | 0.0 |
| DIP_10002395 | putative dna replication complex gins protein psf3 protein [Eutypa lata UCREL1] | 80.2 | 7.0E-98 |
| DIP_10002396 | hypothetical protein PFICI_08472 [Pestalotiopsis fici W106-1] | 78.7 | 2.0E-132 |
| DIP_10002397 | putative amino-acid permease inda1 protein [Eutypa lata UCREL1] | 71.4 | 0.0 |
| DIP_10002398 | NA | | |
| DIP_10002399 | FAD binding domain protein [Aspergillus clavatus NRRL 1] | 47.2 | 3.0E-175 |
| DIP_10002400 | hypothetical protein PFICI_08385 [Pestalotiopsis fici W106-1] | 51.3 | 0.0 |
| DIP_10002401 | putative spindle pole body component alp14 protein [Eutypa lata UCREL1] | 82.1 | 0.0 |
| DIP_10002402 | WD domain-containing protein [Colletotrichum graminicola M1.001] | 84 | 0.0 |
| DIP_10002403 | glycoside hydrolase [Histoplasma capsulatum H143] | 55.1 | 0.0 |
| DIP_10002404 | putative sit4 phosphatase-associated protein [Eutypa lata UCREL1] | 67.4 | 0.0 |
| DIP_10002405 | putative nadp-dependent leukotriene b4 12-hydroxydehydrogenase protein [Eutypa lata UCREL1] | 62.4 | 1.0E-154 |
| DIP_10002406 | NA | | |
| DIP_10002407 | NA | | |
| DIP_10002408 | NA | | |
| DIP_10002409 | hypothetical protein S7711_06140 [Stachybotrys chartarum IBT 7711] | 53.2 | 3.0E-62 |
| DIP_10002410 | NA | | |
| DIP_10002411 | L-ascorbate oxidase [Fusarium verticillioides 7600] | 38.5 | 2.0E-122 |
| DIP_10002412 | hypothetical protein PFICI_08928 [Pestalotiopsis fici W106-1] | 69.6 | 1.0E-103 |
| DIP_10002413 | NA | | |
| DIP_10002414 | putative actin-like protein arp-6 protein [Eutypa lata UCREL1] | 66.6 | 0.0 |
| DIP_10002415 | hypothetical protein PFICI_09247 [Pestalotiopsis fici W106-1] | 31 | 5.0E-81 |
| DIP_10002416 | hypothetical protein PFICI_09254 [Pestalotiopsis fici W106-1] | 50.5 | 8.0E-59 |
| DIP_10002417 | hypothetical protein PFICI_03087 [Pestalotiopsis fici W106-1] | 45.2 | 5.0E-46 |
| DIP_10002418 | NA | | |
| DIP_10002419 | hypothetical protein PFICI_15254 [Pestalotiopsis fici W106-1] | 52.6 | 5.0E-128 |
| DIP_10002420 | hypothetical protein SAPIO_CDS8016 [Scedosporium apiospermum] | 47.6 | 8.0E-112 |
| DIP_10002421 | putative ring finger domain-containing protein [Eutypa lata UCREL1] | 57.7 | 1.0E-56 |
| DIP_10002422 | putative glycosyltransferase family 34 protein [Eutypa lata UCREL1] | 59.6 | 5.0E-148 |
| DIP_10002423 | NA | | |
| DIP_10002424 | hypothetical protein PFICI_09256 [Pestalotiopsis fici W106-1] | 69.3 | 0.0 |
| DIP_10002425 | putative triose-phosphate transporter protein [Eutypa lata UCREL1] | 68.9 | 0.0 |
| DIP_10002426 | putative cytochrome p450 protein [Eutypa lata UCREL1] | 60.4 | 0.0 |
| DIP_10002427 | hypothetical protein PFICI_04479 [Pestalotiopsis fici W106-1] | 37.6 | 1.0E-46 |
| DIP_10002428 | putative 37s ribosomal protein s5 protein [Eutypa lata UCREL1] | 47.8 | 4.0E-129 |
| DIP_10002429 | NA | | |
| DIP_10002430 | hypothetical protein HIM_09351 [Hirsutella minnesotensis 3608] | 46.3 | 8.0E-169 |
| DIP_10002431 | NA | | |
| DIP_10002432 | hypothetical protein PFICI_08942 [Pestalotiopsis fici W106-1] | 89.6 | 1.0E-44 |
| DIP_10002433 | putative cysteine protease domain-containing protein [Eutypa lata UCREL1] | 37.8 | 5.0E-105 |
| DIP_10002434 | hypothetical protein UCREL1_2595 [Eutypa lata UCREL1] | 42.9 | 2.0E-97 |
| DIP_10002435 | hypothetical protein PFICI_11437 [Pestalotiopsis fici W106-1] | 60.1 | 0.0 |
| DIP_10002436 | NA | | |
| DIP_10002437 | YALI0D03949p [Yarrowia lipolytica] | 74.3 | 9.0E-34 |
| DIP_10002438 | putative transcription initiation factor tfiid subunit protein [Eutypa lata UCREL1] | 65.3 | 2.0E-69 |
| DIP_10002439 | integral membrane protein DUF6 [Colletotrichum fioriniae PJ7] | 37.7 | 4.0E-93 |
| DIP_10002440 | RuvB-like helicase 1 [Pestalotiopsis fici W106-1] | 94.1 | 0.0 |
| DIP_10002441 | hypothetical protein PFICI_11445 [Pestalotiopsis fici W106-1] | 33.6 | 2.0E-59 |
| DIP_10002442 | dynactin [Verticillium dahliae VdLs.17] | 60.1 | 0.0 |
| DIP_10002443 | putative signal recognition particle protein sec65 protein [Eutypa lata UCREL1] | 64.8 | 2.0E-102 |
| DIP_10002444 | NA | | |
| DIP_10002445 | GPI-anchored cell wall beta-1,3-endoglucanase EglC [Gaeumannomyces graminis var. tritici R3-111a-1] | 52.6 | 5.0E-110 |
| DIP_10002446 | putative nf-x1 finger and helicase domain protein [Eutypa lata UCREL1] | 48.5 | 8.0E-45 |
| DIP_10002447 | NA | | |
| DIP_10002448 | NA | | |
| DIP_10002449 | hypothetical protein COCSADRAFT_121392 [Bipolaris sorokiniana ND90Pr] | 53.1 | 1.0E-66 |
| DIP_10002450 | hypothetical protein PFICI_11421 [Pestalotiopsis fici W106-1] | 61.9 | 0.0 |
| DIP_10002451 | putative ribonucleoside-diphosphate reductase large chain protein [Eutypa lata UCREL1] | 88.8 | 0.0 |
| DIP_10002452 | NA | | |
| DIP_10002453 | hypothetical protein MYCTH_2316642 [Myceliophthora thermophila ATCC 42464] | 85.9 | 0.0 |
| DIP_10002454 | NA | | |
| DIP_10002455 | NA | | |
| DIP_10002456 | hypothetical protein CMQ_4298 [Grosmannia clavigera kw1407] | 33.8 | 1.0E-101 |
| DIP_10002457 | hypothetical protein GLRG_08070 [Colletotrichum graminicola M1.001] | 35.3 | 5.0E-58 |
| DIP_10002458 | hypothetical protein UCREL1_2595 [Eutypa lata UCREL1] | 33.9 | 6.0E-59 |
| DIP_10002459 | hypothetical protein CFIO01_02569 [Colletotrichum fioriniae PJ7] | 58.3 | 0.0 |
| DIP_10002460 | putative membrane transporter protein [Eutypa lata UCREL1] | 71.1 | 0.0 |
| DIP_10002461 | hypothetical protein PFICI_06124 [Pestalotiopsis fici W106-1] | 38.1 | 1.0E-75 |
| DIP_10002462 | putative negative regulator of dna transposition protein [Eutypa lata UCREL1] | 65.9 | 0.0 |
| DIP_10002463 | conserved hypothetical protein [Histoplasma capsulatum H143] | 52.2 | 1.0E-56 |
| DIP_10002464 | hypothetical protein S40288_08960 [Stachybotrys chartarum IBT 40288] | 43.9 | 2.0E-75 |
| DIP_10002465 | NA | | |
| DIP_10002466 | hypothetical protein PFICI_08831 [Pestalotiopsis fici W106-1] | 54.2 | 2.0E-137 |
| DIP_10002467 | hypothetical protein PFICI_08830 [Pestalotiopsis fici W106-1] | 55.3 | 1.0E-119 |
| DIP_10002468 | putative duf647 domain-containing protein [Eutypa lata UCREL1] | 52.1 | 8.0E-141 |
| DIP_10002469 | hypothetical protein CGLO_03854 [Colletotrichum gloeosporioides Cg-14] | 51.5 | 6.0E-107 |
| DIP_10002470 | putative diphosphomevalonate decarboxylase protein [Eutypa lata UCREL1] | 78.9 | 0.0 |
| DIP_10002471 | argininosuccinate lyase [Gaeumannomyces graminis var. tritici R3-111a-1] | 80.2 | 0.0 |
| DIP_10002472 | hypothetical protein PFICI_08870 [Pestalotiopsis fici W106-1] | 72.8 | 0.0 |
| DIP_10002473 | hypothetical protein PV08_03170 [Exophiala spinifera] | 50.5 | 4.0E-116 |
| DIP_10002474 | putative manganese lipoxygenase protein [Eutypa lata UCREL1] | 42.7 | 6.0E-148 |
| DIP_10002475 | putative rna polymerase ii mediator complex subunit protein [Eutypa lata UCREL1] | 64.2 | 7.0E-126 |
| DIP_10002476 | putative isoamyl alcohol protein [Togninia minima UCRPA7] | 52 | 0.0 |
| DIP_10002477 | putative integral membrane protein [Togninia minima UCRPA7] | 64.3 | 7.0E-118 |
| DIP_10002478 | peroxisomal catalase [Metarhizium robertsii ARSEF 23] | 34.4 | 1.0E-85 |
| DIP_10002479 | putative der1-like family protein [Eutypa lata UCREL1] | 90.5 | 2.0E-155 |
| DIP_10002480 | histone H3-like centromeric protein cse-4 [Colletotrichum fioriniae PJ7] | 78.3 | 3.0E-62 |
| DIP_10002481 | hypothetical protein PFICI_12620 [Pestalotiopsis fici W106-1] | 59.7 | 3.0E-45 |
| DIP_10002482 | X-Prolyl aminopeptidase [Trichoderma reesei QM6a] | 56.8 | 0.0 |
| DIP_10002483 | hypothetical protein PFICI_12618 [Pestalotiopsis fici W106-1] | 40.7 | 0.0 |
| DIP_10002484 | hypothetical protein PFICI_12629 [Pestalotiopsis fici W106-1] | 62.2 | 1.0E-171 |
| DIP_10002485 | putative 30s ribosomal protein s10 protein [Eutypa lata UCREL1] | 74.6 | 7.0E-106 |
| DIP_10002486 | putative cop9 signalosome complex subunit 5 protein [Eutypa lata UCREL1] | 74.7 | 0.0 |
| DIP_10002487 | hypothetical protein PFICI_12632 [Pestalotiopsis fici W106-1] | 55.4 | 0.0 |
| DIP_10002488 | hypothetical protein UCREL1_4915 [Eutypa lata UCREL1] | 79.2 | 0.0 |
| DIP_10002489 | 60S ribosomal protein L32 [Thielavia terrestris NRRL 8126] | 88.6 | 1.0E-84 |
| DIP_10002490 | pyridine nucleotide-disulfide [Colletotrichum gloeosporioides Nara gc5] | 59.6 | 2.0E-179 |
| DIP_10002491 | NA | | |
| DIP_10002492 | putative mitochondrial 54S ribosomal protein MNP1 [Botrytis cinerea B05.10] | 76.9 | 1.0E-69 |
| DIP_10002493 | hypothetical protein PFICI_05341 [Pestalotiopsis fici W106-1] | 66.7 | 2.0E-164 |
| DIP_10002494 | putative ctd kinase subunit gamma protein [Eutypa lata UCREL1] | 69.7 | 1.0E-122 |
| DIP_10002495 | putative g-protein coupled receptor protein [Eutypa lata UCREL1] | 63.3 | 4.0E-58 |
| DIP_10002496 | hypothetical protein NCU08550 [Neurospora crassa OR74A] | 70.1 | 5.0E-141 |
| DIP_10002497 | NA | | |
| DIP_10002498 | hypothetical protein PFICI_07964 [Pestalotiopsis fici W106-1] | 76.4 | 0.0 |
| DIP_10002499 | hypothetical protein PFICI_07970 [Pestalotiopsis fici W106-1] | 63.3 | 3.0E-84 |
| DIP_10002500 | putative palmitoyltransferase akr1 protein [Eutypa lata UCREL1] | 71.5 | 0.0 |
| DIP_10002501 | hypothetical protein FG05_02887 [Fusarium graminearum] | 61.2 | 5.0E-132 |
| DIP_10002502 | methylcrotonoyl-CoA carboxylase beta chain [Pestalotiopsis fici W106-1] | 85.6 | 0.0 |
| DIP_10002503 | Isovaleryl-CoA dehydrogenase 2 [Pestalotiopsis fici W106-1] | 79.4 | 0.0 |
| DIP_10002504 | dienelactone hydrolase [Beauveria bassiana ARSEF 2860] | 66.3 | 2.0E-120 |
| DIP_10002505 | hypothetical protein PFICI_09883 [Pestalotiopsis fici W106-1] | 35.5 | 1.0E-39 |
| DIP_10002506 | hypothetical protein PFICI_09887 [Pestalotiopsis fici W106-1] | 52.6 | 3.0E-121 |
| DIP_10002507 | NA | | |
| DIP_10002508 | hypothetical protein UCREL1_11739 [Eutypa lata UCREL1] | 55.1 | 2.0E-149 |
| DIP_10002509 | transcriptional regulatory protein GAL4 [Exophiala dermatitidis NIH/UT8656] | 56.5 | 0.0 |
| DIP_10002510 | putative acyltransferase protein [Eutypa lata UCREL1] | 42.1 | 2.0E-132 |
| DIP_10002511 | hypothetical protein S40288_07942 [Stachybotrys chartarum IBT 40288] | 45.6 | 1.0E-63 |
| DIP_10002512 | putative amino acid [Diplodia seriata] | 65.5 | 0.0 |
| DIP_10002513 | hypothetical protein MYCFIDRAFT_202262 [Pseudocercospora fijiensis CIRAD86] | 61 | 0.0 |
| DIP_10002514 | CFEM domain-containing protein [Colletotrichum orbiculare MAFF 240422] | 39.3 | 2.0E-102 |
| DIP_10002515 | hypothetical protein PFICI_13245 [Pestalotiopsis fici W106-1] | 49.5 | 1.0E-130 |
| DIP_10002516 | hypothetical protein S7711_10028 [Stachybotrys chartarum IBT 7711] | 32.4 | 3.0E-10 |
| DIP_10002517 | NA | | |
| DIP_10002518 | hypothetical protein PFICI_13598 [Pestalotiopsis fici W106-1] | 74.1 | 3.0E-162 |
| DIP_10002519 | hypothetical protein PFICI_13564 [Pestalotiopsis fici W106-1] | 41 | 8.0E-111 |
| DIP_10002520 | hypothetical protein PV10_03707 [Exophiala mesophila] | 44.9 | 2.0E-31 |
| DIP_10002521 | putative major facilitator superfamily transporter protein [Eutypa lata UCREL1] | 53.5 | 0.0 |
| DIP_10002522 | putative scp-like extracellular protein [Eutypa lata UCREL1] | 45.1 | 9.0E-76 |
| DIP_10002523 | Serine carboxypeptidase [Aspergillus parasiticus SU-1] | 34.4 | 3.0E-50 |
| DIP_10002524 | hypothetical protein OIDMADRAFT_20934 [Oidiodendron maius Zn] | 40.6 | 2.0E-45 |
| DIP_10002525 | putative nadh-ubiquinone oxidoreductase kda subunit protein [Eutypa lata UCREL1] | 88.8 | 2.0E-94 |
| DIP_10002526 | putative nucleoporin nup44 protein [Eutypa lata UCREL1] | 73.6 | 1.0E-179 |
| DIP_10002527 | hypothetical protein FOXB_07196 [Fusarium oxysporum Fo5176] | 60 | 2.0E-09 |
| DIP_10002528 | hypothetical protein PFICI_13767 [Pestalotiopsis fici W106-1] | 64 | 0.0 |
| DIP_10002529 | hypothetical protein NECHADRAFT_102443 [Nectria haematococca mpVI 77-13-4] | 49.3 | 7.0E-60 |
| DIP_10002530 | putative amino acid transporter protein [Eutypa lata UCREL1] | 70.9 | 0.0 |
| DIP_10002531 | hypothetical protein PFICI_09715 [Pestalotiopsis fici W106-1] | 33.9 | 5.0E-37 |
| DIP_10002532 | NA | | |
| DIP_10002533 | putative osmosensor protein [Eutypa lata UCREL1] | 70.3 | 4.0E-137 |
| DIP_10002534 | NA | | |
| DIP_10002535 | putative rho gtpase activator protein [Eutypa lata UCREL1] | 64.8 | 0.0 |
| DIP_10002536 | putative feruloyl esterase-like protein [Eutypa lata UCREL1] | 48.3 | 7.0E-110 |
| DIP_10002537 | hypothetical protein CGLO_15338 [Colletotrichum gloeosporioides Cg-14] | 62.5 | 3.0E-132 |
| DIP_10002538 | NA | | |
| DIP_10002539 | hypothetical protein CCM_03306 [Cordyceps militaris CM01] | 71.4 | 3.0E-15 |
| DIP_10002540 | putative glycoside hydrolase family 17 protein [Erysiphe necator] | 61 | 6.0E-118 |
| DIP_10002541 | putative vacuolar membrane pq loop repeat protein [Eutypa lata UCREL1] | 63.7 | 1.0E-117 |
| DIP_10002542 | putative nuclease s1 protein [Eutypa lata UCREL1] | 65.6 | 1.0E-142 |
| DIP_10002543 | NA | | |
| DIP_10002544 | hypothetical protein PFICI_09846 [Pestalotiopsis fici W106-1] | 55.3 | 2.0E-83 |
| DIP_10002545 | hypothetical protein PV04_08363 [Capronia semiimmersa] | 36.3 | 1.0E-84 |
| DIP_10002546 | hypothetical protein PFICI_13883 [Pestalotiopsis fici W106-1] | 79.5 | 0.0 |
| DIP_10002547 | NA | | |
| DIP_10002548 | hypothetical protein PFICI_13882 [Pestalotiopsis fici W106-1] | 57.7 | 0.0 |
| DIP_10002549 | NA | | |
| DIP_10002550 | putative amidohydrolase family protein [Eutypa lata UCREL1] | 52.6 | 3.0E-170 |
| DIP_10002551 | NA | | |
| DIP_10002552 | putative transcription factor (snd1 p100) protein [Eutypa lata UCREL1] | 77.7 | 0.0 |
| DIP_10002553 | Cu/Zn superoxide dismutase [Chaetomium globosum CBS 148.51] | 80.9 | 3.0E-84 |
| [illegible] | [illegible] | [illegible] | [illegible] |

| | | | |
|---|---|---|---|
| DIP_10002560 | hypothetical protein UCREL1_6079 [Eutypa lata UCREL1] | 35.5 | 7.0E-81 |
| DIP_10002561 | NA | | |
| DIP_10002562 | hypothetical protein PFICI_10538 [Pestalotiopsis fici W106-1] | 69.9 | 0.0 |
| DIP_10002563 | hypothetical protein HIM_05459 [Hirsutella minnesotensis 3608] | 34.9 | 7.0E-97 |
| DIP_10002564 | hypothetical protein HIM_05458 [Hirsutella minnesotensis 3608] | 35.4 | 3.0E-28 |
| DIP_10002565 | hypothetical protein PFICI_10537 [Pestalotiopsis fici W106-1] | 58.4 | 0.0 |
| DIP_10002566 | putative mitochondrial inner membrane magnesium transporter mrs2 protein [Eutypa lata UCREL1] | 83.3 | 0.0 |
| DIP_10002567 | hypothetical protein MAC_00908 [Metarhizium acridum CQMa 102] | 33.2 | 2.0E-14 |
| DIP_10002568 | hypothetical protein PFICI_10546 [Pestalotiopsis fici W106-1] | 54.1 | 0.0 |
| DIP_10002569 | NA | | |
| DIP_10002570 | hypothetical protein PFICI_10545 [Pestalotiopsis fici W106-1] | 53.5 | 2.0E-55 |
| DIP_10002571 | Pre-rRNA-processing protein IPI1 [Pestalotiopsis fici W106-1] | 61.2 | 5.0E-152 |
| DIP_10002572 | NA | | |
| DIP_10002573 | hypothetical protein PFICI_03258 [Pestalotiopsis fici W106-1] | 37.4 | 6.0E-72 |
| DIP_10002574 | NA | | |
| DIP_10002575 | putative atp-dependent rna helicase ded1 protein [Eutypa lata UCREL1] | 58.7 | 0.0 |
| DIP_10002576 | hypothetical protein PFICI_06405 [Pestalotiopsis fici W106-1] | 51.4 | 1.0E-32 |
| DIP_10002577 | hypothetical protein PFICI_10549 [Pestalotiopsis fici W106-1] | 67 | 8.0E-135 |
| DIP_10002578 | hypothetical protein HMPREF1541_05623 [Cyphellophora europaea CBS 101466] | 52.8 | 2.0E-100 |
| DIP_10002579 | putative atp-dependent rna helicase protein [Eutypa lata UCREL1] | 88.6 | 0.0 |
| DIP_10002580 | hypothetical protein PFICI_10555 [Pestalotiopsis fici W106-1] | 52.7 | 6.0E-57 |
| DIP_10002581 | nadh-ubiquinone oxidoreductase 21 kda [Colletotrichum orbiculare MAFF 240422] | 82.3 | 6.0E-111 |
| DIP_10002582 | putative chromosome segregation in meiosis protein 3 protein [Eutypa lata UCREL1] | 56 | 3.0E-109 |
| DIP_10002583 | putative acid protease protein [Eutypa lata UCREL1] | 49.5 | 4.0E-152 |
| DIP_10002584 | putative polyketide synthase protein [Eutypa lata UCREL1] | 47.9 | 0.0 |
| DIP_10002585 | putative polyketide synthase protein [Eutypa lata UCREL1] | 70.2 | 0.0 |
| DIP_10002586 | putative transporter C3H1.06c [Ceratocystis platani] | 44.9 | 7.0E-51 |
| DIP_10002587 | S-adenosyl-L-methionine-dependent methyltransferase [Glarea lozoyensis ATCC 20868] | 31.8 | 9.0E-56 |
| DIP_10002588 | hypothetical protein PFICI_13787 [Pestalotiopsis fici W106-1] | 61.4 | 1.0E-71 |
| DIP_10002589 | hypothetical protein PFICI_09905 [Pestalotiopsis fici W106-1] | 67.8 | 5.0E-53 |
| DIP_10002590 | hypothetical protein PFICI_09906 [Pestalotiopsis fici W106-1] | 50.3 | 0.0 |
| DIP_10002591 | NA | | |
| DIP_10002592 | NA | | |
| DIP_10002593 | putative protein efr3 [Diaporthe ampelina] | 51.3 | 0.0 |
| DIP_10002594 | hypothetical protein PFICI_10667 [Pestalotiopsis fici W106-1] | 64.1 | 4.0E-158 |
| DIP_10002595 | putative acyl- thioesterase ii protein [Eutypa lata UCREL1] | 63.8 | 9.0E-156 |
| DIP_10002596 | hypothetical protein PFICI_10591 [Pestalotiopsis fici W106-1] | 66.5 | 0.0 |
| DIP_10002597 | putative phosphopantetheinyl transferase protein [Eutypa lata UCREL1] | 52.4 | 1.0E-102 |
| DIP_10002598 | putative rna-binding protein [Eutypa lata UCREL1] | 71.2 | 0.0 |
| DIP_10002599 | NA | | |
| DIP_10002600 | ATP synthase subunit d [Pestalotiopsis fici W106-1] | 88.4 | 6.0E-110 |
| DIP_10002601 | NAC domain-containing protein [Colletotrichum fioriniae PJ7] | 83.2 | 2.0E-98 |
| DIP_10002602 | hypothetical protein OIDMADRAFT_39553 [Oidiodendron maius Zn] | 56.5 | 1.0E-139 |
| DIP_10002603 | putative ankyrin repeat protein [Diaporthe ampelina] | 49.2 | 0.0 |
| DIP_10002604 | hypothetical protein PFICI_09903 [Pestalotiopsis fici W106-1] | 53.2 | 0.0 |
| DIP_10002605 | molybdopterin synthase catalytic subunit, partial [Trichoderma harzianum] | 49.1 | 1.0E-51 |
| DIP_10002606 | small oligopeptide opt family [Colletotrichum orbiculare MAFF 240422] | 67.7 | 0.0 |
| DIP_10002607 | hypothetical protein ATEG_10080 [Aspergillus terreus NIH2624] | 35 | 0.0 |
| DIP_10002608 | hypothetical protein ARAM_002428 [Aspergillus rambellii] | 46.3 | 0.0 |
| DIP_10002609 | NA | | |
| DIP_10002610 | putative duf255 domain protein [Eutypa lata UCREL1] | 55.9 | 0.0 |
| DIP_10002611 | hypothetical protein PFICI_13793 [Pestalotiopsis fici W106-1] | 67.2 | 0.0 |
| DIP_10002612 | RNA binding domain protein [Cordyceps militaris CM01] | 66.4 | 1.0E-101 |
| DIP_10002613 | putative svp1-like protein 2 protein [Eutypa lata UCREL1] | 84.8 | 3.0E-81 |
| DIP_10002614 | putative cell division control protein 11 protein [Eutypa lata UCREL1] | 93.2 | 0.0 |
| DIP_10002615 | putative transmembrane bax inhibitor motif-containing protein [Eutypa lata UCREL1] | 80.4 | 4.0E-170 |
| DIP_10002616 | hypothetical protein MGG_14324 [Magnaporthe oryzae 70-15] | 44.8 | 0.0 |
| DIP_10002617 | hypothetical protein PFICI_13788 [Pestalotiopsis fici W106-1] | 65.7 | 0.0 |
| DIP_10002618 | hypothetical protein V500_06418 [Pseudogymnoascus pannorum VKM F-4518 (FW-2643)] | 48.4 | 9.0E-90 |
| DIP_10002619 | NA | | |
| DIP_10002620 | hypothetical protein PFICI_14406 [Pestalotiopsis fici W106-1] | 45 | 1.0E-18 |
| DIP_10002621 | hypothetical protein FOWG_12874 [Fusarium oxysporum f. sp. lycopersici MN25] | 51.6 | 1.0E-35 |
| DIP_10002622 | putative endo-beta- -glucanase protein [Togninia minima UCRPA7] | 56.1 | 0.0 |
| DIP_10002623 | hypothetical protein AOCH_004893 [Aspergillus ochraceoroseus] | 41.4 | 4.0E-07 |
| DIP_10002624 | hypothetical protein TI39_contig4119g00028 [Zymoseptoria brevis] | 70.5 | 1.0E-109 |
| DIP_10002625 | NA | | |
| DIP_10002626 | hypothetical protein UCREL1_2731 [Eutypa lata UCREL1] | 32.4 | 1.0E-37 |
| DIP_10002627 | putative dshct domain-containing protein [Eutypa lata UCREL1] | 80 | 0.0 |
| DIP_10002628 | hypothetical protein CGLO_01710 [Colletotrichum gloeosporioides Cg-14] | 63.1 | 0.0 |
| DIP_10002629 | putative nuclear envelope protein [Eutypa lata UCREL1] | 44.3 | 3.0E-115 |
| DIP_10002630 | putative transcription factor iws1 protein [Eutypa lata UCREL1] | 69 | 0.0 |
| DIP_10002631 | hypothetical protein PFICI_13321 [Pestalotiopsis fici W106-1] | 74.3 | 0.0 |
| DIP_10002632 | hypothetical protein NECHADRAFT_101789 [Nectria haematococca mpVI 77-13-4] | 32.7 | 4.0E-76 |
| DIP_10002633 | hypothetical protein PFICI_03239 [Pestalotiopsis fici W106-1] | 66.4 | 4.0E-125 |
| DIP_10002634 | hypothetical protein PFICI_07037 [Pestalotiopsis fici W106-1] | 57.3 | 4.0E-129 |
| DIP_10002635 | putative c2h2 type zinc finger containing protein [Eutypa lata UCREL1] | 71 | 2.0E-72 |
| DIP_10002636 | Acetyl esterase [Scedosporium apiospermum] | 39.6 | 1.0E-81 |
| DIP_10002637 | hypothetical protein FOMG_07516 [Fusarium oxysporum f. sp. melonis 26406] | 47.8 | 9.0E-96 |
| DIP_10002638 | hypothetical protein PFICI_06771 [Pestalotiopsis fici W106-1] | 43.6 | 2.0E-113 |
| DIP_10002639 | hypothetical protein S40288_01010 [Stachybotrys chartarum IBT 40288] | 39.6 | 4.0E-98 |
| DIP_10002640 | NA | | |
| DIP_10002641 | NA | | |
| DIP_10002642 | putative integral membrane protein [Diaporthe ampelina] | 41.8 | 2.0E-60 |
| DIP_10002643 | hypothetical protein PFICI_06861 [Pestalotiopsis fici W106-1] | 53.7 | 0.0 |
| DIP_10002644 | putative endo-beta- - protein [Eutypa lata UCREL1] | 51.4 | 2.0E-147 |
| DIP_10002645 | hypothetical protein PFICI_13430 [Pestalotiopsis fici W106-1] | 43.1 | 0.0 |
| DIP_10002646 | putative snf7 family protein [Eutypa lata UCREL1] | 85.7 | 1.0E-124 |
| DIP_10002647 | Similar to Protein flp; acc. no. Q8NUZ4 [Pyronema omphalodes CBS 100304] | 30.7 | 5.0E-19 |
| DIP_10002648 | NA | | |
| DIP_10002649 | NA | | |
| DIP_10002650 | putative transcriptional activator hac1 protein [Eutypa lata UCREL1] | 60.3 | 7.0E-150 |
| DIP_10002651 | NA | | |
| DIP_10002652 | plac8 family protein [Ophiostoma piceae UAMH 11346] | 52.7 | 1.0E-53 |
| DIP_10002653 | putative signal peptide peptidase protein [Eutypa lata UCREL1] | 59.8 | 0.0 |
| DIP_10002654 | peptidyl-prolyl cis-trans isomerase-like 1 [Cladophialophora psammophila CBS 110553] | 74.5 | 5.0E-93 |
| DIP_10002655 | putative transcriptional activator protein [Eutypa lata UCREL1] | 71.3 | 3.0E-180 |
| DIP_10002656 | putative gcn5-related n acetyltransferase protein [Eutypa lata UCREL1] | 52.3 | 1.0E-101 |
| DIP_10002657 | Glucokinase [Pestalotiopsis fici W106-1] | 81.6 | 0.0 |
| DIP_10002658 | NA | | |
| DIP_10002659 | predicted protein [Uncinocarpus reesii 1704] | 30.2 | 4.0E-17 |
| DIP_10002660 | NA | | |
| DIP_10002661 | hypothetical protein PFICI_09746 [Pestalotiopsis fici W106-1] | 42.9 | 5.0E-23 |
| DIP_10002662 | hypothetical protein PFICI_10576 [Pestalotiopsis fici W106-1] | 76.7 | 0.0 |
| DIP_10002663 | hypothetical protein PFICI_10575 [Pestalotiopsis fici W106-1] | 35.5 | 2.0E-66 |
| DIP_10002664 | putative bactericidal permeability-increasing protein [Eutypa lata UCREL1] | 71.5 | 0.0 |
| DIP_10002665 | putative cytochrome c oxidase assembly protein [Eutypa lata UCREL1] | 80.5 | 8.0E-73 |
| DIP_10002666 | putative sterol esterase tgl1 protein [Eutypa lata UCREL1] | 82.3 | 0.0 |
| DIP_10002667 | NA | | |
| DIP_10002668 | RecName: Full=Transaldolase; Short=FoTal [Fusarium oxysporum f. sp. lycopersici 4287] | 82 | 0.0 |
| DIP_10002669 | NA | | |
| DIP_10002670 | hypothetical protein SAPIO_CDS6360 [Scedosporium apiospermum] | 33.1 | 3.0E-71 |
| DIP_10002671 | hypothetical protein EPUS_08997 [Endocarpon pusillum Z07020] | 42.9 | 3.0E-149 |
| DIP_10002672 | hypothetical protein PFICI_14060 [Pestalotiopsis fici W106-1] | 41.4 | 1.0E-81 |
| DIP_10002673 | hypothetical protein PFICI_09764 [Pestalotiopsis fici W106-1] | 47.8 | 0.0 |
| DIP_10002674 | NA | | |
| DIP_10002675 | hypothetical protein FOXB_12378 [Fusarium oxysporum Fo5176] | 40.8 | 4.0E-56 |
| DIP_10002676 | putative ankyrin repeat and zinc finger domain-containing protein [Colletotrichum sublineola] | 62.9 | 0.0 |
| DIP_10002677 | NA | | |
| DIP_10002678 | NA | | |
| DIP_10002679 | hypothetical protein PFICI_10530 [Pestalotiopsis fici W106-1] | 44.4 | 9.0E-154 |
| DIP_10002680 | Glucosamine 6-phosphate N-acetyltransferase [Ceratocystis platani] | 75.9 | 8.0E-97 |
| DIP_10002681 | hypothetical protein [Podospora anserina S mat+] | 64 | 0.0 |
| DIP_10002682 | putative 2-deoxy-d-gluconate 3-dehydrogenase protein [Eutypa lata UCREL1] | 82.5 | 2.0E-137 |
| DIP_10002683 | hypothetical protein UCREL1_3959 [Eutypa lata UCREL1] | 53 | 7.0E-102 |
| DIP_10002684 | NA | | |
| DIP_10002685 | putative sorbitol dehydrogenase protein [Eutypa lata UCREL1] | 67.6 | 0.0 |
| DIP_10002686 | putative d-galacturonic acid reductase protein [Eutypa lata UCREL1] | 75.9 | 4.0E-177 |
| DIP_10002687 | hypothetical protein PFICI_07058 [Pestalotiopsis fici W106-1] | 85.5 | 0.0 |
| DIP_10002688 | NA | | |
| DIP_10002689 | hypothetical protein CGGC5_12299 [Colletotrichum gloeosporioides Nara gc5] | 30.8 | 8.0E-40 |
| DIP_10002690 | NA | | |
| DIP_10002691 | hypothetical protein W97_05792 [Coniosporium apollinis CBS 100218] | 30.7 | 4.0E-50 |
| DIP_10002692 | NA | | |
| DIP_10002693 | NA | | |
| DIP_10002694 | NA | | |
| DIP_10002695 | hypothetical protein V496_09577 [Pseudogymnoascus pannorum VKM F-4515 (FW-2607)] | 30.9 | 9.0E-13 |
| DIP_10002696 | NA | | |
| DIP_10002697 | hypothetical protein CGLO_00651 [Colletotrichum gloeosporioides Cg-14] | 38.9 | 7.0E-95 |
| DIP_10002698 | hypothetical protein S7711_00583 [Stachybotrys chartarum IBT 7711] | 84.9 | 0.0 |
| DIP_10002699 | putative acetylglutamate protein [Eutypa lata UCREL1] | 84.5 | 0.0 |
| DIP_10002700 | hypothetical protein NECHADRAFT_104309 [Nectria haematococca mpVI 77-13-4] | 58.6 | 6.0E-121 |
| DIP_10002701 | NA | | |
| DIP_10002702 | NA | | |
| DIP_10002703 | NA | | |
| DIP_10002704 | NA | | |
| DIP_10002705 | putative serine carboxypeptidase s28 protein [Neofusicoccum parvum UCRNP2] | 45 | 1.0E-157 |
| DIP_10002706 | putative major facilitator superfamily transporter protein [Eutypa lata UCREL1] | 81 | 0.0 |
| DIP_10002707 | hypothetical protein PFICI_10665 [Pestalotiopsis fici W106-1] | 51.7 | 0.0 |
| DIP_10002708 | hypothetical protein CFIO01_02518 [Colletotrichum fioriniae PJ7] | 58.5 | 7.0E-70 |
| DIP_10002709 | pirin domain-containing protein [Colletotrichum orbiculare MAFF 240422] | 81.1 | 3.0E-176 |
| DIP_10002710 | amidophosphoribosyltransferase [Talaromyces marneffei ATCC 18224] | 73.4 | 0.0 |
| DIP_10002711 | NA | | |
| DIP_10002712 | hypothetical protein V495_08491 [Pseudogymnoascus pannorum VKM F-4514 (FW-929)] | 54.8 | 4.0E-116 |
| DIP_10002713 | hypothetical protein UCREL1_3939 [Eutypa lata UCREL1] | 42.8 | 2.0E-59 |
| DIP_10002714 | NA | | |
| DIP_10002715 | hypothetical protein PFICI_10526 [Pestalotiopsis fici W106-1] | 61.6 | 3.0E-170 |
| DIP_10002716 | NA | | |
| DIP_10002717 | putative arylsulfotransferase-like protein [Eutypa lata UCREL1] | 57.6 | 0.0 |
| DIP_10002718 | hypothetical protein UCREL1_3944 [Eutypa lata UCREL1] | 30.9 | 3.0E-84 |
| DIP_10002719 | putative 1-aminocyclopropane-1-carboxylate deaminase [Pestalotiopsis fici W106-1] | 66.3 | 0.0 |
| DIP_10002720 | protein of unknown function DUF861, cupin-3 [Penicillium italicum] | 82.9 | 7.0E-81 |
| DIP_10002721 | glutamyl-tRNA(Gln) amidotransferase [Colletotrichum fioriniae PJ7] | 43.2 | 2.0E-67 |
| DIP_10002722 | NA | | |
| DIP_10002723 | hypothetical protein T310_4814 [Rasamsonia emersonii CBS 393.64] | 50.5 | 0.0 |
| DIP_10002724 | hypothetical protein FOVG_07141 [Fusarium oxysporum f. sp. pisi HDV247] | 43.2 | 1.0E-142 |
| DIP_10002725 | hypothetical protein A1O5_13374 [Cladophialophora psammophila CBS 110553] | 65.2 | 0.0 |
| DIP_10002726 | NA | | |
| DIP_10002727 | hypothetical protein FAVG1_02665 [Fusarium avenaceum] | 50.8 | 4.0E-151 |
| DIP_10002728 | putative inosine uridine-preferring nucleoside hydrolase protein [Togninia minima UCRPA7] | 76.2 | 1.0E-175 |
| DIP_10002729 | hypothetical protein UCRPA7_116 [Togninia minima UCRPA7] | 82.3 | 0.0 |
| DIP_10002730 | NA | | |
| DIP_10002731 | NA | | |
| DIP_10002732 | NA | | |
| DIP_10002733 | NA | | |
| DIP_10002734 | NA | | |
| DIP_10002735 | NA | | |
| DIP_10002736 | hypothetical protein PFICI_10532 [Pestalotiopsis fici W106-1] | 67.2 | 0.0 |
| DIP_10002737 | NA | | |
| DIP_10002738 | hypothetical protein HMPREF1624_04634 [Sporothrix schenckii ATCC 58251] | 57.8 | 3.0E-18 |
| DIP_10002739 | hypothetical protein PFICI_12676 [Pestalotiopsis fici W106-1] | 40.7 | 3.0E-78 |
| DIP_10002740 | hypothetical protein PTT_14439 [Pyrenophora teres f. teres 0-1] | 54.6 | 1.0E-109 |
| DIP_10002741 | putative serologically defined colon cancer antigen 1 protein [Eutypa lata UCREL1] | 62.7 | 0.0 |
| DIP_10002742 | hypothetical protein PFICI_10528 [Pestalotiopsis fici W106-1] | 54.1 | 7.0E-136 |
| DIP_10002743 | hypothetical protein PFICI_07065 [Pestalotiopsis fici W106-1] | 72.3 | 0.0 |
| DIP_10002744 | hypothetical protein PTT_15219 [Pyrenophora teres f. teres 0-1] | 44.4 | 4.0E-169 |
| DIP_10002745 | hypothetical protein PFICI_07209 [Pestalotiopsis fici W106-1] | 35.5 | 3.0E-92 |
| DIP_10002746 | dienelactone hydrolase [Colletotrichum higginsianum] | 57.2 | 3.0E-173 |
| DIP_10002747 | hypothetical protein Cob_13179 [Colletotrichum orbiculare MAFF 240422] | 53.9 | 3.0E-61 |
| DIP_10002748 | putative sh3 domain signaling protein [Eutypa lata UCREL1] | 58.1 | 6.0E-123 |
| DIP_10002749 | hypothetical protein CGLO_12737 [Colletotrichum gloeosporioides Cg-14] | 46.6 | 2.0E-104 |
| DIP_10002750 | MFS quinate transporter [Colletotrichum gloeosporioides Nara gc5] | 73.4 | 0.0 |
| DIP_10002751 | C6 transcription factor [Colletotrichum gloeosporioides Nara gc5] | 39.9 | 1.0E-163 |
| DIP_10002752 | NA | | |
| DIP_10002753 | NA | | |
| DIP_10002754 | hypothetical protein PFICI_09907 [Pestalotiopsis fici W106-1] | 52.7 | 0.0 |
| DIP_10002755 | hypothetical protein CFIO01_06542 [Colletotrichum fioriniae PJ7] | 77.1 | 4.0E-51 |
| DIP_10002756 | NA | | |
| DIP_10002757 | GTPase-activating protein [Pestalotiopsis fici W106-1] | 74.1 | 0.0 |
| DIP_10002758 | putative camp-dependent protein kinase catalytic subunit protein [Eutypa lata UCREL1] | 80.3 | 0.0 |
| DIP_10002759 | putative epl1 protein [Eutypa lata UCREL1] | 66 | 2.0E-61 |
| DIP_10002760 | hypothetical protein PFICI_14082 [Pestalotiopsis fici W106-1] | 50.5 | 0.0 |
| DIP_10002761 | hypothetical protein PFICI_14081 [Pestalotiopsis fici W106-1] | 47.1 | 4.0E-21 |
| DIP_10002762 | hypothetical protein CMQ_4996 [Grosmannia clavigera kw1407] | 52.9 | 6.0E-16 |
| DIP_10002763 | putative sterol 24-c-methyltransferase protein [Eutypa lata UCREL1] | 86.8 | 0.0 |
| DIP_10002764 | putative pyruvate dehydrogenase e1 component subunit beta protein [Eutypa lata UCREL1] | 78.6 | 0.0 |
| DIP_10002765 | NA | | |
| DIP_10002766 | NA | | |
| DIP_10002767 | hypothetical protein MYCFIDRAFT_197825 [Pseudocercospora fijiensis CIRAD86] | 45.6 | 1.0E-44 |
| DIP_10002768 | putative prion formation protein 1 protein [Eutypa lata UCREL1] | 83.6 | 0.0 |
| DIP_10002769 | putative upf0311 protein [Eutypa lata UCREL1] | 34.2 | 4.0E-40 |
| DIP_10002770 | NA | | |
| DIP_10002771 | putative initiation-specific alpha- -mannosyltransferase protein [Eutypa lata UCREL1] | 72.1 | 0.0 |
| DIP_10002772 | putative short-chain-fatty-acid- ligase protein [Eutypa lata UCREL1] | 58.7 | 0.0 |
| DIP_10002773 | putative leucine rich repeat domain protein [Eutypa lata UCREL1] | 62 | 6.0E-109 |
| DIP_10002774 | putative phosducin family protein [Eutypa lata UCREL1] | 86.8 | 2.0E-150 |
| DIP_10002775 | 40S ribosomal protein S27 [Verticillium alfalfae VaMs.102] | 97.6 | 2.0E-54 |
| DIP_10002776 | putative ribonucleotide reductase inhibitor protein [Eutypa lata UCREL1] | 43.4 | 6.0E-61 |
| DIP_10002777 | hypothetical protein FOXB_17253 [Fusarium oxysporum Fo5176] | 52.1 | 1.0E-67 |
| DIP_10002778 | hypothetical protein FPSE_04849 [Fusarium pseudograminearum CS3096] | 32.7 | 7.0E-47 |
| DIP_10002779 | predicted protein [Trichoderma reesei QM6a] | 44.7 | 2.0E-93 |
| DIP_10002780 | NA | | |
| DIP_10002781 | hypothetical protein CH063_01843, partial [Colletotrichum higginsianum] | 31.5 | 2.0E-29 |
| DIP_10002782 | NA | | |
| DIP_10002783 | hypothetical protein ANO11243_085510 [fungal sp. No.11243] | 43.3 | 1.0E-142 |
| DIP_10002784 | NAD dependent epimerase/dehydratase [Rasamsonia emersonii CBS 393.64] | 35 | 1.0E-45 |
| DIP_10002785 | NA | | |
| DIP_10002786 | putative serine protease protein [Eutypa lata UCREL1] | 79.1 | 0.0 |
| DIP_10002787 | hypothetical protein THITE_2121132 [Thielavia terrestris NRRL 8126] | 37.6 | 3.0E-32 |
| DIP_10002788 | hypothetical protein THITE_38252 [Thielavia terrestris NRRL 8126] | 51.2 | 0.0 |
| DIP_10002789 | hypothetical protein PFICI_02871 [Pestalotiopsis fici W106-1] | 62.8 | 0.0 |
| DIP_10002790 | hypothetical protein PDE_09148 [Penicillium oxalicum 114-2] | 91.8 | 1.0E-61 |
| DIP_10002791 | Telomerase-binding protein EST1A [Fusarium oxysporum f. sp. cubense race 1] | 41.5 | 3.0E-134 |
| DIP_10002792 | hypothetical protein PFICI_02870 [Pestalotiopsis fici W106-1] | 42.1 | 3.0E-91 |
| DIP_10002793 | hypothetical protein THITE_2085426 [Thielavia terrestris NRRL 8126] | 50.6 | 3.0E-77 |
| DIP_10002794 | uncharacterized protein FFUJ_09831 [Fusarium fujikuroi IMI 58289] | 31.1 | 3.0E-09 |
| DIP_10002795 | NA | | |
| DIP_10002796 | NA | | |
| DIP_10002797 | NA | | |
| DIP_10002798 | putative -like family protein [Eutypa lata UCREL1] | 51.9 | 1.0E-55 |
| DIP_10002799 | NA | | |
| DIP_10002800 | NA | | |
| DIP_10002801 | hypothetical protein PFICI_02873 [Pestalotiopsis fici W106-1] | 45.6 | 2.0E-46 |
| DIP_10002802 | NA | | |
| DIP_10002803 | hypothetical protein MYCTH_2313956 [Myceliophthora thermophila ATCC 42464] | 69.2 | 5.0E-33 |
| DIP_10002804 | putative dolichyl-di-phosphooligosaccharide-protein glycotransferase protein [Eutypa lata UCREL1] | 69.6 | 0.0 |
| DIP_10002805 | putative pre-mrna-splicing factor isy1 protein [Togninia minima UCRPA7] | 78.8 | 9.0E-137 |
| DIP_10002806 | hypothetical protein GGTG_11714 [Gaeumannomyces graminis var. tritici R3-111a-1] | 55.8 | 0.0 |
| DIP_10002807 | Glutamyl-tRNA(Gln) amidotransferase subunit A [Pestalotiopsis fici W106-1] | 62.2 | 0.0 |
| DIP_10002808 | putative pep38 family protein [Togninia minima UCRPA7] | 65.7 | 4.0E-105 |
| DIP_10002809 | NA | | |
| DIP_10002810 | hypothetical protein BC1G_09861 [Botrytis cinerea B05.10] | 70.3 | 0.0 |
| DIP_10002811 | putative wd repeat-containing protein [Eutypa lata UCREL1] | 53.9 | 0.0 |
| DIP_10002812 | putative ubiquitin conjugating enzyme protein [Eutypa lata UCREL1] | 92 | 2.0E-100 |
| DIP_10002813 | hypothetical protein PFICI_03016 [Pestalotiopsis fici W106-1] | 73.9 | 0.0 |
| DIP_10002814 | NA | | |
| DIP_10002815 | hypothetical protein PFICI_03015 [Pestalotiopsis fici W106-1] | 47 | 8.0E-50 |
| DIP_10002816 | hypothetical protein PFICI_02998 [Pestalotiopsis fici W106-1] | 73.5 | 0.0 |
| DIP_10002817 | putative nuclear protein qri2 protein [Eutypa lata UCREL1] | 58.5 | 5.0E-162 |
| DIP_10002818 | NA | | |
| DIP_10002819 | hypothetical protein UCREL1_3309 [Eutypa lata UCREL1] | 65.9 | 2.0E-138 |
| DIP_10002820 | NA | | |
| DIP_10002821 | NA | | |
| DIP_10002822 | NA | | |
| DIP_10002823 | hypothetical protein PFICI_03005 [Pestalotiopsis fici W106-1] | 73.2 | 0.0 |
| DIP_10002824 | hypothetical protein FPSE_06454 [Fusarium pseudograminearum CS3096] | 90 | 5.0E-100 |
| DIP_10002825 | hypothetical protein NECHADRAFT_99586 [Nectria haematococca mpVI 77-13-4] | 41.4 | 1.0E-31 |
| DIP_10002826 | hypothetical protein UCREL1_3314 [Eutypa lata UCREL1] | 41 | 3.0E-45 |
| DIP_10002827 | putative fasciclin domain family protein [Eutypa lata UCREL1] | 53.7 | 5.0E-156 |
| DIP_10002828 | hypothetical protein MYCTH_2297728 [Myceliophthora thermophila ATCC 42464] | 40.2 | 1.0E-64 |
| DIP_10002829 | putative pal1 cell morphology protein [Eutypa lata UCREL1] | 54.1 | 6.0E-126 |
| DIP_10002830 | NA | | |
| DIP_10002831 | hypothetical protein PFICI_03032 [Pestalotiopsis fici W106-1] | 52.5 | 0.0 |
| DIP_10002832 | putative respiratory complex assembly protein rmp1 protein [Eutypa lata UCREL1] | 49.5 | 1.0E-129 |
| DIP_10002833 | hypothetical protein PFICI_03030 [Pestalotiopsis fici W106-1] | 78.3 | 0.0 |
| DIP_10002834 | hypothetical protein PFICI_03029 [Pestalotiopsis fici W106-1] | 60.2 | 2.0E-111 |
| DIP_10002835 | hypothetical protein CTHT_0008200 [Chaetomium thermophilum var. thermophilum DSM 1495] | 34.8 | 1.0E-12 |
| DIP_10002836 | NA | | |
| DIP_10002837 | NA | | |
| DIP_10002838 | hypothetical protein CH063_16048 [Colletotrichum higginsianum] | 48 | 4.0E-71 |
| DIP_10002839 | hypothetical protein VHEMI03690 [Torrubiella hemipterigena] | 40.9 | 5.0E-87 |
| DIP_10002840 | hypothetical protein PFICI_02217 [Pestalotiopsis fici W106-1] | 62.1 | 0.0 |
| DIP_10002841 | hypothetical protein HIM_04699 [Hirsutella minnesotensis 3608] | 62.7 | 3.0E-44 |
| DIP_10002842 | ribosomal protein S14p/S29e [Colletotrichum graminicola M1.001] | 77.1 | 2.0E-62 |
| DIP_10002843 | hypothetical protein HIM_05402 [Hirsutella minnesotensis 3608] | 33.8 | 1.0E-55 |
| DIP_10002844 | hypothetical protein THITE_2112873 [Thielavia terrestris NRRL 8126] | 30.4 | 2.0E-24 |
| DIP_10002845 | putative wd repeat-containing protein [Eutypa lata UCREL1] | 64.4 | 0.0 |
| DIP_10002846 | hypothetical protein PFICI_02221 [Pestalotiopsis fici W106-1] | 60.4 | 0.0 |
| DIP_10002847 | hypothetical protein UCREL1_7612 [Eutypa lata UCREL1] | 54.3 | 2.0E-117 |
| DIP_10002848 | hypothetical protein CFIO01_02910 [Colletotrichum fioriniae PJ7] | 54.9 | 3.0E-113 |
| DIP_10002849 | hypothetical protein MGG_02727 [Magnaporthe oryzae 70-15] | 76.1 | 2.0E-113 |
| DIP_10002850 | RNA recognition domain-containing protein [Colletotrichum fioriniae PJ7] | 50.9 | 1.0E-146 |
| DIP_10002851 | hypothetical protein CTHT_0000820 [Chaetomium thermophilum var. thermophilum DSM 1495] | 36.3 | 2.0E-58 |
| DIP_10002852 | hypothetical protein Z518_05088 [Rhinocladiella mackenziei CBS 650.93] | 69.9 | 6.0E-75 |
| DIP_10002853 | hypothetical protein W97_06966 [Coniosporium apollinis CBS 100218] | 39.5 | 4.0E-89 |
| DIP_10002854 | hypothetical protein PFICI_03036 [Pestalotiopsis fici W106-1] | 59.6 | 8.0E-145 |
| DIP_10002855 | hypothetical protein MYCTH_75197 [Myceliophthora thermophila ATCC 42464] | 41.6 | 6.0E-68 |
| DIP_10002856 | putative coiled-coil domain-containing protein [Eutypa lata UCREL1] | 48.5 | 8.0E-66 |
| DIP_10002857 | hypothetical protein PFICI_02815 [Pestalotiopsis fici W106-1] | 46 | 1.0E-99 |
| DIP_10002858 | NA | | |
| DIP_10002859 | putative trna-specific adenosine deaminase-like protein [Eutypa lata UCREL1] | 56.2 | 1.0E-106 |
| DIP_10002860 | putative set domain-containing protein [Eutypa lata UCREL1] | 55.2 | 0.0 |
| DIP_10002861 | hypothetical protein PFICI_06835 [Pestalotiopsis fici W106-1] | 32.2 | 2.0E-13 |
| DIP_10002862 | ibr domain-containing protein [Colletotrichum gloeosporioides Nara gc5] | 31.8 | 1.0E-11 |
| DIP_10002863 | hypothetical protein CGLO_01249 [Colletotrichum gloeosporioides Cg-14] | 38.2 | 4.0E-10 |
| DIP_10002864 | NA | | |
| DIP_10002865 | NA | | |
| DIP_10002866 | hypothetical protein NEUTE1DRAFT_118559 [Neurospora tetrasperma FGSC 2508] | 49.6 | 5.0E-82 |
| DIP_10002867 | phosphoglycerate mutase [Colletotrichum fioriniae PJ7] | 76.5 | 2.0E-134 |
| DIP_10002868 | NA | | |
| DIP_10002869 | hypothetical protein UCREL1_658 [Eutypa lata UCREL1] | 54.1 | 2.0E-28 |
| DIP_10002870 | putative dicarboxylic amino acid permease protein [Eutypa lata UCREL1] | 72.6 | 0.0 |
| DIP_10002871 | adp-ribose pyrophosphatase like protein [Zymoseptoria brevis] | 72.7 | 7.0E-96 |
| DIP_10002872 | putative ubiquitin carboxyl-terminal hydrolase 6 protein [Togninia minima UCRPA7] | 75.1 | 0.0 |
| DIP_10002873 | conserved hypothetical protein [Sclerotinia sclerotiorum 1980] | 94.7 | 5.0E-115 |
| DIP_10002874 | hypothetical protein PFICI_06583 [Pestalotiopsis fici W106-1] | 58.4 | 0.0 |
| DIP_10002875 | putative protein family protein [Eutypa lata UCREL1] | 76.5 | 3.0E-169 |
| DIP_10002876 | NA | | |
| DIP_10002877 | hypothetical protein CH063_02101 [Colletotrichum higginsianum] | 34.1 | 2.0E-98 |
| DIP_10002878 | putative sulfate adenylyltransferase protein [Eutypa lata UCREL1] | 86.4 | 0.0 |
| DIP_10002879 | hypothetical protein PFICI_09711 [Pestalotiopsis fici W106-1] | 55 | 0.0 |
| DIP_10002880 | subtilisin-like serine protease precursor [Metarhizium robertsii ARSEF 23] | 45.8 | 5.0E-103 |
| DIP_10002881 | NA | | |
| DIP_10002882 | hypothetical protein PFICI_09732 [Pestalotiopsis fici W106-1] | 53 | 6.0E-163 |
| DIP_10002883 | hypothetical protein PFICI_09731 [Pestalotiopsis fici W106-1] | 83 | 0.0 |
| DIP_10002884 | hypothetical protein PFICI_09730 [Pestalotiopsis fici W106-1] | 49.7 | 7.0E-75 |
| DIP_10002885 | putative importin-beta domain-containing protein [Eutypa lata UCREL1] | 85.7 | 0.0 |
| DIP_10002886 | imp-specific 5'-nucleotidase-like protein [Chaetomium thermophilum var. thermophilum DSM 1495] | 78.5 | 0.0 |
| DIP_10002887 | hypothetical protein PFICI_02844 [Pestalotiopsis fici W106-1] | 33.2 | 4.0E-37 |
| DIP_10002888 | hypothetical protein UCREL1_8169 [Eutypa lata UCREL1] | 49.1 | 6.0E-99 |
| DIP_10002889 | putative serine threonine-protein kinase hal4 protein [Eutypa lata UCREL1] | 74.2 | 0.0 |
| DIP_10002890 | putative glycosyltransferase family 39 protein [Eutypa lata UCREL1] | 77.1 | 0.0 |
| DIP_10002891 | hypothetical protein PFICI_02829 [Pestalotiopsis fici W106-1] | 71.5 | 0.0 |
| DIP_10002892 | hypothetical protein PFICI_02828 [Pestalotiopsis fici W106-1] | 66.4 | 0.0 |
| DIP_10002893 | hypothetical protein OIDMADRAFT_50282 [Oidiodendron maius Zn] | 51.7 | 0.0 |
| DIP_10002894 | putative lsm domain-containing protein [Eutypa lata UCREL1] | 89.8 | 6.0E-57 |
| DIP_10002895 | hypothetical protein PFICI_02826 [Pestalotiopsis fici W106-1] | 64.6 | 6.0E-99 |
| DIP_10002896 | hypothetical protein PFICI_02916 [Pestalotiopsis fici W106-1] | 66.1 | 8.0E-177 |
| DIP_10002897 | sorting nexin-3 [Cordyceps militaris CM01] | 83 | 5.0E-105 |
| DIP_10002898 | putative rna mp-1 protein [Togninia minima UCRPA7] | 44.8 | 6.0E-111 |
| DIP_10002899 | hypothetical protein PFICI_13438 [Pestalotiopsis fici W106-1] | 53.9 | 0.0 |
| DIP_10002900 | putative gat domain-containing protein [Eutypa lata UCREL1] | 57.4 | 0.0 |
| DIP_10002901 | F-box domain-containing protein [Colletotrichum graminicola M1.001] | 52.3 | 2.0E-173 |
| DIP_10002902 | putative pre-mrna-splicing factor 38b protein [Togninia minima UCRPA7] | 41.1 | 3.0E-57 |
| DIP_10002903 | hypothetical protein PFICI_07051 [Pestalotiopsis fici W106-1] | 62.2 | 4.0E-118 |
| DIP_10002904 | hypothetical protein PFICI_00504 [Pestalotiopsis fici W106-1] | 67.2 | 0.0 |
| DIP_10002905 | Pre-mRNA-splicing factor cwc24 [Pestalotiopsis fici W106-1] | 68 | 1.0E-160 |
| DIP_10002906 | putative anucleate primary sterigmata protein a protein [Eutypa lata UCREL1] | 47.4 | 0.0 |
| DIP_10002907 | NAD(P)-binding domain protein [Ophiocordyceps sinensis CO18] | 45.7 | 5.0E-49 |
| DIP_10002908 | hypothetical protein PFICI_12538 [Pestalotiopsis fici W106-1] | 42.9 | 0.0 |
| DIP_10002909 | hypothetical protein MYCFIDRAFT_71095 [Pseudocercospora fijiensis CIRAD86] | 70.8 | 8.0E-145 |
| DIP_10002910 | hypothetical protein CGLO_00894 [Colletotrichum gloeosporioides Cg-14] | 63.8 | 0.0 |
| DIP_10002911 | hypothetical protein PV04_02800 [Capronia semiimmersa] | 66.4 | 1.0E-60 |
| DIP_10002912 | hypothetical protein EMCG_01195 [Emmonsia crescens UAMH 3008] | 50.7 | 3.0E-21 |
| DIP_10002913 | hypothetical protein THAR02_07852 [Trichoderma harzianum] | 30.8 | 3.0E-37 |
| DIP_10002914 | Ankyrin repeat-containing domain protein [Metarhizium robertsii ARSEF 23] | 79.7 | 2.0E-78 |
| DIP_10002915 | hypothetical protein PFICI_12559 [Pestalotiopsis fici W106-1] | 70.3 | 8.0E-159 |
| DIP_10002916 | hypothetical protein UCREL1_971 [Eutypa lata UCREL1] | 67.2 | 0.0 |
| DIP_10002917 | putative family protein [Eutypa lata UCREL1] | 67.8 | 8.0E-142 |
| DIP_10002918 | putative protein-l-isoaspartate o-methyltransferase protein [Eutypa lata UCREL1] | 66.9 | 1.0E-129 |
| DIP_10002919 | NA | | |
| DIP_10002920 | hypothetical protein PFICI_03945 [Pestalotiopsis fici W106-1] | 70.5 | 0.0 |
| DIP_10002921 | hypothetical protein PFICI_10605 [Pestalotiopsis fici W106-1] | 47.5 | 8.0E-144 |
| DIP_10002922 | hypothetical protein PFICI_14060 [Pestalotiopsis fici W106-1] | 54 | 2.0E-106 |
| DIP_10002923 | hypothetical protein PFICI_12582 [Pestalotiopsis fici W106-1] | 54.4 | 1.0E-85 |
| DIP_10002924 | glycoside hydrolase family 93 protein [Trichoderma atroviride IMI 206040] | 49.1 | 3.0E-114 |
| DIP_10002925 | putative glycosyltransferase family 25 protein [Diaporthe ampelina] | 61.8 | 7.0E-138 |

| | | | |
|---|---|---|---|
| DIP_10002926 | hypothetical protein UCDDA912_g02479 [Diaporthe ampelina] | 43.6 | 2.0E-24 |
| DIP_10002927 | putative udp-glucose:glycoprotein glucosyltransferase protein [Eutypa lata UCREL1] | 78.7 | 0.0 |
| DIP_10002928 | putative coatomer subunit protein [Eutypa lata UCREL1] | 90.6 | 0.0 |
| DIP_10002929 | putative 26s proteasome non-atpase regulatory subunit 8 protein [Eutypa lata UCREL1] | 61.3 | 4.0E-124 |
| DIP_10002930 | putative chitin synthase activator protein [Eutypa lata UCREL1] | 67.5 | 0.0 |
| DIP_10002931 | NA | | |
| DIP_10002932 | NA | | |
| DIP_10002933 | hypothetical protein PFICI_05851 [Pestalotiopsis fici W106-1] | 46.2 | 3.0E-67 |
| DIP_10002934 | hypothetical protein PFICI_12360 [Pestalotiopsis fici W106-1] | 71.2 | 0.0 |
| DIP_10002935 | hypothetical protein AOL_s00054g931 [Arthrobotrys oligospora ATCC 24927] | 55.8 | 3.0E-165 |
| DIP_10002936 | putative 2-oxoacid dehydrogenase acyltransferase protein [Eutypa lata UCREL1] | 81 | 0.0 |
| DIP_10002937 | complex 1 protein [Colletotrichum higginsianum] | 83.8 | 4.0E-56 |
| DIP_10002938 | histone H2A [Metarhizium acridum CQMa 102] | 95.6 | 7.0E-86 |
| DIP_10002939 | putative histone h2b protein [Eutypa lata UCREL1] | 97.8 | 3.0E-90 |
| DIP_10002940 | Glutaryl-CoA dehydrogenase [Pestalotiopsis fici W106-1] | 77 | 0.0 |
| DIP_10002941 | hypothetical protein PFICI_14637 [Pestalotiopsis fici W106-1] | 79.1 | 0.0 |
| DIP_10002942 | L-xylulose reductase [Verticillium dahliae VdLs.17] | 83.3 | 5.0E-170 |
| DIP_10002943 | putative 26s protease regulatory subunit 6b protein [Eutypa lata UCREL1] | 91.5 | 0.0 |
| DIP_10002944 | hypothetical protein MAC_01461 [Metarhizium acridum CQMa 102] | 62.4 | 1.0E-51 |
| DIP_10002945 | hypothetical protein PFICI_03846 [Pestalotiopsis fici W106-1] | 69.4 | 0.0 |
| DIP_10002946 | putative high-affinity nicotinic acid transporter protein [Eutypa lata UCREL1] | 79.3 | 0.0 |
| DIP_10002947 | hypothetical protein PFICI_11515 [Pestalotiopsis fici W106-1] | 33.3 | 9.0E-18 |
| DIP_10002948 | hypothetical protein THITE_125074 [Thielavia terrestris NRRL 8126] | 51.6 | 2.0E-75 |
| DIP_10002949 | putative protein [Neurospora crassa] | 48 | 1.0E-129 |
| DIP_10002950 | NA | | |
| DIP_10002951 | hypothetical protein PFICI_03309 [Pestalotiopsis fici W106-1] | 31.5 | 7.0E-43 |
| DIP_10002952 | putative protein-tyrosine phosphatase 2 protein [Eutypa lata UCREL1] | 53.4 | 1.0E-163 |
| DIP_10002953 | hypothetical protein PFICI_05858 [Pestalotiopsis fici W106-1] | 46 | 0.0 |
| DIP_10002954 | dash complex subunit spc34 [Colletotrichum orbiculare MAFF 240422] | 61.4 | 1.0E-98 |
| DIP_10002955 | hypothetical protein UCREL1_10482 [Eutypa lata UCREL1] | 53.2 | 7.0E-62 |
| DIP_10002956 | hypothetical protein PFICI_12338 [Pestalotiopsis fici W106-1] | 40.5 | 3.0E-123 |
| DIP_10002957 | NA | | |
| DIP_10002958 | NA | | |
| DIP_10002959 | putative 26s proteasome non-atpase regulatory subunit 3 protein [Eutypa lata UCREL1] | 74.5 | 0.0 |
| DIP_10002960 | putative histone acetyltransferase sas2 [Diaporthe ampelina] | 47.5 | 2.0E-137 |
| DIP_10002961 | F-box only protein [Sphaerulina musiva SO2202] | 37 | 1.0E-119 |
| DIP_10002962 | putative atp synthase subunit 9 protein [Eutypa lata UCREL1] | 82.9 | 1.0E-79 |
| DIP_10002963 | C6 transcription factor [Fusarium sp. FN080326] | 37.6 | 6.0E-146 |
| DIP_10002964 | hypothetical protein FOMG_09668 [Fusarium oxysporum f. sp. melonis 26406] | 63.2 | 0.0 |
| DIP_10002965 | hypothetical protein NECHADRAFT_45518 [Nectria haematococca mpVI 77-13-4] | 50 | 4.0E-114 |
| DIP_10002966 | hypothetical protein PFICI_07330 [Pestalotiopsis fici W106-1] | 34.8 | 9.0E-53 |
| DIP_10002967 | NA | | |
| DIP_10002968 | GTPase-activator protein for ras-like GTPase containing protein [Colletotrichum orbiculare MAFF 240422] | 91.8 | 0.0 |
| DIP_10002969 | hypothetical protein PFICI_11920 [Pestalotiopsis fici W106-1] | 61.7 | 0.0 |
| DIP_10002970 | putative histone chaperone asf1 protein [Eutypa lata UCREL1] | 74.2 | 3.0E-117 |
| DIP_10002971 | hypothetical protein PFICI_03838 [Pestalotiopsis fici W106-1] | 59 | 0.0 |
| DIP_10002972 | nmrA-like family protein [Colletotrichum orbiculare MAFF 240422] | 50.5 | 4.0E-110 |
| DIP_10002973 | putative piwi domain-containing protein [Eutypa lata UCREL1] | 50.7 | 0.0 |
| DIP_10002974 | amino acid permease [Colletotrichum gloeosporioides Cg-14] | 58.8 | 0.0 |
| DIP_10002975 | NA | | |
| DIP_10002976 | NA | | |
| DIP_10002977 | hypothetical protein PFICI_02200 [Pestalotiopsis fici W106-1] | 70.9 | 0.0 |
| DIP_10002978 | related to ARG81-transcription factor involved in arginine metabolism [Fusarium fujikuroi IMI 58289] | 36.6 | 2.0E-152 |
| DIP_10002979 | putative dihydrodipicolinate synthase protein [Eutypa lata UCREL1] | 63.1 | 6.0E-131 |
| DIP_10002980 | fmn-dependent dehydrogenase family protein [Colletotrichum gloeosporioides Nara gc5] | 71.3 | 0.0 |
| DIP_10002981 | ferric reductase like transmembrane component [Colletotrichum gloeosporioides Nara gc5] | 39 | 4.0E-141 |
| DIP_10002982 | hypothetical protein T310_6745 [Rasamsonia emersonii CBS 393.64] | 31.2 | 6.0E-51 |
| DIP_10002983 | putative alpha-galactosidase a protein [Togninia minima UCRPA7] | 31 | 1.0E-12 |
| DIP_10002984 | hypothetical protein PFICI_03849 [Pestalotiopsis fici W106-1] | 69.8 | 3.0E-135 |
| DIP_10002985 | putative secreted aspartic proteinase protein [Eutypa lata UCREL1] | 56.2 | 3.0E-164 |
| DIP_10002986 | putative ammecr1 family protein [Eutypa lata UCREL1] | 65.6 | 4.0E-121 |
| DIP_10002987 | putative dna-repair protein rad2 protein [Eutypa lata UCREL1] | 86.8 | 0.0 |
| DIP_10002988 | NA | | |
| DIP_10002989 | hypothetical protein PFICI_12353 [Pestalotiopsis fici W106-1] | 56.6 | 7.0E-132 |
| DIP_10002990 | putative cytochrome p450 protein [Eutypa lata UCREL1] | 53.2 | 0.0 |
| DIP_10002991 | hypothetical protein PFICI_14390 [Pestalotiopsis fici W106-1] | 54.4 | 0.0 |
| DIP_10002992 | Tubulin binding cofactor C [Ophiocordyceps sinensis CO18] | 52.7 | 3.0E-46 |
| DIP_10002993 | NA | | |
| DIP_10002994 | NA | | |
| DIP_10002995 | NA | | |
| DIP_10002996 | hypothetical protein A1O1_01364 [Capronia coronata CBS 617.96] | 59.2 | 0.0 |
| DIP_10002997 | putative family protein [Eutypa lata UCREL1] | 63.3 | 1.0E-101 |
| DIP_10002998 | hypothetical protein [Podospora anserina S mat+] | 33.2 | 1.0E-23 |
| DIP_10002999 | putative lea domain-containing protein [Eutypa lata UCREL1] | 57.7 | 6.0E-35 |
| DIP_10003000 | putative casein kinase substrate phosphoprotein pp28 protein [Eutypa lata UCREL1] | 66.7 | 5.0E-81 |
| DIP_10003001 | hypothetical protein UCREL1_1627 [Eutypa lata UCREL1] | 57.9 | 2.0E-145 |
| DIP_10003002 | putative mitochondrial translation initiation protein [Eutypa lata UCREL1] | 65.2 | 0.0 |
| DIP_10003003 | hypothetical protein PFICI_03966 [Pestalotiopsis fici W106-1] | 69.8 | 0.0 |
| DIP_10003004 | putative duf367 domain-containing protein [Eutypa lata UCREL1] | 65.5 | 0.0 |
| DIP_10003005 | putative short chain dehydrogenase protein [Eutypa lata UCREL1] | 44.7 | 5.0E-89 |
| DIP_10003006 | phosphatidylinositol n-acetylglucosaminyltransferase subunit gpi1 [Ophiostoma piceae UAMH 11346] | 75 | 3.0E-43 |
| DIP_10003007 | hypothetical protein PFICI_14629 [Pestalotiopsis fici W106-1] | 79.7 | 0.0 |
| DIP_10003008 | hypothetical protein BC1G_07527 [Botrytis cinerea B05.10] | 60.9 | 9.0E-140 |
| DIP_10003009 | putative -bisphosphoglycerate-independent phosphoglycerate mutase protein [Eutypa lata UCREL1] | 82.5 | 0.0 |
| DIP_10003010 | hypothetical protein CH063_05574 [Colletotrichum higginsianum] | 48.2 | 1.0E-28 |
| DIP_10003011 | NA | | |
| DIP_10003012 | hypothetical protein PFICI_03258 [Pestalotiopsis fici W106-1] | 39.1 | 3.0E-87 |
| DIP_10003013 | Glucan endo-1,3-alpha-glucosidase agn1 [Talaromyces marneffei PM1] | 60.5 | 0.0 |
| DIP_10003014 | putative fad binding domain-containing protein [Eutypa lata UCREL1] | 37.3 | 4.0E-79 |
| DIP_10003015 | putative catalase [Colletotrichum sublineola] | 32 | 2.0E-40 |
| DIP_10003016 | putative dna polymerase epsilon protein [Eutypa lata UCREL1] | 79.2 | 0.0 |
| DIP_10003017 | hypothetical protein PFICI_03266 [Pestalotiopsis fici W106-1] | 55.9 | 6.0E-63 |
| DIP_10003018 | hypothetical protein UCREL1_3209 [Eutypa lata UCREL1] | 35.2 | 6.0E-54 |
| DIP_10003019 | hypothetical protein PILCRDRAFT_78838 [Piloderma croceum F 1598] | 31.4 | 1.0E-18 |
| DIP_10003020 | putative aldose 1-epimerase protein [Eutypa lata UCREL1] | 69.9 | 2.0E-171 |
| DIP_10003021 | hypothetical protein PFICI_03263 [Pestalotiopsis fici W106-1] | 64.7 | 0.0 |
| DIP_10003022 | hypothetical protein PFICI_02226 [Pestalotiopsis fici W106-1] | 59.9 | 0.0 |
| DIP_10003023 | NA | | |
| DIP_10003024 | hypothetical protein W97_00979 [Coniosporium apollinis CBS 100218] | 30.3 | 1.0E-11 |
| DIP_10003025 | NA | | |
| DIP_10003026 | hypothetical protein PFICI_05909 [Pestalotiopsis fici W106-1] | 76.6 | 0.0 |
| DIP_10003027 | putative metallo-beta-lactamase domain protein [Eutypa lata UCREL1] | 55.2 | 2.0E-102 |
| DIP_10003028 | putative integral membrane protein [Eutypa lata UCREL1] | 70.8 | 0.0 |
| DIP_10003029 | putative ribosome assembly protein 3 protein [Eutypa lata UCREL1] | 55.8 | 7.0E-16 |
| DIP_10003030 | putative 26s proteasome regulatory subunit rpn9 protein [Eutypa lata UCREL1] | 79.8 | 0.0 |
| DIP_10003031 | NADH-ubiquinone oxidoreductase subunit A [Pestalotiopsis fici W106-1] | 84 | 0.0 |
| DIP_10003032 | hypothetical protein THITE_2117465 [Thielavia terrestris NRRL 8126] | 78.1 | 4.0E-73 |
| DIP_10003033 | hypothetical protein FPSE_12264 [Fusarium pseudograminearum CS3096] | 52.5 | 2.0E-134 |
| DIP_10003034 | putative ran-interacting mog1 protein [Eutypa lata UCREL1] | 65.5 | 2.0E-93 |
| DIP_10003035 | putative glutamine-dependent nad(+) synthetase protein [Eutypa lata UCREL1] | 85.4 | 0.0 |
| DIP_10003036 | NA | | |
| DIP_10003037 | putative aconitate hydratase protein [Eutypa lata UCREL1] | 86.2 | 0.0 |
| DIP_10003038 | putative zinc finger protein [Eutypa lata UCREL1] | 71.1 | 5.0E-58 |
| DIP_10003039 | hypothetical protein PFICI_05918 [Pestalotiopsis fici W106-1] | 70.6 | 3.0E-64 |
| DIP_10003040 | putative rhomboid protein 2 protein [Eutypa lata UCREL1] | 74.1 | 1.0E-136 |
| DIP_10003041 | NA | | |
| DIP_10003042 | putative had superfamily hydrolase protein [Eutypa lata UCREL1] | 73.5 | 2.0E-129 |
| DIP_10003043 | putative eukaryotic translation initiation factor 3 subunit protein [Togninia minima UCRPA7] | 71 | 8.0E-126 |
| DIP_10003044 | hypothetical protein CGLO_15544 [Colletotrichum gloeosporioides Cg-14] | 51.7 | 2.0E-137 |
| DIP_10003045 | NA | | |
| DIP_10003046 | hypothetical protein S40285_04927 [Stachybotrys chlorohalonata IBT 40285] | 46.7 | 5.0E-81 |
| DIP_10003047 | hypothetical protein S40285_04929 [Stachybotrys chlorohalonata IBT 40285] | 40.1 | 2.0E-85 |
| DIP_10003048 | putative letm1-like protein [Eutypa lata UCREL1] | 49.5 | 2.0E-78 |
| DIP_10003049 | NA | | |
| DIP_10003050 | NA | | |
| DIP_10003051 | hypothetical protein THITE_2117298 [Thielavia terrestris NRRL 8126] | 60.3 | 6.0E-64 |
| DIP_10003052 | Calcium/calmodulin-dependent protein kinase type I [Pestalotiopsis fici W106-1] | 82.2 | 0.0 |
| DIP_10003053 | hypothetical protein FVEG_12614 [Fusarium verticillioides 7600] | 74.9 | 0.0 |
| DIP_10003054 | hypothetical protein NECHADRAFT_87379 [Nectria haematococca mpVI 77-13-4] | 56.9 | 0.0 |
| DIP_10003055 | RNA polymerase I, subunit RPA34.5, partial [Metarhizium majus ARSEF 297] | 33.8 | 3.0E-40 |
| DIP_10003056 | hypothetical protein PFICI_14691 [Pestalotiopsis fici W106-1] | 41.2 | 0.0 |
| DIP_10003057 | NA | | |
| DIP_10003058 | hypothetical protein PFICI_12535 [Pestalotiopsis fici W106-1] | 44.1 | 0.0 |
| DIP_10003059 | hypothetical protein PFICI_12537 [Pestalotiopsis fici W106-1] | 40.7 | 0.0 |
| DIP_10003060 | putative family protein [Eutypa lata UCREL1] | 52.8 | 4.0E-145 |
| DIP_10003061 | NA | | |
| DIP_10003062 | NA | | |
| DIP_10003063 | putative alcohol dehydrogenase protein [Eutypa lata UCREL1] | 49.4 | 8.0E-114 |
| DIP_10003064 | putative vacuolar protease a protein [Eutypa lata UCREL1] | 85.7 | 0.0 |
| DIP_10003065 | hypothetical protein PFICI_12379 [Pestalotiopsis fici W106-1] | 50.3 | 0.0 |
| DIP_10003066 | NA | | |
| DIP_10003067 | putative bnr asp-box repeat domain protein [Eutypa lata UCREL1] | 36.5 | 3.0E-48 |
| DIP_10003068 | putative serine hydroxymethyltransferase protein [Eutypa lata UCREL1] | 87 | 0.0 |
| DIP_10003069 | NA | | |
| DIP_10003070 | putative -beta-glucan synthase component gls1 protein [Eutypa lata UCREL1] | 80.7 | 0.0 |
| DIP_10003071 | NA | | |
| DIP_10003072 | putative hmg box-containing protein [Eutypa lata UCREL1] | 40.4 | 1.0E-119 |
| DIP_10003073 | hypothetical protein V501_01790, partial [Pseudogymnoascus pannorum VKM F-4519 (FW-2642)] | 30.9 | 5.0E-62 |
| DIP_10003074 | putative cell wall glucanase protein [Eutypa lata UCREL1] | 65.2 | 3.0E-123 |
| DIP_10003075 | putative serine palmitoyltransferase 1 protein [Eutypa lata UCREL1] | 81.2 | 0.0 |
| DIP_10003076 | Gelsolin [Penicillium expansum] | 48.5 | 2.0E-120 |
| DIP_10003077 | hypothetical protein PFICI_05929 [Pestalotiopsis fici W106-1] | 49.4 | 2.0E-80 |
| DIP_10003078 | hypothetical protein NECHADRAFT_92359 [Nectria haematococca mpVI 77-13-4] | 50 | 0.0 |
| DIP_10003079 | NA | | |
| DIP_10003080 | hypothetical protein PFICI_05927 [Pestalotiopsis fici W106-1] | 66.4 | 0.0 |
| DIP_10003081 | hypothetical protein PFICI_05926 [Pestalotiopsis fici W106-1] | 35.2 | 3.0E-40 |
| DIP_10003082 | putative rta1 domain protein [Eutypa lata UCREL1] | 58.2 | 1.0E-154 |
| DIP_10003083 | hypothetical protein PFICI_03850 [Pestalotiopsis fici W106-1] | 54.6 | 5.0E-101 |
| DIP_10003084 | putative o- protein [Eutypa lata UCREL1] | 47.9 | 2.0E-154 |
| DIP_10003085 | NA | | |
| DIP_10003086 | NA | | |
| DIP_10003087 | hypothetical protein [Podospora anserina S mat+] | 63 | 0.0 |
| DIP_10003088 | putative eukaryotic translation initiation factor 2a protein [Eutypa lata UCREL1] | 72.2 | 0.0 |
| DIP_10003089 | putative l-fucose permease glucose galactose transporter protein [Togninia minima UCRPA7] | 53.6 | 5.0E-177 |
| DIP_10003090 | T-complex protein 1 subunit eta [Pestalotiopsis fici W106-1] | 90.9 | 0.0 |
| DIP_10003091 | putative histone acetyltransferase spt10 protein [Eutypa lata UCREL1] | 74.1 | 0.0 |
| DIP_10003092 | NA | | |
| DIP_10003093 | NA | | |
| DIP_10003094 | putative 37s ribosomal protein rsmp4 protein [Eutypa lata UCREL1] | 51.8 | 1.0E-138 |
| DIP_10003095 | putative mitochondrial ribosomal protein subunit s4 protein [Eutypa lata UCREL1] | 83.2 | 0.0 |
| DIP_10003096 | hypothetical protein S40293_06653 [Stachybotrys chartarum IBT 40293] | 42.8 | 8.0E-142 |
| DIP_10003097 | NA | | |
| DIP_10003098 | hypothetical protein OIDMADRAFT_131398 [Oidiodendron maius Zn] | 46.5 | 2.0E-64 |
| DIP_10003099 | NA | | |
| DIP_10003100 | hypothetical protein PFICI_03162 [Pestalotiopsis fici W106-1] | 53.8 | 0.0 |
| DIP_10003101 | hypothetical protein PFICI_03161 [Pestalotiopsis fici W106-1] | 89.5 | 2.0E-123 |
| DIP_10003102 | NA | | |
| DIP_10003103 | putative f-box domain-containing protein [Eutypa lata UCREL1] | 38.8 | 2.0E-81 |
| DIP_10003104 | NA | | |
| DIP_10003105 | putative membrane transporter protein [Eutypa lata UCREL1] | 55.6 | 9.0E-157 |
| DIP_10003106 | Cytochrome p450 protein [Acremonium chrysogenum ATCC 11550] | 52.2 | 1.0E-180 |
| DIP_10003107 | predicted protein [Trichoderma reesei QM6a] | 41.7 | 1.0E-131 |
| DIP_10003108 | hypothetical protein OCS_04346 [Ophiocordyceps sinensis CO18] | 54.2 | 4.0E-19 |
| DIP_10003109 | hypothetical protein TRIVIDRAFT_36225 [Trichoderma virens Gv29-8] | 30.1 | 6.0E-22 |
| DIP_10003110 | hypothetical protein PFICI_09501 [Pestalotiopsis fici W106-1] | 41.2 | 2.0E-179 |
| DIP_10003111 | hypothetical protein PFICI_14738 [Pestalotiopsis fici W106-1] | 68.5 | 0.0 |
| DIP_10003112 | putative surfeit locus protein 6 protein [Eutypa lata UCREL1] | 62.8 | 0.0 |
| DIP_10003113 | hypothetical protein EPUS_05455 [Endocarpon pusillum Z07020] | 55.4 | 9.0E-105 |
| DIP_10003114 | hypothetical protein PFICI_03452 [Pestalotiopsis fici W106-1] | 48.7 | 1.0E-135 |
| DIP_10003115 | putative transcription factor protein [Eutypa lata UCREL1] | 53.2 | 3.0E-153 |
| DIP_10003116 | hypothetical protein PFICI_03482 [Pestalotiopsis fici W106-1] | 52.5 | 0.0 |
| DIP_10003117 | putative vacuolar protein sorting-associated protein 74 protein [Eutypa lata UCREL1] | 87.5 | 0.0 |
| DIP_10003118 | T-complex protein 1 subunit zeta [Pestalotiopsis fici W106-1] | 92.9 | 0.0 |
| DIP_10003119 | Putative prenylcysteine oxidase [Fusarium oxysporum f. sp. cubense race 4] | 52.3 | 3.0E-165 |
| DIP_10003120 | NA | | |
| DIP_10003121 | hypothetical protein FOZG_02996 [Fusarium oxysporum Fo47] | 46.4 | 9.0E-86 |
| DIP_10003122 | hypothetical protein PV09_05249 [Verruconis gallopava] | 39.5 | 2.0E-40 |
| DIP_10003123 | putative rint-1 family protein [Eutypa lata UCREL1] | 63.8 | 0.0 |
| DIP_10003124 | putative mitochondrial carrier protein [Eutypa lata UCREL1] | 80.5 | 8.0E-167 |
| DIP_10003125 | hypothetical protein PFICI_08649 [Pestalotiopsis fici W106-1] | 39.7 | 5.0E-171 |
| DIP_10003126 | NA | | |
| DIP_10003127 | putative sec34-like family protein [Eutypa lata UCREL1] | 68.9 | 0.0 |
| DIP_10003128 | putative rrna processing protein [Eutypa lata UCREL1] | 69.4 | 0.0 |
| DIP_10003129 | putative amino acid transporter protein [Eutypa lata UCREL1] | 66.6 | 0.0 |
| DIP_10003130 | hypothetical protein CONPUDRAFT_139443 [Coniophora puteana RWD-64-598 SS2] | 44.8 | 5.0E-80 |
| DIP_10003131 | NA | | |
| DIP_10003132 | putative transmembrane alpha-helix domain-containing protein [Eutypa lata UCREL1] | 81 | 4.0E-46 |
| DIP_10003133 | NA | | |
| DIP_10003134 | hypothetical protein GLRG_02029 [Colletotrichum graminicola M1.001] | 81.7 | 2.0E-43 |
| DIP_10003135 | putative exosome complex exonuclease rrp protein [Eutypa lata UCREL1] | 61.3 | 0.0 |
| DIP_10003136 | NA | | |
| DIP_10003137 | conserved hypothetical protein [Talaromyces marneffei ATCC 18224] | 48.9 | 5.0E-41 |
| DIP_10003138 | putative het domain-containing protein [Eutypa lata UCREL1] | 51.6 | 0.0 |
| DIP_10003139 | hypothetical protein UCREL1_7274 [Eutypa lata UCREL1] | 53.3 | 0.0 |
| DIP_10003140 | bifunctional catalase-peroxidase cat2 [Colletotrichum orbiculare MAFF 240422] | 80.9 | 0.0 |
| DIP_10003141 | hypothetical protein PFICI_03278 [Pestalotiopsis fici W106-1] | 46.7 | 5.0E-63 |
| DIP_10003142 | hypothetical protein NEUTE1DRAFT_117873 [Neurospora tetrasperma FGSC 2508] | 63.7 | 6.0E-138 |
| DIP_10003143 | putative prefoldin subunit 3 protein [Eutypa lata UCREL1] | 80.2 | 1.0E-110 |
| DIP_10003144 | putative dna repair and recombination protein rad54 protein [Eutypa lata UCREL1] | 87.8 | 0.0 |
| DIP_10003145 | NA | | |
| DIP_10003146 | hypothetical protein PFICI_05899 [Pestalotiopsis fici W106-1] | 52.9 | 0.0 |
| DIP_10003147 | putative ubiquitin thioesterase otu1 protein [Eutypa lata UCREL1] | 65.5 | 4.0E-158 |
| DIP_10003148 | hypothetical protein PDE_08244 [Penicillium oxalicum 114-2] | 34.7 | 1.0E-32 |
| DIP_10003149 | putative glycerophosphoryl diester phosphodiesterase family protein [Eutypa lata UCREL1] | 42.2 | 4.0E-108 |
| DIP_10003150 | hypothetical protein GGTG_12651 [Gaeumannomyces graminis var. tritici R3-111a-1] | 31.9 | 5.0E-21 |
| DIP_10003151 | putative transmembrane protein [Togninia minima UCRPA7] | 68.1 | 0.0 |
| DIP_10003152 | NA | | |
| DIP_10003153 | hypothetical protein UCREL1_5170 [Eutypa lata UCREL1] | 42.6 | 3.0E-47 |
| DIP_10003154 | putative rrna processing protein [Eutypa lata UCREL1] | 62.8 | 0.0 |
| DIP_10003155 | putative cytochrome p450 monooxygenase protein [Eutypa lata UCREL1] | 67.4 | 6.0E-164 |
| DIP_10003156 | hypothetical protein FOXB_13553 [Fusarium oxysporum Fo5176] | 65.9 | 0.0 |
| DIP_10003157 | hypothetical protein EMCG_03372 [Emmonsia crescens UAMH 3008] | 34.6 | 4.0E-22 |
| DIP_10003158 | NA | | |
| DIP_10003159 | hypothetical protein PFICI_14441 [Pestalotiopsis fici W106-1] | 49.7 | 9.0E-37 |
| DIP_10003160 | hypothetical protein PFICI_14447 [Pestalotiopsis fici W106-1] | 55.4 | 1.0E-148 |
| DIP_10003161 | major facilitator superfamily transporter [Colletotrichum gloeosporioides Cg-14] | 56.5 | 0.0 |
| DIP_10003162 | hypothetical protein CGLO_02494 [Colletotrichum gloeosporioides Cg-14] | 76.3 | 0.0 |
| DIP_10003163 | zn 2cys6 transcription factor [Colletotrichum orbiculare MAFF 240422] | 48.4 | 6.0E-105 |
| DIP_10003164 | hypothetical protein NECHADRAFT_49675 [Nectria haematococca mpVI 77-13-4] | 63.4 | 1.0E-119 |
| DIP_10003165 | amidohydrolase 2 [Colletotrichum orbiculare MAFF 240422] | 58.7 | 1.0E-130 |
| DIP_10003166 | 5'-nucleotidase domain-containing protein, partial [Rhizobium sp. Pop5] | 30 | 1.0E-28 |
| DIP_10003167 | hypothetical protein PFICI_03559 [Pestalotiopsis fici W106-1] | 77.7 | 6.0E-148 |
| DIP_10003168 | predicted protein, partial [Ostreococcus lucimarinus CCE9901] | 30 | 6.0E-10 |
| DIP_10003169 | hypothetical protein PFICI_03557 [Pestalotiopsis fici W106-1] | 63.6 | 0.0 |
| DIP_10003170 | hypothetical protein PFICI_03555 [Pestalotiopsis fici W106-1] | 39.6 | 2.0E-37 |
| DIP_10003171 | Succinyl-CoA:3-ketoacid-coenzyme A transferase 1 [Pestalotiopsis fici W106-1] | 88.1 | 0.0 |
| DIP_10003172 | putative pre-rrna processing protein utp22 protein [Eutypa lata UCREL1] | 58 | 0.0 |
| DIP_10003173 | putative transport protein sec24 protein [Eutypa lata UCREL1] | 79.2 | 0.0 |
| DIP_10003174 | hypothetical protein PFICI_14757 [Pestalotiopsis fici W106-1] | 72.5 | 0.0 |
| DIP_10003175 | hypothetical protein PFICI_12547 [Pestalotiopsis fici W106-1] | 67.2 | 0.0 |
| DIP_10003176 | hypothetical protein PV04_09094 [Capronia semiimmersa] | 33.3 | 5.0E-86 |
| DIP_10003177 | uncharacterized protein FFUJ_12436 [Fusarium fujikuroi IMI 58289] | 40.4 | 1.0E-71 |
| DIP_10003178 | NA | | |
| DIP_10003179 | putative terpene synthase metal-binding domain protein [Eutypa lata UCREL1] | 72.5 | 0.0 |
| DIP_10003180 | hypothetical protein S7711_06216 [Stachybotrys chartarum IBT 7711] | 67.3 | 2.0E-128 |
| DIP_10003181 | hypothetical protein PFICI_11079 [Pestalotiopsis fici W106-1] | 37.7 | 0.0 |
| DIP_10003182 | putative nad dependent epimerase dehydratase protein [Eutypa lata UCREL1] | 77.7 | 4.0E-159 |
| DIP_10003183 | putative cleavage and polyadenylation specificity protein [Togninia minima UCRPA7] | 58.9 | 0.0 |
| DIP_10003184 | putative stress responsive a b barrel domain-containing protein [Eutypa lata UCREL1] | 53.9 | 3.0E-33 |
| DIP_10003185 | hypothetical protein PFICI_14755 [Pestalotiopsis fici W106-1] | 63.2 | 1.0E-126 |
| DIP_10003186 | hypothetical protein PFICI_14756 [Pestalotiopsis fici W106-1] | 58 | 2.0E-86 |
| DIP_10003187 | NA | | |
| DIP_10003188 | hypothetical protein PFICI_14754 [Pestalotiopsis fici W106-1] | 54.7 | 2.0E-104 |
| DIP_10003189 | hypothetical protein PFICI_14753 [Pestalotiopsis fici W106-1] | 76.2 | 9.0E-57 |
| DIP_10003190 | putative delta fatty acid desaturase protein [Eutypa lata UCREL1] | 71.7 | 0.0 |
| DIP_10003191 | putative alpha beta-hydrolase protein [Eutypa lata UCREL1] | 43.5 | 1.0E-164 |
| DIP_10003192 | hypothetical protein HIM_11218 [Hirsutella minnesotensis 3608] | 35.4 | 8.0E-07 |
| DIP_10003193 | NA | | |
| DIP_10003194 | putative cytochrome b5 protein [Eutypa lata UCREL1] | 57.8 | 2.0E-179 |
| DIP_10003195 | cytochrome p450 [Colletotrichum orbiculare MAFF 240422] | 71.3 | 0.0 |
| DIP_10003196 | hypothetical protein UCRPA7_3981 [Togninia minima UCRPA7] | 52.1 | 0.0 |
| DIP_10003197 | phosphoglycerate mutase family protein, putative [Aspergillus flavus NRRL3357] | 62.1 | 1.0E-66 |
| DIP_10003198 | hypothetical protein PFICI_08657 [Pestalotiopsis fici W106-1] | 51.3 | 0.0 |
| DIP_10003199 | NA | | |
| DIP_10003200 | glutamate decarboxylase [Verticillium dahliae VdLs.17] | 64.8 | 0.0 |
| DIP_10003201 | NA | | |
| DIP_10003202 | hypothetical protein MYCTH_2308699 [Myceliophthora thermophila ATCC 42464] | 53.5 | 9.0E-77 |
| DIP_10003203 | putative bromodomain-containing protein [Eutypa lata UCREL1] | 74.9 | 0.0 |
| DIP_10003204 | NA | | |
| DIP_10003205 | hydrophobin 3 precursor [Fusarium culmorum] | 61.3 | 2.0E-12 |
| DIP_10003206 | hypothetical protein V497_03867 [Pseudogymnoascus pannorum VKM F-4516 (FW-969)] | 30.3 | 9.0E-16 |
| DIP_10003207 | hypothetical protein [Podospora anserina S mat+] | 36.6 | 1.0E-38 |
| DIP_10003208 | methyltransferase domain-containing protein [Colletotrichum gloeosporioides Nara gc5] | 54.2 | 6.0E-123 |
| DIP_10003209 | NA | | |
| DIP_10003210 | putative acyl- dehydrogenase protein [Eutypa lata UCREL1] | 87 | 0.0 |
| DIP_10003211 | hypothetical protein FOXB_03045 [Fusarium oxysporum Fo5176] | 38.2 | 3.0E-84 |
| DIP_10003212 | aldo-keto reductase (AKR13), puatative [Neosartorya fischeri NRRL 181] | 57.4 | 2.0E-132 |
| DIP_10003213 | hypothetical protein THAR02_06363 [Trichoderma harzianum] | 30.6 | 2.0E-63 |
| DIP_10003214 | hypothetical protein ARAM_005212 [Aspergillus rambellii] | 34.1 | 2.0E-90 |
| DIP_10003215 | hypothetical protein FAVG1_11619 [Fusarium avenaceum] | 60.5 | 0.0 |
| DIP_10003216 | hypothetical protein FPSE_06091 [Fusarium pseudograminearum CS3096] | 34.1 | 2.0E-37 |
| DIP_10003217 | putative protein sik1 protein [Eutypa lata UCREL1] | 87.3 | 0.0 |
| DIP_10003218 | putative ubiquitin-40s ribosomal protein s31 fusion protein [Eutypa lata UCREL1] | 99.4 | 6.0E-109 |
| DIP_10003219 | 40S ribosomal protein S26E [Neurospora crassa OR74A] | 94.1 | 7.0E-79 |
| DIP_10003220 | hypothetical protein PFICI_14487 [Pestalotiopsis fici W106-1] | 70.6 | 0.0 |
| DIP_10003221 | hypothetical protein UCREL1_7906 [Eutypa lata UCREL1] | 53.4 | 7.0E-106 |
| DIP_10003222 | hypothetical protein PV09_06386 [Verruconis gallopava] | 67.6 | 9.0E-142 |
| DIP_10003223 | hypothetical protein SAPIO_CDS10240 [Scedosporium apiospermum] | 34.2 | 1.0E-47 |
| DIP_10003224 | putative phospholipid:diacylglycerol acyltransferase protein [Togninia minima UCRPA7] | 71.8 | 0.0 |
| DIP_10003225 | hypothetical protein PFICI_03523 [Pestalotiopsis fici W106-1] | 85 | 2.0E-98 |
| DIP_10003226 | hypothetical protein TRIATDRAFT_319051 [Trichoderma atroviride IMI 206040] | 53.2 | 1.0E-110 |
| DIP_10003227 | Protein MSN5 [Beauveria bassiana D1-5] | 49.9 | 0.0 |
| DIP_10003228 | hypothetical protein PFICI_03518 [Pestalotiopsis fici W106-1] | 62.9 | 7.0E-171 |
| DIP_10003229 | hypothetical protein PFICI_03517 [Pestalotiopsis fici W106-1] | 75.3 | 0.0 |
| DIP_10003230 | putative pyruvate decarboxylase protein [Eutypa lata UCREL1] | 67.5 | 0.0 |
| DIP_10003231 | hypothetical protein THITE_2145027 [Thielavia terrestris NRRL 8126] | 62.2 | 2.0E-72 |
| DIP_10003232 | uroporphyrinogen-III synthase [Trichoderma harzianum] | 52.4 | 3.0E-111 |
| DIP_10003233 | putative oxidation resistance protein 1 protein [Eutypa lata UCREL1] | 61.4 | 2.0E-141 |
| DIP_10003234 | mitogen activated protein kinase [Fusarium fujikuroi] | 95.8 | 0.0 |
| DIP_10003235 | putative transport protein sec24 protein [Eutypa lata UCREL1] | 84.2 | 0.0 |
| DIP_10003236 | hypothetical protein GLRG_06775 [Colletotrichum graminicola M1.001] | 48.2 | 2.0E-28 |
| DIP_10003237 | hypothetical protein PFICI_03549 [Pestalotiopsis fici W106-1] | 52.8 | 0.0 |
| DIP_10003238 | hypothetical protein PFICI_03543 [Pestalotiopsis fici W106-1] | 59.8 | 0.0 |
| DIP_10003239 | putative ubiquitin carboxyl-terminal hydrolase protein [Eutypa lata UCREL1] | 40.9 | 8.0E-99 |
| DIP_10003240 | putative phosphatidylserine synthase protein [Eutypa lata UCREL1] | 72.3 | 1.0E-127 |
| DIP_10003241 | hypothetical protein HIM_08201 [Hirsutella minnesotensis 3608] | 50.1 | 0.0 |
| DIP_10003242 | putative nrm1-like protein [Togninia minima UCRPA7] | 66.6 | 3.0E-158 |
| DIP_10003243 | hypothetical protein NEUTE1DRAFT_85981 [Neurospora tetrasperma FGSC 2508] | 38.3 | 5.0E-111 |
| DIP_10003244 | putative caspase domain-containing protein [Togninia minima UCRPA7] | 75.6 | 0.0 |
| DIP_10003245 | NA | | |
| DIP_10003246 | putative f-box domain-containing protein [Eutypa lata UCREL1] | 49.6 | 6.0E-93 |
| DIP_10003247 | f-box domain-containing protein [Colletotrichum orbiculare MAFF 240422] | 42.9 | 3.0E-36 |
| DIP_10003248 | putative tropomyosin-1 alpha chain protein [Eutypa lata UCREL1] | 60.4 | 0.0 |
| DIP_10003249 | putative endochitinase 1 precursor protein [Eutypa lata UCREL1] | 77.6 | 0.0 |
| DIP_10003250 | endosome-associated ubiquitin isopeptidase [Colletotrichum gloeosporioides Nara gc5] | 39.9 | 7.0E-52 |
| DIP_10003251 | hypothetical protein PFICI_14431 [Pestalotiopsis fici W106-1] | 75.8 | 2.0E-60 |
| DIP_10003252 | Vacuolar protein sorting-associated protein 4 [Pestalotiopsis fici W106-1] | 85 | 0.0 |
| DIP_10003253 | subtilisin-like serine protease precursor [Purpureocillium lilacinum] | 55.2 | 5.0E-145 |
| DIP_10003254 | putative atp-dependent rna helicase dbh1 protein [Eutypa lata UCREL1] | 85.9 | 0.0 |
| DIP_10003255 | hypothetical protein PFICI_12225 [Pestalotiopsis fici W106-1] | 58.2 | 9.0E-91 |
| DIP_10003256 | aspartyl/glutamyl-tRNA(Asn/Gln) amidotransferase subunit B [Verticillium alfalfae VaMs.102] | 78.7 | 6.0E-145 |
| DIP_10003257 | putative thump domain containing protein [Eutypa lata UCREL1] | 49.2 | 5.0E-104 |
| DIP_10003258 | hypothetical protein UCREL1_2188 [Eutypa lata UCREL1] | 37.8 | 2.0E-101 |
| DIP_10003259 | NA | | |
| DIP_10003260 | NA | | |
| DIP_10003261 | H/ACA ribonucleoprotein complex subunit 1 [Verticillium dahliae VdLs.17] | 87 | 6.0E-62 |
| DIP_10003262 | putative threonyl-trna synthetase protein [Eutypa lata UCREL1] | 77 | 0.0 |
| DIP_10003263 | NA | | |
| DIP_10003264 | putative nucleolar essential protein 1 protein [Eutypa lata UCREL1] | 90.3 | 3.0E-165 |
| DIP_10003265 | hypothetical protein CSUB01_02877 [Colletotrichum sublineola] | 32.8 | 7.0E-66 |
| DIP_10003266 | hypothetical protein PFICI_03505 [Pestalotiopsis fici W106-1] | 60.9 | 0.0 |
| DIP_10003267 | NA | | |
| DIP_10003268 | hypothetical protein PFICI_03504 [Pestalotiopsis fici W106-1] | 83.8 | 0.0 |
| DIP_10003269 | hypothetical protein NECHADRAFT_38507 [Nectria haematococca mpVI 77-13-4] | 39.9 | 5.0E-69 |
| DIP_10003270 | putative and tpr domain-containing protein [Eutypa lata UCREL1] | 62.3 | 0.0 |
| DIP_10003271 | NA | | |
| DIP_10003272 | mitoferrin [Verticillium dahliae VdLs.17] | 84.5 | 0.0 |
| DIP_10003273 | hypothetical protein PFICI_03529 [Pestalotiopsis fici W106-1] | 63.8 | 0.0 |
| DIP_10003274 | putative component of oligomeric golgi complex 6 protein [Eutypa lata UCREL1] | 71.5 | 0.0 |
| DIP_10003275 | putative mitochondrial-processing peptidase subunit alpha protein [Eutypa lata UCREL1] | 49.9 | 1.0E-138 |
| DIP_10003276 | putative ribosomal protein s17 protein [Eutypa lata UCREL1] | 58.3 | 3.0E-54 |
| DIP_10003277 | hypothetical protein UVI_009370 [Ustilaginoidea virens] | 61.9 | 4.0E-93 |
| DIP_10003278 | putative benzoate 4-monooxygenase cytochrome p450 protein [Eutypa lata UCREL1] | 55.9 | 5.0E-67 |
| DIP_10003279 | putative benzoate 4-monooxygenase cytochrome p450 protein [Eutypa lata UCREL1] | 49.2 | 7.0E-73 |
| DIP_10003280 | putative aaa family atpase protein [Eutypa lata UCREL1] | 49.9 | 3.0E-132 |
| DIP_10003281 | putative dna repair protein [Eutypa lata UCREL1] | 53.9 | 0.0 |
| DIP_10003282 | hypothetical protein PFICI_03509 [Pestalotiopsis fici W106-1] | 59 | 6.0E-96 |
| DIP_10003283 | hypothetical protein H634G_00515 [Metarhizium anisopliae BRIP 53293] | 66.1 | 9.0E-149 |
| DIP_10003284 | putative signal recognition particle 14kd protein [Eutypa lata UCREL1] | 61.3 | 4.0E-38 |
| DIP_10003285 | putative suppressor of glycerol defect protein [Eutypa lata UCREL1] | 57.1 | 0.0 |
| DIP_10003286 | Dihydrolipoyl dehydrogenase [Pestalotiopsis fici W106-1] | 91 | 0.0 |
| DIP_10003287 | putative alpha-galactosidase c protein [Eutypa lata UCREL1] | 71.8 | 0.0 |
| DIP_10003288 | putative rna polymerase ii ctd phosphatase protein [Eutypa lata UCREL1] | 59.8 | 0.0 |
| DIP_10003289 | putative nadh-ubiquinone oxidoreductase subunit grim-19 protein [Eutypa lata UCREL1] | 87.4 | 2.0E-76 |
| DIP_10003290 | putative fad binding domain-containing protein [Eutypa lata UCREL1] | 61.5 | 0.0 |
| DIP_10003291 | hypothetical protein FAVG1_06883 [Fusarium avenaceum] | 46 | 2.0E-17 |

| | | | |
|---|---|---|---|
| DIP_10003292 | putative condensin complex subunit 1 protein [Eutypa lata UCREL1] | 85.1 | 0.0 |
| DIP_10003293 | hypothetical protein PFICI_03977 [Pestalotiopsis fici W106-1] | 61.6 | 0.0 |
| DIP_10003294 | NA | | |
| DIP_10003295 | putative wd repeat-containing protein [Eutypa lata UCREL1] | 73.8 | 0.0 |
| DIP_10003296 | hypothetical protein UCREL1_3816 [Eutypa lata UCREL1] | 42 | 3.0E-82 |
| DIP_10003297 | putative nitrate assimilation regulatory protein nira protein [Togninia minima UCRPA7] | 30.3 | 4.0E-43 |
| DIP_10003298 | hypothetical protein THAR02_01702 [Trichoderma harzianum] | 40.7 | 6.0E-62 |
| DIP_10003299 | hypothetical protein PFICI_03940 [Pestalotiopsis fici W106-1] | 67.2 | 0.0 |
| DIP_10003300 | hypothetical protein HIM_06171 [Hirsutella minnesotensis 3608] | 53 | 0.0 |
| DIP_10003301 | hypothetical protein PFICI_05107 [Pestalotiopsis fici W106-1] | 66.4 | 0.0 |
| DIP_10003302 | iron-sulfur cluster transporter ATM1 [Colletotrichum fioriniae PJ7] | 83.5 | 0.0 |
| DIP_10003303 | hypothetical protein SEPMUDRAFT_149381 [Sphaerulina musiva SO2202] | 31.5 | 4.0E-28 |
| DIP_10003304 | hypothetical protein LOTGIDRAFT_88143, partial [Lottia gigantea] | 32.1 | 3.0E-08 |
| DIP_10003305 | hypothetical protein ACRE_034940 [Acremonium chrysogenum ATCC 11550] | 38.7 | 2.0E-25 |
| DIP_10003306 | pre-mRNA cleavage complex II protein Clp1, partial [Metarhizium brunneum ARSEF 3297] | 31.4 | 8.0E-42 |
| DIP_10003307 | hypothetical protein THITE_2112596 [Thielavia terrestris NRRL 8126] | 55.7 | 2.0E-167 |
| DIP_10003308 | hypothetical protein BAUCODRAFT_88833 [Baudoinia compniacensis UAMH 10762] | 72.2 | 3.0E-165 |
| DIP_10003309 | hypothetical protein AN8316.2 [Aspergillus nidulans FGSC A4] | 34.9 | 4.0E-109 |
| DIP_10003310 | hypothetical protein PFICI_14635 [Pestalotiopsis fici W106-1] | 48 | 4.0E-79 |
| DIP_10003311 | short-chain dehydrogenase/oxidoreductase [Sporothrix schenckii 1099-18] | 46.4 | 1.0E-78 |
| DIP_10003312 | phosphatidylinositol-4-phosphate 5-kinase its3 (PtdIns(4)P-5-kinase) [Gaeumannomyces graminis var. tritici R3-111a-1] | 79.9 | 0.0 |
| DIP_10003313 | putative oxidoreductase-like protein [Eutypa lata UCREL1] | 73 | 5.0E-57 |
| DIP_10003314 | putative integral membrane protein [Eutypa lata UCREL1] | 74 | 6.0E-80 |
| DIP_10003315 | putative g-patch domain-containing protein [Eutypa lata UCREL1] | 63.4 | 5.0E-157 |
| DIP_10003316 | NA | | |
| DIP_10003317 | Cysteine protease ATG4 [Pestalotiopsis fici W106-1] | 70.4 | 0.0 |
| DIP_10003318 | hypothetical protein UCREL1_3407 [Eutypa lata UCREL1] | 68.6 | 1.0E-154 |
| DIP_10003319 | putative cystathionine gamma-synthase protein [Eutypa lata UCREL1] | 61.7 | 0.0 |
| DIP_10003320 | putative cystathionine gamma-synthase protein [Eutypa lata UCREL1] | 60.5 | 0.0 |
| DIP_10003321 | hypothetical protein FAVG1_09479 [Fusarium avenaceum] | 50.5 | 2.0E-99 |
| DIP_10003322 | putative c6 transcription protein [Neofusicoccum parvum UCRNP2] | 31.3 | 4.0E-56 |
| DIP_10003323 | exo-rhamnogalacturonase b [Colletotrichum gloeosporioides Nara gc5] | 66.5 | 0.0 |
| DIP_10003324 | NA | | |
| DIP_10003325 | eukaryotic and archaeal DNA primase small subunit [Colletotrichum graminicola M1.001] | 64.9 | 0.0 |
| DIP_10003326 | hypothetical protein PFICI_14402 [Pestalotiopsis fici W106-1] | 55.5 | 0.0 |
| DIP_10003327 | putative nadh-ubiquinone oxidoreductase 213 kda subunit protein [Eutypa lata UCREL1] | 65.2 | 6.0E-83 |
| DIP_10003328 | putative ubiquitin carboxyl-terminal hydrolase protein [Eutypa lata UCREL1] | 45.9 | 0.0 |
| DIP_10003329 | putative smad nuclear-interacting protein [Eutypa lata UCREL1] | 78.1 | 4.0E-117 |
| DIP_10003330 | hypothetical protein PFICI_14404 [Pestalotiopsis fici W106-1] | 51.6 | 0.0 |
| DIP_10003331 | hypothetical protein GGTG_05774 [Gaeumannomyces graminis var. tritici R3-111a-1] | 41.2 | 2.0E-132 |
| DIP_10003332 | putative quinone oxidoreductase protein [Eutypa lata UCREL1] | 64.8 | 5.0E-154 |
| DIP_10003333 | hypothetical protein, partial [Streptomyces flavidovirens] | 32.4 | 2.0E-16 |
| DIP_10003334 | hypothetical protein W97_08139 [Coniosporium apollinis CBS 100218] | 39.2 | 2.0E-74 |
| DIP_10003335 | major facilitator superfamily transporter [Colletotrichum graminicola M1.001] | 59.6 | 0.0 |
| DIP_10003336 | hypothetical protein PFICI_14412 [Pestalotiopsis fici W106-1] | 58.7 | 0.0 |
| DIP_10003337 | Fungal specific transcription factor domain protein [Scedosporium apiospermum] | 39 | 3.0E-112 |
| DIP_10003338 | hypothetical protein FVEG_05728 [Fusarium verticillioides 7600] | 39.1 | 8.0E-104 |
| DIP_10003339 | endoglucanase [Colletotrichum higginsianum] | 38.1 | 1.0E-80 |
| DIP_10003340 | hypothetical protein PFICI_12911 [Pestalotiopsis fici W106-1] | 41.5 | 2.0E-76 |
| DIP_10003341 | NA | | |
| DIP_10003342 | putative oxysterol binding protein [Eutypa lata UCREL1] | 77.9 | 0.0 |
| DIP_10003343 | hypothetical protein PFICI_14657 [Pestalotiopsis fici W106-1] | 50.9 | 1.0E-125 |
| DIP_10003344 | hypothetical protein TRIVIDRAFT_34075 [Trichoderma virens Gv29-8] | 30.5 | 3.0E-10 |
| DIP_10003345 | nucleoside hydrolase, partial [Neurospora crassa] | 32.7 | 3.0E-07 |
| DIP_10003346 | hypothetical protein PFICI_03252 [Pestalotiopsis fici W106-1] | 81 | 9.0E-23 |
| DIP_10003347 | hypothetical protein PFICI_11526 [Pestalotiopsis fici W106-1] | 67.2 | 0.0 |
| DIP_10003348 | hypothetical protein PFICI_11527 [Pestalotiopsis fici W106-1] | 70.5 | 0.0 |
| DIP_10003349 | NA | | |
| DIP_10003350 | hypothetical protein MPH_06874 [Macrophomina phaseolina MS6] | 44.9 | 2.0E-45 |
| DIP_10003351 | NA | | |
| DIP_10003352 | putative eukaryotic translation initiation factor 3 subunit e protein [Eutypa lata UCREL1] | 85.1 | 0.0 |
| DIP_10003353 | hypothetical protein PFICI_08290 [Pestalotiopsis fici W106-1] | 39.7 | 1.0E-21 |
| DIP_10003354 | hypothetical protein PV06_02584 [Exophiala oligosperma] | 50.9 | 0.0 |
| DIP_10003355 | hexaprenyl pyrophosphate synthetase [Colletotrichum orbiculare MAFF 240422] | 77.7 | 0.0 |
| DIP_10003356 | hypothetical protein CHGG_09599 [Chaetomium globosum CBS 148.51] | 48.4 | 4.0E-84 |
| DIP_10003357 | putative vacuolar er assembly vma12 protein [Eutypa lata UCREL1] | 33.5 | 3.0E-15 |
| DIP_10003358 | hypothetical protein PFICI_02845 [Pestalotiopsis fici W106-1] | 38.1 | 4.0E-20 |
| DIP_10003359 | putative c6 transcription protein [Eutypa lata UCREL1] | 51.3 | 0.0 |
| DIP_10003360 | hypothetical protein [Streptomyces acidiscabies] | 43.1 | 7.0E-140 |
| DIP_10003361 | NA | | |
| DIP_10003362 | NA | | |
| DIP_10003363 | hypothetical protein Z518_00929 [Rhinocladiella mackenziei CBS 650.93] | 30.7 | 1.0E-108 |
| DIP_10003364 | Pyroglutamyl-peptidase 1 [Ceratocystis platani] | 31 | 4.0E-28 |
| DIP_10003365 | hypothetical protein DRE_00183 [Drechslerella stenobrocha 248] | 39.7 | 1.0E-90 |
| DIP_10003366 | NA | | |
| DIP_10003367 | hypothetical protein PFICI_03160 [Pestalotiopsis fici W106-1] | 39.8 | 4.0E-49 |
| DIP_10003368 | histone H1 [Metarhizium album ARSEF 1941] | 68.3 | 2.0E-41 |
| DIP_10003369 | hypothetical protein THITE_2118118 [Thielavia terrestris NRRL 8126] | 31.4 | 1.0E-19 |
| DIP_10003370 | acid phosphatase [Fusarium oxysporum FOSC 3-a] | 54.5 | 0.0 |
| DIP_10003371 | hypothetical protein MGG_14244 [Magnaporthe oryzae 70-15] | 31.2 | 9.0E-19 |
| DIP_10003372 | putative glucan endo- -beta-glucosidase protein [Eutypa lata UCREL1] | 60.2 | 3.0E-166 |
| DIP_10003373 | NA | | |
| DIP_10003374 | hypothetical protein FGSG_10978 [Fusarium graminearum PH-1] | 31.7 | 3.0E-54 |
| DIP_10003375 | NA | | |
| DIP_10003376 | short-chain dehydrogenase, putative [Talaromyces stipitatus ATCC 10500] | 51.8 | 6.0E-100 |
| DIP_10003377 | stress responsive A/B Barrel domain-containing protein [Colletotrichum gloeosporioides Cg-14] | 51.8 | 9.0E-39 |
| DIP_10003378 | hypothetical protein PFICI_14565 [Pestalotiopsis fici W106-1] | 55.7 | 1.0E-100 |
| DIP_10003379 | hypothetical protein ANI_1_408154 [Aspergillus niger CBS 513.88] | 42.8 | 8.0E-70 |
| DIP_10003380 | Protein PNS1 [Pestalotiopsis fici W106-1] | 80.6 | 0.0 |
| DIP_10003381 | conserved hypothetical protein [Talaromyces stipitatus ATCC 10500] | 46.9 | 7.0E-91 |
| DIP_10003382 | hypothetical protein BofuT4_P036730.1 [Botrytis cinerea T4] | 32.4 | 0.0 |
| DIP_10003383 | hypothetical protein GLRG_09249 [Colletotrichum graminicola M1.001] | 52.4 | 0.0 |
| DIP_10003384 | Altered inheritance of mitochondria protein 24 [Pestalotiopsis fici W106-1] | 60 | 2.0E-144 |
| DIP_10003385 | putative riboflavin transporter mch5 protein [Togninia minima UCRPA7] | 59.4 | 0.0 |
| DIP_10003386 | salicylate hydroxylase [Fusarium oxysporum f. sp. radicis-lycopersici 26381] | 61.4 | 0.0 |
| DIP_10003387 | NA | | |
| DIP_10003388 | putative glycosyl hydrolase family 2 protein [Eutypa lata UCREL1] | 63.2 | 0.0 |
| DIP_10003389 | NA | | |
| DIP_10003390 | kinesin family protein [Colletotrichum gloeosporioides Nara gc5] | 53.1 | 0.0 |
| DIP_10003391 | hypothetical protein PFICI_12567 [Pestalotiopsis fici W106-1] | 48.8 | 5.0E-84 |
| DIP_10003392 | hypothetical protein PFICI_12566 [Pestalotiopsis fici W106-1] | 70 | 0.0 |
| DIP_10003393 | hypothetical protein PFICI_12542 [Pestalotiopsis fici W106-1] | 49.5 | 0.0 |
| DIP_10003394 | putative mfs mcp solute carrier family 16 (monocarboxylic acid transporters) member 10 [Diaporthe [illegible] | 57.7 | 1.0E-175 |
| DIP_10003395 | NA | | |
| DIP_10003396 | hypothetical protein SAPIO_CDS4403 [Scedosporium apiospermum] | 86.7 | 1.0E-158 |
| DIP_10003397 | NA | | |
| DIP_10003398 | putative proline-specific permease protein [Eutypa lata UCREL1] | 74.3 | 0.0 |
| DIP_10003399 | putative glycoside hydrolase family 47 protein [Eutypa lata UCREL1] | 50.3 | 0.0 |
| DIP_10003400 | hypothetical protein CGLO_09667 [Colletotrichum gloeosporioides Cg-14] | 30.6 | 8.0E-29 |
| DIP_10003401 | hypothetical protein PFICI_02620 [Pestalotiopsis fici W106-1] | 58.7 | 1.0E-143 |
| DIP_10003402 | predicted protein [Nectria haematococca mpVI 77-13-4] | 69.8 | 3.0E-90 |
| DIP_10003403 | hypothetical protein HMPREF1624_03044 [Sporothrix schenckii ATCC 58251] | 35.6 | 2.0E-46 |
| DIP_10003404 | IBR domain-containing protein [Colletotrichum fioriniae PJ7] | 73.9 | 0.0 |
| DIP_10003405 | hypothetical protein VHEMI07559 [Torrubiella hemipterigena] | 58 | 0.0 |
| DIP_10003406 | putative translation machinery-associated protein 16 protein [Eutypa lata UCREL1] | 66.5 | 1.0E-90 |
| DIP_10003407 | putative regulatory protein suaprga1 protein [Eutypa lata UCREL1] | 70.6 | 4.0E-144 |
| DIP_10003408 | Endo-1,4-beta-xylanase A [Pestalotiopsis fici W106-1] | 60.6 | 3.0E-96 |
| DIP_10003409 | hypothetical protein PFICI_14720 [Pestalotiopsis fici W106-1] | 45 | 4.0E-144 |
| DIP_10003410 | hypothetical protein PFICI_03841 [Pestalotiopsis fici W106-1] | 39.6 | 3.0E-65 |
| DIP_10003411 | putative glycosyl hydrolase protein [Eutypa lata UCREL1] | 35.3 | 2.0E-48 |
| DIP_10003412 | putative arginase protein [Eutypa lata UCREL1] | 85.6 | 0.0 |
| DIP_10003413 | NA | | |
| DIP_10003414 | hypothetical protein PFICI_14820 [Pestalotiopsis fici W106-1] | 67.2 | 4.0E-87 |
| DIP_10003415 | NA | | |
| DIP_10003416 | hypothetical protein UCREL1_2595 [Eutypa lata UCREL1] | 32.8 | 1.0E-58 |
| DIP_10003417 | putative glycosyltransferase family 90 protein [Eutypa lata UCREL1] | 67.1 | 0.0 |
| DIP_10003418 | hypothetical protein Cob_12900 [Colletotrichum orbiculare MAFF 240422] | 41.6 | 3.0E-132 |
| DIP_10003419 | NA | | |
| DIP_10003420 | hypothetical protein PFICI_14824 [Pestalotiopsis fici W106-1] | 57.7 | 1.0E-84 |
| DIP_10003421 | hypothetical protein PFICI_14825 [Pestalotiopsis fici W106-1] | 54.4 | 1.0E-60 |
| DIP_10003422 | hypothetical protein TRIATDRAFT_50863 [Trichoderma atroviride IMI 206040] | 41.4 | 2.0E-71 |
| DIP_10003423 | hypothetical protein FOIG_07558 [Fusarium oxysporum f. sp. cubense tropical race 4 54006] | 33.5 | 5.0E-128 |
| DIP_10003424 | putative duf1479 domain protein [Eutypa lata UCREL1] | 59.5 | 0.0 |
| DIP_10003425 | NA | | |
| DIP_10003426 | Arylsulfatase [Pestalotiopsis fici W106-1] | 59 | 0.0 |
| DIP_10003427 | hypothetical protein TERG_02714 [Trichophyton rubrum CBS 118892] | 36.2 | 1.0E-39 |
| DIP_10003428 | hypothetical protein CHGG_09397 [Chaetomium globosum CBS 148.51] | 46.5 | 2.0E-139 |
| DIP_10003429 | hypothetical protein EPUS_06736 [Endocarpon pusillum Z07020] | 48.5 | 3.0E-99 |
| DIP_10003430 | FAD-binding protein [Glarea lozoyensis ATCC 20868] | 54.8 | 0.0 |
| DIP_10003431 | NA | | |
| DIP_10003432 | hypothetical protein EPUS_06733 [Endocarpon pusillum Z07020] | 41.5 | 4.0E-85 |
| DIP_10003433 | hypothetical protein PFICI_14206 [Pestalotiopsis fici W106-1] | 51 | 0.0 |
| DIP_10003434 | NA | | |
| DIP_10003435 | putative gnat family protein [Eutypa lata UCREL1] | 30.6 | 2.0E-25 |
| DIP_10003436 | putative arid bright dna binding domain-containing protein [Togninia minima UCRPA7] | 46.4 | 0.0 |
| DIP_10003437 | hypothetical protein OIDMADRAFT_150441 [Oidiodendron maius Zn] | 40 | 2.0E-84 |
| DIP_10003438 | putative u3 small nucleolar rna-associated protein 6 protein [Eutypa lata UCREL1] | 65.1 | 0.0 |
| DIP_10003439 | putative calcofluor white hypersensitive protein [Togninia minima UCRPA7] | 55.6 | 3.0E-152 |
| DIP_10003440 | NA | | |
| DIP_10003441 | hypothetical protein PFICI_03230 [Pestalotiopsis fici W106-1] | 44.1 | 0.0 |
| DIP_10003442 | hypothetical protein PFICI_03229 [Pestalotiopsis fici W106-1] | 62.6 | 0.0 |
| DIP_10003443 | 40S ribosomal protein S20 [Magnaporthe oryzae 70-15] | 83.6 | 3.0E-62 |
| DIP_10003444 | hypothetical protein PFICI_03227 [Pestalotiopsis fici W106-1] | 39.1 | 2.0E-98 |
| DIP_10003445 | capsule polysaccharide biosynthesis protein [Trichoderma harzianum] | 50.5 | 7.0E-138 |
| DIP_10003446 | hypothetical protein PFICI_13450 [Pestalotiopsis fici W106-1] | 31.7 | 7.0E-33 |
| DIP_10003447 | hypothetical protein UCDDA912_g00981 [Diaporthe ampelina] | 49.4 | 2.0E-46 |
| DIP_10003448 | hypothetical protein PFICI_15373 [Pestalotiopsis fici W106-1] | 31 | 4.0E-10 |
| DIP_10003449 | hypothetical protein PFICI_03225 [Pestalotiopsis fici W106-1] | 70.7 | 0.0 |
| DIP_10003450 | hypothetical protein PFICI_11938 [Pestalotiopsis fici W106-1] | 38.1 | 1.0E-76 |
| DIP_10003451 | hypothetical protein ANI_1_572134 [Aspergillus niger CBS 513.88] | 33.7 | 6.0E-42 |
| DIP_10003452 | hypothetical protein ASPNIDRAFT_210217 [Aspergillus niger ATCC 1015] | 44.8 | 0.0 |
| DIP_10003453 | hypothetical protein THITE_2116547 [Thielavia terrestris NRRL 8126] | 57.9 | 0.0 |
| DIP_10003454 | NA | | |
| DIP_10003455 | NA | | |
| DIP_10003456 | Oxygen-dependent choline dehydrogenase-like protein [Acremonium chrysogenum ATCC 11550] | 48.1 | 0.0 |
| DIP_10003457 | Six-bladed beta-propeller, TolB-like protein [Ophiocordyceps sinensis CO18] | 50.8 | 5.0E-140 |
| DIP_10003458 | hypothetical protein MAC_05872 [Metarhizium acridum CQMa 102] | 43.8 | 3.0E-69 |
| DIP_10003459 | putative glycosyltransferase family 1 protein [Eutypa lata UCREL1] | 75.4 | 0.0 |
| DIP_10003460 | putative autophagy-related protein 26 protein [Eutypa lata UCREL1] | 59.5 | 2.0E-158 |
| DIP_10003461 | 60S ribosomal protein L27a (L29) [Cordyceps militaris CM01] | 88.6 | 2.0E-90 |
| DIP_10003462 | hypothetical protein PFICI_03958 [Pestalotiopsis fici W106-1] | 69.7 | 0.0 |
| DIP_10003463 | hypothetical protein PFICI_11605 [Pestalotiopsis fici W106-1] | 44.1 | 8.0E-163 |
| DIP_10003464 | hypothetical protein PFICI_14477 [Pestalotiopsis fici W106-1] | 66.6 | 4.0E-171 |
| DIP_10003465 | NA | | |
| DIP_10003466 | glycosyl hydrolase family 92 [Beauveria bassiana ARSEF 2860] | 57.5 | 0.0 |
| DIP_10003467 | NA | | |
| DIP_10003468 | putative allantoinase protein [Eutypa lata UCREL1] | 64.2 | 0.0 |
| DIP_10003469 | alpha/beta-hydrolase [Aspergillus ruber CBS 135680] | 45.3 | 2.0E-149 |
| DIP_10003470 | NA | | |
| DIP_10003471 | conserved hypothetical protein [Aspergillus flavus NRRL3357] | 50.9 | 5.0E-93 |
| DIP_10003472 | NA | | |
| DIP_10003473 | NA | | |
| DIP_10003474 | Pc24g02580 [Penicillium rubens Wisconsin 54-1255] | 67.6 | 2.0E-114 |
| DIP_10003475 | putative metal homeostatis protein bsd2 protein [Togninia minima UCRPA7] | 61.1 | 2.0E-130 |
| DIP_10003476 | NA | | |
| DIP_10003477 | hypothetical protein PFICI_02238 [Pestalotiopsis fici W106-1] | 64.4 | 4.0E-47 |
| DIP_10003478 | putative tricarboxylate transport protein [Eutypa lata UCREL1] | 81.1 | 8.0E-177 |
| DIP_10003479 | hypothetical protein CFIO01_01759 [Colletotrichum fioriniae PJ7] | 41.4 | 1.0E-123 |
| DIP_10003480 | hypothetical protein AKAW_03634 [Aspergillus kawachii IFO 4308] | 49.3 | 8.0E-29 |
| DIP_10003481 | putative antibiotic biosynthesis monooxygenase protein [Eutypa lata UCREL1] | 31.2 | 2.0E-27 |
| DIP_10003482 | hypothetical protein PFICI_00798 [Pestalotiopsis fici W106-1] | 30.2 | 2.0E-18 |
| DIP_10003483 | high-affinity methionine permease [Magnaporthe oryzae P131] | 73.9 | 0.0 |
| DIP_10003484 | hypothetical protein PFICI_03961 [Pestalotiopsis fici W106-1] | 66 | 0.0 |
| DIP_10003485 | NA | | |
| DIP_10003486 | FAD binding domain-containing protein [Colletotrichum gloeosporioides Cg-14] | 47.2 | 3.0E-120 |
| DIP_10003487 | hypothetical protein UVI_062080 [Ustilaginoidea virens] | 42.4 | 3.0E-50 |
| DIP_10003488 | NA | | |
| DIP_10003489 | hypothetical protein NECHADRAFT_52109 [Nectria haematococca mpVI 77-13-4] | 32.7 | 5.0E-11 |
| DIP_10003490 | hypothetical protein NECHADRAFT_81671 [Nectria haematococca mpVI 77-13-4] | 64.5 | 7.0E-141 |
| DIP_10003491 | hypothetical protein AOCH_001416 [Aspergillus ochraceoroseus] | 36.6 | 1.0E-72 |
| DIP_10003492 | hypothetical protein PV09_00159 [Verruconis gallopava] | 42.2 | 2.0E-77 |
| DIP_10003493 | hypothetical protein FOQG_17299 [Fusarium oxysporum f. sp. raphani 54005] | 46.8 | 4.0E-119 |
| DIP_10003494 | NA | | |
| DIP_10003495 | hypothetical protein NECHADRAFT_101111 [Nectria haematococca mpVI 77-13-4] | 65.2 | 3.0E-70 |
| DIP_10003496 | putative 2-haloalkanoic acid dehalogenase protein [Eutypa lata UCREL1] | 63.9 | 7.0E-113 |
| DIP_10003497 | hypothetical protein CHGG_09212 [Chaetomium globosum CBS 148.51] | 39.3 | 2.0E-64 |
| DIP_10003498 | NA | | |
| DIP_10003499 | hypothetical protein PFICI_10129 [Pestalotiopsis fici W106-1] | 51.2 | 1.0E-100 |
| DIP_10003500 | hypothetical protein PFICI_10128 [Pestalotiopsis fici W106-1] | 48.5 | 0.0 |
| DIP_10003501 | hypothetical protein PFICI_05966 [Pestalotiopsis fici W106-1] | 71.1 | 5.0E-99 |
| DIP_10003502 | p-glycoprotein [Daldinia eschscholtzii] | 58.2 | 0.0 |
| DIP_10003503 | hypothetical protein PFICI_14095 [Pestalotiopsis fici W106-1] | 57.5 | 0.0 |
| DIP_10003504 | putative glutamyl-trna amidotransferase subunit a protein [Eutypa lata UCREL1] | 66.9 | 0.0 |
| DIP_10003505 | hypothetical protein SNOG_03804 [Phaeosphaeria nodorum SN15] | 64.7 | 3.0E-92 |
| DIP_10003506 | hypothetical protein PFICI_10736 [Pestalotiopsis fici W106-1] | 75.7 | 2.0E-141 |
| DIP_10003507 | UbiA prenyltransferase family [Penicillium expansum] | 61.8 | 2.0E-133 |
| DIP_10003508 | hypothetical protein UCREL1_9432 [Eutypa lata UCREL1] | 57.8 | 2.0E-18 |
| DIP_10003509 | conserved hypothetical protein [Aspergillus terreus NIH2624] | 56.4 | 1.0E-105 |
| DIP_10003510 | hypothetical protein THAR02_00545 [Trichoderma harzianum] | 40 | 9.0E-89 |
| DIP_10003511 | putative aaa family protein [Eutypa lata UCREL1] | 58.9 | 0.0 |
| DIP_10003512 | putative thioesterase family protein [Eutypa lata UCREL1] | 69.2 | 2.0E-74 |
| DIP_10003513 | hypothetical protein W97_08373 [Coniosporium apollinis CBS 100218] | 36.1 | 2.0E-17 |
| DIP_10003514 | hypothetical protein PFICI_07055 [Pestalotiopsis fici W106-1] | 73.1 | 0.0 |
| DIP_10003515 | putative lactoylglutathione lyase protein [Eutypa lata UCREL1] | 71.9 | 6.0E-175 |
| DIP_10003516 | hypothetical protein PFICI_07217 [Pestalotiopsis fici W106-1] | 30.9 | 3.0E-43 |
| DIP_10003517 | NA | | |
| DIP_10003518 | Vacuolar calcium ion transporter [Pestalotiopsis fici W106-1] | 65.2 | 0.0 |
| DIP_10003519 | hypothetical protein PFICI_10072 [Pestalotiopsis fici W106-1] | 63 | 0.0 |
| DIP_10003520 | DnaJ domain-containing protein [Colletotrichum graminicola M1.001] | 66.6 | 7.0E-128 |
| DIP_10003521 | DRAP deaminase [Neurospora crassa OR74A] | 70.7 | 2.0E-76 |
| DIP_10003522 | integral membrane protein [Metarhizium album ARSEF 1941] | 52.9 | 2.0E-104 |
| DIP_10003523 | Six-hairpin glycosidase-like protein [Macrophomina phaseolina MS6] | 57.1 | 5.0E-140 |
| DIP_10003524 | hypothetical protein UCREL1_257 [Eutypa lata UCREL1] | 40.7 | 3.0E-22 |
| DIP_10003525 | hypothetical protein PFICI_10090 [Pestalotiopsis fici W106-1] | 64.6 | 0.0 |
| DIP_10003526 | putative qde2-like protein [Eutypa lata UCREL1] | 50 | 0.0 |
| DIP_10003527 | hypothetical protein PFICI_10085 [Pestalotiopsis fici W106-1] | 60 | 1.0E-76 |
| DIP_10003528 | hypothetical protein OOU_Y34scaffold00683g24 [Magnaporthe oryzae Y34] | 54.5 | 7.0E-25 |
| DIP_10003529 | hypothetical protein PFICI_10100 [Pestalotiopsis fici W106-1] | 51.4 | 7.0E-102 |
| DIP_10003530 | hypothetical protein UCREL1_2470 [Eutypa lata UCREL1] | 38.6 | 1.0E-18 |
| DIP_10003531 | hypothetical protein PFICI_10098 [Pestalotiopsis fici W106-1] | 57.1 | 3.0E-102 |
| DIP_10003532 | putative cytochrome b5 protein [Eutypa lata UCREL1] | 82.2 | 4.0E-81 |
| DIP_10003533 | hypothetical protein PFICI_08657 [Pestalotiopsis fici W106-1] | 54.7 | 0.0 |
| DIP_10003534 | NA | | |
| DIP_10003535 | F-actin-capping protein subunit alpha [Pestalotiopsis fici W106-1] | 80.5 | 1.0E-164 |
| DIP_10003536 | transketolase [Colletotrichum graminicola M1.001] | 81 | 0.0 |
| DIP_10003537 | putative glycoside hydrolase family 5 protein [Eutypa lata UCREL1] | 84.1 | 0.0 |
| DIP_10003538 | hypothetical protein PFICI_13914 [Pestalotiopsis fici W106-1] | 70.3 | 0.0 |
| DIP_10003539 | NA | | |
| DIP_10003540 | hypothetical protein PFICI_10093 [Pestalotiopsis fici W106-1] | 46.7 | 0.0 |
| DIP_10003541 | putative riboflavin kinase protein [Eutypa lata UCREL1] | 54.8 | 3.0E-82 |
| DIP_10003542 | NA | | |
| DIP_10003543 | hypothetical protein MYCTH_2307682 [Myceliophthora thermophila ATCC 42464] | 38.4 | 7.0E-112 |
| DIP_10003544 | NA | | |
| DIP_10003545 | hypothetical protein PFICI_14529 [Pestalotiopsis fici W106-1] | 53.6 | 0.0 |
| DIP_10003546 | hypothetical protein NECHADRAFT_54189 [Nectria haematococca mpVI 77-13-4] | 31.4 | 2.0E-66 |
| DIP_10003547 | hypothetical protein UCREL1_2971 [Eutypa lata UCREL1] | 32 | 4.0E-16 |
| DIP_10003548 | NA | | |
| DIP_10003549 | hypothetical protein PFICI_10159 [Pestalotiopsis fici W106-1] | 58.2 | 5.0E-97 |
| DIP_10003550 | NA | | |
| DIP_10003551 | NA | | |
| DIP_10003552 | NA | | |
| DIP_10003553 | putative serine threonine-protein phosphatase 2a activator 1 protein [Eutypa lata UCREL1] | 74.4 | 6.0E-164 |
| DIP_10003554 | MFS multidrug transporter [Trichophyton rubrum CBS 118892] | 62.1 | 0.0 |
| DIP_10003555 | hypothetical protein PFICI_10164 [Pestalotiopsis fici W106-1] | 39.9 | 0.0 |
| DIP_10003556 | putative hsp90-like protein [Colletotrichum sublineola] | 38.5 | 0.0 |
| DIP_10003557 | hypothetical protein V502_04944 [Pseudogymnoascus pannorum VKM F-4520 (FW-2644)] | 47.7 | 5.0E-18 |
| DIP_10003558 | hypothetical protein Cob_01176 [Colletotrichum orbiculare MAFF 240422] | 46.7 | 0.0 |
| DIP_10003559 | putative late embryogenesis abundant protein [Eutypa lata UCREL1] | 75.1 | 0.0 |
| DIP_10003560 | hypothetical protein CSUB01_07047 [Colletotrichum sublineola] | 33.6 | 2.0E-34 |
| DIP_10003561 | hypothetical protein UCREL1_8133 [Eutypa lata UCREL1] | 67.4 | 7.0E-25 |
| DIP_10003562 | hypothetical protein UCREL1_8132 [Eutypa lata UCREL1] | 71.3 | 1.0E-107 |
| DIP_10003563 | hypothetical protein OOU_Y34scaffold00140g79 [Magnaporthe oryzae Y34] | 43 | 2.0E-17 |
| DIP_10003564 | hypothetical protein CH063_00959 [Colletotrichum higginsianum] | 38.2 | 4.0E-87 |
| DIP_10003565 | hypothetical protein PFICI_02321 [Pestalotiopsis fici W106-1] | 35.8 | 5.0E-29 |
| DIP_10003566 | N-terminal fungal transcription regulatory domain-containing protein [Metarhizium robertsii ARSEF 23] | 50.9 | 0.0 |
| DIP_10003567 | putative gpi mannosyltransferase protein [Eutypa lata UCREL1] | 78.2 | 0.0 |
| DIP_10003568 | putative mfs transporter protein [Eutypa lata UCREL1] | 63.1 | 0.0 |
| DIP_10003569 | hypothetical protein PFICI_14202 [Pestalotiopsis fici W106-1] | 34.7 | 4.0E-95 |
| DIP_10003570 | hypothetical protein PFICI_14203 [Pestalotiopsis fici W106-1] | 53.9 | 6.0E-87 |
| DIP_10003571 | YT521-B-like family protein [Metarhizium anisopliae] | 35.9 | 2.0E-28 |
| DIP_10003572 | RecName: Full=Carboxypeptidase S1 [Penicillium janthinellum] | 31.7 | 2.0E-57 |
| DIP_10003573 | NA | | |
| DIP_10003574 | hypothetical protein PFICI_09655 [Pestalotiopsis fici W106-1] | 72.2 | 0.0 |
| DIP_10003575 | hypothetical protein OIDMADRAFT_101801 [Oidiodendron maius Zn] | 61.8 | 3.0E-160 |
| DIP_10003576 | Pectate lyase B [Pestalotiopsis fici W106-1] | 74.1 | 1.0E-154 |
| DIP_10003577 | hypothetical protein S40293_01709 [Stachybotrys chartarum IBT 40293] | 58.5 | 0.0 |
| DIP_10003578 | hypothetical protein PFICI_10246 [Pestalotiopsis fici W106-1] | 58.8 | 1.0E-66 |
| DIP_10003579 | NA | | |
| DIP_10003580 | NA | | |
| DIP_10003581 | beta-galactosidase [Metarhizium anisopliae] | 52.3 | 0.0 |
| DIP_10003582 | Aha1 domain family [Ustilaginoidea virens] | 51.5 | 1.0E-113 |
| DIP_10003583 | hypothetical protein PFICI_10184 [Pestalotiopsis fici W106-1] | 55.2 | 0.0 |
| DIP_10003584 | hypothetical protein PFICI_10179 [Pestalotiopsis fici W106-1] | 69.4 | 2.0E-25 |
| DIP_10003585 | putative vacuolar protein sorting-associated protein 26b-b protein [Eutypa lata UCREL1] | 48 | 3.0E-83 |
| DIP_10003586 | hypothetical protein HMPREF1541_05128 [Cyphellophora europaea CBS 101466] | 54.6 | 5.0E-48 |
| DIP_10003587 | serine protease [Streptomyces showdoensis] | 30 | 1.0E-10 |
| DIP_10003588 | NA | | |
| DIP_10003589 | hypothetical protein PFICI_09811 [Pestalotiopsis fici W106-1] | 56.6 | 4.0E-129 |
| DIP_10003590 | hypothetical protein PFICI_05652 [Pestalotiopsis fici W106-1] | 52.6 | 3.0E-117 |
| DIP_10003591 | putative sulfite reductase flavoprotein component protein [Eutypa lata UCREL1] | 74.5 | 0.0 |
| DIP_10003592 | hypothetical protein PFICI_09810 [Pestalotiopsis fici W106-1] | 55.2 | 1.0E-57 |
| DIP_10003593 | NA | | |
| DIP_10003594 | isotrichodermin C-15 hydroxylase [Trichoderma harzianum] | 54.6 | 0.0 |
| DIP_10003595 | hypothetical protein TRIATDRAFT_299408 [Trichoderma atroviride IMI 206040] | 42.9 | 3.0E-80 |
| DIP_10003596 | hypothetical protein PFICI_14787 [Pestalotiopsis fici W106-1] | 64 | 3.0E-134 |
| DIP_10003597 | hypothetical protein PFICI_13489 [Pestalotiopsis fici W106-1] | 34.2 | 5.0E-81 |
| DIP_10003598 | hypothetical protein TRIATDRAFT_88792 [Trichoderma atroviride IMI 206040] | 45.3 | 2.0E-106 |
| DIP_10003599 | cation-transporting ATPase pacS [Verticillium alfalfae VaMs.102] | 44.3 | 1.0E-155 |
| DIP_10003600 | putative c3hc zinc finger domain-containing protein [Eutypa lata UCREL1] | 60.8 | 0.0 |
| DIP_10003601 | putative 3 -bisphosphate nucleotidase protein [Eutypa lata UCREL1] | 77.1 | 0.0 |
| DIP_10003602 | putative 54s ribosomal protein l6 protein [Eutypa lata UCREL1] | 66.2 | 4.0E-118 |
| DIP_10003603 | NA | | |
| DIP_10003604 | putative transcription factor protein [Eutypa lata UCREL1] | 38.6 | 2.0E-76 |
| DIP_10003605 | putative palmitoyltransferase swf1 protein [Eutypa lata UCREL1] | 71.2 | 0.0 |
| DIP_10003606 | hypothetical protein PFICI_10000 [Pestalotiopsis fici W106-1] | 61 | 1.0E-39 |
| DIP_10003607 | putative rcc1 domain-containing protein [Eutypa lata UCREL1] | 62.6 | 0.0 |
| DIP_10003608 | putative dna polymerase sigma protein [Eutypa lata UCREL1] | 54.2 | 4.0E-153 |
| DIP_10003609 | NA | | |
| DIP_10003610 | hypothetical protein PFICI_04975 [Pestalotiopsis fici W106-1] | 50.2 | 9.0E-178 |
| DIP_10003611 | Serine/threonine-protein kinase MST20 [Pestalotiopsis fici W106-1] | 59.5 | 0.0 |
| DIP_10003612 | NA | | |
| DIP_10003613 | hypothetical protein PFICI_09819 [Pestalotiopsis fici W106-1] | 59.8 | 0.0 |
| DIP_10003614 | NA | | |
| DIP_10003615 | NA | | |
| DIP_10003616 | hypothetical protein PFICI_14340 [Pestalotiopsis fici W106-1] | 77.2 | 0.0 |
| DIP_10003617 | hypothetical protein PFICI_06371 [Pestalotiopsis fici W106-1] | 55.3 | 9.0E-100 |
| DIP_10003618 | hypothetical protein PFICI_14360 [Pestalotiopsis fici W106-1] | 44.8 | 0.0 |
| DIP_10003619 | NA | | |
| DIP_10003620 | hypothetical protein PFICI_10254 [Pestalotiopsis fici W106-1] | 75.1 | 0.0 |
| DIP_10003621 | RNA-binding protein Luc7-like 2 [Rhodosporidium toruloides NP11] | 42.7 | 1.0E-64 |
| DIP_10003622 | NA | | |
| DIP_10003623 | S-methyl-5'-thioadenosine phosphorylase [Pestalotiopsis fici W106-1] | 79.1 | 0.0 |
| DIP_10003624 | het domain-containing protein [Colletotrichum orbiculare MAFF 240422] | 42.5 | 6.0E-171 |
| DIP_10003625 | hypothetical protein PFICI_14361 [Pestalotiopsis fici W106-1] | 64 | 0.0 |
| DIP_10003626 | NA | | |
| DIP_10003627 | putative arid/bright DNA binding domain-containing protein [Colletotrichum sublineola] | 36.8 | 2.0E-176 |
| DIP_10003628 | hypothetical protein PFICI_09822 [Pestalotiopsis fici W106-1] | 41.8 | 3.0E-172 |
| DIP_10003629 | putative wd repeat-containing protein [Eutypa lata UCREL1] | 71 | 0.0 |
| DIP_10003630 | hypothetical protein VDAG_05540 [Verticillium dahliae VdLs.17] | 65.3 | 1.0E-39 |
| DIP_10003631 | NA | | |
| DIP_10003632 | hypothetical protein GGTG_00874 [Gaeumannomyces graminis var. tritici R3-111a-1] | 81 | 2.0E-27 |
| DIP_10003633 | putative mannose-1-phosphate guanyltransferase protein [Eutypa lata UCREL1] | 85.1 | 0.0 |
| DIP_10003634 | putative integral membrane protein [Eutypa lata UCREL1] | 36.7 | 3.0E-52 |
| DIP_10003635 | hypothetical protein THITE_2145189 [Thielavia terrestris NRRL 8126] | 44.4 | 2.0E-120 |
| DIP_10003636 | cytochrome P450, partial [Metarhizium majus ARSEF 297] | 45.2 | 1.0E-130 |
| DIP_10003637 | putative ent-kaurene synthase protein [Eutypa lata UCREL1] | 48.9 | 0.0 |
| DIP_10003638 | putative beta-glucosidase M [Pestalotiopsis fici W106-1] | 60.3 | 0.0 |
| DIP_10003639 | NA | | |
| DIP_10003640 | hypothetical protein A1O1_04970 [Capronia coronata CBS 617.96] | 31.3 | 3.0E-46 |
| DIP_10003641 | putative fumarylacetoacetase protein [Eutypa lata UCREL1] | 61.4 | 0.0 |
| DIP_10003642 | hypothetical protein PFICI_15217 [Pestalotiopsis fici W106-1] | 49.3 | 0.0 |
| DIP_10003643 | hypothetical protein PFICI_10126 [Pestalotiopsis fici W106-1] | 71.7 | 0.0 |
| DIP_10003644 | hypothetical protein PFICI_06822 [Pestalotiopsis fici W106-1] | 49.3 | 2.0E-51 |
| DIP_10003645 | hypothetical protein V490_07334 [Pseudogymnoascus pannorum VKM F-3557] | 79 | 2.0E-56 |
| DIP_10003646 | hypothetical protein PFICI_06824 [Pestalotiopsis fici W106-1] | 68.6 | 1.0E-174 |
| DIP_10003647 | 40S ribosomal protein S2 [Neurospora crassa OR74A] | 94.2 | 4.0E-146 |
| DIP_10003648 | NA | | |
| DIP_10003649 | NA | | |
| DIP_10003650 | NA | | |
| DIP_10003651 | hypothetical protein PFICI_06790 [Pestalotiopsis fici W106-1] | 65.9 | 8.0E-125 |
| DIP_10003652 | putative duf52 domain-containing protein [Togninia minima UCRPA7] | 55.5 | 1.0E-130 |
| DIP_10003653 | putative T-complex protein 1 [Colletotrichum sublineola] | 91.1 | 0.0 |
| DIP_10003654 | putative hect domain-containing protein [Eutypa lata UCREL1] | 72 | 5.0E-151 |
| DIP_10003655 | hypothetical protein PFICI_10149 [Pestalotiopsis fici W106-1] | 70 | 0.0 |
| DIP_10003656 | putative long-chain-fatty-acid- ligase protein [Eutypa lata UCREL1] | 67.5 | 0.0 |
| DIP_10003657 | hypothetical protein PFICI_10136 [Pestalotiopsis fici W106-1] | 67.6 | 0.0 |

| | | | |
|---|---|---|---|
| DIP_10003658 | putative VHS domain-containing protein [Colletotrichum sublineola] | 71.4 | 0.0 |
| DIP_10003659 | ubiquitin carboxyl-terminal hydrolase [Beauveria bassiana ARSEF 2860] | 31 | 1.0E-126 |
| DIP_10003660 | putative glucose-regulated protein [Eutypa lata UCREL1] | 84.3 | 0.0 |
| DIP_10003661 | hypothetical protein OIDMADRAFT_172370, partial [Oidiodendron maius Zn] | 37.6 | 7.0E-81 |
| DIP_10003662 | putative integral membrane protein [Colletotrichum sublineola] | 33.9 | 4.0E-70 |
| DIP_10003663 | putative mutator-like element protein [Eutypa lata UCREL1] | 61.7 | 1.0E-106 |
| DIP_10003664 | putative ribose-phosphate pyrophosphokinase protein [Togninia minima UCRPA7] | 72.4 | 0.0 |
| DIP_10003665 | putative formamidopyrimidine-dna glycosylase protein [Eutypa lata UCREL1] | 63.1 | 3.0E-174 |
| DIP_10003666 | hypothetical protein MYCTH_2307131 [Myceliophthora thermophila ATCC 42464] | 75.5 | 0.0 |
| DIP_10003667 | putative methionyl-trna formyltransferase protein [Eutypa lata UCREL1] | 34.8 | 2.0E-54 |
| DIP_10003668 | hypothetical protein PFICI_08447 [Pestalotiopsis fici W106-1] | 42.2 | 3.0E-114 |
| DIP_10003669 | hypothetical protein CGLO_08547 [Colletotrichum gloeosporioides Cg-14] | 45.5 | 5.0E-28 |
| DIP_10003670 | hypothetical protein PFICI_10335 [Pestalotiopsis fici W106-1] | 51.3 | 0.0 |
| DIP_10003671 | putative purine nucleoside protein [Togninia minima UCRPA7] | 54.1 | 7.0E-150 |
| DIP_10003672 | hypothetical protein PFICI_15171 [Pestalotiopsis fici W106-1] | 78.9 | 3.0E-152 |
| DIP_10003673 | hypothetical protein PFICI_15170 [Pestalotiopsis fici W106-1] | 56 | 0.0 |
| DIP_10003674 | putative calcium-dependent protein kinase protein [Eutypa lata UCREL1] | 60.6 | 0.0 |
| DIP_10003675 | benzoate 4-monooxygenase cytochrome P450 [Neosartorya fischeri NRRL 181] | 55.8 | 0.0 |
| DIP_10003676 | hypothetical protein NFIA_055050 [Neosartorya fischeri NRRL 181] | 40.2 | 9.0E-88 |
| DIP_10003677 | hypothetical protein NECHADRAFT_46272 [Nectria haematococca mpVI 77-13-4] | 48.8 | 9.0E-113 |
| DIP_10003678 | hypothetical protein [Podospora anserina S mat+] | 35 | 2.0E-97 |
| DIP_10003679 | unnamed protein product [Fusarium sp. FIESC_5 CS3069] | 41.3 | 6.0E-127 |
| DIP_10003680 | putative duf757 domain-containing protein [Eutypa lata UCREL1] | 66.2 | 3.0E-67 |
| DIP_10003681 | putative mfs multidrug transporter protein [Eutypa lata UCREL1] | 62.6 | 0.0 |
| DIP_10003682 | NA | | |
| DIP_10003683 | hypothetical protein H072_1314 [Dactylellina haptotyla CBS 200.50] | 30.3 | 5.0E-20 |
| DIP_10003684 | NA | | |
| DIP_10003685 | NA | | |
| DIP_10003686 | hypothetical protein MVEG_01292 [Mortierella verticillata NRRL 6337] | 53 | 6.0E-95 |
| DIP_10003687 | NA | | |
| DIP_10003688 | putative mfs transporter protein [Eutypa lata UCREL1] | 58.7 | 0.0 |
| DIP_10003689 | putative pectinesterase [Diaporthe ampelina] | 61.2 | 9.0E-130 |
| DIP_10003690 | hypothetical protein S40285_05484 [Stachybotrys chlorohalonata IBT 40285] | 65.2 | 0.0 |
| DIP_10003691 | hypothetical protein CMQ_3319 [Grosmannia clavigera kw1407] | 43.3 | 2.0E-79 |
| DIP_10003692 | TfdA family Taurine catabolism dioxygenase TauD [Colletotrichum graminicola M1.001] | 68.9 | 0.0 |
| DIP_10003693 | putative tetracenomycin polyketide synthesis o-methyltransferase tcmp protein [Eutypa lata UCREL1] | 50.5 | 3.0E-96 |
| DIP_10003694 | artemis protein [Colletotrichum fioriniae PJ7] | 43.9 | 3.0E-171 |
| DIP_10003695 | putative phospholipase c protein [Eutypa lata UCREL1] | 49 | 0.0 |
| DIP_10003696 | NA | | |
| DIP_10003697 | NA | | |
| DIP_10003698 | putative transcription initiation protein spt5 protein [Eutypa lata UCREL1] | 73.7 | 0.0 |
| DIP_10003699 | hypothetical protein PFICI_15162 [Pestalotiopsis fici W106-1] | 72 | 0.0 |
| DIP_10003700 | putative pantetheine-phosphate adenylyltransferase family protein [Eutypa lata UCREL1] | 62.1 | 2.0E-169 |
| DIP_10003701 | putative short chain dehydrogenase reductase family protein [Eutypa lata UCREL1] | 64.7 | 6.0E-114 |
| DIP_10003702 | putative autophagy-related protein 28 protein [Eutypa lata UCREL1] | 32.9 | 4.0E-84 |
| DIP_10003703 | Protein of unknown function DUF1748 [Beauveria bassiana ARSEF 2860] | 76.7 | 9.0E-46 |
| DIP_10003704 | pyruvate kinase [Colletotrichum orbiculare MAFF 240422] | 80.3 | 0.0 |
| DIP_10003705 | hypothetical protein [Podospora anserina S mat+] | 30.4 | 2.0E-18 |
| DIP_10003706 | integral membrane protein [Colletotrichum orbiculare MAFF 240422] | 47.1 | 1.0E-83 |
| DIP_10003707 | hypothetical protein PFICI_10336 [Pestalotiopsis fici W106-1] | 46 | 6.0E-99 |
| DIP_10003708 | putative hec ndc80p family protein [Eutypa lata UCREL1] | 80.9 | 0.0 |
| DIP_10003709 | brct domain-containing protein [Colletotrichum orbiculare MAFF 240422] | 49.9 | 0.0 |
| DIP_10003710 | hypothetical protein FPSE_09627 [Fusarium pseudograminearum CS3096] | 48.4 | 0.0 |
| DIP_10003711 | NA | | |
| DIP_10003712 | NA | | |
| DIP_10003713 | Putative Calcineurin-like phosphoesterase [Torrubiella hemipterigena] | 67.7 | 0.0 |
| DIP_10003714 | hypothetical protein GGTG_07409 [Gaeumannomyces graminis var. tritici R3-111a-1] | 65 | 1.0E-144 |
| DIP_10003715 | hypothetical protein PFICI_10281 [Pestalotiopsis fici W106-1] | 82.2 | 0.0 |
| DIP_10003716 | putative e3 ubiquitin-protein ligase upl3 protein [Eutypa lata UCREL1] | 73.3 | 0.0 |
| DIP_10003717 | hypothetical protein UCREL1_1506 [Eutypa lata UCREL1] | 40.8 | 1.0E-152 |
| DIP_10003718 | putative toxin biosynthesis protein [Eutypa lata UCREL1] | 69 | 0.0 |
| DIP_10003719 | NADH-ubiquinone oxidoreductase subunit A [Pestalotiopsis fici W106-1] | 83.6 | 3.0E-177 |
| DIP_10003720 | hypothetical protein PFICI_10278 [Pestalotiopsis fici W106-1] | 51.2 | 4.0E-144 |
| DIP_10003721 | hypothetical protein PFICI_10277 [Pestalotiopsis fici W106-1] | 64.6 | 2.0E-95 |
| DIP_10003722 | hypothetical protein UCREL1_1501 [Eutypa lata UCREL1] | 58.2 | 7.0E-74 |
| DIP_10003723 | hypothetical protein CSUB01_03805 [Colletotrichum sublineola] | 57.6 | 8.0E-158 |
| DIP_10003724 | hypothetical protein PFICI_10267 [Pestalotiopsis fici W106-1] | 47.2 | 0.0 |
| DIP_10003725 | NA | | |
| DIP_10003726 | hypothetical protein PFICI_04176 [Pestalotiopsis fici W106-1] | 60.4 | 0.0 |
| DIP_10003727 | NA | | |
| DIP_10003728 | putative nicotinamide mononucleotide adenylyl transferase protein [Eutypa lata UCREL1] | 79.2 | 1.0E-162 |
| DIP_10003729 | putative patched sphingolipid transporter protein [Eutypa lata UCREL1] | 81.5 | 0.0 |
| DIP_10003730 | hypothetical protein PFICI_10262 [Pestalotiopsis fici W106-1] | 37.9 | 9.0E-83 |
| DIP_10003731 | putative nape-hydrolyzing phospholipase d protein [Eutypa lata UCREL1] | 60.4 | 6.0E-176 |
| DIP_10003732 | NA | | |
| DIP_10003733 | hypothetical protein CGLO_04597 [Colletotrichum gloeosporioides Cg-14] | 51.4 | 2.0E-171 |
| DIP_10003734 | methylene-fatty-acyl-phospholipid synthase [Ophiostoma piceae UAMH 11346] | 71.4 | 2.0E-94 |
| DIP_10003735 | putative transporter smf2 protein [Eutypa lata UCREL1] | 68 | 0.0 |
| DIP_10003736 | hypothetical protein CGGC5_3089 [Colletotrichum gloeosporioides Nara gc5] | 31.9 | 5.0E-32 |
| DIP_10003737 | hypothetical protein PFICI_14188 [Pestalotiopsis fici W106-1] | 68.6 | 0.0 |
| DIP_10003738 | NA | | |
| DIP_10003739 | NA | | |
| DIP_10003740 | Mannose-1-phosphate guanyltransferase [Pestalotiopsis fici W106-1] | 87.4 | 0.0 |
| DIP_10003741 | NA | | |
| DIP_10003742 | hypothetical protein VDAG_05600 [Verticillium dahliae VdLs.17] | 78.6 | 6.0E-65 |
| DIP_10003743 | NA | | |
| DIP_10003744 | prenyltransferase and squalene oxidase [Colletotrichum fioriniae PJ7] | 67 | 5.0E-166 |
| DIP_10003745 | NA | | |
| DIP_10003746 | putative c6 finger domain-containing protein [Eutypa lata UCREL1] | 73.9 | 0.0 |
| DIP_10003747 | ubiquitin [Metarhizium acridum CQMa 102] | 100 | 0.0 |
| DIP_10003748 | unnamed protein product [Fusarium sp. FIESC_5 CS3069] | 44.8 | 8.0E-178 |
| DIP_10003749 | putative glycerol-3-phosphate o-acyltransferase protein [Eutypa lata UCREL1] | 67 | 0.0 |
| DIP_10003750 | related to aminopeptidase Y precursor, vacuolar [Sporisorium reilianum SRZ2] | 33.7 | 5.0E-30 |
| DIP_10003751 | putative sucrase ferredoxin domain-containing protein [Eutypa lata UCREL1] | 67.6 | 4.0E-175 |
| DIP_10003752 | acetyl-hydrolase-like protein [Magnaporthe grisea] | 43.7 | 2.0E-146 |
| DIP_10003753 | putative 24-dehydrocholesterol reductase protein [Eutypa lata UCREL1] | 69.6 | 0.0 |
| DIP_10003754 | triacylglycerol lipase v precursor [Fusarium avenaceum] | 44.4 | 3.0E-163 |
| DIP_10003755 | hypothetical protein [Podospora anserina S mat+] | 81.2 | 0.0 |
| DIP_10003756 | hypothetical protein GLRG_08767 [Colletotrichum graminicola M1.001] | 56.6 | 0.0 |
| DIP_10003757 | NA | | |
| DIP_10003758 | NA | | |
| DIP_10003759 | hypothetical protein COCVIDRAFT_29706 [Bipolaris victoriae FI3] | 73.4 | 0.0 |
| DIP_10003760 | hypothetical protein PFICI_14170 [Pestalotiopsis fici W106-1] | 44.5 | 1.0E-117 |
| DIP_10003761 | hypothetical protein ACRE_050920 [Acremonium chrysogenum ATCC 11550] | 48.6 | 2.0E-87 |
| DIP_10003762 | putative homoserine dehydrogenase protein [Eutypa lata UCREL1] | 80.6 | 0.0 |
| DIP_10003763 | NA | | |
| DIP_10003764 | putative 3-ketosteroid reductase protein [Eutypa lata UCREL1] | 54.3 | 8.0E-177 |
| DIP_10003765 | major facilitator superfamily transporter [Ophiostoma piceae UAMH 11346] | 49.3 | 3.0E-159 |
| DIP_10003766 | ubiquinol cytochrome C reductase hinge protein [Epichloe typhina] | 52.8 | 2.0E-44 |
| DIP_10003767 | NA | | |
| DIP_10003768 | hypothetical protein PFICI_14177 [Pestalotiopsis fici W106-1] | 59.3 | 0.0 |
| DIP_10003769 | putative trna (guanine-n -)-methyltransferase protein [Eutypa lata UCREL1] | 70.6 | 2.0E-153 |
| DIP_10003770 | putative intermediate filament protein [Eutypa lata UCREL1] | 84.4 | 0.0 |
| DIP_10003771 | hypothetical protein S7711_03256 [Stachybotrys chartarum IBT 7711] | 78.7 | 2.0E-158 |
| DIP_10003772 | putative glutathione s-transferase ii protein [Togninia minima UCRPA7] | 53.9 | 2.0E-111 |
| DIP_10003773 | hypothetical protein NECHADRAFT_77217 [Nectria haematococca mpVI 77-13-4] | 30.3 | 5.0E-60 |
| DIP_10003774 | hypothetical protein VHEMI05798 [Torrubiella hemipterigena] | 34.5 | 4.0E-25 |
| DIP_10003775 | putative homogentisate -dioxygenase protein [Eutypa lata UCREL1] | 61.7 | 2.0E-68 |
| DIP_10003776 | von willebrand factor, partial [Colletotrichum gloeosporioides Nara gc5] | 30.8 | 3.0E-11 |
| DIP_10003777 | hypothetical protein CGLO_02743 [Colletotrichum gloeosporioides Cg-14] | 52.1 | 0.0 |
| DIP_10003778 | hypothetical protein PDIP_40520 [Penicillium digitatum Pd1] | 48.3 | 7.0E-46 |
| DIP_10003779 | putative homogentisate -dioxygenase protein [Eutypa lata UCREL1] | 83.7 | 0.0 |
| DIP_10003780 | oxidoreductase [Colletotrichum orbiculare MAFF 240422] | 44.7 | 2.0E-107 |
| DIP_10003781 | putative phd finger domain protein [Eutypa lata UCREL1] | 63.9 | 0.0 |
| DIP_10003782 | putative proteasome component y13 protein [Eutypa lata UCREL1] | 95.6 | 2.0E-178 |
| DIP_10003783 | NA | | |
| DIP_10003784 | NA | | |
| DIP_10003785 | hypothetical protein UCREL1_6646 [Eutypa lata UCREL1] | 81.3 | 2.0E-43 |
| DIP_10003786 | putative duf833 domain-containing protein [Eutypa lata UCREL1] | 57.7 | 2.0E-110 |
| DIP_10003787 | putative rna polymerase ii transcription factor b subunit 2 protein [Togninia minima UCRPA7] | 62.5 | 0.0 |
| DIP_10003788 | NA | | |
| DIP_10003789 | NA | | |
| DIP_10003790 | NA | | |
| DIP_10003791 | Copper amine oxidase 1 [Pestalotiopsis fici W106-1] | 74.9 | 0.0 |
| DIP_10003792 | hypothetical protein PFICI_11210 [Pestalotiopsis fici W106-1] | 49.3 | 3.0E-178 |
| DIP_10003793 | ubiquitin-conjugating enzyme [Colletotrichum higginsianum] | 79.1 | 7.0E-156 |
| DIP_10003794 | hypothetical protein PFICI_15185 [Pestalotiopsis fici W106-1] | 65.2 | 0.0 |
| DIP_10003795 | putative glycoside hydrolase protein [Eutypa lata UCREL1] | 45.2 | 2.0E-61 |
| DIP_10003796 | 3-methyl-2-oxobutanoate hydroxymethyltransferase [Gaeumannomyces graminis var. tritici R3-111a-1] | 71 | 2.0E-165 |
| DIP_10003797 | hypothetical protein PFICI_09837 [Pestalotiopsis fici W106-1] | 52.6 | 8.0E-46 |
| DIP_10003798 | putative g1 s regulator protein [Eutypa lata UCREL1] | 55.3 | 0.0 |
| DIP_10003799 | NA | | |
| DIP_10003800 | putative metallopeptidase family m24 protein [Eutypa lata UCREL1] | 72.4 | 0.0 |
| DIP_10003801 | putative helix-loop-helix dna-binding domain-containing protein [Eutypa lata UCREL1] | 46.8 | 2.0E-49 |
| DIP_10003802 | NA | | |
| DIP_10003803 | putative beta- -glucanase protein [Eutypa lata UCREL1] | 63.2 | 0.0 |
| DIP_10003804 | hypothetical protein FOXB_04827 [Fusarium oxysporum Fo5176] | 46.5 | 4.0E-141 |
| DIP_10003805 | hypothetical protein PV09_04479 [Verruconis gallopava] | 78 | 0.0 |
| DIP_10003806 | hypothetical protein UCREL1_3952 [Eutypa lata UCREL1] | 38 | 2.0E-139 |
| DIP_10003807 | hypothetical protein A1O7_07034 [Cladophialophora yegresii CBS 114405] | 46.4 | 8.0E-49 |
| DIP_10003808 | NA | | |
| DIP_10003809 | hypothetical protein PFICI_09995 [Pestalotiopsis fici W106-1] | 43.1 | 1.0E-94 |
| DIP_10003810 | aspartylglucosaminidase [Gaeumannomyces graminis var. tritici R3-111a-1] | 61.1 | 2.0E-136 |
| DIP_10003811 | hypothetical protein UCREL1_2595 [Eutypa lata UCREL1] | 31.2 | 5.0E-53 |
| DIP_10003812 | putative neutral amino acid permease protein [Eutypa lata UCREL1] | 46.4 | 0.0 |
| DIP_10003813 | hypothetical protein PFICI_03071 [Pestalotiopsis fici W106-1] | 66.3 | 0.0 |
| DIP_10003814 | hypothetical protein UCREL1_3436 [Eutypa lata UCREL1] | 51.3 | 7.0E-78 |
| DIP_10003815 | putative mitochondrial import inner membrane translocase subunit tim16 protein [Eutypa lata UCREL1] | 72 | 2.0E-70 |
| DIP_10003816 | hypothetical protein PFICI_10056 [Pestalotiopsis fici W106-1] | 69.1 | 3.0E-130 |
| DIP_10003817 | NA | | |
| DIP_10003818 | hypothetical protein FVEG_09997 [Fusarium verticillioides 7600] | 57.9 | 1.0E-62 |
| DIP_10003819 | hypothetical protein UCREL1_3397 [Eutypa lata UCREL1] | 38.4 | 1.0E-112 |
| DIP_10003820 | hypothetical protein PFICI_06217 [Pestalotiopsis fici W106-1] | 43 | 2.0E-61 |
| DIP_10003821 | hypothetical protein PFICI_14370 [Pestalotiopsis fici W106-1] | 62.4 | 0.0 |
| DIP_10003822 | putative laccase protein [Eutypa lata UCREL1] | 54.6 | 0.0 |
| DIP_10003823 | NA | | |
| DIP_10003824 | NA | | |
| DIP_10003825 | hypothetical protein THITE_2038523 [Thielavia terrestris NRRL 8126] | 68.8 | 0.0 |
| DIP_10003826 | hypothetical protein THITE_70175 [Thielavia terrestris NRRL 8126] | 67.4 | 8.0E-169 |
| DIP_10003827 | hypothetical protein TCE0_013f00909 [Talaromyces cellulolyticus] | 47.2 | 2.0E-123 |
| DIP_10003828 | subtilase [Colletotrichum fioriniae PJ7] | 54.4 | 5.0E-132 |
| DIP_10003829 | NA | | |
| DIP_10003830 | NA | | |
| DIP_10003831 | putative amidohydrolase family protein [Eutypa lata UCREL1] | 59.3 | 0.0 |
| DIP_10003832 | hypothetical protein PFICI_05415 [Pestalotiopsis fici W106-1] | 54 | 0.0 |
| DIP_10003833 | putative sugar transporter protein [Eutypa lata UCREL1] | 66.4 | 0.0 |
| DIP_10003834 | putative d-xylulose reductase a protein [Eutypa lata UCREL1] | 74 | 0.0 |
| DIP_10003835 | hypothetical protein PFICI_05410 [Pestalotiopsis fici W106-1] | 58.6 | 3.0E-108 |
| DIP_10003836 | NA | | |
| DIP_10003837 | putative hva22 family tb2 dp1 protein [Eutypa lata UCREL1] | 59.1 | 3.0E-141 |
| DIP_10003838 | hypothetical protein V492_01557 [Pseudogymnoascus pannorum VKM F-4246] | 48.3 | 6.0E-103 |
| DIP_10003839 | putative biotin-protein ligase ligase protein [Eutypa lata UCREL1] | 66.7 | 0.0 |
| DIP_10003840 | NA | | |
| DIP_10003841 | putative carboxymuconolactone decarboxylase protein [Eutypa lata UCREL1] | 61.1 | 8.0E-110 |
| DIP_10003842 | hypothetical protein HMPREF1541_05620 [Cyphellophora europaea CBS 101466] | 62.5 | 0.0 |
| DIP_10003843 | hypothetical protein PFICI_03603 [Pestalotiopsis fici W106-1] | 57.4 | 0.0 |
| DIP_10003844 | short chain dehydrogenase/reductase [Sporothrix brasiliensis 5110] | 58.4 | 5.0E-114 |
| DIP_10003845 | hypothetical protein PFICI_10044 [Pestalotiopsis fici W106-1] | 54 | 4.0E-79 |
| DIP_10003846 | cellulose-binding gdsl lipase [Colletotrichum orbiculare MAFF 240422] | 39.4 | 6.0E-73 |
| DIP_10003847 | hypothetical protein TRIVIDRAFT_54649 [Trichoderma virens Gv29-8] | 54.8 | 0.0 |
| DIP_10003848 | putative peptidase family t4 protein [Eutypa lata UCREL1] | 56.9 | 1.0E-164 |
| DIP_10003849 | hypothetical protein PFICI_09817 [Pestalotiopsis fici W106-1] | 75.3 | 0.0 |
| DIP_10003850 | alkaline serine protease Alp1 [Colletotrichum fioriniae PJ7] | 47.3 | 2.0E-96 |
| DIP_10003851 | NA | | |
| DIP_10003852 | Fructose-1,6-bisphosphatase [Pestalotiopsis fici W106-1] | 84.8 | 0.0 |
| DIP_10003853 | hypothetical protein GLRG_08757 [Colletotrichum graminicola M1.001] | 63.2 | 2.0E-151 |
| DIP_10003854 | hypothetical protein PV06_02602 [Exophiala oligosperma] | 63.2 | 0.0 |
| DIP_10003855 | NA | | |
| DIP_10003856 | hypothetical protein PFICI_01129 [Pestalotiopsis fici W106-1] | 53.4 | 4.0E-112 |
| DIP_10003857 | putative s-adenosylmethionine-dependent methyltransferase protein [Eutypa lata UCREL1] | 39.7 | 4.0E-89 |
| DIP_10003858 | putative glycoside hydrolase family 47 protein [Eutypa lata UCREL1] | 62.6 | 0.0 |
| DIP_10003859 | hypothetical protein PFICI_14350 [Pestalotiopsis fici W106-1] | 67.1 | 9.0E-172 |
| DIP_10003860 | putative spermine spermidine synthase protein [Eutypa lata UCREL1] | 58.7 | 0.0 |
| DIP_10003861 | NA | | |
| DIP_10003862 | protoheme IX farnesyltransferase [Gaeumannomyces graminis var. tritici R3-111a-1] | 55.5 | 1.0E-154 |
| DIP_10003863 | myb family transcription factor [Colletotrichum gloeosporioides Nara gc5] | 41.8 | 6.0E-38 |
| DIP_10003864 | hypothetical protein THAR02_08632 [Trichoderma harzianum] | 38.6 | 1.0E-70 |
| DIP_10003865 | hypothetical protein PFICI_10068 [Pestalotiopsis fici W106-1] | 43.2 | 1.0E-158 |
| DIP_10003866 | NA | | |
| DIP_10003867 | putative dna mismatch repair protein msh6 protein [Eutypa lata UCREL1] | 82.9 | 0.0 |
| DIP_10003868 | hypothetical protein PFICI_10370 [Pestalotiopsis fici W106-1] | 57.9 | 0.0 |
| DIP_10003869 | hypothetical protein UCREL1_11076 [Eutypa lata UCREL1] | 56.7 | 8.0E-167 |
| DIP_10003870 | NA | | |
| DIP_10003871 | hypothetical protein PFICI_13466 [Pestalotiopsis fici W106-1] | 70.3 | 0.0 |
| DIP_10003872 | putative long-chain fatty acid protein [Eutypa lata UCREL1] | 59 | 0.0 |
| DIP_10003873 | putative diphthamide biosynthesis protein 2 protein [Eutypa lata UCREL1] | 57.6 | 0.0 |
| DIP_10003874 | putative paf acetylhydrolase family protein [Eutypa lata UCREL1] | 42.7 | 7.0E-88 |
| DIP_10003875 | NA | | |
| DIP_10003876 | hypothetical protein PFICI_13752 [Pestalotiopsis fici W106-1] | 70 | 2.0E-84 |
| DIP_10003877 | NA | | |
| DIP_10003878 | NA | | |
| DIP_10003879 | hypothetical protein UCREL1_5513 [Eutypa lata UCREL1] | 30.4 | 4.0E-50 |
| DIP_10003880 | hypothetical protein PFICI_10226 [Pestalotiopsis fici W106-1] | 69.3 | 0.0 |
| DIP_10003881 | cross-pathway control protein 1 [Magnaporthe oryzae Y34] | 48.3 | 6.0E-88 |
| DIP_10003882 | putative abc heavy metal transporter [Diaporthe ampelina] | 70 | 0.0 |
| DIP_10003883 | NA | | |
| DIP_10003884 | hypothetical protein, partial [Ktedonobacter racemifer] | 32 | 7.0E-08 |
| DIP_10003885 | hypothetical protein V495_06827 [Pseudogymnoascus pannorum VKM F-4514 (FW-929)] | 37.6 | 2.0E-26 |
| DIP_10003886 | hypothetical protein PFICI_14161 [Pestalotiopsis fici W106-1] | 44.5 | 0.0 |
| DIP_10003887 | NA | | |
| DIP_10003888 | putative abc transporter protein [Eutypa lata UCREL1] | 88.3 | 0.0 |
| DIP_10003889 | hypothetical protein G647_03136 [Cladophialophora carrionii CBS 160.54] | 49.8 | 7.0E-55 |
| DIP_10003890 | NA | | |
| DIP_10003891 | tubulin beta-2 chain [Colletotrichum fioriniae PJ7] | 98.7 | 0.0 |
| DIP_10003892 | nitrite reductase [Magnaporthe oryzae 70-15] | 53.7 | 0.0 |
| DIP_10003893 | NA | | |
| DIP_10003894 | hypothetical protein PFICI_14166 [Pestalotiopsis fici W106-1] | 76.9 | 0.0 |
| DIP_10003895 | hypothetical protein PFICI_10263 [Pestalotiopsis fici W106-1] | 53 | 2.0E-159 |
| DIP_10003896 | putative bsd domain-containing protein [Eutypa lata UCREL1] | 60.4 | 2.0E-177 |
| DIP_10003897 | putative pre-rrna-processing protein tsr2 protein [Eutypa lata UCREL1] | 59.1 | 5.0E-76 |
| DIP_10003898 | hypothetical protein PFICI_10266 [Pestalotiopsis fici W106-1] | 58.3 | 0.0 |
| DIP_10003899 | hypothetical protein FAVG1_07142 [Fusarium avenaceum] | 73 | 7.0E-27 |
| DIP_10003900 | putative small nucleolar ribonucleoprotein complex subunit utp14 protein [Eutypa lata UCREL1] | 55.1 | 0.0 |
| DIP_10003901 | hypothetical protein PFICI_10253 [Pestalotiopsis fici W106-1] | 57.4 | 0.0 |
| DIP_10003902 | putative potassium:hydrogen antiporter protein [Togninia minima UCRPA7] | 58 | 0.0 |
| DIP_10003903 | NA | | |
| DIP_10003904 | hypothetical protein PFICI_10256 [Pestalotiopsis fici W106-1] | 58.7 | 6.0E-52 |
| DIP_10003905 | hypothetical protein GLRG_08754 [Colletotrichum graminicola M1.001] | 49.6 | 2.0E-72 |
| DIP_10003906 | NA | | |
| DIP_10003907 | hypothetical protein CGLO_09689 [Colletotrichum gloeosporioides Cg-14] | 41.1 | 1.0E-20 |
| DIP_10003908 | hypothetical protein NECHADRAFT_95402 [Nectria haematococca mpVI 77-13-4] | 30.4 | 7.0E-44 |
| DIP_10003909 | hypothetical protein GLRG_10481 [Colletotrichum graminicola M1.001] | 37.4 | 1.0E-13 |
| DIP_10003910 | putative sterol regulatory element binding protein sre1 protein [Eutypa lata UCREL1] | 68.2 | 2.0E-117 |
| DIP_10003911 | hypothetical protein PFICI_14284 [Pestalotiopsis fici W106-1] | 51.6 | 2.0E-163 |
| DIP_10003912 | hypothetical protein PFICI_14285 [Pestalotiopsis fici W106-1] | 61 | 0.0 |
| DIP_10003913 | putative n-acetylglucosamine-6-phosphate deacetylase protein [Eutypa lata UCREL1] | 72.5 | 0.0 |
| DIP_10003914 | hypothetical protein PFICI_14287 [Pestalotiopsis fici W106-1] | 73 | 0.0 |
| DIP_10003915 | putative transcriptional regulator vib-1 protein [Eutypa lata UCREL1] | 65.9 | 2.0E-175 |
| DIP_10003916 | hypothetical protein PFICI_14289 [Pestalotiopsis fici W106-1] | 60.4 | 0.0 |
| DIP_10003917 | NA | | |
| DIP_10003918 | hypothetical protein PFICI_14290 [Pestalotiopsis fici W106-1] | 74.3 | 0.0 |
| DIP_10003919 | putative neutral ceramidase protein [Togninia minima UCRPA7] | 58.2 | 0.0 |
| DIP_10003920 | putative glycoside hydrolase family 31 protein [Togninia minima UCRPA7] | 63.6 | 0.0 |
| DIP_10003921 | NA | | |
| DIP_10003922 | hypothetical protein THAR02_00791 [Trichoderma harzianum] | 39.3 | 4.0E-09 |
| DIP_10003923 | hypothetical protein PFICI_02634 [Pestalotiopsis fici W106-1] | 40.6 | 3.0E-88 |
| DIP_10003924 | NA | | |
| DIP_10003925 | putative cell differentiation protein rcd1 protein [Eutypa lata UCREL1] | 85 | 0.0 |
| DIP_10003926 | hypothetical protein PFICI_14859 [Pestalotiopsis fici W106-1] | 58.7 | 0.0 |
| DIP_10003927 | hypothetical protein UCREL1_6554 [Eutypa lata UCREL1] | 35.6 | 1.0E-57 |
| DIP_10003928 | putative glycerol uptake facilitator protein [Eutypa lata UCREL1] | 52.4 | 0.0 |
| DIP_10003929 | putative pap2 superfamily protein [Eutypa lata UCREL1] | 70.6 | 5.0E-152 |
| DIP_10003930 | hypothetical protein UCREL1_2595 [Eutypa lata UCREL1] | 30.4 | 2.0E-49 |
| DIP_10003931 | putative glucose sorbosone dehydrogenase protein [Eutypa lata UCREL1] | 47.1 | 3.0E-119 |
| DIP_10003932 | hypothetical protein PFICI_14148 [Pestalotiopsis fici W106-1] | 67.6 | 0.0 |
| DIP_10003933 | hypothetical protein H634G_04760 [Metarhizium anisopliae BRIP 53293] | 54.5 | 2.0E-95 |
| DIP_10003934 | NmrA family protein [Streptomyces griseus] | 30.2 | 2.0E-10 |
| DIP_10003935 | hypothetical protein PFICI_14266 [Pestalotiopsis fici W106-1] | 63.5 | 8.0E-100 |
| DIP_10003936 | NA | | |
| DIP_10003937 | predicted protein [Trichoderma reesei QM6a] | 55.5 | 5.0E-71 |
| DIP_10003938 | putative alanine aminotransferase protein [Eutypa lata UCREL1] | 81.5 | 0.0 |
| DIP_10003939 | putative 26s proteasome regulatory subunit rpn7 protein [Eutypa lata UCREL1] | 81.6 | 0.0 |
| DIP_10003940 | hypothetical protein PFICI_14259 [Pestalotiopsis fici W106-1] | 45 | 0.0 |
| DIP_10003941 | hypothetical protein PFICI_14245 [Pestalotiopsis fici W106-1] | 49.9 | 2.0E-129 |
| DIP_10003942 | cupin domain-containing protein [Colletotrichum graminicola M1.001] | 44.9 | 8.0E-47 |
| DIP_10003943 | putative alpha beta hydrolase fold family protein [Eutypa lata UCREL1] | 41.8 | 2.0E-151 |
| DIP_10003944 | NA | | |
| DIP_10003945 | lactonohydrolase [Colletotrichum gloeosporioides Nara gc5] | 53.9 | 1.0E-159 |
| DIP_10003946 | DlpA domain-containing protein [Colletotrichum gloeosporioides Cg-14] | 77 | 1.0E-121 |
| DIP_10003947 | hypothetical protein CGLO_05464 [Colletotrichum gloeosporioides Cg-14] | 70.4 | 0.0 |
| DIP_10003948 | fungal specific transcription factor domain protein [Colletotrichum gloeosporioides Nara gc5] | 44.5 | 2.0E-150 |
| DIP_10003949 | hypothetical protein PFICI_02180 [Pestalotiopsis fici W106-1] | 61.1 | 3.0E-164 |
| DIP_10003950 | putative glycoside hydrolase family 61 protein [Eutypa lata UCREL1] | 48.6 | 8.0E-75 |
| DIP_10003951 | hypothetical protein PFICI_06732 [Pestalotiopsis fici W106-1] | 36.6 | 3.0E-31 |
| DIP_10003952 | hypothetical protein VDAG_07979 [Verticillium dahliae VdLs.17] | 32.9 | 9.0E-08 |
| DIP_10003953 | hypothetical protein PFICI_11209 [Pestalotiopsis fici W106-1] | 54 | 0.0 |
| DIP_10003954 | NA | | |
| DIP_10003955 | putative nudix domain-containing protein [Eutypa lata UCREL1] | 34.9 | 4.0E-35 |
| DIP_10003956 | hypothetical protein UCREL1_2595 [Eutypa lata UCREL1] | 33.2 | 4.0E-62 |
| DIP_10003957 | NA | | |
| DIP_10003958 | hypothetical protein PFICI_14873 [Pestalotiopsis fici W106-1] | 66.2 | 0.0 |
| DIP_10003959 | Mitochondrial import inner membrane translocase subunit tim-17 [Pestalotiopsis fici W106-1] | 93.3 | 3.0E-99 |
| DIP_10003960 | hypothetical protein COCHEDRAFT_1187409 [Bipolaris maydis C5] | 40.7 | 6.0E-50 |
| DIP_10003961 | putative protein transport protein sec13 protein [Eutypa lata UCREL1] | 67.3 | 2.0E-145 |
| DIP_10003962 | hypothetical protein PFICI_00271 [Pestalotiopsis fici W106-1] | 57.6 | 0.0 |
| DIP_10003963 | Dyp-type peroxidase [Penicillium expansum] | 60.3 | 0.0 |
| DIP_10003964 | putative galactose oxidase precursor protein [Botrytis cinerea BcDW1] | 58.1 | 0.0 |
| DIP_10003965 | putative zinc-binding oxidoreductase protein [Eutypa lata UCREL1] | 47.5 | 3.0E-114 |
| DIP_10003966 | hypothetical protein PHACADRAFT_176642 [Phanerochaete carnosa HHB-10118-sp] | 34.3 | 7.0E-19 |
| DIP_10003967 | hypothetical protein PFICI_06802 [Pestalotiopsis fici W106-1] | 54.4 | 1.0E-104 |
| DIP_10003968 | NA | | |
| DIP_10003969 | hypothetical protein CGLO_00621 [Colletotrichum gloeosporioides Cg-14] | 56.2 | 5.0E-155 |
| DIP_10003970 | hypothetical protein M7I_7469 [Glarea lozoyensis 74030] | 39 | 3.0E-23 |
| DIP_10003971 | hypothetical protein UCRPA7_2118 [Togninia minima UCRPA7] | 44.9 | 2.0E-52 |
| DIP_10003972 | subtilisin-like protease [Colletotrichum gloeosporioides Nara gc5] | 59.3 | 2.0E-153 |
| DIP_10003973 | hypothetical protein PFICI_06796 [Pestalotiopsis fici W106-1] | 74.4 | 0.0 |
| DIP_10003974 | hypothetical protein CTHT_0047820 [Chaetomium thermophilum var. thermophilum DSM 1495] | 58.4 | 2.0E-135 |
| DIP_10003975 | putative squalene synthetase protein [Eutypa lata UCREL1] | 72.6 | 0.0 |
| DIP_10003976 | hypothetical protein PFICI_14280 [Pestalotiopsis fici W106-1] | 76.4 | 0.0 |
| DIP_10003977 | putative pantothenate protein [Eutypa lata UCREL1] | 71.5 | 0.0 |
| DIP_10003978 | hypothetical protein FOTG_17884 [Fusarium oxysporum f. sp. vasinfectum 25433] | 57.9 | 5.0E-98 |
| DIP_10003979 | putative transaldolase protein [Eutypa lata UCREL1] | 70.2 | 0.0 |
| DIP_10003980 | hypothetical protein AURDEDRAFT_113503 [Auricularia delicata TFB-10046 SS5] | 45.9 | 3.0E-26 |
| DIP_10003981 | putative mfs hexose protein [Eutypa lata UCREL1] | 69.6 | 0.0 |
| DIP_10003982 | hypothetical protein PFICI_14271 [Pestalotiopsis fici W106-1] | 43.9 | 0.0 |
| DIP_10003983 | hypothetical protein PFICI_14244 [Pestalotiopsis fici W106-1] | 60.1 | 0.0 |
| DIP_10003984 | hypothetical protein SETTUDRAFT_162491 [Setosphaeria turcica Et28A] | 45 | 3.0E-70 |
| DIP_10003985 | NA | | |
| DIP_10003986 | hypothetical protein PFICI_14090 [Pestalotiopsis fici W106-1] | 51.6 | 4.0E-170 |
| DIP_10003987 | hypothetical protein SAPIO_CDS8128 [Scedosporium apiospermum] | 66.4 | 0.0 |
| DIP_10003988 | putative dna replication protein 4 protein [Eutypa lata UCREL1] | 61.8 | 0.0 |
| DIP_10003989 | hypothetical protein PFICI_07014 [Pestalotiopsis fici W106-1] | 35.5 | 8.0E-58 |
| DIP_10003990 | NA | | |
| DIP_10003991 | hypothetical protein GLAREA_11168 [Glarea lozoyensis ATCC 20868] | 47.7 | 2.0E-89 |
| DIP_10003992 | putative glucosamine 6-phosphate acetyltransferase protein [Eutypa lata UCREL1] | 55 | 1.0E-58 |
| DIP_10003993 | NA | | |
| DIP_10003994 | NA | | |
| DIP_10003995 | cysteine synthase A [Trichoderma harzianum] | 67.6 | 0.0 |
| DIP_10003996 | hypothetical protein PFICI_06899 [Pestalotiopsis fici W106-1] | 66.5 | 0.0 |
| DIP_10003997 | putative alanyl-trna synthetase protein [Eutypa lata UCREL1] | 85 | 0.0 |
| DIP_10003998 | NA | | |
| DIP_10003999 | hypothetical protein PFICI_06792 [Pestalotiopsis fici W106-1] | 53.6 | 4.0E-44 |
| DIP_10004000 | hypothetical protein NECHADRAFT_102387 [Nectria haematococca mpVI 77-13-4] | 42.2 | 4.0E-25 |
| DIP_10004001 | putative ubiquitin-protein ligase sel1 protein [Eutypa lata UCREL1] | 67.3 | 0.0 |
| DIP_10004002 | hypothetical protein PFICI_02703 [Pestalotiopsis fici W106-1] | 72.4 | 0.0 |
| DIP_10004003 | hypothetical protein PFICI_02704 [Pestalotiopsis fici W106-1] | 63.8 | 8.0E-123 |
| DIP_10004004 | hypothetical protein PFICI_02705 [Pestalotiopsis fici W106-1] | 58.7 | 1.0E-159 |
| DIP_10004005 | NA | | |
| DIP_10004006 | hypothetical protein CFIO01_10301 [Colletotrichum fioriniae PJ7] | 38.8 | 5.0E-34 |
| DIP_10004007 | hypothetical protein BBA_05318 [Beauveria bassiana ARSEF 2860] | 32 | 2.0E-27 |
| DIP_10004008 | hypothetical protein PFICI_06292 [Pestalotiopsis fici W106-1] | 67.3 | 9.0E-105 |
| DIP_10004009 | NA | | |
| DIP_10004010 | hypothetical protein PFICI_06295 [Pestalotiopsis fici W106-1] | 47.1 | 3.0E-166 |
| DIP_10004011 | putative transcription elongation factor s-ii protein [Eutypa lata UCREL1] | 64.9 | 1.0E-144 |
| DIP_10004012 | major facilitator superfamily transporter [Colletotrichum graminicola M1.001] | 52.8 | 0.0 |
| DIP_10004013 | putative capsule polysaccharide biosynthesis protein [Eutypa lata UCREL1] | 55.2 | 4.0E-116 |
| DIP_10004014 | putative fatty acid elongase protein [Eutypa lata UCREL1] | 74.4 | 2.0E-159 |
| DIP_10004015 | putative cytochrome p450 monooxygenase protein [Eutypa lata UCREL1] | 82.4 | 0.0 |
| DIP_10004016 | hypothetical protein UCREL1_5312 [Eutypa lata UCREL1] | 35.1 | 9.0E-27 |
| DIP_10004017 | putative cytochrome p450 52a12 protein [Eutypa lata UCREL1] | 67 | 0.0 |
| DIP_10004018 | NA | | |
| DIP_10004019 | putative tetrahydrofolylpolyglutamate synthase protein [Eutypa lata UCREL1] | 59.4 | 0.0 |
| DIP_10004020 | putative rhoptry protein [Eutypa lata UCREL1] | 43.6 | 0.0 |
| DIP_10004021 | putative g2 mitotic-specific cyclin-b1 protein [Eutypa lata UCREL1] | 84.1 | 0.0 |
| DIP_10004022 | NA | | |
| DIP_10004023 | NA | | |

| | | | |
|---|---|---|---|
| DIP_10004024 | NA | | |
| DIP_10004025 | hypothetical protein PFICI_06814 [Pestalotiopsis fici W106-1] | 70.4 | 0.0 |
| DIP_10004026 | hypothetical protein PFICI_06815 [Pestalotiopsis fici W106-1] | 65.6 | 9.0E-177 |
| DIP_10004027 | hypothetical protein TRV_00001, partial [Trichophyton verrucosum HKI 0517] | 45.1 | 9.0E-29 |
| DIP_10004028 | putative inosine-5 -monophosphate dehydrogenase imd2 protein [Eutypa lata UCREL1] | 86.7 | 0.0 |
| DIP_10004029 | hypothetical protein PFICI_06812 [Pestalotiopsis fici W106-1] | 67.3 | 0.0 |
| DIP_10004030 | hypothetical protein CFIO01_10965 [Colletotrichum fioriniae PJ7] | 56.7 | 1.0E-122 |
| DIP_10004031 | hypothetical protein CSUB01_10150 [Colletotrichum sublineola] | 52.2 | 4.0E-131 |
| DIP_10004032 | ubiE/COQ5 methyltransferase [Trichophyton tonsurans CBS 112818] | 58 | 4.0E-111 |
| DIP_10004033 | hypothetical protein MYCTH_2306233 [Myceliophthora thermophila ATCC 42464] | 34 | 4.0E-23 |
| DIP_10004034 | NA | | |
| DIP_10004035 | hypothetical protein PFICI_06811 [Pestalotiopsis fici W106-1] | 71.3 | 2.0E-130 |
| DIP_10004036 | hypothetical protein V491_02624 [Pseudogymnoascus pannorum VKM F-3775] | 51.6 | 0.0 |
| DIP_10004037 | hypothetical protein ARB_02111 [Byssochlamys spectabilis No. 5] | 55.3 | 3.0E-122 |
| DIP_10004038 | conserved hypothetical protein [Byssochlamys spectabilis No. 5] | 66.5 | 0.0 |
| DIP_10004039 | malate/L-lactate dehydrogenase [Byssochlamys spectabilis No. 5] | 58.8 | 4.0E-150 |
| DIP_10004040 | putative d-3-phosphoglycerate dehydrogenase protein [Togninia minima UCRPA7] | 65.9 | 2.0E-147 |
| DIP_10004041 | hypothetical protein GGTG_12262 [Gaeumannomyces graminis var. tritici R3-111a-1] | 70.7 | 6.0E-117 |
| DIP_10004042 | putative sulphydryl oxidase protein [Eutypa lata UCREL1] | 58.6 | 8.0E-170 |
| DIP_10004043 | Oxygen-dependent choline dehydrogenase-like protein [Acremonium chrysogenum ATCC 11550] | 44.2 | 8.0E-150 |
| DIP_10004044 | NA | | |
| DIP_10004045 | tannase and feruloyl esterase [Aspergillus ruber CBS 135680] | 69.2 | 0.0 |
| DIP_10004046 | putative karyopherin kap123 protein [Eutypa lata UCREL1] | 73.8 | 0.0 |
| DIP_10004047 | putative covalently-linked cell wall protein [Eutypa lata UCREL1] | 62.3 | 2.0E-99 |
| DIP_10004048 | putative u1 zinc finger domain-containing protein [Eutypa lata UCREL1] | 50.3 | 2.0E-86 |
| DIP_10004049 | NA | | |
| DIP_10004050 | NA | | |
| DIP_10004051 | putative catabolite repression protein creC [Pestalotiopsis fici W106-1] | 73.6 | 0.0 |
| DIP_10004052 | mitochondrial rho gtpase [Colletotrichum gloeosporioides Nara gc5] | 78 | 0.0 |
| DIP_10004053 | hypothetical protein VDAG_08232 [Verticillium dahliae VdLs.17] | 51.5 | 3.0E-77 |
| DIP_10004054 | hypothetical protein PFICI_14155 [Pestalotiopsis fici W106-1] | 60.7 | 0.0 |
| DIP_10004055 | putative ph domain-containing protein [Eutypa lata UCREL1] | 72.2 | 0.0 |
| DIP_10004056 | NA | | |
| DIP_10004057 | hypothetical protein THAR02_03281 [Trichoderma harzianum] | 35.2 | 7.0E-40 |
| DIP_10004058 | hypothetical protein PFICI_14151 [Pestalotiopsis fici W106-1] | 33.2 | 3.0E-07 |
| DIP_10004059 | putative chromosomal organization and dna repair protein [Eutypa lata UCREL1] | 43.4 | 5.0E-94 |
| DIP_10004060 | hypothetical protein PV10_08583 [Exophiala mesophila] | 59.6 | 3.0E-138 |
| DIP_10004061 | hypothetical protein W97_08595 [Coniosporium apollinis CBS 100218] | 41.7 | 6.0E-128 |
| DIP_10004062 | NA | | |
| DIP_10004063 | putative integral membrane protein [Eutypa lata UCREL1] | 48.1 | 9.0E-47 |
| DIP_10004064 | putative l-galactonate dehydratase protein [Eutypa lata UCREL1] | 87.1 | 0.0 |
| DIP_10004065 | NA | | |
| DIP_10004066 | putative dna damage response protein [Eutypa lata UCREL1] | 56.6 | 0.0 |
| DIP_10004067 | cytoplasm protein [Verticillium dahliae VdLs.17] | 41.6 | 1.0E-54 |
| DIP_10004068 | hypothetical protein CHGG_10863 [Chaetomium globosum CBS 148.51] | 39.3 | 4.0E-75 |
| DIP_10004069 | NA | | |
| DIP_10004070 | unnamed protein product [Fusarium sp. FIESC_5 CS3069] | 82.4 | 4.0E-33 |
| DIP_10004071 | Thioredoxin reductase [Pestalotiopsis fici W106-1] | 80.3 | 0.0 |
| DIP_10004072 | putative phosphatidylinositol transfer protein sfh5 protein [Eutypa lata UCREL1] | 72.3 | 1.0E-153 |
| DIP_10004073 | hypothetical protein PFICI_13723 [Pestalotiopsis fici W106-1] | 64.3 | 0.0 |
| DIP_10004074 | putative tyrosyl-trna synthetase protein [Eutypa lata UCREL1] | 56.6 | 0.0 |
| DIP_10004075 | putative succinate dehydrogenase membrane anchor subunit protein [Eutypa lata UCREL1] | 70 | 5.0E-95 |
| DIP_10004076 | hypothetical protein PFICI_13719 [Pestalotiopsis fici W106-1] | 70.7 | 0.0 |
| DIP_10004077 | hypothetical protein PFICI_13720 [Pestalotiopsis fici W106-1] | 67.4 | 0.0 |
| DIP_10004078 | Alpha-glucosidase [Pestalotiopsis fici W106-1] | 65.4 | 0.0 |
| DIP_10004079 | putative zinc finger ccch type domain containing protein [Eutypa lata UCREL1] | 76.7 | 7.0E-123 |
| DIP_10004080 | NA | | |
| DIP_10004081 | hypothetical protein PFICI_13854 [Pestalotiopsis fici W106-1] | 56.5 | 0.0 |
| DIP_10004082 | hypothetical protein PFICI_13852 [Pestalotiopsis fici W106-1] | 55.5 | 3.0E-174 |
| DIP_10004083 | hypothetical protein Z518_07912 [Rhinocladiella mackenziei CBS 650.93] | 40 | 1.0E-19 |
| DIP_10004084 | NA | | |
| DIP_10004085 | AAA family ATPase [Colletotrichum higginsianum] | 43.9 | 8.0E-26 |
| DIP_10004086 | class II Aldolase and Adducin domain-containing protein [Colletotrichum fioriniae PJ7] | 54.7 | 2.0E-115 |
| DIP_10004087 | hypothetical protein UCREL1_2595 [Eutypa lata UCREL1] | 42.9 | 3.0E-60 |
| DIP_10004088 | hypothetical protein VHEMI04314 [Torrubiella hemipterigena] | 52.2 | 2.0E-109 |
| DIP_10004089 | hypothetical protein NEUTE1DRAFT_138203 [Neurospora tetrasperma FGSC 2508] | 31.6 | 2.0E-15 |
| DIP_10004090 | hypothetical protein PFICI_02854 [Pestalotiopsis fici W106-1] | 60.1 | 3.0E-175 |
| DIP_10004091 | hypothetical protein THITE_115282 [Thielavia terrestris NRRL 8126] | 32.8 | 1.0E-77 |
| DIP_10004092 | hypothetical protein THITE_2091470 [Thielavia terrestris NRRL 8126] | 61.6 | 0.0 |
| DIP_10004093 | hypothetical protein GLRG_05294 [Colletotrichum graminicola M1.001] | 70.3 | 0.0 |
| DIP_10004094 | hypothetical protein PFICI_12761 [Pestalotiopsis fici W106-1] | 58.6 | 5.0E-128 |
| DIP_10004095 | hypothetical protein NECHADRAFT_80111 [Nectria haematococca mpVI 77-13-4] | 51.9 | 2.0E-167 |
| DIP_10004096 | putative 2-epi-5-epi-valiolone synthase [Diaporthe ampelina] | 81.8 | 0.0 |
| DIP_10004097 | hypothetical protein PFICI_12760 [Pestalotiopsis fici W106-1] | 74.2 | 1.0E-127 |
| DIP_10004098 | alcohol dehydrogenase [Fusarium oxysporum f. sp. pisi HDV247] | 57.7 | 9.0E-110 |
| DIP_10004099 | hypothetical protein VDAG_08291 [Verticillium dahliae VdLs.17] | 51 | 3.0E-68 |
| DIP_10004100 | NA | | |
| DIP_10004101 | hypothetical protein S40293_04883 [Stachybotrys chartarum IBT 40293] | 65.6 | 0.0 |
| DIP_10004102 | maltose O-acetyltransferase [Verticillium dahliae VdLs.17] | 63.5 | 3.0E-78 |
| DIP_10004103 | hypothetical protein PFICI_07490 [Pestalotiopsis fici W106-1] | 58.3 | 1.0E-47 |
| DIP_10004104 | hypothetical protein VHEMI02051 [Torrubiella hemipterigena] | 41.7 | 2.0E-157 |
| DIP_10004105 | putative zip zinc transporter protein [Eutypa lata UCREL1] | 61.6 | 0.0 |
| DIP_10004106 | putative ssdna binding protein [Eutypa lata UCREL1] | 74.8 | 1.0E-69 |
| DIP_10004107 | putative monooxygenase fad-binding protein [Eutypa lata UCREL1] | 51.7 | 0.0 |
| DIP_10004108 | hypothetical protein A1O7_10085 [Cladophialophora yegresii CBS 114405] | 32.1 | 8.0E-54 |
| DIP_10004109 | putative glycerol proton symporter of the plasma subject to glucose-induced inactivation protein [Eutypa lata UCREL1] | 63 | 0.0 |
| DIP_10004110 | pheromone receptor, partial [Metarhizium anisopliae ARSEF 549] | 31.6 | 4.0E-53 |
| DIP_10004111 | hypothetical protein Cob_12449 [Colletotrichum orbiculare MAFF 240422] | 52.6 | 5.0E-121 |
| DIP_10004112 | putative serine threonine-protein kinase rio1 protein [Eutypa lata UCREL1] | 75.6 | 0.0 |
| DIP_10004113 | NA | | |
| DIP_10004114 | putative translocation protein sec63 protein [Eutypa lata UCREL1] | 72.3 | 0.0 |
| DIP_10004115 | putative protein-s-isoprenylcysteine o-methyltransferase [Diaporthe ampelina] | 60 | 4.0E-112 |
| DIP_10004116 | hypothetical protein PFICI_01440 [Pestalotiopsis fici W106-1] | 57.9 | 6.0E-104 |
| DIP_10004117 | putative rna binding protein [Eutypa lata UCREL1] | 69.6 | 0.0 |
| DIP_10004118 | hypothetical protein PFICI_07523 [Pestalotiopsis fici W106-1] | 73.1 | 0.0 |
| DIP_10004119 | putative methyltransferase domain-containing protein [Togninia minima UCRPA7] | 50.8 | 2.0E-106 |
| DIP_10004120 | putative dtdp-glucose -dehydratase protein [Eutypa lata UCREL1] | 76.9 | 0.0 |
| DIP_10004121 | integral membrane protein [Grosmannia clavigera kw1407] | 59.8 | 2.0E-151 |
| DIP_10004122 | hypothetical protein CMQ_366 [Grosmannia clavigera kw1407] | 41.6 | 1.0E-53 |
| DIP_10004123 | putative glycoside hydrolase family 16 protein [Eutypa lata UCREL1] | 52.1 | 1.0E-87 |
| DIP_10004124 | Sorting nexin-41 [Pestalotiopsis fici W106-1] | 72.4 | 0.0 |
| DIP_10004125 | putative vacuolar atp synthase 16 kda proteolipid subunit protein [Togninia minima UCRPA7] | 83.8 | 6.0E-57 |
| DIP_10004126 | putative vacuolar atp synthase 16 kda proteolipid subunit protein [Eutypa lata UCREL1] | 70.3 | 5.0E-27 |
| DIP_10004127 | putative candidapepsin-4 precursor protein [Eutypa lata UCREL1] | 63.7 | 2.0E-123 |
| DIP_10004128 | NA | | |
| DIP_10004129 | hypothetical protein PFICI_01435 [Pestalotiopsis fici W106-1] | 51.8 | 0.0 |
| DIP_10004130 | sucrose transporter, partial [Colletotrichum fioriniae PJ7] | 54.9 | 0.0 |
| DIP_10004131 | NA | | |
| DIP_10004132 | putative atp phosphoribosyltransferase protein [Eutypa lata UCREL1] | 80 | 0.0 |
| DIP_10004133 | hypothetical protein PFICI_01881 [Pestalotiopsis fici W106-1] | 32.3 | 1.0E-10 |
| DIP_10004134 | hypothetical protein UCREL1_6426 [Eutypa lata UCREL1] | 49.8 | 2.0E-68 |
| DIP_10004135 | hypothetical protein PFICI_01878 [Pestalotiopsis fici W106-1] | 57.8 | 0.0 |
| DIP_10004136 | hypothetical protein THITE_114904 [Thielavia terrestris NRRL 8126] | 44.5 | 3.0E-92 |
| DIP_10004137 | hypothetical protein PFICI_01877 [Pestalotiopsis fici W106-1] | 50.1 | 0.0 |
| DIP_10004138 | hypothetical protein THITE_45297 [Thielavia terrestris NRRL 8126] | 42.9 | 2.0E-51 |
| DIP_10004139 | hypothetical protein UCREL1_11596 [Eutypa lata UCREL1] | 48.8 | 2.0E-37 |
| DIP_10004140 | putative gnat family protein [Eutypa lata UCREL1] | 39 | 2.0E-67 |
| DIP_10004141 | putative uroporphyrinogen decarboxylase protein [Eutypa lata UCREL1] | 81.6 | 0.0 |
| DIP_10004142 | NA | | |
| DIP_10004143 | hypothetical protein PFICI_01871 [Pestalotiopsis fici W106-1] | 62.2 | 0.0 |
| DIP_10004144 | hypothetical protein S40285_03835 [Stachybotrys chlorohalonata IBT 40285] | 47 | 4.0E-66 |
| DIP_10004145 | NA | | |
| DIP_10004146 | NA | | |
| DIP_10004147 | putative tat pathway signal sequence protein [Eutypa lata UCREL1] | 58.5 | 5.0E-138 |
| DIP_10004148 | hypothetical protein PFICI_01869 [Pestalotiopsis fici W106-1] | 76.8 | 0.0 |
| DIP_10004149 | NA | | |
| DIP_10004150 | NA | | |
| DIP_10004151 | hypothetical protein FAVG1_00066 [Fusarium avenaceum] | 56.1 | 0.0 |
| DIP_10004152 | peroxisome biosynthesis protein (PAS8/Peroxin-6), putative [Aspergillus flavus NRRL3357] | 49.6 | 0.0 |
| DIP_10004153 | NA | | |
| DIP_10004154 | NA | | |
| DIP_10004155 | putative 4-coumarate- ligase protein [Eutypa lata UCREL1] | 60.4 | 0.0 |
| DIP_10004156 | hypothetical protein PFICI_01221 [Pestalotiopsis fici W106-1] | 52.6 | 2.0E-107 |
| DIP_10004157 | hypothetical protein PFICI_12182 [Pestalotiopsis fici W106-1] | 67.2 | 0.0 |
| DIP_10004158 | NA | | |
| DIP_10004159 | putative glycerophosphoryl diester phosphodiesterase family protein [Eutypa lata UCREL1] | 36.7 | 6.0E-83 |
| DIP_10004160 | NA | | |
| DIP_10004161 | hypothetical protein PFICI_12181 [Pestalotiopsis fici W106-1] | 54.2 | 0.0 |
| DIP_10004162 | hypothetical protein PFICI_12179 [Pestalotiopsis fici W106-1] | 52 | 0.0 |
| DIP_10004163 | putative major facilitator superfamily protein [Eutypa lata UCREL1] | 65.4 | 0.0 |
| DIP_10004164 | integral membrane protein [Trichoderma harzianum] | 41.9 | 9.0E-28 |
| DIP_10004165 | hypothetical protein CFIO01_04649 [Colletotrichum fioriniae PJ7] | 58.8 | 0.0 |
| DIP_10004166 | hypothetical protein CHGG_02335 [Chaetomium globosum CBS 148.51] | 54 | 0.0 |
| DIP_10004167 | hypothetical protein, variant [Magnaporthe oryzae 70-15] | 60.5 | 9.0E-102 |
| DIP_10004168 | mate efflux family protein [Colletotrichum orbiculare MAFF 240422] | 58 | 0.0 |
| DIP_10004169 | hypothetical protein NECHADRAFT_106298 [Nectria haematococca mpVI 77-13-4] | 35.2 | 2.0E-20 |
| DIP_10004170 | NA | | |
| DIP_10004171 | hypothetical protein TRIATDRAFT_263429 [Trichoderma atroviride IMI 206040] | 67.1 | 6.0E-129 |
| DIP_10004172 | ATP-dependent DNA helicase MPH1 [Pestalotiopsis fici W106-1] | 59 | 0.0 |
| DIP_10004173 | putative mboat family protein [Eutypa lata UCREL1] | 69.2 | 0.0 |
| DIP_10004174 | major facilitator superfamily transporter [Colletotrichum fioriniae PJ7] | 57 | 0.0 |
| DIP_10004175 | putative small glutamine-rich tetratricopeptide repeat-containing protein a protein [Eutypa lata UCREL1] | 76.4 | 0.0 |
| DIP_10004176 | NA | | |
| DIP_10004177 | hypothetical protein DOTSEDRAFT_138875 [Dothistroma septosporum NZE10] | 49.4 | 6.0E-52 |
| DIP_10004178 | NA | | |
| DIP_10004179 | hypothetical protein PFICI_12140 [Pestalotiopsis fici W106-1] | 31 | 2.0E-26 |
| DIP_10004180 | hypothetical protein PFICI_12136 [Pestalotiopsis fici W106-1] | 49.4 | 0.0 |
| DIP_10004181 | NA | | |
| DIP_10004182 | hypothetical protein PFICI_12135 [Pestalotiopsis fici W106-1] | 75.7 | 0.0 |
| DIP_10004183 | NA | | |
| DIP_10004184 | hypothetical protein PFICI_07526 [Pestalotiopsis fici W106-1] | 40.7 | 4.0E-144 |
| DIP_10004185 | putative curved dna-binding protein [Eutypa lata UCREL1] | 77.5 | 0.0 |
| DIP_10004186 | hypothetical protein FVEG_13296 [Fusarium verticillioides 7600] | 62 | 0.0 |
| DIP_10004187 | hypothetical protein PFICI_07528 [Pestalotiopsis fici W106-1] | 63.4 | 0.0 |
| DIP_10004188 | hypothetical protein FOTG_01069 [Fusarium oxysporum f. sp. vasinfectum 25433] | 71.5 | 0.0 |
| DIP_10004189 | putative mrna capping enzyme alpha protein [Eutypa lata UCREL1] | 68.6 | 0.0 |
| DIP_10004190 | putative s-formylglutathione hydrolase protein [Eutypa lata UCREL1] | 73.2 | 9.0E-156 |
| DIP_10004191 | NA | | |
| DIP_10004192 | nucleoside-diphosphate-sugar epimerase, putative [Talaromyces islandicus] | 49.8 | 7.0E-81 |
| DIP_10004193 | NA | | |
| DIP_10004194 | putative duf6 domain protein [Eutypa lata UCREL1] | 55.4 | 4.0E-136 |
| DIP_10004195 | hypothetical protein THITE_2120102 [Thielavia terrestris NRRL 8126] | 47.7 | 6.0E-79 |
| DIP_10004196 | hypothetical protein PFICI_01436 [Pestalotiopsis fici W106-1] | 53.1 | 0.0 |
| DIP_10004197 | hypothetical protein SMAC_02774 [Sordaria macrospora k-hell] | 40.1 | 9.0E-138 |
| DIP_10004198 | putative mrna export factor mex67 protein [Eutypa lata UCREL1] | 47.5 | 0.0 |
| DIP_10004199 | Polyadenylation factor subunit 2 [Pestalotiopsis fici W106-1] | 79.5 | 0.0 |
| DIP_10004200 | NA | | |
| DIP_10004201 | putative gdp-mannose transporter protein [Eutypa lata UCREL1] | 83.9 | 0.0 |
| DIP_10004202 | hypothetical protein S7711_07197 [Stachybotrys chartarum IBT 7711] | 74.7 | 0.0 |
| DIP_10004203 | NA | | |
| DIP_10004204 | putative glutathione synthetase protein [Eutypa lata UCREL1] | 65.9 | 0.0 |
| DIP_10004205 | putative snf2 family atp-dependent chromatin-remodeling factor snf21 protein [Togninia minima UCRPA7] | 33.9 | 0.0 |
| DIP_10004206 | NA | | |
| DIP_10004207 | tyrosinase-like protein [Myceliophthora thermophila ATCC 42464] | 39.3 | 3.0E-81 |
| DIP_10004208 | glutathione-dependent formaldehyde-activating enzyme [Colletotrichum fioriniae PJ7] | 37.9 | 1.0E-55 |
| DIP_10004209 | putative major facilitator superfamily transporter protein [Eutypa lata UCREL1] | 55.6 | 0.0 |
| DIP_10004210 | hypothetical protein M419DRAFT_23250 [Trichoderma reesei RUT C-30] | 49 | 1.0E-62 |
| DIP_10004211 | hypothetical protein MGG_14713 [Magnaporthe oryzae 70-15] | 61.7 | 0.0 |
| DIP_10004212 | NA | | |
| DIP_10004213 | NA | | |
| DIP_10004214 | hypothetical protein PFICI_12449 [Pestalotiopsis fici W106-1] | 66.9 | 0.0 |
| DIP_10004215 | snf7 family protein [Ophiostoma piceae UAMH 11346] | 81.3 | 5.0E-117 |
| DIP_10004216 | putative protein phosphatase 2c protein [Eutypa lata UCREL1] | 74.5 | 0.0 |
| DIP_10004217 | putative c6 zinc finger domain-containing protein [Eutypa lata UCREL1] | 46.5 | 0.0 |
| DIP_10004218 | NA | | |
| DIP_10004219 | putative phosphoadenosine phosphosulfate reductase protein [Togninia minima UCRPA7] | 34.8 | 4.0E-72 |
| DIP_10004220 | similar to C4-dicarboxylate transporter/malic acid transport protein [Leptosphaeria maculans JN3] | 45.6 | 6.0E-86 |
| DIP_10004221 | putative calpain family cysteine protease protein [Eutypa lata UCREL1] | 61.3 | 0.0 |
| DIP_10004222 | NA | | |
| DIP_10004223 | hypothetical protein V499_07102 [Pseudogymnoascus pannorum VKM F-103] | 33.1 | 8.0E-13 |
| DIP_10004224 | hypothetical protein PFICI_09784 [Pestalotiopsis fici W106-1] | 36.2 | 3.0E-68 |
| DIP_10004225 | Superoxide dismutase [Pestalotiopsis fici W106-1] | 59.4 | 2.0E-82 |
| DIP_10004226 | hypothetical protein FOC1_g10010477 [Fusarium oxysporum f. sp. cubense race 1] | 38.5 | 0.0 |
| DIP_10004227 | hypothetical protein NECHADRAFT_87532 [Nectria haematococca mpVI 77-13-4] | 30.4 | 1.0E-17 |
| DIP_10004228 | NA | | |
| DIP_10004229 | hypothetical protein PFICI_03732 [Pestalotiopsis fici W106-1] | 62.4 | 0.0 |
| DIP_10004230 | putative xaa-pro dipeptidase protein [Eutypa lata UCREL1] | 75.2 | 0.0 |
| DIP_10004231 | hypothetical protein PFICI_12190 [Pestalotiopsis fici W106-1] | 63.5 | 0.0 |
| DIP_10004232 | putative fad binding domain-containing protein [Eutypa lata UCREL1] | 68.5 | 0.0 |
| DIP_10004233 | putative -like polyketide cyclase protein [Eutypa lata UCREL1] | 34 | 7.0E-47 |
| DIP_10004234 | putative malic acid transport protein [Eutypa lata UCREL1] | 67.2 | 0.0 |
| DIP_10004235 | NA | | |
| DIP_10004236 | hypothetical protein PFICI_12049 [Pestalotiopsis fici W106-1] | 51.7 | 2.0E-49 |
| DIP_10004237 | hypothetical protein OCS_01967 [Ophiocordyceps sinensis CO18] | 30.6 | 1.0E-08 |
| DIP_10004238 | NA | | |
| DIP_10004239 | NA | | |
| DIP_10004240 | putative fatty-acid amide hydrolase protein [Eutypa lata UCREL1] | 63.6 | 0.0 |
| DIP_10004241 | NA | | |
| DIP_10004242 | putative mfs sugar transporter protein [Eutypa lata UCREL1] | 62.3 | 0.0 |
| DIP_10004243 | hypothetical protein PFICI_11517 [Pestalotiopsis fici W106-1] | 61.1 | 3.0E-154 |
| DIP_10004244 | NA | | |
| DIP_10004245 | hypothetical protein PFICI_03745 [Pestalotiopsis fici W106-1] | 54.1 | 0.0 |
| DIP_10004246 | NA | | |
| DIP_10004247 | probable c-14 sterol reductase ERG-3 [Claviceps purpurea 20.1] | 72.9 | 0.0 |
| DIP_10004248 | putative vacuolar sorting receptor protein [Eutypa lata UCREL1] | 56.6 | 8.0E-138 |
| DIP_10004249 | hypothetical protein PFICI_11713 [Pestalotiopsis fici W106-1] | 31.5 | 3.0E-66 |
| DIP_10004250 | NA | | |
| DIP_10004251 | NA | | |
| DIP_10004252 | putative glycoside hydrolase family 76 protein [Eutypa lata UCREL1] | 61.3 | 2.0E-180 |
| DIP_10004253 | NA | | |
| DIP_10004254 | hypothetical protein PFICI_07324 [Pestalotiopsis fici W106-1] | 54.7 | 3.0E-112 |
| DIP_10004255 | hypothetical protein PFICI_11644 [Pestalotiopsis fici W106-1] | 66.2 | 0.0 |
| DIP_10004256 | hypothetical protein PVAR5_3618 [Byssochlamys spectabilis No. 5] | 50.5 | 8.0E-28 |
| DIP_10004257 | NA | | |
| DIP_10004258 | NA | | |
| DIP_10004259 | NA | | |
| DIP_10004260 | hypothetical protein PFICI_11646 [Pestalotiopsis fici W106-1] | 77.3 | 0.0 |
| DIP_10004261 | hypothetical protein Cob_12448 [Colletotrichum orbiculare MAFF 240422] | 43.4 | 3.0E-115 |
| DIP_10004262 | hypothetical protein UCREL1_6190 [Eutypa lata UCREL1] | 40 | 0.0 |
| DIP_10004263 | putative scf complex subunit skp1 protein [Eutypa lata UCREL1] | 85.2 | 3.0E-103 |
| DIP_10004264 | 60S ribosomal protein L39 [Verticillium alfalfae VaMs.102] | 96.1 | 7.0E-30 |
| DIP_10004265 | probable ARL1-ADP-ribosylation factor [Claviceps purpurea 20.1] | 96.1 | 6.0E-127 |
| DIP_10004266 | catalase [Colletotrichum graminicola M1.001] | 79.3 | 0.0 |
| DIP_10004267 | serine carboxypeptidase [Pseudozyma brasiliensis GHG001] | 55.1 | 0.0 |
| DIP_10004268 | NA | | |
| DIP_10004269 | NA | | |
| DIP_10004270 | NA | | |
| DIP_10004271 | hypothetical protein PFICI_12045 [Pestalotiopsis fici W106-1] | 56.1 | 0.0 |
| DIP_10004272 | NA | | |
| DIP_10004273 | NA | | |
| DIP_10004274 | NA | | |
| DIP_10004275 | putative het-s domain protein [Togninia minima UCRPA7] | 57.6 | 0.0 |
| DIP_10004276 | putative arginyl-trna synthetase protein [Eutypa lata UCREL1] | 77.5 | 0.0 |
| DIP_10004277 | putative cytochrome p450 protein [Eutypa lata UCREL1] | 65.3 | 0.0 |
| DIP_10004278 | Cytochrome c peroxidase [Pestalotiopsis fici W106-1] | 81.2 | 0.0 |
| DIP_10004279 | hypothetical protein TRIVIDRAFT_57654 [Trichoderma virens Gv29-8] | 62.8 | 1.0E-63 |
| DIP_10004280 | putative cell division control protein 48 protein [Eutypa lata UCREL1] | 95 | 0.0 |
| DIP_10004281 | putative vacuolar amino acid transporter 1 protein [Eutypa lata UCREL1] | 66.7 | 0.0 |
| DIP_10004282 | putative coatomer zeta subunit protein [Eutypa lata UCREL1] | 89.8 | 3.0E-129 |
| DIP_10004283 | putative translation initiation protein sua5 protein [Eutypa lata UCREL1] | 50.9 | 2.0E-165 |
| DIP_10004284 | hypothetical protein PFICI_01123 [Pestalotiopsis fici W106-1] | 69.2 | 0.0 |
| DIP_10004285 | hypothetical protein PFICI_09659 [Pestalotiopsis fici W106-1] | 66 | 0.0 |
| DIP_10004286 | DNA-directed RNA polymerases I, II, and III subunit RPABC2-like protein [Acremonium chrysogenum ATCC 11550] | 73.3 | 4.0E-74 |
| DIP_10004287 | putative zinc finger protein 467 protein [Eutypa lata UCREL1] | 50.8 | 0.0 |
| DIP_10004288 | NA | | |
| DIP_10004289 | PRTase-like protein [Glarea lozoyensis ATCC 20868] | 43.6 | 3.0E-179 |
| DIP_10004290 | putative epoxide hydrolase protein [Eutypa lata UCREL1] | 52.4 | 3.0E-119 |
| DIP_10004291 | putative phospholipid-transporting atpase protein [Eutypa lata UCREL1] | 78.9 | 0.0 |
| DIP_10004292 | External alternative NADH-ubiquinone oxidoreductase [Pestalotiopsis fici W106-1] | 75.4 | 0.0 |
| DIP_10004293 | hypothetical protein GGTG_09582 [Gaeumannomyces graminis var. tritici R3-111a-1] | 69.9 | 2.0E-118 |
| DIP_10004294 | NA | | |
| DIP_10004295 | hypothetical protein PFICI_05583 [Pestalotiopsis fici W106-1] | 52.6 | 7.0E-149 |
| DIP_10004296 | putative brct domain containing protein [Eutypa lata UCREL1] | 35.5 | 9.0E-42 |
| DIP_10004297 | synaptobrevin [Colletotrichum gloeosporioides Cg-14] | 39.9 | 4.0E-68 |
| DIP_10004298 | hypothetical protein CH063_05603 [Colletotrichum higginsianum] | 53.9 | 3.0E-132 |
| DIP_10004299 | NA | | |
| DIP_10004300 | hypothetical protein FOXB_01385 [Fusarium oxysporum Fo5176] | 31.5 | 1.0E-52 |
| DIP_10004301 | hypothetical protein UCDDA912_g05135 [Diaporthe ampelina] | 38.5 | 3.0E-63 |
| DIP_10004302 | hypothetical protein UCREL1_8406 [Eutypa lata UCREL1] | 52.5 | 5.0E-103 |
| DIP_10004303 | cyclin-dependent kinase G-1 [Metarhizium guizhouense ARSEF 977] | 55.2 | 0.0 |
| DIP_10004304 | putative duf185 domain-containing protein [Eutypa lata UCREL1] | 68.9 | 0.0 |
| DIP_10004305 | hypothetical protein PFICI_05565 [Pestalotiopsis fici W106-1] | 69.7 | 0.0 |
| DIP_10004306 | putative cyclophilin type peptidyl-prolyl cis-trans isomerase cld protein [Eutypa lata UCREL1] | 76.4 | 0.0 |
| DIP_10004307 | putative transmembrane protein [Togninia minima UCRPA7] | 47.7 | 8.0E-63 |
| DIP_10004308 | elongator complex associated protein Kti2 [Ustilaginoidea virens] | 54.3 | 2.0E-133 |
| DIP_10004309 | hypothetical protein PFICI_12967 [Pestalotiopsis fici W106-1] | 68.3 | 0.0 |
| DIP_10004310 | hypothetical protein PFICI_12965 [Pestalotiopsis fici W106-1] | 58.6 | 0.0 |
| DIP_10004311 | putative coatomer subunit delta protein [Togninia minima UCRPA7] | 82.4 | 0.0 |
| DIP_10004312 | putative glutamic acid-rich protein [Eutypa lata UCREL1] | 32.8 | 6.0E-65 |
| DIP_10004313 | duf803 domain membrane protein [Colletotrichum orbiculare MAFF 240422] | 75.2 | 0.0 |
| DIP_10004314 | 40S ribosomal protein S3 [Gaeumannomyces graminis var. tritici R3-111a-1] | 95.5 | 6.0E-173 |
| DIP_10004315 | hypothetical protein PFICI_12960 [Pestalotiopsis fici W106-1] | 63 | 0.0 |
| DIP_10004316 | putative phenylalanyl-trna beta subunit protein [Eutypa lata UCREL1] | 78.5 | 0.0 |
| DIP_10004317 | putative protein phosphatase pp2a regulatory subunit a protein [Eutypa lata UCREL1] | 87.3 | 0.0 |
| DIP_10004318 | putative s-adenosylmethionine-dependent methyltransferase protein [Eutypa lata UCREL1] | 41.8 | 2.0E-67 |
| DIP_10004319 | NA | | |
| DIP_10004320 | hypothetical protein UCREL1_8930 [Eutypa lata UCREL1] | 39.9 | 1.0E-32 |
| DIP_10004321 | putative beta-glucosidase a protein [Eutypa lata UCREL1] | 63.2 | 0.0 |
| DIP_10004322 | hypothetical protein PFICI_12955 [Pestalotiopsis fici W106-1] | 58.1 | 3.0E-77 |
| DIP_10004323 | hypothetical protein PFICI_12954 [Pestalotiopsis fici W106-1] | 62.9 | 9.0E-177 |
| DIP_10004324 | hypothetical protein GLAREA_01381 [Glarea lozoyensis ATCC 20868] | 31.8 | 2.0E-27 |
| DIP_10004325 | hypothetical protein PFICI_12944 [Pestalotiopsis fici W106-1] | 54.1 | 1.0E-79 |
| DIP_10004326 | hypothetical protein PFICI_12945 [Pestalotiopsis fici W106-1] | 55.4 | 6.0E-159 |
| DIP_10004327 | putative vacuolar protein sorting-associated protein [Eutypa lata UCREL1] | 80.9 | 0.0 |
| DIP_10004328 | NA | | |
| DIP_10004329 | putative rrna-processing protein fcf2 protein [Eutypa lata UCREL1] | 60.5 | 6.0E-82 |
| DIP_10004330 | putative nuclear transcription factor y subunit c-4 protein [Eutypa lata UCREL1] | 71 | 2.0E-119 |
| DIP_10004331 | hypothetical protein PFICI_12941 [Pestalotiopsis fici W106-1] | 86.4 | 0.0 |
| DIP_10004332 | hypothetical protein PFICI_12940 [Pestalotiopsis fici W106-1] | 51 | 2.0E-174 |
| DIP_10004333 | hypothetical protein PFICI_12939 [Pestalotiopsis fici W106-1] | 51.3 | 0.0 |
| DIP_10004334 | NA | | |
| DIP_10004335 | hypothetical protein PFICI_12595 [Pestalotiopsis fici W106-1] | 38.1 | 7.0E-99 |
| DIP_10004336 | hypothetical protein PFICI_12930 [Pestalotiopsis fici W106-1] | 68 | 2.0E-175 |
| DIP_10004337 | putative wd repeat-containing protein 70 protein [Eutypa lata UCREL1] | 79.1 | 0.0 |
| DIP_10004338 | hypothetical protein PFICI_12932 [Pestalotiopsis fici W106-1] | 72.4 | 0.0 |
| DIP_10004339 | hypothetical protein CFIO01_04872 [Colletotrichum fioriniae PJ7] | 34.9 | 1.0E-118 |
| DIP_10004340 | NA | | |
| DIP_10004341 | hypothetical protein PFICI_12936 [Pestalotiopsis fici W106-1] | 85.3 | 0.0 |
| DIP_10004342 | hypothetical protein PFICI_12935 [Pestalotiopsis fici W106-1] | 68 | 0.0 |
| DIP_10004343 | hypothetical protein PFICI_12934 [Pestalotiopsis fici W106-1] | 77 | 0.0 |
| DIP_10004344 | hypothetical protein FGSG_08061 [Fusarium graminearum PH-1] | 59.6 | 5.0E-155 |
| DIP_10004345 | hypothetical protein COCHEDRAFT_1149798, partial [Bipolaris maydis C5] | 36.5 | 1.0E-78 |
| DIP_10004346 | hypothetical protein V500_05890, partial [Pseudogymnoascus pannorum VKM F-4518 (FW-2643)] | 41.4 | 6.0E-21 |
| DIP_10004347 | quinone oxidoreductase, putative [Talaromyces marneffei ATCC 18224] | 61.9 | 8.0E-150 |
| DIP_10004348 | hypothetical protein PFICI_02656 [Pestalotiopsis fici W106-1] | 34.1 | 2.0E-68 |
| DIP_10004349 | hypothetical protein FOMG_07245 [Fusarium oxysporum f. sp. melonis 26406] | 53.3 | 0.0 |
| DIP_10004350 | NA | | |
| DIP_10004351 | hypothetical protein PFICI_12597 [Pestalotiopsis fici W106-1] | 74.4 | 0.0 |
| DIP_10004352 | NA | | |
| DIP_10004353 | NA | | |
| DIP_10004354 | putative rrna biogenesis protein rrp5 protein [Eutypa lata UCREL1] | 68.5 | 0.0 |
| DIP_10004355 | putative vacuolar-sorting protein snf8 protein [Eutypa lata UCREL1] | 84.2 | 1.0E-153 |
| DIP_10004356 | 30S ribosomal protein S9 [Metarhizium acridum CQMa 102] | 93.8 | 5.0E-100 |
| DIP_10004357 | putative eukaryotic ribosome biogenesis protein 1 protein [Eutypa lata UCREL1] | 81.9 | 0.0 |
| DIP_10004358 | unnamed protein product [Penicillium roqueforti FM164] | 52.9 | 0.0 |
| DIP_10004359 | Siderophore iron transporter mirB [Talaromyces islandicus] | 54.4 | 0.0 |
| DIP_10004360 | NA | | |
| DIP_10004361 | NA | | |
| DIP_10004362 | NA | | |
| DIP_10004363 | hypothetical protein PFICI_10109 [Pestalotiopsis fici W106-1] | 31.5 | 8.0E-08 |
| DIP_10004364 | hypothetical protein PFICI_00980 [Pestalotiopsis fici W106-1] | 59.5 | 0.0 |
| DIP_10004365 | Alternative oxidase [Pestalotiopsis fici W106-1] | 73.3 | 0.0 |
| DIP_10004366 | NA | | |
| DIP_10004367 | putative translation initiation factor eif3 subunit protein [Eutypa lata UCREL1] | 69.6 | 1.0E-107 |
| DIP_10004368 | putative mitochondrial distribution and morphology protein 31 protein [Eutypa lata UCREL1] | 77.5 | 0.0 |
| DIP_10004369 | putative pre-rrna-processing protein pno1 protein [Eutypa lata UCREL1] | 69.3 | 7.0E-137 |
| DIP_10004370 | hypothetical protein UCREL1_10209 [Eutypa lata UCREL1] | 34.5 | 4.0E-65 |
| DIP_10004371 | putative u6 snrna-associated sm-like protein lsm6 protein [Eutypa lata UCREL1] | 89.3 | 2.0E-48 |
| DIP_10004372 | putative methyltransferase protein [Eutypa lata UCREL1] | 67.3 | 1.0E-144 |
| DIP_10004373 | putative polysaccharide export protein [Eutypa lata UCREL1] | 62.9 | 2.0E-86 |
| DIP_10004374 | putative peroxisomal membrane protein 4 protein [Eutypa lata UCREL1] | 69.1 | 8.0E-121 |
| DIP_10004375 | NA | | |
| DIP_10004376 | NA | | |
| DIP_10004377 | hypothetical protein UCREL1_10216 [Eutypa lata UCREL1] | 73.4 | 8.0E-107 |
| DIP_10004378 | hypothetical protein PFICI_01568 [Pestalotiopsis fici W106-1] | 62.6 | 9.0E-78 |
| DIP_10004379 | putative mfs protein [Eutypa lata UCREL1] | 61.4 | 0.0 |
| DIP_10004380 | hypothetical protein S40288_04115 [Stachybotrys chartarum IBT 40288] | 53.1 | 3.0E-122 |
| DIP_10004381 | small nuclear ribonucleoprotein lsm2 [Colletotrichum orbiculare MAFF 240422] | 89.8 | 2.0E-60 |
| DIP_10004382 | hypothetical protein PFICI_01567 [Pestalotiopsis fici W106-1] | 65.2 | 0.0 |
| DIP_10004383 | hypothetical protein PFICI_01563 [Pestalotiopsis fici W106-1] | 68.1 | 0.0 |
| DIP_10004384 | putative dna-(apurinic or apyrimidinic site) lyase 1 protein [Eutypa lata UCREL1] | 65.2 | 0.0 |
| DIP_10004385 | putative checkpoint serine threonine-protein kinase bub1 protein [Togninia minima UCRPA7] | 57 | 0.0 |
| DIP_10004386 | NA | | |
| DIP_10004387 | putative transcription regulator protein [Eutypa lata UCREL1] | 52.8 | 8.0E-85 |
| DIP_10004388 | N-glycosylase/DNA lyase [Verticillium alfalfae VaMs.102] | 59.9 | 8.0E-168 |
| DIP_10004389 | hypothetical protein PFICI_01966 [Pestalotiopsis fici W106-1] | 81 | 0.0 |

| | | | |
|---|---|---|---|
| DIP_10004390 | NA | | |
| DIP_10004391 | NA | | |
| DIP_10004392 | putative Nop14-like family protein [Colletotrichum sublineola] | 55.1 | 0.0 |
| DIP_10004393 | putative cytoskeleton-associated protein [Eutypa lata UCREL1] | 55.3 | 2.0E-72 |
| DIP_10004394 | hypothetical protein PFICI_00420 [Pestalotiopsis fici W106-1] | 61.9 | 0.0 |
| DIP_10004395 | hypothetical protein V495_07713 [Pseudogymnoascus pannorum VKM F-4514 (FW-929)] | 72.5 | 3.0E-152 |
| DIP_10004396 | hypothetical protein PFICI_00414 [Pestalotiopsis fici W106-1] | 64.2 | 0.0 |
| DIP_10004397 | NA | | |
| DIP_10004398 | NA | | |
| DIP_10004399 | putative glycoside hydrolase family 38 protein [Eutypa lata UCREL1] | 78.8 | 0.0 |
| DIP_10004400 | capsule polysaccharide biosynthesis [Moniliophthora roreri MCA 2997] | 51 | 6.0E-157 |
| DIP_10004401 | NA | | |
| DIP_10004402 | putative pentatricopeptide repeat domain-containing protein [Eutypa lata UCREL1] | 45.5 | 0.0 |
| DIP_10004403 | hypothetical protein PFICI_00425 [Pestalotiopsis fici W106-1] | 45.8 | 0.0 |
| DIP_10004404 | hypothetical protein NCU02682 [Neurospora crassa OR74A] | 39.7 | 4.0E-40 |
| DIP_10004405 | hypothetical protein PFICI_00430 [Pestalotiopsis fici W106-1] | 80.8 | 0.0 |
| DIP_10004406 | hypothetical protein PFICI_00429 [Pestalotiopsis fici W106-1] | 43.4 | 1.0E-70 |
| DIP_10004407 | hypothetical protein PFICI_00428 [Pestalotiopsis fici W106-1] | 76.7 | 0.0 |
| DIP_10004408 | putative translocation protein sec66 protein [Eutypa lata UCREL1] | 78.3 | 7.0E-125 |
| DIP_10004409 | Carbamoyl-phosphate synthase arginine-specific large chain [Pestalotiopsis fici W106-1] | 85 | 0.0 |
| DIP_10004410 | putative pwwp domain-containing protein [Eutypa lata UCREL1] | 53.1 | 3.0E-153 |
| DIP_10004411 | hypothetical protein CGLO_15894 [Colletotrichum gloeosporioides Cg-14] | 59.5 | 6.0E-89 |
| DIP_10004412 | NA | | |
| DIP_10004413 | Serine/threonine-protein phosphatase 2A activator 2 [Pestalotiopsis fici W106-1] | 76.6 | 0.0 |
| DIP_10004414 | putative exosome complex subunit rrp46 protein [Eutypa lata UCREL1] | 68.3 | 4.0E-124 |
| DIP_10004415 | hypothetical protein PFICI_01898 [Pestalotiopsis fici W106-1] | 83.7 | 0.0 |
| DIP_10004416 | putative homeobox domain-containing protein [Colletotrichum sublineola] | 46.7 | 5.0E-137 |
| DIP_10004417 | putative exocyst complex component protein [Eutypa lata UCREL1] | 80.1 | 0.0 |
| DIP_10004418 | hypothetical protein UCREL1_2392 [Eutypa lata UCREL1] | 69.4 | 2.0E-150 |
| DIP_10004419 | putative dna repair family protein [Eutypa lata UCREL1] | 45.3 | 2.0E-118 |
| DIP_10004420 | tRNA (guanine(37)-N1)-methyltransferase [Pestalotiopsis fici W106-1] | 56.5 | 0.0 |
| DIP_10004421 | hypothetical protein PFICI_01892 [Pestalotiopsis fici W106-1] | 51.5 | 2.0E-67 |
| DIP_10004422 | hypothetical protein PFICI_01441 [Pestalotiopsis fici W106-1] | 42.7 | 0.0 |
| DIP_10004423 | NA | | |
| DIP_10004424 | putative glycoside hydrolase family 17 protein [Eutypa lata UCREL1] | 41.6 | 4.0E-113 |
| DIP_10004425 | hypothetical protein PFICI_01444 [Pestalotiopsis fici W106-1] | 48.7 | 5.0E-161 |
| DIP_10004426 | hypothetical protein VDAG_02808 [Verticillium dahliae VdLs.17] | 34.8 | 4.0E-32 |
| DIP_10004427 | putative chitin synthase 4 protein [Eutypa lata UCREL1] | 80.7 | 0.0 |
| DIP_10004428 | NA | | |
| DIP_10004429 | hypothetical protein PFICI_01447 [Pestalotiopsis fici W106-1] | 56 | 0.0 |
| DIP_10004430 | putative serine threonine protein kinase protein [Eutypa lata UCREL1] | 70.1 | 0.0 |
| DIP_10004431 | putative opa3 domain protein [Eutypa lata UCREL1] | 70.5 | 5.0E-107 |
| DIP_10004432 | putative pro41 protein [Eutypa lata UCREL1] | 79.3 | 7.0E-84 |
| DIP_10004433 | putative shwachman-bodian-diamond syndrome protein [Eutypa lata UCREL1] | 60 | 3.0E-49 |
| DIP_10004434 | hypothetical protein PFICI_01452 [Pestalotiopsis fici W106-1] | 59.9 | 0.0 |
| DIP_10004435 | 60S ribosomal protein L36 [Sclerotinia borealis F-4157] | 89.6 | 1.0E-62 |
| DIP_10004436 | putative ubx domain protein [Eutypa lata UCREL1] | 50 | 8.0E-173 |
| DIP_10004437 | putative proteasome component pup3 protein [Eutypa lata UCREL1] | 93.7 | 8.0E-143 |
| DIP_10004438 | hypothetical protein PFICI_01455 [Pestalotiopsis fici W106-1] | 84 | 0.0 |
| DIP_10004439 | small GTPase RanA [Aspergillus nidulans] | 90.2 | 2.0E-146 |
| DIP_10004440 | hypothetical protein PFICI_01453 [Pestalotiopsis fici W106-1] | 77.1 | 6.0E-128 |
| DIP_10004441 | hypothetical protein UCREL1_11015 [Eutypa lata UCREL1] | 36.1 | 8.0E-19 |
| DIP_10004442 | lipase [Verticillium dahliae VdLs.17] | 51.5 | 1.0E-130 |
| DIP_10004443 | putative 14-3-3 protein [Eutypa lata UCREL1] | 96.1 | 0.0 |
| DIP_10004444 | carboxylesterase [Colletotrichum gloeosporioides Cg-14] | 44.7 | 6.0E-165 |
| DIP_10004445 | NA | | |
| DIP_10004446 | hypothetical protein PFICI_01462 [Pestalotiopsis fici W106-1] | 66.2 | 0.0 |
| DIP_10004447 | putative pre-mrna-processing factor 39 protein [Eutypa lata UCREL1] | 80.3 | 0.0 |
| DIP_10004448 | hypothetical protein PFICI_03831 [Pestalotiopsis fici W106-1] | 69.6 | 0.0 |
| DIP_10004449 | hypothetical protein PFICI_03813 [Pestalotiopsis fici W106-1] | 32.6 | 3.0E-68 |
| DIP_10004450 | hypothetical protein PFICI_03834 [Pestalotiopsis fici W106-1] | 60.3 | 0.0 |
| DIP_10004451 | putative sphingosine n-acyltransferase lac1 protein [Eutypa lata UCREL1] | 76.9 | 0.0 |
| DIP_10004452 | NA | | |
| DIP_10004453 | putative atp synthase delta mitochondrial protein [Eutypa lata UCREL1] | 74.9 | 1.0E-95 |
| DIP_10004454 | NA | | |
| DIP_10004455 | hypothetical protein PFICI_03794 [Pestalotiopsis fici W106-1] | 54.6 | 4.0E-122 |
| DIP_10004456 | putative small monomeric gtpase protein [Eutypa lata UCREL1] | 95.7 | 6.0E-115 |
| DIP_10004457 | putative patatin-like phospholipase domain-containing protein [Togninia minima UCRPA7] | 69.5 | 0.0 |
| DIP_10004458 | putative guanine nucleotide exchange factor-like protein [Eutypa lata UCREL1] | 48.7 | 8.0E-169 |
| DIP_10004459 | conserved hypothetical protein [Verticillium alfalfae VaMs.102] | 48.3 | 3.0E-120 |
| DIP_10004460 | hypothetical protein UCRPA7_3854 [Togninia minima UCRPA7] | 43.5 | 1.0E-107 |
| DIP_10004461 | NA | | |
| DIP_10004462 | catechol -dioxygenase 1 [Colletotrichum orbiculare MAFF 240422] | 61.3 | 5.0E-128 |
| DIP_10004463 | NA | | |
| DIP_10004464 | hypothetical protein PFICI_05322 [Pestalotiopsis fici W106-1] | 73.7 | 0.0 |
| DIP_10004465 | putative superoxide dismutase 1 copper chaperone protein [Eutypa lata UCREL1] | 70.8 | 4.0E-143 |
| DIP_10004466 | putative 50s ribosomal protein l13 protein [Eutypa lata UCREL1] | 86.6 | 2.0E-117 |
| DIP_10004467 | hypothetical protein SS1G_00106 [Sclerotinia sclerotiorum 1980] | 73 | 4.0E-167 |
| DIP_10004468 | hypothetical protein PFICI_05328 [Pestalotiopsis fici W106-1] | 68.1 | 0.0 |
| DIP_10004469 | hypothetical protein PFICI_05321 [Pestalotiopsis fici W106-1] | 67.5 | 0.0 |
| DIP_10004470 | ATP-dependent RNA helicase DBP2 [Pestalotiopsis fici W106-1] | 85.7 | 0.0 |
| DIP_10004471 | NA | | |
| DIP_10004472 | NA | | |
| DIP_10004473 | hypothetical protein CH063_06991 [Colletotrichum higginsianum] | 57 | 3.0E-125 |
| DIP_10004474 | hypothetical protein PFICI_05316 [Pestalotiopsis fici W106-1] | 55.9 | 0.0 |
| DIP_10004475 | hypothetical protein PFICI_00976 [Pestalotiopsis fici W106-1] | 68.5 | 0.0 |
| DIP_10004476 | putative rna polymerase-associated protein [Eutypa lata UCREL1] | 62.6 | 4.0E-140 |
| DIP_10004477 | hypothetical protein PFICI_00973 [Pestalotiopsis fici W106-1] | 77.7 | 0.0 |
| DIP_10004478 | molybdopterin synthase sulfur carrier subunit [Magnaporthe oryzae 70-15] | 57.7 | 2.0E-36 |
| DIP_10004479 | putative 40s ribosomal protein s28 protein [Togninia minima UCRPA7] | 97.1 | 3.0E-41 |
| DIP_10004480 | NA | | |
| DIP_10004481 | hypothetical protein HIM_05662 [Hirsutella minnesotensis 3608] | 54.1 | 0.0 |
| DIP_10004482 | NA | | |
| DIP_10004483 | hypothetical protein HIM_05661 [Hirsutella minnesotensis 3608] | 48.9 | 6.0E-134 |
| DIP_10004484 | hypothetical protein VDAG_00274 [Verticillium dahliae VdLs.17] | 74.8 | 0.0 |
| DIP_10004485 | putative dynactin subunit protein [Eutypa lata UCREL1] | 67.5 | 0.0 |
| DIP_10004486 | hypothetical protein UCREL1_3683 [Eutypa lata UCREL1] | 53.4 | 4.0E-64 |
| DIP_10004487 | Adenylate cyclase [Pestalotiopsis fici W106-1] | 65.4 | 0.0 |
| DIP_10004488 | NA | | |
| DIP_10004489 | putative general substrate transporter protein [Eutypa lata UCREL1] | 84.1 | 0.0 |
| DIP_10004490 | putative fumarylacetoacetate hydrolase protein [Eutypa lata UCREL1] | 83.8 | 0.0 |
| DIP_10004491 | NA | | |
| DIP_10004492 | NA | | |
| DIP_10004493 | hypothetical protein THITE_125074 [Thielavia terrestris NRRL 8126] | 52.6 | 4.0E-65 |
| DIP_10004494 | hypothetical protein COCMIDRAFT_111002 [Bipolaris oryzae ATCC 44560] | 30.1 | 3.0E-19 |
| DIP_10004495 | hypothetical protein UCDDA912_g03680 [Diaporthe ampelina] | 64.3 | 3.0E-178 |
| DIP_10004496 | putative glycosyltransferase family 57 protein [Eutypa lata UCREL1] | 74.6 | 0.0 |
| DIP_10004497 | pH-response regulator protein palC [Colletotrichum graminicola M1.001] | 57.6 | 0.0 |
| DIP_10004498 | putative glycoside hydrolase family 2 protein [Eutypa lata UCREL1] | 63.4 | 0.0 |
| DIP_10004499 | DNA replication licensing factor mcm4 [Pestalotiopsis fici W106-1] | 79 | 0.0 |
| DIP_10004500 | hypothetical protein PV04_07170 [Capronia semiimmersa] | 51 | 4.0E-163 |
| DIP_10004501 | triacylglycerol lipase [Beauveria bassiana ARSEF 2860] | 45.8 | 3.0E-41 |
| DIP_10004502 | putative ku family dna protein [Eutypa lata UCREL1] | 67.1 | 0.0 |
| DIP_10004503 | hypothetical protein PFICI_07342 [Pestalotiopsis fici W106-1] | 65.4 | 1.0E-84 |
| DIP_10004504 | hypothetical protein UCREL1_4620 [Eutypa lata UCREL1] | 47.7 | 6.0E-12 |
| DIP_10004505 | hypothetical protein PFICI_07340 [Pestalotiopsis fici W106-1] | 63.3 | 9.0E-165 |
| DIP_10004506 | hypothetical protein PFICI_07339 [Pestalotiopsis fici W106-1] | 70.5 | 0.0 |
| DIP_10004507 | hypothetical protein PFICI_07338 [Pestalotiopsis fici W106-1] | 70.1 | 5.0E-145 |
| DIP_10004508 | NA | | |
| DIP_10004509 | hypothetical protein PFICI_08173 [Pestalotiopsis fici W106-1] | 62.2 | 0.0 |
| DIP_10004510 | NA | | |
| DIP_10004511 | hypothetical protein VDAG_06514 [Verticillium dahliae VdLs.17] | 62.1 | 0.0 |
| DIP_10004512 | putative upf0261 domain-containing protein [Eutypa lata UCREL1] | 55.7 | 1.0E-147 |
| DIP_10004513 | putative tim-barrel enzyme family protein [Eutypa lata UCREL1] | 64.6 | 3.0E-128 |
| DIP_10004514 | hypothetical protein PFICI_08269 [Pestalotiopsis fici W106-1] | 56.4 | 3.0E-56 |
| DIP_10004515 | putative swr1-complex protein 5 protein [Eutypa lata UCREL1] | 56.3 | 7.0E-107 |
| DIP_10004516 | hypothetical protein TRIATDRAFT_155450 [Trichoderma atroviride IMI 206040] | 62.9 | 5.0E-121 |
| DIP_10004517 | NA | | |
| DIP_10004518 | NA | | |
| DIP_10004519 | NA | | |
| DIP_10004520 | hypothetical protein PFICI_08130 [Pestalotiopsis fici W106-1] | 36.6 | 2.0E-70 |
| DIP_10004521 | NA | | |
| DIP_10004522 | hypothetical protein O988_09707 [Pseudogymnoascus pannorum VKM F-3808] | 31.4 | 1.0E-16 |
| DIP_10004523 | putative 6-phosphogluconolactonase protein [Eutypa lata UCREL1] | 82.8 | 3.0E-161 |
| DIP_10004524 | putative 4-nitrophenylphosphatase protein [Eutypa lata UCREL1] | 80.8 | 0.0 |
| DIP_10004525 | hypothetical protein PV07_09995 [Cladophialophora immunda] | 30.8 | 1.0E-19 |
| DIP_10004526 | hypothetical protein PFICI_08133 [Pestalotiopsis fici W106-1] | 63.3 | 9.0E-87 |
| DIP_10004527 | hypothetical protein PFICI_08142 [Pestalotiopsis fici W106-1] | 56.4 | 7.0E-99 |
| DIP_10004528 | 3-oxoacyl-(acyl-carrier-protein)-synthase [Sordaria macrospora k-hell] | 71.7 | 0.0 |
| DIP_10004529 | hypothetical protein PFICI_08144 [Pestalotiopsis fici W106-1] | 70.9 | 0.0 |
| DIP_10004530 | hypothetical protein PFICI_13806 [Pestalotiopsis fici W106-1] | 31 | 1.0E-49 |
| DIP_10004531 | hypothetical protein CSUB01_06050 [Colletotrichum sublineola] | 35.3 | 2.0E-40 |
| DIP_10004532 | hypothetical protein UCREL1_4545 [Eutypa lata UCREL1] | 52.4 | 5.0E-124 |
| DIP_10004533 | hypothetical protein FOYG_16402 [Fusarium oxysporum FOSC 3-a] | 59.3 | 2.0E-60 |
| DIP_10004534 | NA | | |
| DIP_10004535 | NA | | |
| DIP_10004536 | NA | | |
| DIP_10004537 | putative cysteine desulfurase protein [Eutypa lata UCREL1] | 64.4 | 0.0 |
| DIP_10004538 | ATP-dependent RNA helicase mrh-4 [Pestalotiopsis fici W106-1] | 68.2 | 0.0 |
| DIP_10004539 | NA | | |
| DIP_10004540 | putative duf1212 domain membrane protein prm10 protein [Eutypa lata UCREL1] | 60.9 | 0.0 |
| DIP_10004541 | hypothetical protein MYCTH_2307560 [Myceliophthora thermophila ATCC 42464] | 34.4 | 5.0E-64 |
| DIP_10004542 | hypothetical protein FOZG_05402 [Fusarium oxysporum Fo47] | 36.4 | 3.0E-66 |
| DIP_10004543 | NA | | |
| DIP_10004544 | hypothetical protein PFICI_00963 [Pestalotiopsis fici W106-1] | 59.2 | 0.0 |
| DIP_10004545 | hypothetical protein CFIO01_01905 [Colletotrichum fioriniae PJ7] | 65.7 | 0.0 |
| DIP_10004546 | isochorismatase family protein [Aspergillus fumigatus Af293] | 44.5 | 3.0E-35 |
| DIP_10004547 | hypothetical protein PFICI_08267 [Pestalotiopsis fici W106-1] | 59 | 0.0 |
| DIP_10004548 | putative nuclear protein involved in pre-rrna protein [Eutypa lata UCREL1] | 45.7 | 6.0E-76 |
| DIP_10004549 | putative rrna-processing protein [Eutypa lata UCREL1] | 31.8 | 2.0E-37 |
| DIP_10004550 | putative trna 2 -phosphotransferase 1 protein [Eutypa lata UCREL1] | 62 | 5.0E-110 |
| DIP_10004551 | NA | | |
| DIP_10004552 | hypothetical protein FAVG1_05543 [Fusarium avenaceum] | 42.2 | 4.0E-117 |
| DIP_10004553 | NA | | |
| DIP_10004554 | NA | | |
| DIP_10004555 | hypothetical protein PFICI_08303 [Pestalotiopsis fici W106-1] | 54.7 | 0.0 |
| DIP_10004556 | hypothetical protein PFICI_08302 [Pestalotiopsis fici W106-1] | 72.7 | 0.0 |
| DIP_10004557 | hypothetical protein FGSG_06979 [Fusarium graminearum PH-1] | 82.7 | 0.0 |
| DIP_10004558 | hypothetical protein PFICI_08300 [Pestalotiopsis fici W106-1] | 43.6 | 2.0E-148 |
| DIP_10004559 | hypothetical protein PFICI_08129 [Pestalotiopsis fici W106-1] | 57.9 | 0.0 |
| DIP_10004560 | hypothetical protein PFICI_08149 [Pestalotiopsis fici W106-1] | 66.9 | 0.0 |
| DIP_10004561 | hypothetical protein PFICI_07494 [Pestalotiopsis fici W106-1] | 66 | 0.0 |
| DIP_10004562 | putative ubiquitin carboxyl-terminal hydrolase protein [Eutypa lata UCREL1] | 55.4 | 0.0 |
| DIP_10004563 | putative peptidase m16 inactive domain-containing protein [Eutypa lata UCREL1] | 72.7 | 0.0 |
| DIP_10004564 | hypothetical protein MYCTH_2304725 [Myceliophthora thermophila ATCC 42464] | 38.7 | 9.0E-40 |
| DIP_10004565 | hypothetical protein PFICI_07547 [Pestalotiopsis fici W106-1] | 74.4 | 3.0E-93 |
| DIP_10004566 | putative upf0047 domain protein [Diaporthe ampelina] | 69.1 | 2.0E-100 |
| DIP_10004567 | putative cleavage and polyadenylation specificity factor subunit 5 protein [Eutypa lata UCREL1] | 76.7 | 1.0E-143 |
| DIP_10004568 | conserved hypothetical protein [Talaromyces stipitatus ATCC 10500] | 30.4 | 3.0E-58 |
| DIP_10004569 | hypothetical protein ACRE_012660 [Acremonium chrysogenum ATCC 11550] | 32.4 | 5.0E-28 |
| DIP_10004570 | WD40 repeat-like protein [Neurospora tetrasperma FGSC 2509] | 48.4 | 3.0E-153 |
| DIP_10004571 | hypothetical protein THAR02_06735 [Trichoderma harzianum] | 65 | 0.0 |
| DIP_10004572 | hypothetical protein PFICI_07540 [Pestalotiopsis fici W106-1] | 49.3 | 1.0E-80 |
| DIP_10004573 | putative fatty acid elongase-like protein [Eutypa lata UCREL1] | 67.9 | 0.0 |
| DIP_10004574 | putative long chain fatty alcohol oxidase protein [Eutypa lata UCREL1] | 50.1 | 0.0 |
| DIP_10004575 | hypothetical protein Z518_02514 [Rhinocladiella mackenziei CBS 650.93] | 39.3 | 1.0E-41 |
| DIP_10004576 | putative f-box domain-containing protein [Eutypa lata UCREL1] | 57.1 | 1.0E-123 |
| DIP_10004577 | calcium/proton exchanger [Cladophialophora bantiana CBS 173.52] | 41.7 | 3.0E-96 |
| DIP_10004578 | NA | | |
| DIP_10004579 | NA | | |
| DIP_10004580 | hypothetical protein CGLO_03011 [Colletotrichum gloeosporioides Cg-14] | 76.7 | 0.0 |
| DIP_10004581 | NA | | |
| DIP_10004582 | repetitive proline-rich cell wall protein [Colletotrichum higginsianum] | 38.9 | 8.0E-47 |
| DIP_10004583 | NA | | |
| DIP_10004584 | hypothetical protein PFICI_11639 [Pestalotiopsis fici W106-1] | 50.1 | 0.0 |
| DIP_10004585 | NA | | |
| DIP_10004586 | NA | | |
| DIP_10004587 | hypothetical protein SAPIO_CDS1298 [Scedosporium apiospermum] | 60 | 2.0E-33 |
| DIP_10004588 | NA | | |
| DIP_10004589 | NA | | |
| DIP_10004590 | NA | | |
| DIP_10004591 | Endonuclease/Exonuclease/phosphatase family protein [Metarhizium anisopliae] | 54 | 4.0E-100 |
| DIP_10004592 | hypothetical protein CGGC5_5601 [Colletotrichum gloeosporioides Nara gc5] | 57.5 | 9.0E-121 |
| DIP_10004593 | hypothetical protein PFICI_12121 [Pestalotiopsis fici W106-1] | 73.8 | 0.0 |
| DIP_10004594 | hypothetical protein NCU06464 [Neurospora crassa OR74A] | 64.2 | 5.0E-75 |
| DIP_10004595 | hypothetical protein PFICI_12123 [Pestalotiopsis fici W106-1] | 40.3 | 2.0E-29 |
| DIP_10004596 | hypothetical protein PFICI_12124 [Pestalotiopsis fici W106-1] | 46 | 5.0E-95 |
| DIP_10004597 | putative pentatricopeptide repeat protein [Eutypa lata UCREL1] | 47.1 | 0.0 |
| DIP_10004598 | putative histidine acid phosphatase protein [Eutypa lata UCREL1] | 63.9 | 0.0 |
| DIP_10004599 | putative negative regulator of differentiation 1 protein [Togninia minima UCRPA7] | 78.6 | 0.0 |
| DIP_10004600 | hypothetical protein PFICI_12114 [Pestalotiopsis fici W106-1] | 82.2 | 0.0 |
| DIP_10004601 | cell division control protein [Trichoderma reesei QM6a] | 97.9 | 4.0E-141 |
| DIP_10004602 | hypothetical protein Cob_12478 [Colletotrichum orbiculare MAFF 240422] | 47.5 | 9.0E-59 |
| DIP_10004603 | putative fad-dependent pyridine nucleotide-disulfide oxidoreductase protein [Togninia minima UCRPA7] | 49.8 | 2.0E-179 |
| DIP_10004604 | ataxia telangiectasia mutated [Verticillium alfalfae VaMs.102] | 51 | 0.0 |
| DIP_10004605 | putative nuclear pore complex subunit protein [Eutypa lata UCREL1] | 66.8 | 0.0 |
| DIP_10004606 | putative dicarboxylic amino acid permease protein [Eutypa lata UCREL1] | 77.7 | 0.0 |
| DIP_10004607 | DNA polymerase gamma [Pestalotiopsis fici W106-1] | 72 | 0.0 |
| DIP_10004608 | hypothetical protein GLRG_00056 [Colletotrichum graminicola M1.001] | 39.7 | 1.0E-29 |
| DIP_10004609 | hypothetical protein PFICI_12154 [Pestalotiopsis fici W106-1] | 63.9 | 0.0 |
| DIP_10004610 | putative duf1941 family protein [Eutypa lata UCREL1] | 55.5 | 0.0 |
| DIP_10004611 | NA | | |
| DIP_10004612 | NA | | |
| DIP_10004613 | hypothetical protein PFICI_11514 [Pestalotiopsis fici W106-1] | 50.8 | 5.0E-108 |
| DIP_10004614 | putative glucan -beta-glucosidase protein [Eutypa lata UCREL1] | 67.8 | 0.0 |
| DIP_10004615 | inositol phospholipid synthesis protein Scs3p [Colletotrichum gloeosporioides Cg-14] | 57.2 | 4.0E-117 |
| DIP_10004616 | putative arylsulfatase precursor [Diaporthe ampelina] | 62.6 | 0.0 |
| DIP_10004617 | NA | | |
| DIP_10004618 | hypothetical protein PFICI_08146 [Pestalotiopsis fici W106-1] | 59.9 | 0.0 |
| DIP_10004619 | hypothetical protein UCREL1_11419 [Eutypa lata UCREL1] | 70.7 | 5.0E-74 |
| DIP_10004620 | putative nadp-dependent alcohol dehydrogenase 6 protein [Eutypa lata UCREL1] | 75.6 | 0.0 |
| DIP_10004621 | hypothetical protein GLRG_08000 [Colletotrichum graminicola M1.001] | 33.5 | 1.0E-20 |
| DIP_10004622 | putative 40s ribosomal protein s7 protein [Eutypa lata UCREL1] | 93.1 | 8.0E-135 |
| DIP_10004623 | putative dynactin arp1 p62 subunit ro2 protein [Eutypa lata UCREL1] | 55.8 | 0.0 |
| DIP_10004624 | hypothetical protein PFICI_08116 [Pestalotiopsis fici W106-1] | 53.2 | 0.0 |
| DIP_10004625 | hypothetical protein PFICI_08119 [Pestalotiopsis fici W106-1] | 62.6 | 4.0E-97 |
| DIP_10004626 | putative fad linked oxidase protein [Eutypa lata UCREL1] | 70.3 | 2.0E-120 |
| DIP_10004627 | putative secreted protein [Eutypa lata UCREL1] | 63.6 | 5.0E-122 |
| DIP_10004628 | putative zinc finger domain-containing protein [Eutypa lata UCREL1] | 63.3 | 1.0E-159 |
| DIP_10004629 | hypothetical protein UCREL1_2037 [Eutypa lata UCREL1] | 34.1 | 1.0E-18 |
| DIP_10004630 | conserved hypothetical protein [Verticillium alfalfae VaMs.102] | 64.6 | 1.0E-149 |
| DIP_10004631 | putative fungal zn binuclear cluster domain containing protein [Eutypa lata UCREL1] | 78.4 | 0.0 |
| DIP_10004632 | putative 60s ribosomal protein l10a protein [Eutypa lata UCREL1] | 94.5 | 3.0E-141 |
| DIP_10004633 | hypothetical protein PFICI_08128 [Pestalotiopsis fici W106-1] | 63.2 | 0.0 |
| DIP_10004634 | putative pq loop repeat protein [Eutypa lata UCREL1] | 71 | 5.0E-173 |
| DIP_10004635 | hypothetical protein PFICI_08297 [Pestalotiopsis fici W106-1] | 60.6 | 2.0E-78 |
| DIP_10004636 | putative methyltransferase domain-containing protein [Eutypa lata UCREL1] | 45.2 | 3.0E-41 |
| DIP_10004637 | hypothetical protein PFICI_08295 [Pestalotiopsis fici W106-1] | 56.5 | 0.0 |
| DIP_10004638 | putative peroxisomal -enoyl- isomerase protein [Eutypa lata UCREL1] | 68.3 | 1.0E-139 |
| DIP_10004639 | putative pyruvate dehydrogenase protein [Eutypa lata UCREL1] | 61.4 | 0.0 |
| DIP_10004640 | hypothetical protein UCREL1_8157 [Eutypa lata UCREL1] | 53.6 | 2.0E-86 |
| DIP_10004641 | putative alpha beta hydrolase family protein [Eutypa lata UCREL1] | 61.3 | 4.0E-168 |
| DIP_10004642 | short chain dehydrogenase [Colletotrichum graminicola M1.001] | 48.8 | 1.0E-73 |
| DIP_10004643 | hypothetical protein PFICI_00470 [Pestalotiopsis fici W106-1] | 56 | 0.0 |
| DIP_10004644 | cupin [Colletotrichum fioriniae PJ7] | 61.5 | 5.0E-73 |
| DIP_10004645 | conserved hypothetical protein [Aspergillus clavatus NRRL 1] | 51.9 | 1.0E-52 |
| DIP_10004646 | succinate dehydrogenase flavoprotein subunit [Verticillium dahliae VdLs.17] | 88.3 | 0.0 |
| DIP_10004647 | hypothetical protein PFICI_00251 [Pestalotiopsis fici W106-1] | 70.9 | 0.0 |
| DIP_10004648 | hypothetical protein UCREL1_6932 [Eutypa lata UCREL1] | 53.2 | 7.0E-74 |
| DIP_10004649 | nucleolar protein CgrA [Beauveria bassiana ARSEF 2860] | 74.5 | 3.0E-49 |
| DIP_10004650 | NA | | |
| DIP_10004651 | hypothetical protein BC1G_05787 [Botrytis cinerea B05.10] | 44.7 | 5.0E-53 |
| DIP_10004652 | putative mannan polymerase complexes mnn9 subunit protein [Eutypa lata UCREL1] | 71.2 | 3.0E-163 |
| DIP_10004653 | putative npl4 family protein [Eutypa lata UCREL1] | 76.2 | 0.0 |
| DIP_10004654 | putative sec63 brl domain-containing protein [Eutypa lata UCREL1] | 84.3 | 0.0 |
| DIP_10004655 | hypothetical protein UCREL1_11508 [Eutypa lata UCREL1] | 37 | 3.0E-93 |
| DIP_10004656 | hypothetical protein CFIO01_03011 [Colletotrichum fioriniae PJ7] | 44.7 | 6.0E-49 |
| DIP_10004657 | NA | | |
| DIP_10004658 | hypothetical protein PFICI_00572 [Pestalotiopsis fici W106-1] | 58.5 | 0.0 |
| DIP_10004659 | Replication factor C subunit 2 [Pestalotiopsis fici W106-1] | 82.8 | 0.0 |
| DIP_10004660 | hypothetical protein PFICI_00576 [Pestalotiopsis fici W106-1] | 72.2 | 0.0 |
| DIP_10004661 | putative sterol-4-alpha-carboxylate 3-dehydrogenase protein [Eutypa lata UCREL1] | 72.2 | 0.0 |
| DIP_10004662 | hypothetical protein PFICI_00583 [Pestalotiopsis fici W106-1] | 63.4 | 0.0 |
| DIP_10004663 | putative hmg box protein [Togninia minima UCRPA7] | 41.4 | 2.0E-33 |
| DIP_10004664 | putative pre-mrna-processing atp-dependent rna helicase prp5 protein [Eutypa lata UCREL1] | 68.5 | 0.0 |
| DIP_10004665 | conserved hypothetical protein [Byssochlamys spectabilis No. 5] | 54.7 | 5.0E-29 |
| DIP_10004666 | putative 60s ribosomal protein l35 protein [Eutypa lata UCREL1] | 89.4 | 3.0E-74 |
| DIP_10004667 | hypothetical protein PFICI_01504 [Pestalotiopsis fici W106-1] | 78.7 | 4.0E-134 |
| DIP_10004668 | hypothetical protein THITE_2112968 [Thielavia terrestris NRRL 8126] | 61.9 | 5.0E-142 |
| DIP_10004669 | hypothetical protein PFICI_00856 [Pestalotiopsis fici W106-1] | 62.1 | 0.0 |
| DIP_10004670 | putative thermoresistant gluconokinase protein [Eutypa lata UCREL1] | 76.4 | 2.0E-120 |
| DIP_10004671 | hypothetical protein CGLO_10841 [Colletotrichum gloeosporioides Cg-14] | 41.6 | 5.0E-28 |
| DIP_10004672 | NA | | |
| DIP_10004673 | putative fk506-binding protein 2 precursor protein [Eutypa lata UCREL1] | 63.3 | 1.0E-129 |
| DIP_10004674 | putative pre-mrna branch site protein p14 protein [Eutypa lata UCREL1] | 38.6 | 2.0E-20 |
| DIP_10004675 | Di-copper centre-containing protein [Aureobasidium namibiae CBS 147.97] | 54.3 | 1.0E-159 |
| DIP_10004676 | hypothetical protein PFICI_01374 [Pestalotiopsis fici W106-1] | 39.8 | 2.0E-75 |
| DIP_10004677 | hypothetical protein HIM_07006 [Hirsutella minnesotensis 3608] | 42.2 | 1.0E-24 |
| DIP_10004678 | putative adp-ribosylation factor family protein [Eutypa lata UCREL1] | 93.5 | 2.0E-94 |
| DIP_10004679 | DNA-directed RNA polymerase I subunit RPA2 [Pestalotiopsis fici W106-1] | 82.7 | 0.0 |
| DIP_10004680 | DNA polymerase lambda [Verticillium dahliae VdLs.17] | 45.2 | 0.0 |
| DIP_10004681 | hypothetical protein PFICI_14116 [Pestalotiopsis fici W106-1] | 63.5 | 0.0 |
| DIP_10004682 | putative hnrnp arginine n-methyltransferase protein [Eutypa lata UCREL1] | 86.6 | 0.0 |
| DIP_10004683 | putative n-acetylglucosamine-phosphate mutase protein [Eutypa lata UCREL1] | 86 | 0.0 |
| DIP_10004684 | crossover junction endonuclease MUS81 [Verticillium dahliae VdLs.17] | 59.2 | 0.0 |
| DIP_10004685 | putative riboflavin synthase alpha chain protein [Eutypa lata UCREL1] | 74.1 | 2.0E-127 |
| DIP_10004686 | hypothetical protein PFICI_01389 [Pestalotiopsis fici W106-1] | 65.5 | 7.0E-162 |
| DIP_10004687 | putative zinc finger transcription factor ace1 protein [Eutypa lata UCREL1] | 61 | 0.0 |
| DIP_10004688 | NA | | |
| DIP_10004689 | putative endoribonuclease l-psp protein [Eutypa lata UCREL1] | 75.2 | 7.0E-67 |
| DIP_10004690 | dolichyl-phosphate-mannose-protein mannosyltransferase [Colletotrichum fioriniae PJ7] | 79.5 | 0.0 |
| DIP_10004691 | hypothetical protein CGLO_06814 [Colletotrichum gloeosporioides Cg-14] | 46.3 | 4.0E-112 |
| DIP_10004692 | putative fumarylacetoacetate hydrolase domain-containing protein [Eutypa lata UCREL1] | 81.1 | 1.0E-142 |
| DIP_10004693 | hypothetical protein PFICI_01397 [Pestalotiopsis fici W106-1] | 51.2 | 1.0E-36 |
| DIP_10004694 | hypothetical protein PFICI_12861 [Pestalotiopsis fici W106-1] | 71.5 | 0.0 |
| DIP_10004695 | alternative sulfate transporter [Colletotrichum orbiculare MAFF 240422] | 70.7 | 0.0 |
| DIP_10004696 | putative pwi domain mrna processing protein [Eutypa lata UCREL1] | 86.1 | 1.0E-73 |
| DIP_10004697 | putative histone acetyltransferase subunit 4 protein [Eutypa lata UCREL1] | 60.6 | 3.0E-56 |
| DIP_10004698 | hypothetical protein PFICI_01400 [Pestalotiopsis fici W106-1] | 63.9 | 0.0 |
| DIP_10004699 | NA | | |
| DIP_10004700 | putative atp-dependent rna helicase eif4a protein [Eutypa lata UCREL1] | 98.5 | 0.0 |
| DIP_10004701 | F-box and domain-containing protein [Colletotrichum gloeosporioides Nara gc5] | 76.9 | 9.0E-136 |
| DIP_10004702 | putative tlg2-vesicle protein of 38 kda protein [Eutypa lata UCREL1] | 47.4 | 5.0E-122 |
| DIP_10004703 | putative proline iminopeptidase protein [Eutypa lata UCREL1] | 61.5 | 0.0 |
| DIP_10004704 | hypothetical protein UCREL1_10773 [Eutypa lata UCREL1] | 52.9 | 6.0E-120 |
| DIP_10004705 | hypothetical protein PFICI_02607 [Pestalotiopsis fici W106-1] | 56.3 | 2.0E-131 |
| DIP_10004706 | hypothetical protein PFICI_10997 [Pestalotiopsis fici W106-1] | 71.3 | 2.0E-78 |
| DIP_10004707 | putative c6 zinc finger protein [Eutypa lata UCREL1] | 50.4 | 0.0 |
| DIP_10004708 | NA | | |
| DIP_10004709 | putative kinesin related protein 1 protein [Eutypa lata UCREL1] | 72.4 | 0.0 |
| DIP_10004710 | putative acyltransferase-like protein [Eutypa lata UCREL1] | 65.4 | 3.0E-139 |
| DIP_10004711 | putative dihydroxy-acid dehydratase protein [Eutypa lata UCREL1] | 81.5 | 0.0 |
| DIP_10004712 | conserved hypothetical protein [Talaromyces marneffei ATCC 18224] | 44.6 | 2.0E-72 |
| DIP_10004713 | hypothetical protein UCREL1_10249 [Eutypa lata UCREL1] | 56.1 | 8.0E-50 |
| DIP_10004714 | hypothetical protein SMAC_02566 [Sordaria macrospora k-hell] | 55.2 | 1.0E-11 |
| DIP_10004715 | hypothetical protein TRIVIDRAFT_209932 [Trichoderma virens Gv29-8] | 76.8 | 0.0 |
| DIP_10004716 | hypothetical protein PFICI_11186 [Pestalotiopsis fici W106-1] | 67.4 | 0.0 |
| DIP_10004717 | NA | | |
| DIP_10004718 | hypothetical protein PFICI_11187 [Pestalotiopsis fici W106-1] | 66 | 0.0 |
| DIP_10004719 | putative flavo protein monooxygenase [Diaporthe ampelina] | 42.1 | 1.0E-117 |
| DIP_10004720 | hypothetical protein PFICI_11193 [Pestalotiopsis fici W106-1] | 69.1 | 2.0E-172 |
| DIP_10004721 | hypothetical protein PFICI_14415 [Pestalotiopsis fici W106-1] | 44.4 | 1.0E-142 |
| DIP_10004722 | putative taf5-like protein [Eutypa lata UCREL1] | 68.3 | 0.0 |
| DIP_10004723 | hypothetical protein PFICI_11004 [Pestalotiopsis fici W106-1] | 51.1 | 1.0E-48 |
| DIP_10004724 | hypothetical protein FOMG_02811 [Fusarium oxysporum f. sp. melonis 26406] | 80.6 | 0.0 |
| DIP_10004725 | putative 50s ribosomal protein [Eutypa lata UCREL1] | 53.8 | 2.0E-50 |
| DIP_10004726 | putative flavin reductase like domain-containing protein [Eutypa lata UCREL1] | 34.1 | 4.0E-40 |
| DIP_10004727 | heat shock protein 90 [Grosmannia clavigera kw1407] | 88.7 | 0.0 |
| DIP_10004728 | hypothetical protein PFICI_11165 [Pestalotiopsis fici W106-1] | 38.5 | 2.0E-81 |
| DIP_10004729 | ATP-dependent clp protease proteolytic subunit [Colletotrichum orbiculare MAFF 240422] | 45.9 | 3.0E-74 |
| DIP_10004730 | hypothetical protein GMDG_07437 [Pseudogymnoascus destructans 20631-21] | 39.1 | 2.0E-40 |
| DIP_10004731 | NA | | |
| DIP_10004732 | calcineurin-like phosphoesterase [Colletotrichum fioriniae PJ7] | 45.6 | 3.0E-94 |
| DIP_10004733 | hypothetical protein PFICI_11192 [Pestalotiopsis fici W106-1] | 46.2 | 2.0E-46 |
| DIP_10004734 | putative sterol o-acyltransferase protein [Eutypa lata UCREL1] | 75.5 | 0.0 |
| DIP_10004735 | NA | | |
| DIP_10004736 | putative peptidase family m28 family protein [Togninia minima UCRPA7] | 49.1 | 3.0E-175 |
| DIP_10004737 | hypothetical protein PFICI_06121 [Pestalotiopsis fici W106-1] | 56 | 3.0E-177 |
| DIP_10004738 | putative branchpoint-bridging protein [Eutypa lata UCREL1] | 81.5 | 0.0 |
| DIP_10004739 | putative ras-like gene member b protein [Eutypa lata UCREL1] | 36 | 9.0E-57 |
| DIP_10004740 | hypothetical protein PFICI_00747 [Pestalotiopsis fici W106-1] | 55.2 | 2.0E-100 |
| DIP_10004741 | putative glucan 4-alpha-glucosidase-like protein [Togninia minima UCRPA7] | 53.3 | 0.0 |
| DIP_10004742 | hypothetical protein CGLO_13354 [Colletotrichum gloeosporioides Cg-14] | 41.8 | 8.0E-31 |
| DIP_10004743 | cytokinesis protein sepA [Marssonina brunnea f. sp. 'multigermtubi' MB_m1] | 64.5 | 0.0 |
| DIP_10004744 | hypothetical protein CH063_04514 [Colletotrichum higginsianum] | 53.3 | 2.0E-45 |
| DIP_10004745 | checkpoint protein hus1 [Magnaporthe oryzae 70-15] | 58.3 | 4.0E-138 |
| DIP_10004746 | DEAD-like helicase, partial [Metarhizium anisopliae ARSEF 549] | 41.2 | 0.0 |
| DIP_10004747 | hypothetical protein FOQG_17081 [Fusarium oxysporum f. sp. raphani 54005] | 44.5 | 0.0 |
| DIP_10004748 | putative copii coat assembly protein sec16 protein [Togninia minima UCRPA7] | 56.2 | 0.0 |
| DIP_10004749 | putative sap domain protein [Eutypa lata UCREL1] | 52 | 2.0E-46 |
| DIP_10004750 | NA | | |
| DIP_10004751 | putative leucyl-trna synthetase protein [Eutypa lata UCREL1] | 51.2 | 0.0 |
| DIP_10004752 | putative asc-type amino acid transporter 1 protein [Eutypa lata UCREL1] | 76.4 | 0.0 |
| DIP_10004753 | peptidase M13 [Aspergillus fumigatus var. RP-2014] | 30.2 | 2.0E-57 |
| DIP_10004754 | NA | | |
| DIP_10004755 | NA | | |

| | | | |
|---|---|---|---|
| DIP_10004756 | nuclear pore complex component [Colletotrichum graminicola M1.001] | 52.1 | 5.0E-72 |
| DIP_10004757 | putative amp-dependent synthetase ligase protein [Eutypa lata UCREL1] | 71.2 | 0.0 |
| DIP_10004758 | hypothetical protein PFICI_08183 [Pestalotiopsis fici W106-1] | 69.2 | 5.0E-165 |
| DIP_10004759 | hypothetical protein PFICI_08182 [Pestalotiopsis fici W106-1] | 42.9 | 4.0E-153 |
| DIP_10004760 | putative farnesyl pyrophosphate synthetase protein [Eutypa lata UCREL1] | 81.3 | 0.0 |
| DIP_10004761 | putative anaphase-promoting complex subunit 8 protein [Eutypa lata UCREL1] | 75.4 | 0.0 |
| DIP_10004762 | putative nop16-like protein [Eutypa lata UCREL1] | 52.1 | 4.0E-67 |
| DIP_10004763 | hypothetical protein PFICI_07761 [Pestalotiopsis fici W106-1] | 62.7 | 0.0 |
| DIP_10004764 | NA | | |
| DIP_10004765 | putative glutathione peroxidase [Colletotrichum sublineola] | 84.5 | 1.0E-106 |
| DIP_10004766 | conserved hypothetical protein [Neurospora crassa] | 50.8 | 5.0E-160 |
| DIP_10004767 | dihydrodipicolinate synthase [Gaeumannomyces graminis var. tritici R3-111a-1] | 71.3 | 2.0E-161 |
| DIP_10004768 | putative d-galacturonic acid reductase protein [Eutypa lata UCREL1] | 81.7 | 0.0 |
| DIP_10004769 | hypothetical protein ACRE_014710 [Acremonium chrysogenum ATCC 11550] | 30.4 | 2.0E-43 |
| DIP_10004770 | putative glucose-repressible protein [Eutypa lata UCREL1] | 74.7 | 6.0E-29 |
| DIP_10004771 | hypothetical protein PFICI_07766 [Pestalotiopsis fici W106-1] | 78.2 | 6.0E-116 |
| DIP_10004772 | NA | | |
| DIP_10004773 | NA | | |
| DIP_10004774 | putative nitrilase [Diplodia seriata] | 54.3 | 2.0E-113 |
| DIP_10004775 | NA | | |
| DIP_10004776 | hypothetical protein FOVG_17417 [Fusarium oxysporum f. sp. pisi HDV247] | 51 | 8.0E-57 |
| DIP_10004777 | hypothetical protein CGLO_17107 [Colletotrichum gloeosporioides Cg-14] | 58.2 | 0.0 |
| DIP_10004778 | hypothetical protein FG05_07591 [Fusarium graminearum] | 40.4 | 2.0E-100 |
| DIP_10004779 | hypothetical protein S40293_04787 [Stachybotrys chartarum IBT 40293] | 50.5 | 5.0E-108 |
| DIP_10004780 | hypothetical protein PFICI_11034 [Pestalotiopsis fici W106-1] | 59.3 | 3.0E-104 |
| DIP_10004781 | putative proteasome subunit alpha type-1 protein [Eutypa lata UCREL1] | 86.4 | 2.0E-157 |
| DIP_10004782 | vacuolar protein sorting 55 [Colletotrichum gloeosporioides Cg-14] | 92 | 1.0E-78 |
| DIP_10004783 | hypothetical protein PFICI_08348 [Pestalotiopsis fici W106-1] | 42.2 | 0.0 |
| DIP_10004784 | hypothetical protein PFICI_13116 [Pestalotiopsis fici W106-1] | 39.1 | 8.0E-107 |
| DIP_10004785 | putative bur asp-box repeat domain protein [Eutypa lata UCREL1] | 30.4 | 4.0E-12 |
| DIP_10004786 | CMGC/SRPK protein kinase [Trichophyton equinum CBS 127.97] | 56 | 2.0E-164 |
| DIP_10004787 | alpha/beta-Hydrolase [Glarea lozoyensis ATCC 20868] | 41.8 | 4.0E-47 |
| DIP_10004788 | hypothetical protein V501_00489 [Pseudogymnoascus pannorum VKM F-4519 (FW-2642)] | 35.8 | 2.0E-151 |
| DIP_10004789 | NA | | |
| DIP_10004790 | NA | | |
| DIP_10004791 | NA | | |
| DIP_10004792 | NA | | |
| DIP_10004793 | hypothetical protein V490_08030 [Pseudogymnoascus pannorum VKM F-3557] | 57 | 0.0 |
| DIP_10004794 | hypothetical protein V490_08026 [Pseudogymnoascus pannorum VKM F-3557] | 41.1 | 3.0E-161 |
| DIP_10004795 | NA | | |
| DIP_10004796 | hypothetical protein V500_02669 [Pseudogymnoascus pannorum VKM F-4518 (FW-2643)] | 30.8 | 3.0E-13 |
| DIP_10004797 | putative polyol transporter-like protein [Acremonium chrysogenum ATCC 11550] | 46.7 | 3.0E-165 |
| DIP_10004798 | NA | | |
| DIP_10004799 | putative fad binding domain-containing protein [Eutypa lata UCREL1] | 74.5 | 0.0 |
| DIP_10004800 | NA | | |
| DIP_10004801 | NA | | |
| DIP_10004802 | putative nitroreductase family protein [Eutypa lata UCREL1] | 65.8 | 2.0E-114 |
| DIP_10004803 | hypothetical protein PFICI_07843 [Pestalotiopsis fici W106-1] | 72.3 | 0.0 |
| DIP_10004804 | hypothetical protein HMPREF1624_02333 [Sporothrix schenckii ATCC 58251] | 47.5 | 4.0E-154 |
| DIP_10004805 | hypothetical protein PFICI_07850 [Pestalotiopsis fici W106-1] | 36.5 | 6.0E-26 |
| DIP_10004806 | NA | | |
| DIP_10004807 | NA | | |
| DIP_10004808 | hypothetical protein CFIO01_02477 [Colletotrichum fioriniae PJ7] | 53.3 | 2.0E-10 |
| DIP_10004809 | tRNA modification GTPase TrmE [Fusarium oxysporum f. sp. lycopersici MN25] | 64 | 0.0 |
| DIP_10004810 | putative ribosomal large subunit biogenesis protein [Eutypa lata UCREL1] | 76.2 | 2.0E-172 |
| DIP_10004811 | hypothetical protein CH063_08741 [Colletotrichum higginsianum] | 62.6 | 9.0E-65 |
| DIP_10004812 | putative gtpase-activating protein gyp7 protein [Togninia minima UCRPA7] | 62.5 | 0.0 |
| DIP_10004813 | putative mannosyl-oligosaccharide glucosidase protein [Eutypa lata UCREL1] | 67.9 | 0.0 |
| DIP_10004814 | hypothetical protein A1O7_02822 [Cladophialophora yegresii CBS 114405] | 34 | 2.0E-11 |
| DIP_10004815 | hypothetical protein PFICI_10707 [Pestalotiopsis fici W106-1] | 50.1 | 9.0E-122 |
| DIP_10004816 | plasma membrane proteolipid 3 [Colletotrichum gloeosporioides Nara gc5] | 86.3 | 2.0E-55 |
| DIP_10004817 | hypothetical protein PFICI_14145 [Pestalotiopsis fici W106-1] | 54.2 | 6.0E-174 |
| DIP_10004818 | NA | | |
| DIP_10004819 | hypothetical protein PFICI_12202 [Pestalotiopsis fici W106-1] | 58.3 | 1.0E-172 |
| DIP_10004820 | putative major myo-inositol transporter iolt protein [Eutypa lata UCREL1] | 70.8 | 0.0 |
| DIP_10004821 | NA | | |
| DIP_10004822 | NA | | |
| DIP_10004823 | putative mfs transporter protein [Eutypa lata UCREL1] | 69.2 | 0.0 |
| DIP_10004824 | putative scf e3 ubiquitin ligase complex f-box protein grra protein [Eutypa lata UCREL1] | 73 | 0.0 |
| DIP_10004825 | putative chalcone-flavanone isomerase protein [Eutypa lata UCREL1] | 56.6 | 2.0E-153 |
| DIP_10004826 | NA | | |
| DIP_10004827 | NA | | |
| DIP_10004828 | hypothetical protein PV04_07304 [Capronia semiimmersa] | 42.8 | 2.0E-27 |
| DIP_10004829 | hypothetical protein G647_04757 [Cladophialophora carrionii CBS 160.54] | 43.5 | 2.0E-86 |
| DIP_10004830 | putative glutaredoxin [Aspergillus ruber CBS 135680] | 61.5 | 1.0E-34 |
| DIP_10004831 | BAG domain-containing protein [Colletotrichum gloeosporioides Cg-14] | 51.9 | 1.0E-118 |
| DIP_10004832 | 60s ribosomal protein l16 [Colletotrichum gloeosporioides Nara gc5] | 75.3 | 9.0E-110 |
| DIP_10004833 | NA | | |
| DIP_10004834 | hob3 [Gaeumannomyces graminis var. tritici R3-111a-1] | 94.6 | 0.0 |
| DIP_10004835 | putative peptidyl-trna hydrolase protein [Togninia minima UCRPA7] | 40 | 3.0E-40 |
| DIP_10004836 | NA | | |
| DIP_10004837 | hypothetical protein PFICI_09570 [Pestalotiopsis fici W106-1] | 52.9 | 4.0E-164 |
| DIP_10004838 | putative general amino acid permease protein [Eutypa lata UCREL1] | 70.1 | 0.0 |
| DIP_10004839 | hypothetical protein Z518_04241 [Rhinocladiella mackenziei CBS 650.93] | 45 | 0.0 |
| DIP_10004840 | hypothetical protein UCREL1_8678 [Eutypa lata UCREL1] | 60.2 | 6.0E-38 |
| DIP_10004841 | putative ctr copper transporter protein [Eutypa lata UCREL1] | 59.9 | 5.0E-68 |
| DIP_10004842 | hypothetical protein MGG_13765 [Magnaporthe oryzae 70-15] | 56.4 | 0.0 |
| DIP_10004843 | nitrate assimilation regulatory protein nirA, partial [Metarhizium majus ARSEF 297] | 30.4 | 3.0E-60 |
| DIP_10004844 | hypothetical protein V495_01344 [Pseudogymnoascus pannorum VKM F-4514 (FW-929)] | 42.3 | 1.0E-68 |
| DIP_10004845 | hypothetical protein PFICI_10999 [Pestalotiopsis fici W106-1] | 50.9 | 2.0E-86 |
| DIP_10004846 | putative evolved d-pantonohydrolase protein [Togninia minima UCRPA7] | 58.9 | 7.0E-161 |
| DIP_10004847 | hypothetical protein CGLO_14119 [Colletotrichum gloeosporioides Cg-14] | 54.4 | 2.0E-176 |
| DIP_10004848 | NA | | |
| DIP_10004849 | unnamed protein product [Sordaria macrospora k-hell] | 31.9 | 6.0E-16 |
| DIP_10004850 | hypothetical protein PFICI_10902 [Pestalotiopsis fici W106-1] | 83.6 | 0.0 |
| DIP_10004851 | Cytochrome c oxidase subunit 6 [Pestalotiopsis fici W106-1] | 78 | 1.0E-77 |
| DIP_10004852 | hypothetical protein PFICI_10906 [Pestalotiopsis fici W106-1] | 32.2 | 1.0E-88 |
| DIP_10004853 | hypothetical protein PFICI_00867 [Pestalotiopsis fici W106-1] | 30.3 | 6.0E-18 |
| DIP_10004854 | putative glucose-methanol-choline oxidoreductase protein [Eutypa lata UCREL1] | 66.7 | 0.0 |
| DIP_10004855 | hypothetical protein TD95_005279 [Thielaviopsis punctulata] | 77.6 | 0.0 |
| DIP_10004856 | DNA polymerase delta catalytic subunit [Pestalotiopsis fici W106-1] | 84.8 | 0.0 |
| DIP_10004857 | hypothetical protein PFICI_07635 [Pestalotiopsis fici W106-1] | 78.7 | 0.0 |
| DIP_10004858 | putative 60s ribosomal protein l38 protein [Eutypa lata UCREL1] | 81.9 | 6.0E-41 |
| DIP_10004859 | hypothetical protein CGGC5_11922 [Colletotrichum gloeosporioides Nara gc5] | 30.6 | 8.0E-16 |
| DIP_10004860 | NA | | |
| DIP_10004861 | mevalonate kinase [Colletotrichum fioriniae PJ7] | 71.9 | 0.0 |
| DIP_10004862 | prenyltransferase alpha subunit [Colletotrichum graminicola M1.001] | 46.1 | 7.0E-159 |
| DIP_10004863 | putative nuclear transport factor 2 domain-containing protein [Eutypa lata UCREL1] | 32.6 | 2.0E-35 |
| DIP_10004864 | putative methionyl-trna synthetase protein [Eutypa lata UCREL1] | 44.4 | 1.0E-70 |
| DIP_10004865 | L-serine dehydratase [Gaeumannomyces graminis var. tritici R3-111a-1] | 48.3 | 2.0E-105 |
| DIP_10004866 | methionyl-tRNA synthetase [Magnaporthe oryzae 70-15] | 52.6 | 0.0 |
| DIP_10004867 | hypothetical protein S40285_02262 [Stachybotrys chlorohalonata IBT 40285] | 60 | 0.0 |
| DIP_10004868 | hypothetical protein PFICI_12071 [Pestalotiopsis fici W106-1] | 57.1 | 0.0 |
| DIP_10004869 | NA | | |
| DIP_10004870 | flavin-containing monooxygenase-like protein [Chaetomium thermophilum var. thermophilum DSM 1495] | 32.3 | 1.0E-42 |
| DIP_10004871 | putative lipase protein [Eutypa lata UCREL1] | 56.4 | 6.0E-146 |
| DIP_10004872 | putative glycoside hydrolase family 3 protein [Eutypa lata UCREL1] | 65 | 0.0 |
| DIP_10004873 | hypothetical protein THITE_2107125 [Thielavia terrestris NRRL 8126] | 51.2 | 2.0E-30 |
| DIP_10004874 | NA | | |
| DIP_10004875 | NA | | |
| DIP_10004876 | hypothetical protein PFICI_09630 [Pestalotiopsis fici W106-1] | 52.2 | 2.0E-107 |
| DIP_10004877 | putative centromere microtubule-binding protein cbf5 protein [Eutypa lata UCREL1] | 87.9 | 0.0 |
| DIP_10004878 | putative lpxtg-domain-containing protein [Eutypa lata UCREL1] | 44.9 | 4.0E-101 |
| DIP_10004879 | carbon-nitrogen family protein [Colletotrichum orbiculare MAFF 240422] | 54.8 | 2.0E-118 |
| DIP_10004880 | NA | | |
| DIP_10004881 | hypothetical protein PFICI_09626 [Pestalotiopsis fici W106-1] | 48.5 | 4.0E-77 |
| DIP_10004882 | NA | | |
| DIP_10004883 | hypothetical protein PFICI_09626 [Pestalotiopsis fici W106-1] | 49.1 | 1.0E-77 |
| DIP_10004884 | NA | | |
| DIP_10004885 | carbon-nitrogen family protein [Colletotrichum orbiculare MAFF 240422] | 78.8 | 9.0E-145 |
| DIP_10004886 | NA | | |
| DIP_10004887 | hypothetical protein PFICI_09625 [Pestalotiopsis fici W106-1] | 61.4 | 5.0E-90 |
| DIP_10004888 | Arginine biosynthesis bifunctional protein ArgJ [Pestalotiopsis fici W106-1] | 78.9 | 0.0 |
| DIP_10004889 | activator 1 37 kda subunit [Colletotrichum gloeosporioides Nara gc5] | 93.3 | 0.0 |
| DIP_10004890 | putative imidazoleglycerol-phosphate dehydratase protein [Eutypa lata UCREL1] | 75 | 2.0E-49 |
| DIP_10004891 | NA | | |
| DIP_10004892 | NA | | |
| DIP_10004893 | hypothetical protein THITE_2106802 [Thielavia terrestris NRRL 8126] | 52.8 | 0.0 |
| DIP_10004894 | putative transcription initiation factor iie subunit alpha protein [Eutypa lata UCREL1] | 65.8 | 3.0E-151 |
| DIP_10004895 | NA | | |
| DIP_10004896 | putative smc1 protein [Eutypa lata UCREL1] | 79.2 | 0.0 |
| DIP_10004897 | putative f-box protein [Eutypa lata UCREL1] | 69.4 | 0.0 |
| DIP_10004898 | hypothetical protein PFICI_08351 [Pestalotiopsis fici W106-1] | 89.8 | 0.0 |
| DIP_10004899 | NA | | |
| DIP_10004900 | NA | | |
| DIP_10004901 | hypothetical protein PFICI_11037 [Pestalotiopsis fici W106-1] | 56.4 | 0.0 |
| DIP_10004902 | hypothetical protein UCREL1_1940 [Eutypa lata UCREL1] | 54.1 | 3.0E-112 |
| DIP_10004903 | putative n-acetyltransferase 5 protein [Eutypa lata UCREL1] | 74.7 | 4.0E-109 |
| DIP_10004904 | hypothetical protein CGLO_00847 [Colletotrichum gloeosporioides Cg-14] | 66.4 | 0.0 |
| DIP_10004905 | hypothetical protein V500_09345 [Pseudogymnoascus pannorum VKM F-4518 (FW-2643)] | 45.1 | 2.0E-43 |
| DIP_10004906 | NA | | |
| DIP_10004907 | NA | | |
| DIP_10004908 | hypothetical protein PFICI_07379 [Pestalotiopsis fici W106-1] | 51.5 | 3.0E-133 |
| DIP_10004909 | ubiquitin family protein [Colletotrichum graminicola M1.001] | 38.3 | 3.0E-104 |
| DIP_10004910 | hypothetical protein PFICI_07829 [Pestalotiopsis fici W106-1] | 63.4 | 0.0 |
| DIP_10004911 | putative importin beta-2 protein [Eutypa lata UCREL1] | 74.7 | 0.0 |
| DIP_10004912 | hypothetical protein CTHT_0061320 [Chaetomium thermophilum var. thermophilum DSM 1495] | 61.7 | 0.0 |
| DIP_10004913 | conserved hypothetical protein [Botrytis cinerea B05.10] | 81.7 | 0.0 |
| DIP_10004914 | hypothetical protein PFICI_07824 [Pestalotiopsis fici W106-1] | 45 | 6.0E-170 |
| DIP_10004915 | hypothetical protein PFICI_07776 [Pestalotiopsis fici W106-1] | 76.5 | 3.0E-161 |
| DIP_10004916 | kinase domain protein, partial [Actinomyces sp. oral taxon 171 str. F0337] | 32.7 | 4.0E-06 |
| DIP_10004917 | hypothetical protein PFICI_07218 [Pestalotiopsis fici W106-1] | 45.8 | 1.0E-153 |
| DIP_10004918 | hypothetical protein CSUB01_05497 [Colletotrichum sublineola] | 30.8 | 5.0E-50 |
| DIP_10004919 | putative ubiquitination network signaling protein [Eutypa lata UCREL1] | 70.1 | 0.0 |
| DIP_10004920 | hypothetical protein FOTG_04978 [Fusarium oxysporum f. sp. vasinfectum 25433] | 31.4 | 5.0E-57 |
| DIP_10004921 | hypothetical protein PV08_02243 [Exophiala spinifera] | 43.2 | 6.0E-93 |
| DIP_10004922 | NA | | |
| DIP_10004923 | putative thioesterase thiol ester dehydrase-isomerase protein [Eutypa lata UCREL1] | 54.4 | 8.0E-101 |
| DIP_10004924 | LYR family protein [Magnaporthe oryzae 70-15] | 72.1 | 5.0E-30 |
| DIP_10004925 | putative sam binding domain-containing protein containing protein [Eutypa lata UCREL1] | 67 | 0.0 |
| DIP_10004926 | NA | | |
| DIP_10004927 | hypothetical protein PFICI_07640 [Pestalotiopsis fici W106-1] | 72.2 | 0.0 |
| DIP_10004928 | hypothetical protein PFICI_07641 [Pestalotiopsis fici W106-1] | 53.4 | 1.0E-118 |
| DIP_10004929 | hypothetical protein MYCFIDRAFT_128036, partial [Pseudocercospora fijiensis CIRAD86] | 35.7 | 5.0E-82 |
| DIP_10004930 | NA | | |
| DIP_10004931 | hypothetical protein PFICI_07644 [Pestalotiopsis fici W106-1] | 52 | 2.0E-112 |
| DIP_10004932 | hypothetical protein PFICI_07427 [Pestalotiopsis fici W106-1] | 65.3 | 0.0 |
| DIP_10004933 | AdoMet-dependent rRNA methyltransferase SPB1 [Pestalotiopsis fici W106-1] | 72.8 | 0.0 |
| DIP_10004934 | NA | | |
| DIP_10004935 | NA | | |
| DIP_10004936 | hypothetical protein PFICI_07646 [Pestalotiopsis fici W106-1] | 61.7 | 0.0 |
| DIP_10004937 | hypothetical protein [Podospora anserina S mat+] | 45.1 | 2.0E-69 |
| DIP_10004938 | NA | | |
| DIP_10004939 | putative signal sequence receptor alpha chain protein [Eutypa lata UCREL1] | 62.5 | 2.0E-111 |
| DIP_10004940 | hypothetical protein PFICI_07676 [Pestalotiopsis fici W106-1] | 57.1 | 4.0E-171 |
| DIP_10004941 | putative x-pro dipeptidyl-peptidase protein [Diaporthe ampelina] | 71 | 0.0 |
| DIP_10004942 | hypothetical protein V495_05068 [Pseudogymnoascus pannorum VKM F-4514 (FW-929)] | 35.1 | 1.0E-25 |
| DIP_10004943 | putative aldehyde dehydrogenase [Diaporthe ampelina] | 54.2 | 0.0 |
| DIP_10004944 | hypothetical protein PFICI_07680 [Pestalotiopsis fici W106-1] | 55.6 | 3.0E-141 |
| DIP_10004945 | hypothetical protein PFICI_07650 [Pestalotiopsis fici W106-1] | 56.1 | 0.0 |
| DIP_10004946 | putative ilp is an apoptosis inhibitor protein [Eutypa lata UCREL1] | 68.7 | 2.0E-48 |
| DIP_10004947 | NA | | |
| DIP_10004948 | predicted protein [Nectria haematococca mpVI 77-13-4] | 76.3 | 0.0 |
| DIP_10004949 | NA | | |
| DIP_10004950 | hypothetical protein PFICI_07636 [Pestalotiopsis fici W106-1] | 73 | 0.0 |
| DIP_10004951 | hypothetical protein PFICI_07637 [Pestalotiopsis fici W106-1] | 55.1 | 2.0E-99 |
| DIP_10004952 | hypothetical protein PFICI_07638 [Pestalotiopsis fici W106-1] | 54.5 | 1.0E-15 |
| DIP_10004953 | Importin subunit alpha-1 [Pestalotiopsis fici W106-1] | 94 | 0.0 |
| DIP_10004954 | putative telomere-binding alpha subunit central domain-containing protein [Eutypa lata UCREL1] | 43.3 | 9.0E-169 |
| DIP_10004955 | putative stag domain-containing protein [Eutypa lata UCREL1] | 62.9 | 0.0 |
| DIP_10004956 | putative dna-directed rna polymerase iii protein [Eutypa lata UCREL1] | 56.7 | 2.0E-153 |
| DIP_10004957 | NA | | |
| DIP_10004958 | putative major facilitator superfamily protein [Eutypa lata UCREL1] | 65.8 | 0.0 |
| DIP_10004959 | hypothetical protein PFICI_07655 [Pestalotiopsis fici W106-1] | 59.9 | 0.0 |
| DIP_10004960 | putative g2 mitotic-specific cyclin-b protein [Eutypa lata UCREL1] | 59 | 0.0 |
| DIP_10004961 | putative calcium-binding mitochondrial carrier protein aralar1 protein [Eutypa lata UCREL1] | 76.1 | 0.0 |
| DIP_10004962 | hypothetical protein PFICI_07660 [Pestalotiopsis fici W106-1] | 61.3 | 2.0E-114 |
| DIP_10004963 | putative glucosamine 6-phosphate synthetase protein [Eutypa lata UCREL1] | 74.4 | 0.0 |
| DIP_10004964 | NA | | |
| DIP_10004965 | hypothetical protein MPH_05938 [Macrophomina phaseolina MS6] | 57.7 | 1.0E-94 |
| DIP_10004966 | NA | | |
| DIP_10004967 | putative xpg domain-containing protein [Eutypa lata UCREL1] | 69.3 | 0.0 |
| DIP_10004968 | putative 2og-fe oxygenase protein [Eutypa lata UCREL1] | 66 | 5.0E-87 |
| DIP_10004969 | hypothetical protein PFICI_11017 [Pestalotiopsis fici W106-1] | 63.5 | 0.0 |
| DIP_10004970 | WD domain-containing protein [Colletotrichum fioriniae PJ7] | 47.9 | 3.0E-158 |
| DIP_10004971 | putative vacuolar sorting protein 1 protein [Eutypa lata UCREL1] | 87.8 | 0.0 |
| DIP_10004972 | hypothetical protein UCREL1_9422 [Eutypa lata UCREL1] | 42.7 | 3.0E-63 |
| DIP_10004973 | putative type ii inositol- -trisphosphate 5-phosphatase protein [Eutypa lata UCREL1] | 58.8 | 0.0 |
| DIP_10004974 | putative ammonium transporter mep2 protein [Eutypa lata UCREL1] | 80.6 | 0.0 |
| DIP_10004975 | hypothetical protein PFICI_11027 [Pestalotiopsis fici W106-1] | 62.1 | 0.0 |
| DIP_10004976 | putative hypercellular protein a [Diaporthe ampelina] | 62.1 | 3.0E-71 |
| DIP_10004977 | putative phosphoserine aminotransferase protein [Togninia minima UCRPA7] | 76.2 | 0.0 |
| DIP_10004978 | hypothetical protein PFICI_11026 [Pestalotiopsis fici W106-1] | 77.3 | 0.0 |
| DIP_10004979 | putative uncharacterized conserved protein ucp014753 protein [Togninia minima UCRPA7] | 51 | 9.0E-42 |
| DIP_10004980 | hypothetical protein PFICI_11024 [Pestalotiopsis fici W106-1] | 76.6 | 0.0 |
| DIP_10004981 | hypothetical protein Cob_03205 [Colletotrichum orbiculare MAFF 240422] | 64.1 | 2.0E-69 |
| DIP_10004982 | Nitrogen metabolic regulation protein 1 [Pestalotiopsis fici W106-1] | 75.9 | 0.0 |
| DIP_10004983 | hypothetical protein PFICI_11013 [Pestalotiopsis fici W106-1] | 42.5 | 0.0 |
| DIP_10004984 | hypothetical protein PFICI_11012 [Pestalotiopsis fici W106-1] | 63.1 | 0.0 |
| DIP_10004985 | putative rhomboid family membrane protein [Eutypa lata UCREL1] | 43 | 3.0E-45 |
| DIP_10004986 | NA | | |
| DIP_10004987 | putative ribosome biogenesis protein ytm1 protein [Eutypa lata UCREL1] | 68.3 | 0.0 |
| DIP_10004988 | dna polymerase delta subunit 2 [Fusarium avenaceum] | 63.2 | 0.0 |
| DIP_10004989 | NA | | |
| DIP_10004990 | BTB/POZ domain protein [Beauveria bassiana ARSEF 2860] | 38.1 | 1.0E-18 |
| DIP_10004991 | uncharacterized protein CPUR_05456 [Claviceps purpurea 20.1] | 34.5 | 5.0E-11 |
| DIP_10004992 | NA | | |
| DIP_10004993 | putative solute carrier family 12 member 3 protein [Eutypa lata UCREL1] | 57 | 3.0E-57 |
| DIP_10004994 | amino acid transporter [Aspergillus oryzae 3.042] | 42.5 | 6.0E-78 |
| DIP_10004995 | hypothetical protein PFICI_09439 [Pestalotiopsis fici W106-1] | 66.4 | 0.0 |
| DIP_10004996 | putative dna-directed rna polymerase ii subunit rpb11 protein [Eutypa lata UCREL1] | 79.4 | 6.0E-73 |
| DIP_10004997 | U6 snRNA-associated Sm-like protein LSm5 [Trichoderma harzianum] | 100 | 4.0E-55 |
| DIP_10004998 | hypothetical protein PFICI_09436 [Pestalotiopsis fici W106-1] | 43.1 | 1.0E-120 |
| DIP_10004999 | hypothetical protein PFICI_08052 [Pestalotiopsis fici W106-1] | 73.8 | 0.0 |
| DIP_10005000 | hypothetical protein PFICI_08055 [Pestalotiopsis fici W106-1] | 60.3 | 0.0 |
| DIP_10005001 | putative septum-promoting gtp-binding protein 1 protein [Eutypa lata UCREL1] | 79.6 | 2.0E-176 |
| DIP_10005002 | hypothetical protein PFICI_04195 [Pestalotiopsis fici W106-1] | 76 | 5.0E-95 |
| DIP_10005003 | putative porphobilinogen deaminase protein [Eutypa lata UCREL1] | 65 | 3.0E-158 |
| DIP_10005004 | hypothetical protein CSUB01_05636 [Colletotrichum sublineola] | 35.1 | 4.0E-11 |
| DIP_10005005 | hypothetical protein S40285_08958 [Stachybotrys chlorohalonata IBT 40285] | 43.9 | 4.0E-87 |
| DIP_10005006 | Beta-glucosidase B [Pestalotiopsis fici W106-1] | 65.7 | 0.0 |
| DIP_10005007 | putative glycosyl family protein [Eutypa lata UCREL1] | 43.2 | 4.0E-110 |
| DIP_10005008 | 60S ribosomal protein L17 [Sclerotinia sclerotiorum 1980 UF-70] | 87.7 | 1.0E-119 |
| DIP_10005009 | putative rrna 2 -o-methyltransferase fibrillarin protein [Eutypa lata UCREL1] | 75.8 | 2.0E-130 |
| DIP_10005010 | NA | | |
| DIP_10005011 | cyclic AMP receptor-like protein A [Pestalotiopsis microspora] | 48.3 | 2.0E-141 |
| DIP_10005012 | hypothetical protein CGLO_16779 [Colletotrichum gloeosporioides Cg-14] | 44.7 | 5.0E-12 |
| DIP_10005013 | putative inner membrane protein yieg protein [Eutypa lata UCREL1] | 76.7 | 0.0 |
| DIP_10005014 | hypothetical protein PFICI_09448 [Pestalotiopsis fici W106-1] | 67.3 | 0.0 |
| DIP_10005015 | hypothetical protein PFICI_09447 [Pestalotiopsis fici W106-1] | 69.8 | 8.0E-138 |
| DIP_10005016 | NA | | |
| DIP_10005017 | putative serine threonine-protein phosphatase 5 protein [Eutypa lata UCREL1] | 88.3 | 0.0 |
| DIP_10005018 | predicted protein [Trichoderma reesei QM6a] | 38.9 | 8.0E-28 |
| DIP_10005019 | putative transcription elongation factor spt-4 protein [Eutypa lata UCREL1] | 64 | 9.0E-47 |
| DIP_10005020 | NA | | |
| DIP_10005021 | putative malate dehydrogenase protein [Eutypa lata UCREL1] | 56.1 | 4.0E-106 |
| DIP_10005022 | NA | | |
| DIP_10005023 | hypothetical protein GLRG_04894 [Colletotrichum graminicola M1.001] | 51.5 | 4.0E-147 |
| DIP_10005024 | Periodic tryptophan protein 2 [Pestalotiopsis fici W106-1] | 81.5 | 0.0 |
| DIP_10005025 | putative cell division control protein 12 protein [Eutypa lata UCREL1] | 95.5 | 0.0 |
| DIP_10005026 | hypothetical protein PFICI_08313 [Pestalotiopsis fici W106-1] | 76.9 | 3.0E-146 |
| DIP_10005027 | putative ubiquitin carboxyl-terminal hydrolase protein [Eutypa lata UCREL1] | 74.4 | 0.0 |
| DIP_10005028 | NA | | |
| DIP_10005029 | hypothetical protein CHGG_07250 [Chaetomium globosum CBS 148.51] | 70.6 | 9.0E-81 |
| DIP_10005030 | unnamed protein product [Fusarium sp. FIESC_5 CS3069] | 63.8 | 4.0E-168 |
| DIP_10005031 | hypothetical protein PFICI_00199 [Pestalotiopsis fici W106-1] | 74.2 | 0.0 |
| DIP_10005032 | pre-mRNA-processing protein [Verticillium alfalfae VaMs.102] | 74 | 0.0 |
| DIP_10005033 | hypothetical protein PFICI_08318 [Pestalotiopsis fici W106-1] | 75.6 | 2.0E-117 |
| DIP_10005034 | putative alkaline phosphatase family protein [Eutypa lata UCREL1] | 65.2 | 0.0 |
| DIP_10005035 | putative serine threonine-protein phosphatase 2b catalytic subunit protein [Eutypa lata UCREL1] | 87.7 | 0.0 |
| DIP_10005036 | hypothetical protein PFICI_08325 [Pestalotiopsis fici W106-1] | 58.1 | 2.0E-132 |
| DIP_10005037 | hypothetical protein GGTG_12219 [Gaeumannomyces graminis var. tritici R3-111a-1] | 53.9 | 2.0E-99 |
| DIP_10005038 | hypothetical protein PFICI_08327 [Pestalotiopsis fici W106-1] | 72.3 | 2.0E-69 |
| DIP_10005039 | putative deah-box rna helicase protein [Eutypa lata UCREL1] | 77.2 | 0.0 |
| DIP_10005040 | cutinase precursor [Colletotrichum orbiculare MAFF 240422] | 32.5 | 3.0E-17 |
| DIP_10005041 | hypothetical protein GALMADRAFT_146525 [Galerina marginata CBS 339.88] | 40.5 | 2.0E-63 |
| DIP_10005042 | NA | | |
| DIP_10005043 | hypothetical protein PFICI_08329 [Pestalotiopsis fici W106-1] | 55 | 7.0E-113 |
| DIP_10005044 | putative hatpase c domain-containing protein [Eutypa lata UCREL1] | 68.6 | 0.0 |
| DIP_10005045 | putative histone acetylase complex subunit protein [Eutypa lata UCREL1] | 79.2 | 0.0 |
| DIP_10005046 | hypothetical protein SS1G_00565 [Sclerotinia sclerotiorum 1980] | 63.7 | 4.0E-149 |
| DIP_10005047 | putative asparagine-rich protein [Eutypa lata UCREL1] | 87.7 | 0.0 |
| DIP_10005048 | putative 5 -3 exoribonuclease 1 protein [Eutypa lata UCREL1] | 75.6 | 0.0 |
| DIP_10005049 | hypothetical protein V502_09118 [Pseudogymnoascus pannorum VKM F-4520 (FW-2644)] | 57.4 | 0.0 |
| DIP_10005050 | Histone acetyltransferase type B subunit 2 [Pestalotiopsis fici W106-1] | 72.1 | 0.0 |
| DIP_10005051 | putative aspartate kinase protein [Eutypa lata UCREL1] | 77.6 | 0.0 |
| DIP_10005052 | hypothetical protein TRIVIDRAFT_34449, partial [Trichoderma virens Gv29-8] | 53.5 | 0.0 |
| DIP_10005053 | putative protein yop-1 protein [Eutypa lata UCREL1] | 87 | 4.0E-106 |
| DIP_10005054 | putative translation initiation factor eif4e3 protein [Eutypa lata UCREL1] | 62.1 | 8.0E-142 |
| DIP_10005055 | putative chromatin assembly factor 1 subunit a protein [Eutypa lata UCREL1] | 51.1 | 7.0E-89 |
| DIP_10005056 | hypothetical protein S7711_06615 [Stachybotrys chartarum IBT 7711] | 49.5 | 2.0E-125 |
| DIP_10005057 | Carboxylesterase family protein [Aspergillus parasiticus SU-1] | 64.9 | 0.0 |
| DIP_10005058 | NA | | |
| DIP_10005059 | NA | | |
| DIP_10005060 | NAD(P)-binding Rossmann-fold containing protein [Glarea lozoyensis ATCC 20868] | 48.7 | 3.0E-102 |
| DIP_10005061 | hypothetical protein A1O1_03543 [Capronia coronata CBS 617.96] | 31.7 | 2.0E-98 |
| DIP_10005062 | hypothetical protein PV06_05046 [Exophiala oligosperma] | 42.8 | 3.0E-109 |
| DIP_10005063 | hypothetical protein FPSE_03235 [Fusarium pseudograminearum CS3096] | 51.2 | 1.0E-179 |
| DIP_10005064 | hypothetical protein FVEG_09999 [Fusarium verticillioides 7600] | 32.1 | 9.0E-60 |
| DIP_10005065 | similar to cytochrome P450 [Botrytis cinerea T4] | 38.4 | 6.0E-132 |
| DIP_10005066 | NA | | |
| DIP_10005067 | hypothetical protein TRIATDRAFT_14480, partial [Trichoderma atroviride IMI 206040] | 32 | 2.0E-14 |
| DIP_10005068 | hypothetical protein PFICI_09596 [Pestalotiopsis fici W106-1] | 62.6 | 0.0 |
| DIP_10005069 | hypothetical protein PFICI_09595 [Pestalotiopsis fici W106-1] | 43 | 6.0E-146 |
| DIP_10005070 | NA | | |
| DIP_10005071 | NA | | |
| DIP_10005072 | hypothetical protein CGLO_03301 [Colletotrichum gloeosporioides Cg-14] | 55 | 8.0E-114 |
| DIP_10005073 | hypothetical protein THAR02_02193 [Trichoderma harzianum] | 50.4 | 2.0E-34 |
| DIP_10005074 | putative transaldolase protein [Togninia minima UCRPA7] | 77.4 | 0.0 |
| DIP_10005075 | sugar transporter [Colletotrichum gloeosporioides Nara gc5] | 73.8 | 0.0 |
| DIP_10005076 | NA | | |
| DIP_10005077 | hypothetical protein THITE_2107729 [Thielavia terrestris NRRL 8126] | 31.2 | 3.0E-26 |
| DIP_10005078 | NA | | |
| DIP_10005079 | NA | | |
| DIP_10005080 | hypothetical protein PFICI_11546 [Pestalotiopsis fici W106-1] | 59.3 | 4.0E-119 |
| DIP_10005081 | hypothetical protein FVEG_04537 [Fusarium verticillioides 7600] | 46.9 | 0.0 |
| DIP_10005082 | putative zinc metalloprotease protein [Eutypa lata UCREL1] | 75.7 | 0.0 |
| DIP_10005083 | Cutinase-like protein [Acremonium chrysogenum ATCC 11550] | 45.2 | 2.0E-54 |
| DIP_10005084 | hypothetical protein HIM_02808 [Hirsutella minnesotensis 3608] | 60.9 | 1.0E-135 |
| DIP_10005085 | NA | | |
| DIP_10005086 | NA | | |
| DIP_10005087 | hypothetical protein CGLO_04905 [Colletotrichum gloeosporioides Cg-14] | 34.8 | 1.0E-38 |
| DIP_10005088 | NA | | |
| DIP_10005089 | putative ornithine decarboxylase protein [Eutypa lata UCREL1] | 88.1 | 0.0 |
| DIP_10005090 | putative mitochondrial presequence protease protein [Eutypa lata UCREL1] | 81.4 | 0.0 |
| DIP_10005091 | putative muscle-derived protein (neurite-outgrowth-promoting) protein [Eutypa lata UCREL1] | 71.2 | 0.0 |
| DIP_10005092 | putative low temperature viability protein [Colletotrichum sublineola] | 54.6 | 1.0E-155 |
| DIP_10005093 | putative nucleolar gtp-binding protein [Eutypa lata UCREL1] | 74.7 | 0.0 |
| DIP_10005094 | hypothetical protein PFICI_05776 [Pestalotiopsis fici W106-1] | 47.3 | 2.0E-35 |
| DIP_10005095 | NA | | |
| DIP_10005096 | hypothetical protein [Podospora anserina S mat+] | 34.5 | 2.0E-16 |
| DIP_10005097 | putative periplasmic binding ii protein [Eutypa lata UCREL1] | 78.9 | 2.0E-176 |
| DIP_10005098 | hypothetical protein FOVG_07333 [Fusarium oxysporum f. sp. pisi HDV247] | 36.5 | 4.0E-26 |
| DIP_10005099 | NA | | |
| DIP_10005100 | putative glycoside hydrolase family 31 protein [Eutypa lata UCREL1] | 74.8 | 0.0 |
| DIP_10005101 | nucleoside diphosphate kinase [Magnaporthe oryzae 70-15] | 84 | 2.0E-100 |
| DIP_10005102 | hypothetical protein UCREL1_7895 [Eutypa lata UCREL1] | 33.8 | 3.0E-74 |
| DIP_10005103 | triacylglycerol lipase-like protein [Chaetomium thermophilum var. thermophilum DSM 1495] | 33.9 | 1.0E-19 |
| DIP_10005104 | hypothetical protein UCREL1_10262 [Eutypa lata UCREL1] | 69.5 | 2.0E-114 |
| DIP_10005105 | HAD-superfamily hydrolase subfamily IIA CECR5 [Macrophomina phaseolina MS6] | 82.2 | 0.0 |
| DIP_10005106 | putative glycerol dehydrogenase protein [Neofusicoccum parvum UCRNP2] | 74.5 | 5.0E-180 |
| DIP_10005107 | hypothetical protein PFICI_09593 [Pestalotiopsis fici W106-1] | 65.3 | 0.0 |
| DIP_10005108 | cysteinyl-tRNA synthetase [Metarhizium acridum CQMa 102] | 68.8 | 0.0 |
| DIP_10005109 | hypothetical protein FOYG_17030 [Fusarium oxysporum FOSC 3-a] | 38.8 | 9.0E-44 |
| DIP_10005110 | hypothetical protein THAR02_04740 [Trichoderma harzianum] | 61.2 | 5.0E-166 |
| DIP_10005111 | putative glycoside hydrolase family 18 protein [Eutypa lata UCREL1] | 47.1 | 2.0E-99 |
| DIP_10005112 | hypothetical protein PFICI_11078 [Pestalotiopsis fici W106-1] | 47.3 | 1.0E-55 |
| DIP_10005113 | putative benzoylformate decarboxylase protein [Eutypa lata UCREL1] | 68.1 | 3.0E-149 |
| DIP_10005114 | Tryptophan--tRNA ligase [Pestalotiopsis fici W106-1] | 79.7 | 0.0 |
| DIP_10005115 | putative small nuclear ribonucleoprotein sm d3 protein [Eutypa lata UCREL1] | 100 | 2.0E-79 |
| DIP_10005116 | hypothetical protein PV09_07058 [Verruconis gallopava] | 71 | 9.0E-46 |
| DIP_10005117 | NA | | |
| DIP_10005118 | putative wsc domain-containing protein [Eutypa lata UCREL1] | 55.4 | 0.0 |
| DIP_10005119 | hypothetical protein PFICI_11064 [Pestalotiopsis fici W106-1] | 53 | 5.0E-125 |
| DIP_10005120 | putative sugar transporter stl1 protein [Eutypa lata UCREL1] | 75.7 | 0.0 |
| DIP_10005121 | hypothetical protein PFICI_03962 [Pestalotiopsis fici W106-1] | 47.9 | 0.0 |

| | | | |
|---|---|---|---|
| DIP_10005122 | hypothetical protein MGG_02347 [Magnaporthe oryzae 70-15] | 35.1 | 9.0E-20 |
| DIP_10005123 | putative ubiquitin-conjugating enzyme protein [Eutypa lata UCREL1] | 89.2 | 8.0E-105 |
| DIP_10005124 | hypothetical protein PFICI_11076 [Pestalotiopsis fici W106-1] | 46.7 | 0.0 |
| DIP_10005125 | protease inhibitor [Neurospora crassa OR74A] | 36.2 | 3.0E-26 |
| DIP_10005126 | hypothetical protein NECHADRAFT_102200 [Nectria haematococca mpVI 77-13-4] | 33.3 | 5.0E-55 |
| DIP_10005127 | putative microsomal glutathione s-transferase protein [Eutypa lata UCREL1] | 63.5 | 8.0E-62 |
| DIP_10005128 | putative zinc carboxypeptidase protein [Eutypa lata UCREL1] | 65 | 0.0 |
| DIP_10005129 | hypothetical protein SAPIO_CDS0575 [Scedosporium apiospermum] | 50.9 | 0.0 |
| DIP_10005130 | putative signal recognition particle protein [Eutypa lata UCREL1] | 59.9 | 0.0 |
| DIP_10005131 | NA | | |
| DIP_10005132 | related to transcriptional activator CMR1 [Fusarium fujikuroi IMI 58289] | 30.2 | 1.0E-90 |
| DIP_10005133 | hypothetical protein VHEMI04314 [Torrubiella hemipterigena] | 57.4 | 4.0E-118 |
| DIP_10005134 | NA | | |
| DIP_10005135 | hypothetical protein PFICI_08353 [Pestalotiopsis fici W106-1] | 37.2 | 3.0E-93 |
| DIP_10005136 | hypothetical protein PFICI_08352 [Pestalotiopsis fici W106-1] | 46.5 | 2.0E-85 |
| DIP_10005137 | NA | | |
| DIP_10005138 | NA | | |
| DIP_10005139 | NA | | |
| DIP_10005140 | hypothetical protein HIM_04809 [Hirsutella minnesotensis 3608] | 50.8 | 8.0E-57 |
| DIP_10005141 | NA | | |
| DIP_10005142 | hypothetical protein PFICI_08363 [Pestalotiopsis fici W106-1] | 72.8 | 0.0 |
| DIP_10005143 | Ribosomal protein L11 [Macrophomina phaseolina MS6] | 89.7 | 7.0E-103 |
| DIP_10005144 | putative transport protein sec23 protein [Eutypa lata UCREL1] | 91.5 | 0.0 |
| DIP_10005145 | hypothetical protein PFICI_10996 [Pestalotiopsis fici W106-1] | 44.6 | 2.0E-76 |
| DIP_10005146 | NA | | |
| DIP_10005147 | putative glucose-repressible alcohol dehydrogenase transcriptional effector protein [Eutypa lata UCREL1] | 79.5 | 0.0 |
| DIP_10005148 | NA | | |
| DIP_10005149 | putative guanosine-diphosphatase protein [Eutypa lata UCREL1] | 79.2 | 0.0 |
| DIP_10005150 | hypothetical protein PFICI_07422 [Pestalotiopsis fici W106-1] | 60.3 | 0.0 |
| DIP_10005151 | hypothetical protein UCDDA912_g03530 [Diaporthe ampelina] | 41.6 | 6.0E-104 |
| DIP_10005152 | NA | | |
| DIP_10005153 | putative metallo-hydrolase oxidoreductase protein [Eutypa lata UCREL1] | 58.4 | 7.0E-75 |
| DIP_10005154 | putative ras-like protein rab7 protein [Eutypa lata UCREL1] | 93.1 | 2.0E-143 |
| DIP_10005155 | putative thiamine-repressible mitochondrial transport protein thi74 protein [Eutypa lata UCREL1] | 57.8 | 0.0 |
| DIP_10005156 | NA | | |
| DIP_10005157 | putative heme oxygenase-like protein [Eutypa lata UCREL1] | 38.5 | 3.0E-66 |
| DIP_10005158 | hypothetical protein PFICI_07430 [Pestalotiopsis fici W106-1] | 68.3 | 3.0E-142 |
| DIP_10005159 | hypothetical protein PFICI_03601 [Pestalotiopsis fici W106-1] | 57.8 | 5.0E-180 |
| DIP_10005160 | putative protein transport protein sec61 alpha subunit protein [Eutypa lata UCREL1] | 96.4 | 0.0 |
| DIP_10005161 | NA | | |
| DIP_10005162 | putative atp-dependent bile acid permease protein [Eutypa lata UCREL1] | 73.5 | 0.0 |
| DIP_10005163 | hypothetical protein PFICI_09524 [Pestalotiopsis fici W106-1] | 31.5 | 1.0E-109 |
| DIP_10005164 | putative mitochondrial inner membrane aaa protease protein [Eutypa lata UCREL1] | 66.3 | 0.0 |
| DIP_10005165 | hypothetical protein PFICI_11031 [Pestalotiopsis fici W106-1] | 54 | 3.0E-113 |
| DIP_10005166 | hypothetical protein CFIO01_06797 [Colletotrichum fioriniae PJ7] | 37.6 | 7.0E-39 |
| DIP_10005167 | hypothetical protein TRIVIDRAFT_19037, partial [Trichoderma virens Gv29-8] | 43.4 | 1.0E-30 |
| DIP_10005168 | putative protein transport protein sec22 protein [Eutypa lata UCREL1] | 82.3 | 3.0E-134 |
| DIP_10005169 | hypothetical protein UCREL1_402 [Eutypa lata UCREL1] | 52.5 | 2.0E-136 |
| DIP_10005170 | putative cleft lip and palate transmembrane protein 1 protein [Eutypa lata UCREL1] | 69 | 0.0 |
| DIP_10005171 | NA | | |
| DIP_10005172 | NA | | |
| DIP_10005173 | NA | | |
| DIP_10005174 | hypothetical protein PFICI_07409 [Pestalotiopsis fici W106-1] | 77.1 | 0.0 |
| DIP_10005175 | initiator tRNA phosphoribosyl transferase [Magnaporthe oryzae 70-15] | 45.1 | 8.0E-154 |
| DIP_10005176 | NA | | |
| DIP_10005177 | putative glycosyl hydrolase family 43 protein [Eutypa lata UCREL1] | 41.6 | 3.0E-83 |
| DIP_10005178 | dnaJ heat shock family protein [Colletotrichum gloeosporioides Nara gc5] | 72.2 | 3.0E-179 |
| DIP_10005179 | NA | | |
| DIP_10005180 | ubiquitin interaction domain-containing protein [Colletotrichum gloeosporioides Cg-14] | 36.7 | 8.0E-94 |
| DIP_10005181 | putative ubiquitin-conjugating enzyme protein [Eutypa lata UCREL1] | 75.7 | 8.0E-109 |
| DIP_10005182 | hypothetical protein PFICI_08197 [Pestalotiopsis fici W106-1] | 53.8 | 9.0E-105 |
| DIP_10005183 | hypothetical protein UCDDA912_g03717 [Diaporthe ampelina] | 45.3 | 1.0E-152 |
| DIP_10005184 | fungal chitosanase [Colletotrichum fioriniae PJ7] | 48.5 | 7.0E-79 |
| DIP_10005185 | NmrA-like protein [Macrophomina phaseolina MS6] | 37.6 | 1.0E-65 |
| DIP_10005186 | alcohol dehydrogenase [Pseudozyma brasiliensis GHG001] | 67.2 | 5.0E-177 |
| DIP_10005187 | putative apc13 domain-containing protein [Eutypa lata UCREL1] | 66.3 | 2.0E-39 |
| DIP_10005188 | putative trafficking protein particle complex subunit 5 protein [Eutypa lata UCREL1] | 83.1 | 8.0E-137 |
| DIP_10005189 | putative cral trio domain-containing protein [Eutypa lata UCREL1] | 64.4 | 9.0E-155 |
| DIP_10005190 | DNA polymerase alpha/epsilon subunit B [Colletotrichum fioriniae PJ7] | 52.2 | 0.0 |
| DIP_10005191 | hypothetical protein PFICI_07817 [Pestalotiopsis fici W106-1] | 52 | 0.0 |
| DIP_10005192 | hypothetical protein UCREL1_5444 [Eutypa lata UCREL1] | 52.9 | 4.0E-48 |
| DIP_10005193 | related to microfibril-associated protein [Fusarium fujikuroi IMI 58289] | 54.9 | 2.0E-105 |
| DIP_10005194 | duf250 domain membrane protein [Colletotrichum gloeosporioides Nara gc5] | 44.7 | 3.0E-101 |
| DIP_10005195 | Protein ssh-4 [Pestalotiopsis fici W106-1] | 84.1 | 0.0 |
| DIP_10005196 | putative ribosomal protein 35 protein [Eutypa lata UCREL1] | 68.1 | 0.0 |
| DIP_10005197 | hypothetical protein PFICI_07438 [Pestalotiopsis fici W106-1] | 61.1 | 1.0E-121 |
| DIP_10005198 | hypothetical protein PFICI_09460 [Pestalotiopsis fici W106-1] | 80.7 | 0.0 |
| DIP_10005199 | putative GLE1-like protein [Colletotrichum sublineola] | 50.1 | 3.0E-160 |
| DIP_10005200 | hypothetical protein CGLO_08224 [Colletotrichum gloeosporioides Cg-14] | 62.1 | 2.0E-96 |
| DIP_10005201 | hypothetical protein PFICI_09463 [Pestalotiopsis fici W106-1] | 56.3 | 0.0 |
| DIP_10005202 | hypothetical protein DICSQDRAFT_158575 [Dichomitus squalens LYAD-421 SS1] | 35.7 | 8.0E-09 |
| DIP_10005203 | unnamed protein product [Sordaria macrospora k-hell] | 77.6 | 0.0 |
| DIP_10005204 | hypothetical protein UCREL1_9417 [Eutypa lata UCREL1] | 50 | 3.0E-122 |
| DIP_10005205 | putative atp-dependent dna ligase domain-containing protein [Eutypa lata UCREL1] | 39 | 3.0E-87 |
| DIP_10005206 | hypothetical protein PFICI_09517 [Pestalotiopsis fici W106-1] | 78.9 | 1.0E-160 |
| DIP_10005207 | putative topoisomerase 1-associated factor 1 protein [Eutypa lata UCREL1] | 73.2 | 0.0 |
| DIP_10005208 | formate dehydrogenase [Magnaporthe oryzae Y34] | 78.9 | 0.0 |
| DIP_10005209 | putative fad binding protein [Eutypa lata UCREL1] | 50.8 | 3.0E-143 |
| DIP_10005210 | hypothetical protein BAUCODRAFT_145544 [Baudoinia compniacensis UAMH 10762] | 54.7 | 0.0 |
| DIP_10005211 | putative ctd kinase subunit alpha protein [Eutypa lata UCREL1] | 67.4 | 0.0 |
| DIP_10005212 | NA | | |
| DIP_10005213 | NA | | |
| DIP_10005214 | Uncharacterized protein SAPIO_CDS10439 [Scedosporium apiospermum] | 39.7 | 5.0E-117 |
| DIP_10005215 | hypothetical protein PFICI_09466 [Pestalotiopsis fici W106-1] | 62.7 | 0.0 |
| DIP_10005216 | hypothetical protein UCREL1_6423 [Eutypa lata UCREL1] | 47.7 | 1.0E-61 |
| DIP_10005217 | ndt80 like dna-binding family protein [Colletotrichum orbiculare MAFF 240422] | 51.5 | 0.0 |
| DIP_10005218 | hypothetical protein PFICI_09468 [Pestalotiopsis fici W106-1] | 58.5 | 0.0 |
| DIP_10005219 | putative pap2 domain protein [Eutypa lata UCREL1] | 72.3 | 5.0E-127 |
| DIP_10005220 | putative succinate dehydrogenase cytochrome b subunit protein [Eutypa lata UCREL1] | 49.2 | 6.0E-54 |
| DIP_10005221 | putative transcription initiation factor tfiid subunit 6 protein [Togninia minima UCRPA7] | 70.7 | 0.0 |
| DIP_10005222 | hypothetical protein PFICI_11663 [Pestalotiopsis fici W106-1] | 82 | 5.0E-63 |
| DIP_10005223 | putative nad transhydrogenase protein [Eutypa lata UCREL1] | 78.2 | 0.0 |
| DIP_10005224 | hypothetical protein CGGC5_12162 [Colletotrichum gloeosporioides Nara gc5] | 35.2 | 2.0E-15 |
| DIP_10005225 | hypothetical protein PFICI_08043 [Pestalotiopsis fici W106-1] | 48.5 | 1.0E-116 |
| DIP_10005226 | hypothetical protein PFICI_08042 [Pestalotiopsis fici W106-1] | 40.2 | 9.0E-27 |
| DIP_10005227 | hypothetical protein UCREL1_1103 [Eutypa lata UCREL1] | 60.7 | 3.0E-79 |
| DIP_10005228 | NA | | |
| DIP_10005229 | putative cation diffusion facilitator 1 protein [Eutypa lata UCREL1] | 65.6 | 0.0 |
| DIP_10005230 | glycosyl transferase family 8 [Colletotrichum graminicola M1.001] | 50.3 | 0.0 |
| DIP_10005231 | hypothetical protein PFICI_07570 [Pestalotiopsis fici W106-1] | 69.4 | 0.0 |
| DIP_10005232 | NA | | |
| DIP_10005233 | hypothetical protein PFICI_07568 [Pestalotiopsis fici W106-1] | 71.6 | 0.0 |
| DIP_10005234 | NA | | |
| DIP_10005235 | hypothetical protein PFICI_11107 [Pestalotiopsis fici W106-1] | 84.9 | 0.0 |
| DIP_10005236 | putative atp-dependent permease mdl2 protein [Eutypa lata UCREL1] | 72.9 | 0.0 |
| DIP_10005237 | putative duf618 domain-containing protein [Eutypa lata UCREL1] | 72.6 | 5.0E-164 |
| DIP_10005238 | putative clathrin light chain protein [Eutypa lata UCREL1] | 65.7 | 1.0E-94 |
| DIP_10005239 | hypothetical protein PFICI_07622 [Pestalotiopsis fici W106-1] | 71.7 | 4.0E-27 |
| DIP_10005240 | putative pentatricopeptide repeat domain-containing protein [Eutypa lata UCREL1] | 39.8 | 1.0E-108 |
| DIP_10005241 | hypothetical protein HMPREF1624_00665 [Sporothrix schenckii ATCC 58251] | 67.6 | 2.0E-154 |
| DIP_10005242 | calmodulin [Cladophialophora bantiana CBS 173.52] | 98.7 | 2.0E-104 |
| DIP_10005243 | putative class i alpha-mannosidase protein [Eutypa lata UCREL1] | 59.5 | 0.0 |
| DIP_10005244 | NADH-cytochrome b-5 reductase [Colletotrichum graminicola M1.001] | 37.3 | 1.0E-69 |
| DIP_10005245 | hypothetical protein CHGG_02460 [Chaetomium globosum CBS 148.51] | 33.3 | 2.0E-14 |
| DIP_10005246 | hypothetical protein M419DRAFT_10782 [Trichoderma reesei RUT C-30] | 43.5 | 3.0E-36 |
| DIP_10005247 | hypothetical protein PFICI_11060 [Pestalotiopsis fici W106-1] | 79.2 | 4.0E-58 |
| DIP_10005248 | hypothetical protein PFICI_11061 [Pestalotiopsis fici W106-1] | 61.7 | 0.0 |
| DIP_10005249 | NA | | |
| DIP_10005250 | putative transformer-sr ribonucleo protein [Eutypa lata UCREL1] | 65.4 | 2.0E-116 |
| DIP_10005251 | putative nadph-dependent 1-acyldihydroxyacetone phosphate reductase protein [Eutypa lata UCREL1] | 69.4 | 9.0E-162 |
| DIP_10005252 | putative acyl- thioesterase ii protein [Eutypa lata UCREL1] | 43.9 | 1.0E-83 |
| DIP_10005253 | hypothetical protein TI39_contig332g00013 [Zymoseptoria brevis] | 49.8 | 4.0E-95 |
| DIP_10005254 | hypothetical protein NECHADRAFT_53787 [Nectria haematococca mpVI 77-13-4] | 43.1 | 1.0E-76 |
| DIP_10005255 | hypothetical protein PFICI_10944 [Pestalotiopsis fici W106-1] | 63 | 0.0 |
| DIP_10005256 | hypothetical protein PFICI_10951 [Pestalotiopsis fici W106-1] | 80.7 | 0.0 |
| DIP_10005257 | putative disease resistance protein [Eutypa lata UCREL1] | 61.1 | 1.0E-80 |
| DIP_10005258 | putative chitin- domain 3 protein [Eutypa lata UCREL1] | 44.3 | 3.0E-101 |
| DIP_10005259 | hypothetical protein TRIVIDRAFT_68639 [Trichoderma virens Gv29-8] | 30.6 | 4.0E-153 |
| DIP_10005260 | putative nuclear protein es2 protein [Eutypa lata UCREL1] | 65.4 | 0.0 |
| DIP_10005261 | hypothetical protein PFICI_07442 [Pestalotiopsis fici W106-1] | 78.9 | 0.0 |
| DIP_10005262 | hypothetical protein UCREL1_5910 [Eutypa lata UCREL1] | 52.6 | 3.0E-22 |
| DIP_10005263 | Glycogen synthase [Pestalotiopsis fici W106-1] | 91.2 | 0.0 |
| DIP_10005264 | putative f-box domain-containing protein [Eutypa lata UCREL1] | 62.8 | 0.0 |
| DIP_10005265 | NA | | |
| DIP_10005266 | NA | | |
| DIP_10005267 | DUF1754-domain-containing protein [Ustilaginoidea virens] | 45.1 | 7.0E-30 |
| DIP_10005268 | putative alpha-amylase [Phaeomoniella chlamydospora] | 48.2 | 4.0E-153 |
| DIP_10005269 | hypothetical protein MAJ_03385, partial [Metarhizium majus ARSEF 297] | 30.7 | 5.0E-06 |
| DIP_10005270 | putative phosphatidate cytidylyltransferase protein [Eutypa lata UCREL1] | 64.8 | 0.0 |
| DIP_10005271 | hypothetical protein PFICI_07869 [Pestalotiopsis fici W106-1] | 46.7 | 0.0 |
| DIP_10005272 | hypothetical protein PFICI_07870 [Pestalotiopsis fici W106-1] | 60.5 | 0.0 |
| DIP_10005273 | NA | | |
| DIP_10005274 | hypothetical protein UCREL1_2062 [Eutypa lata UCREL1] | 44.9 | 1.0E-33 |
| DIP_10005275 | NA | | |
| DIP_10005276 | hypothetical protein UCREL1_2063 [Eutypa lata UCREL1] | 32.6 | 5.0E-14 |
| DIP_10005277 | putative nap family protein [Eutypa lata UCREL1] | 57.1 | 4.0E-123 |
| DIP_10005278 | putative mannan endo- -alpha-mannosidase dcw1 protein [Eutypa lata UCREL1] | 52.3 | 5.0E-180 |
| DIP_10005279 | putative translation initiation factor protein [Eutypa lata UCREL1] | 39.4 | 2.0E-42 |
| DIP_10005280 | putative mitochondrial inner membrane protease subunit protein [Eutypa lata UCREL1] | 53 | 4.0E-59 |
| DIP_10005281 | putative vacuolar protein 8 protein [Togninia minima UCRPA7] | 92.9 | 0.0 |
| DIP_10005282 | Nuclear protein SNF4 [Pestalotiopsis fici W106-1] | 73.3 | 2.0E-129 |
| DIP_10005283 | hypothetical protein UCDDA912_g08780 [Diaporthe ampelina] | 41.8 | 5.0E-73 |
| DIP_10005284 | putative 26s protease regulatory subunit 4 protein [Eutypa lata UCREL1] | 94.6 | 0.0 |
| DIP_10005285 | putative succinyl- ligase subunit alpha protein [Eutypa lata UCREL1] | 86.1 | 0.0 |
| DIP_10005286 | hypothetical protein PFICI_07382 [Pestalotiopsis fici W106-1] | 38.6 | 1.0E-104 |
| DIP_10005287 | hypothetical protein Ao3042_10397 [Aspergillus oryzae 3.042] | 42.3 | 2.0E-22 |
| DIP_10005288 | hypothetical protein PFICI_07381 [Pestalotiopsis fici W106-1] | 63.8 | 0.0 |
| DIP_10005289 | hypothetical protein UCREL1_4812 [Eutypa lata UCREL1] | 31.7 | 1.0E-07 |
| DIP_10005290 | putative dihydroxy acid [Diaporthe ampelina] | 63.4 | 0.0 |
| DIP_10005291 | hypothetical protein ACRE_059450 [Acremonium chrysogenum ATCC 11550] | 38.6 | 4.0E-54 |
| DIP_10005292 | aldo/keto reductase [Colletotrichum higginsianum] | 72 | 3.0E-164 |
| DIP_10005293 | putative c6 transcription factor protein [Eutypa lata UCREL1] | 55.6 | 0.0 |
| DIP_10005294 | NA | | |
| DIP_10005295 | NA | | |
| DIP_10005296 | hypothetical protein CFO_g3799 [Ceratocystis platani] | 55.9 | 3.0E-90 |
| DIP_10005297 | hypothetical protein PFICI_07809 [Pestalotiopsis fici W106-1] | 74.7 | 9.0E-41 |
| DIP_10005298 | putative translation initiation factor eif-2b subunit delta protein [Eutypa lata UCREL1] | 68 | 0.0 |
| DIP_10005299 | putative dihydroxyacetone kinase protein [Eutypa lata UCREL1] | 67.7 | 0.0 |
| DIP_10005300 | small nuclear ribonucleoprotein Sm D2 [Gaeumannomyces graminis var. tritici R3-111a-1] | 93.2 | 3.0E-76 |
| DIP_10005301 | putative 3-hydroxyisobutyryl- hydrolase protein [Togninia minima UCRPA7] | 60.8 | 0.0 |
| DIP_10005302 | hypothetical protein PV06_05002 [Exophiala oligosperma] | 40.1 | 3.0E-73 |
| DIP_10005303 | putative gata transcription factor [Diaporthe ampelina] | 64.4 | 7.0E-61 |
| DIP_10005304 | GATA transcription factor (Ams2) [Ophiocordyceps sinensis CO18] | 34.6 | 1.0E-67 |
| DIP_10005305 | NA | | |
| DIP_10005306 | putative replication factor a protein 3 protein [Eutypa lata UCREL1] | 74.3 | 1.0E-55 |
| DIP_10005307 | hypothetical protein FAVG1_00263 [Fusarium avenaceum] | 37.4 | 1.0E-28 |
| DIP_10005308 | predicted protein [Nectria haematococca mpVI 77-13-4] | 72.3 | 2.0E-118 |
| DIP_10005309 | hypothetical protein PFICI_09452 [Pestalotiopsis fici W106-1] | 75.8 | 2.0E-113 |
| DIP_10005310 | putative ribosomal protein l32 protein [Botrytis cinerea BcDW1] | 47.1 | 1.0E-33 |
| DIP_10005311 | hypothetical protein PFICI_10605 [Pestalotiopsis fici W106-1] | 39.7 | 9.0E-90 |
| DIP_10005312 | NA | | |
| DIP_10005313 | NA | | |
| DIP_10005314 | NA | | |
| DIP_10005315 | NA | | |
| DIP_10005316 | hypothetical protein PFICI_11665 [Pestalotiopsis fici W106-1] | 72.1 | 0.0 |
| DIP_10005317 | putative nadh-ubiquinone oxidoreductase subunit protein [Eutypa lata UCREL1] | 79.1 | 4.0E-79 |
| DIP_10005318 | NA | | |
| DIP_10005319 | NA | | |
| DIP_10005320 | hypothetical protein BBA_01908 [Beauveria bassiana ARSEF 2860] | 41.5 | 1.0E-06 |
| DIP_10005321 | NA | | |
| DIP_10005322 | hypothetical protein PFICI_04116 [Pestalotiopsis fici W106-1] | 59.2 | 0.0 |
| DIP_10005323 | hypothetical protein CH063_06757 [Colletotrichum higginsianum] | 49 | 1.0E-133 |
| DIP_10005324 | proline racemase [Colletotrichum gloeosporioides Cg-14] | 58.5 | 8.0E-141 |
| DIP_10005325 | hypothetical protein FVEG_01820 [Fusarium verticillioides 7600] | 41.6 | 7.0E-92 |
| DIP_10005326 | hypothetical protein CFIO01_07022 [Colletotrichum fioriniae PJ7] | 34.1 | 1.0E-89 |
| DIP_10005327 | amino acid transporter, putative [Aspergillus flavus NRRL3357] | 67.2 | 0.0 |
| DIP_10005328 | hypothetical protein PV10_03524 [Exophiala mesophila] | 34.3 | 1.0E-28 |
| DIP_10005329 | hypothetical protein PFICI_04119 [Pestalotiopsis fici W106-1] | 67.8 | 2.0E-129 |
| DIP_10005330 | hypothetical protein MYCTH_2300011 [Myceliophthora thermophila ATCC 42464] | 57.7 | 1.0E-39 |
| DIP_10005331 | putative glc8 protein [Eutypa lata UCREL1] | 62.4 | 3.0E-95 |
| DIP_10005332 | NA | | |
| DIP_10005333 | hypothetical protein PFICI_04122 [Pestalotiopsis fici W106-1] | 55.6 | 0.0 |
| DIP_10005334 | hypothetical protein PFICI_04123 [Pestalotiopsis fici W106-1] | 62.4 | 0.0 |
| DIP_10005335 | hypothetical protein TRIATDRAFT_318140 [Trichoderma atroviride IMI 206040] | 57.2 | 3.0E-93 |
| DIP_10005336 | endonuclease/Exonuclease/phosphatase [Colletotrichum gloeosporioides Cg-14] | 67.6 | 0.0 |
| DIP_10005337 | hypothetical protein CGGC5_12530, partial [Colletotrichum gloeosporioides Nara gc5] | 39.6 | 1.0E-99 |
| DIP_10005338 | Acyl-CoA dehydrogenase-like protein [Acremonium chrysogenum ATCC 11550] | 73.8 | 0.0 |
| DIP_10005339 | Acyl-CoA dehydrogenase family member-like protein [Acremonium chrysogenum ATCC 11550] | 69.7 | 0.0 |
| DIP_10005340 | hypothetical protein UCREL1_4044 [Eutypa lata UCREL1] | 71.9 | 1.0E-68 |
| DIP_10005341 | hypothetical protein PFICI_07777 [Pestalotiopsis fici W106-1] | 69.1 | 0.0 |
| DIP_10005342 | NA | | |
| DIP_10005343 | S-adenosyl-L-methionine-dependent methyltransferase [Glarea lozoyensis ATCC 20868] | 50.3 | 9.0E-141 |
| DIP_10005344 | NA | | |
| DIP_10005345 | NA | | |
| DIP_10005346 | NA | | |
| DIP_10005347 | NA | | |
| DIP_10005348 | putative lccl domain containing protein [Eutypa lata UCREL1] | 65.1 | 0.0 |
| DIP_10005349 | hypothetical protein PFICI_04073 [Pestalotiopsis fici W106-1] | 51.8 | 3.0E-66 |
| DIP_10005350 | NA | | |
| DIP_10005351 | putative alkaline phosphatase protein [Eutypa lata UCREL1] | 62.5 | 0.0 |
| DIP_10005352 | NA | | |
| DIP_10005353 | NA | | |
| DIP_10005354 | putative cs domain-containing protein [Eutypa lata UCREL1] | 58 | 6.0E-147 |
| DIP_10005355 | putative lanosterol synthase protein [Eutypa lata UCREL1] | 77.2 | 0.0 |
| DIP_10005356 | hypothetical protein PFICI_07705 [Pestalotiopsis fici W106-1] | 57.3 | 5.0E-65 |
| DIP_10005357 | putative phosphoinositide-specific phospholipase c protein [Eutypa lata UCREL1] | 69.1 | 0.0 |
| DIP_10005358 | hypothetical protein PFICI_11083 [Pestalotiopsis fici W106-1] | 63.3 | 0.0 |
| DIP_10005359 | putative c2h2 finger domain protein [Eutypa lata UCREL1] | 30.8 | 1.0E-25 |
| DIP_10005360 | NA | | |
| DIP_10005361 | putative glutathione-dependent formaldehyde-activating family gfa protein [Eutypa lata UCREL1] | 70.9 | 3.0E-101 |
| DIP_10005362 | putative origin recognition complex subunit 2 protein [Eutypa lata UCREL1] | 59.6 | 0.0 |
| DIP_10005363 | hypothetical protein PFICI_09638 [Pestalotiopsis fici W106-1] | 61.9 | 1.0E-89 |
| DIP_10005364 | hypothetical protein PFICI_15326 [Pestalotiopsis fici W106-1] | 42.9 | 5.0E-49 |
| DIP_10005365 | putative copper homeostasis protein cutc protein [Eutypa lata UCREL1] | 41 | 6.0E-56 |
| DIP_10005366 | hypothetical protein UCREL1_11294 [Eutypa lata UCREL1] | 36.4 | 2.0E-35 |
| DIP_10005367 | NA | | |
| DIP_10005368 | NA | | |
| DIP_10005369 | NA | | |
| DIP_10005370 | hypothetical protein PFICI_11606 [Pestalotiopsis fici W106-1] | 72 | 0.0 |
| DIP_10005371 | hypothetical protein PFICI_11607 [Pestalotiopsis fici W106-1] | 59.6 | 2.0E-136 |
| DIP_10005372 | putative carboxypeptidase s1 protein [Eutypa lata UCREL1] | 68.2 | 0.0 |
| DIP_10005373 | hypothetical protein UCREL1_8600 [Eutypa lata UCREL1] | 58.9 | 0.0 |
| DIP_10005374 | putative phenylacetaldoxime dehydratase protein [Eutypa lata UCREL1] | 48.7 | 9.0E-127 |
| DIP_10005375 | hypothetical protein EMCG_03165 [Emmonsia crescens UAMH 3008] | 58 | 8.0E-144 |
| DIP_10005376 | putative glycoside hydrolase family 47 protein [Eutypa lata UCREL1] | 59.5 | 0.0 |
| DIP_10005377 | hypothetical protein PFICI_15205 [Pestalotiopsis fici W106-1] | 47.4 | 6.0E-80 |
| DIP_10005378 | hypothetical protein FOPG_17148 [Fusarium oxysporum f. sp. conglutinans race 2 54008] | 30.1 | 1.0E-17 |
| DIP_10005379 | putative duf1665 domain-containing protein [Eutypa lata UCREL1] | 48.5 | 3.0E-88 |
| DIP_10005380 | 2Fe-2S iron-sulfur cluster binding domain-containing protein [Neurospora crassa OR74A] | 83.9 | 5.0E-79 |
| DIP_10005381 | hypothetical protein PFICI_04169 [Pestalotiopsis fici W106-1] | 66.1 | 0.0 |
| DIP_10005382 | NA | | |
| DIP_10005383 | hypothetical protein SAPIO_CDS6239 [Scedosporium apiospermum] | 30.4 | 4.0E-47 |
| DIP_10005384 | Protein kinase-like domain protein, partial [Metarhizium brunneum ARSEF 3297] | 56.6 | 4.0E-156 |
| DIP_10005385 | endoglucanase IV [Neurospora crassa OR74A] | 55.4 | 2.0E-132 |
| DIP_10005386 | hypothetical protein PV07_01793 [Cladophialophora immunda] | 54.8 | 4.0E-99 |
| DIP_10005387 | hypothetical protein PFICI_06672 [Pestalotiopsis fici W106-1] | 34.2 | 3.0E-43 |
| DIP_10005388 | hypothetical protein ACRE_049090 [Acremonium chrysogenum ATCC 11550] | 35.7 | 2.0E-68 |
| DIP_10005389 | putative splicing factor 3a subunit 3 protein [Eutypa lata UCREL1] | 77.9 | 0.0 |
| DIP_10005390 | hypothetical protein CGLO_00551 [Colletotrichum gloeosporioides Cg-14] | 96.8 | 6.0E-106 |
| DIP_10005391 | eukaryotic initiation factor 4A-12 [Neurospora crassa OR74A] | 100 | 4.0E-171 |
| DIP_10005392 | hypothetical protein A1O1_08839 [Capronia coronata CBS 617.96] | 50.6 | 1.0E-46 |
| DIP_10005393 | NA | | |
| DIP_10005394 | hypothetical protein PFICI_03915 [Pestalotiopsis fici W106-1] | 68.4 | 0.0 |
| DIP_10005395 | hypothetical protein PFICI_03914 [Pestalotiopsis fici W106-1] | 61 | 1.0E-145 |
| DIP_10005396 | FAD binding domain protein [Colletotrichum gloeosporioides Nara gc5] | 65.1 | 0.0 |
| DIP_10005397 | putative histone acetyltransferase protein [Eutypa lata UCREL1] | 92.6 | 0.0 |
| DIP_10005398 | hypothetical protein PFICI_08526 [Pestalotiopsis fici W106-1] | 60.3 | 2.0E-139 |
| DIP_10005399 | UBC-like protein [Glarea lozoyensis ATCC 20868] | 59.1 | 5.0E-95 |
| DIP_10005400 | hypothetical protein PFICI_11172 [Pestalotiopsis fici W106-1] | 50 | 3.0E-56 |
| DIP_10005401 | hypothetical protein PFICI_10755 [Pestalotiopsis fici W106-1] | 74.6 | 0.0 |
| DIP_10005402 | NA | | |
| DIP_10005403 | hypothetical protein OIDMADRAFT_44475 [Oidiodendron maius Zn] | 48.9 | 4.0E-82 |
| DIP_10005404 | short-chain dehydrogenase [Neosartorya fischeri NRRL 181] | 69 | 2.0E-163 |
| DIP_10005405 | alpha-1,2-mannosyltransferase (Mnn2) [Aspergillus niger CBS 513.88] | 31.1 | 1.0E-38 |
| DIP_10005406 | hypothetical protein GGTG_06525 [Gaeumannomyces graminis var. tritici R3-111a-1] | 33.2 | 9.0E-17 |
| DIP_10005407 | hypothetical protein Cob_05868 [Colletotrichum orbiculare MAFF 240422] | 45.2 | 6.0E-25 |
| DIP_10005408 | hypothetical protein FAVG1_06112 [Fusarium avenaceum] | 50.8 | 2.0E-38 |
| DIP_10005409 | hypothetical protein UCREL1_5556 [Eutypa lata UCREL1] | 61.9 | 2.0E-43 |
| DIP_10005410 | putative translocation protein sec72 protein [Eutypa lata UCREL1] | 70.5 | 2.0E-81 |
| DIP_10005411 | hypothetical protein VDAG_07217 [Verticillium dahliae VdLs.17] | 30.5 | 2.0E-42 |
| DIP_10005412 | NA | | |
| DIP_10005413 | hypothetical protein FAVG1_06127 [Fusarium avenaceum] | 42.6 | 1.0E-64 |
| DIP_10005414 | putative glycoside hydrolase family 72 protein [Eutypa lata UCREL1] | 58.5 | 0.0 |
| DIP_10005415 | hypothetical protein PFICI_10764 [Pestalotiopsis fici W106-1] | 39.3 | 9.0E-77 |
| DIP_10005416 | putative 1-acyl-sn-glycerol-3-phosphate acyltransferase 2 protein [Eutypa lata UCREL1] | 68.1 | 0.0 |
| DIP_10005417 | hypothetical protein GGTG_06017 [Gaeumannomyces graminis var. tritici R3-111a-1] | 51.9 | 1.0E-87 |
| DIP_10005418 | hypothetical protein V491_01693, partial [Pseudogymnoascus pannorum VKM F-3775] | 60 | 8.0E-14 |
| DIP_10005419 | NADH kinase POS5 [Magnaporthe oryzae Y34] | 58.6 | 0.0 |
| DIP_10005420 | putative rodlet peptide protein [Eutypa lata UCREL1] | 70.3 | 3.0E-63 |
| DIP_10005421 | NA | | |
| DIP_10005422 | NA | | |
| DIP_10005423 | DNA helicase INO80 [Pestalotiopsis fici W106-1] | 74.2 | 0.0 |
| DIP_10005424 | putative atp-binding cassette sub-family f member 2 protein [Eutypa lata UCREL1] | 85.6 | 0.0 |
| DIP_10005425 | hypothetical protein VDAG_07787 [Verticillium dahliae VdLs.17] | 33.3 | 1.0E-22 |
| DIP_10005426 | putative tyrosine phosphatase protein [Eutypa lata UCREL1] | 62.1 | 3.0E-135 |
| DIP_10005427 | hypothetical protein PFICI_10750 [Pestalotiopsis fici W106-1] | 64.5 | 3.0E-155 |
| DIP_10005428 | hypothetical protein PFICI_09186 [Pestalotiopsis fici W106-1] | 43.5 | 1.0E-71 |
| DIP_10005429 | putative proline-specific permease protein [Eutypa lata UCREL1] | 79.4 | 4.0E-167 |
| DIP_10005430 | hypothetical protein PFICI_10737 [Pestalotiopsis fici W106-1] | 57.7 | 2.0E-116 |
| DIP_10005431 | hypothetical protein AN1737.2 [Aspergillus nidulans FGSC A4] | 49 | 7.0E-160 |
| DIP_10005432 | hypothetical protein PTT_09944 [Pyrenophora teres f. teres 0-1] | 70.3 | 0.0 |
| DIP_10005433 | putative dcp1-like decapping family protein [Eutypa lata UCREL1] | 33.2 | 4.0E-24 |
| DIP_10005434 | putative trna pseudouridine synthase b protein [Eutypa lata UCREL1] | 71.1 | 0.0 |
| DIP_10005435 | NA | | |
| DIP_10005436 | hypothetical protein UCREL1_9414 [Eutypa lata UCREL1] | 61.6 | 7.0E-169 |
| DIP_10005437 | hypothetical protein COCMIDRAFT_80258, partial [Bipolaris oryzae ATCC 44560] | 37.7 | 2.0E-70 |
| DIP_10005438 | hypothetical protein PFICI_03441 [Pestalotiopsis fici W106-1] | 55.5 | 6.0E-155 |
| DIP_10005439 | hypothetical protein PV08_07706 [Exophiala spinifera] | 33.2 | 1.0E-58 |
| DIP_10005440 | hypothetical protein SMAC_01541 [Sordaria macrospora k-hell] | 42.1 | 8.0E-46 |
| DIP_10005441 | hypothetical protein TRIATDRAFT_54502 [Trichoderma atroviride IMI 206040] | 60.4 | 3.0E-100 |
| DIP_10005442 | putative fic doc family protein [Eutypa lata UCREL1] | 52.8 | 1.0E-96 |
| DIP_10005443 | hypothetical protein FVEG_12881 [Fusarium verticillioides 7600] | 41.8 | 3.0E-66 |
| DIP_10005444 | putative methyltransferase domain-containing protein [Botrytis cinerea BcDW1] | 48.8 | 6.0E-95 |
| DIP_10005445 | NA | | |
| DIP_10005446 | hypothetical protein V494_00530 [Pseudogymnoascus pannorum VKM F-4513 (FW-928)] | 52.3 | 3.0E-105 |
| DIP_10005447 | putative rna polymerase ii mediator complex component protein [Togninia minima UCRPA7] | 43.5 | 4.0E-89 |
| DIP_10005448 | putative nad dependent epimerase protein [Togninia minima UCRPA7] | 47.5 | 8.0E-82 |
| DIP_10005449 | hypothetical protein V497_04033 [Pseudogymnoascus pannorum VKM F-4516 (FW-969)] | 30.2 | 6.0E-19 |
| DIP_10005450 | hypothetical protein Cob_11492 [Colletotrichum orbiculare MAFF 240422] | 37.6 | 2.0E-166 |
| DIP_10005451 | NA | | |
| DIP_10005452 | Xylose isomerase-like TIM barrel [Aspergillus parasiticus SU-1] | 53.7 | 1.0E-103 |
| DIP_10005453 | Alcohol dehydrogenase GroES-like protein [Penicillium solitum] | 81.8 | 0.0 |
| DIP_10005454 | hypothetical protein CFIO01_07592 [Colletotrichum fioriniae PJ7] | 86.3 | 0.0 |
| DIP_10005455 | methionine permease [Colletotrichum fioriniae PJ7] | 60.7 | 0.0 |
| DIP_10005456 | hypothetical protein VDAG_10462 [Verticillium dahliae VdLs.17] | 71.3 | 1.0E-144 |
| DIP_10005457 | hypothetical protein PFICI_03037 [Pestalotiopsis fici W106-1] | 80.5 | 0.0 |
| DIP_10005458 | NA | | |
| DIP_10005459 | serine/threonine protein kinase [Metarhizium album ARSEF 1941] | 43.7 | 2.0E-80 |
| DIP_10005460 | hypothetical protein PFICI_04744 [Pestalotiopsis fici W106-1] | 35.8 | 1.0E-91 |
| DIP_10005461 | Glucose-6-phosphate isomerase [Pestalotiopsis fici W106-1] | 88.4 | 0.0 |
| DIP_10005462 | putative serine threonine-protein kinase gad8 protein [Eutypa lata UCREL1] | 91.1 | 0.0 |
| DIP_10005463 | putative capsular associated protein [Eutypa lata UCREL1] | 61.3 | 0.0 |
| DIP_10005464 | NA | | |
| DIP_10005465 | putative mannitol-1-phosphate 5-dehydrogenase protein [Eutypa lata UCREL1] | 72.3 | 0.0 |
| DIP_10005466 | putative coatomer beta subunit protein [Eutypa lata UCREL1] | 88.8 | 0.0 |
| DIP_10005467 | hypothetical protein UCREL1_7785 [Eutypa lata UCREL1] | 36.5 | 1.0E-51 |
| DIP_10005468 | hypothetical protein UCDDA912_g02476 [Diaporthe ampelina] | 71.5 | 2.0E-146 |
| DIP_10005469 | NA | | |
| DIP_10005470 | hypothetical protein S40288_10289 [Stachybotrys chartarum IBT 40288] | 52.6 | 9.0E-60 |
| DIP_10005471 | hypothetical protein U727_00431291101 [Penicillium solitum] | 58.3 | 2.0E-30 |
| DIP_10005472 | NA | | |
| DIP_10005473 | ankyrin and het domain-containing protein [Colletotrichum gloeosporioides Nara gc5] | 34.3 | 6.0E-106 |
| DIP_10005474 | NA | | |
| DIP_10005475 | putative integral membrane protein [Eutypa lata UCREL1] | 64.9 | 4.0E-70 |
| DIP_10005476 | NA | | |
| DIP_10005477 | putative aminotransferase family protein [Eutypa lata UCREL1] | 58.3 | 3.0E-177 |
| DIP_10005478 | U-box domain-containing protein [Colletotrichum higginsianum] | 63 | 3.0E-107 |
| DIP_10005479 | putative 2 -cyclic-nucleotide 3 -phosphodiesterase protein [Eutypa lata UCREL1] | 60.1 | 1.0E-91 |
| DIP_10005480 | hypothetical protein FOTG_16244 [Fusarium oxysporum f. sp. vasinfectum 25433] | 58.3 | 0.0 |
| DIP_10005481 | putative peptidase protein [Eutypa lata UCREL1] | 54.4 | 4.0E-154 |
| DIP_10005482 | putative ferulic acid esterase protein [Togninia minima UCRPA7] | 57.6 | 3.0E-126 |
| DIP_10005483 | sure-like protein [Auricularia delicata TFB-10046 SS5] | 38.4 | 8.0E-25 |
| DIP_10005484 | hypothetical protein UCREL1_6206 [Eutypa lata UCREL1] | 46.9 | 6.0E-157 |
| DIP_10005485 | hypothetical protein PFICI_12551 [Pestalotiopsis fici W106-1] | 32.6 | 4.0E-12 |
| DIP_10005486 | hypothetical protein S40293_08154 [Stachybotrys chartarum IBT 40293] | 36.7 | 1.0E-59 |
| DIP_10005487 | hypothetical protein NECHADRAFT_44961 [Nectria haematococca mpVI 77-13-4] | 54.7 | 4.0E-132 |

| | | | |
|---|---|---|---|
| DIP_10005488 | NA | | |
| DIP_10005489 | hsc70 cochaperone [Grosmannia clavigera kw1407] | 65 | 1.0E-156 |
| DIP_10005490 | hypothetical protein PFICI_01137 [Pestalotiopsis fici W106-1] | 57.1 | 0.0 |
| DIP_10005491 | NA | | |
| DIP_10005492 | hypothetical protein PV04_06458 [Capronia semiimmersa] | 57.5 | 3.0E-99 |
| DIP_10005493 | NA | | |
| DIP_10005494 | NA | | |
| DIP_10005495 | NA | | |
| DIP_10005496 | putative mitochondrial chaperone bcs1 protein [Eutypa lata UCREL1] | 72.3 | 0.0 |
| DIP_10005497 | hypothetical protein Z519_02046 [Cladophialophora bantiana CBS 173.52] | 63.4 | 0.0 |
| DIP_10005498 | hypothetical protein NECHADRAFT_97166 [Nectria haematococca mpVI 77-13-4] | 51.5 | 4.0E-107 |
| DIP_10005499 | putative integral membrane protein [Eutypa lata UCREL1] | 58.1 | 7.0E-87 |
| DIP_10005500 | hypothetical protein THITE_2109409 [Thielavia terrestris NRRL 8126] | 74.1 | 0.0 |
| DIP_10005501 | putative zinc finger protein gcs1 protein [Togninia minima UCRPA7] | 65.7 | 6.0E-180 |
| DIP_10005502 | putative rnp domain-containing protein [Eutypa lata UCREL1] | 86.8 | 6.0E-177 |
| DIP_10005503 | hypothetical protein PFICI_05656 [Pestalotiopsis fici W106-1] | 44.5 | 0.0 |
| DIP_10005504 | putative sap domain-containing protein [Eutypa lata UCREL1] | 45.3 | 8.0E-132 |
| DIP_10005505 | NA | | |
| DIP_10005506 | NA | | |
| DIP_10005507 | hypothetical protein PFICI_04719 [Pestalotiopsis fici W106-1] | 43.6 | 0.0 |
| DIP_10005508 | putative gpi-anchor transamidase precursor protein [Eutypa lata UCREL1] | 79.2 | 0.0 |
| DIP_10005509 | hypothetical protein PFICI_04939 [Pestalotiopsis fici W106-1] | 45.5 | 0.0 |
| DIP_10005510 | hypothetical protein UCREL1_9294 [Eutypa lata UCREL1] | 48.2 | 5.0E-140 |
| DIP_10005511 | putative protein bli-3 protein [Togninia minima UCRPA7] | 66.2 | 5.0E-101 |
| DIP_10005512 | putative diphthine synthase protein [Eutypa lata UCREL1] | 88.8 | 0.0 |
| DIP_10005513 | putative c2h2 finger domain-containing protein [Eutypa lata UCREL1] | 65.1 | 2.0E-88 |
| DIP_10005514 | hypothetical protein S40285_07877 [Stachybotrys chlorohalonata IBT 40285] | 64.3 | 8.0E-78 |
| DIP_10005515 | hypothetical protein PFICI_04935 [Pestalotiopsis fici W106-1] | 59.2 | 0.0 |
| DIP_10005516 | hypothetical protein PFICI_04936 [Pestalotiopsis fici W106-1] | 63.9 | 2.0E-150 |
| DIP_10005517 | hypothetical protein MYCTH_2298349 [Myceliophthora thermophila ATCC 42464] | 79.8 | 0.0 |
| DIP_10005518 | NA | | |
| DIP_10005519 | NA | | |
| DIP_10005520 | putative atp-dependent rna helicase dhx8 protein [Eutypa lata UCREL1] | 79.6 | 0.0 |
| DIP_10005521 | putative alpha- -mannosyltransferase protein [Eutypa lata UCREL1] | 65.2 | 0.0 |
| DIP_10005522 | putative golgi apparatus membrane protein tvp23 protein [Eutypa lata UCREL1] | 89.9 | 2.0E-126 |
| DIP_10005523 | NA | | |
| DIP_10005524 | hypothetical protein PFICI_00547 [Pestalotiopsis fici W106-1] | 48 | 5.0E-66 |
| DIP_10005525 | putative cop9 signalosome complex subunit 4 protein [Togninia minima UCRPA7] | 58.1 | 6.0E-169 |
| DIP_10005526 | retinol dehydrogenase 12 [Colletotrichum higginsianum] | 49.9 | 4.0E-104 |
| DIP_10005527 | Galactose-1-phosphate uridylyltransferase [Pestalotiopsis fici W106-1] | 67.6 | 0.0 |
| DIP_10005528 | putative transmembrane domain-containing protein [Eutypa lata UCREL1] | 67.8 | 5.0E-83 |
| DIP_10005529 | putative 3-ketoacyl- reductase protein [Eutypa lata UCREL1] | 72.7 | 0.0 |
| DIP_10005530 | putative nucleoporin nup157 170 protein [Eutypa lata UCREL1] | 68.9 | 0.0 |
| DIP_10005531 | putative atp-dependent protease la protein [Eutypa lata UCREL1] | 77 | 0.0 |
| DIP_10005532 | NA | | |
| DIP_10005533 | hypothetical protein BC1G_16051 [Botrytis cinerea B05.10] | 38.8 | 6.0E-30 |
| DIP_10005534 | hypothetical protein PFICI_13943 [Pestalotiopsis fici W106-1] | 47.8 | 1.0E-99 |
| DIP_10005535 | hypothetical protein PFICI_13942 [Pestalotiopsis fici W106-1] | 30.4 | 9.0E-30 |
| DIP_10005536 | uncharacterized protein FFUJ_12058 [Fusarium fujikuroi IMI 58289] | 32.6 | 3.0E-85 |
| DIP_10005537 | putative RIO1 family protein [Colletotrichum sublineola] | 63.7 | 0.0 |
| DIP_10005538 | putative rna binding protein mrsp-2 protein [Eutypa lata UCREL1] | 59.2 | 0.0 |
| DIP_10005539 | NADPH-cytochrome P450 reductase [Diaporthe amygdali] | 82.2 | 0.0 |
| DIP_10005540 | putative cell morphogenesis protein pag1 protein [Eutypa lata UCREL1] | 74.7 | 0.0 |
| DIP_10005541 | hypothetical protein TRIATDRAFT_51231 [Trichoderma atroviride IMI 206040] | 62.9 | 2.0E-61 |
| DIP_10005542 | putative ribosome biogenesis protein urb1 protein [Eutypa lata UCREL1] | 54.9 | 0.0 |
| DIP_10005543 | hypothetical protein MYCFIDRAFT_47095 [Pseudocercospora fijiensis CIRAD86] | 38.7 | 1.0E-12 |
| DIP_10005544 | neuronal calcium sensor 1 [Colletotrichum gloeosporioides Nara gc5] | 97.9 | 2.0E-134 |
| DIP_10005545 | putative sna41 protein [Metarhizium acridum CQMa 102] | 69.5 | 0.0 |
| DIP_10005546 | hypothetical protein THITE_2115825 [Thielavia terrestris NRRL 8126] | 41.2 | 9.0E-84 |
| DIP_10005547 | NA | | |
| DIP_10005548 | hypothetical protein PFICI_05225 [Pestalotiopsis fici W106-1] | 48.7 | 4.0E-139 |
| DIP_10005549 | NA | | |
| DIP_10005550 | hypothetical protein NECHADRAFT_85878 [Nectria haematococca mpVI 77-13-4] | 43.4 | 0.0 |
| DIP_10005551 | phytanoyl- hydroxylase [Fusarium avenaceum] | 70.4 | 0.0 |
| DIP_10005552 | MFS quinate transporter [Colletotrichum gloeosporioides Nara gc5] | 68.3 | 0.0 |
| DIP_10005553 | hypothetical protein VHEMI09293 [Torrubiella hemipterigena] | 51.3 | 1.0E-84 |
| DIP_10005554 | hypothetical protein PFICI_04842 [Pestalotiopsis fici W106-1] | 61.4 | 3.0E-72 |
| DIP_10005555 | mate efflux family protein [Colletotrichum gloeosporioides Nara gc5] | 60.7 | 0.0 |
| DIP_10005556 | putative isopentenyl-diphosphate delta-isomerase protein [Eutypa lata UCREL1] | 85.1 | 4.0E-161 |
| DIP_10005557 | putative replication protein a 32 kda subunit protein [Eutypa lata UCREL1] | 69.5 | 2.0E-128 |
| DIP_10005558 | NA | | |
| DIP_10005559 | hypothetical protein S7711_03675 [Stachybotrys chartarum IBT 7711] | 45.2 | 1.0E-120 |
| DIP_10005560 | hypothetical protein PFICI_04843 [Pestalotiopsis fici W106-1] | 45.9 | 2.0E-174 |
| DIP_10005561 | NA | | |
| DIP_10005562 | putative arm-like repeat-containing protein [Eutypa lata UCREL1] | 56.2 | 0.0 |
| DIP_10005563 | hypothetical protein UCREL1_11427 [Eutypa lata UCREL1] | 82 | 3.0E-88 |
| DIP_10005564 | ATP-dependent RNA helicase DBP10 [Fusarium graminearum PH-1] | 68.6 | 0.0 |
| DIP_10005565 | putative o- protein [Eutypa lata UCREL1] | 34.8 | 2.0E-91 |
| DIP_10005566 | NA | | |
| DIP_10005567 | putative 26s proteasome regulatory subunit rpn-1 protein [Eutypa lata UCREL1] | 85.7 | 0.0 |
| DIP_10005568 | putative pro-apoptotic serine protease nma111 protein [Eutypa lata UCREL1] | 79.4 | 0.0 |
| DIP_10005569 | hypothetical protein VHEMI03596 [Torrubiella hemipterigena] | 43.8 | 5.0E-31 |
| DIP_10005570 | NA | | |
| DIP_10005571 | hypothetical protein MGG_11160 [Magnaporthe oryzae 70-15] | 32.8 | 8.0E-91 |
| DIP_10005572 | NA | | |
| DIP_10005573 | putative 2og-fe oxygenase protein [Eutypa lata UCREL1] | 61.4 | 3.0E-119 |
| DIP_10005574 | hypothetical protein NEUTE1DRAFT_121091 [Neurospora tetrasperma FGSC 2508] | 40 | 4.0E-100 |
| DIP_10005575 | hypothetical protein PFICI_05067 [Pestalotiopsis fici W106-1] | 36.6 | 5.0E-125 |
| DIP_10005576 | putative mitochondrial dna-directed rna polymerase protein [Eutypa lata UCREL1] | 57.6 | 0.0 |
| DIP_10005577 | hypothetical protein PFICI_05065 [Pestalotiopsis fici W106-1] | 31.1 | 6.0E-06 |
| DIP_10005578 | putative gu4 nucleic-binding protein [Eutypa lata UCREL1] | 75.6 | 0.0 |
| DIP_10005579 | NA | | |
| DIP_10005580 | putative v-atpase proteolipid subunit protein [Eutypa lata UCREL1] | 80.4 | 2.0E-109 |
| DIP_10005581 | putative phosphoinositide 3-kinase regulatory subunit 4 protein [Eutypa lata UCREL1] | 81.7 | 0.0 |
| DIP_10005582 | putative sh3 domain-containing protein [Diplodia seriata] | 45.4 | 5.0E-104 |
| DIP_10005583 | hypothetical protein UCREL1_4423 [Eutypa lata UCREL1] | 80.7 | 4.0E-47 |
| DIP_10005584 | hypothetical protein PFICI_04967 [Pestalotiopsis fici W106-1] | 55.5 | 1.0E-150 |
| DIP_10005585 | hypothetical protein PFICI_04969 [Pestalotiopsis fici W106-1] | 65.9 | 0.0 |
| DIP_10005586 | NA | | |
| DIP_10005587 | NA | | |
| DIP_10005588 | putative translocation protein sec62 protein [Eutypa lata UCREL1] | 76.8 | 4.0E-162 |
| DIP_10005589 | hypothetical protein PFICI_04840 [Pestalotiopsis fici W106-1] | 39.9 | 2.0E-46 |
| DIP_10005590 | hypothetical protein PFICI_07031 [Pestalotiopsis fici W106-1] | 33 | 2.0E-102 |
| DIP_10005591 | putative 3-phytase a protein [Eutypa lata UCREL1] | 62.3 | 0.0 |
| DIP_10005592 | hypothetical protein AKAW_08258 [Aspergillus kawachii IFO 4308] | 30.8 | 4.0E-11 |
| DIP_10005593 | hypothetical protein Cob_10368 [Colletotrichum orbiculare MAFF 240422] | 48.3 | 4.0E-45 |
| DIP_10005594 | hypothetical protein PFICI_05004 [Pestalotiopsis fici W106-1] | 58.3 | 1.0E-131 |
| DIP_10005595 | hypothetical protein PFICI_05005 [Pestalotiopsis fici W106-1] | 42.2 | 1.0E-94 |
| DIP_10005596 | putative major facilitator superfamily transporter protein [Togninia minima UCRPA7] | 69.8 | 0.0 |
| DIP_10005597 | putative gpi-anchored cell wall beta- -endoglucanase protein [Eutypa lata UCREL1] | 56.3 | 1.0E-161 |
| DIP_10005598 | hypothetical protein FPSE_00922 [Fusarium pseudograminearum CS3096] | 39.7 | 0.0 |
| DIP_10005599 | putative family protein [Eutypa lata UCREL1] | 70.6 | 1.0E-152 |
| DIP_10005600 | putative ribosomal protein [Eutypa lata UCREL1] | 52.9 | 3.0E-55 |
| DIP_10005601 | putative mt-a70 family protein [Eutypa lata UCREL1] | 34.7 | 2.0E-51 |
| DIP_10005602 | hypothetical protein TRIVIDRAFT_28473 [Trichoderma virens Gv29-8] | 50.2 | 0.0 |
| DIP_10005603 | putative multispanning membrane protein [Togninia minima UCRPA7] | 72.4 | 0.0 |
| DIP_10005604 | putative pre-60s factor protein [Eutypa lata UCREL1] | 33 | 9.0E-48 |
| DIP_10005605 | hypothetical protein [Podospora anserina S mat+] | 51.8 | 8.0E-80 |
| DIP_10005606 | hypothetical protein PFICI_04597 [Pestalotiopsis fici W106-1] | 73.4 | 2.0E-175 |
| DIP_10005607 | hypothetical protein MGYG_08855 [Microsporum gypseum CBS 118893] | 31.1 | 2.0E-07 |
| DIP_10005608 | hypothetical protein FOTG_13871 [Fusarium oxysporum f. sp. vasinfectum 25433] | 36.6 | 4.0E-24 |
| DIP_10005609 | hypothetical protein PFICI_05615 [Pestalotiopsis fici W106-1] | 49.4 | 9.0E-102 |
| DIP_10005610 | putative rab gdp-dissociation inhibitor [Diaporthe ampelina] | 83.6 | 0.0 |
| DIP_10005611 | putative 50s ribosomal protein l4 protein [Eutypa lata UCREL1] | 59.3 | 2.0E-82 |
| DIP_10005612 | NA | | |
| DIP_10005613 | putative c2h2 transcription factor protein [Eutypa lata UCREL1] | 82.2 | 2.0E-170 |
| DIP_10005614 | hypothetical protein PFICI_05616 [Pestalotiopsis fici W106-1] | 62.7 | 0.0 |
| DIP_10005615 | putative sodium p-type atpase protein [Eutypa lata UCREL1] | 70.6 | 0.0 |
| DIP_10005616 | hypothetical protein PFICI_00681 [Pestalotiopsis fici W106-1] | 67.4 | 2.0E-110 |
| DIP_10005617 | NA | | |
| DIP_10005618 | NA | | |
| DIP_10005619 | NA | | |
| DIP_10005620 | NA | | |
| DIP_10005621 | hypothetical protein PFICI_02022 [Pestalotiopsis fici W106-1] | 62.9 | 7.0E-109 |
| DIP_10005622 | NA | | |
| DIP_10005623 | putative anaphase-promoting complex subunit protein [Eutypa lata UCREL1] | 64.6 | 0.0 |
| DIP_10005624 | NA | | |
| DIP_10005625 | putative arp2 3 complex 34 kda subunit protein [Eutypa lata UCREL1] | 89.7 | 0.0 |
| DIP_10005626 | hypothetical protein PFICI_02013 [Pestalotiopsis fici W106-1] | 77.8 | 0.0 |
| DIP_10005627 | hypothetical protein PFICI_01967 [Pestalotiopsis fici W106-1] | 49 | 1.0E-133 |
| DIP_10005628 | Zinc finger, RING/FYVE/PHD-type, partial [Metarhizium brunneum ARSEF 3297] | 32.3 | 5.0E-07 |
| DIP_10005629 | putative cystathionine beta-lyase protein [Eutypa lata UCREL1] | 84 | 0.0 |
| DIP_10005630 | hypothetical protein PFICI_05191 [Pestalotiopsis fici W106-1] | 80 | 0.0 |
| DIP_10005631 | putative chromosome segregation protein [Eutypa lata UCREL1] | 48.1 | 2.0E-109 |
| DIP_10005632 | hypothetical protein UCREL1_7914 [Eutypa lata UCREL1] | 64.1 | 0.0 |
| DIP_10005633 | hypothetical protein PFICI_01324 [Pestalotiopsis fici W106-1] | 42.6 | 3.0E-99 |
| DIP_10005634 | NA | | |
| DIP_10005635 | putative transcription initiation factor iif protein [Eutypa lata UCREL1] | 62.7 | 3.0E-163 |
| DIP_10005636 | hypothetical protein PFICI_01322 [Pestalotiopsis fici W106-1] | 45.9 | 5.0E-117 |
| DIP_10005637 | NA | | |
| DIP_10005638 | hypothetical protein PFICI_01319 [Pestalotiopsis fici W106-1] | 48.3 | 9.0E-62 |
| DIP_10005639 | predicted protein [Aspergillus terreus NIH2624] | 80 | 1.0E-51 |
| DIP_10005640 | NA | | |
| DIP_10005641 | putative bzip transcription factor protein [Eutypa lata UCREL1] | 71.1 | 0.0 |
| DIP_10005642 | hypothetical protein PFICI_01317 [Pestalotiopsis fici W106-1] | 71.7 | 0.0 |
| DIP_10005643 | putative atp synthase subunit 4 protein [Eutypa lata UCREL1] | 83.1 | 3.0E-148 |
| DIP_10005644 | hypothetical protein PFICI_01315 [Pestalotiopsis fici W106-1] | 35.2 | 1.0E-46 |
| DIP_10005645 | hypothetical protein NECHADRAFT_75698 [Nectria haematococca mpVI 77-13-4] | 32.9 | 6.0E-09 |
| DIP_10005646 | NA | | |
| DIP_10005647 | NA | | |
| DIP_10005648 | NA | | |
| DIP_10005649 | hypothetical protein PFICI_00112 [Pestalotiopsis fici W106-1] | 45.4 | 2.0E-148 |
| DIP_10005650 | hypothetical protein PFICI_00111 [Pestalotiopsis fici W106-1] | 40.1 | 1.0E-60 |
| DIP_10005651 | hypothetical protein PFICI_00106 [Pestalotiopsis fici W106-1] | 59.6 | 6.0E-160 |
| DIP_10005652 | chlorogenic acid esterase precursor [Colletotrichum gloeosporioides Nara gc5] | 47.4 | 3.0E-157 |
| DIP_10005653 | hypothetical protein PFICI_05623 [Pestalotiopsis fici W106-1] | 66.1 | 0.0 |
| DIP_10005654 | putative dna-binding protein [Eutypa lata UCREL1] | 34.7 | 4.0E-77 |
| DIP_10005655 | putative swi snf complex protein [Eutypa lata UCREL1] | 78.6 | 0.0 |
| DIP_10005656 | putative electron transfer flavoprotein subunit alpha protein [Eutypa lata UCREL1] | 82.6 | 0.0 |
| DIP_10005657 | putative nadph:adrenodoxin oxidoreductase protein [Eutypa lata UCREL1] | 59.5 | 0.0 |
| DIP_10005658 | hypothetical protein UCREL1_3331 [Eutypa lata UCREL1] | 45.5 | 4.0E-114 |
| DIP_10005659 | hypothetical protein [Epichloe amarillans] | 56.5 | 1.0E-120 |
| DIP_10005660 | Lon protease [Pestalotiopsis fici W106-1] | 80.6 | 0.0 |
| DIP_10005661 | hypothetical protein PFICI_05199 [Pestalotiopsis fici W106-1] | 62.8 | 0.0 |
| DIP_10005662 | putative acyl- desaturase protein [Eutypa lata UCREL1] | 80 | 0.0 |
| DIP_10005663 | putative flavonol synthase protein [Eutypa lata UCREL1] | 65.5 | 0.0 |
| DIP_10005664 | hypothetical protein PFICI_13681 [Pestalotiopsis fici W106-1] | 66.1 | 0.0 |
| DIP_10005665 | hypothetical protein PFICI_05185 [Pestalotiopsis fici W106-1] | 49.8 | 6.0E-153 |
| DIP_10005666 | hypothetical protein NEUTE1DRAFT_121594 [Neurospora tetrasperma FGSC 2508] | 39.3 | 0.0 |
| DIP_10005667 | hypothetical protein [Podospora anserina S mat+] | 95.5 | 3.0E-174 |
| DIP_10005668 | putative spc97 spc98 family protein [Eutypa lata UCREL1] | 79.9 | 0.0 |
| DIP_10005669 | hypothetical protein PFICI_10635 [Pestalotiopsis fici W106-1] | 77.1 | 2.0E-46 |
| DIP_10005670 | Electron transfer flavoprotein subunit beta [Pestalotiopsis fici W106-1] | 88 | 5.0E-162 |
| DIP_10005671 | putative carbamoyl-phosphate synthase protein [Eutypa lata UCREL1] | 87.4 | 0.0 |
| DIP_10005672 | hypothetical protein FVEG_06980 [Fusarium verticillioides 7600] | 44.4 | 5.0E-89 |
| DIP_10005673 | NA | | |
| DIP_10005674 | hypothetical protein PFICI_05658 [Pestalotiopsis fici W106-1] | 50 | 7.0E-108 |
| DIP_10005675 | related to f-box/wd-repeat protein pof10 [Fusarium fujikuroi IMI 58289] | 50.6 | 0.0 |
| DIP_10005676 | putative c6 finger domain protein [Eutypa lata UCREL1] | 52.1 | 0.0 |
| DIP_10005677 | putative mob1 family protein [Eutypa lata UCREL1] | 68.9 | 0.0 |
| DIP_10005678 | hypothetical protein PFICI_04663 [Pestalotiopsis fici W106-1] | 71.6 | 0.0 |
| DIP_10005679 | hypothetical protein PFICI_04662 [Pestalotiopsis fici W106-1] | 63.9 | 8.0E-178 |
| DIP_10005680 | NA | | |
| DIP_10005681 | hypothetical protein GGTG_03389 [Gaeumannomyces graminis var. tritici R3-111a-1] | 42.3 | 5.0E-76 |
| DIP_10005682 | NA | | |
| DIP_10005683 | hypothetical protein Z517_00226 [Fonsecaea pedrosoi CBS 271.37] | 56 | 2.0E-82 |
| DIP_10005684 | putative abhydrolase domain-containing protein 12 protein [Eutypa lata UCREL1] | 60.9 | 3.0E-174 |
| DIP_10005685 | hypothetical protein PFICI_04984 [Pestalotiopsis fici W106-1] | 62.9 | 0.0 |
| DIP_10005686 | SJCHGC07487 protein [Schistosoma japonicum] | 33 | 9.0E-12 |
| DIP_10005687 | cation transporter [Colletotrichum fioriniae PJ7] | 53.1 | 0.0 |
| DIP_10005688 | putative barwin-like endoglucanase protein [Eutypa lata UCREL1] | 58.7 | 2.0E-39 |
| DIP_10005689 | putative mitochondrial outer membrane protein iml2 protein [Eutypa lata UCREL1] | 61.2 | 0.0 |
| DIP_10005690 | hypothetical protein S7711_01043 [Stachybotrys chartarum IBT 7711] | 68.1 | 2.0E-30 |
| DIP_10005691 | hypothetical protein PFICI_15383 [Pestalotiopsis fici W106-1] | 78.9 | 0.0 |
| DIP_10005692 | putative transcriptional corepressor of histone genes protein [Eutypa lata UCREL1] | 60.4 | 0.0 |
| DIP_10005693 | NA | | |
| DIP_10005694 | hypothetical protein PV07_08453 [Cladophialophora immunda] | 32.3 | 7.0E-35 |
| DIP_10005695 | putative atpase rrpa3 protein [Eutypa lata UCREL1] | 81.2 | 0.0 |
| DIP_10005696 | putative metallothionein-i gene transcription activator protein [Eutypa lata UCREL1] | 74.4 | 7.0E-37 |
| DIP_10005697 | hypothetical protein S40285_02378 [Stachybotrys chlorohalonata IBT 40285] | 74 | 2.0E-77 |
| DIP_10005698 | putative glycerol dehydrogenase protein [Eutypa lata UCREL1] | 82.6 | 0.0 |
| DIP_10005699 | putative atp-dependent rrna helicase rrp3 protein [Eutypa lata UCREL1] | 69.9 | 0.0 |
| DIP_10005700 | putative ribosome biogenesis protein ssf1 protein [Eutypa lata UCREL1] | 70.3 | 0.0 |
| DIP_10005701 | putative major facilitator superfamily domain containing protein 5 protein [Eutypa lata UCREL1] | 39.4 | 5.0E-101 |
| DIP_10005702 | hypothetical protein PFICI_15354 [Pestalotiopsis fici W106-1] | 82.6 | 5.0E-173 |
| DIP_10005703 | hypothetical protein PFICI_15355 [Pestalotiopsis fici W106-1] | 56.6 | 1.0E-110 |
| DIP_10005704 | microsomal signal peptidase 18 kDa subunit [Verticillium dahliae VdLs.17] | 91.9 | 2.0E-114 |
| DIP_10005705 | putative cell cycle control protein [Eutypa lata UCREL1] | 67 | 0.0 |
| DIP_10005706 | putative brf1-like tbp-binding domain-containing protein [Eutypa lata UCREL1] | 59.9 | 0.0 |
| DIP_10005707 | ubiquitin conjugating enzyme [Trichophyton tonsurans CBS 112818] | 51 | 6.0E-55 |
| DIP_10005708 | putative gtpase-activating protein gyp3 protein [Eutypa lata UCREL1] | 66.4 | 0.0 |
| DIP_10005709 | NA | | |
| DIP_10005710 | glutathione-dependent formaldehyde-activating enzyme [Colletotrichum higginsianum] | 36.4 | 3.0E-53 |
| DIP_10005711 | NA | | |
| DIP_10005712 | hypothetical protein MBM_08588 [Marssonina brunnea f. sp. 'multigermtubi' MB_m1] | 32.4 | 8.0E-16 |
| DIP_10005713 | Cytosolic Fe-S cluster assembly factor nar-1 [Pestalotiopsis fici W106-1] | 65.8 | 0.0 |
| DIP_10005714 | hypothetical protein THITE_2121992 [Thielavia terrestris NRRL 8126] | 53.1 | 2.0E-48 |
| DIP_10005715 | hypothetical protein UCREL1_1974 [Eutypa lata UCREL1] | 44.3 | 2.0E-13 |
| DIP_10005716 | NA | | |
| DIP_10005717 | hypothetical protein MGG_13985 [Magnaporthe oryzae 70-15] | 43.2 | 1.0E-34 |
| DIP_10005718 | NA | | |
| DIP_10005719 | hypothetical protein PFICI_00543 [Pestalotiopsis fici W106-1] | 67.5 | 0.0 |
| DIP_10005720 | NA | | |
| DIP_10005721 | hypothetical protein PFICI_00545 [Pestalotiopsis fici W106-1] | 81.9 | 6.0E-61 |
| DIP_10005722 | putative glucose-inducible sam-dependent methyltransferase protein [Eutypa lata UCREL1] | 47.2 | 1.0E-87 |
| DIP_10005723 | hypothetical protein UCREL1_3224 [Eutypa lata UCREL1] | 64.1 | 8.0E-173 |
| DIP_10005724 | hypothetical protein PFICI_05638 [Pestalotiopsis fici W106-1] | 52.6 | 0.0 |
| DIP_10005725 | hypothetical protein PFICI_05637 [Pestalotiopsis fici W106-1] | 65.7 | 0.0 |
| DIP_10005726 | NA | | |
| DIP_10005727 | hypothetical protein PFICI_05617 [Pestalotiopsis fici W106-1] | 54.1 | 0.0 |
| DIP_10005728 | hypothetical protein PFICI_05196 [Pestalotiopsis fici W106-1] | 75.8 | 2.0E-158 |
| DIP_10005729 | putative 6-phosphogluconate dehydrogenase protein [Eutypa lata UCREL1] | 56.2 | 3.0E-119 |
| DIP_10005730 | putative vacuolar atp synthase subunit protein [Eutypa lata UCREL1] | 81.3 | 1.0E-42 |
| DIP_10005731 | hypothetical protein THITE_66503 [Thielavia terrestris NRRL 8126] | 57.5 | 3.0E-177 |
| DIP_10005732 | hypothetical protein PFICI_15370 [Pestalotiopsis fici W106-1] | 70.5 | 4.0E-155 |
| DIP_10005733 | Hydrolase [Scedosporium apiospermum] | 39.1 | 2.0E-59 |
| DIP_10005734 | putative hydrolase protein [Eutypa lata UCREL1] | 72.7 | 3.0E-78 |
| DIP_10005735 | hypothetical protein UCREL1_1326 [Eutypa lata UCREL1] | 45.9 | 6.0E-80 |
| DIP_10005736 | putative inosine-uridine preferring nucleoside hydrolase protein [Eutypa lata UCREL1] | 64.9 | 0.0 |
| DIP_10005737 | putative lsm domain-containing protein [Eutypa lata UCREL1] | 77.6 | 5.0E-50 |
| DIP_10005738 | conserved hypothetical protein [Arthroderma otae CBS 113480] | 31.2 | 1.0E-17 |
| DIP_10005739 | hypothetical protein PFICI_04487 [Pestalotiopsis fici W106-1] | 68.7 | 0.0 |
| DIP_10005740 | putative protein kinase c protein [Neofusicoccum parvum UCRNP2] | 65.1 | 0.0 |
| DIP_10005741 | hypothetical protein PFICI_04260 [Pestalotiopsis fici W106-1] | 53.3 | 1.0E-93 |
| DIP_10005742 | putative phosphoserine phosphatase protein [Eutypa lata UCREL1] | 71 | 5.0E-131 |
| DIP_10005743 | putative methyltransferase domain-containing protein [Colletotrichum sublineola] | 36.4 | 2.0E-47 |
| DIP_10005744 | putative d-amino-acid oxidase protein [Eutypa lata UCREL1] | 62.8 | 2.0E-167 |
| DIP_10005745 | putative glycosyltransferase family 58 protein [Eutypa lata UCREL1] | 67.3 | 0.0 |
| DIP_10005746 | hypothetical protein PFICI_04254 [Pestalotiopsis fici W106-1] | 69.8 | 6.0E-51 |
| DIP_10005747 | putative mfs efflux protein [Eutypa lata UCREL1] | 61.1 | 0.0 |
| DIP_10005748 | hypothetical protein HMPREF1624_07438 [Sporothrix schenckii ATCC 58251] | 44.7 | 3.0E-81 |
| DIP_10005749 | hypothetical protein PFICI_04250 [Pestalotiopsis fici W106-1] | 41.6 | 3.0E-73 |
| DIP_10005750 | hypothetical protein PFICI_04246 [Pestalotiopsis fici W106-1] | 57.1 | 4.0E-131 |
| DIP_10005751 | AP-2 complex subunit sigma [Neurospora crassa OR74A] | 96.5 | 6.0E-102 |
| DIP_10005752 | NA | | |
| DIP_10005753 | hypothetical protein PFICI_04243 [Pestalotiopsis fici W106-1] | 80.7 | 2.0E-96 |
| DIP_10005754 | hypothetical protein PFICI_04242 [Pestalotiopsis fici W106-1] | 48.1 | 0.0 |
| DIP_10005755 | putative spt3-like protein [Eutypa lata UCREL1] | 83.5 | 1.0E-177 |
| DIP_10005756 | hypothetical protein PFICI_04240 [Pestalotiopsis fici W106-1] | 54.8 | 8.0E-126 |
| DIP_10005757 | hypothetical protein PFICI_04241 [Pestalotiopsis fici W106-1] | 49.4 | 2.0E-117 |
| DIP_10005758 | putative pci domain-containing protein [Eutypa lata UCREL1] | 56.1 | 0.0 |
| DIP_10005759 | Acetyl-CoA acetyltransferase [Pestalotiopsis fici W106-1] | 83.8 | 0.0 |
| DIP_10005760 | putative ubiquitin-conjugating enzyme e2 2 protein [Eutypa lata UCREL1] | 98 | 2.0E-110 |
| DIP_10005761 | putative kinesin heavy chain protein [Eutypa lata UCREL1] | 83.9 | 0.0 |
| DIP_10005762 | hypothetical protein CTHT_0034360 [Chaetomium thermophilum var. thermophilum DSM 1495] | 45 | 1.0E-123 |
| DIP_10005763 | putative high-affinity glucose transporter rgt2 protein [Eutypa lata UCREL1] | 80.5 | 0.0 |
| DIP_10005764 | hypothetical protein PFICI_04228 [Pestalotiopsis fici W106-1] | 65.8 | 0.0 |
| DIP_10005765 | putative diacylglycerol kinase catalytic domain-containing protein [Eutypa lata UCREL1] | 53.2 | 7.0E-147 |
| DIP_10005766 | putative -dimethylguanosine trna methyltransferase protein [Eutypa lata UCREL1] | 59.7 | 0.0 |
| DIP_10005767 | putative c6 transcription factor protein [Eutypa lata UCREL1] | 46.6 | 2.0E-44 |
| DIP_10005768 | hypothetical protein PFICI_04436 [Pestalotiopsis fici W106-1] | 72.7 | 0.0 |
| DIP_10005769 | hypothetical protein PFICI_04437 [Pestalotiopsis fici W106-1] | 63.4 | 9.0E-118 |
| DIP_10005770 | hypothetical protein UCREL1_6890 [Eutypa lata UCREL1] | 93.6 | 1.0E-59 |
| DIP_10005771 | hypothetical protein PFICI_04439 [Pestalotiopsis fici W106-1] | 73.5 | 0.0 |
| DIP_10005772 | Histone-lysine N-methyltransferase, H3 lysine-9 specific dim-5 [Pestalotiopsis fici W106-1] | 60.6 | 6.0E-143 |
| DIP_10005773 | putative ubiquitin domain protein [Eutypa lata UCREL1] | 46.7 | 2.0E-46 |
| DIP_10005774 | putative sphingosine kinase protein [Eutypa lata UCREL1] | 55.5 | 0.0 |
| DIP_10005775 | ribosomal RNA-processing protein [Verticillium dahliae VdLs.17] | 30.8 | 1.0E-46 |
| DIP_10005776 | putative carnitinyl- dehydratase protein [Togninia minima UCRPA7] | 54.9 | 3.0E-113 |
| DIP_10005777 | putative sh3 domain protein [Eutypa lata UCREL1] | 45.1 | 1.0E-83 |
| DIP_10005778 | putative snorap assembly factor protein [Eutypa lata UCREL1] | 59.3 | 2.0E-88 |
| DIP_10005779 | hypothetical protein PFICI_04708 [Pestalotiopsis fici W106-1] | 64.3 | 0.0 |
| DIP_10005780 | putative mfs transporter protein [Eutypa lata UCREL1] | 62.4 | 0.0 |
| DIP_10005781 | hypothetical protein PFICI_04297 [Pestalotiopsis fici W106-1] | 57.3 | 1.0E-147 |
| DIP_10005782 | putative dhh family protein [Eutypa lata UCREL1] | 56.9 | 2.0E-169 |
| DIP_10005783 | putative dienelactone hydrolase family protein [Eutypa lata UCREL1] | 34.2 | 2.0E-42 |
| DIP_10005784 | hypothetical protein PFICI_04419 [Pestalotiopsis fici W106-1] | 55.1 | 1.0E-56 |
| DIP_10005785 | hypothetical protein PFICI_04420 [Pestalotiopsis fici W106-1] | 55.5 | 5.0E-130 |
| DIP_10005786 | related to calmodulin [Claviceps purpurea 20.1] | 87.8 | 3.0E-93 |
| DIP_10005787 | hypothetical protein PFICI_04422 [Pestalotiopsis fici W106-1] | 77.8 | 0.0 |
| DIP_10005788 | NA | | |
| DIP_10005789 | Tetrapyrrole biosynthesis, glutamate-1-semialdehyde aminotransferase [Penicillium italicum] | 34.9 | 2.0E-26 |
| DIP_10005790 | putative cel3e secreted beta-glucosidase protein [Eutypa lata UCREL1] | 62.7 | 0.0 |
| DIP_10005791 | Trimeric LpxA-like protein, partial [Metarhizium majus ARSEF 297] | 72.8 | 2.0E-121 |
| DIP_10005792 | NA | | |
| DIP_10005793 | NA | | |
| DIP_10005794 | hypothetical protein HIM_03404 [Hirsutella minnesotensis 3608] | 40.7 | 1.0E-87 |
| DIP_10005795 | hypothetical protein O988_07689 [Pseudogymnoascus pannorum VKM F-3808] | 55 | 3.0E-99 |
| DIP_10005796 | putative mitochondrial import protein mmp37 protein [Eutypa lata UCREL1] | 75.6 | 0.0 |
| DIP_10005797 | putative spindle assembly checkpoint component mad1 protein [Eutypa lata UCREL1] | 69.7 | 0.0 |
| DIP_10005798 | putative saff domain-containing protein [Eutypa lata UCREL1] | 63 | 0.0 |
| DIP_10005799 | hypothetical protein SMAC_08955 [Sordaria macrospora k-hell] | 59.4 | 4.0E-161 |
| DIP_10005800 | putative nad dependent epimerase protein [Eutypa lata UCREL1] | 49.8 | 2.0E-97 |
| DIP_10005801 | putative short chain dehydrogenase protein [Eutypa lata UCREL1] | 67.2 | 2.0E-164 |
| DIP_10005802 | hypothetical protein CFIO01_01290 [Colletotrichum fioriniae PJ7] | 48.5 | 0.0 |
| DIP_10005803 | NA | | |
| DIP_10005804 | hypothetical protein A1O1_04057 [Capronia coronata CBS 617.96] | 41.9 | 6.0E-72 |
| DIP_10005805 | NA | | |
| DIP_10005806 | hypothetical protein PFICI_09891 [Pestalotiopsis fici W106-1] | 62.3 | 0.0 |
| DIP_10005807 | putative palmitoyl-protein thioesterase 1 protein [Eutypa lata UCREL1] | 56.8 | 2.0E-124 |
| DIP_10005808 | putative electron transfer flavoprotein alpha-subunit protein [Eutypa lata UCREL1] | 63.9 | 3.0E-166 |
| DIP_10005809 | putative group protein [Eutypa lata UCREL1] | 46.8 | 0.0 |
| DIP_10005810 | putative transcription factor tos4 protein [Togninia minima UCRPA7] | 68.1 | 0.0 |
| DIP_10005811 | hypothetical protein UCREL1_6804 [Eutypa lata UCREL1] | 37.6 | 8.0E-07 |
| DIP_10005812 | hypothetical protein FG05_02389 [Fusarium graminearum] | 43.6 | 1.0E-75 |
| DIP_10005813 | NA | | |
| DIP_10005814 | NA | | |
| DIP_10005815 | hypothetical protein CFIO01_09000 [Colletotrichum fioriniae PJ7] | 50.2 | 0.0 |
| DIP_10005816 | calcium/proton exchanger [Colletotrichum fioriniae PJ7] | 57.3 | 0.0 |
| DIP_10005817 | NA | | |
| DIP_10005818 | NA | | |
| DIP_10005819 | hypothetical protein MGG_10456 [Magnaporthe oryzae 70-15] | 41.2 | 2.0E-23 |
| DIP_10005820 | NA | | |
| DIP_10005821 | predicted protein [Aspergillus terreus NIH2624] | 37.4 | 1.0E-32 |
| DIP_10005822 | hypothetical protein UCREL1_4645 [Eutypa lata UCREL1] | 49.4 | 9.0E-32 |
| DIP_10005823 | NA | | |
| DIP_10005824 | short chain dehydrogenase/reductase family oxidoreductase, putative [Byssochlamys spectabilis No. 5] | 43.2 | 2.0E-83 |
| DIP_10005825 | NA | | |
| DIP_10005826 | hypothetical protein S7711_06491 [Stachybotrys chartarum IBT 7711] | 30.8 | 4.0E-53 |
| DIP_10005827 | hypothetical protein H072_7639 [Dactylellina haptotyla CBS 200.50] | 57.1 | 0.0 |
| DIP_10005828 | hypothetical protein PFICI_01366 [Pestalotiopsis fici W106-1] | 54.3 | 0.0 |
| DIP_10005829 | putative transmembrane protein 32 protein [Eutypa lata UCREL1] | 76.3 | 4.0E-79 |
| DIP_10005830 | putative haloacid dehalogenase-like hydrolase protein [Eutypa lata UCREL1] | 85 | 4.0E-154 |
| DIP_10005831 | hypothetical protein UCREL1_1668 [Eutypa lata UCREL1] | 45.6 | 2.0E-34 |
| DIP_10005832 | putative methylthioribulose-1-phosphate dehydratase protein [Eutypa lata UCREL1] | 81.1 | 4.0E-161 |
| DIP_10005833 | putative calnexin precursor protein [Eutypa lata UCREL1] | 72.1 | 0.0 |
| DIP_10005834 | NA | | |
| DIP_10005835 | NA | | |
| DIP_10005836 | RNA polymerase Rpb4 family protein [Marssonina brunnea f. sp. 'multigermtubi' MB_m1] | 76.2 | 8.0E-75 |
| DIP_10005837 | putative sybindin-like family protein [Eutypa lata UCREL1] | 68.2 | 3.0E-83 |
| DIP_10005838 | hypothetical protein PFICI_00580 [Pestalotiopsis fici W106-1] | 77.4 | 2.0E-141 |
| DIP_10005839 | hypothetical protein PFICI_00581 [Pestalotiopsis fici W106-1] | 51.6 | 6.0E-88 |
| DIP_10005840 | Translation machinery-associated protein 22 [Pestalotiopsis fici W106-1] | 87.1 | 9.0E-108 |
| DIP_10005841 | hypothetical protein PFICI_01215 [Pestalotiopsis fici W106-1] | 55 | 0.0 |
| DIP_10005842 | NA | | |
| DIP_10005843 | hypothetical protein PFICI_01362 [Pestalotiopsis fici W106-1] | 56.2 | 0.0 |
| DIP_10005844 | NA | | |
| DIP_10005845 | NA | | |
| DIP_10005846 | hypothetical protein PFICI_01361 [Pestalotiopsis fici W106-1] | 55.2 | 1.0E-162 |
| DIP_10005847 | putative adp-ribosylation factor protein [Eutypa lata UCREL1] | 47.4 | 3.0E-141 |
| DIP_10005848 | hypothetical protein MYCTH_45081 [Myceliophthora thermophila ATCC 42464] | 32.8 | 1.0E-08 |
| DIP_10005849 | hypothetical protein PFICI_09824 [Pestalotiopsis fici W106-1] | 58.2 | 0.0 |
| DIP_10005850 | hypothetical protein PFICI_04819 [Pestalotiopsis fici W106-1] | 46.4 | 9.0E-63 |
| DIP_10005851 | NA | | |
| DIP_10005852 | hypothetical protein NECHADRAFT_33721 [Nectria haematococca mpVI 77-13-4] | 35.8 | 2.0E-76 |
| DIP_10005853 | putative methylenetetrahydrofolate reductase 1 protein [Eutypa lata UCREL1] | 81.2 | 0.0 |

| | | | |
|---|---|---|---|
| DIP_10005854 | UDP-glucuronosyl/UDP-glucosyltransferase [Metarhizium guizhouense ARSEF 977] | 47.1 | 6.0E-143 |
| DIP_10005855 | putative formyltetrahydrofolate deformylase protein [Eutypa lata UCREL1] | 78.8 | 4.0E-172 |
| DIP_10005856 | hypothetical protein UCREL1_8107 [Eutypa lata UCREL1] | 66.1 | 5.0E-104 |
| DIP_10005857 | putative tpr domain-containing protein [Eutypa lata UCREL1] | 45.8 | 1.0E-109 |
| DIP_10005858 | hypothetical protein PFICI_00586 [Pestalotiopsis fici W106-1] | 60 | 5.0E-81 |
| DIP_10005859 | putative pumilio domain-containing protein [Eutypa lata UCREL1] | 70.6 | 0.0 |
| DIP_10005860 | NA | | |
| DIP_10005861 | hypothetical protein PFICI_04818 [Pestalotiopsis fici W106-1] | 56.5 | 0.0 |
| DIP_10005862 | molybdopterin binding domain-containing protein [Neurospora crassa OR74A] | 70 | 7.0E-145 |
| DIP_10005863 | putative gmf family protein [Eutypa lata UCREL1] | 89.2 | 4.0E-87 |
| DIP_10005864 | hypothetical protein MAA_06383 [Metarhizium robertsii ARSEF 23] | 83.4 | 0.0 |
| DIP_10005865 | hypothetical protein [Podospora anserina S mat+] | 63.1 | 0.0 |
| DIP_10005866 | NA | | |
| DIP_10005867 | NA | | |
| DIP_10005868 | Ras-like protein [Colletotrichum higginsianum] | 88.3 | 5.0E-132 |
| DIP_10005869 | NA | | |
| DIP_10005870 | putative amino acid transporter protein [Eutypa lata UCREL1] | 73.8 | 0.0 |
| DIP_10005871 | S-adenosyl-L-methionine-dependent methyltransferase [Glarea lozoyensis ATCC 20868] | 51.8 | 1.0E-85 |
| DIP_10005872 | phenylalanine ammonia-lyase [Beauveria bassiana ARSEF 2860] | 68.2 | 0.0 |
| DIP_10005873 | hypothetical protein S40285_03904 [Stachybotrys chlorohalonata IBT 40285] | 66 | 0.0 |
| DIP_10005874 | hypothetical protein NECHADRAFT_90636 [Nectria haematococca mpVI 77-13-4] | 42.8 | 9.0E-177 |
| DIP_10005875 | putative transcription factor-like protein [Acremonium chrysogenum ATCC 11550] | 40.6 | 8.0E-152 |
| DIP_10005876 | putative endosomal integral membrane protein [Eutypa lata UCREL1] | 84.8 | 0.0 |
| DIP_10005877 | hypothetical protein PFICI_04211 [Pestalotiopsis fici W106-1] | 60.6 | 2.0E-146 |
| DIP_10005878 | hypothetical protein ACRE_036200 [Acremonium chrysogenum ATCC 11550] | 38.3 | 2.0E-59 |
| DIP_10005879 | hypothetical protein MYCFIDRAFT_134279 [Pseudocercospora fijiensis CIRAD86] | 53.6 | 2.0E-147 |
| DIP_10005880 | hypothetical protein NECHADRAFT_75198 [Nectria haematococca mpVI 77-13-4] | 34.1 | 7.0E-22 |
| DIP_10005881 | hypothetical protein CFIO01_11292 [Colletotrichum fioriniae PJ7] | 55.3 | 6.0E-170 |
| DIP_10005882 | hypothetical protein V492_07035, partial [Pseudogymnoascus pannorum VKM F-4246] | 67.3 | 4.0E-74 |
| DIP_10005883 | NA | | |
| DIP_10005884 | NA | | |
| DIP_10005885 | hypothetical protein HMPREF1624_05601 [Sporothrix schenckii ATCC 58251] | 53.9 | 3.0E-84 |
| DIP_10005886 | putative ring finger domain-containing protein [Togninia minima UCRPA7] | 42.5 | 5.0E-70 |
| DIP_10005887 | putative cytochrome protein [Eutypa lata UCREL1] | 31.5 | 3.0E-67 |
| DIP_10005888 | Interferon-induced GTP-binding protein Mx1 [Talaromyces marneffei PM1] | 41.4 | 2.0E-137 |
| DIP_10005889 | hypothetical protein PFICI_01326 [Pestalotiopsis fici W106-1] | 60.4 | 0.0 |
| DIP_10005890 | hypothetical protein PFICI_04793 [Pestalotiopsis fici W106-1] | 85.8 | 0.0 |
| DIP_10005891 | hypothetical protein PFICI_04792 [Pestalotiopsis fici W106-1] | 43.3 | 0.0 |
| DIP_10005892 | hypothetical protein BofuT4_P122820.1 [Botrytis cinerea T4] | 42.5 | 1.0E-70 |
| DIP_10005893 | NA | | |
| DIP_10005894 | hypothetical protein TRIVIDRAFT_137343, partial [Trichoderma virens Gv29-8] | 31 | 6.0E-10 |
| DIP_10005895 | putative secreted protein [Eutypa lata UCREL1] | 40.4 | 2.0E-38 |
| DIP_10005896 | hypothetical protein PFICI_10601 [Pestalotiopsis fici W106-1] | 73.6 | 0.0 |
| DIP_10005897 | hypothetical protein PFICI_10602 [Pestalotiopsis fici W106-1] | 49.8 | 0.0 |
| DIP_10005898 | tartrate dehydrogenase/decarboxylase [Pestalotiopsis fici W106-1] | 90.2 | 0.0 |
| DIP_10005899 | hypothetical protein OIDMADRAFT_204094 [Oidiodendron maius Zn] | 34.7 | 9.0E-73 |
| DIP_10005900 | glutathionylspermidine synthase [Colletotrichum fioriniae PJ7] | 49.4 | 1.0E-162 |
| DIP_10005901 | NA | | |
| DIP_10005902 | hypothetical protein CHGG_04337 [Chaetomium globosum CBS 148.51] | 40.4 | 8.0E-72 |
| DIP_10005903 | putative ribosomal protein s21 protein [Eutypa lata UCREL1] | 47.2 | 1.0E-27 |
| DIP_10005904 | putative rna-binding protein rbm25 protein [Eutypa lata UCREL1] | 75.9 | 0.0 |
| DIP_10005905 | putative zn 2cys6 transcription factor protein [Eutypa lata UCREL1] | 73.4 | 3.0E-133 |
| DIP_10005906 | NA | | |
| DIP_10005907 | NA | | |
| DIP_10005908 | NA | | |
| DIP_10005909 | hypothetical protein PFICI_04444 [Pestalotiopsis fici W106-1] | 70.5 | 0.0 |
| DIP_10005910 | hypothetical protein PFICI_04443 [Pestalotiopsis fici W106-1] | 65 | 0.0 |
| DIP_10005911 | hypothetical protein S40285_06944 [Stachybotrys chlorohalonata IBT 40285] | 87.8 | 0.0 |
| DIP_10005912 | putative sodium nucleoside cotransporter 1 protein [Eutypa lata UCREL1] | 74.9 | 0.0 |
| DIP_10005913 | NA | | |
| DIP_10005914 | NA | | |
| DIP_10005915 | NA | | |
| DIP_10005916 | putative glycoside hydrolase family 16 protein [Neofusicoccum parvum UCRNP2] | 64.7 | 2.0E-116 |
| DIP_10005917 | NA | | |
| DIP_10005918 | putative mitochondrial carrier protein [Eutypa lata UCREL1] | 69 | 3.0E-165 |
| DIP_10005919 | hypothetical protein PFICI_15377 [Pestalotiopsis fici W106-1] | 70.1 | 4.0E-99 |
| DIP_10005920 | putative eukaryotic translation initiation factor 3 subunit 3 [Diaporthe ampelina] | 82.1 | 0.0 |
| DIP_10005921 | putative adenosylhomocysteinase protein [Eutypa lata UCREL1] | 90.9 | 0.0 |
| DIP_10005922 | hypothetical protein BGT96224_5132 [Blumeria graminis f. sp. tritici 96224] | 39.8 | 1.0E-104 |
| DIP_10005923 | hypothetical protein GGTG_03825 [Gaeumannomyces graminis var. tritici R3-111a-1] | 35.6 | 2.0E-28 |
| DIP_10005924 | UPF0132 domain-containing protein [Neurospora crassa OR74A] | 52.3 | 3.0E-72 |
| DIP_10005925 | hypothetical protein PFICI_00677 [Pestalotiopsis fici W106-1] | 60.3 | 9.0E-143 |
| DIP_10005926 | hypothetical protein PFICI_00675 [Pestalotiopsis fici W106-1] | 68.1 | 0.0 |
| DIP_10005927 | putative trna ligase protein [Eutypa lata UCREL1] | 65.9 | 0.0 |
| DIP_10005928 | Ankyrin repeat-containing protein-like protein [Acremonium chrysogenum ATCC 11550] | 63.2 | 5.0E-64 |
| DIP_10005929 | ribonuclease p protein subunit [Colletotrichum gloeosporioides Nara gc5] | 50.9 | 2.0E-124 |
| DIP_10005930 | putative rna methylase family protein [Eutypa lata UCREL1] | 62.4 | 0.0 |
| DIP_10005931 | hypothetical protein PFICI_00671 [Pestalotiopsis fici W106-1] | 56.3 | 4.0E-137 |
| DIP_10005932 | Mago nashi protein [Verticillium alfalfae VaMs.102] | 89 | 7.0E-103 |
| DIP_10005933 | hypothetical protein OIDMADRAFT_161428 [Oidiodendron maius Zn] | 87.6 | 0.0 |
| DIP_10005934 | hypothetical protein GMDG_06440 [Pseudogymnoascus destructans 20631-21] | 54.7 | 8.0E-13 |
| DIP_10005935 | putative amylase-binding protein [Eutypa lata UCREL1] | 74.8 | 0.0 |
| DIP_10005936 | putative zinc finger protein [Eutypa lata UCREL1] | 68.8 | 4.0E-163 |
| DIP_10005937 | hypothetical protein NECHADRAFT_67889 [Nectria haematococca mpVI 77-13-4] | 37.7 | 5.0E-72 |
| DIP_10005938 | NA | | |
| DIP_10005939 | hypothetical protein PFICI_00531 [Pestalotiopsis fici W106-1] | 74.4 | 2.0E-138 |
| DIP_10005940 | NA | | |
| DIP_10005941 | putative glutaredoxin-c4 precursor protein [Eutypa lata UCREL1] | 60.1 | 7.0E-91 |
| DIP_10005942 | hypothetical protein UCREL1_1338 [Eutypa lata UCREL1] | 34.8 | 4.0E-36 |
| DIP_10005943 | hypothetical protein FVEG_14803 [Fusarium verticillioides 7600] | 40.6 | 7.0E-07 |
| DIP_10005944 | putative secretory protein [Eutypa lata UCREL1] | 48.1 | 9.0E-161 |
| DIP_10005945 | NA | | |
| DIP_10005946 | hypothetical protein PFICI_01903 [Pestalotiopsis fici W106-1] | 34.3 | 6.0E-90 |
| DIP_10005947 | hypothetical protein FOTG_17775 [Fusarium oxysporum f. sp. vasinfectum 25433] | 62.1 | 3.0E-48 |
| DIP_10005948 | putative nucleoside-diphosphate-sugar epimerase protein [Eutypa lata UCREL1] | 73.2 | 6.0E-174 |
| DIP_10005949 | SMP-30/Gluconolaconase/LRE-like region [Penicillium expansum] | 65.5 | 1.0E-178 |
| DIP_10005950 | guanine nucleotide exchange factor vps9 [Colletotrichum orbiculare MAFF 240422] | 69 | 5.0E-164 |
| DIP_10005951 | putative phospholipid-transporting atpase 1 protein [Eutypa lata UCREL1] | 61.5 | 0.0 |
| DIP_10005952 | hypothetical protein PFICI_02191 [Pestalotiopsis fici W106-1] | 42.9 | 1.0E-38 |
| DIP_10005953 | hypothetical protein ACRE_015940 [Acremonium chrysogenum ATCC 11550] | 35.4 | 2.0E-39 |
| DIP_10005954 | hypothetical protein HIM_02962 [Hirsutella minnesotensis 3608] | 54 | 4.0E-46 |
| DIP_10005955 | NA | | |
| DIP_10005956 | hypothetical protein PFICI_04492 [Pestalotiopsis fici W106-1] | 51.8 | 0.0 |
| DIP_10005957 | putative endoplasmic reticulum protein [Eutypa lata UCREL1] | 74.2 | 8.0E-143 |
| DIP_10005958 | hypothetical protein PFICI_05391 [Pestalotiopsis fici W106-1] | 53.5 | 0.0 |
| DIP_10005959 | NA | | |
| DIP_10005960 | hypothetical protein THITE_120596 [Thielavia terrestris NRRL 8126] | 30.6 | 3.0E-22 |
| DIP_10005961 | hypothetical protein PFICI_14406 [Pestalotiopsis fici W106-1] | 37.3 | 6.0E-19 |
| DIP_10005962 | putative s-layer protein [Eutypa lata UCREL1] | 30.4 | 1.0E-22 |
| DIP_10005963 | putative cytochrome p450 protein [Eutypa lata UCREL1] | 49.5 | 0.0 |
| DIP_10005964 | putative proteasome component pre2 protein [Eutypa lata UCREL1] | 93 | 0.0 |
| DIP_10005965 | hypothetical protein PFICI_01329 [Pestalotiopsis fici W106-1] | 49.8 | 9.0E-169 |
| DIP_10005966 | NA | | |
| DIP_10005967 | hypothetical protein Cob_03355 [Colletotrichum orbiculare MAFF 240422] | 49.2 | 1.0E-109 |
| DIP_10005968 | hypothetical protein [Podospora anserina S mat+] | 85.7 | 0.0 |
| DIP_10005969 | hypothetical protein PFICI_01335 [Pestalotiopsis fici W106-1] | 56.8 | 1.0E-117 |
| DIP_10005970 | NA | | |
| DIP_10005971 | NA | | |
| DIP_10005972 | hypothetical protein PFICI_01327 [Pestalotiopsis fici W106-1] | 64 | 0.0 |
| DIP_10005973 | hypothetical protein PFICI_01328 [Pestalotiopsis fici W106-1] | 74.8 | 0.0 |
| DIP_10005974 | putative 1-phosphatidylinositol phosphodiesterase protein [Eutypa lata UCREL1] | 61.3 | 0.0 |
| DIP_10005975 | 6-phosphogluconate dehydrogenase [Colletotrichum orbiculare MAFF 240422] | 34.9 | 4.0E-62 |
| DIP_10005976 | putative rna binding protein [Eutypa lata UCREL1] | 54.1 | 1.0E-69 |
| DIP_10005977 | putative transient receptor potential ion channel protein [Eutypa lata UCREL1] | 61.5 | 0.0 |
| DIP_10005978 | hypothetical protein UCREL1_6717 [Eutypa lata UCREL1] | 84.3 | 4.0E-49 |
| DIP_10005979 | putative superoxide dismutase protein [Eutypa lata UCREL1] | 51.4 | 2.0E-87 |
| DIP_10005980 | C2H2 finger domain-containing protein [Colletotrichum fioriniae PJ7] | 40.4 | 9.0E-152 |
| DIP_10005981 | hypothetical protein UCREL1_1190 [Eutypa lata UCREL1] | 77.9 | 4.0E-71 |
| DIP_10005982 | hypothetical protein PFICI_01352 [Pestalotiopsis fici W106-1] | 30.2 | 2.0E-28 |
| DIP_10005983 | putative short-chain dehydrogenase reductase sdr protein [Eutypa lata UCREL1] | 76.4 | 6.0E-173 |
| DIP_10005984 | putative integral membrane protein [Eutypa lata UCREL1] | 60.3 | 3.0E-127 |
| DIP_10005985 | NA | | |
| DIP_10005986 | uncharacterized protein FFUJ_10083 [Fusarium fujikuroi IMI 58289] | 51.2 | 4.0E-97 |
| DIP_10005987 | F-box domain-containing protein, variant [Magnaporthe oryzae 70-15] | 33.2 | 2.0E-38 |
| DIP_10005988 | putative carbohydrate binding family 6 protein [Eutypa lata UCREL1] | 54.1 | 4.0E-106 |
| DIP_10005989 | putative fad binding domain protein [Eutypa lata UCREL1] | 59.2 | 0.0 |
| DIP_10005990 | hypothetical protein UCREL1_7211 [Eutypa lata UCREL1] | 56.7 | 6.0E-49 |
| DIP_10005991 | putative alpha beta hydrolase fold-3 domain-containing protein [Eutypa lata UCREL1] | 52.7 | 5.0E-111 |
| DIP_10005992 | hypothetical protein PFICI_01345 [Pestalotiopsis fici W106-1] | 46.6 | 1.0E-73 |
| DIP_10005993 | NA | | |
| DIP_10005994 | hypothetical protein UCREL1_4639 [Eutypa lata UCREL1] | 49.5 | 6.0E-47 |
| DIP_10005995 | putative gpn-loop gtpase 3 like protein [Eutypa lata UCREL1] | 81.2 | 0.0 |
| DIP_10005996 | putative s-adenosylmethionine synthetase protein [Eutypa lata UCREL1] | 87.1 | 0.0 |
| DIP_10005997 | fungal specific transcription factor domain-containing protein [Colletotrichum graminicola M1.001] | 72.9 | 0.0 |
| DIP_10005998 | phosphoethanolamine transferase class N [Ustilaginoidea virens] | 58.3 | 1.0E-140 |
| DIP_10005999 | putative gpi ethanolamine phosphate transferase protein [Eutypa lata UCREL1] | 80.2 | 0.0 |
| DIP_10006000 | putative spindle pole body-associated protein sad1 protein [Eutypa lata UCREL1] | 42.5 | 1.0E-122 |
| DIP_10006001 | hypothetical protein PFICI_04797 [Pestalotiopsis fici W106-1] | 60.8 | 1.0E-108 |
| DIP_10006002 | ATP-dependent RNA helicase dbp-4 [Pestalotiopsis fici W106-1] | 65.9 | 0.0 |
| DIP_10006003 | putative choline oxidase protein [Eutypa lata UCREL1] | 79.4 | 0.0 |
| DIP_10006004 | Betaine aldehyde dehydrogenase [Pestalotiopsis fici W106-1] | 80.1 | 0.0 |
| DIP_10006005 | NA | | |
| DIP_10006006 | hypothetical protein THAR02_08433 [Trichoderma harzianum] | 55.7 | 0.0 |
| DIP_10006007 | putative nad h-dependent d-xylose reductase protein [Eutypa lata UCREL1] | 76 | 0.0 |
| DIP_10006008 | predicted protein [Nectria haematococca mpVI 77-13-4] | 62.6 | 1.0E-99 |
| DIP_10006009 | NA | | |
| DIP_10006010 | hypothetical protein TRIVIDRAFT_72675 [Trichoderma virens Gv29-8] | 54.6 | 0.0 |
| DIP_10006011 | putative iswi chromatin-remodeling complex atpase isw2 protein [Eutypa lata UCREL1] | 79.7 | 0.0 |
| DIP_10006012 | putative transcription initiation factor iia gamma chain protein [Eutypa lata UCREL1] | 91 | 3.0E-73 |
| DIP_10006013 | NA | | |
| DIP_10006014 | NA | | |
| DIP_10006015 | NA | | |
| DIP_10006016 | NA | | |
| DIP_10006017 | Mitochondrial division protein 1 [Pestalotiopsis fici W106-1] | 76.6 | 0.0 |
| DIP_10006018 | putative chorismate mutase protein [Eutypa lata UCREL1] | 81.4 | 1.0E-156 |
| DIP_10006019 | putative bcp1-like protein [Eutypa lata UCREL1] | 70.8 | 2.0E-139 |
| DIP_10006020 | GMP synthase [Talaromyces stipitatus ATCC 10500] | 76.6 | 0.0 |
| DIP_10006021 | hypothetical protein UCREL1_3641 [Eutypa lata UCREL1] | 55.4 | 1.0E-120 |
| DIP_10006022 | NA | | |
| DIP_10006023 | conserved hypothetical protein [Talaromyces stipitatus ATCC 10500] | 44.1 | 3.0E-85 |
| DIP_10006024 | hypothetical protein PFICI_04536 [Pestalotiopsis fici W106-1] | 39.4 | 3.0E-69 |
| DIP_10006025 | hypothetical protein GLRG_08331 [Colletotrichum graminicola M1.001] | 48.9 | 0.0 |
| DIP_10006026 | hypothetical protein UCREL1_3644 [Eutypa lata UCREL1] | 64.2 | 3.0E-134 |
| DIP_10006027 | hypothetical protein MGG_01731 [Magnaporthe oryzae 70-15] | 47.5 | 2.0E-44 |
| DIP_10006028 | NA | | |
| DIP_10006029 | NA | | |
| DIP_10006030 | hypothetical protein MGG_05058 [Magnaporthe oryzae 70-15] | 36.1 | 7.0E-09 |
| DIP_10006031 | NA | | |
| DIP_10006032 | putative u2 small nuclear ribonucleoprotein a protein [Eutypa lata UCREL1] | 84.8 | 9.0E-154 |
| DIP_10006033 | putative histone-like transcription factor and archaeal histone protein [Eutypa lata UCREL1] | 52.5 | 8.0E-69 |
| DIP_10006034 | putative voltage gated chloride channel protein [Eutypa lata UCREL1] | 81.3 | 0.0 |
| DIP_10006035 | hypothetical protein GLRG_05881 [Colletotrichum graminicola M1.001] | 60.4 | 1.0E-58 |
| DIP_10006036 | hypothetical protein PFICI_05058 [Pestalotiopsis fici W106-1] | 69.1 | 2.0E-176 |
| DIP_10006037 | putative atp-binding domain 1 family member b protein [Eutypa lata UCREL1] | 85 | 0.0 |
| DIP_10006038 | hypothetical protein PFICI_05060 [Pestalotiopsis fici W106-1] | 66.7 | 2.0E-64 |
| DIP_10006039 | putative molybdenum cofactor synthesis domain-containing protein [Eutypa lata UCREL1] | 64.3 | 0.0 |
| DIP_10006040 | Actin cytoskeleton-regulatory complex protein end-3 [Pestalotiopsis fici W106-1] | 71.5 | 0.0 |
| DIP_10006041 | putative amid-like mitochondrial oxidoreductase protein [Eutypa lata UCREL1] | 59.7 | 2.0E-159 |
| DIP_10006042 | NA | | |
| DIP_10006043 | NA | | |
| DIP_10006044 | NA | | |
| DIP_10006045 | hypothetical protein DOTSEDRAFT_72430 [Dothistroma septosporum NZE10] | 32 | 6.0E-42 |
| DIP_10006046 | hypothetical protein NECHADRAFT_58149 [Nectria haematococca mpVI 77-13-4] | 67 | 9.0E-43 |
| DIP_10006047 | hypothetical protein PFICI_05012 [Pestalotiopsis fici W106-1] | 49.8 | 0.0 |
| DIP_10006048 | putative disulfide isomerase protein [Eutypa lata UCREL1] | 72.9 | 0.0 |
| DIP_10006049 | hypothetical protein CGLO_08961 [Colletotrichum gloeosporioides Cg-14] | 58.5 | 0.0 |
| DIP_10006050 | putative pre-rrna-processing protein esf1 protein [Eutypa lata UCREL1] | 59.2 | 0.0 |
| DIP_10006051 | putative dual specificity phosphatase protein [Eutypa lata UCREL1] | 52 | 4.0E-118 |
| DIP_10006052 | putative spo7-like protein [Eutypa lata UCREL1] | 67.8 | 0.0 |
| DIP_10006053 | pyridoxal-dependent decarboxylase domain protein [Colletotrichum gloeosporioides Nara gc5] | 60.4 | 0.0 |
| DIP_10006054 | hypothetical protein V496_07208 [Pseudogymnoascus pannorum VKM F-4515 (FW-2607)] | 41.1 | 1.0E-20 |
| DIP_10006055 | hypothetical protein MYCTH_100838 [Myceliophthora thermophila ATCC 42464] | 33.5 | 1.0E-22 |
| DIP_10006056 | NA | | |
| DIP_10006057 | hypothetical protein PFICI_04255 [Pestalotiopsis fici W106-1] | 81.5 | 0.0 |
| DIP_10006058 | hypothetical protein SBOR_9046 [Sclerotinia borealis F-4157] | 36.2 | 2.0E-10 |
| DIP_10006059 | conserved hypothetical protein [Verticillium alfalfae VaMs.102] | 58.5 | 8.0E-66 |
| DIP_10006060 | hypothetical protein PFICI_04258 [Pestalotiopsis fici W106-1] | 57.8 | 0.0 |
| DIP_10006061 | cytochrome P450 [Colletotrichum fioriniae PJ7] | 47.9 | 1.0E-160 |
| DIP_10006062 | hypothetical protein UCREL1_3736 [Eutypa lata UCREL1] | 37.1 | 3.0E-51 |
| DIP_10006063 | NA | | |
| DIP_10006064 | NA | | |
| DIP_10006065 | putative rna polymerase ii-associated protein [Eutypa lata UCREL1] | 75 | 0.0 |
| DIP_10006066 | putative tor signaling pathway regulator protein [Eutypa lata UCREL1] | 73.1 | 0.0 |
| DIP_10006067 | putative mase3 domain-containing protein [Togninia minima UCRPA7] | 42.4 | 0.0 |
| DIP_10006068 | NA | | |
| DIP_10006069 | hypothetical protein Cob_06403 [Colletotrichum orbiculare MAFF 240422] | 35.5 | 4.0E-30 |
| DIP_10006070 | NA | | |
| DIP_10006071 | putative ubiquitin carboxyl-terminal hydrolase, family 1 [Colletotrichum sublineola] | 75.1 | 4.0E-132 |
| DIP_10006072 | glycoside hydrolase family 13 protein [Botrytis cinerea T4] | 51.6 | 0.0 |
| DIP_10006073 | hypothetical protein M7I_1066 [Glarea lozoyensis 74030] | 40.6 | 9.0E-23 |
| DIP_10006074 | hypothetical protein PFICI_00529 [Pestalotiopsis fici W106-1] | 63.1 | 0.0 |
| DIP_10006075 | NA | | |
| DIP_10006076 | NA | | |
| DIP_10006077 | hypothetical protein PFICI_00822 [Pestalotiopsis fici W106-1] | 50.8 | 0.0 |
| DIP_10006078 | putative brix domain-containing protein [Eutypa lata UCREL1] | 78.6 | 0.0 |
| DIP_10006079 | hypothetical protein UCREL1_6246 [Eutypa lata UCREL1] | 68.6 | 4.0E-49 |
| DIP_10006080 | NA | | |
| DIP_10006081 | NA | | |
| DIP_10006082 | NA | | |
| DIP_10006083 | hypothetical protein FOIG_16956, partial [Fusarium oxysporum f. sp. cubense tropical race 4 54006] | 51.9 | 3.0E-61 |
| DIP_10006084 | NA | | |
| DIP_10006085 | putative glucooligosaccharide oxidase protein [Neofusicoccum parvum UCRNP2] | 45.5 | 4.0E-153 |
| DIP_10006086 | hypothetical protein PFICI_01190 [Pestalotiopsis fici W106-1] | 57.9 | 0.0 |
| DIP_10006087 | hypothetical protein PFICI_01188 [Pestalotiopsis fici W106-1] | 64.9 | 7.0E-118 |
| DIP_10006088 | hypothetical protein PFICI_01187 [Pestalotiopsis fici W106-1] | 74.5 | 1.0E-52 |
| DIP_10006089 | hypothetical protein PFICI_01186 [Pestalotiopsis fici W106-1] | 74.5 | 8.0E-176 |
| DIP_10006090 | hypothetical protein PFICI_01185 [Pestalotiopsis fici W106-1] | 40.3 | 1.0E-65 |
| DIP_10006091 | hypothetical protein PFICI_01184 [Pestalotiopsis fici W106-1] | 51.2 | 3.0E-155 |
| DIP_10006092 | hypothetical protein CGLO_04408 [Colletotrichum gloeosporioides Cg-14] | 44.4 | 2.0E-24 |
| DIP_10006093 | NA | | |
| DIP_10006094 | NA | | |
| DIP_10006095 | hypothetical protein PFICI_01181 [Pestalotiopsis fici W106-1] | 63.3 | 4.0E-154 |
| DIP_10006096 | hypothetical protein PFICI_01180 [Pestalotiopsis fici W106-1] | 56.3 | 0.0 |
| DIP_10006097 | hypothetical protein UCREL1_579 [Eutypa lata UCREL1] | 67.6 | 1.0E-32 |
| DIP_10006098 | hypothetical protein PFICI_01178 [Pestalotiopsis fici W106-1] | 52.1 | 0.0 |
| DIP_10006099 | putative nucleolar protein 12 protein [Eutypa lata UCREL1] | 64.1 | 0.0 |
| DIP_10006100 | putative tubulin alpha chain protein [Eutypa lata UCREL1] | 94.1 | 0.0 |
| DIP_10006101 | hypothetical protein PFICI_01174 [Pestalotiopsis fici W106-1] | 48.4 | 1.0E-159 |
| DIP_10006102 | hypothetical protein UCREL1_573 [Eutypa lata UCREL1] | 38.9 | 3.0E-97 |
| DIP_10006103 | meiotic chromosome segregation protein [Magnaporthe oryzae 70-15] | 70.2 | 4.0E-160 |
| DIP_10006104 | putative cystathionine gamma-lyase protein [Eutypa lata UCREL1] | 83.5 | 0.0 |
| DIP_10006105 | hypothetical protein COCHEDRAFT_1191132 [Bipolaris maydis C5] | 63.4 | 1.0E-165 |
| DIP_10006106 | hypothetical protein PFICI_01170 [Pestalotiopsis fici W106-1] | 35.6 | 2.0E-78 |
| DIP_10006107 | uridine permease [Colletotrichum gloeosporioides Nara gc5] | 72.5 | 0.0 |
| DIP_10006108 | hypothetical protein FG05_01135 [Fusarium graminearum] | 86.5 | 2.0E-170 |
| DIP_10006109 | putative aconitate hydratase protein [Eutypa lata UCREL1] | 59.3 | 4.0E-77 |
| DIP_10006110 | putative checkpoint protein [Eutypa lata UCREL1] | 58 | 0.0 |
| DIP_10006111 | 60s ribosomal protein rla2, P2 [Trichoderma reesei QM6a] | 71.6 | 1.0E-34 |
| DIP_10006112 | putative poly polymerase 2 adp-ribosyltransferase 2 protein [Eutypa lata UCREL1] | 67.9 | 0.0 |
| DIP_10006113 | putative nuclear fragile x mental retardation-interacting protein 1 protein [Eutypa lata UCREL1] | 62.6 | 8.0E-97 |
| DIP_10006114 | putative dna-directed rna polymerase iii polypeptide protein [Eutypa lata UCREL1] | 79.8 | 2.0E-114 |
| DIP_10006115 | putative transmembrane protein 111 protein [Eutypa lata UCREL1] | 83.7 | 2.0E-149 |
| DIP_10006116 | hypothetical protein PFICI_01158 [Pestalotiopsis fici W106-1] | 57.3 | 5.0E-90 |
| DIP_10006117 | hypothetical protein TD95_004374 [Thielaviopsis punctulata] | 38.5 | 2.0E-83 |
| DIP_10006118 | putative methylenetetrahydrofolate dehydrogenase protein [Eutypa lata UCREL1] | 85 | 0.0 |
| DIP_10006119 | NA | | |
| DIP_10006120 | hypothetical protein FOIG_01014 [Fusarium oxysporum f. sp. cubense tropical race 4 54006] | 53.7 | 0.0 |
| DIP_10006121 | putative f-box domain-containing protein [Togninia minima UCRPA7] | 35.5 | 3.0E-13 |
| DIP_10006122 | NA | | |
| DIP_10006123 | hypothetical protein VDAG_03221 [Verticillium dahliae VdLs.17] | 34.7 | 1.0E-06 |
| DIP_10006124 | putative glycosyl transferase family 17 protein [Eutypa lata UCREL1] | 63.9 | 3.0E-149 |
| DIP_10006125 | hypothetical protein PFICI_01155 [Pestalotiopsis fici W106-1] | 67.1 | 0.0 |
| DIP_10006126 | putative gcn5-related n-acetyltransferase protein [Eutypa lata UCREL1] | 34.6 | 6.0E-23 |
| DIP_10006127 | putative endoplasmic reticulum-golgi intermediate compartment protein 3 protein [Eutypa lata UCREL1] | 80.6 | 0.0 |
| DIP_10006128 | hypothetical protein VDAG_04255 [Verticillium dahliae VdLs.17] | 53.9 | 0.0 |
| DIP_10006129 | hypothetical protein PFICI_01153 [Pestalotiopsis fici W106-1] | 44 | 1.0E-75 |
| DIP_10006130 | L-allo-threonine aldolase [Colletotrichum higginsianum] | 77 | 1.0E-50 |
| DIP_10006131 | putative proteasome subunit beta type 7 precursor protein [Eutypa lata UCREL1] | 92.9 | 0.0 |
| DIP_10006132 | hypothetical protein FOWG_06907 [Fusarium oxysporum f. sp. lycopersici MN25] | 45.1 | 0.0 |
| DIP_10006133 | hypothetical protein THITE_2121090 [Thielavia terrestris NRRL 8126] | 66.4 | 0.0 |
| DIP_10006134 | putative mitochondrial f0 subunit e protein [Eutypa lata UCREL1] | 73.4 | 8.0E-45 |
| DIP_10006135 | hypothetical protein HMPREF1624_07127 [Sporothrix schenckii ATCC 58251] | 39.7 | 3.0E-63 |
| DIP_10006136 | putative oligosaccharyl transferase stt3 subunit protein [Eutypa lata UCREL1] | 80.8 | 0.0 |
| DIP_10006137 | NA | | |
| DIP_10006138 | hypothetical protein AKAW_10619 [Aspergillus kawachii IFO 4308] | 52.1 | 1.0E-124 |
| DIP_10006139 | cytochrome P450 [Aspergillus niger CBS 513.88] | 71.1 | 0.0 |
| DIP_10006140 | hypothetical protein PFICI_01208 [Pestalotiopsis fici W106-1] | 54 | 2.0E-164 |
| DIP_10006141 | putative dna replication complex gins protein sld5 [Diaporthe ampelina] | 73.8 | 6.0E-112 |
| DIP_10006142 | putative protein phosphatase methylesterase 1 protein [Eutypa lata UCREL1] | 67.7 | 0.0 |
| DIP_10006143 | putative dihydrouridine synthase protein [Eutypa lata UCREL1] | 71.3 | 0.0 |
| DIP_10006144 | hypothetical protein PFICI_01508 [Pestalotiopsis fici W106-1] | 56.2 | 0.0 |
| DIP_10006145 | hypothetical protein PFICI_01507 [Pestalotiopsis fici W106-1] | 49.7 | 0.0 |
| DIP_10006146 | hypothetical protein HMPREF1624_06122 [Sporothrix schenckii ATCC 58251] | 56.5 | 2.0E-47 |
| DIP_10006147 | putative cytochrome c1 protein [Eutypa lata UCREL1] | 86 | 0.0 |
| DIP_10006148 | putative phospho-2-dehydro-3-deoxyheptonate aldolase protein [Eutypa lata UCREL1] | 88.9 | 0.0 |
| DIP_10006149 | hypothetical protein Z517_11623 [Fonsecaea pedrosoi CBS 271.37] | 42.7 | 8.0E-56 |
| DIP_10006150 | NA | | |
| DIP_10006151 | hypothetical protein PFICI_01210 [Pestalotiopsis fici W106-1] | 82.6 | 0.0 |
| DIP_10006152 | hypothetical protein PFICI_01526 [Pestalotiopsis fici W106-1] | 60.6 | 1.0E-96 |
| DIP_10006153 | hypothetical protein PFICI_01209 [Pestalotiopsis fici W106-1] | 54 | 0.0 |
| DIP_10006154 | putative pyroglutamyl peptidase type protein [Eutypa lata UCREL1] | 49.8 | 5.0E-74 |
| DIP_10006155 | hypothetical protein FAVG1_09740 [Fusarium avenaceum] | 37.4 | 1.0E-28 |
| DIP_10006156 | hypothetical protein S7711_02242 [Stachybotrys chartarum IBT 7711] | 66 | 1.0E-21 |
| DIP_10006157 | putative nadph dehydrogenase protein [Eutypa lata UCREL1] | 75.6 | 0.0 |
| DIP_10006158 | hypothetical protein NEUTE1DRAFT_88941 [Neurospora tetrasperma FGSC 2508] | 38.5 | 5.0E-57 |
| DIP_10006159 | hypothetical protein CGGC5_13773 [Colletotrichum gloeosporioides Nara gc5] | 48.4 | 6.0E-115 |
| DIP_10006160 | hypothetical protein PFICI_01511 [Pestalotiopsis fici W106-1] | 56.6 | 2.0E-162 |
| DIP_10006161 | predicted protein [Trichoderma reesei QM6a] | 30.5 | 4.0E-30 |
| DIP_10006162 | NA | | |
| DIP_10006163 | NA | | |
| DIP_10006164 | NA | | |
| DIP_10006165 | putative inorganic pyrophosphatase protein [Eutypa lata UCREL1] | 89.7 | 0.0 |
| DIP_10006166 | putative mitochondrial ribosomal protein l44 protein [Eutypa lata UCREL1] | 67.4 | 5.0E-38 |
| DIP_10006167 | putative sec20 domain-containing protein [Eutypa lata UCREL1] | 55.1 | 2.0E-137 |
| DIP_10006168 | NA | | |
| DIP_10006169 | hypothetical protein PFICI_01200 [Pestalotiopsis fici W106-1] | 55.9 | 0.0 |
| DIP_10006170 | predicted protein [Nectria haematococca mpVI 77-13-4] | 57.1 | 0.0 |
| DIP_10006171 | NA | | |
| DIP_10006172 | NA | | |
| DIP_10006173 | putative solute carrier family 40 (iron-regulated transporter) protein [Eutypa lata UCREL1] | 62 | 2.0E-177 |
| DIP_10006174 | hypothetical protein PFICI_02327 [Pestalotiopsis fici W106-1] | 58.9 | 0.0 |
| DIP_10006175 | hypothetical protein TD95_003377 [Thielaviopsis punctulata] | 56.3 | 6.0E-28 |
| DIP_10006176 | putative g2 mitotic-specific cyclin-b protein [Eutypa lata UCREL1] | 71.1 | 0.0 |
| DIP_10006177 | putative mitochondrial 54s ribosomal protein rml2 protein [Eutypa lata UCREL1] | 90.9 | 0.0 |
| DIP_10006178 | hypothetical protein FPSE_03739 [Fusarium pseudograminearum CS3096] | 42.5 | 2.0E-89 |
| DIP_10006179 | hypothetical protein PFICI_00864 [Pestalotiopsis fici W106-1] | 56.9 | 6.0E-159 |
| DIP_10006180 | hypothetical protein PFICI_00863 [Pestalotiopsis fici W106-1] | 60.5 | 1.0E-155 |
| DIP_10006181 | NA | | |
| DIP_10006182 | putative casein kinase ii beta subunit protein [Eutypa lata UCREL1] | 90.9 | 5.0E-144 |
| DIP_10006183 | hypothetical protein PFICI_01537 [Pestalotiopsis fici W106-1] | 69.3 | 0.0 |
| DIP_10006184 | hypothetical protein PFICI_01536 [Pestalotiopsis fici W106-1] | 70.9 | 3.0E-150 |
| DIP_10006185 | binuclear zinc transcription factor [Colletotrichum orbiculare MAFF 240422] | 61.9 | 0.0 |
| DIP_10006186 | hypothetical protein PFICI_01539 [Pestalotiopsis fici W106-1] | 69.7 | 0.0 |
| DIP_10006187 | hypothetical protein PFICI_01529 [Pestalotiopsis fici W106-1] | 70.8 | 0.0 |
| DIP_10006188 | acetate kinase [Colletotrichum gloeosporioides Nara gc5] | 46.7 | 5.0E-103 |
| DIP_10006189 | hypothetical protein PFICI_01532 [Pestalotiopsis fici W106-1] | 56.3 | 4.0E-80 |
| DIP_10006190 | putative pep31 c terminal domain-containing protein [Eutypa lata UCREL1] | 62.2 | 0.0 |
| DIP_10006191 | putative cell cycle control protein [Eutypa lata UCREL1] | 86.3 | 0.0 |
| DIP_10006192 | hypothetical protein PFICI_01535 [Pestalotiopsis fici W106-1] | 54.3 | 5.0E-162 |
| DIP_10006193 | hypothetical protein FOXB_00234 [Fusarium oxysporum Fo5176] | 32.6 | 3.0E-62 |
| DIP_10006194 | hypothetical protein CFIO01_07293 [Colletotrichum fioriniae PJ7] | 76.7 | 7.0E-125 |
| DIP_10006195 | putative helix-loop-helix dna-binding domain-containing protein [Eutypa lata UCREL1] | 69.4 | 0.0 |
| DIP_10006196 | putative upf0171 domain protein [Eutypa lata UCREL1] | 64 | 0.0 |
| DIP_10006197 | hypothetical protein NEUTE1DRAFT_88612 [Neurospora tetrasperma FGSC 2508] | 42.4 | 2.0E-47 |
| DIP_10006198 | hypothetical protein PFICI_01958 [Pestalotiopsis fici W106-1] | 73.5 | 0.0 |
| DIP_10006199 | hypothetical protein PFICI_01952 [Pestalotiopsis fici W106-1] | 53.5 | 7.0E-169 |
| DIP_10006200 | putative lim domain-containing protein [Togninia minima UCRPA7] | 52.5 | 4.0E-138 |
| DIP_10006201 | putative ubiquitin family protein [Togninia minima UCRPA7] | 86.7 | 2.0E-52 |
| DIP_10006202 | putative nuclear pore complex protein nup107 protein [Eutypa lata UCREL1] | 57.5 | 0.0 |
| DIP_10006203 | hypothetical protein [Podospora anserina S mat+] | 45.7 | 3.0E-79 |
| DIP_10006204 | NA | | |
| DIP_10006205 | hypothetical protein PFICI_01940 [Pestalotiopsis fici W106-1] | 56.8 | 0.0 |
| DIP_10006206 | putative vacuolar atp synthase subunit c 1 protein [Eutypa lata UCREL1] | 72.1 | 0.0 |
| DIP_10006207 | putative adenylyl-sulfate kinase protein [Eutypa lata UCREL1] | 91.4 | 4.0E-116 |
| DIP_10006208 | hypothetical protein OIDMADRAFT_121040 [Oidiodendron maius Zn] | 66.7 | 9.0E-93 |
| DIP_10006209 | putative ribosome biogenesis protein mak21 protein [Eutypa lata UCREL1] | 59 | 0.0 |
| DIP_10006210 | hypothetical protein O988_08820 [Pseudogymnoascus pannorum VKM F-3808] | 39.5 | 3.0E-59 |
| DIP_10006211 | hypothetical protein Cob_07584 [Colletotrichum orbiculare MAFF 240422] | 44.3 | 7.0E-93 |
| DIP_10006212 | hypothetical protein CGLO_09787 [Colletotrichum gloeosporioides Cg-14] | 68.9 | 0.0 |
| DIP_10006213 | putative oligopeptide transporter [Diplodia seriata] | 62.5 | 0.0 |
| DIP_10006214 | hypothetical protein UCREL1_5467 [Eutypa lata UCREL1] | 63.6 | 7.0E-155 |
| DIP_10006215 | putative phd finger containing protein phf1 protein [Eutypa lata UCREL1] | 37.3 | 4.0E-66 |
| DIP_10006216 | hypothetical protein HMPREF1120_07378 [Exophiala dermatitidis NIH/UT8656] | 36.5 | 1.0E-86 |
| DIP_10006217 | unnamed protein product [Penicillium roqueforti FM164] | 32 | 3.0E-55 |
| DIP_10006218 | hypothetical protein NECHADRAFT_67419 [Nectria haematococca mpVI 77-13-4] | 71.1 | 0.0 |
| DIP_10006219 | putative mfs drug transporter protein [Eutypa lata UCREL1] | 68.9 | 0.0 |

| | | | |
|---|---|---|---|
| DIP_10006220 | putative mac1 protein [Eutypa lata UCREL1] | 75.8 | 0.0 |
| DIP_10006221 | hypothetical protein THITE_2114197 [Thielavia terrestris NRRL 8126] | 42 | 6.0E-86 |
| DIP_10006222 | hypothetical protein UCREL1_5375 [Eutypa lata UCREL1] | 66.4 | 0.0 |
| DIP_10006223 | putative kinesin family protein [Eutypa lata UCREL1] | 59.5 | 0.0 |
| DIP_10006224 | hypothetical protein PFICI_05038 [Pestalotiopsis fici W106-1] | 34.9 | 4.0E-55 |
| DIP_10006225 | NA | | |
| DIP_10006226 | NA | | |
| DIP_10006227 | putative hydrophobin precursor protein [Eutypa lata UCREL1] | 59.4 | 5.0E-37 |
| DIP_10006228 | putative nmda receptor-regulated protein 1 protein [Eutypa lata UCREL1] | 73.9 | 0.0 |
| DIP_10006229 | hypothetical protein PFICI_00711 [Pestalotiopsis fici W106-1] | 64.1 | 2.0E-86 |
| DIP_10006230 | 26S protease regulatory subunit 6A [Pestalotiopsis fici W106-1] | 90.2 | 0.0 |
| DIP_10006231 | putative trypsin-like serine protease protein [Eutypa lata UCREL1] | 44 | 9.0E-78 |
| DIP_10006232 | STU1 [Verticillium alfalfae VaMs.102] | 56.5 | 0.0 |
| DIP_10006233 | putative pbsp domain-containing protein [Togninia minima UCRPA7] | 41.1 | 9.0E-71 |
| DIP_10006234 | hypothetical protein UCREL1_5832 [Eutypa lata UCREL1] | 54.1 | 1.0E-106 |
| DIP_10006235 | hypothetical protein PFICI_04269 [Pestalotiopsis fici W106-1] | 62.8 | 4.0E-10 |
| DIP_10006236 | Isocitrate dehydrogenase [Pestalotiopsis fici W106-1] | 82.4 | 0.0 |
| DIP_10006237 | putative exocyst complex component sec5 protein [Eutypa lata UCREL1] | 69.8 | 0.0 |
| DIP_10006238 | hypothetical protein UCRPA7_6831 [Togninia minima UCRPA7] | 66.1 | 2.0E-104 |
| DIP_10006239 | hypothetical protein UCREL1_5829 [Eutypa lata UCREL1] | 48.8 | 8.0E-51 |
| DIP_10006240 | hypothetical protein PFICI_04483 [Pestalotiopsis fici W106-1] | 57.1 | 0.0 |
| DIP_10006241 | hypothetical protein PFICI_04463 [Pestalotiopsis fici W106-1] | 35.3 | 2.0E-178 |
| DIP_10006242 | hypothetical protein PFICI_04462 [Pestalotiopsis fici W106-1] | 64.7 | 0.0 |
| DIP_10006243 | laccase, partial [Daldinia eschscholtzii] | 49.9 | 0.0 |
| DIP_10006244 | putative hsp70-like protein [Eutypa lata UCREL1] | 92.6 | 0.0 |
| DIP_10006245 | hypothetical protein PFICI_04459 [Pestalotiopsis fici W106-1] | 50.3 | 0.0 |
| DIP_10006246 | L-2-hydroxyglutarate dehydrogenase-like protein [Acremonium chrysogenum ATCC 11550] | 54.1 | 1.0E-153 |
| DIP_10006247 | NA | | |
| DIP_10006248 | hypothetical protein G647_04484 [Cladophialophora carrionii CBS 160.54] | 38.9 | 1.0E-118 |
| DIP_10006249 | hypothetical protein BBAD15_g2454 [Beauveria bassiana D1-5] | 31.5 | 2.0E-11 |
| DIP_10006250 | hypothetical protein PFICI_04637 [Pestalotiopsis fici W106-1] | 61.1 | 0.0 |
| DIP_10006251 | hypothetical protein FOC1_g10005019 [Fusarium oxysporum f. sp. cubense race 1] | 46 | 9.0E-45 |
| DIP_10006252 | NRPS-like enzyme, putative [Talaromyces marneffei ATCC 18224] | 39.7 | 1.0E-122 |
| DIP_10006253 | hypothetical protein FOZG_07262 [Fusarium oxysporum Fo47] | 64.8 | 0.0 |
| DIP_10006254 | NA | | |
| DIP_10006255 | putative cop9 signalosome subunit 6 protein [Eutypa lata UCREL1] | 63.3 | 1.0E-173 |
| DIP_10006256 | predicted protein [Nectria haematococca mpVI 77-13-4] | 70.5 | 0.0 |
| DIP_10006257 | NA | | |
| DIP_10006258 | NA | | |
| DIP_10006259 | putative cyclin domain-containing protein [Eutypa lata UCREL1] | 54.3 | 2.0E-121 |
| DIP_10006260 | putative interferon-induced gtp-binding protein mx2 protein [Eutypa lata UCREL1] | 46.5 | 0.0 |
| DIP_10006261 | putative ring-like domain-containing protein [Eutypa lata UCREL1] | 56.1 | 6.0E-154 |
| DIP_10006262 | putative dna repair protein [Eutypa lata UCREL1] | 66.5 | 1.0E-109 |
| DIP_10006263 | hypothetical protein CH063_01111 [Colletotrichum higginsianum] | 43.3 | 1.0E-26 |
| DIP_10006264 | putative gram domain-containing protein [Eutypa lata UCREL1] | 62.5 | 0.0 |
| DIP_10006265 | hypothetical protein UCREL1_9630 [Eutypa lata UCREL1] | 54 | 0.0 |
| DIP_10006266 | hypothetical protein GGTG_11961 [Gaeumannomyces graminis var. tritici R3-111a-1] | 39.5 | 6.0E-77 |
| DIP_10006267 | hypothetical protein PFICI_05127 [Pestalotiopsis fici W106-1] | 68.6 | 0.0 |
| DIP_10006268 | hypothetical protein PFICI_06040 [Pestalotiopsis fici W106-1] | 30.3 | 1.0E-33 |
| DIP_10006269 | hypothetical protein EPUS_07064 [Endocarpon pusillum Z07020] | 36.1 | 9.0E-61 |
| DIP_10006270 | hypothetical protein A1O7_07816 [Cladophialophora yegresii CBS 114405] | 48.9 | 5.0E-161 |
| DIP_10006271 | hypothetical protein SNOG_04807 [Phaeosphaeria nodorum SN15] | 49.9 | 8.0E-126 |
| DIP_10006272 | putative atp synthase f0 protein [Eutypa lata UCREL1] | 82.4 | 2.0E-167 |
| DIP_10006273 | hypothetical protein MYCTH_2298517 [Myceliophthora thermophila ATCC 42464] | 72.1 | 1.0E-105 |
| DIP_10006274 | putative rad5-like protein [Eutypa lata UCREL1] | 66.5 | 0.0 |
| DIP_10006275 | benzoate 4-monooxygenase cytochrome P450 [Verticillium alfalfae VaMs.102] | 49.6 | 0.0 |
| DIP_10006276 | hypothetical protein PFICI_04906 [Pestalotiopsis fici W106-1] | 37.8 | 1.0E-70 |
| DIP_10006277 | hypothetical protein PFICI_04905 [Pestalotiopsis fici W106-1] | 55.5 | 1.0E-118 |
| DIP_10006278 | NA | | |
| DIP_10006279 | putative survival factor 1 protein [Eutypa lata UCREL1] | 72 | 0.0 |
| DIP_10006280 | 54S ribosomal protein L19 [Pestalotiopsis fici W106-1] | 85.6 | 6.0E-92 |
| DIP_10006281 | hypothetical protein SMAC_01810 [Sordaria macrospora k-hell] | 46.1 | 0.0 |
| DIP_10006282 | NA | | |
| DIP_10006283 | chaperonin 10 kDa subunit [Colletotrichum higginsianum] | 84.6 | 1.0E-61 |
| DIP_10006284 | putative glycoside hydrolase family 61 protein [Eutypa lata UCREL1] | 58.5 | 6.0E-140 |
| DIP_10006285 | putative alpha beta hydrolase fold protein [Eutypa lata UCREL1] | 43.3 | 7.0E-51 |
| DIP_10006286 | NA | | |
| DIP_10006287 | NA | | |
| DIP_10006288 | hypothetical protein [Podospora anserina S mat+] | 74.4 | 8.0E-169 |
| DIP_10006289 | putative dynein heavy chain protein [Eutypa lata UCREL1] | 87.3 | 0.0 |
| DIP_10006290 | hypothetical protein PFICI_05672 [Pestalotiopsis fici W106-1] | 56.4 | 2.0E-105 |
| DIP_10006291 | NA | | |
| DIP_10006292 | COPI-coated vesicle protein [Sporothrix brasiliensis 5110] | 79.2 | 1.0E-71 |
| DIP_10006293 | hypothetical protein A1O9_00561 [Exophiala aquamarina CBS 119918] | 34 | 7.0E-58 |
| DIP_10006294 | calcineurin-like phosphoesterase [Colletotrichum graminicola M1.001] | 88.6 | 0.0 |
| DIP_10006295 | Exocyst complex component EXO84 [Pestalotiopsis fici W106-1] | 69.6 | 0.0 |
| DIP_10006296 | putative mitochondrial import receptor subunit tom-20 protein [Eutypa lata UCREL1] | 68.4 | 3.0E-65 |
| DIP_10006297 | NA | | |
| DIP_10006298 | hypothetical protein PFICI_00720 [Pestalotiopsis fici W106-1] | 62.2 | 0.0 |
| DIP_10006299 | hypothetical protein A1O5_12037 [Cladophialophora psammophila CBS 110553] | 45.2 | 8.0E-28 |
| DIP_10006300 | methyltransferase [Colletotrichum higginsianum] | 44.4 | 4.0E-68 |
| DIP_10006301 | putative cyclin-like protein [Eutypa lata UCREL1] | 54.6 | 1.0E-132 |
| DIP_10006302 | hypothetical protein PFICI_08044 [Pestalotiopsis fici W106-1] | 53 | 5.0E-11 |
| DIP_10006303 | putative nuclear segregation protein [Eutypa lata UCREL1] | 77.3 | 0.0 |
| DIP_10006304 | hypothetical protein UCREL1_8822 [Eutypa lata UCREL1] | 66.7 | 3.0E-95 |
| DIP_10006305 | hypothetical protein PFICI_04471 [Pestalotiopsis fici W106-1] | 64.7 | 0.0 |
| DIP_10006306 | putative nol1 nop2 sun family protein [Eutypa lata UCREL1] | 68.4 | 0.0 |
| DIP_10006307 | hypothetical protein PFICI_14909 [Pestalotiopsis fici W106-1] | 55.9 | 5.0E-170 |
| DIP_10006308 | Phosphatidylethanolamine N-methyltransferase [Pestalotiopsis fici W106-1] | 63.6 | 0.0 |
| DIP_10006309 | hypothetical protein PFICI_04467 [Pestalotiopsis fici W106-1] | 55.6 | 1.0E-42 |
| DIP_10006310 | putative eukaryotic translation initiation factor 3 subunit g protein [Eutypa lata UCREL1] | 86.1 | 0.0 |
| DIP_10006311 | NA | | |
| DIP_10006312 | hypothetical protein VHEMI03521 [Torrubiella hemipterigena] | 55.3 | 4.0E-84 |
| DIP_10006313 | putative inorganic phosphate transporter pho84 protein [Eutypa lata UCREL1] | 60 | 0.0 |
| DIP_10006314 | homoserine O-acetyltransferase [Magnaporthe oryzae Y34] | 71 | 0.0 |
| DIP_10006315 | putative tho complex subunit 2 protein [Eutypa lata UCREL1] | 71.1 | 0.0 |
| DIP_10006316 | hypothetical protein PFICI_04644 [Pestalotiopsis fici W106-1] | 52 | 1.0E-90 |
| DIP_10006317 | hypothetical protein UCREL1_135 [Eutypa lata UCREL1] | 42.9 | 0.0 |
| DIP_10006318 | hypothetical protein UCREL1_3148 [Eutypa lata UCREL1] | 53.2 | 3.0E-92 |
| DIP_10006319 | NA | | |
| DIP_10006320 | putative phosphate metabolism protein 7 protein [Eutypa lata UCREL1] | 72.4 | 0.0 |
| DIP_10006321 | putative phosphatidylglycerol phosphatidylinositol transfer protein [Eutypa lata UCREL1] | 55.9 | 4.0E-58 |
| DIP_10006322 | hypothetical protein PFICI_00196 [Pestalotiopsis fici W106-1] | 51.5 | 0.0 |
| DIP_10006323 | hypothetical protein PFICI_00195 [Pestalotiopsis fici W106-1] | 52.5 | 0.0 |
| DIP_10006324 | duf1275 domain protein [Colletotrichum gloeosporioides Nara gc5] | 58.6 | 1.0E-113 |
| DIP_10006325 | putative oxalate decarboxylase oxdc protein [Eutypa lata UCREL1] | 81.6 | 0.0 |
| DIP_10006326 | hypothetical protein VDAG_07365 [Verticillium dahliae VdLs.17] | 44.4 | 3.0E-94 |
| DIP_10006327 | hypothetical protein PFICI_01084 [Pestalotiopsis fici W106-1] | 53.6 | 0.0 |
| DIP_10006328 | hypothetical protein PFICI_00198 [Pestalotiopsis fici W106-1] | 65.6 | 0.0 |
| DIP_10006329 | peptidase family M28 [Colletotrichum fioriniae PJ7] | 72.7 | 0.0 |
| DIP_10006330 | putative ring finger domain-containing protein [Eutypa lata UCREL1] | 44.1 | 8.0E-106 |
| DIP_10006331 | NA | | |
| DIP_10006332 | putative lysophospholipase plb1 protein [Eutypa lata UCREL1] | 47.8 | 0.0 |
| DIP_10006333 | hypothetical protein CGGC5_3556 [Colletotrichum gloeosporioides Nara gc5] | 43.2 | 1.0E-24 |
| DIP_10006334 | hypothetical protein PFICI_10291 [Pestalotiopsis fici W106-1] | 34.7 | 1.0E-176 |
| DIP_10006335 | NA | | |
| DIP_10006336 | hypothetical protein UCREL1_2927 [Eutypa lata UCREL1] | 52.8 | 5.0E-14 |
| DIP_10006337 | putative quinate permease protein [Eutypa lata UCREL1] | 68.2 | 0.0 |
| DIP_10006338 | putative argonaute sirna chaperone complex subunit arb1 protein [Eutypa lata UCREL1] | 44.1 | 6.0E-113 |
| DIP_10006339 | putative inositol polyphosphate multikinase protein [Eutypa lata UCREL1] | 64.3 | 0.0 |
| DIP_10006340 | hypothetical protein PFICI_04198 [Pestalotiopsis fici W106-1] | 53.8 | 0.0 |
| DIP_10006341 | NA | | |
| DIP_10006342 | hypothetical protein PFICI_04830 [Pestalotiopsis fici W106-1] | 77.8 | 0.0 |
| DIP_10006343 | hypothetical protein PFICI_04832 [Pestalotiopsis fici W106-1] | 58.2 | 2.0E-53 |
| DIP_10006344 | putative calcium proton exchanger protein [Togninia minima UCRPA7] | 56.4 | 0.0 |
| DIP_10006345 | hypothetical protein SNOG_16400 [Phaeosphaeria nodorum SN15] | 31.1 | 6.0E-44 |
| DIP_10006346 | NA | | |
| DIP_10006347 | hypothetical protein V499_03595 [Pseudogymnoascus pannorum VKM F-103] | 40 | 2.0E-81 |
| DIP_10006348 | hypothetical protein PFICI_04770 [Pestalotiopsis fici W106-1] | 54.4 | 0.0 |
| DIP_10006349 | oxidoreductase bli-4 [Pestalotiopsis fici W106-1] | 74.5 | 0.0 |
| DIP_10006350 | putative cation diffusion facilitator family metal ion protein [Eutypa lata UCREL1] | 49.6 | 2.0E-138 |
| DIP_10006351 | NA | | |
| DIP_10006352 | hypothetical protein UCREL1_10177 [Eutypa lata UCREL1] | 61.3 | 3.0E-131 |
| DIP_10006353 | putative histidinol phosphate phosphatase family protein [Eutypa lata UCREL1] | 45.7 | 5.0E-104 |
| DIP_10006354 | hypothetical protein PFICI_05430 [Pestalotiopsis fici W106-1] | 40.9 | 2.0E-155 |
| DIP_10006355 | hypothetical protein PFICI_04202 [Pestalotiopsis fici W106-1] | 38.9 | 8.0E-67 |
| DIP_10006356 | hypothetical protein FPSE_05640 [Fusarium pseudograminearum CS3096] | 34.9 | 2.0E-39 |
| DIP_10006357 | hypothetical protein CGGC5_12841 [Colletotrichum gloeosporioides Nara gc5] | 59.9 | 2.0E-156 |
| DIP_10006358 | NA | | |
| DIP_10006359 | NA | | |
| DIP_10006360 | Ctr copper transporter [Aureobasidium melanogenum CBS 110374] | 42.4 | 3.0E-31 |
| DIP_10006361 | putative hd domain-containing protein [Eutypa lata UCREL1] | 72.1 | 9.0E-149 |
| DIP_10006362 | NA | | |
| DIP_10006363 | hypothetical protein UCREL1_6829 [Eutypa lata UCREL1] | 43.3 | 6.0E-72 |
| DIP_10006364 | NA | | |
| DIP_10006365 | hypothetical protein TRIATDRAFT_262278 [Trichoderma atroviride IMI 206040] | 52.4 | 1.0E-109 |
| DIP_10006366 | hypothetical protein CGGC5_4128 [Colletotrichum gloeosporioides Nara gc5] | 47.2 | 8.0E-76 |
| DIP_10006367 | hypothetical protein PFICI_10398 [Pestalotiopsis fici W106-1] | 34.4 | 2.0E-59 |
| DIP_10006368 | putative fad binding domain protein [Eutypa lata UCREL1] | 58.4 | 0.0 |
| DIP_10006369 | alpha/beta-Hydrolase [Glarea lozoyensis ATCC 20868] | 33 | 5.0E-50 |
| DIP_10006370 | hypothetical protein PFICI_13886 [Pestalotiopsis fici W106-1] | 55.9 | 1.0E-170 |
| DIP_10006371 | hypothetical protein PFICI_04695 [Pestalotiopsis fici W106-1] | 50.1 | 0.0 |
| DIP_10006372 | hypothetical protein PFICI_04688 [Pestalotiopsis fici W106-1] | 39.2 | 1.0E-92 |
| DIP_10006373 | NA | | |
| DIP_10006374 | hypothetical protein PFICI_00667 [Pestalotiopsis fici W106-1] | 39.1 | 8.0E-106 |
| DIP_10006375 | hypothetical protein HIM_09373 [Hirsutella minnesotensis 3608] | 52.7 | 8.0E-176 |
| DIP_10006376 | putative tetratricopeptide repeat protein 15 protein [Eutypa lata UCREL1] | 69.1 | 0.0 |
| DIP_10006377 | putative ring finger domain protein [Eutypa lata UCREL1] | 58.3 | 5.0E-64 |
| DIP_10006378 | putative gtp-binding protein rho2 protein [Eutypa lata UCREL1] | 94 | 7.0E-128 |
| DIP_10006379 | hypothetical protein GLAREA_02760 [Glarea lozoyensis ATCC 20868] | 46 | 0.0 |
| DIP_10006380 | NA | | |
| DIP_10006381 | NA | | |
| DIP_10006382 | hypothetical protein NECHADRAFT_104641 [Nectria haematococca mpVI 77-13-4] | 30.8 | 3.0E-25 |
| DIP_10006383 | putative 3-carboxymuconate cyclase protein [Eutypa lata UCREL1] | 46.7 | 1.0E-99 |
| DIP_10006384 | NA | | |
| DIP_10006385 | hypothetical protein PFICI_07441 [Pestalotiopsis fici W106-1] | 60.7 | 1.0E-130 |
| DIP_10006386 | putative pua rna binding domain-containing protein [Eutypa lata UCREL1] | 69.9 | 1.0E-37 |
| DIP_10006387 | hypothetical protein PFICI_00313 [Pestalotiopsis fici W106-1] | 71.2 | 3.0E-105 |
| DIP_10006388 | NA | | |
| DIP_10006389 | putative udp-n-acetylglucosamine-1-p transferase protein [Eutypa lata UCREL1] | 79.5 | 0.0 |
| DIP_10006390 | multidrug resistant protein, partial [Metarhizium majus ARSEF 297] | 76.5 | 0.0 |
| DIP_10006391 | major facilitator superfamily transporter [Colletotrichum gloeosporioides Cg-14] | 53.1 | 0.0 |
| DIP_10006392 | NA | | |
| DIP_10006393 | hypothetical protein PFICI_04723 [Pestalotiopsis fici W106-1] | 45.1 | 8.0E-161 |
| DIP_10006394 | hypothetical protein V498_02167, partial [Pseudogymnoascus pannorum VKM F-4517 (FW-2822)] | 34.6 | 6.0E-16 |
| DIP_10006395 | Na(+)/H(+) antiporter [Gaeumannomyces graminis var. tritici R3-111a-1] | 55.8 | 3.0E-124 |
| DIP_10006396 | hypothetical protein PFICI_04661 [Pestalotiopsis fici W106-1] | 54.5 | 0.0 |
| DIP_10006397 | predicted protein [Naegleria gruberi] | 30.2 | 2.0E-12 |
| DIP_10006398 | hypothetical protein HMPREF1120_05770 [Exophiala dermatitidis NIH/UT8656] | 45.2 | 3.0E-75 |
| DIP_10006399 | NA | | |
| DIP_10006400 | putative transcription initiation factor tfiid 127kd subunit protein [Eutypa lata UCREL1] | 61.5 | 0.0 |
| DIP_10006401 | hypothetical protein ANO11243_075020 [fungal sp. No.11243] | 37 | 2.0E-13 |
| DIP_10006402 | hypothetical protein CTHT_0058510 [Chaetomium thermophilum var. thermophilum DSM 1495] | 39.7 | 8.0E-127 |
| DIP_10006403 | cyclin-dependent kinases regulatory subunit [Ustilaginoidea virens] | 80 | 1.0E-57 |
| DIP_10006404 | putative endoplasmic oxidoreductin-1 protein [Eutypa lata UCREL1] | 68.7 | 0.0 |
| DIP_10006405 | hypothetical protein PFICI_06994 [Pestalotiopsis fici W106-1] | 37.4 | 6.0E-69 |
| DIP_10006406 | putative protein kinase protein [Eutypa lata UCREL1] | 68.8 | 4.0E-78 |
| DIP_10006407 | Protein cft-1 [Pestalotiopsis fici W106-1] | 64.1 | 0.0 |
| DIP_10006408 | hypothetical protein UCREL1_1204 [Eutypa lata UCREL1] | 56.9 | 5.0E-21 |
| DIP_10006409 | NA | | |
| DIP_10006410 | putative rna polymerase ii largest subunit protein [Eutypa lata UCREL1] | 81.7 | 0.0 |
| DIP_10006411 | putative wd domain-containing protein [Eutypa lata UCREL1] | 75.1 | 0.0 |
| DIP_10006412 | tRNAHis guanylyltransferase, putative [Cordyceps militaris CM01] | 72.4 | 5.0E-160 |
| DIP_10006413 | putative dna-directed rna polymerase ii subunit rpb1 protein [Togninia minima UCRPA7] | 40.6 | 5.0E-24 |
| DIP_10006414 | hypothetical protein UCRPA7_3789 [Togninia minima UCRPA7] | 47.2 | 1.0E-159 |
| DIP_10006415 | hypothetical protein PFICI_10293 [Pestalotiopsis fici W106-1] | 44.4 | 5.0E-68 |
| DIP_10006416 | copper fist DNA binding domain-containing protein [Colletotrichum fioriniae PJ7] | 30.9 | 1.0E-18 |
| DIP_10006417 | putative coatomer subunit protein [Eutypa lata UCREL1] | 65.8 | 0.0 |
| DIP_10006418 | putative fructosyl-amino acid oxidase protein [Eutypa lata UCREL1] | 63 | 0.0 |
| DIP_10006419 | hypothetical protein FOXB_08755 [Fusarium oxysporum Fo5176] | 90.6 | 2.0E-166 |
| DIP_10006420 | hypothetical protein Z519_07164 [Cladophialophora bantiana CBS 173.52] | 66.3 | 0.0 |
| DIP_10006421 | hypothetical protein PFICI_06446 [Pestalotiopsis fici W106-1] | 46 | 0.0 |
| DIP_10006422 | NA | | |
| DIP_10006423 | hypothetical protein PFICI_13259 [Pestalotiopsis fici W106-1] | 54.2 | 0.0 |
| DIP_10006424 | putative regulator of gluconeogenesis protein [Eutypa lata UCREL1] | 54.3 | 9.0E-168 |
| DIP_10006425 | hypothetical protein PFICI_07002 [Pestalotiopsis fici W106-1] | 44.1 | 9.0E-108 |
| DIP_10006426 | hypothetical protein UCREL1_3556 [Eutypa lata UCREL1] | 68.3 | 1.0E-159 |
| DIP_10006427 | NA | | |
| DIP_10006428 | putative rad50 protein [Eutypa lata UCREL1] | 57.9 | 0.0 |
| DIP_10006429 | putative zinc finger white collar 2 protein wc-2 protein [Eutypa lata UCREL1] | 63.1 | 0.0 |
| DIP_10006430 | hypothetical protein PFICI_07005 [Pestalotiopsis fici W106-1] | 61.3 | 7.0E-138 |
| DIP_10006431 | hypothetical protein PFICI_07006 [Pestalotiopsis fici W106-1] | 62.5 | 3.0E-59 |
| DIP_10006432 | conserved hypothetical protein [Aspergillus terreus NIH2624] | 35.2 | 8.0E-53 |
| DIP_10006433 | NA | | |
| DIP_10006434 | putative sorbitol dehydrogenase protein [Eutypa lata UCREL1] | 81.5 | 0.0 |
| DIP_10006435 | hypothetical protein PFICI_06193 [Pestalotiopsis fici W106-1] | 61.4 | 4.0E-88 |
| DIP_10006436 | hypothetical protein PFICI_06194 [Pestalotiopsis fici W106-1] | 81.9 | 1.0E-38 |
| DIP_10006437 | naringenin 3-dioxygenase [Verticillium dahliae VdLs.17] | 71.9 | 0.0 |
| DIP_10006438 | putative atp synthase subunit g protein [Eutypa lata UCREL1] | 62.6 | 2.0E-90 |
| DIP_10006439 | hypothetical protein V492_05929 [Pseudogymnoascus pannorum VKM F-4246] | 42.8 | 6.0E-79 |
| DIP_10006440 | NA | | |
| DIP_10006441 | putative transcription factor tfiia complex subunit protein [Eutypa lata UCREL1] | 54.8 | 9.0E-114 |
| DIP_10006442 | putative wd repeat-containing protein slp1 protein [Togninia minima UCRPA7] | 83.2 | 0.0 |
| DIP_10006443 | hypothetical protein PFICI_06907 [Pestalotiopsis fici W106-1] | 76 | 0.0 |
| DIP_10006444 | NA | | |
| DIP_10006445 | putative c2h2 transcription factor protein [Eutypa lata UCREL1] | 73.1 | 0.0 |
| DIP_10006446 | hypothetical protein PFICI_06905 [Pestalotiopsis fici W106-1] | 68.3 | 5.0E-124 |
| DIP_10006447 | hypothetical protein PFICI_13665 [Pestalotiopsis fici W106-1] | 59.6 | 0.0 |
| DIP_10006448 | putative trna-dihydrouridine synthase 3 protein [Eutypa lata UCREL1] | 69.5 | 0.0 |
| DIP_10006449 | putative endoplasmic reticulum transmembrane protein [Eutypa lata UCREL1] | 82.8 | 1.0E-126 |
| DIP_10006450 | hypothetical protein OIDMADRAFT_116903 [Oidiodendron maius Zn] | 56.3 | 3.0E-157 |
| DIP_10006451 | hypothetical protein UCREL1_958 [Eutypa lata UCREL1] | 35.6 | 6.0E-54 |
| DIP_10006452 | hypothetical protein PFICI_13266 [Pestalotiopsis fici W106-1] | 44.7 | 0.0 |
| DIP_10006453 | putative rna recognition domain-containing protein [Eutypa lata UCREL1] | 72.8 | 0.0 |
| DIP_10006454 | putative dynein light chain protein [Eutypa lata UCREL1] | 86.9 | 4.0E-63 |
| DIP_10006455 | putative csm1 regulatory protein [Eutypa lata UCREL1] | 50.3 | 5.0E-176 |
| DIP_10006456 | NA | | |
| DIP_10006457 | hypothetical protein PFICI_13271 [Pestalotiopsis fici W106-1] | 54.7 | 1.0E-124 |
| DIP_10006458 | hypothetical protein OIDMADRAFT_64233, partial [Oidiodendron maius Zn] | 57.6 | 1.0E-107 |
| DIP_10006459 | NA | | |
| DIP_10006460 | NA | | |
| DIP_10006461 | NA | | |
| DIP_10006462 | hypothetical protein PFICI_06198 [Pestalotiopsis fici W106-1] | 58.7 | 0.0 |
| DIP_10006463 | putative dna-directed rna polymerases i ii iii subunit 10 protein [Eutypa lata UCREL1] | 89.6 | 4.0E-47 |
| DIP_10006464 | putative wd domain-containing protein [Eutypa lata UCREL1] | 53.5 | 0.0 |
| DIP_10006465 | putative pig-p domain-containing protein [Eutypa lata UCREL1] | 68.9 | 2.0E-81 |
| DIP_10006466 | hypothetical protein FOIG_13574 [Fusarium oxysporum f. sp. cubense tropical race 4 54006] | 31.9 | 1.0E-06 |
| DIP_10006467 | MADS-box MEF2 type transcription factor [Gaeumannomyces graminis var. tritici R3-111a-1] | 45.4 | 1.0E-150 |
| DIP_10006468 | NA | | |
| DIP_10006469 | putative transcription initiation factor tfiid subunit 1 protein [Eutypa lata UCREL1] | 64 | 0.0 |
| DIP_10006470 | hypothetical protein PFICI_06923 [Pestalotiopsis fici W106-1] | 55.8 | 0.0 |
| DIP_10006471 | putative actin-related arp4 class protein [Togninia minima UCRPA7] | 74.6 | 0.0 |
| DIP_10006472 | NA | | |
| DIP_10006473 | hypothetical protein SAPIO_CDS2471 [Scedosporium apiospermum] | 59.8 | 1.0E-141 |
| DIP_10006474 | conserved hypothetical protein [Talaromyces stipitatus ATCC 10500] | 38.8 | 2.0E-163 |
| DIP_10006475 | NA | | |
| DIP_10006476 | hypothetical protein HMPREF1624_04878 [Sporothrix schenckii ATCC 58251] | 52.3 | 2.0E-174 |
| DIP_10006477 | hypothetical protein PV07_02025 [Cladophialophora immunda] | 46.6 | 1.0E-107 |
| DIP_10006478 | putative f-box protein [Togninia minima UCRPA7] | 44.1 | 2.0E-159 |
| DIP_10006479 | hypothetical protein PFICI_06917 [Pestalotiopsis fici W106-1] | 70.2 | 0.0 |
| DIP_10006480 | putative uracil-dna glycosylase protein [Eutypa lata UCREL1] | 71.1 | 4.0E-163 |
| DIP_10006481 | hypothetical protein PFICI_06919 [Pestalotiopsis fici W106-1] | 64.3 | 0.0 |
| DIP_10006482 | hypothetical protein TRIVIDRAFT_150386 [Trichoderma virens Gv29-8] | 30.2 | 1.0E-64 |
| DIP_10006483 | hypothetical protein S40285_03522 [Stachybotrys chlorohalonata IBT 40285] | 42.4 | 9.0E-78 |
| DIP_10006484 | hypothetical protein THITE_2077439 [Thielavia terrestris NRRL 8126] | 59.7 | 0.0 |
| DIP_10006485 | hypothetical protein F503_05329 [Ophiostoma piceae UAMH 11346] | 47.5 | 6.0E-73 |
| DIP_10006486 | NA | | |
| DIP_10006487 | hypothetical protein PFICI_02080 [Pestalotiopsis fici W106-1] | 46.9 | 2.0E-143 |
| DIP_10006488 | hypothetical protein A1O3_05682 [Capronia epimyces CBS 606.96] | 50.9 | 2.0E-110 |
| DIP_10006489 | hypothetical protein NEUTE1DRAFT_87143 [Neurospora tetrasperma FGSC 2508] | 67.3 | 5.0E-119 |
| DIP_10006490 | hypothetical protein PFICI_10290 [Pestalotiopsis fici W106-1] | 33.2 | 6.0E-31 |
| DIP_10006491 | putative phytanoyl- dioxygenase protein [Eutypa lata UCREL1] | 50.6 | 2.0E-94 |
| DIP_10006492 | putative glycoside hydrolase family 55 protein [Eutypa lata UCREL1] | 63.6 | 0.0 |
| DIP_10006493 | NA | | |
| DIP_10006494 | hypothetical protein ANO11243_049760 [fungal sp. No.11243] | 34.5 | 1.0E-29 |
| DIP_10006495 | hypothetical protein OOU_Y34scaffold00099g19 [Magnaporthe oryzae Y34] | 58 | 0.0 |
| DIP_10006496 | hypothetical protein PFICI_10402 [Pestalotiopsis fici W106-1] | 67.5 | 1.0E-144 |
| DIP_10006497 | hypothetical protein PFICI_14298 [Pestalotiopsis fici W106-1] | 59.1 | 0.0 |
| DIP_10006498 | hypothetical protein P875_00117374 [Aspergillus parasiticus SU-1] | 79 | 5.0E-111 |
| DIP_10006499 | putative duf1237 domain-containing protein [Eutypa lata UCREL1] | 66.9 | 0.0 |
| DIP_10006500 | O-methyltransferase, putative [Talaromyces marneffei ATCC 18224] | 48.5 | 2.0E-132 |
| DIP_10006501 | hypothetical protein PFICI_14296 [Pestalotiopsis fici W106-1] | 62.4 | 0.0 |
| DIP_10006502 | hypothetical protein PFICI_13700 [Pestalotiopsis fici W106-1] | 63.4 | 0.0 |
| DIP_10006503 | fungal specific transcription factor domain-containing protein [Marssonina brunnea f. sp. 'multigermtubi' MB_m1] | 37.8 | 1.0E-110 |
| DIP_10006504 | similar to cytochrome P450 alkane hydroxylase [Botrytis cinerea T4] | 39.2 | 6.0E-129 |
| DIP_10006505 | hypothetical protein PFICI_13666 [Pestalotiopsis fici W106-1] | 31.8 | 3.0E-31 |
| DIP_10006506 | NA | | |
| DIP_10006507 | NA | | |
| DIP_10006508 | hypothetical protein PFICI_13667 [Pestalotiopsis fici W106-1] | 75.6 | 0.0 |
| DIP_10006509 | hypothetical protein PFICI_13668 [Pestalotiopsis fici W106-1] | 50.3 | 1.0E-95 |
| DIP_10006510 | putative ercc4 domain-containing protein [Eutypa lata UCREL1] | 51.3 | 3.0E-136 |
| DIP_10006511 | putative thiazole biosynthetic enzyme protein [Eutypa lata UCREL1] | 73.9 | 1.0E-175 |
| DIP_10006512 | hypothetical protein NEUTE1DRAFT_45744 [Neurospora tetrasperma FGSC 2508] | 75.4 | 1.0E-70 |
| DIP_10006513 | hypothetical protein PFICI_09190 [Pestalotiopsis fici W106-1] | 53.5 | 5.0E-98 |
| DIP_10006514 | hypothetical protein PFICI_13915 [Pestalotiopsis fici W106-1] | 44.5 | 4.0E-35 |
| DIP_10006515 | NA | | |
| DIP_10006516 | hypothetical protein H072_1043 [Dactylellina haptotyla CBS 200.50] | 56.1 | 6.0E-155 |
| DIP_10006517 | NA | | |
| DIP_10006518 | hypothetical protein UCREL1_6370 [Eutypa lata UCREL1] | 46.7 | 1.0E-24 |
| DIP_10006519 | hypothetical protein UCREL1_6369 [Eutypa lata UCREL1] | 34.6 | 6.0E-09 |
| DIP_10006520 | hypothetical protein CH063_01898 [Colletotrichum higginsianum] | 40.3 | 2.0E-34 |
| DIP_10006521 | hypothetical protein UCREL1_6368 [Eutypa lata UCREL1] | 47.4 | 5.0E-55 |
| DIP_10006522 | hypothetical protein F503_08106 [Ophiostoma piceae UAMH 11346] | 34.7 | 6.0E-12 |
| DIP_10006523 | hypothetical protein SPBR_03796 [Sporothrix brasiliensis 5110] | 31.7 | 1.0E-10 |
| DIP_10006524 | putative vesicle-fusing atpase protein [Eutypa lata UCREL1] | 64.8 | 0.0 |
| DIP_10006525 | hypothetical protein FOVG_18506 [Fusarium oxysporum f. sp. pisi HDV247] | 31.2 | 2.0E-10 |
| DIP_10006526 | Aldehyde dehydrogenase family protein [Aspergillus flavus AF70] | 73.4 | 0.0 |
| DIP_10006527 | hypothetical protein P034_00512841 [Aspergillus flavus AF70] | 38.7 | 4.0E-58 |
| DIP_10006528 | hypothetical protein Moror_11332 [Moniliophthora roreri MCA 2997] | 30.5 | 2.0E-06 |
| DIP_10006529 | putative killer kp4 protein [Eutypa lata UCREL1] | 38.3 | 8.0E-16 |
| DIP_10006530 | putative killer kp4 protein [Eutypa lata UCREL1] | 39.1 | 1.0E-16 |
| DIP_10006531 | putative dihydrodipicolinate synthetase protein [Eutypa lata UCREL1] | 71.6 | 7.0E-140 |
| DIP_10006532 | aspartate aminotransferase [Byssochlamys spectabilis No. 5] | 58.1 | 3.0E-176 |
| DIP_10006533 | putative proline oxidase [Diaporthe ampelina] | 61.6 | 0.0 |
| DIP_10006534 | hypothetical protein PFICI_09742 [Pestalotiopsis fici W106-1] | 47 | 0.0 |
| DIP_10006535 | putative pyrroline-5-carboxylate reductase [Diaporthe ampelina] | 46.4 | 1.0E-80 |
| DIP_10006536 | hypothetical protein NECHADRAFT_15955 [Nectria haematococca mpVI 77-13-4] | 58 | 0.0 |
| DIP_10006537 | NA | | |
| DIP_10006538 | hypothetical protein PFICI_02612 [Pestalotiopsis fici W106-1] | 45.3 | 0.0 |
| DIP_10006539 | putative kinase binding protein cgi-121 protein [Eutypa lata UCREL1] | 65.8 | 7.0E-97 |
| DIP_10006540 | hypothetical protein PFICI_02540 [Pestalotiopsis fici W106-1] | 54.8 | 0.0 |
| DIP_10006541 | hypothetical protein PFICI_02541 [Pestalotiopsis fici W106-1] | 72.3 | 3.0E-73 |
| DIP_10006542 | conserved hypothetical protein [Verticillium alfalfae VaMs.102] | 47.4 | 1.0E-35 |
| DIP_10006543 | hypothetical protein PFICI_02544 [Pestalotiopsis fici W106-1] | 60.6 | 0.0 |
| DIP_10006544 | hypothetical protein MGG_14899 [Magnaporthe oryzae 70-15] | 40.5 | 4.0E-120 |
| DIP_10006545 | putative mfs quinate transporter protein [Eutypa lata UCREL1] | 70.9 | 0.0 |
| DIP_10006546 | NA | | |
| DIP_10006547 | hypothetical protein PFICI_13689 [Pestalotiopsis fici W106-1] | 45.8 | 5.0E-48 |
| DIP_10006548 | NA | | |
| DIP_10006549 | NA | | |
| DIP_10006550 | hypothetical protein CGLO_15908 [Colletotrichum gloeosporioides Cg-14] | 52.5 | 0.0 |
| DIP_10006551 | hypothetical protein GGTG_05621 [Gaeumannomyces graminis var. tritici R3-111a-1] | 43.4 | 1.0E-26 |
| DIP_10006552 | NA | | |
| [illegible] | [illegible] | [illegible] | [illegible] |

| | | | |
|---|---|---|---|
| DIP_10006586 | putative short-chain dehydrogenase [Colletotrichum sublineola] | 64.2 | 7.0E-117 |
| DIP_10006587 | putative amidase protein [Eutypa lata UCREL1] | 64.1 | 0.0 |
| DIP_10006588 | Alcohol dehydrogenase GroES-like domain-containing protein [Scedosporium apiospermum] | 50.8 | 7.0E-127 |
| DIP_10006589 | polysaccharide deacetylase family protein [Colletotrichum gloeosporioides Nara gc5] | 68.9 | 4.0E-152 |
| DIP_10006590 | putative transmembrane amino acid transporter protein [Eutypa lata UCREL1] | 75.4 | 0.0 |
| DIP_10006591 | Nucleolar protein 58 [Pestalotiopsis fici W106-1] | 76.4 | 0.0 |
| DIP_10006592 | vacuolar ATP synthase subunit d [Grosmannia clavigera kw1407] | 89.6 | 0.0 |
| DIP_10006593 | hypothetical protein PFICI_09168 [Pestalotiopsis fici W106-1] | 57 | 2.0E-112 |
| DIP_10006594 | hypothetical protein FPSE_09799 [Fusarium pseudograminearum CS3096] | 79.7 | 2.0E-169 |
| DIP_10006595 | hypothetical protein S7711_09064 [Stachybotrys chartarum IBT 7711] | 62.6 | 6.0E-33 |
| DIP_10006596 | hypothetical protein PFICI_02610 [Pestalotiopsis fici W106-1] | 60.2 | 0.0 |
| DIP_10006597 | Protein transport protein Sec61 subunit beta [Hirsutella minnesotensis 3608] | 86.7 | 8.0E-55 |
| DIP_10006598 | manganese ion homeostasis [Colletotrichum gloeosporioides Nara gc5] | 44.2 | 2.0E-162 |
| DIP_10006599 | glucose transporter rco-3 [Pestalotiopsis fici W106-1] | 72 | 0.0 |
| DIP_10006600 | NA | | |
| DIP_10006601 | hypothetical protein PFICI_06189 [Pestalotiopsis fici W106-1] | 90.5 | 0.0 |
| DIP_10006602 | hypothetical protein CSUB01_02918 [Colletotrichum sublineola] | 59.6 | 1.0E-73 |
| DIP_10006603 | NA | | |
| DIP_10006604 | hypothetical protein S40288_07505 [Stachybotrys chartarum IBT 40288] | 49.8 | 2.0E-63 |
| DIP_10006605 | NA | | |
| DIP_10006606 | putative short-chain dehydrogenase reductase protein [Eutypa lata UCREL1] | 54.4 | 1.0E-97 |
| DIP_10006607 | NA | | |
| DIP_10006608 | putative alcohol dehydrogenase protein [Eutypa lata UCREL1] | 68.3 | 2.0E-176 |
| DIP_10006609 | hypothetical protein PFICI_02550 [Pestalotiopsis fici W106-1] | 58 | 1.0E-69 |
| DIP_10006610 | hypothetical protein CHGG_02287 [Chaetomium globosum CBS 148.51] | 57.3 | 0.0 |
| DIP_10006611 | hypothetical protein PFICI_02548 [Pestalotiopsis fici W106-1] | 72.2 | 0.0 |
| DIP_10006612 | hypothetical protein EMCG_07456 [Emmonsia crescens UAMH 3008] | 43.9 | 2.0E-99 |
| DIP_10006613 | hypothetical protein PFICI_02615 [Pestalotiopsis fici W106-1] | 53.2 | 4.0E-116 |
| DIP_10006614 | putative disulfide isomerase protein [Eutypa lata UCREL1] | 58.3 | 0.0 |
| DIP_10006615 | exosome complex exonuclease RRP41 [Magnaporthe oryzae 70-15] | 65.8 | 2.0E-131 |
| DIP_10006616 | putative 3 exoribonuclease family protein [Eutypa lata UCREL1] | 61.2 | 1.0E-123 |
| DIP_10006617 | Mitochondrial import inner membrane translocase subunit TIM50 [Pestalotiopsis fici W106-1] | 65.7 | 0.0 |
| DIP_10006618 | hypothetical protein TRIVIDRAFT_31506 [Trichoderma virens Gv29-8] | 57.7 | 1.0E-173 |
| DIP_10006619 | hypothetical protein FOCG_13486 [Fusarium oxysporum f. sp. radicis-lycopersici 26381] | 34.7 | 8.0E-96 |
| DIP_10006620 | putative phthalate transporter protein [Eutypa lata UCREL1] | 59.2 | 1.0E-135 |
| DIP_10006621 | NA | | |
| DIP_10006622 | NA | | |
| DIP_10006623 | hypothetical protein CPC735_061790 [Byssochlamys spectabilis No. 5] | 52.2 | 4.0E-112 |
| DIP_10006624 | putative allantoate permease protein [Eutypa lata UCREL1] | 80.9 | 0.0 |
| DIP_10006625 | putative dna polymerase delta protein [Togninia minima UCRPA7] | 48.8 | 1.0E-61 |
| DIP_10006626 | putative upf0553 protein c protein [Togninia minima UCRPA7] | 68 | 0.0 |
| DIP_10006627 | putative family protein [Eutypa lata UCREL1] | 68.9 | 0.0 |
| DIP_10006628 | putative 3-oxoacyl-(acyl-carrier-protein) reductase protein [Eutypa lata UCREL1] | 51.7 | 2.0E-75 |
| DIP_10006629 | putative long-chain base protein 3 protein [Togninia minima UCRPA7] | 62.4 | 0.0 |
| DIP_10006630 | NA | | |
| DIP_10006631 | hypothetical protein FOZG_17149 [Fusarium oxysporum Fo47] | 35.5 | 4.0E-18 |
| DIP_10006632 | hypothetical protein PFICI_11335 [Pestalotiopsis fici W106-1] | 58.2 | 0.0 |
| DIP_10006633 | putative iq calmodulin-binding motif domain protein [Eutypa lata UCREL1] | 70.9 | 0.0 |
| DIP_10006634 | DNA repair protein [Colletotrichum graminicola M1.001] | 71.9 | 0.0 |
| DIP_10006635 | hypothetical protein PFICI_08728 [Pestalotiopsis fici W106-1] | 57.5 | 3.0E-109 |
| DIP_10006636 | putative zinc metalloproteinase protein [Eutypa lata UCREL1] | 56.9 | 8.0E-132 |
| DIP_10006637 | putative er lumen protein retaining receptor protein [Togninia minima UCRPA7] | 83.3 | 4.0E-134 |
| DIP_10006638 | NA | | |
| DIP_10006639 | hypothetical protein SNOG_06459 [Phaeosphaeria nodorum SN15] | 39.5 | 2.0E-10 |
| DIP_10006640 | capsule polysaccharide synthase [Talaromyces cellulolyticus] | 36.6 | 5.0E-90 |
| DIP_10006641 | NA | | |
| DIP_10006642 | NA | | |
| DIP_10006643 | putative udp- transferase protein [Eutypa lata UCREL1] | 40.4 | 9.0E-27 |
| DIP_10006644 | putative mitochondrial distribution and morphology protein 38 protein [Eutypa lata UCREL1] | 73.1 | 0.0 |
| DIP_10006645 | hypothetical protein UCREL1_5651 [Eutypa lata UCREL1] | 55.7 | 0.0 |
| DIP_10006646 | hypothetical protein PFICI_14295 [Pestalotiopsis fici W106-1] | 63.8 | 0.0 |
| DIP_10006647 | NA | | |
| DIP_10006648 | hypothetical protein A1O1_00271 [Capronia coronata CBS 617.96] | 54.8 | 4.0E-132 |
| DIP_10006649 | hypothetical protein PFICI_12004 [Pestalotiopsis fici W106-1] | 58.4 | 0.0 |
| DIP_10006650 | putative nad-binding rossmann fold oxidoreductase family protein [Eutypa lata UCREL1] | 83.7 | 0.0 |
| DIP_10006651 | NADH-cytochrome b5 reductase 2 [Pestalotiopsis fici W106-1] | 73.1 | 5.0E-175 |
| DIP_10006652 | hypothetical protein PFICI_13696 [Pestalotiopsis fici W106-1] | 52 | 0.0 |
| DIP_10006653 | NA | | |
| DIP_10006654 | putative d-lactate dehydrogenase protein [Eutypa lata UCREL1] | 79.3 | 0.0 |
| DIP_10006655 | hypothetical protein CH063_02427 [Colletotrichum higginsianum] | 50.9 | 2.0E-142 |
| DIP_10006656 | putative spry domain-containing protein [Eutypa lata UCREL1] | 65 | 6.0E-122 |
| DIP_10006657 | NA | | |
| DIP_10006658 | hypothetical protein UCREL1_1646 [Eutypa lata UCREL1] | 42.9 | 9.0E-128 |
| DIP_10006659 | conserved hypothetical protein [Chaetomium globosum CBS 148.51] | 81.5 | 7.0E-104 |
| DIP_10006660 | putative cdk-activating kinase assembly factor mat1 protein [Eutypa lata UCREL1] | 54 | 3.0E-137 |
| DIP_10006661 | hypothetical protein PFICI_06844 [Pestalotiopsis fici W106-1] | 67.5 | 0.0 |
| DIP_10006662 | putative aspartyl-trna synthetase protein [Eutypa lata UCREL1] | 45.8 | 0.0 |
| DIP_10006663 | putative cytochrome p450 6a1 protein [Eutypa lata UCREL1] | 60.2 | 0.0 |
| DIP_10006664 | putative guanine nucleotide exchange factor lte1 protein [Eutypa lata UCREL1] | 64.1 | 2.0E-176 |
| DIP_10006665 | NA | | |
| DIP_10006666 | NA | | |
| DIP_10006667 | BTB/POZ domain-containing protein [Colletotrichum fioriniae PJ7] | 58.1 | 0.0 |
| DIP_10006668 | hypothetical protein PFICI_04214 [Pestalotiopsis fici W106-1] | 56.9 | 0.0 |
| DIP_10006669 | hypothetical protein UCREL1_106 [Eutypa lata UCREL1] | 55.6 | 0.0 |
| DIP_10006670 | putative duf1212 domain membrane protein prm10 protein [Eutypa lata UCREL1] | 62.5 | 0.0 |
| DIP_10006671 | putative perilipin mpl1-like protein [Eutypa lata UCREL1] | 64.1 | 8.0E-86 |
| DIP_10006672 | probable heterokaryon incompatibility Het-C protein [Fusarium fujikuroi IMI 58289] | 64.5 | 0.0 |
| DIP_10006673 | putative c6 zinc finger domain containing protein [Eutypa lata UCREL1] | 32.4 | 4.0E-27 |
| DIP_10006674 | endoglucanase [Clonostachys rosea] | 43 | 4.0E-60 |
| DIP_10006675 | putative serine threonine protein kinase protein [Eutypa lata UCREL1] | 66.4 | 0.0 |
| DIP_10006676 | hypothetical protein PFICI_09050 [Pestalotiopsis fici W106-1] | 53.4 | 0.0 |
| DIP_10006677 | putative intracellular protein transport protein [Eutypa lata UCREL1] | 60.4 | 0.0 |
| DIP_10006678 | putative 5 -3 exoribonuclease 2 protein [Eutypa lata UCREL1] | 78.6 | 0.0 |
| DIP_10006679 | hypothetical protein THITE_2124231 [Thielavia terrestris NRRL 8126] | 60.2 | 0.0 |
| DIP_10006680 | putative a g-specific adenine glycosylase protein [Eutypa lata UCREL1] | 47.5 | 3.0E-110 |
| DIP_10006681 | NA | | |
| DIP_10006682 | NA | | |
| DIP_10006683 | NA | | |
| DIP_10006684 | hypothetical protein PFICI_02594 [Pestalotiopsis fici W106-1] | 55 | 2.0E-67 |
| DIP_10006685 | putative golgi to endosome transport protein [Togninia minima UCRPA7] | 63.7 | 0.0 |
| DIP_10006686 | putative peptidyl-prolyl cis-trans isomerase protein [Eutypa lata UCREL1] | 90.1 | 5.0E-121 |
| DIP_10006687 | putative dna-directed rna polymerase iii subunit rpc6 protein [Eutypa lata UCREL1] | 59.3 | 3.0E-152 |
| DIP_10006688 | hypothetical protein THITE_2052956 [Thielavia terrestris NRRL 8126] | 33.5 | 7.0E-29 |
| DIP_10006689 | hypothetical protein SETTUDRAFT_156535 [Setosphaeria turcica Et28A] | 30.7 | 1.0E-34 |
| DIP_10006690 | hypothetical protein PFICI_11184 [Pestalotiopsis fici W106-1] | 63.1 | 7.0E-137 |
| DIP_10006691 | putative iron sulfur assembly protein 1 protein [Eutypa lata UCREL1] | 59 | 5.0E-99 |
| DIP_10006692 | hypothetical protein PFICI_08629 [Pestalotiopsis fici W106-1] | 76 | 0.0 |
| DIP_10006693 | putative indole-diterpene biosynthesis protein [Eutypa lata UCREL1] | 37.8 | 2.0E-63 |
| DIP_10006694 | NA | | |
| DIP_10006695 | hypothetical protein PFICI_09294 [Pestalotiopsis fici W106-1] | 50.5 | 0.0 |
| DIP_10006696 | hypothetical protein FOMG_10203 [Fusarium oxysporum f. sp. melonis 26406] | 38.9 | 2.0E-170 |
| DIP_10006697 | Fructose-2,6-bisphosphatase [Pestalotiopsis fici W106-1] | 84.3 | 0.0 |
| DIP_10006698 | NA | | |
| DIP_10006699 | ATP-dependent RNA helicase HAS1 [Fusarium verticillioides 7600] | 69.9 | 0.0 |
| DIP_10006700 | NA | | |
| DIP_10006701 | NA | | |
| DIP_10006702 | hypothetical protein G647_06610 [Cladophialophora carrionii CBS 160.54] | 51.1 | 2.0E-59 |
| DIP_10006703 | hypothetical protein PFICI_13680 [Pestalotiopsis fici W106-1] | 69 | 0.0 |
| DIP_10006704 | hypothetical protein PFICI_02635 [Pestalotiopsis fici W106-1] | 44.4 | 0.0 |
| DIP_10006705 | putative major facilitator superfamily transporter protein [Eutypa lata UCREL1] | 56.3 | 0.0 |
| DIP_10006706 | hypothetical protein S40285_01066 [Stachybotrys chlorohalonata IBT 40285] | 50.3 | 1.0E-178 |
| DIP_10006707 | hypothetical protein S7711_01288 [Stachybotrys chartarum IBT 7711] | 58.2 | 6.0E-171 |
| DIP_10006708 | putative camk protein kinase protein [Eutypa lata UCREL1] | 62.2 | 0.0 |
| DIP_10006709 | hypothetical protein PFICI_02568 [Pestalotiopsis fici W106-1] | 51.8 | 6.0E-85 |
| DIP_10006710 | hypothetical protein UCDDA912_g06664 [Diaporthe ampelina] | 44.2 | 5.0E-19 |
| DIP_10006711 | hypothetical protein V496_00650 [Pseudogymnoascus pannorum VKM F-4515 (FW-2607)] | 54.3 | 7.0E-61 |
| DIP_10006712 | Alkaline proteinase-like protein [Acremonium chrysogenum ATCC 11550] | 45.1 | 6.0E-117 |
| DIP_10006713 | hypothetical protein UCREL1_9109 [Eutypa lata UCREL1] | 65.4 | 4.0E-99 |
| DIP_10006714 | hypothetical protein PFICI_13612 [Pestalotiopsis fici W106-1] | 68.1 | 0.0 |
| DIP_10006715 | hypothetical protein PFICI_13606 [Pestalotiopsis fici W106-1] | 63 | 0.0 |
| DIP_10006716 | NA | | |
| DIP_10006717 | putative cytochrome p450 protein [Eutypa lata UCREL1] | 41.2 | 2.0E-152 |
| DIP_10006718 | hypothetical protein ACRE_045670 [Acremonium chrysogenum ATCC 11550] | 45 | 4.0E-79 |
| DIP_10006719 | hypothetical protein PFICI_13603 [Pestalotiopsis fici W106-1] | 66.9 | 2.0E-177 |
| DIP_10006720 | NA | | |
| DIP_10006721 | putative 3-hydroxyacyl- dehydrogenase protein [Eutypa lata UCREL1] | 70.2 | 3.0E-133 |
| DIP_10006722 | hypothetical protein UCREL1_5636 [Eutypa lata UCREL1] | 79.1 | 3.0E-102 |
| DIP_10006723 | hypothetical protein PFICI_13662 [Pestalotiopsis fici W106-1] | 62.9 | 0.0 |
| DIP_10006724 | putative acetyl- acetyltransferase protein [Eutypa lata UCREL1] | 84.5 | 0.0 |
| DIP_10006725 | putative 116 kda u5 small nuclear ribonucleoprotein component protein [Eutypa lata UCREL1] | 76.4 | 0.0 |
| DIP_10006726 | NA | | |
| DIP_10006727 | hypothetical protein PFICI_13658 [Pestalotiopsis fici W106-1] | 62.9 | 0.0 |
| DIP_10006728 | hypothetical protein PFICI_13659 [Pestalotiopsis fici W106-1] | 67.8 | 8.0E-88 |
| DIP_10006729 | hypothetical protein PFICI_09619 [Pestalotiopsis fici W106-1] | 42 | 7.0E-169 |
| DIP_10006730 | hypothetical protein PFICI_09618 [Pestalotiopsis fici W106-1] | 49.7 | 0.0 |
| DIP_10006731 | NA | | |
| DIP_10006732 | probable deoxyribodipyrimidine photo-lyase PHR [Claviceps purpurea 20.1] | 49.3 | 0.0 |
| DIP_10006733 | 40S ribosomal protein S21 [Nectria haematococca mpVI 77-13-4] | 86.2 | 3.0E-54 |
| DIP_10006734 | putative structure-specific endonuclease subunit slx4 [Diaporthe ampelina] | 66.7 | 3.0E-46 |
| DIP_10006735 | hypothetical protein PFICI_03084 [Pestalotiopsis fici W106-1] | 71.1 | 0.0 |
| DIP_10006736 | NA | | |
| DIP_10006737 | NA | | |
| DIP_10006738 | putative c6 zinc finger domain-containing protein [Eutypa lata UCREL1] | 47.1 | 3.0E-94 |
| DIP_10006739 | putative duf726 domain-containing protein [Eutypa lata UCREL1] | 66.1 | 0.0 |
| DIP_10006740 | NA | | |
| DIP_10006741 | hypothetical protein PFICI_03077 [Pestalotiopsis fici W106-1] | 49.7 | 0.0 |
| DIP_10006742 | putative peptide methionine sulfoxide reductase msrb msra protein [Eutypa lata UCREL1] | 73.7 | 2.0E-111 |
| DIP_10006743 | putative bax inhibitor family protein [Eutypa lata UCREL1] | 78.7 | 0.0 |
| DIP_10006744 | NA | | |
| DIP_10006745 | hypothetical protein PFICI_03074 [Pestalotiopsis fici W106-1] | 72.6 | 0.0 |
| DIP_10006746 | hypothetical protein PFICI_03075 [Pestalotiopsis fici W106-1] | 58.7 | 6.0E-155 |
| DIP_10006747 | putative regulator of g-protein signaling domain-containing protein [Eutypa lata UCREL1] | 82.7 | 0.0 |
| DIP_10006748 | 60S ribosomal protein L31 [Zymoseptoria tritici IPO323] | 81.8 | 5.0E-71 |
| DIP_10006749 | Ubiquitin-conjugating enzyme [Pestalotiopsis fici W106-1] | 85.6 | 1.0E-111 |
| DIP_10006750 | hypothetical protein PFICI_13672 [Pestalotiopsis fici W106-1] | 72.1 | 0.0 |
| DIP_10006751 | alpha-ketoglutarate dependent xanthine dioxygenase [Colletotrichum orbiculare MAFF 240422] | 56.5 | 4.0E-148 |
| DIP_10006752 | NA | | |
| DIP_10006753 | NA | | |
| DIP_10006754 | NA | | |
| DIP_10006755 | putative ankyrin domain protein [Eutypa lata UCREL1] | 47.3 | 7.0E-90 |
| DIP_10006756 | LolM [Epichloe aotearoae] | 38.3 | 7.0E-47 |
| DIP_10006757 | NA | | |
| DIP_10006758 | NA | | |
| DIP_10006759 | putative camk camk1 protein kinase protein [Eutypa lata UCREL1] | 77 | 0.0 |
| DIP_10006760 | arginine metabolism regulation protein iii [Colletotrichum gloeosporioides Nara gc5] | 33.3 | 7.0E-36 |
| DIP_10006761 | putative sodium hydrogen exchanger family protein [Eutypa lata UCREL1] | 69.1 | 0.0 |
| DIP_10006762 | hypothetical protein PFICI_13633 [Pestalotiopsis fici W106-1] | 56.6 | 3.0E-111 |
| DIP_10006763 | NA | | |
| DIP_10006764 | hypothetical protein PFICI_13589 [Pestalotiopsis fici W106-1] | 56.8 | 0.0 |
| DIP_10006765 | NA | | |
| DIP_10006766 | hypothetical protein ACRE_087140 [Acremonium chrysogenum ATCC 11550] | 39.7 | 5.0E-14 |
| DIP_10006767 | putative chitin binding protein [Eutypa lata UCREL1] | 63.6 | 8.0E-88 |
| DIP_10006768 | hypothetical protein NECHADRAFT_51427 [Nectria haematococca mpVI 77-13-4] | 53.5 | 0.0 |
| DIP_10006769 | hypothetical protein GLRG_09975 [Colletotrichum graminicola M1.001] | 42.7 | 6.0E-71 |
| DIP_10006770 | NA | | |
| DIP_10006771 | hypothetical protein PFICI_11442 [Pestalotiopsis fici W106-1] | 65 | 0.0 |
| DIP_10006772 | putative gcd10p family protein [Eutypa lata UCREL1] | 60.3 | 0.0 |
| DIP_10006773 | putative arp2 3 complex 21 kda subunit protein [Eutypa lata UCREL1] | 88.6 | 1.0E-122 |
| DIP_10006774 | NA | | |
| DIP_10006775 | hypothetical protein S40293_04129 [Stachybotrys chartarum IBT 40293] | 30.8 | 7.0E-30 |
| DIP_10006776 | NA | | |
| DIP_10006777 | putative glycosyltransferase family 31 protein [Eutypa lata UCREL1] | 66.9 | 0.0 |
| DIP_10006778 | putative ph-response regulator protein pala rim20 protein [Eutypa lata UCREL1] | 70 | 0.0 |
| DIP_10006779 | putative hydroxymethylglutaryl- lyase protein [Eutypa lata UCREL1] | 65.5 | 3.0E-179 |
| DIP_10006780 | NA | | |
| DIP_10006781 | hypothetical protein CFIO01_08041 [Colletotrichum fioriniae PJ7] | 31.8 | 4.0E-08 |
| DIP_10006782 | putative aldose 1-epimerase protein [Eutypa lata UCREL1] | 72.8 | 0.0 |
| DIP_10006783 | putative vacuolar atp synthase subunit b protein [Eutypa lata UCREL1] | 92 | 0.0 |
| DIP_10006784 | Diphthamide biosynthesis protein 1 [Pestalotiopsis fici W106-1] | 80.5 | 0.0 |
| DIP_10006785 | NA | | |
| DIP_10006786 | hypothetical protein CGLO_13478 [Colletotrichum gloeosporioides Cg-14] | 45.4 | 0.0 |
| DIP_10006787 | NA | | |
| DIP_10006788 | hypothetical protein Z518_00043 [Rhinocladiella mackenziei CBS 650.93] | 46.1 | 2.0E-161 |
| DIP_10006789 | hypothetical protein FPSE_08489 [Fusarium pseudograminearum CS3096] | 40.8 | 1.0E-120 |
| DIP_10006790 | Serine/threonine-protein phosphatase ppe1 [Pestalotiopsis fici W106-1] | 81.1 | 0.0 |
| DIP_10006791 | NA | | |
| DIP_10006792 | putative dash complex subunit dad1 protein [Togninia minima UCRPA7] | 81.2 | 8.0E-42 |
| DIP_10006793 | putative gtp-binding protein rho4 protein [Eutypa lata UCREL1] | 74.1 | 1.0E-164 |
| DIP_10006794 | transcription elongation factor B, polypeptide 1 [Trichoderma harzianum] | 56.2 | 3.0E-34 |
| DIP_10006795 | putative wsc domain protein [Eutypa lata UCREL1] | 61.1 | 3.0E-156 |
| DIP_10006796 | putative lariat debranching enzyme protein [Togninia minima UCRPA7] | 64.2 | 0.0 |
| DIP_10006797 | NA | | |
| DIP_10006798 | hypothetical protein TCE0_044r16159 [Talaromyces cellulolyticus] | 46.8 | 2.0E-85 |
| DIP_10006799 | mannitol dehydrogenase, putative [Byssochlamys spectabilis No. 5] | 72.2 | 0.0 |
| DIP_10006800 | NA | | |
| DIP_10006801 | hypothetical protein V496_03190 [Pseudogymnoascus pannorum VKM F-4515 (FW-2607)] | 40.2 | 4.0E-136 |
| DIP_10006802 | putative atp-dependent rna helicase rok1 protein [Eutypa lata UCREL1] | 62.7 | 0.0 |
| DIP_10006803 | D-xylulose reductase A-like protein [Acremonium chrysogenum ATCC 11550] | 71.3 | 0.0 |
| DIP_10006804 | putative glycoside hydrolase family 2 protein [Eutypa lata UCREL1] | 47.3 | 0.0 |
| DIP_10006805 | Ras family protein [Colletotrichum higginsianum] | 74.7 | 7.0E-162 |
| DIP_10006806 | putative chromatin assembly factor 1 subunit b protein [Eutypa lata UCREL1] | 73.3 | 3.0E-174 |
| DIP_10006807 | hypothetical protein PFICI_06225 [Pestalotiopsis fici W106-1] | 55.9 | 0.0 |
| DIP_10006808 | NA | | |
| DIP_10006809 | NA | | |
| DIP_10006810 | hypothetical protein PFICI_11353 [Pestalotiopsis fici W106-1] | 63.9 | 0.0 |
| DIP_10006811 | hypothetical protein OOU_Y34scaffold00126g54 [Magnaporthe oryzae Y34] | 55 | 9.0E-19 |
| DIP_10006812 | NA | | |
| DIP_10006813 | hypothetical protein PFICI_11355 [Pestalotiopsis fici W106-1] | 66.6 | 1.0E-165 |
| DIP_10006814 | hypothetical protein VDAG_05358 [Verticillium dahliae VdLs.17] | 46.5 | 2.0E-111 |
| DIP_10006815 | Bifunctional solanapyrone synthase-like protein [Acremonium chrysogenum ATCC 11550] | 50.2 | 3.0E-174 |
| DIP_10006816 | hypothetical protein PFICI_14419 [Pestalotiopsis fici W106-1] | 55.6 | 8.0E-32 |
| DIP_10006817 | putative glycerophosphodiester phosphodiesterase gde1 protein [Eutypa lata UCREL1] | 70.6 | 0.0 |
| DIP_10006818 | NA | | |
| DIP_10006819 | hypothetical protein PFICI_11350 [Pestalotiopsis fici W106-1] | 32.2 | 9.0E-68 |
| DIP_10006820 | hypothetical protein FOQG_18203 [Fusarium oxysporum f. sp. raphani 54005] | 43.1 | 2.0E-157 |
| DIP_10006821 | hypothetical protein TD95_000249 [Thielaviopsis punctulata] | 71.9 | 4.0E-87 |
| DIP_10006822 | putative small nuclear ribonucleo protein [Eutypa lata UCREL1] | 90.9 | 9.0E-56 |
| DIP_10006823 | hypothetical protein PFICI_11347 [Pestalotiopsis fici W106-1] | 67.8 | 0.0 |
| DIP_10006824 | putative prefoldin subunit 4 protein [Eutypa lata UCREL1] | 83 | 1.0E-74 |
| DIP_10006825 | hypothetical protein PFICI_11345 [Pestalotiopsis fici W106-1] | 70.2 | 0.0 |
| DIP_10006826 | hypothetical protein PFICI_11352 [Pestalotiopsis fici W106-1] | 68.3 | 0.0 |
| DIP_10006827 | hypothetical protein PFICI_11356 [Pestalotiopsis fici W106-1] | 53.5 | 3.0E-144 |
| DIP_10006828 | NA | | |
| DIP_10006829 | NA | | |
| DIP_10006830 | NA | | |
| DIP_10006831 | putative extracellular protein [Eutypa lata UCREL1] | 55 | 8.0E-80 |
| DIP_10006832 | hypothetical protein FPSE_04728 [Fusarium pseudograminearum CS3096] | 51.1 | 0.0 |
| DIP_10006833 | hypothetical protein PFICI_08775 [Pestalotiopsis fici W106-1] | 36.6 | 9.0E-91 |
| DIP_10006834 | hypothetical protein PFICI_08774 [Pestalotiopsis fici W106-1] | 67.6 | 0.0 |
| DIP_10006835 | hypothetical protein PFICI_08783 [Pestalotiopsis fici W106-1] | 46.9 | 7.0E-72 |
| DIP_10006836 | hypothetical protein UCRPA7_5141 [Togninia minima UCRPA7] | 39.5 | 1.0E-48 |
| DIP_10006837 | Tubulin alpha-B chain [Pestalotiopsis fici W106-1] | 94.9 | 0.0 |
| DIP_10006838 | UBA/TS-N domain-containing protein [Beauveria bassiana ARSEF 2860] | 49.4 | 0.0 |
| DIP_10006839 | NA | | |
| DIP_10006840 | hypothetical protein UCREL1_7686 [Eutypa lata UCREL1] | 32.4 | 9.0E-18 |
| DIP_10006841 | hypothetical protein PFICI_06211 [Pestalotiopsis fici W106-1] | 60.3 | 0.0 |
| DIP_10006842 | NA | | |
| DIP_10006843 | putative epoxide hydrolase protein [Eutypa lata UCREL1] | 47.8 | 1.0E-114 |
| DIP_10006844 | putative rna-binding protein 8a protein [Eutypa lata UCREL1] | 89.7 | 3.0E-84 |
| DIP_10006845 | hypothetical protein PFICI_02085 [Pestalotiopsis fici W106-1] | 43.2 | 7.0E-36 |
| DIP_10006846 | hypothetical protein Z520_09639 [Fonsecaea multimorphosa CBS 102226] | 44.5 | 6.0E-129 |
| DIP_10006847 | unnamed protein product [Aspergillus oryzae RIB40] | 40.8 | 8.0E-92 |
| DIP_10006848 | hypothetical protein FGSG_03112 [Fusarium graminearum PH-1] | 47.3 | 3.0E-48 |
| DIP_10006849 | putative transmembrane and coiled-coil domain-containing protein [Eutypa lata UCREL1] | 57.9 | 0.0 |
| DIP_10006850 | putative 3-amino-3-carboxypropyl transferase protein [Eutypa lata UCREL1] | 70.4 | 0.0 |
| DIP_10006851 | putative alanyl-trna synthetase protein [Eutypa lata UCREL1] | 54.4 | 9.0E-103 |
| DIP_10006852 | putative vacuolar protein sorting-associated protein 28 protein [Eutypa lata UCREL1] | 82.6 | 1.0E-138 |
| DIP_10006853 | putative nucleolar complex protein 2 protein [Eutypa lata UCREL1] | 68.7 | 0.0 |
| DIP_10006854 | hypothetical protein NECHADRAFT_86687 [Nectria haematococca mpVI 77-13-4] | 44.3 | 1.0E-18 |
| DIP_10006855 | hypothetical protein MYCTH_2073132 [Myceliophthora thermophila ATCC 42464] | 56.1 | 7.0E-142 |
| DIP_10006856 | hypothetical protein PFICI_07160 [Pestalotiopsis fici W106-1] | 56.2 | 2.0E-123 |
| DIP_10006857 | putative glycoside hydrolase family 36 protein [Togninia minima UCRPA7] | 53.9 | 5.0E-118 |
| DIP_10006858 | putative mitochondrial-processing peptidase subunit beta protein [Eutypa lata UCREL1] | 87.9 | 0.0 |
| DIP_10006859 | putative metal tolerance protein 3 protein [Eutypa lata UCREL1] | 38.6 | 1.0E-21 |
| DIP_10006860 | aldehyde reductase [Trichoderma harzianum] | 51.2 | 1.0E-117 |
| DIP_10006861 | hypothetical protein PFICI_06289 [Pestalotiopsis fici W106-1] | 59.5 | 1.0E-109 |
| DIP_10006862 | hypothetical protein CSUB01_05169 [Colletotrichum sublineola] | 38.7 | 4.0E-106 |
| DIP_10006863 | hypothetical protein BAUCODRAFT_150372 [Baudoinia compniacensis UAMH 10762] | 46.2 | 3.0E-56 |
| DIP_10006864 | NA | | |
| DIP_10006865 | major facilitator superfamily transporter [Colletotrichum gloeosporioides Nara gc5] | 37.3 | 5.0E-106 |
| DIP_10006866 | hypothetical protein V499_07555 [Pseudogymnoascus pannorum VKM F-103] | 35.3 | 1.0E-27 |
| DIP_10006867 | 4,5-DOPA dioxygenase extradiol-like protein [Fusarium oxysporum f. sp. cubense race 4] | 52.5 | 8.0E-105 |
| DIP_10006868 | NA | | |
| DIP_10006869 | NA | | |
| DIP_10006870 | putative serine threonine-protein phosphatase dullard protein [Eutypa lata UCREL1] | 91.1 | 2.0E-131 |
| DIP_10006871 | NA | | |
| DIP_10006872 | putative ABC transporter protein [Chaetomium thermophilum var. thermophilum DSM 1495] | 69.2 | 0.0 |
| DIP_10006873 | NA | | |
| DIP_10006874 | hypothetical protein PFICI_00680 [Pestalotiopsis fici W106-1] | 73.5 | 0.0 |
| DIP_10006875 | hypothetical protein UCREL1_9746 [Eutypa lata UCREL1] | 39.6 | 3.0E-49 |
| DIP_10006876 | putative 26s proteasome regulatory subunit protein [Eutypa lata UCREL1] | 92 | 0.0 |
| DIP_10006877 | hypothetical protein UCREL1_9749 [Eutypa lata UCREL1] | 40.7 | 4.0E-21 |
| DIP_10006878 | hypothetical protein THITE_2120651 [Thielavia terrestris NRRL 8126] | 48.6 | 4.0E-66 |
| DIP_10006879 | putative histone deacetylase rpd3 protein [Eutypa lata UCREL1] | 77.2 | 0.0 |
| DIP_10006880 | 60S ribosomal protein L23 [Magnaporthe oryzae 70-15] | 100 | 8.0E-87 |
| DIP_10006881 | hypothetical protein UCREL1_9753 [Eutypa lata UCREL1] | 44.7 | 1.0E-153 |
| DIP_10006882 | hypothetical protein PFICI_13545 [Pestalotiopsis fici W106-1] | 54.5 | 5.0E-178 |
| DIP_10006883 | putative nad-dependent malic enzyme protein [Togninia minima UCRPA7] | 73 | 0.0 |
| DIP_10006884 | hypothetical protein PFICI_13532 [Pestalotiopsis fici W106-1] | 56.2 | 2.0E-45 |
| DIP_10006885 | hypothetical protein MYCTH_2307917 [Myceliophthora thermophila ATCC 42464] | 38.6 | 5.0E-135 |
| DIP_10006886 | hypothetical protein PFICI_13579 [Pestalotiopsis fici W106-1] | 52.9 | 0.0 |
| DIP_10006887 | hypothetical protein PFICI_13578 [Pestalotiopsis fici W106-1] | 41.2 | 2.0E-54 |
| DIP_10006888 | alpha-ketoglutarate-dependent taurine dioxygenase [Verticillium dahliae VdLs.17] | 68.1 | 0.0 |
| DIP_10006889 | NA | | |
| DIP_10006890 | putative n-acetyltransferase gcn5 protein [Togninia minima UCRPA7] | 45.4 | 4.0E-54 |
| DIP_10006891 | NA | | |
| DIP_10006892 | hypothetical protein UCREL1_7627 [Eutypa lata UCREL1] | 52.5 | 3.0E-153 |
| DIP_10006893 | hypothetical protein PFICI_14322 [Pestalotiopsis fici W106-1] | 91.1 | 5.0E-79 |
| DIP_10006894 | hypothetical protein A1O1_01673 [Capronia coronata CBS 617.96] | 68.7 | 0.0 |
| DIP_10006895 | hypothetical protein HMPREF1624_05337 [Sporothrix schenckii ATCC 58251] | 78 | 0.0 |
| DIP_10006896 | hypothetical protein PFICI_14365 [Pestalotiopsis fici W106-1] | 64 | 0.0 |
| DIP_10006897 | NA | | |
| DIP_10006898 | putative saf domain-containing protein [Eutypa lata UCREL1] | 88.7 | 0.0 |
| DIP_10006899 | putative mfs transporter protein [Eutypa lata UCREL1] | 68.6 | 0.0 |
| DIP_10006900 | hypothetical protein PFICI_06218 [Pestalotiopsis fici W106-1] | 40.8 | 0.0 |
| DIP_10006901 | hypothetical protein ANO11243_054780 [fungal sp. No.11243] | 83.8 | 4.0E-109 |
| DIP_10006902 | Protein CSN12 [Pestalotiopsis fici W106-1] | 76.1 | 0.0 |
| DIP_10006903 | predicted protein [Nectria haematococca mpVI 77-13-4] | 75.4 | 4.0E-141 |
| DIP_10006904 | hypothetical protein PFICI_05983 [Pestalotiopsis fici W106-1] | 76.8 | 0.0 |
| DIP_10006905 | Centromere protein V-like protein [Acremonium chrysogenum ATCC 11550] | 67.6 | 1.0E-67 |
| DIP_10006906 | hypothetical protein PFICI_05985 [Pestalotiopsis fici W106-1] | 36.9 | 9.0E-54 |
| DIP_10006907 | hypothetical protein FAVG1_06564 [Fusarium avenaceum] | 35.6 | 9.0E-89 |
| DIP_10006908 | hypothetical protein TRIATDRAFT_42240 [Trichoderma atroviride IMI 206040] | 55.7 | 1.0E-58 |
| DIP_10006909 | putative choline-sulfatase protein [Eutypa lata UCREL1] | 84 | 0.0 |
| DIP_10006910 | putative mannose-p-dolichol utilization defect 1 protein [Eutypa lata UCREL1] | 84.8 | 9.0E-152 |
| DIP_10006911 | NA | | |
| DIP_10006912 | hypothetical protein PFICI_07145 [Pestalotiopsis fici W106-1] | 55.1 | 6.0E-145 |
| DIP_10006913 | hypothetical protein CFO_g2496 [Ceratocystis platani] | 31.5 | 2.0E-81 |
| DIP_10006914 | putative aldose 1-epimerase protein [Eutypa lata UCREL1] | 67.7 | 8.0E-154 |
| DIP_10006915 | hypothetical protein CSUB01_11809 [Colletotrichum sublineola] | 67.7 | 2.0E-33 |
| DIP_10006916 | hypothetical protein PFICI_06228 [Pestalotiopsis fici W106-1] | 59.5 | 0.0 |
| DIP_10006917 | hypothetical protein ACRE_053950 [Acremonium chrysogenum ATCC 11550] | 52.9 | 9.0E-105 |
| DIP_10006918 | putative RNase P subunit p30 [Colletotrichum sublineola] | 55.5 | 2.0E-103 |
| DIP_10006919 | NA | | |
| DIP_10006920 | NA | | |
| DIP_10006921 | hypothetical protein PFICI_07149 [Pestalotiopsis fici W106-1] | 70.2 | 5.0E-49 |
| DIP_10006922 | hypothetical protein [Podospora anserina S mat+] | 75.5 | 0.0 |
| DIP_10006923 | hypothetical protein PFICI_05981 [Pestalotiopsis fici W106-1] | 59.4 | 1.0E-121 |
| DIP_10006924 | putative lysophospholipase plb1 protein [Eutypa lata UCREL1] | 64 | 0.0 |
| DIP_10006925 | hypothetical protein PFICI_05979 [Pestalotiopsis fici W106-1] | 46.3 | 0.0 |
| DIP_10006926 | putative cobalt uptake protein cot1 protein [Eutypa lata UCREL1] | 68 | 0.0 |
| DIP_10006927 | putative glycosyl hydrolase protein [Eutypa lata UCREL1] | 40.6 | 1.0E-76 |
| DIP_10006928 | putative nadix domain containing protein [Eutypa lata UCREL1] | 67.6 | 9.0E-77 |
| DIP_10006929 | hypothetical protein PFICI_15261 [Pestalotiopsis fici W106-1] | 54 | 1.0E-100 |
| DIP_10006930 | hypothetical protein UCREL1_9541 [Eutypa lata UCREL1] | 41.5 | 3.0E-23 |
| DIP_10006931 | hypothetical protein UCRPA7_5250 [Togninia minima UCRPA7] | 39 | 8.0E-31 |
| DIP_10006932 | NA | | |
| DIP_10006933 | putative glutathione s-transferase protein [Eutypa lata UCREL1] | 52.9 | 7.0E-95 |
| DIP_10006934 | putative 2og-fe oxygenase protein [Eutypa lata UCREL1] | 58.4 | 6.0E-123 |
| DIP_10006935 | hypothetical protein CFIO01_10899 [Colletotrichum fioriniae PJ7] | 32.8 | 1.0E-59 |
| DIP_10006936 | hypothetical protein UCREL1_9548 [Eutypa lata UCREL1] | 43.2 | 6.0E-25 |
| DIP_10006937 | hypothetical protein A1O9_11199 [Exophiala aquamarina CBS 119918] | 58.3 | 1.0E-85 |
| DIP_10006938 | putative arylsulfotransferase-like protein [Eutypa lata UCREL1] | 59.3 | 1.0E-89 |
| DIP_10006939 | hypothetical protein FGSG_03263 [Fusarium graminearum PH-1] | 66.3 | 7.0E-129 |
| DIP_10006940 | hypothetical protein PFICI_09836 [Pestalotiopsis fici W106-1] | 74.2 | 0.0 |
| DIP_10006941 | putative phospho-2-dehydro-3-deoxyheptonate aldolase protein [Eutypa lata UCREL1] | 83.1 | 0.0 |
| DIP_10006942 | hypothetical protein UCREL1_4760 [Eutypa lata UCREL1] | 57.9 | 6.0E-53 |
| DIP_10006943 | hypothetical protein UCREL1_4759 [Eutypa lata UCREL1] | 52.8 | 6.0E-92 |
| DIP_10006944 | hypothetical protein PFICI_10386 [Pestalotiopsis fici W106-1] | 51.9 | 9.0E-102 |
| DIP_10006945 | hypothetical protein UVI_039450 [Ustilaginoidea virens] | 37.8 | 2.0E-27 |
| DIP_10006946 | putative 60s ribosomal protein l13 protein [Eutypa lata UCREL1] | 77.8 | 9.0E-116 |
| DIP_10006947 | hypothetical protein PFICI_10371 [Pestalotiopsis fici W106-1] | 38.4 | 7.0E-145 |
| DIP_10006948 | putative dolichyl-phosphate mannosyltransferase polypeptide 3 protein [Eutypa lata UCREL1] | 84.8 | 2.0E-52 |
| DIP_10006949 | hypothetical protein PFICI_15281 [Pestalotiopsis fici W106-1] | 32.5 | 6.0E-62 |
| DIP_10006950 | putative oxidoreductase-like protein [Eutypa lata UCREL1] | 56.8 | 4.0E-113 |
| DIP_10006951 | putative hypoxanthine guanine phosphoribosyltransferase protein [Eutypa lata UCREL1] | 79.7 | 1.0E-121 |

| | | | |
|---|---|---|---|
| DIP_10006952 | hypothetical protein PFICI_08429 [Pestalotiopsis fici W106-1] | 45.6 | 6.0E-156 |
| DIP_10006953 | hypothetical protein UCREL1_2949 [Eutypa lata UCREL1] | 47.3 | 2.0E-107 |
| DIP_10006954 | 5-methyltetrahydropteroyltriglutamate-homocysteine methyltransferase [Colletotrichum fioriniae PJ7] | 79 | 0.0 |
| DIP_10006955 | hypothetical protein COCCADRAFT_3067 [Bipolaris zeicola 26-R-13] | 39.6 | 2.0E-37 |
| DIP_10006956 | hypothetical protein PFICI_10366 [Pestalotiopsis fici W106-1] | 66.2 | 0.0 |
| DIP_10006957 | NA | | |
| DIP_10006958 | hypothetical protein PFICI_10361 [Pestalotiopsis fici W106-1] | 83.5 | 0.0 |
| DIP_10006959 | FAD-dependent oxidoreductase-like enzyme [Metarhizium robertsii ARSEF 23] | 44.7 | 3.0E-59 |
| DIP_10006960 | leucine-rich repeat-containing protein 40 [Gaeumannomyces graminis var. tritici R3-111a-1] | 46.6 | 8.0E-122 |
| DIP_10006961 | NA | | |
| DIP_10006962 | hypothetical protein V494_01357 [Pseudogymnoascus pannorum VKM F-4513 (FW-928)] | 37.6 | 2.0E-18 |
| DIP_10006963 | NA | | |
| DIP_10006964 | hypothetical protein THAR02_08299 [Trichoderma harzianum] | 42.7 | 1.0E-11 |
| DIP_10006965 | hypothetical protein FAVG1_07534 [Fusarium avenaceum] | 42.8 | 0.0 |
| DIP_10006966 | putative malate l-lactate protein [Eutypa lata UCREL1] | 79.7 | 0.0 |
| DIP_10006967 | putative fumarylacetoacetate hydrolase protein [Eutypa lata UCREL1] | 71.8 | 2.0E-160 |
| DIP_10006968 | hypothetical protein UCREL1_11604 [Eutypa lata UCREL1] | 48 | 8.0E-30 |
| DIP_10006969 | NA | | |
| DIP_10006970 | hypothetical protein PFICI_09201 [Pestalotiopsis fici W106-1] | 62.1 | 2.0E-172 |
| DIP_10006971 | putative 5-methyltetrahydropteroyltriglutamate-homocysteine s-methyltransferase protein [Togninia minima UCRPA7] | 82.9 | 0.0 |
| DIP_10006972 | hypothetical protein [Podospora anserina S mat+] | 31.9 | 8.0E-09 |
| DIP_10006973 | hypothetical protein PFICI_08693 [Pestalotiopsis fici W106-1] | 75 | 0.0 |
| DIP_10006974 | duf974 domain-containing protein [Colletotrichum orbiculare MAFF 240422] | 63 | 2.0E-139 |
| DIP_10006975 | protein phosphatase inhibitor [Colletotrichum graminicola M1.001] | 50 | 8.0E-24 |
| DIP_10006976 | hypothetical protein PFICI_08713 [Pestalotiopsis fici W106-1] | 58 | 0.0 |
| DIP_10006977 | putative sad1 interacting factor 1 protein [Eutypa lata UCREL1] | 52.9 | 4.0E-88 |
| DIP_10006978 | hypothetical protein PFICI_08714 [Pestalotiopsis fici W106-1] | 72.6 | 0.0 |
| DIP_10006979 | hypothetical protein UV8b_734 [Ustilaginoidea virens] | 34.4 | 4.0E-15 |
| DIP_10006980 | hypothetical protein PFICI_08716 [Pestalotiopsis fici W106-1] | 45.6 | 4.0E-170 |
| DIP_10006981 | hypothetical protein UCREL1_5913 [Eutypa lata UCREL1] | 57.9 | 3.0E-94 |
| DIP_10006982 | putative denn domain-containing protein [Togninia minima UCRPA7] | 82.8 | 0.0 |
| DIP_10006983 | hypothetical protein UCREL1_4703 [Eutypa lata UCREL1] | 30.2 | 3.0E-20 |
| DIP_10006984 | Actin-like protein 3 [Metarhizium robertsii ARSEF 23] | 94.5 | 0.0 |
| DIP_10006985 | putative asparaginase family protein [Togninia minima UCRPA7] | 56.1 | 3.0E-166 |
| DIP_10006986 | putative rna polymerase ii transcription factor b subunit 4 protein [Eutypa lata UCREL1] | 62.1 | 2.0E-160 |
| DIP_10006987 | hypothetical protein PFICI_03683 [Pestalotiopsis fici W106-1] | 59.9 | 0.0 |
| DIP_10006988 | hypothetical protein PFICI_03681 [Pestalotiopsis fici W106-1] | 55.8 | 0.0 |
| DIP_10006989 | NA | | |
| DIP_10006990 | putative gamma-glutamyltranspeptidase protein [Eutypa lata UCREL1] | 54.3 | 0.0 |
| DIP_10006991 | NA | | |
| DIP_10006992 | hypothetical protein PFICI_04026 [Pestalotiopsis fici W106-1] | 44.4 | 3.0E-91 |
| DIP_10006993 | putative nuclear envelope protein [Eutypa lata UCREL1] | 56.4 | 0.0 |
| DIP_10006994 | putative snare domain-containing protein [Eutypa lata UCREL1] | 68.5 | 2.0E-109 |
| DIP_10006995 | putative mitochondrial export translocase oxa2 protein [Eutypa lata UCREL1] | 36 | 3.0E-66 |
| DIP_10006996 | putative alpha- -mannosyltransferase alg11 protein [Eutypa lata UCREL1] | 72.2 | 0.0 |
| DIP_10006997 | hypothetical protein PFICI_04327 [Pestalotiopsis fici W106-1] | 60 | 2.0E-111 |
| DIP_10006998 | hypothetical protein NECHADRAFT_89859 [Nectria haematococca mpVI 77-13-4] | 76.9 | 0.0 |
| DIP_10006999 | Zn(2)-Cys(6) zinc finger domain protein [Metarhizium robertsii] | 39.2 | 2.0E-95 |
| DIP_10007000 | putative cfem domain-containing protein [Eutypa lata UCREL1] | 31.7 | 3.0E-50 |
| DIP_10007001 | NA | | |
| DIP_10007002 | hypothetical protein X797_003556 [Metarhizium robertsii] | 71.6 | 7.0E-51 |
| DIP_10007003 | hypothetical protein [Chromobacterium sp. C-61] | 30.1 | 8.0E-25 |
| DIP_10007004 | hypothetical protein PFICI_12594 [Pestalotiopsis fici W106-1] | 65.2 | 2.0E-108 |
| DIP_10007005 | putative meiotic recombination protein rec14 protein [Eutypa lata UCREL1] | 63.6 | 3.0E-153 |
| DIP_10007006 | mitochondrial ribosomal protein subunit L32 [Neurospora crassa OR74A] | 65.6 | 8.0E-37 |
| DIP_10007007 | putative cell division control protein 10 protein [Eutypa lata UCREL1] | 94.6 | 0.0 |
| DIP_10007008 | putative calcium permease protein [Eutypa lata UCREL1] | 72.9 | 0.0 |
| DIP_10007009 | NA | | |
| DIP_10007010 | putative 26s protease regulatory subunit 8 protein [Eutypa lata UCREL1] | 98.7 | 0.0 |
| DIP_10007011 | putative kh domain-containing protein [Eutypa lata UCREL1] | 76.4 | 0.0 |
| DIP_10007012 | putative -beta-glucanosyltransferase gel2 protein [Eutypa lata UCREL1] | 65.1 | 0.0 |
| DIP_10007013 | NA | | |
| DIP_10007014 | putative chromosome-associated kinesin kif4 protein [Eutypa lata UCREL1] | 69.4 | 0.0 |
| DIP_10007015 | NA | | |
| DIP_10007016 | hypothetical protein PFICI_13677 [Pestalotiopsis fici W106-1] | 53.8 | 5.0E-75 |
| DIP_10007017 | hypothetical protein PFICI_13676 [Pestalotiopsis fici W106-1] | 44.4 | 5.0E-171 |
| DIP_10007018 | putative udp-glucoronosyl and udp-glucosyl transferase family protein [Eutypa lata UCREL1] | 36.6 | 1.0E-62 |
| DIP_10007019 | hypothetical protein [Podospora anserina S mat+] | 38.9 | 1.0E-18 |
| DIP_10007020 | hypothetical protein MYCTH_2306098 [Myceliophthora thermophila ATCC 42464] | 38.1 | 3.0E-24 |
| DIP_10007021 | hypothetical protein PFICI_11360 [Pestalotiopsis fici W106-1] | 67.1 | 0.0 |
| DIP_10007022 | NA | | |
| DIP_10007023 | NA | | |
| DIP_10007024 | hypothetical protein UCREL1_503 [Eutypa lata UCREL1] | 54.2 | 2.0E-94 |
| DIP_10007025 | hypothetical protein PFICI_07979 [Pestalotiopsis fici W106-1] | 46.9 | 0.0 |
| DIP_10007026 | NA | | |
| DIP_10007027 | NA | | |
| DIP_10007028 | hypothetical protein CH063_01843, partial [Colletotrichum higginsianum] | 34.6 | 3.0E-42 |
| DIP_10007029 | NA | | |
| DIP_10007030 | putative mfs sugar transporter protein [Eutypa lata UCREL1] | 67.4 | 0.0 |
| DIP_10007031 | putative conserved hypothetical protein [Colletotrichum sublineola] | 44 | 1.0E-149 |
| DIP_10007032 | hypothetical protein PFICI_09115 [Pestalotiopsis fici W106-1] | 43.7 | 1.0E-108 |
| DIP_10007033 | glucan 1,4-alpha-maltohexaosidase [Verticillium dahliae VdLs.17] | 46.7 | 0.0 |
| DIP_10007034 | NA | | |
| DIP_10007035 | NA | | |
| DIP_10007036 | hypothetical protein FAVG1_09026 [Fusarium avenaceum] | 61.6 | 0.0 |
| DIP_10007037 | NA | | |
| DIP_10007038 | NA | | |
| DIP_10007039 | hypothetical protein NECHADRAFT_25501 [Nectria haematococca mpVI 77-13-4] | 67.4 | 0.0 |
| DIP_10007040 | hypothetical protein HMPREF1624_05631 [Sporothrix schenckii ATCC 58251] | 56.9 | 0.0 |
| DIP_10007041 | putative benzoate 4-monooxygenase protein [Eutypa lata UCREL1] | 61 | 0.0 |
| DIP_10007042 | putative sda1 domain-containing protein [Eutypa lata UCREL1] | 73.1 | 0.0 |
| DIP_10007043 | putative cytochrome b2 protein [Eutypa lata UCREL1] | 69.8 | 0.0 |
| DIP_10007044 | putative hormone-sensitive lipase protein [Eutypa lata UCREL1] | 73.1 | 0.0 |
| DIP_10007045 | Acetylornithine aminotransferase [Pestalotiopsis fici W106-1] | 72.4 | 0.0 |
| DIP_10007046 | predicted protein [Nectria haematococca mpVI 77-13-4] | 73.5 | 0.0 |
| DIP_10007047 | hypothetical protein PFICI_09104 [Pestalotiopsis fici W106-1] | 46.6 | 0.0 |
| DIP_10007048 | Glutamate decarboxylase [Pestalotiopsis fici W106-1] | 75.3 | 0.0 |
| DIP_10007049 | putative run and tbc1 domain containing protein [Neofusicoccum parvum UCRNP2] | 34.4 | 4.0E-135 |
| DIP_10007050 | hypothetical protein PFICI_09044 [Pestalotiopsis fici W106-1] | 43.1 | 1.0E-154 |
| DIP_10007051 | peroxisomal membrane protein pmp47b [Colletotrichum orbiculare MAFF 240422] | 71.8 | 1.0E-164 |
| DIP_10007052 | putative mak10 subunit protein [Eutypa lata UCREL1] | 58.2 | 0.0 |
| DIP_10007053 | hypothetical protein PFICI_09047 [Pestalotiopsis fici W106-1] | 77.3 | 8.0E-33 |
| DIP_10007054 | putative cytochrome b-c1 complex subunit 2 protein [Eutypa lata UCREL1] | 71.5 | 0.0 |
| DIP_10007055 | putative rna polymerase iii subunit rpc82 protein [Eutypa lata UCREL1] | 61.2 | 0.0 |
| DIP_10007056 | putative fad dependent oxidoreductase protein [Eutypa lata UCREL1] | 35.3 | 3.0E-107 |
| DIP_10007057 | putative adenosine deaminase protein [Eutypa lata UCREL1] | 38.6 | 3.0E-99 |
| DIP_10007058 | hypothetical protein GMDG_07797 [Pseudogymnoascus destructans 20631-21] | 30.9 | 3.0E-37 |
| DIP_10007059 | putative set domain-containing protein [Diaporthe ampelina] | 34.5 | 2.0E-51 |
| DIP_10007060 | hypothetical protein BBA_03720 [Beauveria bassiana ARSEF 2860] | 45.4 | 6.0E-34 |
| DIP_10007061 | hypothetical protein P875_00053242 [Aspergillus parasiticus SU-1] | 34.9 | 2.0E-48 |
| DIP_10007062 | NA | | |
| DIP_10007063 | Clc chloride channel [Glarea lozoyensis ATCC 20868] | 61.3 | 0.0 |
| DIP_10007064 | hypothetical protein FOZG_14865 [Fusarium oxysporum Fo47] | 38.3 | 5.0E-25 |
| DIP_10007065 | hypothetical protein VDAG_06952 [Verticillium dahliae VdLs.17] | 49.4 | 3.0E-52 |
| DIP_10007066 | hypothetical protein UCREL1_5298 [Eutypa lata UCREL1] | 46 | 2.0E-36 |
| DIP_10007067 | hypothetical protein UCREL1_5299 [Eutypa lata UCREL1] | 62.2 | 4.0E-57 |
| DIP_10007068 | cytidylyltransferase [Colletotrichum fioriniae PJ7] | 51 | 2.0E-123 |
| DIP_10007069 | putative phosphate transporter protein [Eutypa lata UCREL1] | 52.7 | 0.0 |
| DIP_10007070 | NA | | |
| DIP_10007071 | hypothetical protein UCREL1_7013 [Eutypa lata UCREL1] | 35.4 | 6.0E-87 |
| DIP_10007072 | plectin/S10 domain-containing protein [Beauveria bassiana ARSEF 2860] | 88.1 | 2.0E-94 |
| DIP_10007073 | hypothetical protein MAC_06527 [Metarhizium acridum CQMa 102] | 55.7 | 2.0E-108 |
| DIP_10007074 | Cytochrome c oxidase subunit 6B [Pestalotiopsis fici W106-1] | 91.6 | 4.0E-55 |
| DIP_10007075 | hypothetical protein UCREL1_8514 [Eutypa lata UCREL1] | 74.4 | 1.0E-39 |
| DIP_10007076 | putative nuclear cap-binding protein [Eutypa lata UCREL1] | 54.6 | 8.0E-60 |
| DIP_10007077 | hypothetical protein FOZG_03681 [Fusarium oxysporum Fo47] | 59.5 | 0.0 |
| DIP_10007078 | hypothetical protein CGLO_16183 [Colletotrichum gloeosporioides Cg-14] | 34.6 | 9.0E-47 |
| DIP_10007079 | hypothetical protein MYCTH_2070906 [Myceliophthora thermophila ATCC 42464] | 41.6 | 4.0E-58 |
| DIP_10007080 | NA | | |
| DIP_10007081 | NA | | |
| DIP_10007082 | Glyceraldehyde-3-phosphate dehydrogenase [Pestalotiopsis fici W106-1] | 86.8 | 0.0 |
| DIP_10007083 | RuvB-like helicase 2 [Colletotrichum higginsianum] | 86.2 | 0.0 |
| DIP_10007084 | peptidyl-trna hydrolase 2 [Colletotrichum orbiculare MAFF 240422] | 57.2 | 6.0E-95 |
| DIP_10007085 | hypothetical protein PFICI_09354 [Pestalotiopsis fici W106-1] | 54.4 | 1.0E-147 |
| DIP_10007086 | putative ubiquitin fusion degradation protein [Eutypa lata UCREL1] | 75.7 | 0.0 |
| DIP_10007087 | NA | | |
| DIP_10007088 | hypothetical protein UCREL1_3189 [Eutypa lata UCREL1] | 43.3 | 4.0E-153 |
| DIP_10007089 | hypothetical protein UCREL1_6569 [Eutypa lata UCREL1] | 43.6 | 5.0E-86 |
| DIP_10007090 | hypothetical protein PFICI_09338 [Pestalotiopsis fici W106-1] | 45.5 | 0.0 |
| DIP_10007091 | hypothetical protein SMAC_02373 [Sordaria macrospora k-hell] | 37.9 | 2.0E-36 |
| DIP_10007092 | integral membrane protein [Ophiostoma piceae UAMH 11346] | 42.5 | 9.0E-123 |
| DIP_10007093 | hypothetical protein PFICI_11341 [Pestalotiopsis fici W106-1] | 59.2 | 2.0E-144 |
| DIP_10007094 | hypothetical protein PFICI_11342 [Pestalotiopsis fici W106-1] | 88.6 | 1.0E-43 |
| DIP_10007095 | UPF0075-domain-containing protein [Sphaerulina musiva SO2202] | 72.1 | 0.0 |
| DIP_10007096 | NA | | |
| DIP_10007097 | putative erythrocyte band 7 integral membrane protein [Diaporthe ampelina] | 75.9 | 0.0 |
| DIP_10007098 | putative atp-dependent rna helicase spb4 protein [Eutypa lata UCREL1] | 59.9 | 0.0 |
| DIP_10007099 | ADP-ribose pyrophosphatase [Coniosporium apollinis CBS 100218] | 50.2 | 4.0E-67 |
| DIP_10007100 | NA | | |
| DIP_10007101 | NA | | |
| DIP_10007102 | hypothetical protein FGSG_03733 [Fusarium graminearum PH-1] | 44.2 | 1.0E-113 |
| DIP_10007103 | Glucan endo-1,3-beta-glucosidase-like protein [Acremonium chrysogenum ATCC 11550] | 37.2 | 2.0E-61 |
| DIP_10007104 | NA | | |
| DIP_10007105 | hypothetical protein V493_00035 [Pseudogymnoascus pannorum VKM F-4281 (FW-2241)] | 41.7 | 8.0E-42 |
| DIP_10007106 | putative atp citrate lyase protein [Eutypa lata UCREL1] | 87 | 0.0 |
| DIP_10007107 | putative atp-citrate synthase subunit 1 protein [Eutypa lata UCREL1] | 90.7 | 0.0 |
| DIP_10007108 | similar to 50S ribosomal protein L30 [Leptosphaeria maculans JN3] | 57.5 | 2.0E-30 |
| DIP_10007109 | putative keratinolytic protein [Eutypa lata UCREL1] | 73.1 | 4.0E-101 |
| DIP_10007110 | hypothetical protein PFICI_08535 [Pestalotiopsis fici W106-1] | 63.5 | 3.0E-62 |
| DIP_10007111 | hypothetical protein TRIATDRAFT_30449 [Trichoderma atroviride IMI 206040] | 50.5 | 3.0E-60 |
| DIP_10007112 | putative peptidyl-trna hydrolase protein [Eutypa lata UCREL1] | 63.7 | 4.0E-118 |
| DIP_10007113 | hypothetical protein PFICI_08538 [Pestalotiopsis fici W106-1] | 38 | 1.0E-50 |
| DIP_10007114 | NA | | |
| DIP_10007115 | NA | | |
| DIP_10007116 | NA | | |
| DIP_10007117 | NA | | |
| DIP_10007118 | hypothetical protein NECHADRAFT_97328 [Nectria haematococca mpVI 77-13-4] | 46.8 | 2.0E-55 |
| DIP_10007119 | hypothetical protein SMAC_06982 [Sordaria macrospora k-hell] | 45.5 | 1.0E-67 |
| DIP_10007120 | NA | | |
| DIP_10007121 | NA | | |
| DIP_10007122 | hypothetical protein LRAMOSA05471 [Absidia idahoensis var. thermophila] | 39 | 2.0E-12 |
| DIP_10007123 | NA | | |
| DIP_10007124 | putative pre-mrna splicing factor atp-dependent rna helicase prp43 protein [Eutypa lata UCREL1] | 78.9 | 0.0 |
| DIP_10007125 | hypothetical protein UCREL1_524 [Eutypa lata UCREL1] | 73.3 | 0.0 |
| DIP_10007126 | hypothetical protein PFICI_11338 [Pestalotiopsis fici W106-1] | 46.3 | 7.0E-84 |
| DIP_10007127 | putative anaphase-promoting complex subunit 10 protein [Eutypa lata UCREL1] | 80.2 | 5.0E-144 |
| DIP_10007128 | putative aminopeptidase 2 protein [Eutypa lata UCREL1] | 81.6 | 0.0 |
| DIP_10007129 | Mitochondrial GTP/GDP carrier protein 1 [Pestalotiopsis fici W106-1] | 87.8 | 0.0 |
| DIP_10007130 | putative chromosome segregation atpase protein [Eutypa lata UCREL1] | 58.6 | 0.0 |
| DIP_10007131 | seryl-tRNA synthetase [Gaeumannomyces graminis var. tritici R3-111a-1] | 51.4 | 9.0E-160 |
| DIP_10007132 | hypothetical protein FPSE_05742 [Fusarium pseudograminearum CS3096] | 32.2 | 6.0E-60 |
| DIP_10007133 | putative tyrosinase central domain protein [Eutypa lata UCREL1] | 50.6 | 1.0E-126 |
| DIP_10007134 | integral membrane protein [Colletotrichum gloeosporioides Nara gc5] | 43.7 | 7.0E-75 |
| DIP_10007135 | NA | | |
| DIP_10007136 | hypothetical protein Z517_02350 [Fonsecaea pedrosoi CBS 271.37] | 31.8 | 5.0E-77 |
| DIP_10007137 | hypothetical protein CFIO01_07275 [Colletotrichum fioriniae PJ7] | 36.8 | 3.0E-16 |
| DIP_10007138 | putative cruciform cutting endonuclease mitochondrial protein [Eutypa lata UCREL1] | 38.6 | 5.0E-65 |
| DIP_10007139 | putative glutathione reductase protein [Eutypa lata UCREL1] | 76.7 | 0.0 |
| DIP_10007140 | hypothetical protein PFICI_08762 [Pestalotiopsis fici W106-1] | 69.5 | 0.0 |
| DIP_10007141 | hypothetical protein PFICI_08761 [Pestalotiopsis fici W106-1] | 58.5 | 0.0 |
| DIP_10007142 | NA | | |
| DIP_10007143 | putative 3-oxo-5-alpha-steroid 4-dehydrogenase protein [Eutypa lata UCREL1] | 67.6 | 1.0E-150 |
| DIP_10007144 | putative traf-like signal protein [Eutypa lata UCREL1] | 50.1 | 2.0E-138 |
| DIP_10007145 | hypothetical protein PFICI_00325 [Pestalotiopsis fici W106-1] | 39.2 | 5.0E-90 |
| DIP_10007146 | NA | | |
| DIP_10007147 | hypothetical protein H633G_09760 [Metarhizium anisopliae BRIP 53284] | 59.5 | 6.0E-81 |
| DIP_10007148 | hypothetical protein MGG_16147 [Magnaporthe oryzae 70-15] | 43.2 | 6.0E-32 |
| DIP_10007149 | NA | | |
| DIP_10007150 | hypothetical protein UCREL1_2082 [Eutypa lata UCREL1] | 60.5 | 4.0E-48 |
| DIP_10007151 | putative secreted aspartic proteinase precursor protein [Eutypa lata UCREL1] | 51.3 | 2.0E-151 |
| DIP_10007152 | hypothetical protein PFICI_08797 [Pestalotiopsis fici W106-1] | 59.6 | 0.0 |
| DIP_10007153 | hypothetical protein PFICI_06380 [Pestalotiopsis fici W106-1] | 68.4 | 9.0E-125 |
| DIP_10007154 | hypothetical protein THAR02_07317 [Trichoderma harzianum] | 40.5 | 4.0E-74 |
| DIP_10007155 | putative 40s ribosomal protein s5 protein [Eutypa lata UCREL1] | 94.8 | 1.0E-150 |
| DIP_10007156 | NA | | |
| DIP_10007157 | NA | | |
| DIP_10007158 | alcohol dehydrogenase II [Uncinocarpus reesii 1704] | 58.6 | 5.0E-151 |
| DIP_10007159 | putative cytochrome b5-like heme steroid binding domain-containing protein [Eutypa lata UCREL1] | 63 | 2.0E-124 |
| DIP_10007160 | putative extracellular serine-rich protein [Eutypa lata UCREL1] | 48.7 | 3.0E-57 |
| DIP_10007161 | hypothetical protein UCREL1_11160 [Eutypa lata UCREL1] | 46.5 | 4.0E-39 |
| DIP_10007162 | putative carrier protein [Eutypa lata UCREL1] | 93.2 | 0.0 |
| DIP_10007163 | hypothetical protein AOR_1_4054 [Aspergillus oryzae RIB40] | 42 | 2.0E-24 |
| DIP_10007164 | Ribosomal protein S25 [Metarhizium robertsii ARSEF 23] | 88.5 | 4.0E-51 |
| DIP_10007165 | hypothetical protein PFICI_08806 [Pestalotiopsis fici W106-1] | 53.6 | 0.0 |
| DIP_10007166 | hypothetical protein UCREL1_11171 [Eutypa lata UCREL1] | 37.8 | 3.0E-178 |
| DIP_10007167 | 60S ribosomal protein L26 [Pestalotiopsis fici W106-1] | 89.7 | 2.0E-80 |
| DIP_10007168 | hypothetical protein VHEMI01762 [Torrubiella hemipterigena] | 57.6 | 1.0E-79 |
| DIP_10007169 | putative 50s ribosomal protein l3 protein [Eutypa lata UCREL1] | 77.5 | 0.0 |
| DIP_10007170 | coiled-coil domain-containing protein 25 [Magnaporthe oryzae Y34] | 75.3 | 3.0E-122 |
| DIP_10007171 | Galactose-binding protein [Glarea lozoyensis ATCC 20868] | 71.2 | 2.0E-114 |
| DIP_10007172 | NA | | |
| DIP_10007173 | hypothetical protein THITE_2123640 [Thielavia terrestris NRRL 8126] | 70.9 | 3.0E-133 |
| DIP_10007174 | putative mitotic-spindle organizing protein 1 protein [Eutypa lata UCREL1] | 80.3 | 2.0E-37 |
| DIP_10007175 | NA | | |
| DIP_10007176 | CHY zinc finger [Colletotrichum fioriniae PJ7] | 52.3 | 8.0E-144 |
| DIP_10007177 | hypothetical protein S7711_07228 [Stachybotrys chartarum IBT 7711] | 50 | 9.0E-13 |
| DIP_10007178 | hypothetical protein PFICI_08613 [Pestalotiopsis fici W106-1] | 83.3 | 0.0 |
| DIP_10007179 | hypothetical protein NEUTE1DRAFT_122957 [Neurospora tetrasperma FGSC 2508] | 91.8 | 4.0E-156 |
| DIP_10007180 | putative six-hairpin glycosidase-like protein [Eutypa lata UCREL1] | 68.7 | 4.0E-68 |
| DIP_10007181 | putative six-hairpin glycosidase-like protein [Togninia minima UCRPA7] | 52 | 2.0E-33 |
| DIP_10007182 | hypothetical protein THAR02_08461 [Trichoderma harzianum] | 43 | 0.0 |
| DIP_10007183 | hypothetical protein OIDMADRAFT_16881 [Oidiodendron maius Zn] | 64.7 | 1.0E-69 |
| DIP_10007184 | hypothetical protein S7711_02079 [Stachybotrys chartarum IBT 7711] | 52.2 | 1.0E-143 |
| DIP_10007185 | putative nudix family protein [Eutypa lata UCREL1] | 82.2 | 2.0E-92 |
| DIP_10007186 | NA | | |
| DIP_10007187 | NA | | |
| DIP_10007188 | hypothetical protein SBOR_5371 [Sclerotinia borealis F-4157] | 47 | 6.0E-120 |
| DIP_10007189 | hypothetical protein Z518_05360 [Rhinocladiella mackenziei CBS 650.93] | 44.3 | 2.0E-133 |
| DIP_10007190 | hypothetical protein PFICI_08604 [Pestalotiopsis fici W106-1] | 57.2 | 7.0E-154 |
| DIP_10007191 | Cytochrome c [Pestalotiopsis fici W106-1] | 92.4 | 4.0E-68 |
| DIP_10007192 | hypothetical protein PFICI_08605 [Pestalotiopsis fici W106-1] | 53.8 | 4.0E-78 |
| DIP_10007193 | putative aaa family atpase 60s ribosome export protein [Eutypa lata UCREL1] | 66.2 | 0.0 |
| DIP_10007194 | putative snf7 family protein [Eutypa lata UCREL1] | 73 | 3.0E-97 |
| DIP_10007195 | hypothetical protein BC1G_08512 [Botrytis cinerea B05.10] | 62.6 | 3.0E-101 |
| DIP_10007196 | hypothetical protein CGLO_15297 [Colletotrichum gloeosporioides Cg-14] | 73.5 | 1.0E-114 |
| DIP_10007197 | hypothetical protein PFICI_08591 [Pestalotiopsis fici W106-1] | 82.9 | 0.0 |
| DIP_10007198 | hypothetical protein PFICI_08594 [Pestalotiopsis fici W106-1] | 61.3 | 0.0 |
| DIP_10007199 | hypothetical protein CFIO01_10168 [Colletotrichum fioriniae PJ7] | 36.6 | 3.0E-146 |
| DIP_10007200 | hypothetical protein PFICI_09394 [Pestalotiopsis fici W106-1] | 49.7 | 2.0E-38 |
| DIP_10007201 | hypothetical protein PV04_08125 [Capronia semiimmersa] | 30.3 | 6.0E-19 |
| DIP_10007202 | putative vesicular-fusion protein sec17 protein [Eutypa lata UCREL1] | 73.6 | 2.0E-174 |
| DIP_10007203 | putative pre-rrna processing protein rrp12 protein [Eutypa lata UCREL1] | 77.1 | 0.0 |
| DIP_10007204 | hypothetical protein FAVG1_06634 [Fusarium avenaceum] | 73.4 | 4.0E-118 |
| DIP_10007205 | hypothetical protein PFICI_09399 [Pestalotiopsis fici W106-1] | 70.9 | 0.0 |
| DIP_10007206 | hypothetical protein PFICI_09400 [Pestalotiopsis fici W106-1] | 73.1 | 6.0E-135 |
| DIP_10007207 | hypothetical protein PFICI_08813 [Pestalotiopsis fici W106-1] | 32.7 | 2.0E-47 |
| DIP_10007208 | hypothetical protein PFICI_02701 [Pestalotiopsis fici W106-1] | 65.4 | 0.0 |
| DIP_10007209 | putative arylsulfotransferase-like protein [Eutypa lata UCREL1] | 38.8 | 3.0E-104 |
| DIP_10007210 | histidine kinase, variant 2 [Trichoderma atroviride IMI 206040] | 50.7 | 0.0 |
| DIP_10007211 | NA | | |
| DIP_10007212 | hypothetical protein UCREL1_10063 [Eutypa lata UCREL1] | 51.7 | 2.0E-53 |
| DIP_10007213 | hypothetical protein PFICI_07173 [Pestalotiopsis fici W106-1] | 53.2 | 1.0E-40 |
| DIP_10007214 | hypothetical protein PFICI_07174 [Pestalotiopsis fici W106-1] | 50.7 | 0.0 |
| DIP_10007215 | NA | | |
| DIP_10007216 | hypothetical protein PFICI_13319 [Pestalotiopsis fici W106-1] | 55.3 | 0.0 |
| DIP_10007217 | Putative 40S ribosomal protein S11 [Torrubiella hemipterigena] | 96.3 | 1.0E-112 |
| DIP_10007218 | hypothetical protein PFICI_13317 [Pestalotiopsis fici W106-1] | 58.7 | 3.0E-133 |
| DIP_10007219 | hypothetical protein UCREL1_4569 [Eutypa lata UCREL1] | 43.8 | 4.0E-45 |
| DIP_10007220 | hypothetical protein PFICI_08649 [Pestalotiopsis fici W106-1] | 41.2 | 8.0E-139 |
| DIP_10007221 | hypothetical protein DOTSEDRAFT_22137 [Dothistroma septosporum NZE10] | 35.7 | 6.0E-36 |
| DIP_10007222 | predicted protein [Ostreococcus lucimarinus CCE9901] | 30.8 | 3.0E-19 |
| DIP_10007223 | hypothetical protein S40288_08811 [Stachybotrys chartarum IBT 40288] | 67.1 | 4.0E-104 |
| DIP_10007224 | hypothetical protein Cob_10880 [Colletotrichum orbiculare MAFF 240422] | 79.3 | 8.0E-26 |
| DIP_10007225 | hypothetical protein PFICI_14381 [Pestalotiopsis fici W106-1] | 53.7 | 0.0 |
| DIP_10007226 | NA | | |
| DIP_10007227 | NA | | |
| DIP_10007228 | putative glucosidase 2 subunit beta protein [Eutypa lata UCREL1] | 67.7 | 0.0 |
| DIP_10007229 | hypothetical protein FGSG_02955 [Fusarium graminearum PH-1] | 83.8 | 8.0E-40 |
| DIP_10007230 | NA | | |
| DIP_10007231 | putative duf602 domain-containing protein [Eutypa lata UCREL1] | 55.3 | 5.0E-101 |
| DIP_10007232 | hypothetical protein TRIVIDRAFT_56056 [Trichoderma virens Gv29-8] | 77.9 | 0.0 |
| DIP_10007233 | putative u-box domain-containing protein [Eutypa lata UCREL1] | 37.8 | 1.0E-147 |
| DIP_10007234 | NA | | |
| DIP_10007235 | hypothetical protein PFICI_14391 [Pestalotiopsis fici W106-1] | 48.2 | 0.0 |
| DIP_10007236 | NA | | |
| DIP_10007237 | putative aaa atpase domain containing protein [Eutypa lata UCREL1] | 40.1 | 2.0E-44 |
| DIP_10007238 | putative alcohol dehydrogenase protein [Eutypa lata UCREL1] | 51.6 | 9.0E-108 |
| DIP_10007239 | hypothetical protein PFICI_10312 [Pestalotiopsis fici W106-1] | 79.5 | 1.0E-62 |
| DIP_10007240 | putative beta-lactamase-type transpeptidase fold domain containing protein [Eutypa lata UCREL1] | 68.5 | 0.0 |
| DIP_10007241 | NA | | |
| DIP_10007242 | hypothetical protein PFICI_10319 [Pestalotiopsis fici W106-1] | 51 | 2.0E-113 |
| DIP_10007243 | putative cell division control protein [Togninia minima UCRPA7] | 62.7 | 0.0 |
| DIP_10007244 | putative bet v1-like protein [Eutypa lata UCREL1] | 85.2 | 8.0E-90 |
| DIP_10007245 | putative fad binding domain-containing protein [Eutypa lata UCREL1] | 54.8 | 0.0 |
| DIP_10007246 | putative carbohydrate-binding module family 1 protein [Eutypa lata UCREL1] | 57.7 | 1.0E-106 |
| DIP_10007247 | putative nonribosomal peptide synthetase 10 protein [Eutypa lata UCREL1] | 69.6 | 0.0 |
| DIP_10007248 | NA | | |
| DIP_10007249 | electron transfer flavoprotein-ubiquinone oxidoreductase [Pestalotiopsis fici W106-1] | 79.6 | 0.0 |
| DIP_10007250 | C6 finger domain-containing protein [Blastomyces dermatitidis ATCC 18188] | 32.1 | 1.0E-68 |
| DIP_10007251 | hypothetical protein UCDDA912_g00666 [Diaporthe ampelina] | 43.9 | 5.0E-148 |
| DIP_10007252 | hypothetical protein PFICI_09939 [Pestalotiopsis fici W106-1] | 57.2 | 0.0 |
| DIP_10007253 | putative polyadenylate-binding protein [Eutypa lata UCREL1] | 83 | 0.0 |
| DIP_10007254 | hypothetical protein TCE0_033f08694, partial [Talaromyces cellulolyticus] | 31 | 3.0E-25 |
| DIP_10007255 | putative iq and hect domain protein [Eutypa lata UCREL1] | 63.9 | 0.0 |
| DIP_10007256 | hypothetical protein MANI_018607 [Metarhizium anisopliae] | 51.9 | 4.0E-48 |
| DIP_10007257 | hypothetical protein FAVG1_02406 [Fusarium avenaceum] | 35.9 | 5.0E-112 |
| DIP_10007258 | putative cytochrome b2 protein [Eutypa lata UCREL1] | 65 | 0.0 |
| DIP_10007259 | hypothetical protein M436DRAFT_82503 [Aureobasidium namibiae CBS 147.97] | 61 | 2.0E-25 |
| DIP_10007260 | 3-ketoacyl-CoA thiolase [Pestalotiopsis fici W106-1] | 86 | 0.0 |
| DIP_10007261 | putative ornithine cyclodeaminase mu-crystallin family protein [Togninia minima UCRPA7] | 45.9 | 2.0E-145 |
| DIP_10007262 | NA | | |
| DIP_10007263 | carbonic anhydrase [Colletotrichum higginsianum] | 50.8 | 5.0E-56 |
| DIP_10007264 | putative 60s ribosomal protein l8 protein [Eutypa lata UCREL1] | 91.2 | 3.0E-177 |
| DIP_10007265 | Cytoplasmic tRNA 2-thiolation protein 1 [Pestalotiopsis fici W106-1] | 78.4 | 0.0 |
| DIP_10007266 | hypothetical protein PFICI_09936 [Pestalotiopsis fici W106-1] | 75.6 | 1.0E-39 |
| DIP_10007267 | putative histone acetyltransferase protein [Eutypa lata UCREL1] | 58.3 | 0.0 |
| DIP_10007268 | putative nucleoporin sonb protein [Eutypa lata UCREL1] | 61.5 | 0.0 |
| DIP_10007269 | putative cell division control protein cdc31 protein [Eutypa lata UCREL1] | 83.6 | 2.0E-120 |
| DIP_10007270 | putative mitochondrial 2-oxoglutarate malate carrier protein [Diaporthe ampelina] | 82.5 | 0.0 |
| DIP_10007271 | hypothetical protein FGSG_04457 [Fusarium graminearum PH-1] | 37 | 6.0E-75 |
| DIP_10007272 | hypothetical protein PFICI_09953 [Pestalotiopsis fici W106-1] | 47.8 | 1.0E-129 |
| DIP_10007273 | hypothetical protein PFICI_09943 [Pestalotiopsis fici W106-1] | 58.9 | 0.0 |
| DIP_10007274 | putative f-box domain containing protein [Eutypa lata UCREL1] | 38.6 | 1.0E-119 |
| DIP_10007275 | putative mitochondrial 2-oxodicarboxylate transporter [Diaporthe ampelina] | 66.1 | 2.0E-164 |
| DIP_10007276 | hypothetical protein MYCTH_2307046 [Myceliophthora thermophila ATCC 42464] | 32.1 | 1.0E-36 |
| DIP_10007277 | putative enth domain-containing protein [Eutypa lata UCREL1] | 73.4 | 0.0 |
| DIP_10007278 | putative glutaminyl-peptide cyclotransferase [Diaporthe ampelina] | 64 | 4.0E-157 |
| DIP_10007279 | hypothetical protein UCREL1_10541 [Eutypa lata UCREL1] | 33.9 | 2.0E-53 |
| DIP_10007280 | putative endo- -beta- protein [Eutypa lata UCREL1] | 68.2 | 1.0E-125 |
| DIP_10007281 | NA | | |
| DIP_10007282 | hypothetical protein UCREL1_9147 [Eutypa lata UCREL1] | 85.6 | 0.0 |
| DIP_10007283 | similar to MFS sugar transporter [Botrytis cinerea T4] | 65.5 | 7.0E-142 |
| DIP_10007284 | hypothetical protein MAJ_08926, partial [Metarhizium majus ARSEF 297] | 45.8 | 5.0E-40 |
| DIP_10007285 | hypothetical protein PFICI_02699 [Pestalotiopsis fici W106-1] | 69.9 | 0.0 |
| DIP_10007286 | hypothetical protein PFICI_02698 [Pestalotiopsis fici W106-1] | 56.8 | 0.0 |
| DIP_10007287 | hypothetical protein PFICI_02696 [Pestalotiopsis fici W106-1] | 83.9 | 1.0E-174 |
| DIP_10007288 | RNA polymerase I specific transcription initiation factor [Colletotrichum graminicola M1.001] | 31.6 | 2.0E-59 |
| DIP_10007289 | putative pescadillo development protein [Togninia minima UCRPA7] | 68.5 | 0.0 |
| DIP_10007290 | glycosylphosphatidylinositol transamidase 2 [Neurospora crassa OR74A] | 73.6 | 0.0 |
| DIP_10007291 | NA | | |
| DIP_10007292 | hypothetical protein MYCFIDRAFT_211918 [Pseudocercospora fijiensis CIRAD86] | 46.2 | 2.0E-91 |
| DIP_10007293 | hypothetical protein HMPREF1120_08290 [Exophiala dermatitidis NIH/UT8656] | 45.5 | 2.0E-117 |
| DIP_10007294 | NA | | |
| DIP_10007295 | hypothetical protein PFICI_02695 [Pestalotiopsis fici W106-1] | 82.3 | 5.0E-175 |
| DIP_10007296 | hypothetical protein PFICI_02694 [Pestalotiopsis fici W106-1] | 66 | 0.0 |
| DIP_10007297 | hypothetical protein PFICI_06206 [Pestalotiopsis fici W106-1] | 60.3 | 3.0E-80 |
| DIP_10007298 | NA | | |
| DIP_10007299 | hypothetical protein NEUTE1DRAFT_132883 [Neurospora tetrasperma FGSC 2508] | 54.2 | 3.0E-108 |
| DIP_10007300 | putative isoleucyl-trna synthetase protein [Eutypa lata UCREL1] | 61.9 | 0.0 |
| DIP_10007301 | putative kinesin-ii 85 kda subunit protein [Eutypa lata UCREL1] | 65.8 | 0.0 |
| DIP_10007302 | hypothetical protein PFICI_06932 [Pestalotiopsis fici W106-1] | 69.6 | 0.0 |
| DIP_10007303 | putative quinoprotein amine dehydrogenase beta chain-like protein [Eutypa lata UCREL1] | 34.9 | 5.0E-43 |
| DIP_10007304 | Heat shock protein SSB1 [Pestalotiopsis fici W106-1] | 91 | 0.0 |
| DIP_10007305 | hypothetical protein [Podospora anserina S mat+] | 49.7 | 6.0E-97 |
| DIP_10007306 | lysosomal cobalamin transporter [Pestalotiopsis fici W106-1] | 79.6 | 0.0 |
| DIP_10007307 | hypothetical protein PFICI_06962 [Pestalotiopsis fici W106-1] | 42.6 | 5.0E-39 |
| DIP_10007308 | putative glycine-rich protein [Eutypa lata UCREL1] | 63.5 | 1.0E-73 |
| DIP_10007309 | putative g u mismatch-specific uracil dna glycosylase protein [Eutypa lata UCREL1] | 66 | 1.0E-100 |
| DIP_10007310 | putative udp-n-acetylglucosamine pyrophosphorylase protein [Eutypa lata UCREL1] | 65.1 | 0.0 |
| DIP_10007311 | putative taurine dioxygenase family protein [Eutypa lata UCREL1] | 73.7 | 0.0 |
| DIP_10007312 | putative allantoate permease protein [Eutypa lata UCREL1] | 67.4 | 0.0 |
| DIP_10007313 | hypothetical protein PFICI_11543 [Pestalotiopsis fici W106-1] | 66.7 | 4.0E-125 |
| DIP_10007314 | hypothetical protein PV09_04387 [Verruconis gallopava] | 48.8 | 6.0E-121 |
| DIP_10007315 | putative glycoside hydrolase family 47 protein [Eutypa lata UCREL1] | 61 | 0.0 |
| DIP_10007316 | putative general amino acid permease agp2 protein [Togninia minima UCRPA7] | 60.5 | 0.0 |
| DIP_10007317 | acetolactate synthase small subunit [Gaeumannomyces graminis var. tritici R3-111a-1] | 77.6 | 2.0E-180 |

| | | | |
|---|---|---|---|
| DIP_10007318 | ornithine carbamoyltransferase [Magnaporthe oryzae 70-15] | 65.4 | 5.0E-163 |
| DIP_10007319 | hypothetical protein PFICI_08691 [Pestalotiopsis fici W106-1] | 57.1 | 0.0 |
| DIP_10007320 | NA | | |
| DIP_10007321 | NA | | |
| DIP_10007322 | putative sart-1 family protein [Eutypa lata UCREL1] | 68.3 | 0.0 |
| DIP_10007323 | putative glycoside hydrolase family 15 protein [Eutypa lata UCREL1] | 68.1 | 0.0 |
| DIP_10007324 | hypothetical protein PFICI_14952 [Pestalotiopsis fici W106-1] | 50.8 | 1.0E-96 |
| DIP_10007325 | hypothetical protein PFICI_14953 [Pestalotiopsis fici W106-1] | 72.4 | 1.0E-179 |
| DIP_10007326 | NA | | |
| DIP_10007327 | putative succinyl- ligase beta-chain protein [Togninia minima UCRPA7] | 86.9 | 0.0 |
| DIP_10007328 | NA | | |
| DIP_10007329 | NA | | |
| DIP_10007330 | flocculation suppression protein [Magnaporthe oryzae Y34] | 59.8 | 2.0E-104 |
| DIP_10007331 | NA | | |
| DIP_10007332 | NA | | |
| DIP_10007333 | hypothetical protein PFICI_05571 [Pestalotiopsis fici W106-1] | 43.6 | 4.0E-52 |
| DIP_10007334 | putative integral membrane protein [Eutypa lata UCREL1] | 51.4 | 3.0E-125 |
| DIP_10007335 | putative minichromosome loss protein 1 protein [Eutypa lata UCREL1] | 63.6 | 0.0 |
| DIP_10007336 | WD repeat protein [Neurospora crassa OR74A] | 57.9 | 4.0E-163 |
| DIP_10007337 | hypothetical protein UCREL1_1053 [Eutypa lata UCREL1] | 70.9 | 1.0E-121 |
| DIP_10007338 | putative gtp-binding protein ypt1 protein [Eutypa lata UCREL1] | 88.2 | 7.0E-151 |
| DIP_10007339 | hypothetical protein ACRE_057760 [Acremonium chrysogenum ATCC 11550] | 30.8 | 2.0E-139 |
| DIP_10007340 | NA | | |
| DIP_10007341 | putative carnitine acetyl transferase protein [Eutypa lata UCREL1] | 65.7 | 0.0 |
| DIP_10007342 | putative mfs multidrug transporter protein [Eutypa lata UCREL1] | 50.9 | 0.0 |
| DIP_10007343 | hypothetical protein NECHADRAFT_75126 [Nectria haematococca mpVI 77-13-4] | 32.8 | 3.0E-37 |
| DIP_10007344 | NA | | |
| DIP_10007345 | putative dna repair protein rad57 protein [Togninia minima UCRPA7] | 56 | 0.0 |
| DIP_10007346 | putative dna-directed rna polymerase iii 130 kda polypeptide protein [Eutypa lata UCREL1] | 89.1 | 0.0 |
| DIP_10007347 | hypothetical protein BofuT4_P095590.1 [Botrytis cinerea T4] | 65.7 | 1.0E-68 |
| DIP_10007348 | putative pumilio domain-containing protein [Eutypa lata UCREL1] | 66.3 | 0.0 |
| DIP_10007349 | NA | | |
| DIP_10007350 | 60S ribosomal protein L15 [Neurospora crassa OR74A] | 94.1 | 4.0E-141 |
| DIP_10007351 | NA | | |
| DIP_10007352 | cupin [Colletotrichum fioriniae PJ7] | 37.8 | 3.0E-95 |
| DIP_10007353 | ribonuclease H2 subunit A [Gaeumannomyces graminis var. tritici R3-111a-1] | 49.2 | 1.0E-120 |
| DIP_10007354 | Protein SEY1 [Pestalotiopsis fici W106-1] | 79.9 | 0.0 |
| DIP_10007355 | hypothetical protein FAVG1_08079 [Fusarium avenaceum] | 75.5 | 0.0 |
| DIP_10007356 | putative ribosome biogenesis protein tsr1 protein [Eutypa lata UCREL1] | 74.7 | 0.0 |
| DIP_10007357 | urease accessory protein UreG [Sporothrix schenckii ATCC 58251] | 71.6 | 1.0E-135 |
| DIP_10007358 | mitochondrial ribosomal protein s19 [Grosmannia clavigera kw1407] | 84.4 | 2.0E-50 |
| DIP_10007359 | putative transcription initiation factor tfiid subunit 10 protein [Eutypa lata UCREL1] | 82.7 | 3.0E-97 |
| DIP_10007360 | 40S ribosomal protein S6-B [Fusarium verticillioides 7600] | 92.1 | 2.0E-163 |
| DIP_10007361 | NA | | |
| DIP_10007362 | putative ribose-phosphate pyrophosphokinase ii protein [Eutypa lata UCREL1] | 91.6 | 0.0 |
| DIP_10007363 | putative mfs sugar protein [Eutypa lata UCREL1] | 75.1 | 0.0 |
| DIP_10007364 | NA | | |
| DIP_10007365 | hypothetical protein BC1G_04261 [Botrytis cinerea B05.10] | 39.3 | 0.0 |
| DIP_10007366 | 40S ribosomal protein S8-B [Pestalotiopsis fici W106-1] | 91.1 | 6.0E-136 |
| DIP_10007367 | GTPase-activating protein GYP5 [Verticillium dahliae VdLs.17] | 61.9 | 0.0 |
| DIP_10007368 | putative sarcosine oxidase protein [Eutypa lata UCREL1] | 57.8 | 7.0E-180 |
| DIP_10007369 | conserved hypothetical protein [Byssochlamys spectabilis No. 5] | 34.8 | 5.0E-10 |
| DIP_10007370 | hypothetical protein Z519_05474 [Cladophialophora bantiana CBS 173.52] | 36 | 5.0E-13 |
| DIP_10007371 | hypothetical protein FGSG_01837 [Fusarium graminearum PH-1] | 30.7 | 2.0E-14 |
| DIP_10007372 | hypothetical protein V493_02396 [Pseudogymnoascus pannorum VKM F-4281 (FW-2241)] | 35.8 | 2.0E-08 |
| DIP_10007373 | NA | | |
| DIP_10007374 | hypothetical protein UCREL1_4792 [Eutypa lata UCREL1] | 32.1 | 4.0E-20 |
| DIP_10007375 | NA | | |
| DIP_10007376 | myb DNA-binding protein [Grosmannia clavigera kw1407] | 33.8 | 2.0E-80 |
| DIP_10007377 | hypothetical protein PFICI_08512 [Pestalotiopsis fici W106-1] | 46 | 0.0 |
| DIP_10007378 | hypothetical protein PFICI_15032 [Pestalotiopsis fici W106-1] | 61.9 | 0.0 |
| DIP_10007379 | hypothetical protein CHGG_08212 [Chaetomium globosum CBS 148.51] | 53 | 1.0E-144 |
| DIP_10007380 | predicted protein [Nectria haematococca mpVI 77-13-4] | 53.1 | 8.0E-120 |
| DIP_10007381 | hypothetical protein UCREL1_8272 [Eutypa lata UCREL1] | 71.2 | 9.0E-54 |
| DIP_10007382 | hypothetical protein CHGG_08048 [Chaetomium globosum CBS 148.51] | 47.1 | 1.0E-63 |
| DIP_10007383 | putative r3h domain-containing protein [Eutypa lata UCREL1] | 76.1 | 0.0 |
| DIP_10007384 | hypothetical protein UCREL1_3608 [Eutypa lata UCREL1] | 54.6 | 0.0 |
| DIP_10007385 | NA | | |
| DIP_10007386 | hypothetical protein UCREL1_3609 [Eutypa lata UCREL1] | 32.5 | 2.0E-12 |
| DIP_10007387 | hypothetical protein PFICI_09391 [Pestalotiopsis fici W106-1] | 73 | 0.0 |
| DIP_10007388 | putative pro11 protein [Diaporthe ampelina] | 67.8 | 4.0E-78 |
| DIP_10007389 | putative thioesterase superfamily protein [Eutypa lata UCREL1] | 48.5 | 5.0E-49 |
| DIP_10007390 | Histone H2A.Z [Pestalotiopsis fici W106-1] | 91.7 | 3.0E-75 |
| DIP_10007391 | putative replication factor c subunit 3 protein [Eutypa lata UCREL1] | 87 | 0.0 |
| DIP_10007392 | hypothetical protein PFICI_03691 [Pestalotiopsis fici W106-1] | 36.7 | 2.0E-91 |
| DIP_10007393 | RNA-dependent RNA polymerase [Rosellinia necatrix] | 47.8 | 0.0 |
| DIP_10007394 | hypothetical protein PFICI_03048 [Pestalotiopsis fici W106-1] | 52.2 | 0.0 |
| DIP_10007395 | hypothetical protein UCRNP2_3252 [Neofusicoccum parvum UCRNP2] | 39.8 | 6.0E-11 |
| DIP_10007396 | deuterolysin metalloprotease [Colletotrichum fioriniae PJ7] | 46.9 | 1.0E-109 |
| DIP_10007397 | putative quinoprotein amine dehydrogenase beta chain-like protein [Eutypa lata UCREL1] | 33.1 | 7.0E-23 |
| DIP_10007398 | hypothetical protein S7711_10443 [Stachybotrys chartarum IBT 7711] | 33.5 | 2.0E-25 |
| DIP_10007399 | hypothetical protein, partial [Actinoplanes globisporus] | 61 | 4.0E-160 |
| DIP_10007400 | hypothetical protein BAUCODRAFT_26468 [Baudoinia compniacensis UAMH 10762] | 41.2 | 2.0E-18 |
| DIP_10007401 | hypothetical protein PFICI_14901 [Pestalotiopsis fici W106-1] | 56.2 | 0.0 |
| DIP_10007402 | hypothetical protein S40285_01779 [Stachybotrys chlorohalonata IBT 40285] | 36 | 2.0E-113 |
| DIP_10007403 | hypothetical protein PFICI_14903 [Pestalotiopsis fici W106-1] | 40.9 | 8.0E-55 |
| DIP_10007404 | putative mdix domain-containing protein [Eutypa lata UCREL1] | 42.1 | 3.0E-44 |
| DIP_10007405 | putative glucose galactose transporter protein [Eutypa lata UCREL1] | 75.7 | 0.0 |
| DIP_10007406 | hypothetical protein Z519_07848 [Cladophialophora bantiana CBS 173.52] | 41.9 | 3.0E-67 |
| DIP_10007407 | Mitochondrial phosphate carrier protein [Pestalotiopsis fici W106-1] | 88.6 | 0.0 |
| DIP_10007408 | hypothetical protein PTT_14830 [Pyrenophora teres f. teres 0-1] | 48.1 | 4.0E-116 |
| DIP_10007409 | putative xyloglucan-specific endo-beta- -glucanase precursor protein [Togninia minima UCRPA7] | 70.1 | 9.0E-100 |
| DIP_10007410 | NA | | |
| DIP_10007411 | putative 3-oxoacyl-(acyl-carrier-protein) reductase protein [Eutypa lata UCREL1] | 72.2 | 1.0E-165 |
| DIP_10007412 | NA | | |
| DIP_10007413 | hypothetical protein PFICI_14885 [Pestalotiopsis fici W106-1] | 46.9 | 8.0E-79 |
| DIP_10007414 | NA | | |
| DIP_10007415 | NA | | |
| DIP_10007416 | NA | | |
| DIP_10007417 | hypothetical protein PTT_10149 [Pyrenophora teres f. teres 0-1] | 33.6 | 4.0E-21 |
| DIP_10007418 | hypothetical protein PFICI_14881 [Pestalotiopsis fici W106-1] | 62.8 | 2.0E-154 |
| DIP_10007419 | hypothetical protein VDAG_07723 [Verticillium dahliae VdLs.17] | 48.3 | 1.0E-159 |
| DIP_10007420 | putative autophagy protein atg27 protein [Eutypa lata UCREL1] | 50 | 9.0E-112 |
| DIP_10007421 | hypothetical protein TRIVIDRAFT_40129 [Trichoderma virens Gv29-8] | 39.2 | 1.0E-112 |
| DIP_10007422 | NA | | |
| DIP_10007423 | putative secretory pathway protein [Eutypa lata UCREL1] | 57.8 | 3.0E-76 |
| DIP_10007424 | putative origin recognition complex subunit 6 protein [Eutypa lata UCREL1] | 53.6 | 1.0E-136 |
| DIP_10007425 | hypothetical protein PFICI_08584 [Pestalotiopsis fici W106-1] | 33.3 | 3.0E-17 |
| DIP_10007426 | putative paf-acetylhydrolase family member protein [Eutypa lata UCREL1] | 45.1 | 5.0E-125 |
| DIP_10007427 | aldehyde dehydrogenase [Verticillium dahliae VdLs.17] | 83.3 | 0.0 |
| DIP_10007428 | NA | | |
| DIP_10007429 | hypothetical protein PFICI_06593 [Pestalotiopsis fici W106-1] | 52.7 | 0.0 |
| DIP_10007430 | putative gpi-anchored wall transfer protein [Eutypa lata UCREL1] | 70.3 | 0.0 |
| DIP_10007431 | hypothetical protein PFICI_13564 [Pestalotiopsis fici W106-1] | 65.5 | 0.0 |
| DIP_10007432 | hypothetical protein PFICI_13563 [Pestalotiopsis fici W106-1] | 38.1 | 8.0E-80 |
| DIP_10007433 | polysaccharide synthase [Colletotrichum gloeosporioides Nara gc5] | 71.4 | 0.0 |
| DIP_10007434 | hypothetical protein PFICI_06246 [Pestalotiopsis fici W106-1] | 50.1 | 1.0E-105 |
| DIP_10007435 | putative nima-interacting protein [Eutypa lata UCREL1] | 77.3 | 0.0 |
| DIP_10007436 | putative high mobility group protein [Eutypa lata UCREL1] | 65.7 | 8.0E-128 |
| DIP_10007437 | urease Ure [Neurospora crassa OR74A] | 68.1 | 0.0 |
| DIP_10007438 | transcriptional regulator [Colletotrichum gloeosporioides Nara gc5] | 65.7 | 0.0 |
| DIP_10007439 | hypothetical protein PFICI_09630 [Pestalotiopsis fici W106-1] | 49.2 | 2.0E-102 |
| DIP_10007440 | NA | | |
| DIP_10007441 | putative phosphate transporter protein [Eutypa lata UCREL1] | 69.1 | 1.0E-92 |
| DIP_10007442 | putative dna repair protein rad14 protein [Eutypa lata UCREL1] | 66.7 | 0.0 |
| DIP_10007443 | putative bifunctional p-450:nadph-p450 reductase protein [Eutypa lata UCREL1] | 63 | 0.0 |
| DIP_10007444 | putative duf1479 domain-containing protein [Eutypa lata UCREL1] | 50.1 | 0.0 |
| DIP_10007445 | hypothetical protein S40285_04828 [Stachybotrys chlorohalonata IBT 40285] | 56.2 | 2.0E-49 |
| DIP_10007446 | RNAse P Rpr2/Rpp21/SNM1 subunit domain-containing protein [Colletotrichum fioriniae PJ7] | 40.2 | 4.0E-40 |
| DIP_10007447 | hypothetical protein PFICI_03711 [Pestalotiopsis fici W106-1] | 76.6 | 0.0 |
| DIP_10007448 | hypothetical protein, partial [Streptomyces sp. NRRL F-2305] | 33.9 | 4.0E-12 |
| DIP_10007449 | hypothetical protein NECHADRAFT_82397 [Nectria haematococca mpVI 77-13-4] | 46.9 | 3.0E-67 |
| DIP_10007450 | hypothetical protein UCREL1_11475 [Eutypa lata UCREL1] | 36.8 | 5.0E-58 |
| DIP_10007451 | hypothetical protein TRIATDRAFT_172730, partial [Trichoderma atroviride IMI 206040] | 47.5 | 1.0E-121 |
| DIP_10007452 | hypothetical protein MGG_01874 [Magnaporthe oryzae 70-15] | 67.5 | 0.0 |
| DIP_10007453 | putative steroid 5 alpha-reductase protein [Eutypa lata UCREL1] | 73.3 | 5.0E-156 |
| DIP_10007454 | NA | | |
| DIP_10007455 | NA | | |
| DIP_10007456 | hypothetical protein PISL3812_07800 [Talaromyces islandicus] | 36.9 | 5.0E-87 |
| DIP_10007457 | hypothetical protein CFIO01_10981 [Colletotrichum fioriniae PJ7] | 44.7 | 2.0E-164 |
| DIP_10007458 | putative flap structure-specific endonuclease protein [Eutypa lata UCREL1] | 60.8 | 0.0 |
| DIP_10007459 | hypothetical protein PFICI_09032 [Pestalotiopsis fici W106-1] | 66.3 | 4.0E-153 |
| DIP_10007460 | hypothetical protein UCREL1_8851 [Eutypa lata UCREL1] | 67.5 | 6.0E-144 |
| DIP_10007461 | NA | | |
| DIP_10007462 | hypothetical protein PFICI_09029 [Pestalotiopsis fici W106-1] | 66 | 0.0 |
| DIP_10007463 | NA | | |
| DIP_10007464 | putative swi snf complex protein [Eutypa lata UCREL1] | 65 | 0.0 |
| DIP_10007465 | putative short chain dehydrogenase reductase protein [Eutypa lata UCREL1] | 56.5 | 1.0E-104 |
| DIP_10007466 | putative conidiophore development protein hyma protein [Eutypa lata UCREL1] | 92 | 0.0 |
| DIP_10007467 | putative isoleucyl-trna synthetase protein [Eutypa lata UCREL1] | 82.7 | 0.0 |
| DIP_10007468 | hypothetical protein AOR_1_880194 [Aspergillus oryzae RIB40] | 75.9 | 1.0E-25 |
| DIP_10007469 | NA | | |
| DIP_10007470 | NA | | |
| DIP_10007471 | hypothetical protein PFICI_09022 [Pestalotiopsis fici W106-1] | 58.5 | 0.0 |
| DIP_10007472 | WSC-domain-containing protein [Aureobasidium namibiae CBS 147.97] | 51.9 | 2.0E-146 |
| DIP_10007473 | oligopeptide transporter [Acanthamoeba castellanii str. Neff] | 38.2 | 5.0E-102 |
| DIP_10007474 | heat shock protein [Grosmannia clavigera kw1407] | 89.3 | 0.0 |
| DIP_10007475 | hypothetical protein UCREL1_8863 [Eutypa lata UCREL1] | 50 | 1.0E-47 |
| DIP_10007476 | NA | | |
| DIP_10007477 | Mitochondrial escape protein 2 [Pestalotiopsis fici W106-1] | 72.2 | 0.0 |
| DIP_10007478 | putative psp1 domain-containing protein [Diaporthe ampelina] | 63.1 | 1.0E-146 |
| DIP_10007479 | putative hira-interacting protein 5 protein [Eutypa lata UCREL1] | 70.1 | 1.0E-160 |
| DIP_10007480 | putative zinc finger (hit type) family protein [Eutypa lata UCREL1] | 36.4 | 8.0E-39 |
| DIP_10007481 | NA | | |
| DIP_10007482 | NA | | |
| DIP_10007483 | hypothetical protein UCREL1_4997 [Eutypa lata UCREL1] | 61.4 | 7.0E-151 |
| DIP_10007484 | putative glycosyl hydrolase family 71 protein [Eutypa lata UCREL1] | 51.3 | 1.0E-123 |
| DIP_10007485 | NA | | |
| DIP_10007486 | hypothetical protein NECHADRAFT_77482 [Nectria haematococca mpVI 77-13-4] | 40.6 | 4.0E-90 |
| DIP_10007487 | NA | | |
| DIP_10007488 | putative cys met metabolism pyridoxal phosphate-dependent enzyme protein [Eutypa lata UCREL1] | 42.7 | 5.0E-49 |
| DIP_10007489 | putative AT hook domain-containing protein [Colletotrichum sublineola] | 38.2 | 2.0E-105 |
| DIP_10007490 | NA | | |
| DIP_10007491 | putative er lumen protein retaining receptor protein [Eutypa lata UCREL1] | 68.6 | 2.0E-169 |
| DIP_10007492 | hypothetical protein Moror_2614 [Moniliophthora roreri MCA 2997] | 32.3 | 2.0E-19 |
| DIP_10007493 | hypothetical protein SBOR_0325 [Sclerotinia borealis F-4157] | 36.3 | 8.0E-95 |
| DIP_10007494 | hypothetical protein HIM_05423 [Hirsutella minnesotensis 3608] | 34.4 | 5.0E-42 |
| DIP_10007495 | NA | | |
| DIP_10007496 | NA | | |
| DIP_10007497 | hypothetical protein PFICI_13313 [Pestalotiopsis fici W106-1] | 49.8 | 0.0 |
| DIP_10007498 | putative lysyl-trna synthetase protein [Eutypa lata UCREL1] | 57.8 | 9.0E-176 |
| DIP_10007499 | putative mitochondrial import inner membrane translocase subunit tim10 protein [Eutypa lata UCREL1] | 87.8 | 2.0E-49 |
| DIP_10007500 | putative adenylate kinase protein [Eutypa lata UCREL1] | 73 | 2.0E-150 |
| DIP_10007501 | NA | | |
| DIP_10007502 | putative immunogenic protein [Eutypa lata UCREL1] | 62.7 | 7.0E-171 |
| DIP_10007503 | Adenine phosphoribosyltransferase [Fusarium oxysporum f. sp. cubense race 1] | 51.2 | 3.0E-61 |
| DIP_10007504 | NA | | |
| DIP_10007505 | hypothetical protein CH063_00371 [Colletotrichum higginsianum] | 60.7 | 0.0 |
| DIP_10007506 | putative peroxisomal biogenesis factor 2 protein [Eutypa lata UCREL1] | 72.3 | 0.0 |
| DIP_10007507 | putative importin-beta domain-containing protein [Eutypa lata UCREL1] | 59.3 | 0.0 |
| DIP_10007508 | putative membrane protein [Eutypa lata UCREL1] | 78.8 | 3.0E-59 |
| DIP_10007509 | hypothetical protein PV11_04388 [Exophiala sideris] | 32.9 | 1.0E-09 |
| DIP_10007510 | putative tos1 protein [Eutypa lata UCREL1] | 58.3 | 0.0 |
| DIP_10007511 | NA | | |
| DIP_10007512 | NA | | |
| DIP_10007513 | NA | | |
| DIP_10007514 | hypothetical protein CGLO_08867 [Colletotrichum gloeosporioides Cg-14] | 55.4 | 1.0E-14 |
| DIP_10007515 | putative purine-cytosine permease protein [Togninia minima UCRPA7] | 50 | 0.0 |
| DIP_10007516 | hypothetical protein PFICI_13299 [Pestalotiopsis fici W106-1] | 72.8 | 3.0E-175 |
| DIP_10007517 | putative nmral1 protein [Eutypa lata UCREL1] | 65.3 | 1.0E-144 |
| DIP_10007518 | hypothetical protein PFICI_14721 [Pestalotiopsis fici W106-1] | 55.7 | 9.0E-129 |
| DIP_10007519 | DNA excision repair protein [Colletotrichum fioriniae PJ7] | 66.8 | 0.0 |
| DIP_10007520 | hypothetical protein PFICI_06981 [Pestalotiopsis fici W106-1] | 71.2 | 5.0E-54 |
| DIP_10007521 | hypothetical protein PFICI_06985 [Pestalotiopsis fici W106-1] | 58.6 | 0.0 |
| DIP_10007522 | hypothetical protein UCREL1_7505 [Eutypa lata UCREL1] | 52.2 | 4.0E-50 |
| DIP_10007523 | putative short chain dehydrogenase protein [Eutypa lata UCREL1] | 74.3 | 9.0E-153 |
| DIP_10007524 | hypothetical protein PFICI_13290 [Pestalotiopsis fici W106-1] | 45 | 0.0 |
| DIP_10007525 | Sorting nexin-4 [Pestalotiopsis fici W106-1] | 72 | 0.0 |
| DIP_10007526 | hypothetical protein PFICI_06256 [Pestalotiopsis fici W106-1] | 55.7 | 0.0 |
| DIP_10007527 | hypothetical protein PFICI_06255 [Pestalotiopsis fici W106-1] | 58.5 | 0.0 |
| DIP_10007528 | hypothetical protein PFICI_06254 [Pestalotiopsis fici W106-1] | 62.1 | 9.0E-41 |
| DIP_10007529 | putative septum formation protein maf protein [Eutypa lata UCREL1] | 60.1 | 1.0E-109 |
| DIP_10007530 | hypothetical protein MGG_00752 [Magnaporthe oryzae 70-15] | 75.4 | 4.0E-95 |
| DIP_10007531 | putative px domain-containing protein [Eutypa lata UCREL1] | 43.6 | 1.0E-60 |
| DIP_10007532 | hypothetical protein PFICI_06980 [Pestalotiopsis fici W106-1] | 48.2 | 4.0E-116 |
| DIP_10007533 | hypothetical protein CFIO01_01589 [Colletotrichum fioriniae PJ7] | 33.8 | 4.0E-63 |
| DIP_10007534 | NA | | |
| DIP_10007535 | hypothetical protein PFICI_06971 [Pestalotiopsis fici W106-1] | 71.5 | 5.0E-137 |
| DIP_10007536 | DNA repair protein [Colletotrichum orbiculare MAFF 240422] | 65.8 | 5.0E-33 |
| DIP_10007537 | putative histone acetyltransferase gcn5 protein [Eutypa lata UCREL1] | 90 | 0.0 |
| DIP_10007538 | NA | | |
| DIP_10007539 | hypothetical protein PV05_01623 [Exophiala xenobiotica] | 46.9 | 2.0E-116 |
| DIP_10007540 | hypothetical protein UCREL1_10156 [Eutypa lata UCREL1] | 64.4 | 3.0E-111 |
| DIP_10007541 | putative estradiol 17-beta-dehydrogenase protein [Eutypa lata UCREL1] | 36.5 | 3.0E-55 |
| DIP_10007542 | glutaminyl-peptide cyclotransferase [Ophiostoma piceae UAMH 11346] | 50.5 | 5.0E-124 |
| DIP_10007543 | hypothetical protein PFICI_06978 [Pestalotiopsis fici W106-1] | 77.5 | 0.0 |
| DIP_10007544 | hypothetical protein PITC_054370 [Penicillium italicum] | 31.9 | 4.0E-14 |
| DIP_10007545 | putative cdp-alcohol phosphatidyltransferase protein [Eutypa lata UCREL1] | 49 | 3.0E-57 |
| DIP_10007546 | Sphingolipid long chain base-responsive protein PIL1 [Pestalotiopsis fici W106-1] | 88.4 | 0.0 |
| DIP_10007547 | putative set domain-containing protein [Diaporthe ampelina] | 38.7 | 6.0E-107 |
| DIP_10007548 | serine threonine-protein phosphatase pp-z [Colletotrichum gloeosporioides Nara gc5] | 79.1 | 0.0 |
| DIP_10007549 | NA | | |
| DIP_10007550 | putative integral membrane protein [Eutypa lata UCREL1] | 51.7 | 4.0E-124 |
| DIP_10007551 | hypothetical protein PFICI_13272 [Pestalotiopsis fici W106-1] | 33.9 | 0.0 |
| DIP_10007552 | hypothetical protein UCREL1_198 [Eutypa lata UCREL1] | 50.8 | 1.0E-62 |
| DIP_10007553 | putative glutathione s- protein [Eutypa lata UCREL1] | 66.2 | 1.0E-102 |
| DIP_10007554 | NA | | |
| DIP_10007555 | NA | | |
| DIP_10007556 | hypothetical protein PFICI_13286 [Pestalotiopsis fici W106-1] | 53.9 | 4.0E-74 |
| DIP_10007557 | succinate-semialdehyde dehydrogenase [Pestalotiopsis fici W106-1] | 78.1 | 0.0 |
| DIP_10007558 | hypothetical protein PFICI_13282 [Pestalotiopsis fici W106-1] | 60 | 8.0E-113 |
| DIP_10007559 | hypothetical protein PFICI_13281 [Pestalotiopsis fici W106-1] | 73.1 | 0.0 |
| DIP_10007560 | hypothetical protein Cob_13062 [Colletotrichum orbiculare MAFF 240422] | 34.7 | 3.0E-13 |
| DIP_10007561 | hypothetical protein PFICI_13275 [Pestalotiopsis fici W106-1] | 59.9 | 0.0 |
| DIP_10007562 | NA | | |
| DIP_10007563 | hypothetical protein PFICI_13273 [Pestalotiopsis fici W106-1] | 64.4 | 0.0 |
| DIP_10007564 | hypothetical protein PFICI_13289 [Pestalotiopsis fici W106-1] | 57.5 | 0.0 |
| DIP_10007565 | hypothetical protein CGLO_17074 [Colletotrichum gloeosporioides Cg-14] | 31.9 | 3.0E-16 |
| DIP_10007566 | putative pheromone processing endoprotease protein [Eutypa lata UCREL1] | 67.4 | 0.0 |
| DIP_10007567 | NA | | |
| DIP_10007568 | hypothetical protein PFICI_15291 [Pestalotiopsis fici W106-1] | 74.6 | 2.0E-146 |
| DIP_10007569 | zn 2cys6 transcription factor [Colletotrichum gloeosporioides Nara gc5] | 41.3 | 2.0E-155 |
| DIP_10007570 | hypothetical protein PFICI_06260 [Pestalotiopsis fici W106-1] | 59.1 | 6.0E-83 |
| DIP_10007571 | putative u3 snornp-associated protein utp11 protein [Eutypa lata UCREL1] | 54.9 | 4.0E-85 |
| DIP_10007572 | hypothetical protein CFIO01_08378 [Colletotrichum fioriniae PJ7] | 49.9 | 8.0E-119 |
| DIP_10007573 | NA | | |
| DIP_10007574 | NA | | |
| DIP_10007575 | hypothetical protein PFICI_06078 [Pestalotiopsis fici W106-1] | 66 | 0.0 |
| DIP_10007576 | hypothetical protein THITE_2057447 [Thielavia terrestris NRRL 8126] | 48.5 | 2.0E-159 |
| DIP_10007577 | putative 1-phosphatidylinositol-3-phosphate 5-kinase fab1 protein [Togninia minima UCRPA7] | 63.1 | 0.0 |
| DIP_10007578 | NA | | |
| DIP_10007579 | NA | | |
| DIP_10007580 | putative eukaryotic translation initiation factor 4e-1 protein [Eutypa lata UCREL1] | 83.6 | 3.0E-155 |
| DIP_10007581 | putative 2-succinylbenzoate- ligase protein [Eutypa lata UCREL1] | 61 | 0.0 |
| DIP_10007582 | hypothetical protein UCREL1_8443 [Eutypa lata UCREL1] | 77.7 | 1.0E-55 |
| DIP_10007583 | putative phosphoinositide phosphatase protein [Eutypa lata UCREL1] | 76.9 | 0.0 |
| DIP_10007584 | putative short chain dehydrogenase protein [Togninia minima UCRPA7] | 53.7 | 1.0E-99 |
| DIP_10007585 | putative gtp-binding protein ypt5 protein [Eutypa lata UCREL1] | 92.8 | 3.0E-151 |
| DIP_10007586 | putative mitochondrion protein [Eutypa lata UCREL1] | 64.9 | 0.0 |
| DIP_10007587 | putative upf0183 domain containing protein [Eutypa lata UCREL1] | 38.2 | 6.0E-82 |
| DIP_10007588 | transcription factor AATF/Che-1 [Gaeumannomyces graminis var. tritici R3-111a-1] | 47 | 2.0E-152 |
| DIP_10007589 | hypothetical protein PFICI_09957 [Pestalotiopsis fici W106-1] | 78.9 | 6.0E-176 |
| DIP_10007590 | hypothetical protein PFICI_09958 [Pestalotiopsis fici W106-1] | 71.7 | 2.0E-144 |
| DIP_10007591 | NA | | |
| DIP_10007592 | NA | | |
| DIP_10007593 | putative carboxylesterase protein [Eutypa lata UCREL1] | 50.7 | 6.0E-159 |
| DIP_10007594 | hypothetical protein THAR02_03909 [Trichoderma harzianum] | 34.7 | 4.0E-69 |
| DIP_10007595 | putative -beta- protein [Eutypa lata UCREL1] | 67.1 | 0.0 |
| DIP_10007596 | Glutathione-dependent formaldehyde-activating family GFA [Macrophomina phaseolina MS6] | 52.6 | 5.0E-48 |
| DIP_10007597 | cupin 2 domain-containing protein [Colletotrichum orbiculare MAFF 240422] | 58.5 | 6.0E-46 |
| DIP_10007598 | hypothetical protein PFICI_10174 [Pestalotiopsis fici W106-1] | 30.1 | 5.0E-25 |
| DIP_10007599 | hypothetical protein PFICI_04133 [Pestalotiopsis fici W106-1] | 50.9 | 0.0 |
| DIP_10007600 | hypothetical protein GLRG_08804 [Colletotrichum graminicola M1.001] | 48.6 | 8.0E-112 |
| DIP_10007601 | ubiquitin-conjugating enzyme h [Grosmannia clavigera kw1407] | 83.8 | 3.0E-119 |
| DIP_10007602 | hypothetical protein PFICI_15155 [Pestalotiopsis fici W106-1] | 65.6 | 0.0 |
| DIP_10007603 | hypothetical protein CGLO_16018 [Colletotrichum gloeosporioides Cg-14] | 72.1 | 2.0E-163 |
| DIP_10007604 | putative brefeldin a resistance protein [Eutypa lata UCREL1] | 81 | 0.0 |
| DIP_10007605 | putative casein kinase ii subunit beta-1 protein [Eutypa lata UCREL1] | 87.4 | 0.0 |
| DIP_10007606 | hypothetical protein COCMIDRAFT_31446 [Bipolaris oryzae ATCC 44560] | 63.4 | 2.0E-71 |
| DIP_10007607 | hypothetical protein PFICI_14395 [Pestalotiopsis fici W106-1] | 74.5 | 0.0 |
| DIP_10007608 | hypothetical protein UCREL1_9047 [Eutypa lata UCREL1] | 53.3 | 8.0E-70 |
| DIP_10007609 | putative branched-chain-amino-acid aminotransferase protein [Eutypa lata UCREL1] | 61.1 | 2.0E-173 |
| DIP_10007610 | SCY1 protein kinase [Gaeumannomyces graminis var. tritici R3-111a-1] | 73.6 | 0.0 |
| DIP_10007611 | putative histone deacetylase protein [Eutypa lata UCREL1] | 65.7 | 0.0 |
| DIP_10007612 | putative calcium p-type atpase protein [Eutypa lata UCREL1] | 78.7 | 0.0 |
| DIP_10007613 | hypothetical protein PFICI_10317 [Pestalotiopsis fici W106-1] | 63.8 | 0.0 |
| DIP_10007614 | hypothetical protein PFICI_14811 [Pestalotiopsis fici W106-1] | 41.4 | 4.0E-98 |
| DIP_10007615 | NA | | |
| DIP_10007616 | hypothetical protein UCRPA7_748 [Togninia minima UCRPA7] | 49.2 | 1.0E-126 |
| DIP_10007617 | hypothetical protein SBOR_5584 [Sclerotinia borealis F-4157] | 57.4 | 1.0E-44 |
| DIP_10007618 | hypothetical protein Cob_07731 [Colletotrichum orbiculare MAFF 240422] | 44.5 | 9.0E-110 |
| DIP_10007619 | putative dna replication protein [Eutypa lata UCREL1] | 55.6 | 0.0 |
| DIP_10007620 | hypothetical protein UCREL1_1885 [Eutypa lata UCREL1] | 58.4 | 0.0 |
| DIP_10007621 | hypothetical protein GGTG_04818 [Gaeumannomyces graminis var. tritici R3-111a-1] | 31.5 | 4.0E-27 |
| DIP_10007622 | hypothetical protein MGG_11897 [Magnaporthe oryzae 70-15] | 83.3 | 5.0E-84 |
| DIP_10007623 | nuclear transport factor NTF-2, putative [Talaromyces stipitatus ATCC 10500] | 39.3 | 5.0E-30 |
| DIP_10007624 | NA | | |
| DIP_10007625 | putative cue domain-containing protein [Eutypa lata UCREL1] | 66.2 | 2.0E-173 |
| DIP_10007626 | hypothetical protein THITE_2123668 [Thielavia terrestris NRRL 8126] | 72.6 | 0.0 |
| DIP_10007627 | hypothetical protein MANI_007873 [Metarhizium anisopliae] | 44.6 | 2.0E-159 |
| DIP_10007628 | hypothetical protein PFICI_08628 [Pestalotiopsis fici W106-1] | 77.3 | 2.0E-102 |
| DIP_10007629 | hypothetical protein PFICI_09417 [Pestalotiopsis fici W106-1] | 45.9 | 2.0E-124 |
| DIP_10007630 | hypothetical protein UCDDA912_g01855 [Diaporthe ampelina] | 39.2 | 3.0E-43 |
| DIP_10007631 | Ankyrin repeat protein nuc-2 [Pestalotiopsis fici W106-1] | 77.6 | 0.0 |
| DIP_10007632 | Exportin-1 [Pestalotiopsis fici W106-1] | 83.2 | 0.0 |
| DIP_10007633 | NA | | |
| DIP_10007634 | putative alanine-glyoxylate aminotransferase protein [Eutypa lata UCREL1] | 77.5 | 0.0 |
| DIP_10007635 | putative vacuolar protein sorting-associated protein [Eutypa lata UCREL1] | 83.6 | 6.0E-125 |
| DIP_10007636 | hypothetical protein PFICI_09414 [Pestalotiopsis fici W106-1] | 44 | 4.0E-132 |
| DIP_10007637 | hypothetical protein PFICI_09415 [Pestalotiopsis fici W106-1] | 71.3 | 2.0E-62 |
| DIP_10007638 | NA | | |
| DIP_10007639 | hypothetical protein PFICI_11383 [Pestalotiopsis fici W106-1] | 72.8 | 1.0E-122 |
| DIP_10007640 | Allantoicase [Pestalotiopsis fici W106-1] | 83.7 | 0.0 |
| DIP_10007641 | putative dil and ankyrin domain-containing protein [Eutypa lata UCREL1] | 80.9 | 0.0 |
| DIP_10007642 | Pc13g02230 [Penicillium rubens Wisconsin 54-1255] | 38.1 | 4.0E-55 |
| DIP_10007643 | NA | | |
| DIP_10007644 | hypothetical protein PFICI_11384 [Pestalotiopsis fici W106-1] | 52.4 | 0.0 |
| DIP_10007645 | glyoxalase family protein [Zymoseptoria brevis] | 43.5 | 9.0E-35 |
| DIP_10007646 | hypothetical protein PFICI_07177 [Pestalotiopsis fici W106-1] | 45.9 | 5.0E-63 |
| DIP_10007647 | NA | | |
| DIP_10007648 | hypothetical protein PFICI_08471 [Pestalotiopsis fici W106-1] | 42 | 0.0 |
| DIP_10007649 | NA | | |
| DIP_10007650 | putative flavohemo protein [Eutypa lata UCREL1] | 46.4 | 1.0E-109 |
| DIP_10007651 | putative tetraspanin tsp3 protein [Eutypa lata UCREL1] | 36.7 | 4.0E-39 |
| DIP_10007652 | NA | | |
| DIP_10007653 | leucine aminopeptidase 2 [Coccidioides immitis RS] | 44.6 | 3.0E-80 |
| DIP_10007654 | NA | | |
| DIP_10007655 | hypothetical protein CFIO01_05492 [Colletotrichum fioriniae PJ7] | 84.1 | 0.0 |
| DIP_10007656 | NA | | |
| DIP_10007657 | putative ribonuclease 3 protein [Eutypa lata UCREL1] | 46.5 | 5.0E-73 |
| DIP_10007658 | hypothetical protein PFICI_08516 [Pestalotiopsis fici W106-1] | 63.7 | 2.0E-40 |
| DIP_10007659 | hypothetical protein PFICI_08520 [Pestalotiopsis fici W106-1] | 38.9 | 2.0E-45 |
| DIP_10007660 | putative protein phosphatase 2c protein [Eutypa lata UCREL1] | 81.3 | 0.0 |
| DIP_10007661 | NA | | |
| DIP_10007662 | NA | | |
| DIP_10007663 | hypothetical protein NECHADRAFT_37789 [Nectria haematococca mpVI 77-13-4] | 46.9 | 5.0E-91 |
| DIP_10007664 | putative cytokinesis regulator protein [Togninia minima UCRPA7] | 63.9 | 6.0E-125 |
| DIP_10007665 | hypothetical protein PFICI_09387 [Pestalotiopsis fici W106-1] | 41.9 | 3.0E-70 |
| DIP_10007666 | NA | | |
| DIP_10007667 | hypothetical protein UCREL1_281 [Eutypa lata UCREL1] | 70.3 | 0.0 |
| DIP_10007668 | putative ubiquitin carrier protein [Eutypa lata UCREL1] | 48 | 8.0E-92 |
| DIP_10007669 | NA | | |
| DIP_10007670 | NADH-ubiquinone oxidoreductase subunit A [Pestalotiopsis fici W106-1] | 76.1 | 0.0 |
| DIP_10007671 | predicted protein [Nectria haematococca mpVI 77-13-4] | 72.7 | 2.0E-76 |
| DIP_10007672 | putative wd40 repeat-like-containing domain protein [Eutypa lata UCREL1] | 59.6 | 0.0 |
| DIP_10007673 | hypothetical protein TRIREDRAFT_29346 [Trichoderma reesei QM6a] | 40.3 | 8.0E-68 |
| DIP_10007674 | NA | | |
| DIP_10007675 | NA | | |
| DIP_10007676 | NA | | |
| DIP_10007677 | putative short-chain dehydrogenase reductase [Diaporthe ampelina] | 51.7 | 5.0E-88 |
| DIP_10007678 | putative heat shock protein 60 protein [Eutypa lata UCREL1] | 92.4 | 0.0 |
| DIP_10007679 | cofilin/tropomyosin-type actin-binding protein [Colletotrichum graminicola M1.001] | 85.6 | 1.0E-94 |
| DIP_10007680 | hypothetical protein UCRPA7_185 [Togninia minima UCRPA7] | 41.3 | 4.0E-51 |
| DIP_10007681 | putative trna (cytosine-5-)-methyltransferase ncl1 protein [Eutypa lata UCREL1] | 70.8 | 0.0 |
| DIP_10007682 | putative 4-coumarate:coenzyme a ligase protein [Eutypa lata UCREL1] | 60.6 | 2.0E-38 |
| DIP_10007683 | NA | | |

| | | | |
|---|---|---|---|
| DIP_10007684 | NA | | |
| DIP_10007685 | NA | | |
| DIP_10007686 | NA | | |
| DIP_10007687 | hypothetical protein FAVG1_05533 [Fusarium avenaceum] | 44.3 | 6.0E-80 |
| DIP_10007688 | alpha-ketoglutarate-dependent taurine dioxygenase [Colletotrichum gloeosporioides Nara gc5] | 80.5 | 0.0 |
| DIP_10007689 | hypothetical protein PFICI_10661 [Pestalotiopsis fici W106-1] | 55.7 | 1.0E-161 |
| DIP_10007690 | hypothetical protein PFICI_10660 [Pestalotiopsis fici W106-1] | 62.4 | 2.0E-146 |
| DIP_10007691 | hypothetical protein PFICI_14262 [Pestalotiopsis fici W106-1] | 60.3 | 0.0 |
| DIP_10007692 | putative high-affinity nicotinic acid [Phaeomoniella chlamydospora] | 69.4 | 0.0 |
| DIP_10007693 | NA | | |
| DIP_10007694 | NA | | |
| DIP_10007695 | nacht and ankyrin domain protein [Colletotrichum gloeosporioides Nara gc5] | 30.4 | 2.0E-121 |
| DIP_10007696 | NA | | |
| DIP_10007697 | NA | | |
| DIP_10007698 | immunomodulatory protein [Dichomitus squalens LYAD-421 SS1] | 55.7 | 1.0E-33 |
| DIP_10007699 | MFS multidrug transporter [Colletotrichum orbiculare MAFF 240422] | 55.3 | 0.0 |
| DIP_10007700 | NA | | |
| DIP_10007701 | hypothetical protein TRIATDRAFT_91724 [Trichoderma atroviride IMI 206040] | 50 | 5.0E-18 |
| DIP_10007702 | hypothetical protein PFICI_15255 [Pestalotiopsis fici W106-1] | 52.4 | 0.0 |
| DIP_10007703 | hypothetical protein G647_05761 [Cladophialophora carrionii CBS 160.54] | 59.4 | 2.0E-52 |
| DIP_10007704 | NA | | |
| DIP_10007705 | NA | | |
| DIP_10007706 | predicted protein [Nectria haematococca mpVI 77-13-4] | 87.7 | 0.0 |
| DIP_10007707 | putative duf1168 domain protein [Togninia minima UCRPA7] | 51.7 | 5.0E-58 |
| DIP_10007708 | putative phosphoglycerate mutase family domain-containing protein [Eutypa lata UCREL1] | 63.1 | 0.0 |
| DIP_10007709 | Casein kinase II subunit alpha [Pestalotiopsis fici W106-1] | 94.7 | 0.0 |
| DIP_10007710 | NA | | |
| DIP_10007711 | hypothetical protein PFICI_07169 [Pestalotiopsis fici W106-1] | 63 | 0.0 |
| DIP_10007712 | NA | | |
| DIP_10007713 | putative peptidyl-prolyl cis-trans isomerase h protein [Togninia minima UCRPA7] | 85.7 | 1.0E-114 |
| DIP_10007714 | hypothetical protein FOXB_12030 [Fusarium oxysporum Fo5176] | 49.7 | 2.0E-91 |
| DIP_10007715 | hypothetical protein UCREL1_10070 [Eutypa lata UCREL1] | 46.1 | 4.0E-81 |
| DIP_10007716 | putative golgi complex component protein [Eutypa lata UCREL1] | 71 | 0.0 |
| DIP_10007717 | putative class ii aldolase protein [Eutypa lata UCREL1] | 72.2 | 4.0E-165 |
| DIP_10007718 | NA | | |
| DIP_10007719 | hypothetical protein PFICI_14962 [Pestalotiopsis fici W106-1] | 58.2 | 1.0E-36 |
| DIP_10007720 | NA | | |
| DIP_10007721 | NA | | |
| DIP_10007722 | putative ccr4-not transcription complex subunit protein [Eutypa lata UCREL1] | 70.7 | 0.0 |
| DIP_10007723 | zinc-binding dehydrogenase [Colletotrichum higginsianum] | 57.9 | 1.0E-129 |
| DIP_10007724 | hypothetical protein PFICI_10358 [Pestalotiopsis fici W106-1] | 60.6 | 0.0 |
| DIP_10007725 | putative methyltransferase oms1 protein [Eutypa lata UCREL1] | 53.3 | 1.0E-144 |
| DIP_10007726 | hypothetical protein UCREL1_4114 [Eutypa lata UCREL1] | 65.4 | 7.0E-145 |
| DIP_10007727 | hypothetical protein S40285_09494 [Stachybotrys chlorohalonata IBT 40285] | 42 | 2.0E-148 |
| DIP_10007728 | hypothetical protein PFICI_10353 [Pestalotiopsis fici W106-1] | 72.8 | 0.0 |
| DIP_10007729 | NA | | |
| DIP_10007730 | NA | | |
| DIP_10007731 | hypothetical protein CSUB01_02401 [Colletotrichum sublineola] | 30.1 | 9.0E-17 |
| DIP_10007732 | NA | | |
| DIP_10007733 | putative transcription factor protein [Eutypa lata UCREL1] | 74.7 | 0.0 |
| DIP_10007734 | hypothetical protein PFICI_10349 [Pestalotiopsis fici W106-1] | 47.8 | 6.0E-87 |
| DIP_10007735 | putative glycoside hydrolase family 16 protein [Eutypa lata UCREL1] | 56.4 | 5.0E-137 |
| DIP_10007736 | Lysine--tRNA ligase [Pestalotiopsis fici W106-1] | 80.3 | 0.0 |
| DIP_10007737 | putative cytochrome c oxidase assembly protein [Eutypa lata UCREL1] | 65.2 | 0.0 |
| DIP_10007738 | cation efflux family protein [Colletotrichum fioriniae PJ7] | 75.7 | 0.0 |
| DIP_10007739 | putative catechol dioxygenase protein [Eutypa lata UCREL1] | 83.6 | 7.0E-168 |
| DIP_10007740 | putative 3-carboxy- -muconate cyclase protein [Eutypa lata UCREL1] | 83.3 | 0.0 |
| DIP_10007741 | hypothetical protein TRIVIDRAFT_222107 [Trichoderma virens Gv29-8] | 30.5 | 2.0E-20 |
| DIP_10007742 | putative duf21 and cbs domain protein [Eutypa lata UCREL1] | 64.3 | 0.0 |
| DIP_10007743 | hypothetical protein PFICI_10340 [Pestalotiopsis fici W106-1] | 76.7 | 0.0 |
| DIP_10007744 | putative copper fist dna binding domain protein [Eutypa lata UCREL1] | 59.6 | 5.0E-75 |
| DIP_10007745 | putative fmn dependent protein [Neofusicoccum parvum UCRNP2] | 59.8 | 0.0 |
| DIP_10007746 | NA | | |
| DIP_10007747 | hypothetical protein PFICI_14304 [Pestalotiopsis fici W106-1] | 60.2 | 3.0E-74 |
| DIP_10007748 | hypothetical protein PFICI_10411 [Pestalotiopsis fici W106-1] | 61.5 | 0.0 |
| DIP_10007749 | putative heterochromatin protein one protein [Eutypa lata UCREL1] | 60.3 | 2.0E-104 |
| DIP_10007750 | hypothetical protein FVEG_01875 [Fusarium verticillioides 7600] | 44.4 | 1.0E-111 |
| DIP_10007751 | alkaline serine protease [Hirsutella rhossiliensis] | 46.1 | 5.0E-96 |
| DIP_10007752 | NA | | |
| DIP_10007753 | hypothetical protein S40285_09395 [Stachybotrys chlorohalonata IBT 40285] | 85.1 | 5.0E-170 |
| DIP_10007754 | NADP-dependent L-serine/L-allo-threonine dehydrogenase ydfG [Colletotrichum fioriniae PJ7] | 53.4 | 2.0E-89 |
| DIP_10007755 | putative cytochrome p450 protein [Eutypa lata UCREL1] | 41.7 | 2.0E-30 |
| DIP_10007756 | hypothetical protein FOXB_08604 [Fusarium oxysporum Fo5176] | 42.5 | 3.0E-113 |
| DIP_10007757 | LysP [Aspergillus parasiticus SU-1] | 62 | 2.0E-160 |
| DIP_10007758 | hypothetical protein PV07_07617 [Cladophialophora immunda] | 76 | 1.0E-151 |
| DIP_10007759 | hypothetical protein PFICI_00210 [Pestalotiopsis fici W106-1] | 65.3 | 0.0 |
| DIP_10007760 | hypothetical protein A1O5_09608 [Cladophialophora psammophila CBS 110553] | 48.1 | 3.0E-155 |
| DIP_10007761 | hypothetical protein PV11_05447 [Exophiala sideris] | 71.8 | 1.0E-142 |
| DIP_10007762 | Sugar/inositol transporter [Penicillium solitum] | 82.8 | 0.0 |
| DIP_10007763 | cutinase [Colletotrichum gloeosporioides Cg-14] | 61.1 | 7.0E-81 |
| DIP_10007764 | putative phospholipid metabolism enzyme regulator protein [Eutypa lata UCREL1] | 56.3 | 2.0E-159 |
| DIP_10007765 | Serine/threonine-protein kinase SSN3 [Pestalotiopsis fici W106-1] | 77.4 | 0.0 |
| DIP_10007766 | hypothetical protein SAPIO_CDS2687 [Scedosporium apiospermum] | 91.6 | 0.0 |
| DIP_10007767 | hypothetical protein PFICI_05970 [Pestalotiopsis fici W106-1] | 50.9 | 2.0E-76 |
| DIP_10007768 | putative mfs glucose protein [Eutypa lata UCREL1] | 68.8 | 0.0 |
| DIP_10007769 | ADP-ribosylation factor 6 [Magnaporthe oryzae 70-15] | 85.3 | 2.0E-116 |
| DIP_10007770 | NA | | |
| DIP_10007771 | NA | | |
| DIP_10007772 | putative c2h2 finger domain-containing protein [Eutypa lata UCREL1] | 48.8 | 1.0E-70 |
| DIP_10007773 | hypothetical protein PFICI_07109 [Pestalotiopsis fici W106-1] | 47.7 | 6.0E-172 |
| DIP_10007774 | NADH-ubiquinone oxidoreductase subunit A [Pestalotiopsis fici W106-1] | 83.4 | 3.0E-165 |
| DIP_10007775 | putative merozoite capping protein-1 protein [Eutypa lata UCREL1] | 75.1 | 2.0E-98 |
| DIP_10007776 | ketol-acid mitochondrial precursor [Colletotrichum orbiculare MAFF 240422] | 84.7 | 0.0 |
| DIP_10007777 | ubiquitin-conjugating enzyme [Colletotrichum graminicola M1.001] | 82.4 | 9.0E-114 |
| DIP_10007778 | hypothetical protein PFICI_07195 [Pestalotiopsis fici W106-1] | 46.5 | 0.0 |
| DIP_10007779 | hypothetical protein ASPNIDRAFT_45042 [Aspergillus niger ATCC 1015] | 52.7 | 4.0E-124 |
| DIP_10007780 | succinate-semialdehyde dehydrogenase [Colletotrichum gloeosporioides Nara gc5] | 60.9 | 0.0 |
| DIP_10007781 | Zn(II)2Cys6 transcription factor [Aspergillus oryzae RIB40] | 45.4 | 8.0E-139 |
| DIP_10007782 | hypothetical protein CGLO_05822 [Colletotrichum gloeosporioides Cg-14] | 58.8 | 3.0E-108 |
| DIP_10007783 | flavin containing amine oxidoreductase [Colletotrichum gloeosporioides Cg-14] | 63.5 | 0.0 |
| DIP_10007784 | hypothetical protein PFICI_06399 [Pestalotiopsis fici W106-1] | 66.7 | 8.0E-179 |
| DIP_10007785 | putative gtp-binding protein [Eutypa lata UCREL1] | 71.4 | 3.0E-118 |
| DIP_10007786 | putative hd domain-containing protein [Eutypa lata UCREL1] | 70.7 | 2.0E-102 |
| DIP_10007787 | putative methionine aminopeptidase protein [Eutypa lata UCREL1] | 78.4 | 0.0 |
| DIP_10007788 | NA | | |
| DIP_10007789 | putative poly + rna export protein [Eutypa lata UCREL1] | 83.2 | 0.0 |
| DIP_10007790 | hypothetical protein PFICI_06006 [Pestalotiopsis fici W106-1] | 53.2 | 4.0E-139 |
| DIP_10007791 | Tubulin gamma chain [Pestalotiopsis fici W106-1] | 94.2 | 0.0 |
| DIP_10007792 | hypothetical protein [Podospora anserina S mat+] | 71.4 | 8.0E-30 |
| DIP_10007793 | NA | | |
| DIP_10007794 | putative nuclear envelope protein cut8 protein [Eutypa lata UCREL1] | 76.3 | 7.0E-154 |
| DIP_10007795 | putative 5 -nucleotidase protein [Eutypa lata UCREL1] | 64.1 | 0.0 |
| DIP_10007796 | putative snare domain-containing protein [Eutypa lata UCREL1] | 66.7 | 2.0E-121 |
| DIP_10007797 | pal1-like protein [Grosmannia clavigera kw1407] | 33 | 3.0E-73 |
| DIP_10007798 | hypothetical protein CFIO01_03591 [Colletotrichum fioriniae PJ7] | 46.3 | 7.0E-39 |
| DIP_10007799 | hypothetical protein PFICI_06158 [Pestalotiopsis fici W106-1] | 69 | 0.0 |
| DIP_10007800 | oxidoreductase, partial [Klebsiella pneumoniae] | 38.2 | 3.0E-07 |
| DIP_10007801 | putative duf6 domain protein [Eutypa lata UCREL1] | 63.5 | 0.0 |
| DIP_10007802 | putative f-box domain-containing protein [Eutypa lata UCREL1] | 41.9 | 2.0E-90 |
| DIP_10007803 | putative saga complex subunit sgf29 protein [Eutypa lata UCREL1] | 61.8 | 3.0E-130 |
| DIP_10007804 | putative mannose-6-phosphate isomerase protein [Eutypa lata UCREL1] | 80.8 | 0.0 |
| DIP_10007805 | hypothetical protein SAPIO_CDS1601 [Scedosporium apiospermum] | 42.9 | 3.0E-77 |
| DIP_10007806 | putative cfem domain-containing protein [Eutypa lata UCREL1] | 38.6 | 1.0E-27 |
| DIP_10007807 | putative capsule polysaccharide biosynthesis protein [Eutypa lata UCREL1] | 61.2 | 3.0E-151 |
| DIP_10007808 | hypothetical protein UCREL1_8994 [Eutypa lata UCREL1] | 59.1 | 9.0E-19 |
| DIP_10007809 | putative dynactin subunit 5 protein [Togninia minima UCRPA7] | 72.5 | 1.0E-107 |
| DIP_10007810 | hypothetical protein PFICI_06860 [Pestalotiopsis fici W106-1] | 60.1 | 8.0E-156 |
| DIP_10007811 | hypothetical protein MGG_07245 [Magnaporthe oryzae 70-15] | 42.2 | 2.0E-11 |
| DIP_10007812 | hypothetical protein PFICI_03985 [Pestalotiopsis fici W106-1] | 44.5 | 4.0E-144 |
| DIP_10007813 | putative mitochondrial outer membrane protein [Eutypa lata UCREL1] | 37.8 | 2.0E-33 |
| DIP_10007814 | hypothetical protein PFICI_07188 [Pestalotiopsis fici W106-1] | 62.3 | 2.0E-112 |
| DIP_10007815 | hypothetical protein PFICI_07190 [Pestalotiopsis fici W106-1] | 46.7 | 2.0E-63 |
| DIP_10007816 | NA | | |
| DIP_10007817 | hypothetical protein SBOR_9046 [Sclerotinia borealis F-4157] | 44.2 | 5.0E-24 |
| DIP_10007818 | hypothetical protein HMPREF1541_06228 [Cyphellophora europaea CBS 101466] | 66.1 | 3.0E-131 |
| DIP_10007819 | hypothetical protein PFICI_13384 [Pestalotiopsis fici W106-1] | 48.1 | 4.0E-149 |
| DIP_10007820 | putative replication factor-a protein 1 protein [Eutypa lata UCREL1] | 72.3 | 0.0 |
| DIP_10007821 | hypothetical protein PFICI_13386 [Pestalotiopsis fici W106-1] | 68.4 | 6.0E-170 |
| DIP_10007822 | putative zinc-binding alcohol dehydrogenase domain-containing protein 2 protein [Eutypa lata UCREL1] | 56.9 | 4.0E-143 |
| DIP_10007823 | NA | | |
| DIP_10007824 | putative heterokaryon incompatibility protein [Eutypa lata UCREL1] | 41.9 | 4.0E-152 |
| DIP_10007825 | hypothetical protein PFICI_06300 [Pestalotiopsis fici W106-1] | 58.9 | 0.0 |
| DIP_10007826 | NA | | |
| DIP_10007827 | hypothetical protein HIM_09547 [Hirsutella minnesotensis 3608] | 54.1 | 0.0 |
| DIP_10007828 | NA | | |
| DIP_10007829 | hypothetical protein PFICI_13390 [Pestalotiopsis fici W106-1] | 66.3 | 0.0 |
| DIP_10007830 | hypothetical protein PFICI_06305 [Pestalotiopsis fici W106-1] | 61.9 | 0.0 |
| DIP_10007831 | hypothetical protein MYCTH_2300175 [Myceliophthora thermophila ATCC 42464] | 46.7 | 4.0E-37 |
| DIP_10007832 | peptidase-like protein [Myceliophthora thermophila ATCC 42464] | 56.7 | 0.0 |
| DIP_10007833 | NA | | |
| DIP_10007834 | hypothetical protein PFICI_13350 [Pestalotiopsis fici W106-1] | 73.2 | 0.0 |
| DIP_10007835 | putative isoflavone reductase family protein [Eutypa lata UCREL1] | 49.6 | 7.0E-93 |
| DIP_10007836 | hypothetical protein VDAG_08633 [Verticillium dahliae VdLs.17] | 73.2 | 4.0E-89 |
| DIP_10007837 | Cysteine desulfurase [Pestalotiopsis fici W106-1] | 79.2 | 0.0 |
| DIP_10007838 | putative bar domain-containing protein [Eutypa lata UCREL1] | 69.2 | 0.0 |
| DIP_10007839 | NA | | |
| DIP_10007840 | NA | | |
| DIP_10007841 | NA | | |
| DIP_10007842 | NA | | |
| DIP_10007843 | hypothetical protein PFICI_13392 [Pestalotiopsis fici W106-1] | 55.1 | 4.0E-93 |
| DIP_10007844 | putative dihydrofolate synthetase fol3 protein [Eutypa lata UCREL1] | 64.8 | 5.0E-162 |
| DIP_10007845 | NA | | |
| DIP_10007846 | NA | | |
| DIP_10007847 | hypothetical protein UCREL1_6744 [Eutypa lata UCREL1] | 49.5 | 5.0E-40 |
| DIP_10007848 | putative glutamine synthetase protein [Eutypa lata UCREL1] | 83.9 | 0.0 |
| DIP_10007849 | MOSC domain-containing protein [Colletotrichum graminicola M1.001] | 42.3 | 1.0E-171 |
| DIP_10007850 | NA | | |
| DIP_10007851 | NA | | |
| DIP_10007852 | putative meiotic recombination protein dmc1 protein [Eutypa lata UCREL1] | 60.1 | 5.0E-136 |
| DIP_10007853 | putative haloacid dehalogenase protein [Togninia minima UCRPA7] | 55.4 | 1.0E-103 |
| DIP_10007854 | predicted protein [Nectria haematococca mpVI 77-13-4] | 66.3 | 4.0E-39 |
| DIP_10007855 | NA | | |
| DIP_10007856 | hypothetical protein PFICI_06401 [Pestalotiopsis fici W106-1] | 61.1 | 5.0E-125 |
| DIP_10007857 | putative tat pathway signal sequence protein [Eutypa lata UCREL1] | 67.7 | 0.0 |
| DIP_10007858 | putative retinoblastoma-binding protein [Eutypa lata UCREL1] | 72.4 | 0.0 |
| DIP_10007859 | hypothetical protein FAVG1_00889 [Fusarium avenaceum] | 84.4 | 0.0 |
| DIP_10007860 | putative hydroxyisourate hydrolase protein [Eutypa lata UCREL1] | 80.6 | 2.0E-73 |
| DIP_10007861 | putative short chain dehydrogenase protein [Eutypa lata UCREL1] | 48.7 | 5.0E-78 |
| DIP_10007862 | hypothetical protein UCREL1_7948 [Eutypa lata UCREL1] | 55.6 | 0.0 |
| DIP_10007863 | NA | | |
| DIP_10007864 | NA | | |
| DIP_10007865 | hypothetical protein H634G_00133 [Metarhizium anisopliae BRIP 53293] | 87.8 | 0.0 |
| DIP_10007866 | putative nuclear pore complex subunit nup85 protein [Eutypa lata UCREL1] | 59.9 | 0.0 |
| DIP_10007867 | hypothetical protein UCREL1_6746 [Eutypa lata UCREL1] | 45.7 | 2.0E-100 |
| DIP_10007868 | allergen Asp F7 [Colletotrichum higginsianum] | 32.6 | 3.0E-33 |
| DIP_10007869 | hypothetical protein GGTG_00396 [Gaeumannomyces graminis var. tritici R3-111a-1] | 66.5 | 1.0E-108 |
| DIP_10007870 | calmodulin [Drechslerella stenobrocha 248] | 32.1 | 1.0E-11 |
| DIP_10007871 | putative transmembrane glyco protein [Eutypa lata UCREL1] | 37.9 | 1.0E-161 |
| DIP_10007872 | Protein phosphatase 2C [Pestalotiopsis fici W106-1] | 78.1 | 0.0 |
| DIP_10007873 | NA | | |
| DIP_10007874 | putative cytosolic non-specific dipeptidase protein [Eutypa lata UCREL1] | 83.2 | 0.0 |
| DIP_10007875 | hypothetical protein PFICI_04705 [Pestalotiopsis fici W106-1] | 60.1 | 1.0E-127 |
| DIP_10007876 | hypothetical protein GLRG_04721 [Colletotrichum graminicola M1.001] | 41.7 | 2.0E-21 |
| DIP_10007877 | hypothetical protein [Podospora anserina S mat+] | 47 | 1.0E-134 |
| DIP_10007878 | NA | | |
| DIP_10007879 | NA | | |
| DIP_10007880 | hypothetical protein PFICI_13082 [Pestalotiopsis fici W106-1] | 73.4 | 3.0E-59 |
| DIP_10007881 | putative dimethyladenosine transferase protein [Eutypa lata UCREL1] | 71.4 | 0.0 |
| DIP_10007882 | putative mrna splicing factor (prp1 zer1) protein [Eutypa lata UCREL1] | 83.4 | 0.0 |
| DIP_10007883 | hypothetical protein SPBR_04742 [Sporothrix brasiliensis 5110] | 36.2 | 2.0E-135 |
| DIP_10007884 | GTP cyclohydrolase II [Colletotrichum graminicola M1.001] | 71.9 | 0.0 |
| DIP_10007885 | NA | | |
| DIP_10007886 | hypothetical protein FVEG_11171 [Fusarium verticillioides 7600] | 38.3 | 1.0E-66 |
| DIP_10007887 | NA | | |
| DIP_10007888 | NADH dehydrogenase [Pestalotiopsis fici W106-1] | 72.9 | 7.0E-39 |
| DIP_10007889 | cytochrome P450 [Colletotrichum fioriniae PJ7] | 43.5 | 3.0E-159 |
| DIP_10007890 | NA | | |
| DIP_10007891 | hypothetical protein NEUTE2DRAFT_84984 [Neurospora tetrasperma FGSC 2509] | 42.5 | 1.0E-18 |
| DIP_10007892 | hypothetical protein TRIVIDRAFT_33427 [Trichoderma virens Gv29-8] | 51.3 | 1.0E-48 |
| DIP_10007893 | putative isoflavone reductase family protein [Eutypa lata UCREL1] | 51.7 | 2.0E-111 |
| DIP_10007894 | NA | | |
| DIP_10007895 | glutathionylspermidine synthase [Colletotrichum gloeosporioides Nara gc5] | 58.4 | 0.0 |
| DIP_10007896 | NA | | |
| DIP_10007897 | hypothetical protein CGGC5_13896 [Colletotrichum gloeosporioides Nara gc5] | 54 | 4.0E-36 |
| DIP_10007898 | hypothetical protein PFICI_06187 [Pestalotiopsis fici W106-1] | 75 | 0.0 |
| DIP_10007899 | steroid-binding protein-like protein [Acremonium chrysogenum ATCC 11550] | 74.6 | 9.0E-65 |
| DIP_10007900 | hypothetical protein PFICI_06311 [Pestalotiopsis fici W106-1] | 51.9 | 1.0E-178 |
| DIP_10007901 | NA | | |
| DIP_10007902 | NA | | |
| DIP_10007903 | hypothetical protein MYCTH_2300355 [Myceliophthora thermophila ATCC 42464] | 47.4 | 6.0E-95 |
| DIP_10007904 | putative cytochrome b5 [Diaporthe ampelina] | 72.6 | 7.0E-42 |
| DIP_10007905 | hypothetical protein CSUB01_03248 [Colletotrichum sublineola] | 58.3 | 2.0E-21 |
| DIP_10007906 | putative deoxycytidylate deaminase protein [Eutypa lata UCREL1] | 53.2 | 3.0E-142 |
| DIP_10007907 | Ubiquitin-like modifier-activating enzyme atg-7 [Pestalotiopsis fici W106-1] | 75.6 | 0.0 |
| DIP_10007908 | NA | | |
| DIP_10007909 | hypothetical protein PFICI_13627 [Pestalotiopsis fici W106-1] | 67.3 | 0.0 |
| DIP_10007910 | putative sugar transporter protein [Eutypa lata UCREL1] | 74.5 | 0.0 |
| DIP_10007911 | hypothetical protein PFICI_07093 [Pestalotiopsis fici W106-1] | 50.4 | 6.0E-127 |
| DIP_10007912 | hypothetical protein CSUB01_12395 [Colletotrichum sublineola] | 51.7 | 0.0 |
| DIP_10007913 | NA | | |
| DIP_10007914 | hypothetical protein MYCTH_2089701 [Myceliophthora thermophila ATCC 42464] | 33.9 | 1.0E-79 |
| DIP_10007915 | hypothetical protein FPSE_06597 [Fusarium pseudograminearum CS3096] | 47 | 2.0E-108 |
| DIP_10007916 | conserved hypothetical protein [Neurospora crassa] | 57.3 | 3.0E-125 |
| DIP_10007917 | hypothetical protein NECHADRAFT_102246 [Nectria haematococca mpVI 77-13-4] | 43.5 | 4.0E-101 |
| DIP_10007918 | putative gpi-anchored cell wall organization protein ecm33 protein [Eutypa lata UCREL1] | 50.1 | 3.0E-128 |
| DIP_10007919 | putative g-patch dna repair protein [Eutypa lata UCREL1] | 57.1 | 1.0E-151 |
| DIP_10007920 | NA | | |
| DIP_10007921 | putative dimeric alpha-beta barrel protein [Eutypa lata UCREL1] | 67.9 | 4.0E-52 |
| DIP_10007922 | catabolic 3-dehydroquinase [Colletotrichum higginsianum] | 68.3 | 1.0E-70 |
| DIP_10007923 | cytochrome p450 monooxygenase [Colletotrichum gloeosporioides Nara gc5] | 36.6 | 1.0E-119 |
| DIP_10007924 | hypothetical protein BBA_00045 [Beauveria bassiana ARSEF 2860] | 37.1 | 3.0E-29 |
| DIP_10007925 | hypothetical protein NEUTE1DRAFT_75018 [Neurospora tetrasperma FGSC 2508] | 78.5 | 0.0 |
| DIP_10007926 | putative ubiquinol-cytochrome c reductase complex 14 kda protein [Eutypa lata UCREL1] | 79 | 1.0E-70 |
| DIP_10007927 | NAD+ transporter [Verticillium alfalfae VaMs.102] | 54.1 | 6.0E-118 |
| DIP_10007928 | hypothetical protein TD95_001049 [Thielaviopsis punctulata] | 72.1 | 0.0 |
| DIP_10007929 | putative mep domain-containing protein [Eutypa lata UCREL1] | 70.8 | 3.0E-130 |
| DIP_10007930 | putative UBA/TS-N domain-containing protein [Colletotrichum sublineola] | 57.7 | 6.0E-105 |
| DIP_10007931 | hypothetical protein COCMIDRAFT_100113 [Bipolaris oryzae ATCC 44560] | 68.3 | 1.0E-37 |
| DIP_10007932 | hypothetical protein PFICI_06723 [Pestalotiopsis fici W106-1] | 69.9 | 0.0 |
| DIP_10007933 | glutathione S-transferase [Sphaerulina musiva SO2202] | 73.4 | 4.0E-134 |
| DIP_10007934 | hypothetical protein FOZG_17414 [Fusarium oxysporum Fo47] | 41.5 | 2.0E-32 |
| DIP_10007935 | hypothetical protein HIM_12655 [Hirsutella minnesotensis 3608] | 59 | 1.0E-92 |
| DIP_10007936 | NA | | |
| DIP_10007937 | hypothetical protein PFICI_06772 [Pestalotiopsis fici W106-1] | 47.1 | 4.0E-53 |
| DIP_10007938 | hypothetical protein Z517_07573 [Fonsecaea pedrosoi CBS 271.37] | 32.5 | 8.0E-28 |
| DIP_10007939 | hypothetical protein BBAD15_g3662 [Beauveria bassiana D1-5] | 30.6 | 9.0E-08 |
| DIP_10007940 | hypothetical protein PFICI_06774 [Pestalotiopsis fici W106-1] | 39.8 | 5.0E-96 |
| DIP_10007941 | hypothetical protein PFICI_06775 [Pestalotiopsis fici W106-1] | 80 | 0.0 |
| DIP_10007942 | hypothetical protein Z518_05715 [Rhinocladiella mackenziei CBS 650.93] | 30.2 | 1.0E-41 |
| DIP_10007943 | hypothetical protein VDAG_07984 [Verticillium dahliae VdLs.17] | 48.4 | 1.0E-115 |
| DIP_10007944 | hypothetical protein M7I_5696 [Glarea lozoyensis 74030] | 34.7 | 9.0E-32 |
| DIP_10007945 | hypothetical protein CGLO_08610 [Colletotrichum gloeosporioides Cg-14] | 37.1 | 3.0E-136 |
| DIP_10007946 | hypothetical protein PFICI_12802 [Pestalotiopsis fici W106-1] | 59 | 0.0 |
| DIP_10007947 | putative lipid phosphate phosphatase 2 protein [Eutypa lata UCREL1] | 64.5 | 3.0E-175 |
| DIP_10007948 | hypothetical protein UCREL1_8834 [Eutypa lata UCREL1] | 40.4 | 3.0E-34 |
| DIP_10007949 | hypothetical protein NECHADRAFT_18425 [Nectria haematococca mpVI 77-13-4] | 38.7 | 2.0E-16 |
| DIP_10007950 | hypothetical protein PFICI_07256 [Pestalotiopsis fici W106-1] | 70.5 | 2.0E-149 |
| DIP_10007951 | NADH-ubiquinone oxidoreductase subunit A [Pestalotiopsis fici W106-1] | 62.8 | 3.0E-85 |
| DIP_10007952 | putative duf1715 domain-containing protein [Eutypa lata UCREL1] | 50.5 | 1.0E-63 |
| DIP_10007953 | methionine--tRNA ligase [Pestalotiopsis fici W106-1] | 77.5 | 0.0 |
| DIP_10007954 | Cytochrome b-c1 complex subunit 8 [Pestalotiopsis fici W106-1] | 76.2 | 6.0E-57 |
| DIP_10007955 | SPFH domain/Band 7 family protein [Colletotrichum graminicola M1.001] | 87.8 | 7.0E-172 |
| DIP_10007956 | hypothetical protein PFICI_07242 [Pestalotiopsis fici W106-1] | 47.2 | 0.0 |
| DIP_10007957 | hypothetical protein UCREL1_7561 [Eutypa lata UCREL1] | 36.8 | 5.0E-17 |
| DIP_10007958 | hypothetical protein UCREL1_7560 [Eutypa lata UCREL1] | 49.3 | 3.0E-33 |
| DIP_10007959 | hypothetical protein TCE0_018r06137 [Talaromyces cellulolyticus] | 34.6 | 8.0E-10 |
| DIP_10007960 | hypothetical protein NECHADRAFT_61241 [Nectria haematococca mpVI 77-13-4] | 46.3 | 0.0 |
| DIP_10007961 | hypothetical protein GLRG_07496 [Colletotrichum graminicola M1.001] | 53.3 | 5.0E-174 |
| DIP_10007962 | hypothetical protein THITE_2106660 [Thielavia terrestris NRRL 8126] | 54.1 | 6.0E-133 |
| DIP_10007963 | putative an1-type zinc finger protein [Eutypa lata UCREL1] | 91.7 | 4.0E-44 |
| DIP_10007964 | NA | | |
| DIP_10007965 | similar to propionyl-CoA carboxylase [Leptosphaeria maculans JN3] | 51 | 0.0 |
| DIP_10007966 | putative carboxylase:pyruvate acetylpropionyl [Diaporthe ampelina] | 67.7 | 1.0E-134 |
| DIP_10007967 | hypothetical protein PFICI_07119 [Pestalotiopsis fici W106-1] | 66.6 | 0.0 |
| DIP_10007968 | integral membrane protein [Colletotrichum orbiculare MAFF 240422] | 38.5 | 3.0E-94 |
| DIP_10007969 | NA | | |
| DIP_10007970 | putative autophagy protein 5 protein [Eutypa lata UCREL1] | 87.1 | 5.0E-163 |
| DIP_10007971 | hypothetical protein FOPG_03233 [Fusarium oxysporum f. sp. conglutinans race 2 54008] | 38.2 | 7.0E-131 |
| DIP_10007972 | hypothetical protein NEUTE1DRAFT_77490 [Neurospora tetrasperma FGSC 2508] | 43.2 | 3.0E-25 |
| DIP_10007973 | hypothetical protein PFICI_07125 [Pestalotiopsis fici W106-1] | 61.3 | 0.0 |
| DIP_10007974 | NA | | |
| DIP_10007975 | putative acyl- dehydrogenase protein [Eutypa lata UCREL1] | 77.7 | 0.0 |
| DIP_10007976 | NA | | |
| DIP_10007977 | amine oxidase [Gaeumannomyces graminis var. tritici R3-111a-1] | 47.3 | 0.0 |
| DIP_10007978 | putative autophagy-like protein 2 protein [Eutypa lata UCREL1] | 64.1 | 1.0E-97 |
| DIP_10007979 | Atg2p [Colletotrichum higginsianum] | 76.7 | 0.0 |
| DIP_10007980 | hypothetical protein UCREL1_7940 [Eutypa lata UCREL1] | 55 | 4.0E-114 |
| DIP_10007981 | putative alpha beta-hydrolase protein [Eutypa lata UCREL1] | 63.4 | 1.0E-159 |
| DIP_10007982 | NA | | |
| DIP_10007983 | putative aspartyl-trna cytoplasmic protein [Eutypa lata UCREL1] | 72.1 | 0.0 |
| DIP_10007984 | NA | | |
| DIP_10007985 | unnamed protein product [Aspergillus oryzae RIB40] | 51.1 | 5.0E-41 |
| DIP_10007986 | hypothetical protein NECHADRAFT_1046 [Nectria haematococca mpVI 77-13-4] | 44.1 | 5.0E-85 |
| DIP_10007987 | hypothetical protein CGLO_11761 [Colletotrichum gloeosporioides Cg-14] | 70.1 | 0.0 |
| DIP_10007988 | NA | | |
| DIP_10007989 | hypothetical protein NECHADRAFT_36347 [Nectria haematococca mpVI 77-13-4] | 34.3 | 2.0E-70 |
| DIP_10007990 | hypothetical protein PFICI_10063 [Pestalotiopsis fici W106-1] | 49.3 | 2.0E-127 |
| DIP_10007991 | hypothetical protein PVAR5_3618 [Byssochlamys spectabilis No. 5] | 59.6 | 4.0E-35 |
| DIP_10007992 | hypothetical protein PVAR5_3618 [Byssochlamys spectabilis No. 5] | 50 | 4.0E-27 |
| DIP_10007993 | NA | | |
| DIP_10007994 | hypothetical protein PFICI_10609 [Pestalotiopsis fici W106-1] | 71 | 8.0E-142 |
| DIP_10007995 | hypothetical protein PFICI_09074 [Pestalotiopsis fici W106-1] | 42.7 | 2.0E-47 |
| DIP_10007996 | hypothetical protein S40288_06076 [Stachybotrys chartarum IBT 40288] | 73.6 | 1.0E-152 |
| DIP_10007997 | hypothetical protein PFICI_06457 [Pestalotiopsis fici W106-1] | 70 | 1.0E-174 |
| DIP_10007998 | hypothetical protein PFICI_06456 [Pestalotiopsis fici W106-1] | 70.1 | 0.0 |
| DIP_10007999 | hypothetical protein PFICI_06376 [Pestalotiopsis fici W106-1] | 54.3 | 1.0E-163 |
| DIP_10008000 | hypothetical protein PFICI_06137 [Pestalotiopsis fici W106-1] | 52.6 | 1.0E-88 |
| DIP_10008001 | Monothiol glutaredoxin-5 [Pestalotiopsis fici W106-1] | 70.1 | 3.0E-68 |
| DIP_10008002 | putative gamma-glutamyltranspeptidase protein [Eutypa lata UCREL1] | 59.6 | 0.0 |
| DIP_10008003 | hypothetical protein PFICI_06090 [Pestalotiopsis fici W106-1] | 35.7 | 2.0E-110 |
| DIP_10008004 | hypothetical protein CGLO_09640 [Colletotrichum gloeosporioides Cg-14] | 72.2 | 0.0 |
| DIP_10008005 | putative fimbrin protein [Togninia minima UCRPA7] | 88.4 | 0.0 |
| DIP_10008006 | hypothetical protein UCREL1_760 [Eutypa lata UCREL1] | 46.7 | 5.0E-52 |
| DIP_10008007 | NA | | |
| DIP_10008008 | hypothetical protein PFICI_07204 [Pestalotiopsis fici W106-1] | 80.9 | 3.0E-90 |
| DIP_10008009 | conidiation protein 6 [Colletotrichum graminicola M1.001] | 81.9 | 3.0E-42 |
| DIP_10008010 | putative c2 domain containing protein [Eutypa lata UCREL1] | 81.8 | 4.0E-97 |
| DIP_10008011 | 3'-5' exonuclease [Metarhizium anisopliae] | 37.8 | 2.0E-48 |
| DIP_10008012 | hypothetical protein PFICI_07257 [Pestalotiopsis fici W106-1] | 83 | 1.0E-134 |
| DIP_10008013 | NA | | |
| DIP_10008014 | putative initiation factor 2 subunit family protein [Eutypa lata UCREL1] | 59.7 | 3.0E-174 |
| DIP_10008015 | hypothetical protein PFICI_07260 [Pestalotiopsis fici W106-1] | 60.6 | 0.0 |
| DIP_10008016 | dienelactone hydrolase family protein [Neosartorya fischeri NRRL 181] | 54.5 | 1.0E-95 |
| DIP_10008017 | hypothetical protein PFICI_07261 [Pestalotiopsis fici W106-1] | 84.6 | 8.0E-90 |
| DIP_10008018 | putative h aca ribonucleoprotein complex subunit 2 protein [Eutypa lata UCREL1] | 72.7 | 2.0E-88 |
| DIP_10008019 | hypothetical protein PFICI_07270 [Pestalotiopsis fici W106-1] | 44.2 | 3.0E-165 |
| DIP_10008020 | hypothetical protein PFICI_07266 [Pestalotiopsis fici W106-1] | 51.9 | 0.0 |
| DIP_10008021 | NA | | |
| DIP_10008022 | 60S ribosomal protein L5 [Pestalotiopsis fici W106-1] | 83 | 0.0 |
| DIP_10008023 | Putative hydrolase [Podospora anserina S mat+] | 66.4 | 2.0E-136 |
| DIP_10008024 | putative rossmann-fold nad (+)-binding protein [Eutypa lata UCREL1] | 45 | 1.0E-167 |
| DIP_10008025 | putative ost3 ost6 family protein [Eutypa lata UCREL1] | 65.9 | 2.0E-164 |
| DIP_10008026 | putative exoglucanase 3 precursor protein [Eutypa lata UCREL1] | 77.1 | 0.0 |
| DIP_10008027 | hypothetical protein PFICI_12437 [Pestalotiopsis fici W106-1] | 37.8 | 4.0E-78 |
| DIP_10008028 | putative ribosomal rna methyltransferase mrm2 protein [Eutypa lata UCREL1] | 53.7 | 2.0E-62 |
| DIP_10008029 | putative npp1 domain protein [Diplodia seriata] | 63.5 | 5.0E-95 |
| DIP_10008030 | NA | | |
| DIP_10008031 | putative fad dependent oxidoreductase protein [Eutypa lata UCREL1] | 55.4 | 0.0 |
| DIP_10008032 | Argininosuccinate synthase [Ceratocystis platani] | 85.4 | 0.0 |
| DIP_10008033 | putative camk camkl ampk protein kinase protein [Eutypa lata UCREL1] | 68.2 | 0.0 |
| DIP_10008034 | hypothetical protein UCREL1_7575 [Eutypa lata UCREL1] | 76 | 7.0E-76 |
| DIP_10008035 | DNA polymerase epsilon subunit b [Colletotrichum orbiculare MAFF 240422] | 61.3 | 0.0 |
| DIP_10008036 | putative delta(24(24 ))-sterol reductase protein [Eutypa lata UCREL1] | 78.9 | 0.0 |
| DIP_10008037 | NA | | |
| DIP_10008038 | NA | | |
| DIP_10008039 | branched-chain amino acid aminotransferase [Colletotrichum orbiculare MAFF 240422] | 69.2 | 0.0 |
| DIP_10008040 | putative gtpase-activating protein gyp1 [Diaporthe ampelina] | 63.8 | 0.0 |
| DIP_10008041 | heterogeneous nuclear ribonucleoprotein HRP1 [Ophiocordyceps sinensis CO18] | 64.4 | 2.0E-155 |
| DIP_10008042 | putative phytanoyl- dioxygenase protein [Eutypa lata UCREL1] | 73.2 | 9.0E-172 |
| DIP_10008043 | hypothetical protein PFICI_13352 [Pestalotiopsis fici W106-1] | 71.7 | 0.0 |
| DIP_10008044 | putative camp-dependent protein kinase regulatory subunit protein [Eutypa lata UCREL1] | 79.3 | 0.0 |
| DIP_10008045 | putative bcas2 family protein [Eutypa lata UCREL1] | 51.6 | 2.0E-63 |
| DIP_10008046 | NA | | |
| DIP_10008047 | putative neutral ceramidase protein [Eutypa lata UCREL1] | 57.3 | 1.0E-176 |
| DIP_10008048 | putative glutaminase protein [Eutypa lata UCREL1] | 64.3 | 0.0 |
| DIP_10008049 | putative extracellular serine-threonine rich protein [Diaporthe ampelina] | 31.2 | 1.0E-11 |

| | | | |
|---|---|---|---|
| DIP_10008050 | predicted protein [Trichoderma reesei QM6a] | 61.8 | 7.0E-10 |
| DIP_10008051 | hypothetical protein HMPREF1541_01906 [Cyphellophora europaea CBS 101466] | 47.9 | 6.0E-163 |
| DIP_10008052 | hypothetical protein UCREL1_6288 [Eutypa lata UCREL1] | 50.2 | 3.0E-62 |
| DIP_10008053 | putative outer mitochondrial membrane protein porin protein [Eutypa lata UCREL1] | 91.2 | 0.0 |
| DIP_10008054 | putative asparagine synthetase protein [Eutypa lata UCREL1] | 85.1 | 0.0 |
| DIP_10008055 | putative sumo-conjugating enzyme ubc9 protein [Eutypa lata UCREL1] | 89.8 | 6.0E-107 |
| DIP_10008056 | putative pex19-domain-containing protein [Eutypa lata UCREL1] | 65.5 | 4.0E-107 |
| DIP_10008057 | hypothetical protein PFICI_06692 [Pestalotiopsis fici W106-1] | 50.4 | 2.0E-63 |
| DIP_10008058 | ubiquinone biosynthesis [Colletotrichum orbiculare MAFF 240422] | 73 | 0.0 |
| DIP_10008059 | hypothetical protein SAPIO_CDS5572 [Scedosporium apiospermum] | 44.7 | 2.0E-175 |
| DIP_10008060 | hypothetical protein PV08_02253 [Exophiala spinifera] | 63.5 | 1.0E-83 |
| DIP_10008061 | hypothetical protein PFICI_06696 [Pestalotiopsis fici W106-1] | 71 | 9.0E-123 |
| DIP_10008062 | NA | | |
| DIP_10008063 | putative fam86a protein [Eutypa lata UCREL1] | 61.7 | 8.0E-159 |
| DIP_10008064 | 40S ribosomal protein S24 [Neurospora crassa OR74A] | 88.2 | 8.0E-86 |
| DIP_10008065 | NA | | |
| DIP_10008066 | Form_Nir_trans-domain-containing protein [Sphaerulina musiva SO2202] | 66 | 3.0E-143 |
| DIP_10008067 | bicyclomycin resistance protein, putative [Byssochlamys spectabilis No. 5] | 63.3 | 0.0 |
| DIP_10008068 | putative methylase involved in ubiquinone menaquinone biosynthesis protein [Eutypa lata UCREL1] | 60.1 | 3.0E-116 |
| DIP_10008069 | hypothetical protein PFICI_06117 [Pestalotiopsis fici W106-1] | 69.2 | 0.0 |
| DIP_10008070 | putative serine-type carboxypeptidase protein [Eutypa lata UCREL1] | 68.6 | 0.0 |
| DIP_10008071 | hypothetical protein PFICI_06113 [Pestalotiopsis fici W106-1] | 66.8 | 5.0E-120 |
| DIP_10008072 | NA | | |
| DIP_10008073 | hypothetical protein PFICI_06638 [Pestalotiopsis fici W106-1] | 73 | 0.0 |
| DIP_10008074 | putative adaptor complexes medium subunit family protein [Eutypa lata UCREL1] | 81 | 0.0 |
| DIP_10008075 | hypothetical protein PFICI_06640 [Pestalotiopsis fici W106-1] | 62.2 | 0.0 |
| DIP_10008076 | putative longin-like protein [Eutypa lata UCREL1] | 52.2 | 6.0E-65 |
| DIP_10008077 | putative mitochondrial atpase protein [Eutypa lata UCREL1] | 52.9 | 0.0 |
| DIP_10008078 | putative nuclear pore complex subunit nup133 protein [Eutypa lata UCREL1] | 60.6 | 0.0 |
| DIP_10008079 | NA | | |
| DIP_10008080 | hypothetical protein THITE_2106340 [Thielavia terrestris NRRL 8126] | 72.7 | 3.0E-24 |
| DIP_10008081 | putative f-box domain-containing protein [Eutypa lata UCREL1] | 54.7 | 4.0E-71 |
| DIP_10008082 | hypothetical protein PFICI_06643 [Pestalotiopsis fici W106-1] | 69.6 | 0.0 |
| DIP_10008083 | putative exosome complex exonuclease protein [Eutypa lata UCREL1] | 73.6 | 3.0E-40 |
| DIP_10008084 | NA | | |
| DIP_10008085 | putative glutamine synthetase protein [Eutypa lata UCREL1] | 58.2 | 0.0 |
| DIP_10008086 | hypothetical protein SS1G_00871 [Sclerotinia sclerotiorum 1980] | 40.3 | 3.0E-60 |
| DIP_10008087 | putative rna recognition domain-containing protein [Eutypa lata UCREL1] | 65.4 | 9.0E-163 |
| DIP_10008088 | hypothetical protein CGLO_04298 [Colletotrichum gloeosporioides Cg-14] | 62.2 | 5.0E-106 |
| DIP_10008089 | hypothetical protein UCREL1_10147 [Eutypa lata UCREL1] | 52.4 | 3.0E-44 |
| DIP_10008090 | MSF1 protein [Verticillium dahliae VdLs.17] | 79.2 | 1.0E-114 |
| DIP_10008091 | short chain dehydrogenase reductase family [Colletotrichum gloeosporioides Nara gc5] | 71.3 | 3.0E-144 |
| DIP_10008092 | putative d-xylulose reductase a protein [Eutypa lata UCREL1] | 54.7 | 1.0E-78 |
| DIP_10008093 | cell division control protein [Verticillium dahliae VdLs.17] | 73.3 | 0.0 |
| DIP_10008094 | sodium/phosphate symporter, putative [Neosartorya fischeri NRRL 181] | 75.7 | 0.0 |
| DIP_10008095 | putative rodlet peptide protein [Eutypa lata UCREL1] | 63.4 | 4.0E-56 |
| DIP_10008096 | putative citrate synthase protein [Eutypa lata UCREL1] | 78.8 | 0.0 |
| DIP_10008097 | hypothetical protein UCREL1_6741 [Eutypa lata UCREL1] | 45.4 | 3.0E-179 |
| DIP_10008098 | hypothetical protein UCREL1_8718 [Eutypa lata UCREL1] | 48.3 | 1.0E-11 |
| DIP_10008099 | putative actin-like protein [Eutypa lata UCREL1] | 94.7 | 0.0 |
| DIP_10008100 | hypothetical protein CSUB01_09801 [Colletotrichum sublineola] | 31.7 | 6.0E-20 |
| DIP_10008101 | adenylosuccinate lyase [Sporothrix brasiliensis 5110] | 75.6 | 0.0 |
| DIP_10008102 | hypothetical protein THITE_2110720 [Thielavia terrestris NRRL 8126] | 63.6 | 0.0 |
| DIP_10008103 | hypothetical protein THITE_2061232 [Thielavia terrestris NRRL 8126] | 81 | 4.0E-92 |
| DIP_10008104 | putative yippee family protein [Eutypa lata UCREL1] | 37.9 | 5.0E-52 |
| DIP_10008105 | NA | | |
| DIP_10008106 | putative ornithine decarboxylase antizyme protein [Eutypa lata UCREL1] | 63.4 | 9.0E-83 |
| DIP_10008107 | Putative Imidazole glycerol phosphate synthase hisHF [Torrubiella hemipterigena] | 74.9 | 0.0 |
| DIP_10008108 | hypothetical protein PFICI_13102 [Pestalotiopsis fici W106-1] | 37.4 | 3.0E-35 |
| DIP_10008109 | camp-regulated phosphoprotein family protein [Colletotrichum orbiculare MAFF 240422] | 68.3 | 2.0E-65 |
| DIP_10008110 | hypothetical protein V502_03871 [Pseudogymnoascus pannorum VKM F-4520 (FW-2644)] | 69.5 | 0.0 |
| DIP_10008111 | hypothetical protein UCREL1_6716 [Eutypa lata UCREL1] | 43.1 | 1.0E-69 |
| DIP_10008112 | putative duf431 domain-containing protein [Eutypa lata UCREL1] | 73.1 | 1.0E-107 |
| DIP_10008113 | NA | | |
| DIP_10008114 | hypothetical protein PFICI_13039 [Pestalotiopsis fici W106-1] | 59.6 | 2.0E-130 |
| DIP_10008115 | hypothetical protein PFICI_13038 [Pestalotiopsis fici W106-1] | 61.8 | 0.0 |
| DIP_10008116 | putative zinc knuckle transcription factor protein [Eutypa lata UCREL1] | 54.8 | 1.0E-143 |
| DIP_10008117 | putative wd repeat-containing protein 2 protein [Eutypa lata UCREL1] | 70.5 | 0.0 |
| DIP_10008118 | putative glycosyltransferase family protein [Eutypa lata UCREL1] | 36.3 | 1.0E-31 |
| DIP_10008119 | putative mitochondrial ribosomal small subunit component protein [Eutypa lata UCREL1] | 61.5 | 4.0E-110 |
| DIP_10008120 | predicted protein [Trichoderma reesei QM6a] | 96.5 | 2.0E-31 |
| DIP_10008121 | hypothetical protein PFICI_06409 [Pestalotiopsis fici W106-1] | 71.3 | 2.0E-125 |
| DIP_10008122 | putative recombination hotspot-binding protein [Eutypa lata UCREL1] | 64.6 | 9.0E-108 |
| DIP_10008123 | Uncharacterized protein SAPIO_CDS6188 [Scedosporium apiospermum] | 44.1 | 4.0E-121 |
| DIP_10008124 | allantoate permease [Ophiostoma piceae UAMH 11346] | 76.9 | 0.0 |
| DIP_10008125 | putative transcriptional activator protein acu-15 protein [Togninia minima UCRPA7] | 71.8 | 0.0 |
| DIP_10008126 | SET domain-containing protein [Colletotrichum gloeosporioides Cg-14] | 48.2 | 2.0E-136 |
| DIP_10008127 | putative gtp-binding protein [Eutypa lata UCREL1] | 88.7 | 0.0 |
| DIP_10008128 | NA | | |
| DIP_10008129 | hypothetical protein PFICI_05487 [Pestalotiopsis fici W106-1] | 61.7 | 0.0 |
| DIP_10008130 | hypothetical protein PFICI_07294 [Pestalotiopsis fici W106-1] | 45.5 | 1.0E-96 |
| DIP_10008131 | putative vacuolar fusion protein ccz1 protein [Eutypa lata UCREL1] | 61.8 | 4.0E-168 |
| DIP_10008132 | putative methyltransferase domain-containing protein [Eutypa lata UCREL1] | 63.8 | 2.0E-165 |
| DIP_10008133 | putative rta1 domain protein [Eutypa lata UCREL1] | 47.6 | 7.0E-96 |
| DIP_10008134 | hypothetical protein PFICI_13117 [Pestalotiopsis fici W106-1] | 58.2 | 2.0E-134 |
| DIP_10008135 | hypothetical protein UCREL1_10667 [Eutypa lata UCREL1] | 41.3 | 6.0E-33 |
| DIP_10008136 | putative carboxypeptidase kex1 protein [Eutypa lata UCREL1] | 71.8 | 0.0 |
| DIP_10008137 | NA | | |
| DIP_10008138 | mitochondrial phosphate carrier protein [Aspergillus terreus NIH2624] | 83.1 | 0.0 |
| DIP_10008139 | putative calcium-related spray protein [Eutypa lata UCREL1] | 77.1 | 0.0 |
| DIP_10008140 | hypothetical protein NECHADRAFT_52132 [Nectria haematococca mpVI 77-13-4] | 51.4 | 8.0E-92 |
| DIP_10008141 | PDA-like cytochrome P450 monooxygenase, putative [Aspergillus fumigatus A1163] | 41.8 | 1.0E-155 |
| DIP_10008142 | putative mitogen-activated protein kinase maf1 protein [Eutypa lata UCREL1] | 70.5 | 4.0E-179 |
| DIP_10008143 | putative clathrin coat assembly protein ap19 protein [Eutypa lata UCREL1] | 95.7 | 1.0E-97 |
| DIP_10008144 | glutamate-cysteine ligase modifier subunit [Magnaporthe oryzae 70-15] | 38.7 | 6.0E-82 |
| DIP_10008145 | putative serine threonine-protein kinase bud32 protein [Eutypa lata UCREL1] | 55.3 | 4.0E-97 |
| DIP_10008146 | NA | | |
| DIP_10008147 | NA | | |
| DIP_10008148 | NA | | |
| DIP_10008149 | NA | | |
| DIP_10008150 | hypothetical protein MBR_01525, partial [Metarhizium brunneum ARSEF 3297] | 53.6 | 1.0E-84 |
| DIP_10008151 | hypothetical protein H634G_01149 [Metarhizium anisopliae BRIP 53293] | 67.2 | 7.0E-85 |
| DIP_10008152 | putative osmolarity two-component system protein sln1 protein [Togninia minima UCRPA7] | 50.6 | 0.0 |
| DIP_10008153 | aconitase a/isopropylmalate dehydratase [Grosmannia clavigera kw1407] | 75.6 | 0.0 |
| DIP_10008154 | hypothetical protein CIMG_09013 [Byssochlamys spectabilis No. 5] | 30 | 2.0E-75 |
| DIP_10008155 | putative mitochondrial precursor protein import receptor tom70 protein [Eutypa lata UCREL1] | 80.4 | 0.0 |
| DIP_10008156 | NA | | |
| DIP_10008157 | hypothetical protein CIMG_08725 [Coccidioides immitis RS] | 30.4 | 6.0E-12 |
| DIP_10008158 | putative 4-amino-4-deoxychorismate protein [Chaetomium thermophilum var. thermophilum DSM 1495] | 33.2 | 1.0E-54 |
| DIP_10008159 | hypothetical protein UCREL1_6761 [Eutypa lata UCREL1] | 59.6 | 0.0 |
| DIP_10008160 | hypothetical protein PFICI_06544 [Pestalotiopsis fici W106-1] | 34.4 | 6.0E-65 |
| DIP_10008161 | NA | | |
| DIP_10008162 | hypothetical protein GLRG_05865 [Colletotrichum graminicola M1.001] | 68.4 | 0.0 |
| DIP_10008163 | gluconolactonase [Gaeumannomyces graminis var. tritici R3-111a-1] | 65.5 | 6.0E-157 |
| DIP_10008164 | NA | | |
| DIP_10008165 | hypothetical protein PFICI_06543 [Pestalotiopsis fici W106-1] | 42.4 | 4.0E-35 |
| DIP_10008166 | hypothetical protein FGSG_13456 [Fusarium graminearum PH-1] | 53.5 | 6.0E-122 |
| DIP_10008167 | hypothetical protein PFICI_13096 [Pestalotiopsis fici W106-1] | 65.1 | 4.0E-126 |
| DIP_10008168 | hypothetical protein SNOG_14689 [Phaeosphaeria nodorum SN15] | 50 | 3.0E-17 |
| DIP_10008169 | hypothetical protein PFICI_06542 [Pestalotiopsis fici W106-1] | 61 | 1.0E-169 |
| DIP_10008170 | putative ifrd domain-containing protein [Eutypa lata UCREL1] | 49.8 | 9.0E-143 |
| DIP_10008171 | NA | | |
| DIP_10008172 | hypothetical protein V491_04246 [Pseudogymnoascus pannorum VKM F-3775] | 35.2 | 4.0E-12 |
| DIP_10008173 | NA | | |
| DIP_10008174 | NA | | |
| DIP_10008175 | hypothetical protein UCREL1_8722 [Eutypa lata UCREL1] | 54.3 | 3.0E-43 |
| DIP_10008176 | NA | | |
| DIP_10008177 | NA | | |
| DIP_10008178 | hypothetical protein CGLO_07713 [Colletotrichum gloeosporioides Cg-14] | 48 | 2.0E-81 |
| DIP_10008179 | NA | | |
| DIP_10008180 | Isocitrate dehydrogenase [Pestalotiopsis fici W106-1] | 85.2 | 0.0 |
| DIP_10008181 | hypothetical protein PFICI_13108 [Pestalotiopsis fici W106-1] | 52.5 | 3.0E-165 |
| DIP_10008182 | hypothetical protein PFICI_13110 [Pestalotiopsis fici W106-1] | 71.3 | 6.0E-60 |
| DIP_10008183 | Fe-S cluster assembly protein dre2 [Pestalotiopsis fici W106-1] | 70.6 | 2.0E-156 |
| DIP_10008184 | NA | | |
| DIP_10008185 | hypothetical protein H634G_06463 [Metarhizium anisopliae BRIP 53293] | 39.8 | 6.0E-64 |
| DIP_10008186 | hypothetical protein PFICI_13113 [Pestalotiopsis fici W106-1] | 58.8 | 0.0 |
| DIP_10008187 | NA | | |
| DIP_10008188 | Neutral trehalase [Pestalotiopsis fici W106-1] | 70.9 | 0.0 |
| DIP_10008189 | isocitrate lyase family protein [Colletotrichum graminicola M1.001] | 78.5 | 0.0 |
| DIP_10008190 | NA | | |
| DIP_10008191 | putative glucan endo- -beta-glucosidase protein [Eutypa lata UCREL1] | 58.3 | 0.0 |
| DIP_10008192 | putative glycosyltransferase family 2 protein [Eutypa lata UCREL1] | 62.1 | 0.0 |
| DIP_10008193 | putative ribosomal protein [Eutypa lata UCREL1] | 44.1 | 1.0E-44 |
| DIP_10008194 | hypothetical protein PFICI_06179 [Pestalotiopsis fici W106-1] | 63.6 | 5.0E-114 |
| DIP_10008195 | related to trichodiene oxygenase cytochrome P450 [Fusarium fujikuroi IMI 58289] | 46.6 | 0.0 |
| DIP_10008196 | hypothetical protein THITE_2120545 [Thielavia terrestris NRRL 8126] | 52.5 | 1.0E-140 |
| DIP_10008197 | hypothetical protein UCREL1_716 [Eutypa lata UCREL1] | 68 | 0.0 |
| DIP_10008198 | NA | | |
| DIP_10008199 | peptidyl-prolyl cis-trans isomerase [Pestalotiopsis fici W106-1] | 66.7 | 0.0 |
| DIP_10008200 | hypothetical protein FOTG_15295 [Fusarium oxysporum f. sp. vasinfectum 25433] | 42.2 | 2.0E-66 |
| DIP_10008201 | putative mfs sugar transporter protein [Eutypa lata UCREL1] | 73.5 | 0.0 |
| DIP_10008202 | putative low molecular weight phosphotyrosine protein phosphatase protein [Eutypa lata UCREL1] | 65.7 | 1.0E-75 |
| DIP_10008203 | putative map kinase protein [Eutypa lata UCREL1] | 71.4 | 0.0 |
| DIP_10008204 | NA | | |
| DIP_10008205 | IgE-binging protein [Colletotrichum higginsianum] | 73.2 | 4.0E-128 |
| DIP_10008206 | hypothetical protein PFICI_12002 [Pestalotiopsis fici W106-1] | 43 | 2.0E-145 |
| DIP_10008207 | NA | | |
| DIP_10008208 | putative aconitate hydratase protein [Eutypa lata UCREL1] | 83.1 | 0.0 |
| DIP_10008209 | hypothetical protein FOMG_19363 [Fusarium oxysporum f. sp. melonis 26406] | 41.6 | 9.0E-85 |
| DIP_10008210 | NA | | |
| DIP_10008211 | hypothetical protein W97_07788 [Coniosporium apollinis CBS 100218] | 57.2 | 9.0E-54 |
| DIP_10008212 | NA | | |
| DIP_10008213 | Chitin synthase 3 [Pestalotiopsis fici W106-1] | 79.3 | 0.0 |
| DIP_10008214 | hypothetical protein PFICI_07207 [Pestalotiopsis fici W106-1] | 47.7 | 0.0 |
| DIP_10008215 | NA | | |
| DIP_10008216 | NA | | |
| DIP_10008217 | hypothetical protein FAVG1_00863 [Fusarium avenaceum] | 55.6 | 7.0E-64 |
| DIP_10008218 | putative solute carrier family 25 member 42 protein [Eutypa lata UCREL1] | 86.5 | 0.0 |
| DIP_10008219 | Ribosomal proteins L15p and L18e [Glarea lozoyensis ATCC 20868] | 84.2 | 4.0E-110 |
| DIP_10008220 | putative duf159 domain protein [Eutypa lata UCREL1] | 59.8 | 1.0E-127 |
| DIP_10008221 | putative small nucleolar ribonucleoprotein complex subunit protein [Eutypa lata UCREL1] | 75.4 | 0.0 |
| DIP_10008222 | putative delta( )-delta( )-dienoyl- isomerase protein [Eutypa lata UCREL1] | 70.3 | 4.0E-155 |
| DIP_10008223 | uncharacterized protein CPUR_05874 [Claviceps purpurea 20.1] | 70.8 | 6.0E-30 |
| DIP_10008224 | NA | | |
| DIP_10008225 | putative voltage-gated k channel beta protein [Eutypa lata UCREL1] | 68.2 | 0.0 |
| DIP_10008226 | nodulation protein L [Ophiocordyceps sinensis CO18] | 59.1 | 3.0E-51 |
| DIP_10008227 | hypothetical protein PFICI_06026 [Pestalotiopsis fici W106-1] | 54 | 0.0 |
| DIP_10008228 | putative pentatricopeptide repeat protein [Eutypa lata UCREL1] | 31.1 | 5.0E-26 |
| DIP_10008229 | hypothetical protein PFICI_06024 [Pestalotiopsis fici W106-1] | 47 | 7.0E-57 |
| DIP_10008230 | hypothetical protein GLRG_03463, partial [Colletotrichum graminicola M1.001] | 55.3 | 4.0E-23 |
| DIP_10008231 | hypothetical protein PFICI_06023 [Pestalotiopsis fici W106-1] | 51.9 | 0.0 |
| DIP_10008232 | Eukaryotic translation initiation factor 3 subunit F [Pestalotiopsis fici W106-1] | 78.2 | 0.0 |
| DIP_10008233 | putative digestive-organ expansion factor protein [Eutypa lata UCREL1] | 70 | 0.0 |
| DIP_10008234 | mitochondrial 54S ribosomal protein YmL8 [Chaetomium globosum CBS 148.51] | 70.4 | 2.0E-86 |
| DIP_10008235 | hypothetical protein PFICI_06010 [Pestalotiopsis fici W106-1] | 40.4 | 8.0E-54 |
| DIP_10008236 | flo11-like protein [Ophiocordyceps sinensis CO18] | 57.7 | 6.0E-129 |
| DIP_10008237 | hypothetical protein PFICI_14109 [Pestalotiopsis fici W106-1] | 60.2 | 0.0 |
| DIP_10008238 | hypothetical protein PFICI_14108 [Pestalotiopsis fici W106-1] | 36.8 | 4.0E-101 |
| DIP_10008239 | NA | | |
| DIP_10008240 | putative glutamine synthetase protein [Eutypa lata UCREL1] | 52.3 | 0.0 |
| DIP_10008241 | hypothetical protein UCREL1_6267 [Eutypa lata UCREL1] | 37.8 | 3.0E-27 |
| DIP_10008242 | hypothetical protein PFICI_06057 [Pestalotiopsis fici W106-1] | 53.2 | 6.0E-111 |
| DIP_10008243 | NA | | |
| DIP_10008244 | putative glycosyltransferase family 31 protein [Eutypa lata UCREL1] | 61 | 0.0 |
| DIP_10008245 | putative histone h1-binding protein [Eutypa lata UCREL1] | 62.5 | 1.0E-171 |
| DIP_10008246 | putative lem3 cdc50 family protein [Eutypa lata UCREL1] | 66 | 0.0 |
| DIP_10008247 | hypothetical protein PFICI_12437 [Pestalotiopsis fici W106-1] | 32.1 | 1.0E-49 |
| DIP_10008248 | NA | | |
| DIP_10008249 | putative fad binding domain-containing protein [Eutypa lata UCREL1] | 47.3 | 0.0 |
| DIP_10008250 | putative cell division control protein 14 protein [Eutypa lata UCREL1] | 67.5 | 5.0E-148 |
| DIP_10008251 | putative homoserine kinase protein [Eutypa lata UCREL1] | 86.4 | 0.0 |
| DIP_10008252 | hypothetical protein EURHEDRAFT_412205 [Aspergillus ruber CBS 135680] | 35.7 | 2.0E-56 |
| DIP_10008253 | putative ran-binding protein [Eutypa lata UCREL1] | 55.3 | 0.0 |
| DIP_10008254 | putative duf1711 domain protein [Eutypa lata UCREL1] | 57.8 | 5.0E-80 |
| DIP_10008255 | NA | | |
| DIP_10008256 | putative indoleamine -dioxygenase family protein [Eutypa lata UCREL1] | 72.8 | 0.0 |
| DIP_10008257 | putative alpha beta hydrolase fold-3 domain containing protein [Diaporthe ampelina] | 57 | 1.0E-145 |
| DIP_10008258 | NA | | |
| DIP_10008259 | hypothetical protein PFICI_07181 [Pestalotiopsis fici W106-1] | 72.5 | 2.0E-48 |
| DIP_10008260 | hypothetical protein V491_05985 [Pseudogymnoascus pannorum VKM F-3775] | 76.9 | 2.0E-140 |
| DIP_10008261 | hypothetical protein PFICI_06384 [Pestalotiopsis fici W106-1] | 73 | 0.0 |
| DIP_10008262 | hypothetical protein FOPG_12819 [Fusarium oxysporum f. sp. conglutinans race 2 54008] | 48.1 | 1.0E-64 |
| DIP_10008263 | 60S ribosomal protein L22 [Pestalotiopsis fici W106-1] | 85.2 | 5.0E-74 |
| DIP_10008264 | putative fatty aldehyde dehydrogenase protein [Eutypa lata UCREL1] | 64.8 | 0.0 |
| DIP_10008265 | putative exosome complex exonuclease rrp4 protein [Eutypa lata UCREL1] | 69.3 | 1.0E-175 |
| DIP_10008266 | putative integral peroxisomal membrane peroxin protein [Eutypa lata UCREL1] | 70.4 | 0.0 |
| DIP_10008267 | NA | | |
| DIP_10008268 | putative geranylgeranyl pyrophosphate synthetase protein [Eutypa lata UCREL1] | 67.7 | 0.0 |
| DIP_10008269 | oxidoreductase FAD dependent [Aspergillus fumigatus var. RP-2014] | 44.4 | 1.0E-36 |
| DIP_10008270 | putative fad binding domain-containing protein [Eutypa lata UCREL1] | 66.6 | 1.0E-147 |
| DIP_10008271 | hypothetical protein CH063_00079 [Colletotrichum higginsianum] | 67.5 | 0.0 |
| DIP_10008272 | putative wd-repeat protein pop3 [Diaporthe ampelina] | 79.9 | 0.0 |
| DIP_10008273 | NA | | |
| DIP_10008274 | eukaryotic aspartyl protease [Colletotrichum fioriniae PJ7] | 59.1 | 0.0 |
| DIP_10008275 | hypothetical protein VDAG_00281 [Verticillium dahliae VdLs.17] | 54.4 | 0.0 |
| DIP_10008276 | hypothetical protein PFICI_06881 [Pestalotiopsis fici W106-1] | 50 | 4.0E-102 |
| DIP_10008277 | putative copii-coated vesicle protein [Eutypa lata UCREL1] | 74 | 0.0 |
| DIP_10008278 | hypothetical protein PFICI_06892 [Pestalotiopsis fici W106-1] | 57.6 | 0.0 |
| DIP_10008279 | putative mrc1-like domain-containing protein [Eutypa lata UCREL1] | 49.5 | 0.0 |
| DIP_10008280 | hypothetical protein S40288_08707 [Stachybotrys chartarum IBT 40288] | 48.2 | 1.0E-99 |
| DIP_10008281 | hypothetical protein S40288_08706 [Stachybotrys chartarum IBT 40288] | 50.7 | 3.0E-92 |
| DIP_10008282 | NA | | |
| DIP_10008283 | NA | | |
| DIP_10008284 | hypothetical protein PFICI_06059 [Pestalotiopsis fici W106-1] | 59.5 | 1.0E-156 |
| DIP_10008285 | putative non-catalytic module family expansin protein [Togninia minima UCRPA7] | 40.7 | 3.0E-20 |
| DIP_10008286 | putative vacuolar atp synthase 16 kda proteolipid subunit protein [Eutypa lata UCREL1] | 97.6 | 3.0E-109 |
| DIP_10008287 | L30e-like protein [Aureobasidium melanogenum CBS 110374] | 80.2 | 1.0E-71 |
| DIP_10008288 | predicted protein [Nectria haematococca mpVI 77-13-4] | 42.5 | 0.0 |
| DIP_10008289 | NA | | |
| DIP_10008290 | hypothetical protein S40285_06230 [Stachybotrys chlorohalonata IBT 40285] | 82.3 | 0.0 |
| DIP_10008291 | hypothetical protein UCREL1_7351 [Eutypa lata UCREL1] | 35.7 | 2.0E-25 |
| DIP_10008292 | putative protein kinase protein [Eutypa lata UCREL1] | 67.5 | 0.0 |
| DIP_10008293 | hypothetical protein PFICI_06050 [Pestalotiopsis fici W106-1] | 61.7 | 0.0 |
| DIP_10008294 | hypothetical protein PFICI_06048 [Pestalotiopsis fici W106-1] | 57.4 | 1.0E-170 |
| DIP_10008295 | hypothetical protein PFICI_09586 [Pestalotiopsis fici W106-1] | 42.4 | 2.0E-102 |
| DIP_10008296 | hypothetical protein MGCH7_ch7g154 [Magnaporthe oryzae 70-15] | 35.9 | 3.0E-34 |
| DIP_10008297 | multiple ankyrin repeats single kh domain protein, putative [Talaromyces stipitatus ATCC 10500] | 31.7 | 1.0E-24 |
| DIP_10008298 | integral membrane protein [Colletotrichum gloeosporioides Cg-14] | 50 | 1.0E-89 |
| DIP_10008299 | putative glycoside hydrolase family 31 protein [Eutypa lata UCREL1] | 70.8 | 0.0 |
| DIP_10008300 | hypothetical protein GLRG_03536 [Colletotrichum graminicola M1.001] | 58.9 | 0.0 |
| DIP_10008301 | high affinity glucose transporter [Neurospora crassa OR74A] | 74.7 | 0.0 |
| DIP_10008302 | hypothetical protein PFICI_08420 [Pestalotiopsis fici W106-1] | 31.2 | 2.0E-15 |
| DIP_10008303 | putative fact complex subunit spt-16 protein [Eutypa lata UCREL1] | 84.8 | 0.0 |
| DIP_10008304 | putative opy2 protein [Eutypa lata UCREL1] | 55.7 | 7.0E-113 |
| DIP_10008305 | putative chitin synthase 1 protein [Eutypa lata UCREL1] | 84.5 | 0.0 |
| DIP_10008306 | hypothetical protein S40285_01252 [Stachybotrys chlorohalonata IBT 40285] | 48.1 | 4.0E-90 |
| DIP_10008307 | hypothetical protein TRIVIDRAFT_150386 [Trichoderma virens Gv29-8] | 36.2 | 9.0E-152 |
| DIP_10008308 | hypothetical protein PFICI_07198 [Pestalotiopsis fici W106-1] | 60.8 | 6.0E-169 |
| DIP_10008309 | hypothetical protein PFICI_09527 [Pestalotiopsis fici W106-1] | 63.2 | 0.0 |
| DIP_10008310 | glutamate-5-semialdehyde dehydrogenase [Colletotrichum higginsianum] | 70.3 | 0.0 |
| DIP_10008311 | hypothetical protein UCREL1_217 [Eutypa lata UCREL1] | 66 | 0.0 |
| DIP_10008312 | TPR domain-containing protein [Magnaporthe oryzae 70-15] | 32.6 | 2.0E-76 |
| DIP_10008313 | hypothetical protein PFICI_06445 [Pestalotiopsis fici W106-1] | 42.3 | 3.0E-136 |
| DIP_10008314 | Inositol-3-phosphate synthase [Pestalotiopsis fici W106-1] | 82.5 | 0.0 |
| DIP_10008315 | hypothetical protein PFICI_07272 [Pestalotiopsis fici W106-1] | 51.6 | 2.0E-135 |
| DIP_10008316 | hydrolase C26A3.11 [Pestalotiopsis fici W106-1] | 60.8 | 1.0E-155 |
| DIP_10008317 | hypothetical protein PFICI_06629 [Pestalotiopsis fici W106-1] | 76.3 | 0.0 |
| DIP_10008318 | hypothetical protein EMCG_03187 [Emmonsia crescens UAMH 3008] | 31.3 | 3.0E-27 |
| DIP_10008319 | hypothetical protein BDFG_02891 [Blastomyces dermatitidis ATCC 26199] | 37.4 | 1.0E-16 |
| DIP_10008320 | hypothetical protein PFICI_07884 [Pestalotiopsis fici W106-1] | 70.4 | 0.0 |
| DIP_10008321 | Leukotriene A-4 hydrolase [Pestalotiopsis fici W106-1] | 73.8 | 0.0 |
| DIP_10008322 | hypothetical protein PFICI_06425 [Pestalotiopsis fici W106-1] | 47.6 | 7.0E-91 |
| DIP_10008323 | putative ring finger-like protein [Eutypa lata UCREL1] | 68.5 | 2.0E-127 |
| DIP_10008324 | hypothetical protein UCREL1_8969 [Eutypa lata UCREL1] | 81.9 | 2.0E-47 |
| DIP_10008325 | hypothetical protein S40285_04100 [Stachybotrys chlorohalonata IBT 40285] | 93.2 | 1.0E-154 |
| DIP_10008326 | putative proteasome subunit alpha type 6 protein [Eutypa lata UCREL1] | 46.2 | 1.0E-84 |
| DIP_10008327 | NA | | |
| DIP_10008328 | NA | | |
| DIP_10008329 | putative lysine-sensitive aspartokinase 3 protein [Eutypa lata UCREL1] | 73 | 3.0E-100 |
| DIP_10008330 | putative chromatin remodelling complex atpase chain isw1 protein [Eutypa lata UCREL1] | 85.6 | 0.0 |
| DIP_10008331 | uncharacterized protein FFUJ_07230 [Fusarium fujikuroi IMI 58289] | 48.9 | 9.0E-69 |
| DIP_10008332 | dipeptidyl-peptidase V [Gaeumannomyces graminis var. tritici R3-111a-1] | 61.6 | 0.0 |
| DIP_10008333 | NA | | |
| DIP_10008334 | putative aminopeptidase y protein [Eutypa lata UCREL1] | 64.8 | 0.0 |
| DIP_10008335 | NA | | |
| DIP_10008336 | hypothetical protein MPER_00053 [Moniliophthora perniciosa FA553] | 41.3 | 2.0E-16 |
| DIP_10008337 | hypothetical protein PTRG_02552 [Pyrenophora tritici-repentis Pt-1C-BFP] | 67.6 | 2.0E-31 |
| DIP_10008338 | Delta-aminolevulinic acid dehydratase [Pestalotiopsis fici W106-1] | 91.8 | 0.0 |
| DIP_10008339 | putative major facilitator superfamily transporter protein [Eutypa lata UCREL1] | 74.9 | 0.0 |
| DIP_10008340 | hypothetical protein Cob_10247 [Colletotrichum orbiculare MAFF 240422] | 68 | 7.0E-26 |
| DIP_10008341 | putative tyrosine phosphatase protein [Eutypa lata UCREL1] | 58.7 | 7.0E-113 |
| DIP_10008342 | hypothetical protein PFICI_06065 [Pestalotiopsis fici W106-1] | 55.8 | 0.0 |
| DIP_10008343 | NA | | |
| DIP_10008344 | NA | | |
| DIP_10008345 | NA | | |
| DIP_10008346 | NA | | |
| DIP_10008347 | NA | | |
| DIP_10008348 | NA | | |
| DIP_10008349 | hypothetical protein PFICI_06620 [Pestalotiopsis fici W106-1] | 46 | 0.0 |
| DIP_10008350 | ribosomal protein S18 [Colletotrichum higginsianum] | 36.9 | 2.0E-38 |
| DIP_10008351 | U3 small nucleolar ribonucleoprotein IMP3 [Pestalotiopsis fici W106-1] | 79.8 | 1.0E-106 |
| DIP_10008352 | putative dna excision repair protein [Eutypa lata UCREL1] | 62.3 | 0.0 |
| DIP_10008353 | hypothetical protein UCREL1_1360 [Eutypa lata UCREL1] | 55.7 | 5.0E-109 |
| DIP_10008354 | hypothetical protein PFICI_00147 [Pestalotiopsis fici W106-1] | 44.9 | 6.0E-25 |
| DIP_10008355 | NA | | |
| DIP_10008356 | putative glycoside hydrolase family 72 protein [Eutypa lata UCREL1] | 70.2 | 0.0 |
| DIP_10008357 | NA | | |
| DIP_10008358 | hypothetical protein F503_06844 [Ophiostoma piceae UAMH 11346] | 34.3 | 2.0E-28 |
| DIP_10008359 | sodium/calcium transporter [Grosmannia clavigera kw1407] | 46.1 | 7.0E-113 |
| DIP_10008360 | NA | | |
| DIP_10008361 | NA | | |
| DIP_10008362 | hypothetical protein TCE0_017f04024 [Talaromyces cellulolyticus] | 36.6 | 4.0E-172 |
| DIP_10008363 | hypothetical protein FPSE_04701 [Fusarium pseudograminearum CS3096] | 45.6 | 7.0E-132 |
| DIP_10008364 | predicted protein [Aspergillus terreus NIH2624] | 36.6 | 2.0E-108 |
| DIP_10008365 | hypothetical protein THAR02_11017 [Trichoderma harzianum] | 38 | 7.0E-63 |
| DIP_10008366 | hypothetical protein PFICI_06348 [Pestalotiopsis fici W106-1] | 70.2 | 1.0E-75 |
| DIP_10008367 | putative non-classical export protein [Eutypa lata UCREL1] | 60.3 | 9.0E-71 |
| DIP_10008368 | NA | | |
| DIP_10008369 | hypothetical protein NECHADRAFT_42279 [Nectria haematococca mpVI 77-13-4] | 44 | 6.0E-49 |
| DIP_10008370 | putative peptide-n4-(n-acetyl-beta-glucosaminyl)asparagine amidase a protein [Togninia minima UCRPA7] | 56 | 0.0 |
| DIP_10008371 | hypothetical protein FOYG_08284 [Fusarium oxysporum FOSC 3-a] | 57.3 | 1.0E-21 |
| DIP_10008372 | hypothetical protein PFICI_06634 [Pestalotiopsis fici W106-1] | 57.4 | 2.0E-91 |
| DIP_10008373 | NA | | |
| DIP_10008374 | hypothetical protein PFICI_07298 [Pestalotiopsis fici W106-1] | 58.5 | 0.0 |
| DIP_10008375 | monodehydroascorbate reductase [Verticillium alfalfae VaMs.102] | 70.6 | 0.0 |
| DIP_10008376 | PREDICTED: extensin-like [Myotis brandtii] | 32.8 | 2.0E-06 |
| DIP_10008377 | NA | | |
| DIP_10008378 | putative rna polymerase 2 transcription elongation factor protein [Eutypa lata UCREL1] | 61.3 | 0.0 |
| DIP_10008379 | putative ubiquinone biosynthesis protein coq7 protein [Togninia minima UCRPA7] | 88.2 | 1.0E-67 |
| DIP_10008380 | putative gnat family [Diaporthe ampelina] | 46.2 | 2.0E-67 |
| DIP_10008381 | NA | | |
| DIP_10008382 | NA | | |
| DIP_10008383 | GabA permease [Magnaporthe oryzae 70-15] | 75.4 | 0.0 |
| DIP_10008384 | NA | | |
| DIP_10008385 | NA | | |
| DIP_10008386 | hypothetical protein COCCADRAFT_22070 [Bipolaris zeicola 26-R-13] | 35.6 | 1.0E-13 |
| DIP_10008387 | hypothetical protein PFICI_06330 [Pestalotiopsis fici W106-1] | 53.4 | 0.0 |
| DIP_10008388 | NA | | |
| DIP_10008389 | MFS multidrug transporter [Colletotrichum orbiculare MAFF 240422] | 52 | 0.0 |
| DIP_10008390 | multicopper oxidase [Metarhizium robertsii ARSEF 23] | 45.1 | 0.0 |
| DIP_10008391 | putative oxidoreductase nad-binding rossmann fold protein [Eutypa lata UCREL1] | 60.6 | 4.0E-150 |
| DIP_10008392 | hypothetical protein PFICI_13380 [Pestalotiopsis fici W106-1] | 78.4 | 3.0E-87 |
| DIP_10008393 | DRAP deaminase [Magnaporthe oryzae 70-15] | 69.5 | 0.0 |
| DIP_10008394 | putative bzip transcription factor protein [Eutypa lata UCREL1] | 58 | 2.0E-90 |
| DIP_10008395 | hypothetical protein PFICI_06644 [Pestalotiopsis fici W106-1] | 42.1 | 2.0E-97 |
| DIP_10008396 | hypothetical protein PFICI_06645 [Pestalotiopsis fici W106-1] | 57.9 | 3.0E-117 |
| DIP_10008397 | putative het domain-containing protein [Eutypa lata UCREL1] | 45.6 | 2.0E-116 |
| DIP_10008398 | putative duf636 domain protein [Eutypa lata UCREL1] | 80 | 2.0E-80 |
| DIP_10008399 | hypothetical protein FOTG_03078 [Fusarium oxysporum f. sp. vasinfectum 25433] | 50 | 4.0E-45 |
| DIP_10008400 | putative multiprotein-bridging factor 1 protein [Eutypa lata UCREL1] | 58.4 | 1.0E-51 |
| DIP_10008401 | NA | | |
| DIP_10008402 | hypothetical protein PFICI_13377 [Pestalotiopsis fici W106-1] | 45.5 | 5.0E-135 |
| DIP_10008403 | NA | | |
| DIP_10008404 | hypothetical protein HIM_06798 [Hirsutella minnesotensis 3608] | 53.7 | 5.0E-108 |
| DIP_10008405 | mis12-mtw1 family protein [Grosmannia clavigera kw1407] | 45 | 8.0E-101 |
| DIP_10008406 | hypothetical protein PFICI_13365 [Pestalotiopsis fici W106-1] | 53.6 | 4.0E-68 |
| DIP_10008407 | putative endoplasmic reticulum vesicle protein 25 protein [Eutypa lata UCREL1] | 78.3 | 1.0E-129 |
| DIP_10008408 | U1 small nuclear ribonucleoprotein C [Beauveria bassiana D1-5] | 78.2 | 1.0E-51 |
| DIP_10008409 | NA | | |
| DIP_10008410 | hypothetical protein MGG_07429, partial [Magnaporthe oryzae 70-15] | 35.7 | 1.0E-81 |
| DIP_10008411 | NA | | |
| DIP_10008412 | hypothetical protein PFICI_13368 [Pestalotiopsis fici W106-1] | 45.9 | 0.0 |
| DIP_10008413 | NA | | |
| DIP_10008414 | putative translation elongation factor eef-1b gamma protein [Eutypa lata UCREL1] | 42 | 1.0E-43 |
| DIP_10008415 | NA | | |

| | | | |
|---|---|---|---|
| DIP_10008416 | hypothetical protein CSUB01_02794 [Colletotrichum sublineola] | 37.1 | 4.0E-71 |
| DIP_10008417 | hypothetical protein PFICI_13026 [Pestalotiopsis fici W106-1] | 47.5 | 1.0E-173 |
| DIP_10008418 | NA | | |
| DIP_10008419 | putative snf2 family domain-containing protein [Eutypa lata UCREL1] | 55.3 | 0.0 |
| DIP_10008420 | NA | | |
| DIP_10008421 | NA | | |
| DIP_10008422 | putative dihydroorotase [Diaporthe ampelina] | 62.7 | 6.0E-151 |
| DIP_10008423 | NA | | |
| DIP_10008424 | NA | | |
| DIP_10008425 | putative rn serotype protein [Eutypa lata UCREL1] | 64.9 | 0.0 |
| DIP_10008426 | 60S ribosomal protein L30 [Pestalotiopsis fici W106-1] | 92.5 | 3.0E-69 |
| DIP_10008427 | putative 60s ribosomal protein l10 protein [Eutypa lata UCREL1] | 91.7 | 8.0E-153 |
| DIP_10008428 | hypothetical protein UCREL1_5713 [Eutypa lata UCREL1] | 58.2 | 3.0E-123 |
| DIP_10008429 | hypothetical protein PFICI_13036 [Pestalotiopsis fici W106-1] | 54.6 | 0.0 |
| DIP_10008430 | hypothetical protein FOXB_09220 [Fusarium oxysporum Fo5176] | 80.5 | 5.0E-175 |
| DIP_10008431 | Regulator of nonsense transcripts 1 [Pestalotiopsis fici W106-1] | 82.7 | 0.0 |
| DIP_10008432 | SesA [Nectria haematococca] | 33.1 | 6.0E-09 |
| DIP_10008433 | hypothetical protein TCE0_022r06540 [Talaromyces cellulolyticus] | 31.3 | 8.0E-21 |
| DIP_10008434 | metal dependent amidohydrolase [Verticillium alfalfae VaMs.102] | 41.4 | 3.0E-75 |
| DIP_10008435 | hypothetical protein UCREL1_7342 [Eutypa lata UCREL1] | 58 | 9.0E-130 |
| DIP_10008436 | NA | | |
| DIP_10008437 | protein kinase domain-containing protein [Colletotrichum gloeosporioides Nara gc5] | 32.8 | 5.0E-31 |
| DIP_10008438 | hypothetical protein CH063_00363 [Colletotrichum higginsianum] | 31.2 | 1.0E-34 |
| DIP_10008439 | putative swr1-complex protein 4 protein [Eutypa lata UCREL1] | 57.3 | 0.0 |
| DIP_10008440 | hypothetical protein PFICI_06034 [Pestalotiopsis fici W106-1] | 74 | 6.0E-127 |
| DIP_10008441 | Casein kinase I [Pestalotiopsis fici W106-1] | 90.3 | 0.0 |
| DIP_10008442 | NA | | |
| DIP_10008443 | NA | | |
| DIP_10008444 | NA | | |
| DIP_10008445 | hypothetical protein PFICI_10747 [Pestalotiopsis fici W106-1] | 51.4 | 5.0E-125 |
| DIP_10008446 | hypothetical protein A1O5_13467 [Cladophialophora psammophila CBS 110553] | 62.8 | 4.0E-29 |
| DIP_10008447 | NA | | |
| DIP_10008448 | hypothetical protein CFIO01_08792 [Colletotrichum fioriniae PJ7] | 55 | 1.0E-96 |
| DIP_10008449 | NA | | |
| DIP_10008450 | NA | | |
| DIP_10008451 | hypothetical protein PFICI_06041 [Pestalotiopsis fici W106-1] | 57 | 6.0E-138 |
| DIP_10008452 | putative vitamin h transporter 1 biotin symporter vht1 protein [Eutypa lata UCREL1] | 72.1 | 0.0 |
| DIP_10008453 | NA | | |
| DIP_10008454 | NA | | |
| DIP_10008455 | hypothetical protein MGYG_07227 [Microsporum gypseum CBS 118893] | 68.5 | 5.0E-133 |
| DIP_10008456 | hypothetical protein PFICI_06850 [Pestalotiopsis fici W106-1] | 92.2 | 2.0E-113 |
| DIP_10008457 | NA | | |
| DIP_10008458 | putative ubiquitin c-terminal hydrolase protein [Eutypa lata UCREL1] | 77.7 | 0.0 |
| DIP_10008459 | hypothetical protein THITE_2038147, partial [Thielavia terrestris NRRL 8126] | 34.6 | 3.0E-34 |
| DIP_10008460 | zinc knuckle domain protein [Colletotrichum gloeosporioides Nara gc5] | 66.5 | 2.0E-76 |
| DIP_10008461 | putative ddhd domain-containing protein [Eutypa lata UCREL1] | 66 | 0.0 |
| DIP_10008462 | putative dna-directed rna polymerase iii subunit rpc1 protein [Eutypa lata UCREL1] | 75.5 | 0.0 |
| DIP_10008463 | NA | | |
| DIP_10008464 | hypothetical protein PFICI_07103 [Pestalotiopsis fici W106-1] | 74.3 | 4.0E-146 |
| DIP_10008465 | NA | | |
| DIP_10008466 | NA | | |
| DIP_10008467 | polyketide synthase [Daldinia eschscholtzii] | 71.3 | 0.0 |
| DIP_10008468 | putative iron transport multicopper oxidase fet3 protein [Eutypa lata UCREL1] | 55.3 | 0.0 |
| DIP_10008469 | integral membrane protein DUF92 [Metarhizium robertsii ARSEF 23] | 50.3 | 4.0E-100 |
| DIP_10008470 | hypothetical protein AOCH_005191 [Aspergillus ochraceoroseus] | 43.7 | 0.0 |
| DIP_10008471 | putative transcription elongation factor spt6 protein [Eutypa lata UCREL1] | 75.8 | 0.0 |
| DIP_10008472 | hypothetical protein PFICI_06334 [Pestalotiopsis fici W106-1] | 82 | 0.0 |
| DIP_10008473 | haloacid dehalogenase-like hydrolase [Colletotrichum higginsianum] | 66.4 | 2.0E-108 |
| DIP_10008474 | hypothetical protein PFICI_06331 [Pestalotiopsis fici W106-1] | 59.9 | 3.0E-161 |
| DIP_10008475 | pisatin demethylase [Verticillium dahliae VdLs.17] | 45.8 | 3.0E-170 |
| DIP_10008476 | putative integral membrane protein [Eutypa lata UCREL1] | 55.4 | 1.0E-163 |
| DIP_10008477 | putative GDSL-like Lipase/Acylhydrolase [Colletotrichum sublineola] | 64 | 2.0E-118 |
| DIP_10008478 | putative c2h2 transcription factor protein [Eutypa lata UCREL1] | 39.5 | 1.0E-76 |
| DIP_10008479 | putative atp-dependent rna helicase dhx8 protein [Eutypa lata UCREL1] | 86.5 | 0.0 |
| DIP_10008480 | hypothetical protein PFICI_06660 [Pestalotiopsis fici W106-1] | 69 | 0.0 |
| DIP_10008481 | hypothetical protein UCREL1_4922 [Eutypa lata UCREL1] | 44 | 9.0E-94 |
| DIP_10008482 | hypothetical protein, partial [Neisseria lactamica] | 39 | 4.0E-07 |
| DIP_10008483 | NA | | |
| DIP_10008484 | sexual development transcription factor NsdD [Verticillium alfalfae VaMs.102] | 59.2 | 3.0E-55 |
| DIP_10008485 | NA | | |
| DIP_10008486 | NA | | |
| DIP_10008487 | Glutamate--cysteine ligase [Pestalotiopsis fici W106-1] | 84.7 | 0.0 |
| DIP_10008488 | hypothetical protein PFICI_06866 [Pestalotiopsis fici W106-1] | 36.5 | 6.0E-104 |
| DIP_10008489 | putative oleate-induced peroxisomal protein [Eutypa lata UCREL1] | 86.5 | 4.0E-75 |
| DIP_10008490 | putative 1 domain-containing protein [Eutypa lata UCREL1] | 35 | 6.0E-80 |
| DIP_10008491 | putative NAD dependent epimerase/dehydratase [Colletotrichum sublineola] | 46 | 4.0E-83 |
| DIP_10008492 | hypothetical protein PFICI_07226 [Pestalotiopsis fici W106-1] | 65.6 | 0.0 |
| DIP_10008493 | NA | | |
| DIP_10008494 | NA | | |
| DIP_10008495 | putative spermidine synthase protein [Eutypa lata UCREL1] | 93.8 | 0.0 |
| DIP_10008496 | NA | | |
| DIP_10008497 | putative guanine nucleotide-binding protein alpha-2 subunit protein [Eutypa lata UCREL1] | 87.3 | 0.0 |
| DIP_10008498 | beta-glucosidase M [Pestalotiopsis fici W106-1] | 64 | 0.0 |
| DIP_10008499 | putative fungal specific transcription protein [Eutypa lata UCREL1] | 50 | 0.0 |
| DIP_10008500 | putative spx domain-containing protein [Eutypa lata UCREL1] | 54.8 | 1.0E-176 |
| DIP_10008501 | hypothetical protein PFICI_05038 [Pestalotiopsis fici W106-1] | 35.2 | 5.0E-48 |
| DIP_10008502 | hypothetical protein PFICI_02775 [Pestalotiopsis fici W106-1] | 55.5 | 0.0 |
| DIP_10008503 | hypothetical protein PFICI_07224 [Pestalotiopsis fici W106-1] | 55.8 | 0.0 |
| DIP_10008504 | NA | | |
| DIP_10008505 | putative carbohydrate binding family 6 protein [Eutypa lata UCREL1] | 50.4 | 5.0E-79 |
| DIP_10008506 | Cytochrome P450 [Macrophomina phaseolina MS6] | 57.7 | 2.0E-69 |
| DIP_10008507 | cytochrome P450 46A1 [Fomitiporia mediterranea MF3/22] | 53.3 | 3.0E-112 |
| DIP_10008508 | putative flavodoxin reductase family protein [Eutypa lata UCREL1] | 41.2 | 2.0E-155 |
| DIP_10008509 | hypothetical protein PFICI_07275 [Pestalotiopsis fici W106-1] | 54.1 | 2.0E-173 |
| DIP_10008510 | NA | | |
| DIP_10008511 | hypothetical protein UCREL1_1138 [Eutypa lata UCREL1] | 57.7 | 7.0E-16 |
| DIP_10008512 | hypothetical protein PFICI_07279 [Pestalotiopsis fici W106-1] | 31.7 | 3.0E-49 |
| DIP_10008513 | putative atp-dependent rna helicase dbp5 protein [Eutypa lata UCREL1] | 79 | 0.0 |
| DIP_10008514 | hypothetical protein PFICI_07277 [Pestalotiopsis fici W106-1] | 75.6 | 0.0 |
| DIP_10008515 | NA | | |
| DIP_10008516 | NA | | |
| DIP_10008517 | NA | | |
| DIP_10008518 | hypothetical protein PV05_05092 [Exophiala xenobiotica] | 32.2 | 3.0E-19 |
| DIP_10008519 | putative vacuolar-sorting protein snf7 protein [Togninia minima UCRPA7] | 82.3 | 8.0E-112 |
| DIP_10008520 | NA | | |
| DIP_10008521 | NA | | |
| DIP_10008522 | hypothetical protein A1O3_08737 [Capronia epimyces CBS 606.96] | 60 | 3.0E-117 |
| DIP_10008523 | hypothetical protein GLAREA_06763 [Glarea lozoyensis ATCC 20868] | 41 | 5.0E-87 |
| DIP_10008524 | putative eukaryotic translation initiation factor 3 subunit 2 protein [Eutypa lata UCREL1] | 89.6 | 0.0 |
| DIP_10008525 | putative c-1-tetrahydrofolate synthase protein [Eutypa lata UCREL1] | 85 | 0.0 |
| DIP_10008526 | NA | | |
| DIP_10008527 | cytochrome p450 3a21 [Colletotrichum orbiculare MAFF 240422] | 37.6 | 1.0E-140 |
| DIP_10008528 | predicted protein [Uncinocarpus reesii 1704] | 50 | 3.0E-108 |
| DIP_10008529 | hypothetical protein TRIATDRAFT_301412 [Trichoderma atroviride IMI 206040] | 30.1 | 2.0E-29 |
| DIP_10008530 | hypothetical protein PFICI_06592 [Pestalotiopsis fici W106-1] | 67.8 | 0.0 |
| DIP_10008531 | hypothetical protein UCREL1_10113 [Eutypa lata UCREL1] | 65.3 | 1.0E-165 |
| DIP_10008532 | NA | | |
| DIP_10008533 | NA | | |
| DIP_10008534 | hypothetical protein PFICI_06595 [Pestalotiopsis fici W106-1] | 61.1 | 0.0 |
| DIP_10008535 | hypothetical protein AOL_s00083g347 [Arthrobotrys oligospora ATCC 24927] | 97.2 | 5.0E-70 |
| DIP_10008536 | hypothetical protein PFICI_06598 [Pestalotiopsis fici W106-1] | 59.1 | 1.0E-139 |
| DIP_10008537 | NA | | |
| DIP_10008538 | hypothetical protein MYCTH_2307312 [Myceliophthora thermophila ATCC 42464] | 52.3 | 2.0E-75 |
| DIP_10008539 | NA | | |
| DIP_10008540 | putative outer membrane autotransporter protein [Eutypa lata UCREL1] | 65.7 | 0.0 |
| DIP_10008541 | NA | | |
| DIP_10008542 | putative hydrogen voltage-gated channel 1 protein [Eutypa lata UCREL1] | 60.6 | 1.0E-73 |
| DIP_10008543 | hypothetical protein MYCTH_2302739 [Myceliophthora thermophila ATCC 42464] | 42.8 | 3.0E-89 |
| DIP_10008544 | hypothetical protein PFICI_13147 [Pestalotiopsis fici W106-1] | 59.5 | 0.0 |
| DIP_10008545 | hypothetical protein PFICI_13137 [Pestalotiopsis fici W106-1] | 56.3 | 2.0E-157 |
| DIP_10008546 | hypothetical protein OIDMADRAFT_155773, partial [Oidiodendron maius Zn] | 35.9 | 3.0E-17 |
| DIP_10008547 | NA | | |
| DIP_10008548 | putative choline-phosphate cytidylyltransferase protein [Eutypa lata UCREL1] | 70.4 | 0.0 |
| DIP_10008549 | putative f-box domain-containing protein [Eutypa lata UCREL1] | 46.8 | 2.0E-85 |
| DIP_10008550 | NA | | |
| DIP_10008551 | putative dynein light chain protein [Eutypa lata UCREL1] | 80.8 | 1.0E-94 |
| DIP_10008552 | vacuolar transporter chaperone 1 [Verticillium alfalfae VaMs.102] | 86.7 | 5.0E-73 |
| DIP_10008553 | hypothetical protein UCREL1_10677 [Eutypa lata UCREL1] | 31.7 | 4.0E-17 |
| DIP_10008554 | hypothetical protein FOXB_09243 [Fusarium oxysporum Fo5176] | 43.6 | 1.0E-70 |
| DIP_10008555 | putative E3 ubiquitin-protein ligase hulA [Pestalotiopsis fici W106-1] | 83.4 | 0.0 |
| DIP_10008556 | putative polyadenylate-binding protein 2 protein [Eutypa lata UCREL1] | 77.9 | 3.0E-83 |
| DIP_10008557 | pentafunctional AROM polypeptide [Verticillium dahliae VdLs.17] | 65.5 | 0.0 |
| DIP_10008558 | putative fungal specific transcription factor protein [Eutypa lata UCREL1] | 71.6 | 2.0E-141 |
| DIP_10008559 | putative ferric reductase like transmembrane component protein [Eutypa lata UCREL1] | 45.2 | 4.0E-115 |
| DIP_10008560 | NA | | |
| DIP_10008561 | hypothetical protein [Podospora anserina S mat+] | 64.2 | 0.0 |
| DIP_10008562 | putative mitochondrial nadh-ubiquinone oxidoreductase 20 kd subunit protein [Eutypa lata UCREL1] | 74.7 | 6.0E-118 |
| DIP_10008563 | hypothetical protein PFICI_13141 [Pestalotiopsis fici W106-1] | 59.8 | 0.0 |
| DIP_10008564 | hypothetical protein PFICI_13142 [Pestalotiopsis fici W106-1] | 56.3 | 0.0 |
| DIP_10008565 | NA | | |
| DIP_10008566 | putative exopolygalacturonase protein [Eutypa lata UCREL1] | 67.2 | 0.0 |
| DIP_10008567 | hypothetical protein DOTSEDRAFT_88845 [Dothistroma septosporum NZE10] | 42.3 | 1.0E-88 |
| DIP_10008568 | major facilitator superfamily transporter [Colletotrichum gloeosporioides Cg-14] | 63.5 | 0.0 |
| DIP_10008569 | hypothetical protein PFICI_13005 [Pestalotiopsis fici W106-1] | 61.2 | 3.0E-127 |
| DIP_10008570 | Negative regulator of the PHO system [Pestalotiopsis fici W106-1] | 85.2 | 0.0 |
| DIP_10008571 | hypothetical protein PFICI_13008 [Pestalotiopsis fici W106-1] | 63.2 | 2.0E-167 |
| DIP_10008572 | putative o- protein [Eutypa lata UCREL1] | 52.6 | 2.0E-149 |
| DIP_10008573 | hypothetical protein SAPIO_CDS4477 [Scedosporium apiospermum] | 46.4 | 2.0E-67 |
| DIP_10008574 | hypothetical protein O988_05619 [Pseudogymnoascus pannorum VKM F-3808] | 49.1 | 1.0E-57 |
| DIP_10008575 | hypothetical protein SAPIO_CDS5889 [Scedosporium apiospermum] | 40.2 | 3.0E-110 |
| DIP_10008576 | hypothetical protein PV06_09747 [Exophiala oligosperma] | 30.6 | 5.0E-13 |
| DIP_10008577 | NA | | |
| DIP_10008578 | hypothetical protein CFIO01_11370 [Colletotrichum fioriniae PJ7] | 36.6 | 3.0E-109 |
| DIP_10008579 | NA | | |
| DIP_10008580 | putative aspartyl aminopeptidase protein [Eutypa lata UCREL1] | 84.9 | 0.0 |
| DIP_10008581 | putative mfs transporter of unknown specificity protein [Eutypa lata UCREL1] | 65.7 | 0.0 |
| DIP_10008582 | NA | | |
| DIP_10008583 | putative fluconazole resistance protein 1 protein [Eutypa lata UCREL1] | 72.1 | 0.0 |
| DIP_10008584 | hypothetical protein M437DRAFT_64485 [Aureobasidium melanogenum CBS 110374] | 37.7 | 3.0E-08 |
| DIP_10008585 | Exportin-T [Pestalotiopsis fici W106-1] | 66.9 | 0.0 |
| DIP_10008586 | GPI mannosyltransferase 4 [Pestalotiopsis fici W106-1] | 74.9 | 0.0 |
| DIP_10008587 | putative gpi ethanolamine phosphate transferase 2 protein [Eutypa lata UCREL1] | 60.1 | 0.0 |
| DIP_10008588 | NA | | |
| DIP_10008589 | putative nucleolar protein 4 protein [Eutypa lata UCREL1] | 57.5 | 0.0 |
| DIP_10008590 | hypothetical protein PFICI_15402 [Pestalotiopsis fici W106-1] | 74.5 | 0.0 |
| DIP_10008591 | NA | | |
| DIP_10008592 | putative rhomboid family membrane protein [Eutypa lata UCREL1] | 63.7 | 0.0 |
| DIP_10008593 | putative catabolic 3-dehydroquinase protein [Eutypa lata UCREL1] | 44.8 | 4.0E-86 |
| DIP_10008594 | putative heat repeat protein [Eutypa lata UCREL1] | 75.2 | 0.0 |
| DIP_10008595 | hypothetical protein PFICI_15396 [Pestalotiopsis fici W106-1] | 44.4 | 1.0E-88 |
| DIP_10008596 | putative mitochondrial cytochrome c oxidase assembly factor protein [Togninia minima UCRPA7] | 71.4 | 1.0E-31 |
| DIP_10008597 | metallophosphoesterase domain-containing protein 1 [Salpingoeca rosetta] | 31.4 | 4.0E-17 |
| DIP_10008598 | putative ribosome assembly protein rrb1 protein [Eutypa lata UCREL1] | 75.2 | 0.0 |
| DIP_10008599 | dihydroxy-acid dehydratase [Pestalotiopsis fici W106-1] | 75.8 | 0.0 |
| DIP_10008600 | NA | | |
| DIP_10008601 | NA | | |
| DIP_10008602 | hypothetical protein HIM_08526 [Hirsutella minnesotensis 3608] | 50.4 | 4.0E-83 |
| DIP_10008603 | NA | | |
| DIP_10008604 | putative glycoside hydrolase family 16 protein [Eutypa lata UCREL1] | 56.7 | 1.0E-170 |
| DIP_10008605 | putative carboxy- -muconate cyclase protein [Eutypa lata UCREL1] | 40.1 | 1.0E-92 |
| DIP_10008606 | hypothetical protein PFICI_12699 [Pestalotiopsis fici W106-1] | 35.9 | 8.0E-35 |
| DIP_10008607 | hypothetical protein COCHEDRAFT_1158996 [Bipolaris maydis C5] | 41.2 | 4.0E-40 |
| DIP_10008608 | putative cell cycle control protein [Eutypa lata UCREL1] | 72 | 0.0 |
| DIP_10008609 | NA | | |
| DIP_10008610 | hypothetical protein PFICI_12764 [Pestalotiopsis fici W106-1] | 48 | 0.0 |
| DIP_10008611 | hypothetical protein PFICI_12767 [Pestalotiopsis fici W106-1] | 76.8 | 0.0 |
| DIP_10008612 | putative -diketo-d-gluconic acid reductase a protein [Eutypa lata UCREL1] | 49.6 | 2.0E-115 |
| DIP_10008613 | NA | | |
| DIP_10008614 | hypothetical protein PFICI_07685 [Pestalotiopsis fici W106-1] | 61.4 | 0.0 |
| DIP_10008615 | hypothetical protein THAR02_04870 [Trichoderma harzianum] | 55.2 | 3.0E-139 |
| DIP_10008616 | hypothetical protein GGTG_01384 [Gaeumannomyces graminis var. tritici R3-111a-1] | 73.1 | 6.0E-32 |
| DIP_10008617 | hypothetical protein PFICI_12988 [Pestalotiopsis fici W106-1] | 56.8 | 6.0E-110 |
| DIP_10008618 | hypothetical protein PFICI_12994 [Pestalotiopsis fici W106-1] | 56.2 | 0.0 |
| DIP_10008619 | hypothetical protein OIDMADRAFT_147639, partial [Oidiodendron maius Zn] | 35.6 | 3.0E-10 |
| DIP_10008620 | NA | | |
| DIP_10008621 | hypothetical protein CSUB01_04119 [Colletotrichum sublineola] | 45.6 | 6.0E-66 |
| DIP_10008622 | hypothetical protein PFICI_12999 [Pestalotiopsis fici W106-1] | 54.8 | 3.0E-94 |
| DIP_10008623 | GTP-binding protein gtr2 [Pestalotiopsis fici W106-1] | 84.4 | 0.0 |
| DIP_10008624 | putative nol1 nop2 sun family protein [Eutypa lata UCREL1] | 61.2 | 0.0 |
| DIP_10008625 | hypothetical protein BofuT4_P125200.1 [Botrytis cinerea T4] | 32.4 | 1.0E-14 |
| DIP_10008626 | similar to Pc21g09640 [Aspergillus kawachii IFO 4308] | 31.2 | 7.0E-59 |
| DIP_10008627 | putative wsc domain protein [Eutypa lata UCREL1] | 54.8 | 3.0E-149 |
| DIP_10008628 | hypothetical protein CMQ_2398 [Grosmannia clavigera kw1407] | 51.8 | 4.0E-38 |
| DIP_10008629 | putative cell division cycle protein 123 protein [Eutypa lata UCREL1] | 66.9 | 1.0E-175 |
| DIP_10008630 | Pre-mRNA-splicing factor CWC21 [Pestalotiopsis fici W106-1] | 61.4 | 2.0E-72 |
| DIP_10008631 | hypothetical protein PFICI_13013 [Pestalotiopsis fici W106-1] | 56.8 | 1.0E-162 |
| DIP_10008632 | NA | | |
| DIP_10008633 | NA | | |
| DIP_10008634 | hypothetical protein PFICI_13012 [Pestalotiopsis fici W106-1] | 51.7 | 0.0 |
| DIP_10008635 | putative mind kinetochore complex component protein [Togninia minima UCRPA7] | 51.5 | 2.0E-45 |
| DIP_10008636 | putative peroxisomal adenine nucleotide transporter 1 protein [Eutypa lata UCREL1] | 58.2 | 3.0E-80 |
| DIP_10008637 | NA | | |
| DIP_10008638 | hypothetical protein SAPIO_CDS9291 [Scedosporium apiospermum] | 45.8 | 0.0 |
| DIP_10008639 | hypothetical protein FAVG1_02766 [Fusarium avenaceum] | 54.4 | 2.0E-107 |
| DIP_10008640 | NA | | |
| DIP_10008641 | hypothetical protein SAPIO_CDS9286 [Scedosporium apiospermum] | 53 | 0.0 |
| DIP_10008642 | hypothetical protein SNOG_14107 [Phaeosphaeria nodorum SN15] | 57.7 | 1.0E-106 |
| DIP_10008643 | NA | | |
| DIP_10008644 | Cytochrome P450 3A31 [Scedosporium apiospermum] | 58 | 0.0 |
| DIP_10008645 | hypothetical protein SAPIO_CDS9287 [Scedosporium apiospermum] | 51.7 | 1.0E-119 |
| DIP_10008646 | putative ntf2 and rrm domain-containing protein [Eutypa lata UCREL1] | 63.6 | 0.0 |
| DIP_10008647 | hypothetical protein PFICI_13004 [Pestalotiopsis fici W106-1] | 37.4 | 6.0E-122 |
| DIP_10008648 | NA | | |
| DIP_10008649 | putative poly polymerase pla1 protein [Eutypa lata UCREL1] | 68.3 | 0.0 |
| DIP_10008650 | carbohydrate esterase family 1 protein [Trichoderma virens Gv29-8] | 41.1 | 2.0E-72 |
| DIP_10008651 | hypothetical protein PV04_08862 [Capronia semiimmersa] | 35.2 | 9.0E-90 |
| DIP_10008652 | putative dienelactone hydrolase protein [Eutypa lata UCREL1] | 57.5 | 7.0E-99 |
| DIP_10008653 | hypothetical protein PFICI_02753 [Pestalotiopsis fici W106-1] | 66.4 | 0.0 |
| DIP_10008654 | putative guanine nucleotide-binding protein alpha subunit protein [Eutypa lata UCREL1] | 94.3 | 0.0 |
| DIP_10008655 | hypothetical protein PFICI_12986 [Pestalotiopsis fici W106-1] | 59.7 | 2.0E-103 |
| DIP_10008656 | putative nuclear pore complex protein [Eutypa lata UCREL1] | 36.9 | 8.0E-85 |
| DIP_10008657 | putative sas10 utp3 family protein [Togninia minima UCRPA7] | 57.9 | 0.0 |
| DIP_10008658 | putative 3-ketosphinganine reductase protein [Eutypa lata UCREL1] | 66.8 | 1.0E-179 |
| DIP_10008659 | aldo/keto reductase, putative [Talaromyces stipitatus ATCC 10500] | 44.1 | 5.0E-90 |
| DIP_10008660 | hypothetical protein THAR02_00084 [Trichoderma harzianum] | 67.4 | 0.0 |
| DIP_10008661 | NA | | |
| DIP_10008662 | hypothetical protein UCREL1_5205 [Eutypa lata UCREL1] | 63.7 | 2.0E-117 |
| DIP_10008663 | putative glucanase b protein [Eutypa lata UCREL1] | 67.1 | 6.0E-179 |
| DIP_10008664 | putative glycoside hydrolase family 16 protein [Eutypa lata UCREL1] | 60.8 | 4.0E-149 |
| DIP_10008665 | NA | | |
| DIP_10008666 | Transcription factor, fungi [Metarhizium album ARSEF 1941] | 69.6 | 0.0 |
| DIP_10008667 | NA | | |
| DIP_10008668 | hypothetical protein PFICI_05887 [Pestalotiopsis fici W106-1] | 73 | 0.0 |
| DIP_10008669 | putative glycine-rich cell wall structural protein 1 protein [Eutypa lata UCREL1] | 47.6 | 2.0E-56 |
| DIP_10008670 | hypothetical protein PFICI_05884 [Pestalotiopsis fici W106-1] | 47.4 | 3.0E-127 |
| DIP_10008671 | putative duf1740 domain containing protein [Eutypa lata UCREL1] | 50.2 | 0.0 |
| DIP_10008672 | hypothetical protein PFICI_05893 [Pestalotiopsis fici W106-1] | 71.7 | 0.0 |
| DIP_10008673 | hypothetical protein AOCH_007090 [Aspergillus ochraceoroseus] | 42.8 | 6.0E-101 |
| DIP_10008674 | hypothetical protein Z520_06644 [Fonsecaea multimorphosa CBS 102226] | 41.9 | 1.0E-65 |
| DIP_10008675 | NA | | |
| DIP_10008676 | putative gln pro-rich protein [Eutypa lata UCREL1] | 86.3 | 4.0E-128 |
| DIP_10008677 | hypothetical protein UCREL1_5741 [Eutypa lata UCREL1] | 50.9 | 7.0E-33 |
| DIP_10008678 | putative eukaryotic translation initiation factor 3 subunit 8 protein [Eutypa lata UCREL1] | 81.1 | 0.0 |
| DIP_10008679 | NA | | |
| DIP_10008680 | hypothetical protein V490_01099 [Pseudogymnoascus pannorum VKM F-3557] | 30.4 | 3.0E-15 |
| DIP_10008681 | hypothetical protein PFICI_04056 [Pestalotiopsis fici W106-1] | 60.8 | 2.0E-72 |
| DIP_10008682 | hypothetical protein [Podospora anserina S mat+] | 68.7 | 0.0 |
| DIP_10008683 | hypothetical protein PFICI_02742 [Pestalotiopsis fici W106-1] | 62.7 | 0.0 |
| DIP_10008684 | hypothetical protein PFICI_02741 [Pestalotiopsis fici W106-1] | 57.8 | 3.0E-74 |
| DIP_10008685 | NA | | |
| DIP_10008686 | putative family protein [Eutypa lata UCREL1] | 43.2 | 2.0E-72 |
| DIP_10008687 | NA | | |
| DIP_10008688 | putative zinc carboxypeptidase a protein [Eutypa lata UCREL1] | 57.4 | 5.0E-168 |
| DIP_10008689 | hypothetical protein ACRE_051800 [Acremonium chrysogenum ATCC 11550] | 57.5 | 1.0E-60 |
| DIP_10008690 | hypothetical protein T310_8293 [Rasamsonia emersonii CBS 393.64] | 33.8 | 3.0E-35 |
| DIP_10008691 | putative prephenate dehydrogenase protein [Eutypa lata UCREL1] | 69.3 | 0.0 |
| DIP_10008692 | hypothetical protein PFICI_02789 [Pestalotiopsis fici W106-1] | 50.4 | 8.0E-129 |
| DIP_10008693 | hypothetical protein PFICI_02788 [Pestalotiopsis fici W106-1] | 57.4 | 5.0E-173 |
| DIP_10008694 | hypothetical protein PFICI_02787 [Pestalotiopsis fici W106-1] | 39.9 | 6.0E-121 |
| DIP_10008695 | hypothetical protein PFICI_11511 [Pestalotiopsis fici W106-1] | 64.5 | 0.0 |
| DIP_10008696 | hypothetical protein V502_07561 [Pseudogymnoascus pannorum VKM F-4520 (FW-2644)] | 33.5 | 1.0E-37 |
| DIP_10008697 | hypothetical protein FOTG_15699 [Fusarium oxysporum f. sp. vasinfectum 25433] | 55.8 | 6.0E-170 |
| DIP_10008698 | hypothetical protein PFICI_13497 [Pestalotiopsis fici W106-1] | 39.2 | 3.0E-136 |
| DIP_10008699 | hypothetical protein FOMG_15775 [Fusarium oxysporum f. sp. melonis 26406] | 51.2 | 3.0E-120 |
| DIP_10008700 | hypothetical protein CFIO01_00636 [Colletotrichum fioriniae PJ7] | 53.6 | 6.0E-62 |
| DIP_10008701 | hypothetical protein PFICI_11762 [Pestalotiopsis fici W106-1] | 55 | 0.0 |
| DIP_10008702 | hypothetical protein PFICI_12992 [Pestalotiopsis fici W106-1] | 55 | 0.0 |
| DIP_10008703 | putative cop9 signalosome subunit 1 protein [Eutypa lata UCREL1] | 59.4 | 0.0 |
| DIP_10008704 | putative zn2+-binding protein melusin contains chord domain protein [Eutypa lata UCREL1] | 51 | 1.0E-120 |
| DIP_10008705 | conserved hypothetical protein [Aspergillus fumigatus Af293] | 30.2 | 5.0E-16 |
| DIP_10008706 | putative Xaa-Pro aminopeptidase [Pestalotiopsis fici W106-1] | 54.5 | 0.0 |
| DIP_10008707 | putative lectin family integral membrane protein [Eutypa lata UCREL1] | 49.7 | 3.0E-154 |
| DIP_10008708 | putative dolichyl pyrophosphate glc1man9 c2 alpha- -glucosyltransferase protein [Togninia minima UCRPA7] | 83.1 | 0.0 |
| DIP_10008709 | hypothetical protein VHEMI02858 [Torrubiella hemipterigena] | 48.7 | 0.0 |
| DIP_10008710 | hypothetical protein UCREL1_7186 [Eutypa lata UCREL1] | 67.4 | 2.0E-46 |
| DIP_10008711 | oligopeptide transporter [Verticillium alfalfae VaMs.102] | 52.6 | 0.0 |
| DIP_10008712 | hypothetical protein PFICI_12684 [Pestalotiopsis fici W106-1] | 51.9 | 1.0E-137 |
| DIP_10008713 | hypothetical protein Z517_00309 [Fonsecaea pedrosoi CBS 271.37] | 32.9 | 6.0E-28 |
| DIP_10008714 | 37S ribosomal protein MRP17, mitochondrial [Fusarium oxysporum f. sp. cubense race 4] | 53.9 | 3.0E-34 |
| DIP_10008715 | hypothetical protein UCREL1_1590 [Eutypa lata UCREL1] | 54 | 0.0 |
| DIP_10008716 | Cytochrome c oxidase subunit 4 [Pestalotiopsis fici W106-1] | 72.2 | 2.0E-92 |
| DIP_10008717 | NA | | |
| DIP_10008718 | hypothetical protein HMPREF1624_05578 [Sporothrix schenckii ATCC 58251] | 46.8 | 1.0E-76 |
| DIP_10008719 | hypothetical protein M419DRAFT_136368 [Trichoderma reesei RUT C-30] | 56.2 | 0.0 |
| DIP_10008720 | putative mfs drug transporter protein [Eutypa lata UCREL1] | 72.2 | 0.0 |
| DIP_10008721 | putative cysteine synthase protein [Eutypa lata UCREL1] | 72.4 | 0.0 |
| DIP_10008722 | hypothetical protein UCREL1_5669 [Eutypa lata UCREL1] | 54.9 | 0.0 |
| DIP_10008723 | hypothetical protein GMDG_02458 [Pseudogymnoascus destructans 20631-21] | 81.8 | 1.0E-48 |
| DIP_10008724 | NA | | |
| DIP_10008725 | hypothetical protein UCREL1_6685 [Eutypa lata UCREL1] | 36.9 | 2.0E-50 |
| DIP_10008726 | putative n-acetylglucosaminyl-phosphatidylinositol de-n-acetylase protein [Eutypa lata UCREL1] | 50.7 | 2.0E-89 |
| DIP_10008727 | hypothetical protein H634G_00418 [Metarhizium anisopliae BRIP 53293] | 48.1 | 2.0E-75 |
| DIP_10008728 | hypothetical protein FAVG1_09950 [Fusarium avenaceum] | 58.6 | 8.0E-94 |
| DIP_10008729 | putative glycoside hydrolase family 18 protein [Eutypa lata UCREL1] | 54.8 | 1.0E-134 |
| DIP_10008730 | putative polysaccharide biosynthesis protein vipa tvib protein [Eutypa lata UCREL1] | 44.4 | 2.0E-132 |
| DIP_10008731 | putative glycosyltransferase family 2 protein [Eutypa lata UCREL1] | 61.3 | 0.0 |
| DIP_10008732 | hypothetical protein MGG_14966 [Magnaporthe oryzae 70-15] | 35.9 | 6.0E-46 |
| DIP_10008733 | putative mannan polymerase ii complex anp1 subunit protein [Eutypa lata UCREL1] | 36.2 | 1.0E-66 |
| DIP_10008734 | putative mfs monosaccharide protein [Eutypa lata UCREL1] | 69.4 | 0.0 |
| DIP_10008735 | NA | | |
| DIP_10008736 | predicted protein [Nectria haematococca mpVI 77-13-4] | 37.4 | 3.0E-21 |
| DIP_10008737 | putative mitochondrial inheritance component mdm10 protein [Eutypa lata UCREL1] | 66.6 | 0.0 |
| DIP_10008738 | putative plasma membrane low affinity zinc ion protein [Eutypa lata UCREL1] | 69.3 | 9.0E-179 |
| DIP_10008739 | 40S ribosomal protein S19 [Ceratocystis platani] | 81.2 | 2.0E-75 |
| DIP_10008740 | Mitochondrial inner membrane organizing system protein [Hirsutella minnesotensis 3608] | 86.8 | 3.0E-49 |
| DIP_10008741 | hypothetical protein PFICI_12461 [Pestalotiopsis fici W106-1] | 32.5 | 3.0E-81 |
| DIP_10008742 | putative urb2 npa2 family protein [Eutypa lata UCREL1] | 46.2 | 7.0E-136 |
| DIP_10008743 | putative fad binding domain protein [Eutypa lata UCREL1] | 63.4 | 0.0 |
| DIP_10008744 | Short-chain dehydrogenase/reductase SDR [Penicillium expansum] | 35.2 | 7.0E-75 |
| DIP_10008745 | NA | | |
| DIP_10008746 | hypothetical protein PFICI_12457 [Pestalotiopsis fici W106-1] | 59.2 | 0.0 |
| DIP_10008747 | NA | | |
| DIP_10008748 | NA | | |
| DIP_10008749 | hypothetical protein PFICI_12456 [Pestalotiopsis fici W106-1] | 51 | 0.0 |
| DIP_10008750 | hypothetical protein ACRE_074940 [Acremonium chrysogenum ATCC 11550] | 35.3 | 4.0E-21 |
| DIP_10008751 | NA | | |
| DIP_10008752 | hypothetical protein CH063_06670 [Colletotrichum higginsianum] | 60.8 | 2.0E-101 |
| DIP_10008753 | hypothetical protein FAVG1_05597 [Fusarium avenaceum] | 41.6 | 8.0E-146 |
| DIP_10008754 | putative heat repeat protein [Eutypa lata UCREL1] | 43.5 | 1.0E-112 |
| DIP_10008755 | NA | | |
| DIP_10008756 | NA | | |
| DIP_10008757 | hypothetical protein PFICI_03775 [Pestalotiopsis fici W106-1] | 51.2 | 0.0 |
| DIP_10008758 | putative transcription factor protein [Eutypa lata UCREL1] | 66.7 | 0.0 |
| DIP_10008759 | transcription factor [Colletotrichum gloeosporioides Nara gc5] | 56.5 | 2.0E-101 |
| DIP_10008760 | NA | | |
| DIP_10008761 | hypothetical protein PFICI_12749 [Pestalotiopsis fici W106-1] | 83.4 | 0.0 |
| DIP_10008762 | NA | | |
| DIP_10008763 | hypothetical protein PFICI_05547 [Pestalotiopsis fici W106-1] | 58.5 | 0.0 |
| DIP_10008764 | hypothetical protein PFICI_12651 [Pestalotiopsis fici W106-1] | 49.6 | 6.0E-116 |
| DIP_10008765 | NA | | |
| DIP_10008766 | putative sir2 family histone protein [Eutypa lata UCREL1] | 71.4 | 3.0E-145 |
| DIP_10008767 | putative histidine acid protein [Eutypa lata UCREL1] | 70.7 | 0.0 |
| DIP_10008768 | NA | | |
| DIP_10008769 | hypothetical protein PFICI_05286 [Pestalotiopsis fici W106-1] | 54.2 | 0.0 |
| DIP_10008770 | hypothetical protein PFICI_05287 [Pestalotiopsis fici W106-1] | 42.9 | 0.0 |
| DIP_10008771 | hypothetical protein UCREL1_6216 [Eutypa lata UCREL1] | 51.2 | 4.0E-123 |
| DIP_10008772 | putative brct domain-containing protein [Eutypa lata UCREL1] | 43.5 | 9.0E-156 |
| DIP_10008773 | NA | | |
| DIP_10008774 | putative zinc-binding dehydrogenase protein [Eutypa lata UCREL1] | 49.7 | 7.0E-116 |
| DIP_10008775 | putative sugar transporter stl1 protein [Eutypa lata UCREL1] | 79.7 | 0.0 |
| DIP_10008776 | NA | | |
| DIP_10008777 | membrane transporter [Colletotrichum orbiculare MAFF 240422] | 74.8 | 0.0 |
| DIP_10008778 | hypothetical protein PFICI_03344 [Pestalotiopsis fici W106-1] | 47.8 | 2.0E-152 |
| DIP_10008779 | putative catalase 1 protein [Eutypa lata UCREL1] | 74.9 | 0.0 |
| DIP_10008780 | hypothetical protein A1O1_06689 [Capronia coronata CBS 617.96] | 48.6 | 6.0E-83 |
| DIP_10008781 | hypothetical protein PFICI_03342 [Pestalotiopsis fici W106-1] | 57.6 | 0.0 |

| | | | |
|---|---|---|---|
| DIP_10008782 | NA | | |
| DIP_10008783 | ferric reductase transmembrane component [Neurospora crassa OR74A] | 43.4 | 0.0 |
| DIP_10008784 | putative riboflavin kinase protein [Eutypa lata UCREL1] | 47.8 | 5.0E-176 |
| DIP_10008785 | NA | | |
| DIP_10008786 | putative kinase activator protein [Eutypa lata UCREL1] | 68.8 | 0.0 |
| DIP_10008787 | NA | | |
| DIP_10008788 | putative integral membrane mpv17 pmp22 protein [Eutypa lata UCREL1] | 63.3 | 6.0E-91 |
| DIP_10008789 | putative secreted protein [Eutypa lata UCREL1] | 30.5 | 3.0E-13 |
| DIP_10008790 | hypothetical protein BAUCODRAFT_122802 [Baudoinia compniacensis UAMH 10762] | 42.6 | 1.0E-145 |
| DIP_10008791 | phthalate transporter [Colletotrichum orbiculare MAFF 240422] | 63.1 | 0.0 |
| DIP_10008792 | hypothetical protein PFICI_13780 [Pestalotiopsis fici W106-1] | 42.2 | 6.0E-56 |
| DIP_10008793 | hypothetical protein PFICI_03134 [Pestalotiopsis fici W106-1] | 65.1 | 0.0 |
| DIP_10008794 | putative integral membrane protein [Eutypa lata UCREL1] | 53.6 | 5.0E-44 |
| DIP_10008795 | NA | | |
| DIP_10008796 | zinc-binding oxidoreductase ToxD, putative [Aspergillus clavatus NRRL 1] | 39.4 | 2.0E-71 |
| DIP_10008797 | hypothetical protein PFICI_03135 [Pestalotiopsis fici W106-1] | 52.1 | 3.0E-27 |
| DIP_10008798 | putative gpi anchored protein [Eutypa lata UCREL1] | 71.6 | 0.0 |
| DIP_10008799 | putative mitochondrial import receptor subunit tom-22 protein [Eutypa lata UCREL1] | 69.6 | 4.0E-73 |
| DIP_10008800 | putative na(+) h(+) antiporter 2 protein [Eutypa lata UCREL1] | 62.2 | 0.0 |
| DIP_10008801 | hypothetical protein FPSE_04503 [Fusarium pseudograminearum CS3096] | 49.2 | 6.0E-111 |
| DIP_10008802 | scytalone dehydratase, partial [Daldinia eschscholtzii] | 78.5 | 5.0E-104 |
| DIP_10008803 | NA | | |
| DIP_10008804 | putative autophagy protein 16 protein [Eutypa lata UCREL1] | 52.2 | 2.0E-59 |
| DIP_10008805 | putative dot1 protein [Eutypa lata UCREL1] | 61.5 | 0.0 |
| DIP_10008806 | NA | | |
| DIP_10008807 | hypothetical protein FOXB_14067 [Fusarium oxysporum Fo5176] | 38.5 | 6.0E-33 |
| DIP_10008808 | hypothetical protein PFICI_08107 [Pestalotiopsis fici W106-1] | 60.6 | 0.0 |
| DIP_10008809 | putative transport protein particle subunit bet5 protein [Eutypa lata UCREL1] | 62 | 2.0E-87 |
| DIP_10008810 | hypothetical protein PFICI_08104 [Pestalotiopsis fici W106-1] | 59.4 | 0.0 |
| DIP_10008811 | putative duf1649 domain protein [Eutypa lata UCREL1] | 66 | 8.0E-93 |
| DIP_10008812 | putative forkhead box protein l2 protein [Eutypa lata UCREL1] | 67.6 | 0.0 |
| DIP_10008813 | NA | | |
| DIP_10008814 | hypothetical protein THITE_2118793 [Thielavia terrestris NRRL 8126] | 67.8 | 0.0 |
| DIP_10008815 | NA | | |
| DIP_10008816 | NA | | |
| DIP_10008817 | NA | | |
| DIP_10008818 | hypothetical protein PFICI_07472 [Pestalotiopsis fici W106-1] | 58.9 | 1.0E-164 |
| DIP_10008819 | NA | | |
| DIP_10008820 | putative dna-directed rna polymerase i and iii 14 kda polypeptide protein [Eutypa lata UCREL1] | 66.3 | 3.0E-64 |
| DIP_10008821 | NADH-ubiquinone oxidoreductase subunit A [Pestalotiopsis fici W106-1] | 83.9 | 8.0E-75 |
| DIP_10008822 | putative cell cycle control protein cwf14 protein [Eutypa lata UCREL1] | 80.3 | 3.0E-63 |
| DIP_10008823 | hypothetical protein FOZG_05362 [Fusarium oxysporum Fo47] | 51.3 | 2.0E-78 |
| DIP_10008824 | hypothetical protein PFICI_07477 [Pestalotiopsis fici W106-1] | 45.2 | 7.0E-156 |
| DIP_10008825 | putative adenosine kinase protein [Eutypa lata UCREL1] | 81.7 | 0.0 |
| DIP_10008826 | putative 60s ribosomal protein l2 protein [Eutypa lata UCREL1] | 91.6 | 3.0E-171 |
| DIP_10008827 | hypothetical protein PFICI_07480 [Pestalotiopsis fici W106-1] | 47 | 6.0E-77 |
| DIP_10008828 | hypothetical protein PFICI_11540 [Pestalotiopsis fici W106-1] | 57.5 | 0.0 |
| DIP_10008829 | GPI anchored protein [Aspergillus fumigatus Af293] | 30.6 | 4.0E-16 |
| DIP_10008830 | putative insig domain-containing protein [Togninia minima UCRPA7] | 47.8 | 1.0E-113 |
| DIP_10008831 | hypothetical protein PFICI_11538 [Pestalotiopsis fici W106-1] | 53.4 | 0.0 |
| DIP_10008832 | hypothetical protein TRIATDRAFT_283367 [Trichoderma atroviride IMI 206040] | 44.2 | 0.0 |
| DIP_10008833 | hypothetical protein PFICI_11536 [Pestalotiopsis fici W106-1] | 45.3 | 6.0E-75 |
| DIP_10008834 | hypothetical protein PFICI_00826 [Pestalotiopsis fici W106-1] | 70.5 | 0.0 |
| DIP_10008835 | peroxisomal 2,4-dienoyl-CoA reductase SPS19 [Verticillium alfalfae VaMs.102] | 71.6 | 1.0E-167 |
| DIP_10008836 | transcriptional regulator [Aspergillus kawachii IFO 4308] | 63.4 | 3.0E-112 |
| DIP_10008837 | hypothetical protein EURHEDRAFT_531356 [Aspergillus ruber CBS 135680] | 50.4 | 1.0E-80 |
| DIP_10008838 | hypothetical protein UCREL1_8152 [Eutypa lata UCREL1] | 57.3 | 1.0E-74 |
| DIP_10008839 | NA | | |
| DIP_10008840 | putative px domain-containing protein [Eutypa lata UCREL1] | 39.4 | 6.0E-78 |
| DIP_10008841 | putative elmo ced-12 family protein [Togninia minima UCRPA7] | 70.3 | 0.0 |
| DIP_10008842 | NA | | |
| DIP_10008843 | putative sugar transporter protein [Eutypa lata UCREL1] | 83.4 | 0.0 |
| DIP_10008844 | hypothetical protein PFICI_00476 [Pestalotiopsis fici W106-1] | 69.5 | 3.0E-74 |
| DIP_10008845 | hypothetical protein MYCGRDRAFT_73708 [Zymoseptoria tritici IPO323] | 60.7 | 8.0E-151 |
| DIP_10008846 | putative pumilio domain-containing protein [Eutypa lata UCREL1] | 58.9 | 0.0 |
| DIP_10008847 | hypothetical protein UCREL1_2206 [Eutypa lata UCREL1] | 67.9 | 8.0E-78 |
| DIP_10008848 | hypothetical protein PFICI_07154 [Pestalotiopsis fici W106-1] | 64.6 | 9.0E-143 |
| DIP_10008849 | putative gpi ethanolamine phosphate transferase protein [Togninia minima UCRPA7] | 62.3 | 0.0 |
| DIP_10008850 | putative c6 transcription factor protein [Eutypa lata UCREL1] | 60.3 | 1.0E-87 |
| DIP_10008851 | putative leptomycin b resistance protein pmd1 protein [Eutypa lata UCREL1] | 76.2 | 0.0 |
| DIP_10008852 | NA | | |
| DIP_10008853 | transmembrane protein [Beauveria bassiana ARSEF 2860] | 48.6 | 8.0E-53 |
| DIP_10008854 | putative ring-12 protein [Eutypa lata UCREL1] | 61.7 | 0.0 |
| DIP_10008855 | putative transcription factor tfiiib component protein [Eutypa lata UCREL1] | 61.4 | 7.0E-101 |
| DIP_10008856 | hypothetical protein OIDMADRAFT_154449 [Oidiodendron maius Zn] | 63 | 5.0E-148 |
| DIP_10008857 | hypothetical protein GLRG_02256 [Colletotrichum graminicola M1.001] | 38.4 | 4.0E-72 |
| DIP_10008858 | 4-aminobutyrate aminotransferase [Pestalotiopsis fici W106-1] | 77.5 | 0.0 |
| DIP_10008859 | putative histone h3 lysine 36 methyltransferase protein [Eutypa lata UCREL1] | 60.8 | 0.0 |
| DIP_10008860 | predicted protein [Chaetomium globosum CBS 148.51] | 42.2 | 7.0E-27 |
| DIP_10008861 | hypothetical protein UCREL1_2189 [Eutypa lata UCREL1] | 61 | 2.0E-38 |
| DIP_10008862 | hypothetical protein UCREL1_2189 [Eutypa lata UCREL1] | 56.6 | 6.0E-37 |
| DIP_10008863 | hypothetical protein UCREL1_2190 [Eutypa lata UCREL1] | 45.8 | 1.0E-151 |
| DIP_10008864 | putative dna damage tolerance protein rad31 protein [Eutypa lata UCREL1] | 74.8 | 0.0 |
| DIP_10008865 | hypothetical protein OIDMADRAFT_38295 [Oidiodendron maius Zn] | 44.3 | 8.0E-137 |
| DIP_10008866 | eukaryotic translation initiation factor 6 [Colletotrichum gloeosporioides Nara gc5] | 95.5 | 4.0E-174 |
| DIP_10008867 | hypothetical protein PFICI_05353 [Pestalotiopsis fici W106-1] | 58.9 | 0.0 |
| DIP_10008868 | putative dna repair helicase rad25 protein [Eutypa lata UCREL1] | 85.5 | 0.0 |
| DIP_10008869 | hypothetical protein PFICI_05525 [Pestalotiopsis fici W106-1] | 50.9 | 9.0E-119 |
| DIP_10008870 | putative ph domain-containing protein [Eutypa lata UCREL1] | 53.9 | 2.0E-91 |
| DIP_10008871 | putative ribonucleoside-diphosphate reductase small subunit protein [Eutypa lata UCREL1] | 84.7 | 0.0 |
| DIP_10008872 | hypothetical protein GLRG_08432 [Colletotrichum graminicola M1.001] | 55.1 | 1.0E-35 |
| DIP_10008873 | WD repeat protein [Colletotrichum orbiculare MAFF 240422] | 61.9 | 4.0E-169 |
| DIP_10008874 | NA | | |
| DIP_10008875 | Glycoside hydrolase family 28 [Macrophomina phaseolina MS6] | 67.4 | 0.0 |
| DIP_10008876 | hypothetical protein PFICI_05531 [Pestalotiopsis fici W106-1] | 72.1 | 0.0 |
| DIP_10008877 | Ribosomal protein S8 [Glarea lozoyensis ATCC 20868] | 93.1 | 4.0E-88 |
| DIP_10008878 | putative 40s ribosomal protein s12 protein [Eutypa lata UCREL1] | 96.6 | 6.0E-100 |
| DIP_10008879 | Peptidyl-prolyl cis-trans isomerase PIN4 [Ceratocystis platani] | 51.6 | 3.0E-34 |
| DIP_10008880 | hypothetical protein PFICI_12448 [Pestalotiopsis fici W106-1] | 57.8 | 6.0E-93 |
| DIP_10008881 | hypothetical protein PFICI_12447 [Pestalotiopsis fici W106-1] | 58.6 | 0.0 |
| DIP_10008882 | hypothetical protein PFICI_05510 [Pestalotiopsis fici W106-1] | 69.8 | 0.0 |
| DIP_10008883 | predicted protein [Fibroporia radiculosa] | 32.1 | 4.0E-24 |
| DIP_10008884 | NA | | |
| DIP_10008885 | hypothetical protein PFICI_02452 [Pestalotiopsis fici W106-1] | 48.9 | 0.0 |
| DIP_10008886 | hypothetical protein PFICI_14549 [Pestalotiopsis fici W106-1] | 50.9 | 0.0 |
| DIP_10008887 | putative DNA repair and recombination protein pif1 [Colletotrichum sublineola] | 58.6 | 0.0 |
| DIP_10008888 | DNA ligase I [Colletotrichum fioriniae PJ7] | 55.8 | 0.0 |
| DIP_10008889 | hypothetical protein PFICI_14547 [Pestalotiopsis fici W106-1] | 75.8 | 6.0E-122 |
| DIP_10008890 | hypothetical protein PFICI_14548 [Pestalotiopsis fici W106-1] | 58.7 | 0.0 |
| DIP_10008891 | NA | | |
| DIP_10008892 | hypothetical protein UCREL1_486 [Eutypa lata UCREL1] | 71.9 | 0.0 |
| DIP_10008893 | hypothetical protein [Podospora anserina S mat+] | 62.6 | 9.0E-103 |
| DIP_10008894 | Cytochrome P450 [Macrophomina phaseolina MS6] | 42.8 | 6.0E-148 |
| DIP_10008895 | hypothetical protein PFICI_12509 [Pestalotiopsis fici W106-1] | 49.2 | 0.0 |
| DIP_10008896 | ATP-dependent RNA helicase DRS1 [Pestalotiopsis fici W106-1] | 67.1 | 0.0 |
| DIP_10008897 | cytochrome P450, putative [Talaromyces stipitatus ATCC 10500] | 65.6 | 0.0 |
| DIP_10008898 | predicted protein [Blastomyces dermatitidis ER-3] | 36.3 | 3.0E-35 |
| DIP_10008899 | hypothetical protein PFICI_03840 [Pestalotiopsis fici W106-1] | 56.1 | 0.0 |
| DIP_10008900 | putative microtubule-associated protein rp eb family member 1 protein [Togninia minima UCRPA7] | 85.2 | 3.0E-148 |
| DIP_10008901 | NA | | |
| DIP_10008902 | NA | | |
| DIP_10008903 | NA | | |
| DIP_10008904 | NA | | |
| DIP_10008905 | hypothetical protein PFICI_14497 [Pestalotiopsis fici W106-1] | 68.1 | 3.0E-156 |
| DIP_10008906 | NA | | |
| DIP_10008907 | putative dyp-type peroxidase protein [Eutypa lata UCREL1] | 71.1 | 4.0E-110 |
| DIP_10008908 | hypothetical protein PFICI_05781 [Pestalotiopsis fici W106-1] | 63.4 | 0.0 |
| DIP_10008909 | hypothetical protein PFICI_05782 [Pestalotiopsis fici W106-1] | 48.4 | 0.0 |
| DIP_10008910 | hypothetical protein THITE_2171621 [Thielavia terrestris NRRL 8126] | 56.1 | 2.0E-146 |
| DIP_10008911 | putative abc multidrug transporter protein [Eutypa lata UCREL1] | 69.3 | 0.0 |
| DIP_10008912 | putative siderophore iron transporter protein [Eutypa lata UCREL1] | 67.7 | 0.0 |
| DIP_10008913 | hypothetical protein PFICI_05789 [Pestalotiopsis fici W106-1] | 67.4 | 1.0E-161 |
| DIP_10008914 | putative amp-binding enzyme protein [Eutypa lata UCREL1] | 77.5 | 2.0E-134 |
| DIP_10008915 | NA | | |
| DIP_10008916 | glycoside hydrolase family 35 protein [Thielavia terrestris NRRL 8126] | 54.8 | 0.0 |
| DIP_10008917 | NA | | |
| DIP_10008918 | putative aerobactin siderophore biosynthesis protein iucb protein [Eutypa lata UCREL1] | 67.8 | 0.0 |
| DIP_10008919 | putative pyridine nucleotide-disulfide oxidoreductase amid- protein [Eutypa lata UCREL1] | 63.2 | 4.0E-156 |
| DIP_10008920 | putative gtp-binding protein 1 protein [Eutypa lata UCREL1] | 81.2 | 0.0 |
| DIP_10008921 | fatty acid desaturase [Colletotrichum fioriniae PJ7] | 60.6 | 0.0 |
| DIP_10008922 | putative lmbr1 domain-containing protein [Eutypa lata UCREL1] | 68.1 | 7.0E-174 |
| DIP_10008923 | hypothetical protein PFICI_05770 [Pestalotiopsis fici W106-1] | 67.2 | 0.0 |
| DIP_10008924 | putative dna repair protein rad5 [Diaporthe ampelina] | 74.1 | 0.0 |
| DIP_10008925 | hypothetical protein PFICI_02653 [Pestalotiopsis fici W106-1] | 57.1 | 0.0 |
| DIP_10008926 | hypothetical protein S7711_11393 [Stachybotrys chartarum IBT 7711] | 33.3 | 4.0E-07 |
| DIP_10008927 | NA | | |
| DIP_10008928 | NA | | |
| DIP_10008929 | putative tyrosinase protein [Eutypa lata UCREL1] | 55.6 | 4.0E-126 |
| DIP_10008930 | NA | | |
| DIP_10008931 | NA | | |
| DIP_10008932 | putative chitin synthase export chaperone protein [Eutypa lata UCREL1] | 77.8 | 0.0 |
| DIP_10008933 | hypothetical protein UCREL1_2348 [Eutypa lata UCREL1] | 47.3 | 2.0E-36 |
| DIP_10008934 | hypothetical protein UCDDA912_g06106 [Diaporthe ampelina] | 52.2 | 1.0E-75 |
| DIP_10008935 | NA | | |
| DIP_10008936 | putative pre-mrna-splicing factor 38b protein [Eutypa lata UCREL1] | 51.1 | 3.0E-107 |
| DIP_10008937 | RAB5-like protein [Colletotrichum orbiculare MAFF 240422] | 86.1 | 2.0E-142 |
| DIP_10008938 | putative ribosome biogenesis protein kri1 protein [Eutypa lata UCREL1] | 60.7 | 0.0 |
| DIP_10008939 | NA | | |
| DIP_10008940 | hypothetical protein UCREL1_7521 [Eutypa lata UCREL1] | 40.3 | 5.0E-45 |
| DIP_10008941 | 3-oxo-5-alpha-steroid 4-dehydrogenase [Colletotrichum higginsianum] | 50 | 3.0E-110 |
| DIP_10008942 | NA | | |
| DIP_10008943 | ankyrin repeat protein [Verticillium dahliae VdLs.17] | 43.4 | 0.0 |
| DIP_10008944 | GTP-binding protein [Colletotrichum orbiculare MAFF 240422] | 92.4 | 0.0 |
| DIP_10008945 | NA | | |
| DIP_10008946 | hypothetical protein PFICI_03782 [Pestalotiopsis fici W106-1] | 41 | 1.0E-114 |
| DIP_10008947 | NA | | |
| DIP_10008948 | PXA domain-containing protein [Colletotrichum graminicola M1.001] | 52.7 | 2.0E-137 |
| DIP_10008949 | Pre-mRNA-splicing factor rse-1 [Pestalotiopsis fici W106-1] | 86.8 | 0.0 |
| DIP_10008950 | hypothetical protein PFICI_03784 [Pestalotiopsis fici W106-1] | 64.5 | 0.0 |
| DIP_10008951 | hypothetical protein PFICI_03786 [Pestalotiopsis fici W106-1] | 38.4 | 2.0E-58 |
| DIP_10008952 | putative protein pbn1 protein [Eutypa lata UCREL1] | 49 | 4.0E-175 |
| DIP_10008953 | hypothetical protein PFICI_07986 [Pestalotiopsis fici W106-1] | 56.7 | 1.0E-120 |
| DIP_10008954 | hypothetical protein UCREL1_8241 [Eutypa lata UCREL1] | 58 | 5.0E-146 |
| DIP_10008955 | hypothetical protein UCREL1_8242 [Eutypa lata UCREL1] | 44.7 | 4.0E-68 |
| DIP_10008956 | putative heat shock protein hsp98 protein [Eutypa lata UCREL1] | 86.8 | 0.0 |
| DIP_10008957 | hypothetical protein PFICI_07982 [Pestalotiopsis fici W106-1] | 52.8 | 0.0 |
| DIP_10008958 | putative l-ascorbic acid binding protein [Togninia minima UCRPA7] | 36.2 | 1.0E-63 |
| DIP_10008959 | NA | | |
| DIP_10008960 | putative tetratricopeptide repeat protein 1 protein [Eutypa lata UCREL1] | 72.6 | 0.0 |
| DIP_10008961 | putative ste ste20 paka protein kinase protein [Eutypa lata UCREL1] | 81.3 | 0.0 |
| DIP_10008962 | NA | | |
| DIP_10008963 | hypothetical protein FOQG_15881 [Fusarium oxysporum f. sp. raphani 54005] | 45.3 | 3.0E-142 |
| DIP_10008964 | putative 2og-fe oxygenase family protein [Eutypa lata UCREL1] | 42.1 | 2.0E-70 |
| DIP_10008965 | NA | | |
| DIP_10008966 | NA | | |
| DIP_10008967 | NA | | |
| DIP_10008968 | NA | | |
| DIP_10008969 | predicted protein [Chaetomium globosum CBS 148.51] | 52.3 | 1.0E-28 |
| DIP_10008970 | hypothetical protein PFICI_08002 [Pestalotiopsis fici W106-1] | 71.6 | 0.0 |
| DIP_10008971 | Histone-fold domain containing protein [Cordyceps militaris CM01] | 52 | 6.0E-47 |
| DIP_10008972 | hypothetical protein PFICI_08029 [Pestalotiopsis fici W106-1] | 66 | 0.0 |
| DIP_10008973 | hypothetical protein THITE_2118925 [Thielavia terrestris NRRL 8126] | 69.7 | 0.0 |
| DIP_10008974 | hypothetical protein PFICI_08009 [Pestalotiopsis fici W106-1] | 79.2 | 0.0 |
| DIP_10008975 | hypothetical protein PFICI_08010 [Pestalotiopsis fici W106-1] | 69 | 0.0 |
| DIP_10008976 | hypothetical protein UCDDA912_g06677 [Diaporthe ampelina] | 51 | 2.0E-44 |
| DIP_10008977 | hypothetical protein PFICI_08012 [Pestalotiopsis fici W106-1] | 49.6 | 0.0 |
| DIP_10008978 | NA | | |
| DIP_10008979 | hypothetical protein PFICI_08013 [Pestalotiopsis fici W106-1] | 48.8 | 0.0 |
| DIP_10008980 | glycyl-trna synthetase [Colletotrichum orbiculare MAFF 240422] | 73.6 | 0.0 |
| DIP_10008981 | putative brix domain-containing protein [Eutypa lata UCREL1] | 77 | 0.0 |
| DIP_10008982 | NA | | |
| DIP_10008983 | hypothetical protein PFICI_08020 [Pestalotiopsis fici W106-1] | 61.6 | 0.0 |
| DIP_10008984 | putative nuclear cap binding protein subunit 2 protein [Eutypa lata UCREL1] | 77 | 8.0E-88 |
| DIP_10008985 | putative mitochondrial 54s ribosomal protein 31 protein [Togninia minima UCRPA7] | 86.4 | 5.0E-67 |
| DIP_10008986 | hypothetical protein MYCTH_2305195 [Myceliophthora thermophila ATCC 42464] | 61 | 1.0E-84 |
| DIP_10008987 | histone H4 [Botrytis cinerea B05.10] | 100 | 1.0E-64 |
| DIP_10008988 | putative 20s cyclosome subunit ( nuc2 cdc27) protein [Eutypa lata UCREL1] | 60.6 | 0.0 |
| DIP_10008989 | hypothetical protein TRIVIDRAFT_216889 [Trichoderma virens Gv29-8] | 63 | 5.0E-72 |
| DIP_10008990 | hypothetical protein MYCTH_2305190, partial [Myceliophthora thermophila ATCC 42464] | 50.5 | 4.0E-64 |
| DIP_10008991 | putative nadh-cytochrome b5 reductase 1 protein [Eutypa lata UCREL1] | 83.9 | 0.0 |
| DIP_10008992 | hypothetical protein PFICI_07989 [Pestalotiopsis fici W106-1] | 39.4 | 2.0E-64 |
| DIP_10008993 | hypothetical protein HIM_04554 [Hirsutella minnesotensis 3608] | 42.6 | 8.0E-162 |
| DIP_10008994 | hypothetical protein NECHADRAFT_47076 [Nectria haematococca mpVI 77-13-4] | 46.4 | 2.0E-66 |
| DIP_10008995 | NA | | |
| DIP_10008996 | NA | | |
| DIP_10008997 | hypothetical protein A1O5_00623 [Cladophialophora psammophila CBS 110553] | 33.1 | 2.0E-41 |
| DIP_10008998 | hypothetical protein PFICI_08005 [Pestalotiopsis fici W106-1] | 58.2 | 0.0 |
| DIP_10008999 | hypothetical protein PFICI_07998 [Pestalotiopsis fici W106-1] | 49.8 | 0.0 |
| DIP_10009000 | hypothetical protein PFICI_07996 [Pestalotiopsis fici W106-1] | 49.8 | 8.0E-84 |
| DIP_10009001 | hypothetical protein BC1G_10424 [Botrytis cinerea B05.10] | 67.1 | 1.0E-91 |
| DIP_10009002 | Exosome complex component rrp45 [Pestalotiopsis fici W106-1] | 75 | 3.0E-170 |
| DIP_10009003 | hypothetical protein PFICI_07993 [Pestalotiopsis fici W106-1] | 68.1 | 7.0E-132 |
| DIP_10009004 | hypothetical protein PFICI_07992 [Pestalotiopsis fici W106-1] | 51.2 | 3.0E-166 |
| DIP_10009005 | hypothetical protein PFICI_08038 [Pestalotiopsis fici W106-1] | 34.8 | 9.0E-126 |
| DIP_10009006 | NA | | |
| DIP_10009007 | putative carboxypeptidase y protein [Eutypa lata UCREL1] | 81.7 | 0.0 |
| DIP_10009008 | putative shwachman-bodian-diamond syndrome protein [Eutypa lata UCREL1] | 81.4 | 0.0 |
| DIP_10009009 | putative 40s ribosomal protein s18 protein [Eutypa lata UCREL1] | 95.8 | 2.0E-97 |
| DIP_10009010 | hypothetical protein PFICI_08041 [Pestalotiopsis fici W106-1] | 39.8 | 8.0E-13 |
| DIP_10009011 | hypothetical protein MGG_14658 [Magnaporthe oryzae 70-15] | 32.4 | 2.0E-58 |
| DIP_10009012 | hypothetical protein V491_00738 [Pseudogymnoascus pannorum VKM F-3775] | 45.9 | 4.0E-72 |
| DIP_10009013 | putative a49-like rna polymerase i associated factor protein [Eutypa lata UCREL1] | 60.9 | 0.0 |
| DIP_10009014 | NA | | |
| DIP_10009015 | hypothetical protein CHGG_05593 [Chaetomium globosum CBS 148.51] | 61.9 | 0.0 |
| DIP_10009016 | Ornithine aminotransferase [Pestalotiopsis fici W106-1] | 80.9 | 0.0 |
| DIP_10009017 | hypothetical protein SMAC_02235 [Sordaria macrospora k-hell] | 40.8 | 2.0E-129 |
| DIP_10009018 | hypothetical protein PFICI_11798 [Pestalotiopsis fici W106-1] | 56.9 | 0.0 |
| DIP_10009019 | putative n amino acid transport system protein [Eutypa lata UCREL1] | 71.2 | 0.0 |
| DIP_10009020 | MFS general substrate transporter [Glarea lozoyensis ATCC 20868] | 63 | 0.0 |
| DIP_10009021 | NA | | |
| DIP_10009022 | putative carboxyphosphonoenolpyruvate protein [Eutypa lata UCREL1] | 65.6 | 3.0E-141 |
| DIP_10009023 | putative ubiquitin ligase subunit protein [Eutypa lata UCREL1] | 60.1 | 0.0 |
| DIP_10009024 | Ydr124wp-like protein [Metarhizium acridum CQMa 102] | 37.5 | 2.0E-88 |
| DIP_10009025 | NA | | |
| DIP_10009026 | hypothetical protein PFICI_11770 [Pestalotiopsis fici W106-1] | 79 | 0.0 |
| DIP_10009027 | hypothetical protein PFICI_11768 [Pestalotiopsis fici W106-1] | 55.6 | 5.0E-95 |
| DIP_10009028 | hypothetical protein PFICI_11767 [Pestalotiopsis fici W106-1] | 48.7 | 0.0 |
| DIP_10009029 | hypothetical protein UCREL1_1414 [Eutypa lata UCREL1] | 64.9 | 2.0E-29 |
| DIP_10009030 | putative amino-acid permease inda1 protein [Eutypa lata UCREL1] | 65.3 | 0.0 |
| DIP_10009031 | NA | | |
| DIP_10009032 | hypothetical protein S40285_09378 [Stachybotrys chlorohalonata IBT 40285] | 47.5 | 9.0E-114 |
| DIP_10009033 | putative mitochondrial cytochrome protein [Eutypa lata UCREL1] | 63.3 | 0.0 |
| DIP_10009034 | putative serine threonine-protein kinase ppk30 protein [Eutypa lata UCREL1] | 78.3 | 9.0E-163 |
| DIP_10009035 | hypothetical protein PFICI_11775 [Pestalotiopsis fici W106-1] | 51.4 | 2.0E-111 |
| DIP_10009036 | hypothetical protein PFICI_11776 [Pestalotiopsis fici W106-1] | 69.6 | 0.0 |
| DIP_10009037 | putative membrane protein [Eutypa lata UCREL1] | 51.3 | 6.0E-48 |
| DIP_10009038 | hypothetical protein TRIATDRAFT_28757 [Trichoderma atroviride IMI 206040] | 39.8 | 8.0E-06 |
| DIP_10009039 | hypothetical protein PFICI_04150 [Pestalotiopsis fici W106-1] | 40.2 | 1.0E-38 |
| DIP_10009040 | hypothetical protein UCREL1_10786 [Eutypa lata UCREL1] | 52.5 | 3.0E-56 |
| DIP_10009041 | nuclear divisionprotein rft1 [Sporothrix brasiliensis 5110] | 52.3 | 3.0E-80 |
| DIP_10009042 | pyruvate dehydrogenase E1 component alpha subunit [Metarhizium album ARSEF 1941] | 81.9 | 0.0 |
| DIP_10009043 | putative prephenate dehydratase protein [Eutypa lata UCREL1] | 48.3 | 2.0E-91 |
| DIP_10009044 | hypothetical protein FOXB_17523 [Fusarium oxysporum Fo5176] | 87.4 | 0.0 |
| DIP_10009045 | hypothetical protein UCREL1_10523 [Eutypa lata UCREL1] | 62.8 | 0.0 |
| DIP_10009046 | hypothetical protein PFICI_11782 [Pestalotiopsis fici W106-1] | 63.1 | 2.0E-76 |
| DIP_10009047 | putative pentatricopeptide repeat protein [Eutypa lata UCREL1] | 37 | 4.0E-121 |
| DIP_10009048 | putative dna polymerase alpha catalytic subunit protein [Eutypa lata UCREL1] | 77.4 | 0.0 |
| DIP_10009049 | MIF4-like, type 1/2/3 [Metarhizium guizhouense ARSEF 977] | 52.8 | 0.0 |
| DIP_10009050 | putative intron-binding protein aquarius protein [Eutypa lata UCREL1] | 76.4 | 0.0 |
| DIP_10009051 | putative tho complex subunit protein [Eutypa lata UCREL1] | 56.1 | 0.0 |
| DIP_10009052 | hypothetical protein CSUB01_03025 [Colletotrichum sublineola] | 76.2 | 0.0 |
| DIP_10009053 | putative nuclear and cytoplasmic polyadenylated rna-binding protein pub1 protein [Eutypa lata UCREL1] | 89.6 | 0.0 |
| DIP_10009054 | hypothetical protein PFICI_07483 [Pestalotiopsis fici W106-1] | 37.9 | 1.0E-55 |
| DIP_10009055 | putative cellulase family protein [Eutypa lata UCREL1] | 68.3 | 0.0 |
| DIP_10009056 | hypothetical protein PFICI_07488 [Pestalotiopsis fici W106-1] | 69.4 | 4.0E-77 |
| DIP_10009057 | hypothetical protein VHEMI02754 [Torrubiella hemipterigena] | 38.8 | 6.0E-60 |
| DIP_10009058 | putative microtubule associated protein [Eutypa lata UCREL1] | 82.4 | 0.0 |
| DIP_10009059 | putative glycoside hydrolase family 128 protein [Diaporthe ampelina] | 42.4 | 7.0E-71 |
| DIP_10009060 | putative isoflavone reductase protein [Eutypa lata UCREL1] | 45.7 | 6.0E-84 |
| DIP_10009061 | hypothetical protein GYMLUDRAFT_116993, partial [Gymnopus luxurians FD-317 M1] | 30.3 | 7.0E-13 |
| DIP_10009062 | hypothetical protein FGSG_06945 [Fusarium graminearum PH-1] | 73 | 0.0 |
| DIP_10009063 | hypothetical protein MYCTH_2303482 [Myceliophthora thermophila ATCC 42464] | 76 | 0.0 |
| DIP_10009064 | hypothetical protein PFICI_00829 [Pestalotiopsis fici W106-1] | 65.3 | 0.0 |
| DIP_10009065 | hypothetical protein PFICI_00830 [Pestalotiopsis fici W106-1] | 65.5 | 0.0 |
| DIP_10009066 | putative pt repeat family protein [Eutypa lata UCREL1] | 52.8 | 7.0E-157 |
| DIP_10009067 | putative heat shock protein 78 protein [Eutypa lata UCREL1] | 86.6 | 0.0 |
| DIP_10009068 | hypothetical protein TRIATDRAFT_298891 [Trichoderma atroviride IMI 206040] | 78.2 | 4.0E-124 |
| DIP_10009069 | putative bifunctional purine biosynthesis protein [Eutypa lata UCREL1] | 86.2 | 0.0 |
| DIP_10009070 | DNA replication complex GINS protein PSF1 [Metarhizium acridum CQMa 102] | 86.2 | 5.0E-135 |
| DIP_10009071 | hypothetical protein PFICI_00834 [Pestalotiopsis fici W106-1] | 66.5 | 0.0 |
| DIP_10009072 | putative tigr02452 family protein [Eutypa lata UCREL1] | 45.4 | 2.0E-103 |
| DIP_10009073 | hypothetical protein PFICI_01498 [Pestalotiopsis fici W106-1] | 64.6 | 4.0E-174 |
| DIP_10009074 | hypothetical protein UCREL1_596 [Eutypa lata UCREL1] | 70.2 | 0.0 |
| DIP_10009075 | putative glycine cleavage system t protein [Eutypa lata UCREL1] | 63.3 | 0.0 |
| DIP_10009076 | RAS-like GTPase, Rho2 subfamily [Metarhizium robertsii ARSEF 23] | 98.9 | 4.0E-134 |
| DIP_10009077 | putative endosomal protein p24b protein [Eutypa lata UCREL1] | 79.3 | 2.0E-120 |
| DIP_10009078 | putative feruloyl esterase b protein [Eutypa lata UCREL1] | 48.6 | 1.0E-175 |
| DIP_10009079 | hypothetical protein SMAC_05728 [Sordaria macrospora k-hell] | 62 | 4.0E-168 |
| DIP_10009080 | putative 60s ribosomal protein l6 protein [Eutypa lata UCREL1] | 84.5 | 1.0E-123 |
| DIP_10009081 | putative peptidase family m49 protein [Eutypa lata UCREL1] | 78.9 | 0.0 |
| DIP_10009082 | 26S protease regulatory subunit 7 [Pestalotiopsis fici W106-1] | 97.5 | 0.0 |
| DIP_10009083 | T-complex protein 1 subunit delta [Pestalotiopsis fici W106-1] | 91.8 | 0.0 |
| DIP_10009084 | hypothetical protein PFICI_05635 [Pestalotiopsis fici W106-1] | 61 | 0.0 |
| DIP_10009085 | hypothetical protein PFICI_05634 [Pestalotiopsis fici W106-1] | 53.7 | 0.0 |
| DIP_10009086 | NA | | |
| DIP_10009087 | hypothetical protein PFICI_05632 [Pestalotiopsis fici W106-1] | 56.6 | 9.0E-176 |
| DIP_10009088 | F-box domain containing protein [Colletotrichum gloeosporioides Nara gc5] | 43.2 | 0.0 |
| DIP_10009089 | putative pkr1-domain-containing protein [Eutypa lata UCREL1] | 67.7 | 3.0E-58 |
| DIP_10009090 | putative small nuclear ribonucleoprotein sm d1 protein [Eutypa lata UCREL1] | 91.9 | 4.0E-61 |
| DIP_10009091 | putative n -glutamine methyltransferase mtq2 protein [Eutypa lata UCREL1] | 52.7 | 1.0E-90 |
| DIP_10009092 | hypothetical protein PFICI_01964 [Pestalotiopsis fici W106-1] | 69.3 | 0.0 |
| DIP_10009093 | endonuclease LCL3 [Pestalotiopsis fici W106-1] | 69.7 | 6.0E-132 |
| DIP_10009094 | putative cellular morphogenesis protein [Eutypa lata UCREL1] | 55.9 | 0.0 |
| DIP_10009095 | hypothetical protein PFICI_02017 [Pestalotiopsis fici W106-1] | 66.1 | 5.0E-149 |
| DIP_10009096 | NA | | |
| DIP_10009097 | putative u2 snrnp component prp10 protein [Eutypa lata UCREL1] | 88.3 | 0.0 |
| DIP_10009098 | hypothetical protein PFICI_01748 [Pestalotiopsis fici W106-1] | 64.5 | 0.0 |
| DIP_10009099 | putative g-patch domain protein [Togninia minima UCRPA7] | 42.7 | 3.0E-80 |
| DIP_10009100 | hypothetical protein PFICI_01749 [Pestalotiopsis fici W106-1] | 54.3 | 0.0 |
| DIP_10009101 | hypothetical protein PFICI_00416 [Pestalotiopsis fici W106-1] | 68.4 | 0.0 |
| DIP_10009102 | putative wd repeat domain-containing protein [Eutypa lata UCREL1] | 44.3 | 3.0E-145 |
| DIP_10009103 | DnaJ domain-containing protein [Colletotrichum fioriniae PJ7] | 75.7 | 0.0 |
| DIP_10009104 | NA | | |
| DIP_10009105 | putative rna polymerase ii- paf1 protein [Eutypa lata UCREL1] | 73.2 | 0.0 |
| DIP_10009106 | putative rer1 protein [Eutypa lata UCREL1] | 88.4 | 5.0E-116 |
| DIP_10009107 | putative enoyl- hydratase protein [Togninia minima UCRPA7] | 78.5 | 3.0E-172 |
| DIP_10009108 | hypothetical protein GLRG_02308 [Colletotrichum graminicola M1.001] | 47.3 | 0.0 |
| DIP_10009109 | hypothetical protein UCREL1_1801 [Eutypa lata UCREL1] | 50 | 2.0E-39 |
| DIP_10009110 | putative trna pseudouridine synthase d protein [Eutypa lata UCREL1] | 45.5 | 0.0 |
| DIP_10009111 | hypothetical protein PFICI_00335 [Pestalotiopsis fici W106-1] | 38 | 5.0E-44 |
| DIP_10009112 | putative f-box wd repeat-containing protein 7 protein [Eutypa lata UCREL1] | 68.6 | 0.0 |
| DIP_10009113 | hypothetical protein PFICI_12645 [Pestalotiopsis fici W106-1] | 66.5 | 7.0E-112 |
| DIP_10009114 | putative histone acetyltransferase type b catalytic subunit protein [Eutypa lata UCREL1] | 56.6 | 0.0 |
| DIP_10009115 | pyrroline-5-carboxylate reductase [Gaeumannomyces graminis var. tritici R3-111a-1] | 67.3 | 9.0E-151 |
| DIP_10009116 | hypothetical protein FGSG_00095 [Fusarium graminearum PH-1] | 56.6 | 8.0E-160 |
| DIP_10009117 | hypothetical protein PFICI_05275 [Pestalotiopsis fici W106-1] | 42.1 | 0.0 |
| DIP_10009118 | Protein VTS1 [Scedosporium apiospermum] | 61.1 | 0.0 |
| DIP_10009119 | 60S ribosomal protein L29 [Pestalotiopsis fici W106-1] | 95.4 | 5.0E-39 |
| DIP_10009120 | hypothetical protein PFICI_05272 [Pestalotiopsis fici W106-1] | 81.6 | 0.0 |
| DIP_10009121 | hypothetical protein PFICI_05270 [Pestalotiopsis fici W106-1] | 52.8 | 0.0 |
| DIP_10009122 | NA | | |
| DIP_10009123 | hypothetical protein PFICI_05266 [Pestalotiopsis fici W106-1] | 69.9 | 3.0E-151 |
| DIP_10009124 | hypothetical protein VDAG_06612, partial [Verticillium dahliae VdLs.17] | 49.1 | 1.0E-12 |
| DIP_10009125 | putative alpha-l-rhamnosidase c protein [Eutypa lata UCREL1] | 54.2 | 2.0E-144 |
| DIP_10009126 | integral membrane protein [Gaeumannomyces graminis var. tritici R3-111a-1] | 71.4 | 0.0 |
| DIP_10009127 | hypothetical protein UVI_037400 [Ustilaginoidea virens] | 86.2 | 2.0E-67 |
| DIP_10009128 | hypothetical protein PFICI_05261 [Pestalotiopsis fici W106-1] | 69.1 | 0.0 |
| DIP_10009129 | NA | | |
| DIP_10009130 | hypothetical protein PFICI_05262 [Pestalotiopsis fici W106-1] | 33.7 | 4.0E-84 |
| DIP_10009131 | putative eukaryotic translation initiation factor 3 subunit clu1 protein [Eutypa lata UCREL1] | 76.2 | 0.0 |
| DIP_10009132 | hypothetical protein CGLO_08416 [Colletotrichum gloeosporioides Cg-14] | 50.9 | 2.0E-85 |
| DIP_10009133 | hypothetical protein FVEG_07145 [Fusarium verticillioides 7600] | 46.9 | 2.0E-166 |
| DIP_10009134 | hypothetical protein V500_07426 [Pseudogymnoascus pannorum VKM F-4518 (FW-2643)] | 33.1 | 5.0E-07 |
| DIP_10009135 | putative glycoside hydrolase family 79 protein [Togninia minima UCRPA7] | 31.8 | 3.0E-61 |
| DIP_10009136 | putative alkyl hydroperoxide reductase subunit c thiol specific antioxidant protein [Eutypa lata UCREL1] | 68.3 | 8.0E-98 |
| DIP_10009137 | putative misato segment ii myosin-like domain-containing protein [Eutypa lata UCREL1] | 65.8 | 0.0 |
| DIP_10009138 | NA | | |
| DIP_10009139 | STE12-like protein [Trichoderma atroviride IMI 206040] | 60.2 | 0.0 |
| DIP_10009140 | uncharacterized protein FFUJ_03148 [Fusarium fujikuroi IMI 58289] | 39.7 | 5.0E-55 |
| DIP_10009141 | hypothetical protein CFIO01_05689 [Colletotrichum fioriniae PJ7] | 68 | 2.0E-136 |
| DIP_10009142 | putative coronin-6 protein [Eutypa lata UCREL1] | 75.7 | 0.0 |
| DIP_10009143 | putative fact complex subunit pob-3 protein [Eutypa lata UCREL1] | 84.4 | 0.0 |
| DIP_10009144 | hypothetical protein PFICI_05254 [Pestalotiopsis fici W106-1] | 61.6 | 0.0 |
| DIP_10009145 | putative atp-dependent rna helicase dbp9 protein [Togninia minima UCRPA7] | 62 | 0.0 |
| DIP_10009146 | putative tata-binding protein-associated factor mot1 protein [Eutypa lata UCREL1] | 83.7 | 0.0 |
| DIP_10009147 | putative chromosome segregation protein [Eutypa lata UCREL1] | 83.2 | 0.0 |

| | | | |
|---|---|---|---|
| DIP_10009148 | putative gnat family protein [Eutypa lata UCREL1] | 54.4 | 1.0E-84 |
| DIP_10009149 | putative hsp90 co-chaperone cdc37 protein [Eutypa lata UCREL1] | 68 | 0.0 |
| DIP_10009150 | hypothetical protein CFIO01_05678 [Colletotrichum fioriniae PJ7] | 57.4 | 1.0E-134 |
| DIP_10009151 | UPF0045 protein ECM15 [Pestalotiopsis fici W106-1] | 81.7 | 5.0E-63 |
| DIP_10009152 | putative duf1682 domain protein [Eutypa lata UCREL1] | 68.1 | 0.0 |
| DIP_10009153 | hypothetical protein PFICI_05533 [Pestalotiopsis fici W106-1] | 54.9 | 0.0 |
| DIP_10009154 | putative catalase 3 protein [Eutypa lata UCREL1] | 77.5 | 0.0 |
| DIP_10009155 | hypothetical protein UCREL1_4769 [Eutypa lata UCREL1] | 40.1 | 3.0E-26 |
| DIP_10009156 | hypothetical protein PFICI_12668 [Pestalotiopsis fici W106-1] | 78.6 | 0.0 |
| DIP_10009157 | MFS general substrate transporter [Glarea lozoyensis ATCC 20868] | 62.3 | 0.0 |
| DIP_10009158 | NA | | |
| DIP_10009159 | NA | | |
| DIP_10009160 | putative sporulation protein [Eutypa lata UCREL1] | 46.5 | 4.0E-58 |
| DIP_10009161 | hypothetical protein PFICI_12674 [Pestalotiopsis fici W106-1] | 36.5 | 3.0E-48 |
| DIP_10009162 | putative peroxisomal-coenzyme a synthetase protein [Eutypa lata UCREL1] | 67.1 | 0.0 |
| DIP_10009163 | NA | | |
| DIP_10009164 | NA | | |
| DIP_10009165 | NA | | |
| DIP_10009166 | NA | | |
| DIP_10009167 | NA | | |
| DIP_10009168 | NA | | |
| DIP_10009169 | Cysteine synthase [Pestalotiopsis fici W106-1] | 76.1 | 0.0 |
| DIP_10009170 | hypothetical protein UCREL1_1796 [Eutypa lata UCREL1] | 45.7 | 3.0E-93 |
| DIP_10009171 | putative ribosomal protein l19 protein [Eutypa lata UCREL1] | 64.4 | 6.0E-92 |
| DIP_10009172 | NA | | |
| DIP_10009173 | hypothetical protein UCRPA7_6335 [Togninia minima UCRPA7] | 50.1 | 2.0E-92 |
| DIP_10009174 | putative nadh:ubiquinone oxidoreductase subunit protein [Eutypa lata UCREL1] | 74 | 7.0E-45 |
| DIP_10009175 | putative ccr4-not transcription complex subunit 7 protein [Togninia minima UCRPA7] | 63.9 | 0.0 |
| DIP_10009176 | hypothetical protein UVI_012060 [Ustilaginoidea virens] | 49.1 | 8.0E-124 |
| DIP_10009177 | putative helix-loop-helix dna binding protein [Eutypa lata UCREL1] | 56.2 | 1.0E-94 |
| DIP_10009178 | putative eukaryotic translation initiation factor 5 protein [Eutypa lata UCREL1] | 79.6 | 0.0 |
| DIP_10009179 | NA | | |
| DIP_10009180 | putative dephospho- kinase protein [Eutypa lata UCREL1] | 78.1 | 1.0E-157 |
| DIP_10009181 | putative phosphoribosylformylglycinamidine cyclo-ligase protein [Eutypa lata UCREL1] | 74.7 | 0.0 |
| DIP_10009182 | hypothetical protein PV05_09809 [Exophiala xenobiotica] | 47.5 | 2.0E-142 |
| DIP_10009183 | NA | | |
| DIP_10009184 | lipase atg15 [Pestalotiopsis fici W106-1] | 72.5 | 0.0 |
| DIP_10009185 | hypothetical protein PFICI_11906 [Pestalotiopsis fici W106-1] | 59.3 | 0.0 |
| DIP_10009186 | hypothetical protein PFICI_11908 [Pestalotiopsis fici W106-1] | 82.2 | 0.0 |
| DIP_10009187 | D-lactate dehydrogenase [Pestalotiopsis fici W106-1] | 68.6 | 0.0 |
| DIP_10009188 | putative mitochondrial import inner membrane translocase subunit tim8 protein [Eutypa lata UCREL1] | 66.7 | 5.0E-38 |
| DIP_10009189 | putative 60s ribosomal protein l25 protein [Eutypa lata UCREL1] | 67.3 | 1.0E-63 |
| DIP_10009190 | hypothetical protein PFICI_11912 [Pestalotiopsis fici W106-1] | 68.7 | 1.0E-143 |
| DIP_10009191 | short-chain dehydrogenase, putative [Byssochlamys spectabilis No. 5] | 48.8 | 1.0E-97 |
| DIP_10009192 | putative -diketo-5-methylthio-1-phosphopentane phosphatase protein [Eutypa lata UCREL1] | 72.9 | 3.0E-129 |
| DIP_10009193 | putative serine threonine protein [Eutypa lata UCREL1] | 68.1 | 0.0 |
| DIP_10009194 | putative qde-2-interacting protein [Eutypa lata UCREL1] | 36.3 | 3.0E-97 |
| DIP_10009195 | mitochondrial import inner membrane translocase subunit tim-14 [Ophiostoma piceae UAMH 11346] | 86.5 | 2.0E-60 |
| DIP_10009196 | Ribosome biogenesis protein RLP24 [Pestalotiopsis fici W106-1] | 75.6 | 1.0E-114 |
| DIP_10009197 | hypothetical protein PFICI_07350 [Pestalotiopsis fici W106-1] | 47.9 | 1.0E-115 |
| DIP_10009198 | hypothetical protein PFICI_07349 [Pestalotiopsis fici W106-1] | 59 | 3.0E-156 |
| DIP_10009199 | phd finger domain-containing protein [Colletotrichum gloeosporioides Nara gc5] | 47.9 | 2.0E-120 |
| DIP_10009200 | NA | | |
| DIP_10009201 | hypothetical protein UCREL1_3087 [Eutypa lata UCREL1] | 63.9 | 4.0E-35 |
| DIP_10009202 | NA | | |
| DIP_10009203 | putative dolichyl-phosphate mannosyltransferase polypeptide 2 protein [Eutypa lata UCREL1] | 93.3 | 3.0E-44 |
| DIP_10009204 | putative cell cycle control protein [Eutypa lata UCREL1] | 87.9 | 0.0 |
| DIP_10009205 | hypothetical protein UCREL1_10346 [Eutypa lata UCREL1] | 48.7 | 2.0E-117 |
| DIP_10009206 | putative dihydrouridine synthase family protein [Togninia minima UCRPA7] | 62.3 | 0.0 |
| DIP_10009207 | hypothetical protein PFICI_08275 [Pestalotiopsis fici W106-1] | 58.8 | 0.0 |
| DIP_10009208 | hypothetical protein V492_06517 [Pseudogymnoascus pannorum VKM F-4246] | 43.7 | 0.0 |
| DIP_10009209 | hypothetical protein NECHADRAFT_34128 [Nectria haematococca mpVI 77-13-4] | 51.2 | 4.0E-86 |
| DIP_10009210 | hypothetical protein CFIO01_05694 [Colletotrichum fioriniae PJ7] | 42.5 | 5.0E-12 |
| DIP_10009211 | putative aaa family protein [Eutypa lata UCREL1] | 59.3 | 0.0 |
| DIP_10009212 | hypothetical protein HIM_04537 [Hirsutella minnesotensis 3608] | 38.4 | 2.0E-76 |
| DIP_10009213 | coiled-coil domain-containing protein 55, partial [Metarhizium majus ARSEF 297] | 51.9 | 1.0E-104 |
| DIP_10009214 | putative mitochondrial import inner membrane translocase subunit tim13 protein [Eutypa lata UCREL1] | 70 | 1.0E-39 |
| DIP_10009215 | putative serine threonine-protein kinase protein [Eutypa lata UCREL1] | 77.9 | 0.0 |
| DIP_10009216 | peptidase M76 family protein [Colletotrichum graminicola M1.001] | 72.4 | 3.0E-127 |
| DIP_10009217 | hypothetical protein PFICI_07502 [Pestalotiopsis fici W106-1] | 76.4 | 0.0 |
| DIP_10009218 | NA | | |
| DIP_10009219 | hypothetical protein TD95_002439 [Thielaviopsis punctulata] | 85 | 4.0E-121 |
| DIP_10009220 | putative nuclear export protein noc3 protein [Eutypa lata UCREL1] | 63 | 0.0 |
| DIP_10009221 | NA | | |
| DIP_10009222 | NA | | |
| DIP_10009223 | NA | | |
| DIP_10009224 | ubiquitin carboxyl-terminal hydrolase, partial [Colletotrichum higginsianum] | 45.7 | 0.0 |
| DIP_10009225 | hypothetical protein PFICI_07329 [Pestalotiopsis fici W106-1] | 45.1 | 2.0E-93 |
| DIP_10009226 | hypothetical protein UCREL1_3080 [Eutypa lata UCREL1] | 46.1 | 4.0E-89 |
| DIP_10009227 | putative mitotic checkpoint protein pccc protein [Eutypa lata UCREL1] | 49.2 | 4.0E-112 |
| DIP_10009228 | NA | | |
| DIP_10009229 | alcohol dehydrogenase [Colletotrichum fioriniae PJ7] | 54 | 1.0E-126 |
| DIP_10009230 | putative nucleoside transporter family protein [Eutypa lata UCREL1] | 57.2 | 6.0E-110 |
| DIP_10009231 | Nucleoside transporter family [Aspergillus flavus NRRL3357] | 54.9 | 5.0E-47 |
| DIP_10009232 | putative inositol-pentakisphosphate 2-kinase protein [Eutypa lata UCREL1] | 41.3 | 2.0E-80 |
| DIP_10009233 | NA | | |
| DIP_10009234 | beta-ketoacyl synthase domain-containing protein [Colletotrichum graminicola M1.001] | 37.1 | 0.0 |
| DIP_10009235 | O-methyltransferase, putative [Aspergillus clavatus NRRL 1] | 41 | 2.0E-107 |
| DIP_10009236 | putative mitochondrial distribution and morphology protein 34 protein [Eutypa lata UCREL1] | 56.6 | 0.0 |
| DIP_10009237 | NA | | |
| DIP_10009238 | integral membrane protein [Fusarium avenaceum] | 59.3 | 2.0E-169 |
| DIP_10009239 | uncharacterized protein FFUJ_09198 [Fusarium fujikuroi IMI 58289] | 82.4 | 0.0 |
| DIP_10009240 | predicted protein [Nectria haematococca mpVI 77-13-4] | 74.1 | 0.0 |
| DIP_10009241 | hypothetical protein FGSG_03935 [Fusarium graminearum PH-1] | 68.8 | 9.0E-138 |
| DIP_10009242 | NA | | |
| DIP_10009243 | putative transcription factor iiic-like protein [Eutypa lata UCREL1] | 46.8 | 5.0E-134 |
| DIP_10009244 | hypothetical protein HIM_04337 [Hirsutella minnesotensis 3608] | 83.9 | 3.0E-128 |
| DIP_10009245 | hypothetical protein UCREL1_8317 [Eutypa lata UCREL1] | 44.7 | 4.0E-91 |
| DIP_10009246 | putative phosducin protein [Eutypa lata UCREL1] | 72.4 | 5.0E-152 |
| DIP_10009247 | Guanine nucleotide-binding protein subunit beta [Pestalotiopsis fici W106-1] | 98.3 | 0.0 |
| DIP_10009248 | NA | | |
| DIP_10009249 | putative acetoacetate- ligase protein [Eutypa lata UCREL1] | 80.1 | 3.0E-172 |
| DIP_10009250 | putative serine palmitoyltransferase 2 protein [Eutypa lata UCREL1] | 81.2 | 0.0 |
| DIP_10009251 | putative protein kinase regulator ste50 protein [Eutypa lata UCREL1] | 76.2 | 0.0 |
| DIP_10009252 | putative anthranilate synthase component 2 protein [Eutypa lata UCREL1] | 76 | 0.0 |
| DIP_10009253 | hypothetical protein UCREL1_8313 [Eutypa lata UCREL1] | 56.8 | 0.0 |
| DIP_10009254 | putative mitochondrial import inner membrane translocase subunit tim-9 protein [Eutypa lata UCREL1] | 87.3 | 8.0E-48 |
| DIP_10009255 | putative mrna turnover protein 4 protein [Eutypa lata UCREL1] | 66 | 8.0E-129 |
| DIP_10009256 | putative u1 small nuclear ribonucleo protein [Eutypa lata UCREL1] | 78.5 | 2.0E-103 |
| DIP_10009257 | putative mitochondrial 54s ribosomal protein l51 protein [Eutypa lata UCREL1] | 75.9 | 9.0E-77 |
| DIP_10009258 | putative triosephosphate isomerase protein [Eutypa lata UCREL1] | 78.3 | 2.0E-144 |
| DIP_10009259 | hypothetical protein UCDDA912_g09871 [Diaporthe ampelina] | 30.8 | 6.0E-16 |
| DIP_10009260 | MIP family channel protein [Colletotrichum fioriniae PJ7] | 82.1 | 0.0 |
| DIP_10009261 | NA | | |
| DIP_10009262 | NA | | |
| DIP_10009263 | predicted protein [Chaetomium globosum CBS 148.51] | 43 | 6.0E-17 |
| DIP_10009264 | NA | | |
| DIP_10009265 | hypothetical protein OIDMADRAFT_59005 [Oidiodendron maius Zn] | 37 | 5.0E-24 |
| DIP_10009266 | Putative Transcriptional repressor tup1 [Torrubiella hemipterigena] | 68.8 | 0.0 |
| DIP_10009267 | hypothetical protein MBM_08953 [Marssonina brunnea f. sp. 'multigermtubi' MB_m1] | 46.7 | 8.0E-64 |
| DIP_10009268 | putative c-5 sterol desaturase protein [Eutypa lata UCREL1] | 71.4 | 0.0 |
| DIP_10009269 | putative protein precursor protein [Eutypa lata UCREL1] | 69.6 | 5.0E-61 |
| DIP_10009270 | putative 60s ribosomal protein l28 protein [Eutypa lata UCREL1] | 61.2 | 5.0E-55 |
| DIP_10009271 | putative malate dehydrogenase protein [Eutypa lata UCREL1] | 89.7 | 0.0 |
| DIP_10009272 | putative bet3 family protein [Togninia minima UCRPA7] | 65.4 | 6.0E-114 |
| DIP_10009273 | Zinc finger, MIZ-type, partial [Metarhizium majus ARSEF 297] | 45.1 | 3.0E-129 |
| DIP_10009274 | hypothetical protein PFICI_03753 [Pestalotiopsis fici W106-1] | 55.4 | 6.0E-33 |
| DIP_10009275 | hypothetical protein CGLO_14113 [Colletotrichum gloeosporioides Cg-14] | 49.1 | 8.0E-47 |
| DIP_10009276 | putative yl1 nuclear protein [Eutypa lata UCREL1] | 76.1 | 5.0E-135 |
| DIP_10009277 | NA | | |
| DIP_10009278 | hypothetical protein PFICI_03751 [Pestalotiopsis fici W106-1] | 64.8 | 6.0E-99 |
| DIP_10009279 | NA | | |
| DIP_10009280 | hypothetical protein CHGG_02936 [Chaetomium globosum CBS 148.51] | 53.5 | 0.0 |
| DIP_10009281 | NA | | |
| DIP_10009282 | putative syf2 splicing factor protein [Eutypa lata UCREL1] | 82.1 | 2.0E-142 |
| DIP_10009283 | hypothetical protein PFICI_03738 [Pestalotiopsis fici W106-1] | 71.1 | 1.0E-158 |
| DIP_10009284 | hypothetical protein PFICI_03737 [Pestalotiopsis fici W106-1] | 74.6 | 1.0E-139 |
| DIP_10009285 | hypothetical protein G647_07523 [Cladophialophora carrionii CBS 160.54] | 64.8 | 9.0E-104 |
| DIP_10009286 | nonribosomal peptide synthetase 12 [Fomitiporia mediterranea MF3/22] | 44.5 | 0.0 |
| DIP_10009287 | enoyl- hydratase [Colletotrichum gloeosporioides Nara gc5] | 78.2 | 1.0E-143 |
| DIP_10009288 | hypothetical protein MGG_01186 [Magnaporthe oryzae 70-15] | 45.5 | 7.0E-71 |
| DIP_10009289 | hypothetical protein SAPIO_CDS4767 [Scedosporium apiospermum] | 60.5 | 0.0 |
| DIP_10009290 | hypothetical protein CH063_15077 [Colletotrichum higginsianum] | 48.9 | 1.0E-117 |
| DIP_10009291 | putative ww domain-containing protein [Eutypa lata UCREL1] | 58.8 | 1.0E-102 |
| DIP_10009292 | glycoside hydrolase family 64 protein [Dothistroma septosporum NZE10] | 51.5 | 4.0E-156 |
| DIP_10009293 | hypothetical protein PFICI_05329 [Pestalotiopsis fici W106-1] | 47.3 | 0.0 |
| DIP_10009294 | hypothetical protein PFICI_05330 [Pestalotiopsis fici W106-1] | 62.4 | 0.0 |
| DIP_10009295 | hypothetical protein PFICI_05332 [Pestalotiopsis fici W106-1] | 55.4 | 1.0E-125 |
| DIP_10009296 | NA | | |
| DIP_10009297 | NA | | |
| DIP_10009298 | hypothetical protein PFICI_03810 [Pestalotiopsis fici W106-1] | 61.5 | 3.0E-155 |
| DIP_10009299 | hypothetical protein UCREL1_2497 [Eutypa lata UCREL1] | 39.9 | 1.0E-68 |
| DIP_10009300 | hypothetical protein CHGG_05757 [Chaetomium globosum CBS 148.51] | 64.5 | 0.0 |
| DIP_10009301 | NA | | |
| DIP_10009302 | NA | | |
| DIP_10009303 | putative bromodomain containing 1 protein [Eutypa lata UCREL1] | 67.2 | 0.0 |
| DIP_10009304 | putative glycoside hydrolase family 18 protein [Eutypa lata UCREL1] | 44.7 | 1.0E-100 |
| DIP_10009305 | hypothetical protein PFICI_02607 [Pestalotiopsis fici W106-1] | 58.4 | 8.0E-144 |
| DIP_10009306 | putative fad binding domain containing protein [Eutypa lata UCREL1] | 83.3 | 0.0 |
| DIP_10009307 | NA | | |
| DIP_10009308 | hypothetical protein V498_07008 [Pseudogymnoascus pannorum VKM F-4517 (FW-2822)] | 98 | 0.0 |
| DIP_10009309 | NA | | |
| DIP_10009310 | putative selenoprotein domain-containing protein [Eutypa lata UCREL1] | 65.1 | 1.0E-59 |
| DIP_10009311 | hypothetical protein PFICI_03802 [Pestalotiopsis fici W106-1] | 97.9 | 3.0E-63 |
| DIP_10009312 | putative eukaryotic translation initiation factor 3 subunit a protein [Eutypa lata UCREL1] | 79.5 | 0.0 |
| DIP_10009313 | putative stress-activated map kinase-interacting protein [Eutypa lata UCREL1] | 62.6 | 0.0 |
| DIP_10009314 | NA | | |
| DIP_10009315 | hypothetical protein PFICI_05552 [Pestalotiopsis fici W106-1] | 65.1 | 1.0E-148 |
| DIP_10009316 | Eukaryotic translation initiation factor 3 subunit L [Pestalotiopsis fici W106-1] | 88.1 | 0.0 |
| DIP_10009317 | putative elongation factor 2 protein [Eutypa lata UCREL1] | 75.7 | 0.0 |
| DIP_10009318 | putative thymine dioxygenase protein [Eutypa lata UCREL1] | 75.4 | 0.0 |
| DIP_10009319 | putative uracil-5-carboxylate decarboxylase protein [Eutypa lata UCREL1] | 74.9 | 0.0 |
| DIP_10009320 | NA | | |
| DIP_10009321 | hypothetical protein MYCTH_2305555 [Myceliophthora thermophila ATCC 42464] | 69.5 | 3.0E-115 |
| DIP_10009322 | hypothetical protein PFICI_05863 [Pestalotiopsis fici W106-1] | 37.1 | 1.0E-33 |
| DIP_10009323 | hypothetical protein GGTG_05621 [Gaeumannomyces graminis var. tritici R3-111a-1] | 35.3 | 9.0E-21 |
| DIP_10009324 | ste ste7 mkk protein kinase [Fusarium avenaceum] | 79.9 | 0.0 |
| DIP_10009325 | putative atp-dependent rna helicase dbp8 protein [Eutypa lata UCREL1] | 73.5 | 0.0 |
| DIP_10009326 | hypothetical protein UCREL1_11630 [Eutypa lata UCREL1] | 64.7 | 3.0E-78 |
| DIP_10009327 | putative folate-binding protein [Eutypa lata UCREL1] | 43.3 | 1.0E-96 |
| DIP_10009328 | NA | | |
| DIP_10009329 | heterokaryon incompatibility protein [Pyrenophora tritici-repentis Pt-1C-BFP] | 47.7 | 1.0E-39 |
| DIP_10009330 | NA | | |
| DIP_10009331 | hypothetical protein PV07_07502 [Cladophialophora immunda] | 39.9 | 2.0E-60 |
| DIP_10009332 | putative mitochondrial carrier protein [Eutypa lata UCREL1] | 64.5 | 0.0 |
| DIP_10009333 | hypothetical protein UCRPA7_8291 [Togninia minima UCRPA7] | 47.1 | 0.0 |
| DIP_10009334 | NA | | |
| DIP_10009335 | hypothetical protein PFICI_03363 [Pestalotiopsis fici W106-1] | 71.6 | 0.0 |
| DIP_10009336 | hypothetical protein NECHADRAFT_77499 [Nectria haematococca mpVI 77-13-4] | 73 | 0.0 |
| DIP_10009337 | hypothetical protein PFICI_03677 [Pestalotiopsis fici W106-1] | 67.2 | 0.0 |
| DIP_10009338 | hypothetical protein PFICI_01086 [Pestalotiopsis fici W106-1] | 65.3 | 0.0 |
| DIP_10009339 | putative major facilitator superfamily transporter protein [Eutypa lata UCREL1] | 69.1 | 0.0 |
| DIP_10009340 | hypothetical protein BC1G_02359 [Botrytis cinerea B05.10] | 36.8 | 3.0E-33 |
| DIP_10009341 | putative glutamate carboxypeptidase protein [Neofusicoccum parvum UCRNP2] | 56.5 | 0.0 |
| DIP_10009342 | hypothetical protein GLRG_07931 [Colletotrichum graminicola M1.001] | 73.3 | 2.0E-71 |
| DIP_10009343 | NA | | |
| DIP_10009344 | PHF5-like protein [Beauveria bassiana ARSEF 2860] | 98.2 | 1.0E-75 |
| DIP_10009345 | hypothetical protein PFICI_05573 [Pestalotiopsis fici W106-1] | 65.7 | 0.0 |
| DIP_10009346 | caf1 family ribonuclease [Ophiostoma piceae UAMH 11346] | 30.8 | 4.0E-35 |
| DIP_10009347 | putative peroxin 11c protein [Eutypa lata UCREL1] | 53.8 | 1.0E-116 |
| DIP_10009348 | hypothetical protein NECHADRAFT_17138 [Nectria haematococca mpVI 77-13-4] | 46.8 | 2.0E-116 |
| DIP_10009349 | hypothetical protein PFICI_03656 [Pestalotiopsis fici W106-1] | 39.8 | 2.0E-92 |
| DIP_10009350 | Peptidyl-prolyl cis-trans isomerase ssp-like protein [Acremonium chrysogenum ATCC 11550] | 70.9 | 2.0E-85 |
| DIP_10009351 | DNA-directed RNA polymerase [Metarhizium robertsii] | 81.4 | 1.0E-74 |
| DIP_10009352 | hypothetical protein PFICI_03653 [Pestalotiopsis fici W106-1] | 94.7 | 8.0E-87 |
| DIP_10009353 | putative histone-lysine n-methyltransferase set9 protein [Eutypa lata UCREL1] | 60.8 | 0.0 |
| DIP_10009354 | hypothetical protein PFICI_09214 [Pestalotiopsis fici W106-1] | 38.5 | 7.0E-117 |
| DIP_10009355 | NA | | |
| DIP_10009356 | putative elicitor protein [Eutypa lata UCREL1] | 68.9 | 9.0E-101 |
| DIP_10009357 | NA | | |
| DIP_10009358 | hypothetical protein CH063_12375 [Colletotrichum higginsianum] | 53 | 3.0E-120 |
| DIP_10009359 | hypothetical protein UCREL1_1803 [Eutypa lata UCREL1] | 32.7 | 3.0E-38 |
| DIP_10009360 | hypothetical protein UCREL1_1804 [Eutypa lata UCREL1] | 49.9 | 3.0E-165 |
| DIP_10009361 | microsomal epoxide hydrolase [Capronia epimyces CBS 606.96] | 54.5 | 8.0E-161 |
| DIP_10009362 | NA | | |
| DIP_10009363 | NA | | |
| DIP_10009364 | putative mediator of RNA polymerase II transcription subunit [Metarhizium anisopliae] | 54.8 | 2.0E-56 |
| DIP_10009365 | 3-isopropylmalate dehydrogenase [Pestalotiopsis fici W106-1] | 84.4 | 0.0 |
| DIP_10009366 | PHD-finger domain-containing protein [Colletotrichum fioriniae PJ7] | 33.9 | 2.0E-41 |
| DIP_10009367 | putative protein tyrosine phosphatase protein [Eutypa lata UCREL1] | 67 | 1.0E-90 |
| DIP_10009368 | hypothetical protein UCREL1_11624 [Eutypa lata UCREL1] | 65.8 | 3.0E-170 |
| DIP_10009369 | hypothetical protein PFICI_11748 [Pestalotiopsis fici W106-1] | 52.3 | 7.0E-128 |
| DIP_10009370 | hypothetical protein FOWG_14719 [Fusarium oxysporum f. sp. lycopersici MN25] | 31 | 3.0E-48 |
| DIP_10009371 | putative dna repair and recombination protein rad26 protein [Togninia minima UCRPA7] | 66.9 | 0.0 |
| DIP_10009372 | putative 40s ribosomal protein s14 protein [Eutypa lata UCREL1] | 88.5 | 1.0E-82 |
| DIP_10009373 | 60S ribosomal protein L7, partial [Metarhizium brunneum ARSEF 3297] | 82 | 1.0E-135 |
| DIP_10009374 | hypothetical protein SMAC_02107 [Sordaria macrospora k-hell] | 46.2 | 5.0E-69 |
| DIP_10009375 | hypothetical protein PFICI_05811 [Pestalotiopsis fici W106-1] | 37.9 | 1.0E-52 |
| DIP_10009376 | NA | | |
| DIP_10009377 | hypothetical protein PFICI_05810 [Pestalotiopsis fici W106-1] | 85.1 | 2.0E-145 |
| DIP_10009378 | MYB DNA-binding domain-containing protein [Gaeumannomyces graminis var. tritici R3-111a-1] | 45.4 | 1.0E-92 |
| DIP_10009379 | hypothetical protein PFICI_05809 [Pestalotiopsis fici W106-1] | 44.4 | 4.0E-129 |
| DIP_10009380 | Pre-mRNA-processing-splicing factor 8 [Pestalotiopsis fici W106-1] | 93 | 0.0 |
| DIP_10009381 | hypothetical protein TRIVIDRAFT_91260 [Trichoderma virens Gv29-8] | 88.8 | 5.0E-73 |
| DIP_10009382 | hypothetical protein UCDDA912_g08548 [Diaporthe ampelina] | 67.1 | 3.0E-76 |
| DIP_10009383 | hypothetical protein THITE_2119209 [Thielavia terrestris NRRL 8126] | 46.3 | 2.0E-154 |
| DIP_10009384 | NA | | |
| DIP_10009385 | glucose oxidase [Trichoderma reesei QM6a] | 46.4 | 3.0E-133 |
| DIP_10009386 | god1, glucose oxidase, partial sequence [Botrytis cinerea T4] | 40.6 | 7.0E-13 |
| DIP_10009387 | NA | | |
| DIP_10009388 | hypothetical protein CGLO_14616 [Colletotrichum gloeosporioides Cg-14] | 42.6 | 1.0E-55 |
| DIP_10009389 | NA | | |
| DIP_10009390 | hypothetical protein PV10_07542 [Exophiala mesophila] | 42 | 4.0E-70 |
| DIP_10009391 | hypothetical protein PFICI_14516 [Pestalotiopsis fici W106-1] | 65.9 | 0.0 |
| DIP_10009392 | NA | | |
| DIP_10009393 | NA | | |
| DIP_10009394 | hypothetical protein NECHADRAFT_38606 [Nectria haematococca mpVI 77-13-4] | 40.7 | 2.0E-81 |
| DIP_10009395 | hypothetical protein PFICI_09898 [Pestalotiopsis fici W106-1] | 58.6 | 4.0E-114 |
| DIP_10009396 | NA | | |
| DIP_10009397 | putative beta-glucosidase 1 precursor protein [Eutypa lata UCREL1] | 73 | 0.0 |
| DIP_10009398 | hypothetical protein TI39_contig56g00005 [Zymoseptoria brevis] | 47.2 | 3.0E-76 |
| DIP_10009399 | hypothetical protein CGGC5_15449 [Colletotrichum gloeosporioides Nara gc5] | 58.1 | 1.0E-161 |
| DIP_10009400 | hypothetical protein FPSE_09109 [Fusarium pseudograminearum CS3096] | 85.7 | 0.0 |
| DIP_10009401 | putative hmg box protein [Eutypa lata UCREL1] | 56.1 | 1.0E-111 |
| DIP_10009402 | sporulation protein rmd1 [Colletotrichum gloeosporioides Nara gc5] | 64.9 | 0.0 |
| DIP_10009403 | putative saccharopine dehydrogenase protein [Eutypa lata UCREL1] | 61.8 | 0.0 |
| DIP_10009404 | putative 60s ribosomal protein l43 protein [Eutypa lata UCREL1] | 91.4 | 5.0E-53 |
| DIP_10009405 | phd and ring finger domain protein [Sporothrix brasiliensis 5110] | 42.1 | 1.0E-139 |
| DIP_10009406 | hypothetical protein CFIO01_00042 [Colletotrichum fioriniae PJ7] | 63.6 | 1.0E-117 |
| DIP_10009407 | 6-hydroxy-D-nicotine oxidase [Colletotrichum fioriniae PJ7] | 66.3 | 0.0 |
| DIP_10009408 | putative cofilin tropomyosin-type actin-binding protein [Eutypa lata UCREL1] | 69.3 | 3.0E-173 |
| DIP_10009409 | putative succinate fumarate mitochondrial transporter protein [Eutypa lata UCREL1] | 82.9 | 0.0 |
| DIP_10009410 | hypothetical protein PFICI_03352 [Pestalotiopsis fici W106-1] | 64.2 | 0.0 |
| DIP_10009411 | Rhamnogalacturonate lyase A [Pestalotiopsis fici W106-1] | 51.2 | 3.0E-172 |
| DIP_10009412 | Glutamyl-tRNA(Gln) amidotransferase subunit A, mitochondrial [Talaromyces islandicus] | 52.6 | 0.0 |
| DIP_10009413 | hypothetical protein THITE_2051079 [Thielavia terrestris NRRL 8126] | 41.8 | 7.0E-87 |
| DIP_10009414 | hypothetical protein T310_3018 [Rasamsonia emersonii CBS 393.64] | 44 | 1.0E-66 |
| DIP_10009415 | hypothetical protein THITE_117302 [Thielavia terrestris NRRL 8126] | 66.9 | 0.0 |
| DIP_10009416 | hypothetical protein BAUCODRAFT_84564 [Baudoinia compniacensis UAMH 10762] | 56.4 | 8.0E-88 |
| DIP_10009417 | hypothetical protein PFICI_01073 [Pestalotiopsis fici W106-1] | 54.4 | 2.0E-103 |
| DIP_10009418 | predicted protein [Aspergillus terreus NIH2624] | 34.9 | 6.0E-32 |
| DIP_10009419 | hypothetical protein COCVIDRAFT_83616 [Bipolaris victoriae FI3] | 46.3 | 2.0E-166 |
| DIP_10009420 | putative branched-chain-amino-acid aminotransferase 1 protein [Eutypa lata UCREL1] | 77.9 | 0.0 |
| DIP_10009421 | hypothetical protein VHEMI02162 [Torrubiella hemipterigena] | 42.9 | 6.0E-58 |
| DIP_10009422 | hypothetical protein PFICI_01029 [Pestalotiopsis fici W106-1] | 55.2 | 0.0 |
| DIP_10009423 | NA | | |
| DIP_10009424 | NA | | |
| DIP_10009425 | NA | | |
| DIP_10009426 | hypothetical protein PFICI_01028 [Pestalotiopsis fici W106-1] | 78.5 | 0.0 |
| DIP_10009427 | putative glycosyltransferase family 20 protein [Eutypa lata UCREL1] | 73.7 | 0.0 |
| DIP_10009428 | hypothetical protein SMAC_02762 [Sordaria macrospora k-hell] | 34.4 | 4.0E-104 |
| DIP_10009429 | inner membrane transport protein yfaV [Colletotrichum fioriniae PJ7] | 68.2 | 0.0 |
| DIP_10009430 | hypothetical protein NECHADRAFT_42143 [Nectria haematococca mpVI 77-13-4] | 72.5 | 4.0E-178 |
| DIP_10009431 | male sterility protein [Colletotrichum higginsianum] | 60.9 | 0.0 |
| DIP_10009432 | putative short-chain dehydrogenase reductase protein [Eutypa lata UCREL1] | 57.1 | 4.0E-106 |
| DIP_10009433 | hypothetical protein ACRE_066260 [Acremonium chrysogenum ATCC 11550] | 37.5 | 6.0E-93 |
| DIP_10009434 | hypothetical protein PFICI_01023 [Pestalotiopsis fici W106-1] | 57.1 | 0.0 |
| DIP_10009435 | NA | | |
| DIP_10009436 | putative dna mismatch repair protein msh3 protein [Eutypa lata UCREL1] | 72.1 | 0.0 |
| DIP_10009437 | hypothetical protein MGG_09305 [Magnaporthe oryzae 70-15] | 30 | 9.0E-36 |
| DIP_10009438 | mPR-type GPCR [Trichoderma reesei QM6a] | 44.4 | 9.0E-96 |
| DIP_10009439 | putative ubiquinone biosynthesis monooxygenase coq6 protein [Eutypa lata UCREL1] | 65.3 | 0.0 |
| DIP_10009440 | hypothetical protein FVEG_00954 [Fusarium verticillioides 7600] | 57.5 | 1.0E-107 |
| DIP_10009441 | hypothetical protein [Podospora anserina S mat+] | 72.9 | 0.0 |
| DIP_10009442 | hypothetical protein THITE_2114058 [Thielavia terrestris NRRL 8126] | 39.3 | 1.0E-47 |
| DIP_10009443 | hypothetical protein PFICI_01101 [Pestalotiopsis fici W106-1] | 61.3 | 0.0 |
| DIP_10009444 | putative serine threonine-protein kinase nak1 protein [Eutypa lata UCREL1] | 61.8 | 0.0 |
| DIP_10009445 | NA | | |
| DIP_10009446 | hypothetical protein PFICI_01099 [Pestalotiopsis fici W106-1] | 70 | 0.0 |
| DIP_10009447 | NA | | |
| DIP_10009448 | NA | | |
| DIP_10009449 | hypothetical protein SAPIO_CDS0747 [Scedosporium apiospermum] | 44.9 | 0.0 |
| DIP_10009450 | NA | | |
| DIP_10009451 | putative ribosome biogenesis protein [Eutypa lata UCREL1] | 68.4 | 2.0E-76 |
| DIP_10009452 | putative mitochondrial ribosomal protein [Eutypa lata UCREL1] | 58 | 0.0 |
| DIP_10009453 | putative dna replication licensing factor mcm7 protein [Eutypa lata UCREL1] | 85 | 0.0 |
| DIP_10009454 | putative geranylgeranyltransferase beta subunit protein [Eutypa lata UCREL1] | 80.1 | 0.0 |
| DIP_10009455 | short chain dehydrogenase reductase [Grosmannia clavigera kw1407] | 56 | 3.0E-99 |
| DIP_10009456 | putative dna-directed rna polymerases i and iii 40 kda polypeptide protein [Eutypa lata UCREL1] | 73.8 | 0.0 |
| DIP_10009457 | NA | | |
| DIP_10009458 | NA | | |
| DIP_10009459 | NA | | |
| DIP_10009460 | hypothetical protein PFICI_01112 [Pestalotiopsis fici W106-1] | 62.5 | 0.0 |
| DIP_10009461 | putative hexokinase protein [Eutypa lata UCREL1] | 85.3 | 0.0 |
| DIP_10009462 | putative arginine n-methyltransferase 2 protein [Eutypa lata UCREL1] | 64.6 | 0.0 |
| DIP_10009463 | putative vacuolar transporter chaperone 4 protein [Eutypa lata UCREL1] | 72.2 | 0.0 |
| DIP_10009464 | hypothetical protein PFICI_01107 [Pestalotiopsis fici W106-1] | 54.9 | 0.0 |
| DIP_10009465 | NA | | |
| DIP_10009466 | NADH-ubiquinone oxidoreductase subunit A [Pestalotiopsis fici W106-1] | 89.3 | 0.0 |
| DIP_10009467 | hypothetical protein PFICI_12546 [Pestalotiopsis fici W106-1] | 47.3 | 1.0E-61 |
| DIP_10009468 | putative copii-coated vesicle protein 4 erv29 protein [Eutypa lata UCREL1] | 89.7 | 0.0 |
| DIP_10009469 | hypothetical protein PFICI_00905 [Pestalotiopsis fici W106-1] | 42.9 | 3.0E-116 |
| DIP_10009470 | NA | | |
| DIP_10009471 | major facilitator superfamily transporter [Colletotrichum gloeosporioides Cg-14] | 60.7 | 0.0 |
| DIP_10009472 | diacylglycerol pyrophosphate phosphatase [Byssochlamys spectabilis No. 5] | 47.1 | 7.0E-93 |
| DIP_10009473 | NA | | |
| DIP_10009474 | NA | | |
| DIP_10009475 | putative hexose transporter protein [Eutypa lata UCREL1] | 74.3 | 0.0 |
| DIP_10009476 | hypothetical protein PFICI_01128 [Pestalotiopsis fici W106-1] | 73.7 | 5.0E-32 |
| DIP_10009477 | hypothetical protein PFICI_01125 [Pestalotiopsis fici W106-1] | 55.5 | 6.0E-30 |
| DIP_10009478 | hypothetical protein PFICI_01124 [Pestalotiopsis fici W106-1] | 58.6 | 4.0E-134 |
| DIP_10009479 | putative plc-like phosphodiesterase protein [Eutypa lata UCREL1] | 49 | 8.0E-137 |
| DIP_10009480 | hypothetical protein UCDDA912_g07896 [Diaporthe ampelina] | 63.3 | 0.0 |
| DIP_10009481 | aldo/keto reductase [Colletotrichum graminicola M1.001] | 74.5 | 0.0 |
| DIP_10009482 | putative trna processing endoribonuclease protein [Eutypa lata UCREL1] | 57.5 | 0.0 |
| DIP_10009483 | hypothetical protein UCREL1_6874 [Eutypa lata UCREL1] | 46.2 | 2.0E-131 |
| DIP_10009484 | C6 zinc finger domain-containing protein [Verticillium alfalfae VaMs.102] | 30.2 | 1.0E-31 |
| DIP_10009485 | mitochondrial transport protein fsf1 [Pestalotiopsis fici W106-1] | 78.5 | 0.0 |
| DIP_10009486 | hypothetical protein GLRG_02174 [Colletotrichum graminicola M1.001] | 68.3 | 0.0 |
| DIP_10009487 | NA | | |
| DIP_10009488 | hypothetical protein NEUTE1DRAFT_82727 [Neurospora tetrasperma FGSC 2508] | 45.2 | 2.0E-142 |
| DIP_10009489 | NA | | |
| DIP_10009490 | putative acetyl- carboxylase protein [Eutypa lata UCREL1] | 81.9 | 0.0 |
| DIP_10009491 | small secreted protein [Macrophomina phaseolina MS6] | 64.9 | 6.0E-67 |
| DIP_10009492 | Nuclear distribution protein [Pestalotiopsis fici W106-1] | 65.5 | 0.0 |
| DIP_10009493 | NA | | |
| DIP_10009494 | putative dna repair protein rad5 protein [Eutypa lata UCREL1] | 40.1 | 2.0E-53 |
| DIP_10009495 | l-serine dehydratase [Colletotrichum gloeosporioides Nara gc5] | 66.6 | 0.0 |
| DIP_10009496 | putative arp8-like protein [Eutypa lata UCREL1] | 75.6 | 0.0 |
| DIP_10009497 | hypothetical protein PFICI_02786 [Pestalotiopsis fici W106-1] | 41.3 | 2.0E-34 |
| DIP_10009498 | hypothetical protein UCRPA7_350 [Togninia minima UCRPA7] | 75 | 0.0 |
| DIP_10009499 | hypothetical protein PFICI_07529 [Pestalotiopsis fici W106-1] | 68.9 | 2.0E-154 |
| DIP_10009500 | NA | | |
| DIP_10009501 | hypothetical protein PFICI_07537 [Pestalotiopsis fici W106-1] | 69.3 | 0.0 |
| DIP_10009502 | hypothetical protein ACRE_032140 [Acremonium chrysogenum ATCC 11550] | 45.8 | 3.0E-42 |
| DIP_10009503 | camp-dependent protein kinase catalytic subunit [Metarhizium robertsii ARSEF 23] | 68.3 | 0.0 |
| DIP_10009504 | putative vesicular integral-membrane protein vip36 protein [Eutypa lata UCREL1] | 77.5 | 0.0 |
| DIP_10009505 | hypothetical protein PFICI_11701 [Pestalotiopsis fici W106-1] | 65.4 | 0.0 |
| DIP_10009506 | NA | | |
| DIP_10009507 | NA | | |
| DIP_10009508 | hypothetical protein PDE_02267 [Penicillium oxalicum 114-2] | 62.8 | 4.0E-50 |
| DIP_10009509 | hypothetical protein PFICI_07511 [Pestalotiopsis fici W106-1] | 73.2 | 4.0E-160 |
| DIP_10009510 | putative 5-aminolevulinate synthase protein [Eutypa lata UCREL1] | 77.6 | 0.0 |
| DIP_10009511 | isochorismatase domain-containing protein [Colletotrichum orbiculare MAFF 240422] | 57.4 | 3.0E-70 |
| DIP_10009512 | putative gcn5-related n-acetyltransferase protein [Eutypa lata UCREL1] | 48.1 | 1.0E-70 |
| DIP_10009513 | putative hydroxyethylthiazole kinase protein [Eutypa lata UCREL1] | 64.3 | 0.0 |

| | | | |
|---|---|---|---|
| DIP_10009514 | hypothetical protein PFICI_02732 [Pestalotiopsis fici W106-1] | 67.3 | 0.0 |
| DIP_10009515 | U3 small nucleolar RNA-associated protein 10 [Pestalotiopsis fici W106-1] | 60.8 | 0.0 |
| DIP_10009516 | NA | | |
| DIP_10009517 | putative spo76 protein [Eutypa lata UCREL1] | 66 | 0.0 |
| DIP_10009518 | NA | | |
| DIP_10009519 | hypothetical protein CFIO01_11808 [Colletotrichum fioriniae PJ7] | 42.7 | 7.0E-54 |
| DIP_10009520 | putative dna ligase protein [Eutypa lata UCREL1] | 76.1 | 0.0 |
| DIP_10009521 | Major facilitator superfamily [Macrophomina phaseolina MS6] | 42.7 | 1.0E-176 |
| DIP_10009522 | NA | | |
| DIP_10009523 | hypothetical protein [Podospora anserina S mat+] | 30.8 | 6.0E-10 |
| DIP_10009524 | putative cutinase negative acting protein [Eutypa lata UCREL1] | 67 | 2.0E-98 |
| DIP_10009525 | putative nadh-ubiquinone oxidoreductase 12 kda subunit protein [Eutypa lata UCREL1] | 90.1 | 4.0E-65 |
| DIP_10009526 | hypothetical protein CHGG_00837 [Chaetomium globosum CBS 148.51] | 42.2 | 8.0E-47 |
| DIP_10009527 | putative transcription initiation factor tfiid subunit 7 protein [Eutypa lata UCREL1] | 53 | 2.0E-166 |
| DIP_10009528 | NA | | |
| DIP_10009529 | putative bromodomain associated domain protein [Eutypa lata UCREL1] | 58.5 | 1.0E-85 |
| DIP_10009530 | hypothetical protein PFICI_01886 [Pestalotiopsis fici W106-1] | 62.2 | 0.0 |
| DIP_10009531 | NA | | |
| DIP_10009532 | hypothetical protein PFICI_01883 [Pestalotiopsis fici W106-1] | 60.9 | 0.0 |
| DIP_10009533 | putative endoglucanase e1 protein [Eutypa lata UCREL1] | 81.8 | 0.0 |
| DIP_10009534 | putative phosphatidylethanolamine-binding protein [Eutypa lata UCREL1] | 30.3 | 3.0E-15 |
| DIP_10009535 | hypothetical protein CGLO_09326 [Colletotrichum gloeosporioides Cg-14] | 58.5 | 2.0E-125 |
| DIP_10009536 | NA | | |
| DIP_10009537 | NA | | |
| DIP_10009538 | putative cell surface spherulin 4-like protein [Eutypa lata UCREL1] | 45.7 | 6.0E-78 |
| DIP_10009539 | salicylate 1-monooxygenase SalA [Colletotrichum higginsianum] | 31.3 | 2.0E-52 |
| DIP_10009540 | hypothetical protein SAPIO_CDS5256 [Scedosporium apiospermum] | 32.2 | 8.0E-11 |
| DIP_10009541 | NA | | |
| DIP_10009542 | hypothetical protein UCREL1_1764 [Eutypa lata UCREL1] | 48.1 | 5.0E-54 |
| DIP_10009543 | NA | | |
| DIP_10009544 | unnamed protein product [Aspergillus niger] | 34.2 | 7.0E-53 |
| DIP_10009545 | putative udp-glucuronosyl udp-glucosyltransferase protein [Eutypa lata UCREL1] | 43.3 | 5.0E-111 |
| DIP_10009546 | hypothetical protein FAVG1_12170 [Fusarium avenaceum] | 61.2 | 3.0E-126 |
| DIP_10009547 | NA | | |
| DIP_10009548 | NA | | |
| DIP_10009549 | hypothetical protein COCCADRAFT_37061 [Bipolaris zeicola 26-R-13] | 32.7 | 2.0E-12 |
| DIP_10009550 | hypothetical protein PFICI_05595 [Pestalotiopsis fici W106-1] | 59.8 | 4.0E-139 |
| DIP_10009551 | putative g protein-coupled receptor protein [Eutypa lata UCREL1] | 78.4 | 0.0 |
| DIP_10009552 | hypothetical protein PFICI_04912 [Pestalotiopsis fici W106-1] | 41.4 | 2.0E-92 |
| DIP_10009553 | putative mitochondrial oxaloacetate transport protein [Eutypa lata UCREL1] | 84.8 | 0.0 |
| DIP_10009554 | hypothetical protein PFICI_04914 [Pestalotiopsis fici W106-1] | 58.3 | 2.0E-148 |
| DIP_10009555 | NA | | |
| DIP_10009556 | putative mitochondrial-processing peptidase subunit alpha protein [Eutypa lata UCREL1] | 59.8 | 2.0E-167 |
| DIP_10009557 | Mitochondrial-processing peptidase subunit alpha [Pestalotiopsis fici W106-1] | 76.5 | 0.0 |
| DIP_10009558 | hypothetical protein PFICI_05598 [Pestalotiopsis fici W106-1] | 52.1 | 0.0 |
| DIP_10009559 | hypothetical protein H109_03279 [Trichophyton interdigitale MR816] | 37.2 | 2.0E-64 |
| DIP_10009560 | putative nad-specific glutamate dehydrogenase protein [Togninia minima UCRPA7] | 78.5 | 0.0 |
| DIP_10009561 | Kynureninase 1 [Pestalotiopsis fici W106-1] | 63.8 | 0.0 |
| DIP_10009562 | hypothetical protein TRIATDRAFT_127183 [Trichoderma atroviride IMI 206040] | 73 | 2.0E-66 |
| DIP_10009563 | putative proteasome component pre6 protein [Eutypa lata UCREL1] | 94.2 | 0.0 |
| DIP_10009564 | hypothetical protein UCREL1_3119 [Eutypa lata UCREL1] | 75.4 | 0.0 |
| DIP_10009565 | putative microsomal signal peptidase 12kda subunit protein [Eutypa lata UCREL1] | 69.7 | 8.0E-50 |
| DIP_10009566 | putative ring-box protein 1 protein [Eutypa lata UCREL1] | 92.6 | 5.0E-79 |
| DIP_10009567 | hypothetical protein THITE_2119660 [Thielavia terrestris NRRL 8126] | 40.2 | 2.0E-50 |
| DIP_10009568 | hypothetical protein PFICI_12612 [Pestalotiopsis fici W106-1] | 48.1 | 2.0E-176 |
| DIP_10009569 | hypothetical protein A1O3_01084 [Capronia epimyces CBS 606.96] | 51.6 | 4.0E-80 |
| DIP_10009570 | hypothetical protein PFICI_12614 [Pestalotiopsis fici W106-1] | 70.9 | 7.0E-120 |
| DIP_10009571 | Glucose-inhibited division protein A-related protein [Macrophomina phaseolina MS6] | 53.7 | 2.0E-65 |
| DIP_10009572 | FAD/NAD(P)-binding protein [Glarea lozoyensis ATCC 20868] | 74.7 | 0.0 |
| DIP_10009573 | Vacuolar protein sorting-associated protein 45 [Pestalotiopsis fici W106-1] | 84.1 | 0.0 |
| DIP_10009574 | Maintenance of mitochondrial morphology protein 1 [Pestalotiopsis fici W106-1] | 74.4 | 0.0 |
| DIP_10009575 | hypothetical protein PFICI_12628 [Pestalotiopsis fici W106-1] | 68.9 | 1.0E-146 |
| DIP_10009576 | hypothetical protein PFICI_12627 [Pestalotiopsis fici W106-1] | 42.6 | 7.0E-109 |
| DIP_10009577 | putative mboat family protein [Eutypa lata UCREL1] | 71.9 | 0.0 |
| DIP_10009578 | putative gdp-mannose transporter protein [Eutypa lata UCREL1] | 87.3 | 0.0 |
| DIP_10009579 | hypothetical protein CTHT_0023920 [Chaetomium thermophilum var. thermophilum DSM 1495] | 71.7 | 0.0 |
| DIP_10009580 | putative -dihydroxy-3-keto-5-methylthiopentene dioxygenase protein [Eutypa lata UCREL1] | 81.3 | 9.0E-100 |
| DIP_10009581 | hypothetical protein SAPIO_CDS10315 [Scedosporium apiospermum] | 74.8 | 3.0E-66 |
| DIP_10009582 | NA | | |
| DIP_10009583 | putative cfem domain-containing protein [Eutypa lata UCREL1] | 50.5 | 5.0E-60 |
| DIP_10009584 | hypothetical protein UVI_048190 [Ustilaginoidea virens] | 57.6 | 2.0E-131 |
| DIP_10009585 | hypothetical protein CFIO01_09231 [Colletotrichum fioriniae PJ7] | 60.9 | 2.0E-132 |
| DIP_10009586 | putative vacuolar membrane protein [Eutypa lata UCREL1] | 65.9 | 0.0 |
| DIP_10009587 | hypothetical protein THITE_2124650 [Thielavia terrestris NRRL 8126] | 75.2 | 0.0 |
| DIP_10009588 | putative lipoyltransferase-like protein [Eutypa lata UCREL1] | 57.9 | 1.0E-137 |
| DIP_10009589 | putative cytochrome p450 monooxygenase [Diaporthe ampelina] | 59 | 0.0 |
| DIP_10009590 | putative vacuolar atp synthase 98 kda subunit protein [Eutypa lata UCREL1] | 88 | 0.0 |
| DIP_10009591 | predicted protein [Nectria haematococca mpVI 77-13-4] | 54.7 | 0.0 |
| DIP_10009592 | hypothetical protein PFICI_09313 [Pestalotiopsis fici W106-1] | 42.8 | 1.0E-74 |
| DIP_10009593 | hypothetical protein PFICI_09314 [Pestalotiopsis fici W106-1] | 48.2 | 5.0E-153 |
| DIP_10009594 | hypothetical protein TRIVIDRAFT_217122 [Trichoderma virens Gv29-8] | 60.2 | 7.0E-95 |
| DIP_10009595 | hypothetical protein PFICI_08931 [Pestalotiopsis fici W106-1] | 65.5 | 0.0 |
| DIP_10009596 | hypothetical protein PFICI_08930 [Pestalotiopsis fici W106-1] | 75.5 | 2.0E-149 |
| DIP_10009597 | putative rna polymerase ii subunit 3 protein [Eutypa lata UCREL1] | 80.8 | 0.0 |
| DIP_10009598 | hypothetical protein UCREL1_10662 [Eutypa lata UCREL1] | 53.7 | 2.0E-159 |
| DIP_10009599 | hypothetical protein FOVG_14411 [Fusarium oxysporum f. sp. pisi HDV247] | 42.1 | 1.0E-57 |
| DIP_10009600 | putative secretory protein [Eutypa lata UCREL1] | 63.3 | 0.0 |
| DIP_10009601 | hypothetical protein PFICI_02744 [Pestalotiopsis fici W106-1] | 69.4 | 0.0 |
| DIP_10009602 | putative isopenicillin n protein [Togninia minima UCRPA7] | 82 | 0.0 |
| DIP_10009603 | hypothetical protein PFICI_01095 [Pestalotiopsis fici W106-1] | 54.6 | 6.0E-32 |
| DIP_10009604 | related transcriptional activator [Fusarium fujikuroi IMI 58289] | 48.6 | 0.0 |
| DIP_10009605 | putative Urea amidolyase [Glarea lozoyensis 74030] | 54.1 | 4.0E-97 |
| DIP_10009606 | NA | | |
| DIP_10009607 | NA | | |
| DIP_10009608 | hypothetical protein NECHADRAFT_69107 [Nectria haematococca mpVI 77-13-4] | 72.8 | 1.0E-112 |
| DIP_10009609 | hypothetical protein PFICI_14617 [Pestalotiopsis fici W106-1] | 57.3 | 3.0E-157 |
| DIP_10009610 | hypothetical protein PFICI_05312 [Pestalotiopsis fici W106-1] | 59.3 | 0.0 |
| DIP_10009611 | hypothetical protein PFICI_04763 [Pestalotiopsis fici W106-1] | 69.2 | 0.0 |
| DIP_10009612 | putative gyf domain-containing protein [Eutypa lata UCREL1] | 63.4 | 0.0 |
| DIP_10009613 | hypothetical protein PFICI_05314 [Pestalotiopsis fici W106-1] | 54.2 | 0.0 |
| DIP_10009614 | inosine-uridine preferring nucleoside hydrolase [Metarhizium robertsii ARSEF 23] | 46.2 | 6.0E-124 |
| DIP_10009615 | hypothetical protein PFICI_12799 [Pestalotiopsis fici W106-1] | 52.5 | 0.0 |
| DIP_10009616 | hypothetical protein PFICI_05244 [Pestalotiopsis fici W106-1] | 63.2 | 3.0E-135 |
| DIP_10009617 | vacuolar segregation protein pep7 [Metarhizium anisopliae] | 54.4 | 0.0 |
| DIP_10009618 | hypothetical protein PFICI_14605 [Pestalotiopsis fici W106-1] | 33.4 | 3.0E-45 |
| DIP_10009619 | hypothetical protein NCU16722 [Neurospora crassa OR74A] | 35.8 | 2.0E-20 |
| DIP_10009620 | putative peptide synthetase protein [Eutypa lata UCREL1] | 62.3 | 0.0 |
| DIP_10009621 | hypothetical protein AOR_1_1334024 [Aspergillus oryzae RIB40] | 30.1 | 4.0E-09 |
| DIP_10009622 | carbohydrate esterase family 1 protein [Myceliophthora thermophila ATCC 42464] | 52 | 7.0E-108 |
| DIP_10009623 | hypothetical protein UCREL1_7110 [Eutypa lata UCREL1] | 47.9 | 2.0E-85 |
| DIP_10009624 | putative glycosyltransferase family 4 protein [Eutypa lata UCREL1] | 52.4 | 0.0 |
| DIP_10009625 | putative coenzyme a protein [Eutypa lata UCREL1] | 70.4 | 0.0 |
| DIP_10009626 | putative mitochondrial protein [Eutypa lata UCREL1] | 70.5 | 1.0E-142 |
| DIP_10009627 | hypothetical protein PFICI_14602 [Pestalotiopsis fici W106-1] | 73.7 | 7.0E-167 |
| DIP_10009628 | hypothetical protein PFICI_12824 [Pestalotiopsis fici W106-1] | 57.6 | 0.0 |
| DIP_10009629 | hypothetical protein PFICI_12823 [Pestalotiopsis fici W106-1] | 72.3 | 0.0 |
| DIP_10009630 | putative laccase-1 precursor protein [Eutypa lata UCREL1] | 69.9 | 0.0 |
| DIP_10009631 | putative cytochrome p450 protein [Eutypa lata UCREL1] | 42.5 | 3.0E-31 |
| DIP_10009632 | hypothetical protein PFICI_14609 [Pestalotiopsis fici W106-1] | 78.2 | 4.0E-102 |
| DIP_10009633 | hypothetical protein PFICI_14608 [Pestalotiopsis fici W106-1] | 33.9 | 3.0E-59 |
| DIP_10009634 | putative start control protein cdc10 protein [Eutypa lata UCREL1] | 56.8 | 0.0 |
| DIP_10009635 | hypothetical protein V499_03507 [Pseudogymnoascus pannorum VKM F-103] | 56.2 | 0.0 |
| DIP_10009636 | hypothetical protein TCE0_013f01073 [Talaromyces cellulolyticus] | 30.1 | 2.0E-08 |
| DIP_10009637 | NA | | |
| DIP_10009638 | NA | | |
| DIP_10009639 | hypothetical protein CH063_06317 [Colletotrichum higginsianum] | 37.8 | 7.0E-51 |
| DIP_10009640 | putative dna repair protein rad26 protein [Eutypa lata UCREL1] | 54.4 | 0.0 |
| DIP_10009641 | putative phosphatidate cytidylyltransferase protein [Eutypa lata UCREL1] | 76 | 0.0 |
| DIP_10009642 | hypothetical protein PFICI_05246 [Pestalotiopsis fici W106-1] | 32.2 | 3.0E-43 |
| DIP_10009643 | NA | | |
| DIP_10009644 | hypothetical protein FOMG_10740 [Fusarium oxysporum f. sp. melonis 26406] | 58.8 | 1.0E-142 |
| DIP_10009645 | hypothetical protein T310_1538 [Rasamsonia emersonii CBS 393.64] | 65.2 | 0.0 |
| DIP_10009646 | hypothetical protein NEUTE1DRAFT_120620 [Neurospora tetrasperma FGSC 2508] | 53.7 | 6.0E-175 |
| DIP_10009647 | hypothetical protein PFICI_05305 [Pestalotiopsis fici W106-1] | 32.9 | 6.0E-31 |
| DIP_10009648 | 2og-fe oxygenase superfamily protein [Colletotrichum gloeosporioides Nara gc5] | 82.6 | 0.0 |
| DIP_10009649 | major facilitator superfamily transporter [Colletotrichum graminicola M1.001] | 73.7 | 0.0 |
| DIP_10009650 | hypothetical protein PFICI_09983 [Pestalotiopsis fici W106-1] | 52.7 | 4.0E-84 |
| DIP_10009651 | NA | | |
| DIP_10009652 | hypothetical protein PFICI_02078 [Pestalotiopsis fici W106-1] | 56.3 | 7.0E-128 |
| DIP_10009653 | putative glycoside hydrolase family 5 protein [Eutypa lata UCREL1] | 69.3 | 2.0E-173 |
| DIP_10009654 | hypothetical protein PFICI_05502 [Pestalotiopsis fici W106-1] | 67.5 | 0.0 |
| DIP_10009655 | NA | | |
| DIP_10009656 | calcium-translocating p-type atpase [Colletotrichum gloeosporioides Nara gc5] | 67.1 | 0.0 |
| DIP_10009657 | putative mfs quinate protein [Eutypa lata UCREL1] | 76.4 | 0.0 |
| DIP_10009658 | NA | | |
| DIP_10009659 | hypothetical protein OIDMADRAFT_60751 [Oidiodendron maius Zn] | 44.8 | 0.0 |
| DIP_10009660 | hypothetical protein UCREL1_4234 [Eutypa lata UCREL1] | 45.3 | 8.0E-23 |
| DIP_10009661 | hypothetical protein PFICI_05508 [Pestalotiopsis fici W106-1] | 60.3 | 8.0E-104 |
| DIP_10009662 | NA | | |
| DIP_10009663 | putative mitochondrial intermembrane space import and assembly protein 40 protein [Eutypa lata UCREL1] | 69.8 | 2.0E-64 |
| DIP_10009664 | l-amino acid oxidase [Colletotrichum gloeosporioides Nara gc5] | 64 | 0.0 |
| DIP_10009665 | hypothetical protein FAVG1_10859 [Fusarium avenaceum] | 59.9 | 5.0E-56 |
| DIP_10009666 | dna repair and transcription factor [Ophiostoma piceae UAMH 11346] | 36.7 | 4.0E-32 |
| DIP_10009667 | hypothetical protein AOL_s00043g185 [Arthrobotrys oligospora ATCC 24927] | 30.5 | 2.0E-09 |
| DIP_10009668 | NA | | |
| DIP_10009669 | cytochrome p450 [Colletotrichum orbiculare MAFF 240422] | 53.8 | 0.0 |
| DIP_10009670 | putative fad binding domain-containing protein [Eutypa lata UCREL1] | 41.3 | 9.0E-111 |
| DIP_10009671 | hypothetical protein OIDMADRAFT_20084 [Oidiodendron maius Zn] | 33 | 1.0E-24 |
| DIP_10009672 | putative poly(3-hydroxybutyrate) depolymerase protein [Eutypa lata UCREL1] | 44.3 | 1.0E-79 |
| DIP_10009673 | glutamine--tRNA ligase [Pestalotiopsis fici W106-1] | 79.5 | 0.0 |
| DIP_10009674 | hypothetical protein UCREL1_7019 [Eutypa lata UCREL1] | 32 | 3.0E-43 |
| DIP_10009675 | hypothetical protein PFICI_08840 [Pestalotiopsis fici W106-1] | 52 | 1.0E-79 |
| DIP_10009676 | NA | | |
| DIP_10009677 | Protein kinase-like domain protein, partial [Metarhizium majus ARSEF 297] | 57.3 | 0.0 |
| DIP_10009678 | hypothetical protein VDAG_02753 [Verticillium dahliae VdLs.17] | 65.4 | 2.0E-21 |
| DIP_10009679 | putative glutathione synthetase protein [Eutypa lata UCREL1] | 43.5 | 3.0E-139 |
| DIP_10009680 | NA | | |
| DIP_10009681 | Pc21g22090 [Penicillium rubens Wisconsin 54-1255] | 75.7 | 4.0E-179 |
| DIP_10009682 | hypothetical protein HMPREF1624_02443 [Sporothrix schenckii ATCC 58251] | 74.9 | 0.0 |
| DIP_10009683 | hypothetical protein PFICI_05498 [Pestalotiopsis fici W106-1] | 56.4 | 0.0 |
| DIP_10009684 | Malate dehydrogenase [Pestalotiopsis fici W106-1] | 89.4 | 0.0 |
| DIP_10009685 | NA | | |
| DIP_10009686 | hypothetical protein PISL3812_03853 [Talaromyces islandicus] | 53.2 | 0.0 |
| DIP_10009687 | hypothetical protein PFICI_05303 [Pestalotiopsis fici W106-1] | 69.9 | 0.0 |
| DIP_10009688 | hypothetical protein V493_02047 [Pseudogymnoascus pannorum VKM F-4281 (FW-2241)] | 40.3 | 1.0E-34 |
| DIP_10009689 | hypothetical protein SCLCIDRAFT_32114 [Scleroderma citrinum Foug A] | 50 | 3.0E-21 |
| DIP_10009690 | biotin synthase [Cyphellophora europaea CBS 101466] | 77.8 | 0.0 |
| DIP_10009691 | hypothetical protein PFICI_05463 [Pestalotiopsis fici W106-1] | 49.9 | 0.0 |
| DIP_10009692 | 8-amino-7-oxononanoate synthase [Scedosporium apiospermum] | 48.1 | 2.0E-135 |
| DIP_10009693 | NA | | |
| DIP_10009694 | sugar transporter [Aspergillus oryzae RIB40] | 40.8 | 1.0E-138 |
| DIP_10009695 | Putative hydrolase Mb2248c [Talaromyces islandicus] | 59.8 | 0.0 |
| DIP_10009696 | putative carbonic anhydrase protein [Togninia minima UCRPA7] | 84.7 | 3.0E-143 |
| DIP_10009697 | NA | | |
| DIP_10009698 | hypothetical protein PFICI_12831 [Pestalotiopsis fici W106-1] | 80.8 | 0.0 |
| DIP_10009699 | NA | | |
| DIP_10009700 | putative mitochondrial intermembrane space protein mia40 protein [Eutypa lata UCREL1] | 58.2 | 6.0E-93 |
| DIP_10009701 | hypothetical protein M436DRAFT_16579, partial [Aureobasidium namibiae CBS 147.97] | 48.7 | 2.0E-28 |
| DIP_10009702 | hypothetical protein CHGG_05177 [Chaetomium globosum CBS 148.51] | 39.6 | 4.0E-58 |
| DIP_10009703 | putative transporter bos1 protein [Eutypa lata UCREL1] | 71 | 8.0E-126 |
| DIP_10009704 | hypothetical protein DOTSEDRAFT_138875 [Dothistroma septosporum NZE10] | 42.3 | 7.0E-33 |
| DIP_10009705 | hypothetical protein DOTSEDRAFT_138875 [Dothistroma septosporum NZE10] | 34.6 | 1.0E-11 |
| DIP_10009706 | NA | | |
| DIP_10009707 | hypothetical protein HMPREF1541_00778 [Cyphellophora europaea CBS 101466] | 68.6 | 6.0E-161 |
| DIP_10009708 | putative nad-dependent 15-hydroxyprostaglandin dehydrogenase protein [Eutypa lata UCREL1] | 58.1 | 6.0E-131 |
| DIP_10009709 | hypothetical protein PFICI_04789 [Pestalotiopsis fici W106-1] | 48.1 | 0.0 |
| DIP_10009710 | NA | | |
| DIP_10009711 | hypothetical protein G647_01075 [Cladophialophora carrionii CBS 160.54] | 44.3 | 2.0E-65 |
| DIP_10009712 | hypothetical protein MYCFIDRAFT_30146 [Pseudocercospora fijiensis CIRAD86] | 57.1 | 0.0 |
| DIP_10009713 | NA | | |
| DIP_10009714 | candidapepsin-8 [Magnaporthe oryzae Y34] | 45.6 | 1.0E-117 |
| DIP_10009715 | hypothetical protein PFICI_09124 [Pestalotiopsis fici W106-1] | 51.9 | 6.0E-145 |
| DIP_10009716 | NA | | |
| DIP_10009717 | NA | | |
| DIP_10009718 | Elongation factor 3 [Pestalotiopsis fici W106-1] | 84.4 | 0.0 |
| DIP_10009719 | NA | | |
| DIP_10009720 | Sir2 family protein [Cordyceps militaris CM01] | 37.8 | 9.0E-124 |
| DIP_10009721 | major facilitator superfamily transporter [Colletotrichum fioriniae PJ7] | 60.4 | 0.0 |
| DIP_10009722 | NA | | |
| DIP_10009723 | MACRO domain-containing protein 1 [Metarhizium guizhouense ARSEF 977] | 51.9 | 1.0E-69 |
| DIP_10009724 | hypothetical protein PFICI_12776 [Pestalotiopsis fici W106-1] | 77.6 | 0.0 |
| DIP_10009725 | hypothetical protein UCREL1_5802 [Eutypa lata UCREL1] | 38.7 | 5.0E-75 |
| DIP_10009726 | NA | | |
| DIP_10009727 | putative potassium ion channel yvc1 protein [Eutypa lata UCREL1] | 70.9 | 0.0 |
| DIP_10009728 | hypothetical protein CHGG_05075 [Chaetomium globosum CBS 148.51] | 63.4 | 0.0 |
| DIP_10009729 | putative dna-directed rna polymerase iii subunit protein [Eutypa lata UCREL1] | 48.1 | 4.0E-66 |
| DIP_10009730 | NA | | |
| DIP_10009731 | hypothetical protein UCREL1_5757 [Eutypa lata UCREL1] | 54.1 | 3.0E-101 |
| DIP_10009732 | putative guanyl-nucleotide exchange factor protein [Eutypa lata UCREL1] | 80.2 | 0.0 |
| DIP_10009733 | putative pyruvate dehydrogenase complex dihydrolipoamide acetyltransferase protein [Eutypa lata UCREL1] | 77.7 | 0.0 |
| DIP_10009734 | hypothetical protein MAC_05134 [Metarhizium acridum CQMa 102] | 73.4 | 0.0 |
| DIP_10009735 | putative c6 transcription protein [Eutypa lata UCREL1] | 40.7 | 4.0E-151 |
| DIP_10009736 | inosine monophosphate dehydrogenase [Sphaerulina musiva SO2202] | 64.4 | 2.0E-141 |
| DIP_10009737 | hypothetical protein NECHADRAFT_33373 [Nectria haematococca mpVI 77-13-4] | 50.7 | 2.0E-134 |
| DIP_10009738 | hypothetical protein, partial [Ktedonobacter racemifer] | 31 | 4.0E-07 |
| DIP_10009739 | hypothetical protein FVEG_12428 [Fusarium verticillioides 7600] | 61 | 5.0E-34 |
| DIP_10009740 | hypothetical protein PFICI_05218 [Pestalotiopsis fici W106-1] | 83.7 | 0.0 |
| DIP_10009741 | hypothetical protein CTHT_0039510 [Chaetomium thermophilum var. thermophilum DSM 1495] | 37.6 | 4.0E-15 |
| DIP_10009742 | putative proteasome maturation factor protein [Eutypa lata UCREL1] | 66.4 | 5.0E-65 |
| DIP_10009743 | hypothetical protein CH063_09363 [Colletotrichum higginsianum] | 46.9 | 3.0E-110 |
| DIP_10009744 | hypothetical protein PFICI_05239 [Pestalotiopsis fici W106-1] | 65 | 0.0 |
| DIP_10009745 | hypothetical protein COCSADRAFT_223755 [Bipolaris sorokiniana ND90Pr] | 58.1 | 3.0E-147 |
| DIP_10009746 | NA | | |
| DIP_10009747 | putative aldehyde reductase protein [Eutypa lata UCREL1] | 59.5 | 4.0E-134 |
| DIP_10009748 | putative trna nucleotidyltransferase protein [Eutypa lata UCREL1] | 71.8 | 2.0E-140 |
| DIP_10009749 | hypothetical protein PFICI_11672 [Pestalotiopsis fici W106-1] | 76.9 | 0.0 |
| DIP_10009750 | putative trna nucleotidyltransferase protein [Eutypa lata UCREL1] | 39.5 | 4.0E-64 |
| DIP_10009751 | hypothetical protein PFICI_05216 [Pestalotiopsis fici W106-1] | 65.9 | 0.0 |
| DIP_10009752 | Pc18g02620 [Penicillium rubens Wisconsin 54-1255] | 71.1 | 0.0 |
| DIP_10009753 | putative transmembrane protein [Eutypa lata UCREL1] | 76.5 | 5.0E-98 |
| DIP_10009754 | hypothetical protein PFICI_09021 [Pestalotiopsis fici W106-1] | 35.4 | 7.0E-75 |
| DIP_10009755 | putative glyoxalase family protein [Eutypa lata UCREL1] | 69.9 | 9.0E-170 |
| DIP_10009756 | hypothetical protein PFICI_05465 [Pestalotiopsis fici W106-1] | 67 | 2.0E-92 |
| DIP_10009757 | o-methyltransferase [Colletotrichum gloeosporioides Nara gc5] | 56.4 | 4.0E-74 |
| DIP_10009758 | putative sec14 cytosolic factor protein [Eutypa lata UCREL1] | 69.2 | 0.0 |
| DIP_10009759 | putative flavin-binding monooxygenase-like protein [Eutypa lata UCREL1] | 64.6 | 0.0 |
| DIP_10009760 | hypothetical protein PFICI_05037 [Pestalotiopsis fici W106-1] | 57.8 | 2.0E-115 |
| DIP_10009761 | putative transcription initiation factor iif subunit alpha protein [Eutypa lata UCREL1] | 64.7 | 1.0E-180 |
| DIP_10009762 | putative caffeine-induced death protein 1 [Diaporthe ampelina] | 59.8 | 3.0E-142 |
| DIP_10009763 | hypothetical protein VDAG_04926 [Verticillium dahliae VdLs.17] | 39 | 3.0E-10 |
| DIP_10009764 | gpr1 fun34 -class plasma membrane protein [Colletotrichum orbiculare MAFF 240422] | 60.5 | 6.0E-102 |
| DIP_10009765 | hypothetical protein SBOR_1086 [Sclerotinia borealis F-4157] | 44.9 | 2.0E-67 |
| DIP_10009766 | vacuolar ATP synthase subunit D [Metarhizium anisopliae] | 87.1 | 4.0E-163 |
| DIP_10009767 | putative pre-mrna-splicing factor syf1 protein [Eutypa lata UCREL1] | 86.5 | 0.0 |
| DIP_10009768 | putative parasitic phase-specific protein psp-1 protein [Eutypa lata UCREL1] | 34.4 | 2.0E-46 |
| DIP_10009769 | NA | | |
| DIP_10009770 | NA | | |
| DIP_10009771 | hypothetical protein MAC_06163 [Metarhizium acridum CQMa 102] | 31.8 | 2.0E-41 |
| DIP_10009772 | putative rna polymerase i-specific transcription initiation factor rrn3 protein [Eutypa lata UCREL1] | 70.3 | 0.0 |
| DIP_10009773 | putative nuclear transcription factor y subunit b-3 protein [Eutypa lata UCREL1] | 75.8 | 5.0E-99 |
| DIP_10009774 | hypothetical protein THITE_2121748 [Thielavia terrestris NRRL 8126] | 81.2 | 3.0E-64 |
| DIP_10009775 | hypothetical protein PFICI_02072 [Pestalotiopsis fici W106-1] | 68.5 | 4.0E-41 |
| DIP_10009776 | putative upf0136 domain protein [Eutypa lata UCREL1] | 78.9 | 8.0E-53 |
| DIP_10009777 | oligosaccharyl transferase [Trichoderma atroviride IMI 206040] | 67.7 | 5.0E-65 |
| DIP_10009778 | hypothetical protein CHGG_00947 [Chaetomium globosum CBS 148.51] | 63.6 | 1.0E-146 |
| DIP_10009779 | putative rna methylase family protein [Eutypa lata UCREL1] | 45 | 2.0E-59 |
| DIP_10009780 | hypothetical protein PFICI_01467 [Pestalotiopsis fici W106-1] | 67.1 | 2.0E-119 |
| DIP_10009781 | putative rna binding protein [Eutypa lata UCREL1] | 77.6 | 1.0E-95 |
| DIP_10009782 | hypothetical protein GGTG_09603 [Gaeumannomyces graminis var. tritici R3-111a-1] | 58.4 | 2.0E-28 |
| DIP_10009783 | hypothetical protein PFICI_01854 [Pestalotiopsis fici W106-1] | 65.4 | 0.0 |
| DIP_10009784 | hypothetical protein PFICI_01856 [Pestalotiopsis fici W106-1] | 65.3 | 0.0 |
| DIP_10009785 | putative sphingomyelinase family protein [Eutypa lata UCREL1] | 55.2 | 0.0 |
| DIP_10009786 | putative tyrosyl-trna synthetase protein [Eutypa lata UCREL1] | 80.4 | 0.0 |
| DIP_10009787 | hypothetical protein FGSG_12256 [Fusarium graminearum PH-1] | 30.6 | 5.0E-09 |
| DIP_10009788 | related to NAD(P)H-dependent oxidoreductase [Fusarium fujikuroi IMI 58289] | 64.5 | 2.0E-142 |
| DIP_10009789 | hypothetical protein PFICI_14580 [Pestalotiopsis fici W106-1] | 77.2 | 0.0 |
| DIP_10009790 | putative ribonuclease p protein subunit p29 protein [Eutypa lata UCREL1] | 55.4 | 8.0E-88 |
| DIP_10009791 | hypothetical protein PFICI_14579 [Pestalotiopsis fici W106-1] | 50.7 | 1.0E-154 |
| DIP_10009792 | putative c6 zinc finger domain-containing protein [Eutypa lata UCREL1] | 47 | 3.0E-133 |
| DIP_10009793 | hypothetical protein PFICI_14577 [Pestalotiopsis fici W106-1] | 71.3 | 3.0E-150 |
| DIP_10009794 | hypothetical protein MYCTH_2299554 [Myceliophthora thermophila ATCC 42464] | 61.9 | 4.0E-108 |
| DIP_10009795 | hypothetical protein TRIVIDRAFT_180756 [Trichoderma virens Gv29-8] | 34 | 3.0E-64 |
| DIP_10009796 | putative snf7 family protein [Eutypa lata UCREL1] | 56.4 | 6.0E-169 |
| DIP_10009797 | putative dna polymerase subunit cdc27 protein [Eutypa lata UCREL1] | 50.9 | 1.0E-73 |
| DIP_10009798 | hypothetical protein UCREL1_4139 [Eutypa lata UCREL1] | 78 | 0.0 |
| DIP_10009799 | putative ferulic acid esterase protein [Eutypa lata UCREL1] | 51.4 | 3.0E-110 |
| DIP_10009800 | hypothetical protein UCREL1_4131 [Eutypa lata UCREL1] | 70.7 | 0.0 |
| DIP_10009801 | putative serine threonine-protein phosphatase pp2a catalytic subunit protein [Eutypa lata UCREL1] | 96.3 | 0.0 |
| DIP_10009802 | putative RHO protein GDP dissociation inhibitor [Colletotrichum sublineola] | 88.6 | 6.0E-126 |
| DIP_10009803 | putative -n domain-containing protein [Eutypa lata UCREL1] | 70 | 0.0 |
| DIP_10009804 | hypothetical protein PFICI_05450 [Pestalotiopsis fici W106-1] | 67.3 | 2.0E-166 |
| DIP_10009805 | putative arm repeat-containing protein [Diaporthe ampelina] | 55.5 | 0.0 |
| DIP_10009806 | hypothetical protein CGGC5_12094 [Colletotrichum gloeosporioides Nara gc5] | 31 | 3.0E-35 |
| DIP_10009807 | Nuf2 family protein [Colletotrichum gloeosporioides Cg-14] | 76.1 | 0.0 |
| DIP_10009808 | hypothetical protein PDE_04445 [Penicillium oxalicum 114-2] | 46.8 | 2.0E-67 |
| DIP_10009809 | NA | | |
| DIP_10009810 | NA | | |
| DIP_10009811 | hypothetical protein OIDMADRAFT_125386 [Oidiodendron maius Zn] | 36.8 | 1.0E-34 |
| DIP_10009812 | hypothetical protein PFICI_05455 [Pestalotiopsis fici W106-1] | 67 | 1.0E-50 |
| DIP_10009813 | hypothetical protein OIDMADRAFT_183570 [Oidiodendron maius Zn] | 56.5 | 1.0E-89 |
| DIP_10009814 | arsenite efflux transporter ArsB-like, putative [Aspergillus clavatus NRRL 1] | 62.4 | 0.0 |
| DIP_10009815 | hypothetical protein UCREL1_10878 [Eutypa lata UCREL1] | 55.2 | 1.0E-69 |
| DIP_10009816 | NA | | |
| DIP_10009817 | hypothetical protein PFICI_04959 [Pestalotiopsis fici W106-1] | 57.5 | 0.0 |
| DIP_10009818 | hypothetical protein PFICI_04960 [Pestalotiopsis fici W106-1] | 52.5 | 7.0E-44 |
| DIP_10009819 | putative fungal specific transcription factor domain-containing protein [Eutypa lata UCREL1] | 56.4 | 0.0 |
| DIP_10009820 | major facilitator superfamily transporter [Colletotrichum graminicola M1.001] | 53.3 | 0.0 |
| DIP_10009821 | amidase-like protein [Beauveria bassiana ARSEF 2860] | 49.8 | 2.0E-78 |
| DIP_10009822 | NA | | |
| DIP_10009823 | NA | | |
| DIP_10009824 | putative transglutaminase-like superfamily protein [Eutypa lata UCREL1] | 63.6 | 5.0E-169 |
| DIP_10009825 | putative arrestin protein [Eutypa lata UCREL1] | 46.8 | 3.0E-144 |
| DIP_10009826 | zip zinc transporter [Colletotrichum orbiculare MAFF 240422] | 80.4 | 0.0 |
| DIP_10009827 | NA | | |
| DIP_10009828 | putative dna-directed rna polymerase iii complex subunit rpc37 protein [Eutypa lata UCREL1] | 56.1 | 6.0E-132 |
| DIP_10009829 | putative glycolipid 2-alpha-mannosyltransferase protein [Eutypa lata UCREL1] | 51.4 | 3.0E-139 |
| DIP_10009830 | hypothetical protein PFICI_01990 [Pestalotiopsis fici W106-1] | 74.9 | 0.0 |
| DIP_10009831 | hypothetical protein PFICI_01991 [Pestalotiopsis fici W106-1] | 54.2 | 1.0E-132 |
| DIP_10009832 | putative lyar-type c2hc zinc finger protein [Eutypa lata UCREL1] | 52.4 | 3.0E-137 |
| DIP_10009833 | bem46 family protein [Colletotrichum orbiculare MAFF 240422] | 73.1 | 0.0 |
| DIP_10009834 | hypothetical protein PFICI_05000 [Pestalotiopsis fici W106-1] | 42.5 | 3.0E-80 |
| DIP_10009835 | NA | | |
| DIP_10009836 | hypothetical protein UCREL1_9793 [Eutypa lata UCREL1] | 81.9 | 4.0E-49 |
| DIP_10009837 | hypothetical protein PFICI_01988 [Pestalotiopsis fici W106-1] | 68.3 | 1.0E-89 |
| DIP_10009838 | putative topoisomerase ii protein [Eutypa lata UCREL1] | 61 | 0.0 |
| DIP_10009839 | putative alpha- mannosyltransferase ktr1 protein [Eutypa lata UCREL1] | 76.9 | 0.0 |
| DIP_10009840 | hypothetical protein MYCTH_2096341 [Myceliophthora thermophila ATCC 42464] | 77.7 | 3.0E-89 |
| DIP_10009841 | Short/branched chain specific acyl-CoA dehydrogenase [Pestalotiopsis fici W106-1] | 78.4 | 0.0 |
| DIP_10009842 | hypothetical protein PFICI_04715 [Pestalotiopsis fici W106-1] | 45.8 | 0.0 |
| DIP_10009843 | hypothetical protein FOZG_01096 [Fusarium oxysporum Fo47] | 35.2 | 5.0E-14 |
| DIP_10009844 | hypothetical protein PFICI_00964 [Pestalotiopsis fici W106-1] | 59.3 | 0.0 |
| DIP_10009845 | putative mfs transporter protein [Neofusicoccum parvum UCRNP2] | 60.7 | 0.0 |
| DIP_10009846 | putative dihydrodipicolinate synthetase family protein [Diplodia seriata] | 68.2 | 1.0E-154 |
| DIP_10009847 | Putative Enolase C-terminal [Torrubiella hemipterigena] | 73.1 | 0.0 |
| DIP_10009848 | NA | | |
| DIP_10009849 | hypothetical protein PFICI_04677 [Pestalotiopsis fici W106-1] | 59.2 | 0.0 |
| DIP_10009850 | Ferrochelatase [Pestalotiopsis fici W106-1] | 75.2 | 0.0 |
| DIP_10009851 | putative ku70 protein [Eutypa lata UCREL1] | 62.1 | 0.0 |
| DIP_10009852 | hypothetical protein PFICI_04681 [Pestalotiopsis fici W106-1] | 36.4 | 4.0E-107 |
| DIP_10009853 | putative guanine deaminase protein [Eutypa lata UCREL1] | 56 | 0.0 |
| DIP_10009854 | putative lipase 2 protein [Eutypa lata UCREL1] | 42.7 | 2.0E-64 |
| DIP_10009855 | Uncharacterized protein SAPIO_CDS2556 [Scedosporium apiospermum] | 42.9 | 2.0E-114 |
| DIP_10009856 | putative fatty acid synthase beta subunit dehydratase protein [Eutypa lata UCREL1] | 86.7 | 0.0 |
| DIP_10009857 | putative fatty acid synthase subunit alpha protein [Eutypa lata UCREL1] | 83.2 | 0.0 |
| DIP_10009858 | NA | | |
| DIP_10009859 | NA | | |
| DIP_10009860 | hypothetical protein PFICI_10024 [Pestalotiopsis fici W106-1] | 76.9 | 3.0E-156 |
| DIP_10009861 | hypothetical protein PFICI_04963 [Pestalotiopsis fici W106-1] | 51.4 | 5.0E-142 |
| DIP_10009862 | NA | | |
| DIP_10009863 | putative dihydroxy-acid dehydratase protein [Eutypa lata UCREL1] | 64.7 | 9.0E-131 |
| DIP_10009864 | putative pq loop repeat protein [Eutypa lata UCREL1] | 71.8 | 4.0E-117 |
| DIP_10009865 | trihydroxynaphthalene reductase, partial [Daldinia eschscholtzii] | 87.1 | 2.0E-179 |
| DIP_10009866 | putative chitinase 18-11 protein [Eutypa lata UCREL1] | 69.9 | 4.0E-180 |
| DIP_10009867 | hypothetical protein PFICI_14543 [Pestalotiopsis fici W106-1] | 34.9 | 8.0E-105 |
| DIP_10009868 | short-chain dehydrogenase, putative [Cordyceps militaris CM01] | 56.9 | 8.0E-89 |
| DIP_10009869 | putative urea active transporter [Diaporthe ampelina] | 72.6 | 0.0 |
| DIP_10009870 | NA | | |
| DIP_10009871 | putative c6 zinc finger domain containing protein [Eutypa lata UCREL1] | 63.3 | 6.0E-34 |
| DIP_10009872 | hypothetical protein PFICI_02066 [Pestalotiopsis fici W106-1] | 65 | 0.0 |
| DIP_10009873 | putative vacuolar fusion protein mon1 protein [Eutypa lata UCREL1] | 72.2 | 0.0 |
| DIP_10009874 | NA | | |
| DIP_10009875 | NA | | |
| DIP_10009876 | putative protein disulfide isomerase protein [Eutypa lata UCREL1] | 77.6 | 0.0 |
| DIP_10009877 | putative aromatic amino acid aminotransferase protein [Eutypa lata UCREL1] | 69.9 | 0.0 |
| DIP_10009878 | hypothetical protein PFICI_00042 [Pestalotiopsis fici W106-1] | 71.2 | 0.0 |
| DIP_10009879 | putative glycoside hydrolase family 20 protein [Eutypa lata UCREL1] | 63.8 | 0.0 |

| | | | |
|---|---|---|---|
| DIP_10009880 | hypothetical protein G647_01580 [Cladophialophora carrionii CBS 160.54] | 70.9 | 0.0 |
| DIP_10009881 | putative nuclear polyadenylated rna-binding protein nab2 protein [Eutypa lata UCREL1] | 62.6 | 0.0 |
| DIP_10009882 | putative protoporphyrinogen oxidase protein [Eutypa lata UCREL1] | 42.5 | 9.0E-159 |
| DIP_10009883 | hypothetical protein PFICI_01795 [Pestalotiopsis fici W106-1] | 50.6 | 1.0E-93 |
| DIP_10009884 | putative rna exonuclease protein [Eutypa lata UCREL1] | 69.6 | 0.0 |
| DIP_10009885 | hypothetical protein FOC4_g10000207 [Fusarium oxysporum f. sp. cubense race 4] | 46 | 8.0E-74 |
| DIP_10009886 | hypothetical protein PFICI_01788 [Pestalotiopsis fici W106-1] | 73.5 | 0.0 |
| DIP_10009887 | nucleic acid-binding protein [Ophiostoma piceae UAMH 11346] | 44.5 | 6.0E-58 |
| DIP_10009888 | hypothetical protein THITE_2112332 [Thielavia terrestris NRRL 8126] | 55.6 | 6.0E-88 |
| DIP_10009889 | putative gnat family protein [Eutypa lata UCREL1] | 64.6 | 3.0E-86 |
| DIP_10009890 | hypothetical protein PFICI_01791 [Pestalotiopsis fici W106-1] | 39.1 | 2.0E-117 |
| DIP_10009891 | hypothetical protein MGYG_00374 [Microsporum gypseum CBS 118893] | 30.9 | 5.0E-19 |
| DIP_10009892 | hypothetical protein PFICI_01800 [Pestalotiopsis fici W106-1] | 56.7 | 0.0 |
| DIP_10009893 | hypothetical protein PFICI_01796 [Pestalotiopsis fici W106-1] | 55.6 | 4.0E-89 |
| DIP_10009894 | putative utp-glucose-1-phosphate uridylyltransferase protein [Togninia minima UCRPA7] | 90.6 | 0.0 |
| DIP_10009895 | putative phd-finger domain-containing protein [Togninia minima UCRPA7] | 57.4 | 2.0E-72 |
| DIP_10009896 | putative histone deacetylase phd1 protein [Eutypa lata UCREL1] | 78.8 | 0.0 |
| DIP_10009897 | NA | | |
| DIP_10009898 | hypothetical protein PFICI_01776 [Pestalotiopsis fici W106-1] | 61.7 | 0.0 |
| DIP_10009899 | conserved hypothetical protein [Neurospora crassa] | 61.2 | 2.0E-101 |
| DIP_10009900 | extracellular matrix protein [Colletotrichum higginsianum] | 62.9 | 7.0E-33 |
| DIP_10009901 | putative 5-azacytidine resistance protein azr1 protein [Eutypa lata UCREL1] | 71.7 | 4.0E-174 |
| DIP_10009902 | Succinate dehydrogenase [ubiquinone] iron-sulfur subunit [Pestalotiopsis fici W106-1] | 86.2 | 0.0 |
| DIP_10009903 | putative tubulin-specific chaperone e protein [Togninia minima UCRPA7] | 53.6 | 0.0 |
| DIP_10009904 | hypothetical protein PFICI_01782 [Pestalotiopsis fici W106-1] | 86.4 | 0.0 |
| DIP_10009905 | NA | | |
| DIP_10009906 | putative mitochondrial cytochrome b2 protein [Eutypa lata UCREL1] | 74.3 | 0.0 |
| DIP_10009907 | putative aldo-keto reductase protein [Eutypa lata UCREL1] | 58.5 | 1.0E-117 |
| DIP_10009908 | hypothetical protein FOVG_17486 [Fusarium oxysporum f. sp. pisi HDV247] | 32.7 | 2.0E-83 |
| DIP_10009909 | putative enolase protein [Eutypa lata UCREL1] | 93.8 | 0.0 |
| DIP_10009910 | putative 40s ribosomal protein s13 protein [Eutypa lata UCREL1] | 93.4 | 5.0E-103 |
| DIP_10009911 | hypothetical protein HMPREF1624_07648 [Sporothrix schenckii ATCC 58251] | 50.8 | 6.0E-29 |
| DIP_10009912 | hypothetical protein PFICI_00327 [Pestalotiopsis fici W106-1] | 75.6 | 0.0 |
| DIP_10009913 | NA | | |
| DIP_10009914 | uncharacterized protein FFUJ_09279 [Fusarium fujikuroi IMI 58289] | 32.5 | 3.0E-54 |
| DIP_10009915 | putative dolichyl-phosphate beta-glucosyltransferase [Metarhizium anisopliae] | 53.3 | 4.0E-161 |
| DIP_10009916 | hypothetical protein PFICI_04283 [Pestalotiopsis fici W106-1] | 63.3 | 0.0 |
| DIP_10009917 | hypothetical protein UCREL1_10240 [Eutypa lata UCREL1] | 37.1 | 5.0E-77 |
| DIP_10009918 | hypothetical protein GLRG_02830 [Colletotrichum graminicola M1.001] | 46.2 | 7.0E-17 |
| DIP_10009919 | hypothetical protein CSUB01_02812 [Colletotrichum sublineola] | 55.3 | 6.0E-179 |
| DIP_10009920 | Eukaryotic translation initiation factor 2 subunit alpha [Pestalotiopsis fici W106-1] | 90.5 | 0.0 |
| DIP_10009921 | Lysophospholipase NTE1 [Pestalotiopsis fici W106-1] | 76 | 0.0 |
| DIP_10009922 | putative carbohydrate esterase family 3 protein [Eutypa lata UCREL1] | 54.8 | 9.0E-77 |
| DIP_10009923 | NA | | |
| DIP_10009924 | hypothetical protein PFICI_15303 [Pestalotiopsis fici W106-1] | 51.9 | 1.0E-80 |
| DIP_10009925 | putative periodic tryptophan protein 1 protein [Eutypa lata UCREL1] | 75.4 | 0.0 |
| DIP_10009926 | putative pre-mrna-splicing factor atp-dependent rna helicase prp16 protein [Eutypa lata UCREL1] | 85.8 | 0.0 |
| DIP_10009927 | NA | | |
| DIP_10009928 | hypothetical protein PFICI_00450 [Pestalotiopsis fici W106-1] | 57.9 | 7.0E-130 |
| DIP_10009929 | NA | | |
| DIP_10009930 | hypothetical protein MYCTH_2296698 [Myceliophthora thermophila ATCC 42464] | 64.1 | 3.0E-128 |
| DIP_10009931 | putative -like protein [Eutypa lata UCREL1] | 70.9 | 3.0E-97 |
| DIP_10009932 | hypothetical protein GGTG_01840 [Gaeumannomyces graminis var. tritici R3-111a-1] | 68.4 | 3.0E-60 |
| DIP_10009933 | hypothetical protein FAVG1_01901 [Fusarium avenaceum] | 72.8 | 1.0E-175 |
| DIP_10009934 | hypothetical protein PFICI_00453 [Pestalotiopsis fici W106-1] | 72.7 | 0.0 |
| DIP_10009935 | NA | | |
| DIP_10009936 | NA | | |
| DIP_10009937 | hypothetical protein PFICI_00452 [Pestalotiopsis fici W106-1] | 72.5 | 0.0 |
| DIP_10009938 | hypothetical protein FPSE_05226 [Fusarium pseudograminearum CS3096] | 78.3 | 2.0E-25 |
| DIP_10009939 | NA | | |
| DIP_10009940 | NA | | |
| DIP_10009941 | hypothetical protein ACRE_087730 [Acremonium chrysogenum ATCC 11550] | 33.3 | 1.0E-48 |
| DIP_10009942 | hypothetical protein CFIO01_11955 [Colletotrichum fioriniae PJ7] | 65.3 | 0.0 |
| DIP_10009943 | hypothetical protein PFICI_00005 [Pestalotiopsis fici W106-1] | 46.5 | 5.0E-98 |
| DIP_10009944 | hypothetical protein FAVG1_01908 [Fusarium avenaceum] | 58.2 | 1.0E-31 |
| DIP_10009945 | putative transcription factor tfiiic complex subunit protein [Eutypa lata UCREL1] | 36.4 | 5.0E-133 |
| DIP_10009946 | hypothetical protein PFICI_06703 [Pestalotiopsis fici W106-1] | 77.2 | 0.0 |
| DIP_10009947 | hypothetical protein PFICI_06704 [Pestalotiopsis fici W106-1] | 77.1 | 0.0 |
| DIP_10009948 | putative aryl-alcohol dehydrogenase protein [Eutypa lata UCREL1] | 67.7 | 0.0 |
| DIP_10009949 | putative nad-dependent epimerase dehydratase protein [Eutypa lata UCREL1] | 72.5 | 4.0E-156 |
| DIP_10009950 | hypothetical protein PFICI_06707 [Pestalotiopsis fici W106-1] | 59.1 | 0.0 |
| DIP_10009951 | hypothetical protein PFICI_06708 [Pestalotiopsis fici W106-1] | 75.8 | 0.0 |
| DIP_10009952 | hypothetical protein PFICI_06709 [Pestalotiopsis fici W106-1] | 48.6 | 2.0E-149 |
| DIP_10009953 | hypothetical protein PFICI_06710 [Pestalotiopsis fici W106-1] | 78.3 | 0.0 |
| DIP_10009954 | Acetolactate synthase catalytic subunit [Pestalotiopsis fici W106-1] | 82.1 | 0.0 |
| DIP_10009955 | hypothetical protein OCS_00408 [Ophiocordyceps sinensis CO18] | 34.4 | 3.0E-66 |
| DIP_10009956 | putative mitochondrial f1f0 atp synthase subunit f protein [Eutypa lata UCREL1] | 69.3 | 4.0E-52 |
| DIP_10009957 | putative nadh-ubiquinone oxidoreductase 21 kda subunit protein [Eutypa lata UCREL1] | 68.7 | 2.0E-95 |
| DIP_10009958 | hypothetical protein PFICI_14553 [Pestalotiopsis fici W106-1] | 66.5 | 2.0E-105 |
| DIP_10009959 | hypothetical protein PFICI_14552 [Pestalotiopsis fici W106-1] | 70.2 | 0.0 |
| DIP_10009960 | putative adenylosuccinate synthetase protein [Eutypa lata UCREL1] | 55.2 | 4.0E-113 |
| DIP_10009961 | hypothetical protein PFICI_12850 [Pestalotiopsis fici W106-1] | 71.3 | 0.0 |
| DIP_10009962 | hypothetical protein PFICI_12849 [Pestalotiopsis fici W106-1] | 52.6 | 1.0E-56 |
| DIP_10009963 | putative histone acetyltransferase esa1 protein [Eutypa lata UCREL1] | 77.6 | 0.0 |
| DIP_10009964 | putative classes i and ii family protein [Eutypa lata UCREL1] | 45.3 | 4.0E-145 |
| DIP_10009965 | duf1237 domain protein [Colletotrichum orbiculare MAFF 240422] | 66 | 0.0 |
| DIP_10009966 | putative pantoate-beta-alanine ligase protein [Eutypa lata UCREL1] | 72.5 | 0.0 |
| DIP_10009967 | NA | | |
| DIP_10009968 | putative phosphomevalonate kinase protein [Eutypa lata UCREL1] | 64.8 | 0.0 |
| DIP_10009969 | hypothetical protein UCREL1_2850 [Eutypa lata UCREL1] | 59.1 | 3.0E-32 |
| DIP_10009970 | putative aldehyde dehydrogenase protein [Eutypa lata UCREL1] | 80.3 | 0.0 |
| DIP_10009971 | putative pre-rrna-processing protein esf2 protein [Eutypa lata UCREL1] | 62.1 | 3.0E-138 |
| DIP_10009972 | NA | | |
| DIP_10009973 | 60S ribosomal export protein NMD3 [Pestalotiopsis fici W106-1] | 74.5 | 0.0 |
| DIP_10009974 | hypothetical protein PFICI_11673 [Pestalotiopsis fici W106-1] | 53.8 | 0.0 |
| DIP_10009975 | hypothetical protein PISL3812_08299 [Talaromyces islandicus] | 54.3 | 1.0E-76 |
| DIP_10009976 | NA | | |
| DIP_10009977 | putative alpha-ketoglutarate-dependent taurine dioxygenase protein [Togninia minima UCRPA7] | 78.7 | 0.0 |
| DIP_10009978 | hypothetical protein OIDMADRAFT_180581 [Oidiodendron maius Zn] | 44.4 | 1.0E-33 |
| DIP_10009979 | hypothetical protein PFICI_13846 [Pestalotiopsis fici W106-1] | 43 | 9.0E-82 |
| DIP_10009980 | putative multidrug transporter of the major facilitator superfamily protein [Eutypa lata UCREL1] | 66.5 | 0.0 |
| DIP_10009981 | putative short-chain dehydrogenase reductase protein [Eutypa lata UCREL1] | 47.4 | 2.0E-105 |
| DIP_10009982 | hypothetical protein CFIO01_10901 [Colletotrichum fioriniae PJ7] | 55.8 | 2.0E-142 |
| DIP_10009983 | putative metalloprotease mep1 protein [Eutypa lata UCREL1] | 51.5 | 3.0E-78 |
| DIP_10009984 | NA | | |
| DIP_10009985 | hypothetical protein PFICI_02094 [Pestalotiopsis fici W106-1] | 44.2 | 9.0E-152 |
| DIP_10009986 | hypothetical protein PFICI_03087 [Pestalotiopsis fici W106-1] | 37.6 | 3.0E-30 |
| DIP_10009987 | hypothetical protein PFICI_02096 [Pestalotiopsis fici W106-1] | 42.1 | 0.0 |
| DIP_10009988 | conserved hypothetical protein [Chaetomium globosum CBS 148.51] | 85.7 | 0.0 |
| DIP_10009989 | Eukaryotic translation initiation factor 3 subunit B [Hirsutella minnesotensis 3608] | 72.8 | 0.0 |
| DIP_10009990 | NA | | |
| DIP_10009991 | unnamed protein product [Aspergillus niger] | 30.6 | 4.0E-22 |
| DIP_10009992 | NA | | |
| DIP_10009993 | hypothetical protein PFICI_02101 [Pestalotiopsis fici W106-1] | 61.3 | 0.0 |
| DIP_10009994 | hypothetical protein MYCTH_2309477 [Myceliophthora thermophila ATCC 42464] | 38.1 | 3.0E-43 |
| DIP_10009995 | NA | | |
| DIP_10009996 | putative integral membrane protein [Diaporthe ampelina] | 32.4 | 1.0E-73 |
| DIP_10009997 | putative nucleoside-diphosphate-sugar epimerase [Diaporthe ampelina] | 41.4 | 3.0E-72 |
| DIP_10009998 | hypothetical protein PFICI_00034 [Pestalotiopsis fici W106-1] | 65.2 | 0.0 |
| DIP_10009999 | putative glucose-6-phosphate 1-dehydrogenase protein [Eutypa lata UCREL1] | 92.9 | 0.0 |
| DIP_10010000 | 60S ribosomal protein L33-A [Colletotrichum fioriniae PJ7] | 87.2 | 4.0E-68 |
| DIP_10010001 | NA | | |
| DIP_10010002 | hypothetical protein PFICI_00033 [Pestalotiopsis fici W106-1] | 55.4 | 5.0E-107 |
| DIP_10010003 | putative vma21-like domain-containing protein [Eutypa lata UCREL1] | 66.4 | 6.0E-44 |
| DIP_10010004 | putative snf2 family helicase protein [Eutypa lata UCREL1] | 68.4 | 0.0 |
| DIP_10010005 | putative aldo-keto reductase protein [Eutypa lata UCREL1] | 60.8 | 9.0E-120 |
| DIP_10010006 | hypothetical protein PFICI_01634 [Pestalotiopsis fici W106-1] | 61.6 | 1.0E-49 |
| DIP_10010007 | murein transglycosylase [Fusarium verticillioides 7600] | 35.8 | 5.0E-85 |
| DIP_10010008 | NA | | |
| DIP_10010009 | Autophagy-related protein 22 [Pestalotiopsis fici W106-1] | 63.3 | 0.0 |
| DIP_10010010 | F-actin-capping protein subunit beta [Pestalotiopsis fici W106-1] | 85 | 1.0E-170 |
| DIP_10010011 | Splicing factor U2AF subunit A [Pestalotiopsis fici W106-1] | 91.9 | 2.0E-148 |
| DIP_10010012 | putative oxysterol-binding protein [Eutypa lata UCREL1] | 63.2 | 0.0 |
| DIP_10010013 | hypothetical protein UCREL1_7789 [Eutypa lata UCREL1] | 68.9 | 0.0 |
| DIP_10010014 | conserved hypothetical protein [Verticillium alfalfae VaMs.102] | 38.3 | 1.0E-30 |
| DIP_10010015 | hypothetical protein PFICI_01052 [Pestalotiopsis fici W106-1] | 59 | 0.0 |
| DIP_10010016 | Putative Yos1-like protein [Torrubiella hemipterigena] | 81 | 1.0E-43 |
| DIP_10010017 | hypothetical protein PFICI_01050 [Pestalotiopsis fici W106-1] | 69 | 0.0 |
| DIP_10010018 | hypothetical protein FOVG_14904 [Fusarium oxysporum f. sp. pisi HDV247] | 44.8 | 8.0E-168 |
| DIP_10010019 | nad dependent epimerase dehydratase family protein [Colletotrichum orbiculare MAFF 240422] | 72.7 | 0.0 |
| DIP_10010020 | hypothetical protein UCREL1_3235 [Eutypa lata UCREL1] | 45.3 | 9.0E-60 |
| DIP_10010021 | putative cystathionine beta-synthase protein [Eutypa lata UCREL1] | 73.2 | 2.0E-80 |
| DIP_10010022 | putative nrps-like enzyme protein [Eutypa lata UCREL1] | 40.7 | 2.0E-142 |
| DIP_10010023 | putative serine threonine protein kinase protein [Eutypa lata UCREL1] | 38.1 | 4.0E-70 |
| DIP_10010024 | putative c2 domain-containing protein [Eutypa lata UCREL1] | 74.3 | 0.0 |
| DIP_10010025 | NA | | |
| DIP_10010026 | hypothetical protein PFICI_03314 [Pestalotiopsis fici W106-1] | 60.2 | 2.0E-141 |
| DIP_10010027 | NA | | |
| DIP_10010028 | NA | | |
| DIP_10010029 | hypothetical protein PFICI_00043 [Pestalotiopsis fici W106-1] | 85 | 1.0E-68 |
| DIP_10010030 | NA | | |
| DIP_10010031 | hypothetical protein PFICI_01741 [Pestalotiopsis fici W106-1] | 59.4 | 7.0E-117 |
| DIP_10010032 | hypothetical protein PFICI_02004 [Pestalotiopsis fici W106-1] | 68 | 9.0E-91 |
| DIP_10010033 | putative helicase swr-1 [Diaporthe ampelina] | 57.4 | 0.0 |
| DIP_10010034 | NA | | |
| DIP_10010035 | NA | | |
| DIP_10010036 | NA | | |
| DIP_10010037 | hypothetical protein UCREL1_2338 [Eutypa lata UCREL1] | 64.1 | 0.0 |
| DIP_10010038 | NA | | |
| DIP_10010039 | LPS-induced tumor necrosis factor alpha factor, partial [Metarhizium majus ARSEF 297] | 45.8 | 4.0E-28 |
| DIP_10010040 | putative acyl- n-acyltransferase [Diplodia seriata] | 39 | 9.0E-38 |
| DIP_10010041 | 2og-fe oxygenase [Fusarium avenaceum] | 73.2 | 0.0 |
| DIP_10010042 | major facilitator superfamily transporter [Colletotrichum gloeosporioides Cg-14] | 78 | 0.0 |
| DIP_10010043 | NA | | |
| DIP_10010044 | putative had-superfamily hydrolase subfamily iib protein [Togninia minima UCRPA7] | 82 | 3.0E-150 |
| DIP_10010045 | hypothetical protein PFICI_02030 [Pestalotiopsis fici W106-1] | 65 | 0.0 |
| DIP_10010046 | putative dihydrodipicolinate synthetase family protein [Eutypa lata UCREL1] | 72.2 | 7.0E-163 |
| DIP_10010047 | putative c6 zinc finger domain protein [Diplodia seriata] | 30 | 9.0E-75 |
| DIP_10010048 | unnamed protein product [Fusarium sp. FIESC_5 CS3069] | 30 | 2.0E-12 |
| DIP_10010049 | NA | | |
| DIP_10010050 | putative carbamoyl-phosphate synthase protein [Eutypa lata UCREL1] | 78.6 | 0.0 |
| DIP_10010051 | hypothetical protein PFICI_05143 [Pestalotiopsis fici W106-1] | 51.7 | 6.0E-94 |
| DIP_10010052 | NA | | |
| DIP_10010053 | hypothetical protein UCREL1_6071 [Eutypa lata UCREL1] | 51.7 | 1.0E-108 |
| DIP_10010054 | NA | | |
| DIP_10010055 | predicted protein [Nectria haematococca mpVI 77-13-4] | 64.6 | 0.0 |
| DIP_10010056 | NA | | |
| DIP_10010057 | hypothetical protein TI39_contig391g00011 [Zymoseptoria brevis] | 35.4 | 2.0E-138 |
| DIP_10010058 | hypothetical protein DOTSEDRAFT_72985 [Dothistroma septosporum NZE10] | 44.5 | 2.0E-140 |
| DIP_10010059 | NA | | |
| DIP_10010060 | NA | | |
| DIP_10010061 | NA | | |
| DIP_10010062 | RTA1 domain-containing protein [Trichoderma harzianum] | 56 | 3.0E-129 |
| DIP_10010063 | hypothetical protein MYCFIDRAFT_100731, partial [Pseudocercospora fijiensis CIRAD86] | 33.3 | 3.0E-07 |
| DIP_10010064 | NA | | |
| DIP_10010065 | G protein alpha subunit [Rosellinia necatrix] | 89.3 | 0.0 |
| DIP_10010066 | glycogen phosphorylase [Colletotrichum higginsianum] | 82.6 | 0.0 |
| DIP_10010067 | putative mitogen-activated protein kinase protein [Eutypa lata UCREL1] | 95.2 | 0.0 |
| DIP_10010068 | Vacuolar protein-sorting protein BRO1 [Pestalotiopsis fici W106-1] | 77.7 | 0.0 |
| DIP_10010069 | hypothetical protein S7711_07391 [Stachybotrys chartarum IBT 7711] | 74.4 | 4.0E-163 |
| DIP_10010070 | putative chromosome transmission fidelity protein [Eutypa lata UCREL1] | 71.5 | 0.0 |
| DIP_10010071 | hypothetical protein PFICI_04674 [Pestalotiopsis fici W106-1] | 58.2 | 7.0E-85 |
| DIP_10010072 | putative pci domain-containing protein [Eutypa lata UCREL1] | 51.6 | 3.0E-54 |
| DIP_10010073 | putative arabinogalactan endo- -beta-galactosidase protein [Eutypa lata UCREL1] | 53.9 | 4.0E-167 |
| DIP_10010074 | putative gpi-anchored cell wall beta- -endoglucanase protein [Eutypa lata UCREL1] | 65.3 | 2.0E-152 |
| DIP_10010075 | NA | | |
| DIP_10010076 | NA | | |
| DIP_10010077 | hypothetical protein PFICI_00335 [Pestalotiopsis fici W106-1] | 33.1 | 1.0E-26 |
| DIP_10010078 | hypothetical protein NEUTE2DRAFT_100873 [Neurospora tetrasperma FGSC 2509] | 46.6 | 2.0E-64 |
| DIP_10010079 | putative proteasome component c5 protein [Eutypa lata UCREL1] | 86.7 | 9.0E-171 |
| DIP_10010080 | hypothetical protein PFICI_00332 [Pestalotiopsis fici W106-1] | 70.2 | 0.0 |
| DIP_10010081 | NA | | |
| DIP_10010082 | NA | | |
| DIP_10010083 | putative rna polymerase ii transcription factor siii subunit a protein [Eutypa lata UCREL1] | 50.7 | 3.0E-55 |
| DIP_10010084 | putative histidyl-trna synthetase protein [Eutypa lata UCREL1] | 83 | 0.0 |
| DIP_10010085 | hypothetical protein PFICI_00884 [Pestalotiopsis fici W106-1] | 40.4 | 1.0E-80 |
| DIP_10010086 | putative calcium calmodulin-dependent protein kinase protein [Togninia minima UCRPA7] | 80 | 0.0 |
| DIP_10010087 | hypothetical protein SAPIO_CDS3634 [Scedosporium apiospermum] | 79.9 | 2.0E-91 |
| DIP_10010088 | flavin containing amine oxidoreductase [Colletotrichum fioriniae PJ7] | 53.6 | 0.0 |
| DIP_10010089 | hypothetical protein UCDDA912_g09820 [Diaporthe ampelina] | 44.5 | 2.0E-103 |
| DIP_10010090 | hypothetical protein PFICI_07069 [Pestalotiopsis fici W106-1] | 66.4 | 5.0E-177 |
| DIP_10010091 | Cuticle-degrading protease [Scedosporium apiospermum] | 62.6 | 2.0E-161 |
| DIP_10010092 | hypothetical protein SNOG_08094 [Phaeosphaeria nodorum SN15] | 60.7 | 0.0 |
| DIP_10010093 | NA | | |
| DIP_10010094 | proteasome component PRE3 [Magnaporthe oryzae 70-15] | 89.9 | 3.0E-155 |
| DIP_10010095 | Erv1/Alr family protein [Colletotrichum fioriniae PJ7] | 56.3 | 1.0E-70 |
| DIP_10010096 | cyclin [Colletotrichum gloeosporioides Cg-14] | 56.7 | 5.0E-164 |
| DIP_10010097 | putative ATPase [Colletotrichum sublineola] | 56 | 0.0 |
| DIP_10010098 | hypothetical protein PFICI_12780 [Pestalotiopsis fici W106-1] | 72.9 | 0.0 |
| DIP_10010099 | NA | | |
| DIP_10010100 | putative chromo domain-containing protein [Eutypa lata UCREL1] | 35.9 | 2.0E-58 |
| DIP_10010101 | putative paxneb protein [Eutypa lata UCREL1] | 63 | 2.0E-159 |
| DIP_10010102 | RNA polymerase Rpb8 [Colletotrichum fioriniae PJ7] | 69.4 | 5.0E-69 |
| DIP_10010103 | putative nudix hydrolase 20 protein [Eutypa lata UCREL1] | 59 | 1.0E-136 |
| DIP_10010104 | hypothetical protein PFICI_05699 [Pestalotiopsis fici W106-1] | 58.5 | 2.0E-76 |
| DIP_10010105 | putative golgi apyrase protein [Eutypa lata UCREL1] | 67.5 | 0.0 |
| DIP_10010106 | hypothetical protein HIM_00693 [Hirsutella minnesotensis 3608] | 33.4 | 5.0E-46 |
| DIP_10010107 | hypothetical protein FG05_12281 [Fusarium graminearum] | 49.2 | 3.0E-67 |
| DIP_10010108 | conserved hypothetical protein [Verticillium alfalfae VaMs.102] | 37.6 | 1.0E-118 |
| DIP_10010109 | putative nedd8-activating enzyme e1 regulatory subunit protein [Eutypa lata UCREL1] | 66.9 | 0.0 |
| DIP_10010110 | major facilitator superfamily transporter [Colletotrichum fioriniae PJ7] | 51.2 | 0.0 |
| DIP_10010111 | mannan polymerase II complex ANP1 subunit [Magnaporthe oryzae 70-15] | 74.6 | 0.0 |
| DIP_10010112 | hypothetical protein GMDG_03741 [Pseudogymnoascus destructans 20631-21] | 58.1 | 0.0 |
| DIP_10010113 | putative small nuclear ribonucleo protein [Eutypa lata UCREL1] | 72.5 | 5.0E-124 |
| DIP_10010114 | hypothetical protein UCREL1_2106 [Eutypa lata UCREL1] | 55.2 | 1.0E-177 |
| DIP_10010115 | putative trna-splicing endonuclease subunit sen34 protein [Eutypa lata UCREL1] | 47 | 6.0E-85 |
| DIP_10010116 | putative amino acid sensor-independent protein [Eutypa lata UCREL1] | 57.5 | 0.0 |
| DIP_10010117 | putative 3 exoribonuclease protein [Eutypa lata UCREL1] | 68.8 | 2.0E-134 |
| DIP_10010118 | hypothetical protein MYCTH_2298745 [Myceliophthora thermophila ATCC 42464] | 73 | 2.0E-139 |
| DIP_10010119 | hypothetical protein HIM_02719 [Hirsutella minnesotensis 3608] | 44.5 | 5.0E-97 |
| DIP_10010120 | NA | | |
| DIP_10010121 | hypothetical protein UCREL1_10150 [Eutypa lata UCREL1] | 48.2 | 1.0E-44 |
| DIP_10010122 | hypothetical protein PFICI_04884 [Pestalotiopsis fici W106-1] | 35.4 | 2.0E-60 |
| DIP_10010123 | putative ribose-phosphate pyrophosphokinase protein [Eutypa lata UCREL1] | 81.3 | 0.0 |
| DIP_10010124 | NA | | |
| DIP_10010125 | putative carbohydrate esterase family 3 protein [Eutypa lata UCREL1] | 53.3 | 2.0E-85 |
| DIP_10010126 | putative epoxide hydrolase-like protein [Eutypa lata UCREL1] | 42 | 5.0E-122 |
| DIP_10010127 | conserved hypothetical protein [Byssochlamys spectabilis No. 5] | 46.7 | 0.0 |
| DIP_10010128 | hypothetical protein Z518_07363 [Rhinocladiella mackenziei CBS 650.93] | 50.1 | 5.0E-117 |
| DIP_10010129 | hypothetical protein BAUCODRAFT_151097 [Baudoinia compniacensis UAMH 10762] | 35.6 | 6.0E-113 |
| DIP_10010130 | calcineurin-like phosphoesterase [Ophiostoma piceae UAMH 11346] | 51.3 | 0.0 |
| DIP_10010131 | NA | | |
| DIP_10010132 | hypothetical protein PFICI_01417 [Pestalotiopsis fici W106-1] | 53.6 | 1.0E-56 |
| DIP_10010133 | hypothetical protein PFICI_01418 [Pestalotiopsis fici W106-1] | 43.5 | 2.0E-100 |
| DIP_10010134 | kinesin motor domain-containing protein, partial [Colletotrichum fioriniae PJ7] | 73.8 | 0.0 |
| DIP_10010135 | hypothetical protein PFICI_01420 [Pestalotiopsis fici W106-1] | 47.1 | 0.0 |
| DIP_10010136 | putative valyl-trna synthetase protein [Eutypa lata UCREL1] | 79.6 | 0.0 |
| DIP_10010137 | hypothetical protein UCREL1_94 [Eutypa lata UCREL1] | 45.1 | 1.0E-85 |
| DIP_10010138 | probable cytochrome P450 51 (eburicol 14 alpha-demethylase) [Claviceps purpurea 20.1] | 76.6 | 0.0 |
| DIP_10010139 | NA | | |
| DIP_10010140 | hypothetical protein PFICI_01406 [Pestalotiopsis fici W106-1] | 48.4 | 3.0E-122 |
| DIP_10010141 | hypothetical protein SAPIO_CDS3491 [Scedosporium apiospermum] | 58.2 | 3.0E-149 |
| DIP_10010142 | hypothetical protein PFICI_01404 [Pestalotiopsis fici W106-1] | 52.5 | 0.0 |
| DIP_10010143 | hypothetical protein PFICI_01403 [Pestalotiopsis fici W106-1] | 36.8 | 2.0E-104 |
| DIP_10010144 | hypothetical protein PFICI_01402 [Pestalotiopsis fici W106-1] | 84.7 | 0.0 |
| DIP_10010145 | hypothetical protein HIM_05844 [Hirsutella minnesotensis 3608] | 63 | 1.0E-39 |
| DIP_10010146 | zinc-containing alcohol dehydrogenase, putative [Talaromyces marneffei ATCC 18224] | 39.4 | 8.0E-86 |
| DIP_10010147 | hypothetical protein CSUB01_08716 [Colletotrichum sublineola] | 52.4 | 0.0 |
| DIP_10010148 | hypothetical protein CH063_09345 [Colletotrichum higginsianum] | 51.6 | 6.0E-131 |
| DIP_10010149 | NA | | |
| DIP_10010150 | NA | | |
| DIP_10010151 | small nuclear ribonucleoprotein [Neurospora crassa OR74A] | 90.4 | 2.0E-53 |
| DIP_10010152 | putative pentatricopeptide repeat protein [Eutypa lata UCREL1] | 50.9 | 1.0E-142 |
| DIP_10010153 | putative cell division control protein 3 protein [Eutypa lata UCREL1] | 86.1 | 0.0 |
| DIP_10010154 | putative 30s ribosomal subunit protein [Eutypa lata UCREL1] | 49.5 | 4.0E-136 |
| DIP_10010155 | MFS general substrate transporter [Glarea lozoyensis ATCC 20868] | 53.8 | 9.0E-93 |
| DIP_10010156 | putative rna-binding la domain protein [Eutypa lata UCREL1] | 53.4 | 1.0E-122 |
| DIP_10010157 | hypothetical protein [Podospora anserina S mat+] | 71.3 | 0.0 |
| DIP_10010158 | hypothetical protein PFICI_05114 [Pestalotiopsis fici W106-1] | 37.7 | 3.0E-162 |
| DIP_10010159 | NA | | |
| DIP_10010160 | hypothetical protein UCREL1_10235 [Eutypa lata UCREL1] | 48.7 | 3.0E-07 |
| DIP_10010161 | putative aromatic-l-amino-acid decarboxylase protein [Eutypa lata UCREL1] | 63.8 | 0.0 |
| DIP_10010162 | putative polysaccharide deacetylase protein [Eutypa lata UCREL1] | 78.6 | 2.0E-135 |
| DIP_10010163 | putative cystathionine gamma-synthase protein [Eutypa lata UCREL1] | 58 | 0.0 |
| DIP_10010164 | hypothetical protein CHGG_02734 [Chaetomium globosum CBS 148.51] | 45.6 | 0.0 |
| DIP_10010165 | putative cation efflux family protein [Eutypa lata UCREL1] | 73.6 | 0.0 |
| DIP_10010166 | Phosphoribosylglycinamide formyltransferase [Ceratocystis platani] | 68.2 | 7.0E-106 |
| DIP_10010167 | hypothetical protein PFICI_00266 [Pestalotiopsis fici W106-1] | 47.9 | 0.0 |
| DIP_10010168 | 6,7-dimethyl-8-ribityllumazine synthase [Colletotrichum graminicola M1.001] | 76.7 | 2.0E-104 |
| DIP_10010169 | hypothetical protein THITE_2088630 [Thielavia terrestris NRRL 8126] | 55.3 | 6.0E-94 |
| DIP_10010170 | putative rhodanese-related sulfurtransferase protein [Neofusicoccum parvum UCRNP2] | 48.4 | 2.0E-31 |
| DIP_10010171 | Ubiquitin-like protein 1 [Pestalotiopsis fici W106-1] | 93.5 | 3.0E-45 |
| DIP_10010172 | putative prohibitin-2 protein [Eutypa lata UCREL1] | 82.6 | 0.0 |
| DIP_10010173 | NA | | |
| DIP_10010174 | hypothetical protein PFICI_00893 [Pestalotiopsis fici W106-1] | 76.2 | 6.0E-103 |
| DIP_10010175 | hypothetical protein CONPUDRAFT_134719 [Coniophora puteana RWD-64-598 SS2] | 51 | 2.0E-82 |
| DIP_10010176 | putative -like mg2+ transporter protein [Eutypa lata UCREL1] | 63.8 | 0.0 |
| DIP_10010177 | tryptophanase [Colletotrichum gloeosporioides Nara gc5] | 60.5 | 0.0 |
| DIP_10010178 | hypothetical protein PISL3812_02079 [Talaromyces islandicus] | 50.9 | 0.0 |
| DIP_10010179 | hypothetical protein TCE0_039f13133 [Talaromyces cellulolyticus] | 38.3 | 4.0E-66 |
| DIP_10010180 | hypothetical protein PISL3812_06646 [Talaromyces islandicus] | 46.9 | 2.0E-82 |
| DIP_10010181 | cytochrome P450 CYP2 subfamily [Aspergillus oryzae 100-8] | 56.7 | 7.0E-159 |
| DIP_10010182 | putative mediator of rna polymerase ii transcription subunit 14 protein [Eutypa lata UCREL1] | 50.9 | 0.0 |
| DIP_10010183 | putative polyketide synthase protein [Eutypa lata UCREL1] | 46.1 | 4.0E-48 |
| DIP_10010184 | hypothetical protein PFICI_00903 [Pestalotiopsis fici W106-1] | 45 | 4.0E-77 |
| DIP_10010185 | NA | | |
| DIP_10010186 | hypothetical protein PFICI_00901 [Pestalotiopsis fici W106-1] | 74.4 | 0.0 |
| DIP_10010187 | hypothetical protein PFICI_00900 [Pestalotiopsis fici W106-1] | 65.9 | 0.0 |
| DIP_10010188 | putative integral membrane family protein [Eutypa lata UCREL1] | 35.2 | 3.0E-42 |
| DIP_10010189 | hypothetical protein CGLO_01545 [Colletotrichum gloeosporioides Cg-14] | 33.3 | 8.0E-31 |
| DIP_10010190 | hypothetical protein PFICI_14670 [Pestalotiopsis fici W106-1] | 71.7 | 0.0 |
| DIP_10010191 | hypothetical protein [Podospora anserina S mat+] | 62 | 0.0 |
| DIP_10010192 | NA | | |
| DIP_10010193 | hypothetical protein THITE_2107752 [Thielavia terrestris NRRL 8126] | 33.8 | 2.0E-26 |
| DIP_10010194 | hypothetical protein NCU05314 [Neurospora crassa OR74A] | 39.8 | 1.0E-55 |
| DIP_10010195 | Mitochondria fission 1 protein [Pestalotiopsis fici W106-1] | 90.7 | 4.0E-99 |
| DIP_10010196 | kinetochore protein SPC24 [Pestalotiopsis fici W106-1] | 62 | 1.0E-90 |
| DIP_10010197 | hypothetical protein PFICI_05177 [Pestalotiopsis fici W106-1] | 68.7 | 8.0E-172 |
| DIP_10010198 | hypothetical protein PFICI_05178 [Pestalotiopsis fici W106-1] | 67.4 | 0.0 |
| DIP_10010199 | NA | | |
| DIP_10010200 | putative cut9-interacting protein scn1 protein [Eutypa lata UCREL1] | 59.9 | 1.0E-162 |
| DIP_10010201 | hypothetical protein UCREL1_5513 [Eutypa lata UCREL1] | 31.2 | 6.0E-36 |
| DIP_10010202 | hypothetical protein PFICI_05176 [Pestalotiopsis fici W106-1] | 62.9 | 0.0 |
| DIP_10010203 | hypothetical protein FGSG_09802 [Fusarium graminearum PH-1] | 47.7 | 3.0E-71 |
| DIP_10010204 | putative tpr repeat protein [Eutypa lata UCREL1] | 61.2 | 0.0 |
| DIP_10010205 | hypothetical protein PFICI_05171 [Pestalotiopsis fici W106-1] | 74.1 | 0.0 |
| DIP_10010206 | hypothetical protein PFICI_05170 [Pestalotiopsis fici W106-1] | 57.6 | 5.0E-165 |
| DIP_10010207 | putative glycoside hydrolase family 43 protein [Eutypa lata UCREL1] | 53 | 5.0E-118 |
| DIP_10010208 | hypothetical protein PFICI_04788 [Pestalotiopsis fici W106-1] | 53.3 | 4.0E-137 |
| DIP_10010209 | hypothetical protein CFIO01_10398 [Colletotrichum fioriniae PJ7] | 69.8 | 0.0 |
| DIP_10010210 | hypothetical protein PFICI_05018 [Pestalotiopsis fici W106-1] | 76.8 | 0.0 |
| DIP_10010211 | Chitin synthase D [Pestalotiopsis fici W106-1] | 72.6 | 0.0 |
| DIP_10010212 | hypothetical protein PFICI_05221 [Pestalotiopsis fici W106-1] | 84.4 | 0.0 |
| DIP_10010213 | putative nadh-ubiquinone oxidoreductase kda subunit protein [Eutypa lata UCREL1] | 55.9 | 8.0E-82 |
| DIP_10010214 | hypothetical protein PFICI_05732 [Pestalotiopsis fici W106-1] | 31.5 | 5.0E-36 |
| DIP_10010215 | hypothetical protein PFICI_05729 [Pestalotiopsis fici W106-1] | 83.5 | 2.0E-161 |
| DIP_10010216 | NA | | |
| DIP_10010217 | putative embryonic ectoderm development protein [Eutypa lata UCREL1] | 44.4 | 6.0E-110 |
| DIP_10010218 | NA | | |
| DIP_10010219 | hypothetical protein FOXB_07976 [Fusarium oxysporum Fo5176] | 51.2 | 1.0E-76 |
| DIP_10010220 | hypothetical protein SS1G_13293 [Sclerotinia sclerotiorum 1980] | 46.4 | 1.0E-99 |
| DIP_10010221 | hypothetical protein PFICI_05434 [Pestalotiopsis fici W106-1] | 83.5 | 0.0 |
| DIP_10010222 | hypothetical protein MYCTH_94484 [Myceliophthora thermophila ATCC 42464] | 33.1 | 2.0E-45 |
| DIP_10010223 | NA | | |
| DIP_10010224 | hypothetical protein SMAC_03817 [Sordaria macrospora k-hell] | 34 | 2.0E-43 |
| DIP_10010225 | hypothetical protein UCREL1_3500 [Eutypa lata UCREL1] | 51.7 | 6.0E-58 |
| DIP_10010226 | putative sulfite reductase beta subunit protein [Eutypa lata UCREL1] | 82.2 | 0.0 |
| DIP_10010227 | putative ribose 5-phosphate isomerase a protein [Eutypa lata UCREL1] | 75.5 | 1.0E-150 |
| DIP_10010228 | putative ras-like protein rab-6a protein [Togninia minima UCRPA7] | 87.1 | 1.0E-130 |
| DIP_10010229 | putative 60s ribosomal biogenesis protein [Eutypa lata UCREL1] | 74.1 | 0.0 |
| DIP_10010230 | hypothetical protein OIDMADRAFT_121941, partial [Oidiodendron maius Zn] | 44.4 | 2.0E-52 |
| DIP_10010231 | putative mitochondrial nadh dehydrogenase protein [Eutypa lata UCREL1] | 76.6 | 0.0 |
| DIP_10010232 | hypothetical protein PV08_00537 [Exophiala spinifera] | 36.5 | 3.0E-115 |
| DIP_10010233 | hypothetical protein SAPIO_CDS9216 [Scedosporium apiospermum] | 47.7 | 3.0E-146 |
| DIP_10010234 | DNA replication licensing factor mcm6 [Pestalotiopsis fici W106-1] | 79.1 | 0.0 |
| DIP_10010235 | NA | | |
| DIP_10010236 | putative suppressor of mck1 protein [Eutypa lata UCREL1] | 66.8 | 0.0 |
| DIP_10010237 | hypothetical protein PFICI_04553 [Pestalotiopsis fici W106-1] | 60.9 | 8.0E-153 |
| DIP_10010238 | hypothetical protein PFICI_01542 [Pestalotiopsis fici W106-1] | 79.4 | 0.0 |
| DIP_10010239 | putative amino-acid transporter arg-13 protein [Eutypa lata UCREL1] | 71.6 | 9.0E-156 |
| DIP_10010240 | hypothetical protein PFICI_01540 [Pestalotiopsis fici W106-1] | 65.5 | 1.0E-89 |
| DIP_10010241 | hypothetical protein PFICI_01961 [Pestalotiopsis fici W106-1] | 78.2 | 0.0 |
| DIP_10010242 | hypothetical protein FOYG_07046 [Fusarium oxysporum FOSC 3-a] | 47.8 | 2.0E-128 |
| DIP_10010243 | hypothetical protein S40285_00809 [Stachybotrys chlorohalonata IBT 40285] | 35.8 | 1.0E-31 |
| DIP_10010244 | putative uridine nucleosidase protein [Eutypa lata UCREL1] | 61.9 | 6.0E-151 |
| DIP_10010245 | hypothetical protein PFICI_01849 [Pestalotiopsis fici W106-1] | 59.6 | 3.0E-176 |

| | | | |
|---|---|---|---|
| DIP_10010246 | putative dna polymerase iv [Diplodia seriata] | 54.2 | 0.0 |
| DIP_10010247 | hypothetical protein W97_07652 [Coniosporium apollinis CBS 100218] | 35.3 | 3.0E-09 |
| DIP_10010248 | hypothetical protein PFICI_01862 [Pestalotiopsis fici W106-1] | 74.9 | 0.0 |
| DIP_10010249 | hypothetical protein O988_07131 [Pseudogymnoascus pannorum VKM F-3808] | 37.9 | 2.0E-21 |
| DIP_10010250 | putative uncharacterized thoc5 family protein [Eutypa lata UCREL1] | 76.4 | 2.0E-112 |
| DIP_10010251 | hypothetical protein CFIO01_01375 [Colletotrichum fioriniae PJ7] | 69.4 | 0.0 |
| DIP_10010252 | putative family protein [Eutypa lata UCREL1] | 51.4 | 6.0E-61 |
| DIP_10010253 | putative fk506-binding protein 4 protein [Eutypa lata UCREL1] | 68.1 | 0.0 |
| DIP_10010254 | hypothetical protein PFICI_01857 [Pestalotiopsis fici W106-1] | 57.5 | 0.0 |
| DIP_10010255 | WSC domain protein, putative [Aspergillus clavatus NRRL 1] | 34.4 | 1.0E-17 |
| DIP_10010256 | hypothetical protein FOC4_g10015205 [Fusarium oxysporum f. sp. cubense race 4] | 50 | 9.0E-112 |
| DIP_10010257 | hypothetical protein MYCTH_2295918 [Myceliophthora thermophila ATCC 42464] | 70.2 | 1.0E-152 |
| DIP_10010258 | putative mitochondrial protein [Eutypa lata UCREL1] | 56.1 | 9.0E-95 |
| DIP_10010259 | Mitochondrial intermediate peptidase [Pestalotiopsis fici W106-1] | 62.3 | 0.0 |
| DIP_10010260 | putative mitotic check point protein [Eutypa lata UCREL1] | 72.8 | 0.0 |
| DIP_10010261 | putative sulfate permease protein [Eutypa lata UCREL1] | 65.2 | 0.0 |
| DIP_10010262 | hypothetical protein PFICI_05726 [Pestalotiopsis fici W106-1] | 31.5 | 8.0E-19 |
| DIP_10010263 | hypothetical protein PFICI_05725 [Pestalotiopsis fici W106-1] | 51.6 | 7.0E-46 |
| DIP_10010264 | NA | | |
| DIP_10010265 | enolase C-terminal domain-like protein [Coniophora puteana RWD-64-598 SS2] | 72.1 | 0.0 |
| DIP_10010266 | sugar transporter, putative [Aspergillus flavus NRRL3357] | 58.9 | 0.0 |
| DIP_10010267 | hypothetical protein W97_03560 [Coniosporium apollinis CBS 100218] | 43.6 | 3.0E-82 |
| DIP_10010268 | hypothetical protein CGLO_07143 [Colletotrichum gloeosporioides Cg-14] | 38.3 | 2.0E-82 |
| DIP_10010269 | NA | | |
| DIP_10010270 | NA | | |
| DIP_10010271 | glucan endo-1,3-alpha-glucosidase agn1 precursor, putative [Talaromyces marneffei ATCC 18224] | 35.3 | 6.0E-72 |
| DIP_10010272 | NA | | |
| DIP_10010273 | hypothetical protein MAC_04934 [Metarhizium acridum CQMa 102] | 38.1 | 3.0E-66 |
| DIP_10010274 | F-box domain-containing protein [Colletotrichum higginsianum] | 40.2 | 2.0E-55 |
| DIP_10010275 | NA | | |
| DIP_10010276 | NA | | |
| DIP_10010277 | F-box domain-containing protein [Ophiocordyceps sinensis CO18] | 31 | 2.0E-18 |
| DIP_10010278 | hypothetical protein UCREL1_2595 [Eutypa lata UCREL1] | 33.4 | 2.0E-63 |
| DIP_10010279 | Protein transport [Pestalotiopsis fici W106-1] | 84.5 | 1.0E-78 |
| DIP_10010280 | hypothetical protein CGLO_07591 [Colletotrichum gloeosporioides Cg-14] | 61 | 8.0E-72 |
| DIP_10010281 | unnamed protein product [Sordaria macrospora k-hell] | 91.7 | 3.0E-42 |
| DIP_10010282 | hypothetical protein PFICI_05722 [Pestalotiopsis fici W106-1] | 79.6 | 0.0 |
| DIP_10010283 | putative gnat family protein [Eutypa lata UCREL1] | 39.2 | 7.0E-48 |
| DIP_10010284 | hypothetical protein V491_07660 [Pseudogymnoascus pannorum VKM F-3775] | 43.4 | 1.0E-87 |
| DIP_10010285 | hypothetical protein PFICI_04628 [Pestalotiopsis fici W106-1] | 71.2 | 0.0 |
| DIP_10010286 | hypothetical protein BofuT4_P001530.1 [Botrytis cinerea T4] | 56.1 | 1.0E-82 |
| DIP_10010287 | hypothetical protein TRIVIDRAFT_150591 [Trichoderma virens Gv29-8] | 39.4 | 0.0 |
| DIP_10010288 | putative 26s protease subunit rpt4 protein [Eutypa lata UCREL1] | 92.6 | 0.0 |
| DIP_10010289 | hypothetical protein PFICI_01754 [Pestalotiopsis fici W106-1] | 39.8 | 1.0E-42 |
| DIP_10010290 | hypothetical protein PFICI_01760 [Pestalotiopsis fici W106-1] | 47.4 | 0.0 |
| DIP_10010291 | Putative DUF1275 domain protein [Torrubiella hemipterigena] | 64.8 | 1.0E-100 |
| DIP_10010292 | hypothetical protein PFICI_02008 [Pestalotiopsis fici W106-1] | 64.5 | 0.0 |
| DIP_10010293 | putative transporter Avl9 [Colletotrichum sublineola] | 72.8 | 0.0 |
| DIP_10010294 | putative glucosamine-fructose-6-phosphate aminotransferase protein [Eutypa lata UCREL1] | 93.9 | 0.0 |
| DIP_10010295 | hypothetical protein PFICI_01757 [Pestalotiopsis fici W106-1] | 77 | 1.0E-156 |
| DIP_10010296 | hypothetical protein CGGC5_9544 [Colletotrichum gloeosporioides Nara gc5] | 51.9 | 1.0E-175 |
| DIP_10010297 | hypothetical protein CH063_01240 [Colletotrichum higginsianum] | 52.6 | 0.0 |
| DIP_10010298 | putative casein kinase ii beta 2 subunit protein [Eutypa lata UCREL1] | 53.8 | 5.0E-147 |
| DIP_10010299 | hypothetical protein PFICI_00411 [Pestalotiopsis fici W106-1] | 70.4 | 1.0E-20 |
| DIP_10010300 | hypothetical protein NEUTE1DRAFT_106582 [Neurospora tetrasperma FGSC 2508] | 61.6 | 6.0E-106 |
| DIP_10010301 | hypothetical protein PFICI_01192 [Pestalotiopsis fici W106-1] | 70.3 | 0.0 |
| DIP_10010302 | hypothetical protein PFICI_01193 [Pestalotiopsis fici W106-1] | 48.4 | 3.0E-91 |
| DIP_10010303 | hypothetical protein NFIA_047850 [Neosartorya fischeri NRRL 181] | 39.6 | 2.0E-16 |
| DIP_10010304 | hypothetical protein NECHADRAFT_30482 [Nectria haematococca mpVI 77-13-4] | 50.9 | 4.0E-98 |
| DIP_10010305 | putative sun domain-containing protein [Eutypa lata UCREL1] | 57.1 | 2.0E-148 |
| DIP_10010306 | hypothetical protein SS1G_04802 [Sclerotinia sclerotiorum 1980] | 40.5 | 2.0E-10 |
| DIP_10010307 | NA | | |
| DIP_10010308 | 40S ribosomal protein S15 [Colletotrichum fioriniae PJ7] | 93.5 | 1.0E-103 |
| DIP_10010309 | hypothetical protein PFICI_01805 [Pestalotiopsis fici W106-1] | 58.5 | 3.0E-64 |
| DIP_10010310 | hypothetical protein PFICI_01806 [Pestalotiopsis fici W106-1] | 47.6 | 2.0E-115 |
| DIP_10010311 | glycoprotease family protein [Colletotrichum orbiculare MAFF 240422] | 43.4 | 0.0 |
| DIP_10010312 | hypothetical protein PV06_11049 [Exophiala oligosperma] | 30.6 | 6.0E-39 |
| DIP_10010313 | hypothetical protein AFUB_102330 [Aspergillus fumigatus A1163] | 50.7 | 9.0E-90 |
| DIP_10010314 | NA | | |
| DIP_10010315 | putative 3-hydroxyanthranilate -dioxygenase protein [Eutypa lata UCREL1] | 84.4 | 6.0E-118 |
| DIP_10010316 | hypothetical protein FPSE_02725 [Fusarium pseudograminearum CS3096] | 46.5 | 8.0E-59 |
| DIP_10010317 | hypothetical protein PFICI_01809 [Pestalotiopsis fici W106-1] | 62.4 | 2.0E-145 |
| DIP_10010318 | hypothetical protein GLRG_02422 [Colletotrichum graminicola M1.001] | 32.4 | 8.0E-22 |
| DIP_10010319 | NA | | |
| DIP_10010320 | hypothetical protein PFICI_01812 [Pestalotiopsis fici W106-1] | 73.8 | 0.0 |
| DIP_10010321 | hypothetical protein FOZG_05602 [Fusarium oxysporum Fo47] | 59 | 0.0 |
| DIP_10010322 | putative c2 domain containing protein [Eutypa lata UCREL1] | 71.2 | 0.0 |
| DIP_10010323 | Hybrid signal transduction histidine kinase J [Pestalotiopsis fici W106-1] | 82.3 | 0.0 |
| DIP_10010324 | hypothetical protein PAAG_00844 [Paracoccidioides sp. 'lutzii' Pb01] | 57.4 | 7.0E-12 |
| DIP_10010325 | HMG box protein [Colletotrichum graminicola M1.001] | 68.2 | 1.0E-112 |
| DIP_10010326 | putative condensin complex component cnd2 protein [Eutypa lata UCREL1] | 70.3 | 0.0 |
| DIP_10010327 | sodium/calcium exchanger protein [Colletotrichum fioriniae PJ7] | 48.5 | 0.0 |
| DIP_10010328 | NA | | |
| DIP_10010329 | Putative Phosphomannomutase [Torrubiella hemipterigena] | 75.3 | 2.0E-149 |
| DIP_10010330 | hypothetical protein PFICI_01820 [Pestalotiopsis fici W106-1] | 71.7 | 0.0 |
| DIP_10010331 | hypothetical protein OIDMADRAFT_18236 [Oidiodendron maius Zn] | 63.6 | 4.0E-41 |
| DIP_10010332 | hypothetical protein PFICI_01823 [Pestalotiopsis fici W106-1] | 75.9 | 0.0 |
| DIP_10010333 | hypothetical protein CTHT_0009340 [Chaetomium thermophilum var. thermophilum DSM 1495] | 48.6 | 0.0 |
| DIP_10010334 | putative atp-binding endoribonuclease protein [Eutypa lata UCREL1] | 43.2 | 0.0 |
| DIP_10010335 | hypothetical protein PFICI_01825 [Pestalotiopsis fici W106-1] | 51.6 | 4.0E-159 |
| DIP_10010336 | NA | | |
| DIP_10010337 | hypothetical protein H633G_04799 [Metarhizium anisopliae BRIP 53284] | 39.5 | 5.0E-120 |
| DIP_10010338 | hypothetical protein PEXP_103390 [Penicillium expansum] | 45.8 | 1.0E-88 |
| DIP_10010339 | DNA-binding protein SMUBP-2 [Metarhizium robertsii ARSEF 23] | 49 | 9.0E-77 |
| DIP_10010340 | putative dna-binding protein smubp-2 protein [Eutypa lata UCREL1] | 72.4 | 0.0 |
| DIP_10010341 | hypothetical protein CSUB01_09578 [Colletotrichum sublineola] | 33.5 | 2.0E-52 |
| DIP_10010342 | hypothetical protein CH063_03137 [Colletotrichum higginsianum] | 39.3 | 4.0E-15 |
| DIP_10010343 | hypothetical protein PFICI_01833 [Pestalotiopsis fici W106-1] | 53.5 | 0.0 |
| DIP_10010344 | putative er membrane protein sh3 protein [Eutypa lata UCREL1] | 70.2 | 4.0E-110 |
| DIP_10010345 | NA | | |
| DIP_10010346 | hypothetical protein H633G_05594, partial [Metarhizium anisopliae BRIP 53284] | 32.3 | 9.0E-16 |
| DIP_10010347 | NA | | |
| DIP_10010348 | hypothetical protein FOCG_01027 [Fusarium oxysporum f. sp. radicis-lycopersici 26381] | 34.7 | 1.0E-29 |
| DIP_10010349 | putative sam and ph domain-containing protein [Eutypa lata UCREL1] | 58 | 0.0 |
| DIP_10010350 | NA | | |
| DIP_10010351 | putative transcriptional regulatory protein pro-1 protein [Eutypa lata UCREL1] | 74.8 | 0.0 |
| DIP_10010352 | NA | | |
| DIP_10010353 | carboxypeptidase D [Fusarium oxysporum f. sp. melonis 26406] | 34.3 | 2.0E-71 |
| DIP_10010354 | hypothetical protein PVAR5_5638 [Byssochlamys spectabilis No. 5] | 41.8 | 2.0E-14 |
| DIP_10010355 | NA | | |
| DIP_10010356 | putative superoxide dismutase protein [Eutypa lata UCREL1] | 47.9 | 6.0E-94 |
| DIP_10010357 | hypothetical protein FOXB_00337 [Fusarium oxysporum Fo5176] | 66.2 | 0.0 |
| DIP_10010358 | hypothetical protein W97_03981 [Coniosporium apollinis CBS 100218] | 33.2 | 4.0E-56 |
| DIP_10010359 | NADP/FAD dependent oxidoreductase [Colletotrichum higginsianum] | 53 | 0.0 |
| DIP_10010360 | zn 2cys6 transcription factor [Colletotrichum gloeosporioides Nara gc5] | 44.4 | 4.0E-126 |
| DIP_10010361 | NA | | |
| DIP_10010362 | putative nad binding rossmann fold protein [Eutypa lata UCREL1] | 76.6 | 0.0 |
| DIP_10010363 | NA | | |
| DIP_10010364 | hypothetical protein GGTG_11692 [Gaeumannomyces graminis var. tritici R3-111a-1] | 43.4 | 3.0E-53 |
| DIP_10010365 | NA | | |
| DIP_10010366 | NA | | |
| DIP_10010367 | NA | | |
| DIP_10010368 | NA | | |
| DIP_10010369 | NA | | |
| DIP_10010370 | hypothetical protein PFICI_00841 [Pestalotiopsis fici W106-1] | 59.1 | 5.0E-60 |
| DIP_10010371 | NA | | |
| DIP_10010372 | hypothetical protein MAC_05253 [Metarhizium acridum CQMa 102] | 33.1 | 7.0E-19 |
| DIP_10010373 | putative domain containing protein [Eutypa lata UCREL1] | 79.3 | 0.0 |
| DIP_10010374 | NA | | |
| DIP_10010375 | hypothetical protein THITE_2114103 [Thielavia terrestris NRRL 8126] | 88.2 | 0.0 |
| DIP_10010376 | NA | | |
| DIP_10010377 | NA | | |
| DIP_10010378 | hypothetical protein UCREL1_2325 [Eutypa lata UCREL1] | 35.1 | 3.0E-17 |
| DIP_10010379 | NA | | |
| DIP_10010380 | NA | | |
| DIP_10010381 | putative dipeptidyl aminopeptidase b protein [Eutypa lata UCREL1] | 76 | 0.0 |
| DIP_10010382 | putative polysaccharide deacetylase protein [Eutypa lata UCREL1] | 70.6 | 0.0 |
| DIP_10010383 | hypothetical protein H072_7350 [Dactylellina haptotyla CBS 200.50] | 66.9 | 3.0E-63 |
| DIP_10010384 | NA | | |
| DIP_10010385 | putative cullulin 3 protein [Eutypa lata UCREL1] | 68.1 | 0.0 |
| DIP_10010386 | DNL zinc finger domain-containing protein [Verticillium dahliae VdLs.17] | 47.2 | 7.0E-55 |
| DIP_10010387 | hypothetical protein PFICI_01229 [Pestalotiopsis fici W106-1] | 59.9 | 0.0 |
| DIP_10010388 | putative cwfj domain-containing protein [Eutypa lata UCREL1] | 58.4 | 0.0 |
| DIP_10010389 | hypothetical protein PFICI_01227 [Pestalotiopsis fici W106-1] | 52.9 | 0.0 |
| DIP_10010390 | NA | | |
| DIP_10010391 | methyltransferase BUD23 [Pestalotiopsis fici W106-1] | 81.6 | 3.0E-164 |
| DIP_10010392 | putative transmembrane adaptor erv26 protein [Eutypa lata UCREL1] | 53.4 | 5.0E-60 |
| DIP_10010393 | hypothetical protein PFICI_01007 [Pestalotiopsis fici W106-1] | 48.6 | 0.0 |
| DIP_10010394 | hypothetical protein PFICI_01012 [Pestalotiopsis fici W106-1] | 62.1 | 0.0 |
| DIP_10010395 | hypothetical protein PFICI_01013 [Pestalotiopsis fici W106-1] | 64.7 | 0.0 |
| DIP_10010396 | putative mfs transporter protein [Eutypa lata UCREL1] | 65 | 0.0 |
| DIP_10010397 | putative metalloprotease 1 protein [Eutypa lata UCREL1] | 58.6 | 5.0E-102 |
| DIP_10010398 | hypothetical protein SPSK_00060 [Sporothrix schenckii 1099-18] | 49.7 | 3.0E-46 |
| DIP_10010399 | hypothetical protein PFICI_01432 [Pestalotiopsis fici W106-1] | 51.6 | 2.0E-59 |
| DIP_10010400 | hypothetical protein PFICI_01433 [Pestalotiopsis fici W106-1] | 65 | 5.0E-150 |
| DIP_10010401 | hypothetical protein CHGG_05981 [Chaetomium globosum CBS 148.51] | 62.4 | 0.0 |
| DIP_10010402 | putative eukaryotic translation initiation factor 3 subunit d protein [Eutypa lata UCREL1] | 84.2 | 0.0 |
| DIP_10010403 | putative ribulose-phosphate 3-epimerase protein [Eutypa lata UCREL1] | 86 | 4.0E-158 |
| DIP_10010404 | hypothetical protein CFIO01_12761 [Colletotrichum fioriniae PJ7] | 63 | 0.0 |
| DIP_10010405 | mitochondrial ATPase [Colletotrichum orbiculare MAFF 240422] | 67.4 | 0.0 |
| DIP_10010406 | NA | | |
| DIP_10010407 | hypothetical protein UCREL1_5420 [Eutypa lata UCREL1] | 72.2 | 0.0 |
| DIP_10010408 | putative peptidase family m28 protein [Eutypa lata UCREL1] | 76.9 | 0.0 |
| DIP_10010409 | phospholipase D Active site motif protein [Magnaporthe oryzae P131] | 53.2 | 0.0 |
| DIP_10010410 | NA | | |
| DIP_10010411 | putative gpi anchored protein [Eutypa lata UCREL1] | 30.9 | 4.0E-11 |
| DIP_10010412 | NA | | |
| DIP_10010413 | hypothetical protein H634G_08154 [Metarhizium anisopliae BRIP 53293] | 36.5 | 1.0E-53 |
| DIP_10010414 | NA | | |
| DIP_10010415 | hypothetical protein THAR02_10002 [Trichoderma harzianum] | 53.7 | 2.0E-101 |
| DIP_10010416 | vacuolar protein-sorting-associated protein [Verticillium dahliae VdLs.17] | 57 | 0.0 |
| DIP_10010417 | putative sac3 ganp domain protein [Eutypa lata UCREL1] | 74.4 | 0.0 |
| DIP_10010418 | putative scd2 ral3 protein [Eutypa lata UCREL1] | 72.4 | 0.0 |
| DIP_10010419 | putative centromere binding protein b protein [Eutypa lata UCREL1] | 72.3 | 1.0E-170 |
| DIP_10010420 | Thymidylate synthase [Pestalotiopsis fici W106-1] | 75.7 | 0.0 |
| DIP_10010421 | hypothetical protein NCU06602 [Neurospora crassa OR74A] | 38.5 | 6.0E-75 |
| DIP_10010422 | hypothetical protein PFICI_05486 [Pestalotiopsis fici W106-1] | 67.3 | 0.0 |
| DIP_10010423 | hypothetical protein PFICI_05485 [Pestalotiopsis fici W106-1] | 44.8 | 3.0E-86 |
| DIP_10010424 | NA | | |
| DIP_10010425 | hypothetical protein PFICI_05483 [Pestalotiopsis fici W106-1] | 63.1 | 0.0 |
| DIP_10010426 | putative an1-type zinc finger protein 1 protein [Eutypa lata UCREL1] | 51.9 | 1.0E-97 |
| DIP_10010427 | hypothetical protein UCREL1_11435 [Eutypa lata UCREL1] | 75.5 | 4.0E-168 |
| DIP_10010428 | hypothetical protein UCREL1_11436 [Eutypa lata UCREL1] | 70.7 | 6.0E-31 |
| DIP_10010429 | hypothetical protein MGG_04155 [Magnaporthe oryzae 70-15] | 56.3 | 2.0E-128 |
| DIP_10010430 | putative aspartate aminotransferase protein [Eutypa lata UCREL1] | 86 | 0.0 |
| DIP_10010431 | hypothetical protein PFICI_12796 [Pestalotiopsis fici W106-1] | 65.3 | 0.0 |
| DIP_10010432 | hypothetical protein PFICI_05078 [Pestalotiopsis fici W106-1] | 61.7 | 8.0E-151 |
| DIP_10010433 | NA | | |
| DIP_10010434 | hypothetical protein PFICI_01552 [Pestalotiopsis fici W106-1] | 37.4 | 1.0E-94 |
| DIP_10010435 | NA | | |
| DIP_10010436 | putative exported protein [Eutypa lata UCREL1] | 50.8 | 0.0 |
| DIP_10010437 | putative pci domain containing protein [Eutypa lata UCREL1] | 69.6 | 0.0 |
| DIP_10010438 | conserved hypothetical protein [Chaetomium globosum CBS 148.51] | 59.5 | 2.0E-134 |
| DIP_10010439 | hypothetical protein TCE0_034r12021 [Talaromyces cellulolyticus] | 49.2 | 0.0 |
| DIP_10010440 | DnaJ [Pestalotiopsis fici W106-1] | 69.2 | 0.0 |
| DIP_10010441 | asparagine synthase [Colletotrichum fioriniae PJ7] | 52.4 | 0.0 |
| DIP_10010442 | hypothetical protein BAUCODRAFT_37640 [Baudoinia compniacensis UAMH 10762] | 32.6 | 7.0E-07 |
| DIP_10010443 | putative annexin a7 protein [Eutypa lata UCREL1] | 66.4 | 6.0E-176 |
| DIP_10010444 | hypothetical protein PFICI_01143 [Pestalotiopsis fici W106-1] | 54.6 | 0.0 |
| DIP_10010445 | hypothetical protein PFICI_01551 [Pestalotiopsis fici W106-1] | 42 | 2.0E-130 |
| DIP_10010446 | hypothetical protein PFICI_01547 [Pestalotiopsis fici W106-1] | 66.7 | 0.0 |
| DIP_10010447 | putative pre-mrna-splicing factor cef-1 protein [Eutypa lata UCREL1] | 70 | 0.0 |
| DIP_10010448 | hypothetical protein PFICI_04508 [Pestalotiopsis fici W106-1] | 81.1 | 0.0 |
| DIP_10010449 | putative plasma membrane channel protein [Eutypa lata UCREL1] | 69.3 | 0.0 |
| DIP_10010450 | phosphoric ester hydrolase, putative [Colletotrichum orbiculare MAFF 240422] | 49.2 | 1.0E-103 |
| DIP_10010451 | NA | | |
| DIP_10010452 | putative endo- -beta- protein [Eutypa lata UCREL1] | 44.7 | 5.0E-148 |
| DIP_10010453 | hypothetical protein TRIVIDRAFT_213380 [Trichoderma virens Gv29-8] | 77.6 | 0.0 |
| DIP_10010454 | putative adenylyl cyclase-associated protein [Eutypa lata UCREL1] | 72.8 | 0.0 |
| DIP_10010455 | putative mak32 protein [Eutypa lata UCREL1] | 66.7 | 0.0 |
| DIP_10010456 | putative dna replication licensing factor mcm3 protein [Eutypa lata UCREL1] | 80.9 | 0.0 |
| DIP_10010457 | hypothetical protein HMPREF1624_01285 [Sporothrix schenckii ATCC 58251] | 55.2 | 0.0 |
| DIP_10010458 | Pc20g09640 [Penicillium rubens Wisconsin 54-1255] | 78.1 | 0.0 |
| DIP_10010459 | putative exocyst complex protein exo70 protein [Eutypa lata UCREL1] | 80 | 0.0 |
| DIP_10010460 | hypothetical protein NCU08013 [Neurospora crassa OR74A] | 71.8 | 6.0E-14 |
| DIP_10010461 | putative fumarylacetoacetate hydrolase family protein [Eutypa lata UCREL1] | 61.9 | 9.0E-121 |
| DIP_10010462 | putative duf455 domain-containing protein [Eutypa lata UCREL1] | 64.2 | 0.0 |
| DIP_10010463 | hypothetical protein PFICI_11937 [Pestalotiopsis fici W106-1] | 68.1 | 0.0 |
| DIP_10010464 | hypothetical protein UCREL1_10880 [Eutypa lata UCREL1] | 65 | 5.0E-129 |
| DIP_10010465 | hypothetical protein PFICI_04932 [Pestalotiopsis fici W106-1] | 72.9 | 0.0 |
| DIP_10010466 | hypothetical protein PV11_07976 [Exophiala sideris] | 53.2 | 0.0 |
| DIP_10010467 | NA | | |
| DIP_10010468 | hypothetical protein PFICI_01745 [Pestalotiopsis fici W106-1] | 62.8 | 0.0 |
| DIP_10010469 | putative dna polymerase protein [Eutypa lata UCREL1] | 61.6 | 0.0 |
| DIP_10010470 | NA | | |
| DIP_10010471 | putative short chain oxidoreductase protein [Eutypa lata UCREL1] | 46.3 | 4.0E-56 |
| DIP_10010472 | NA | | |
| DIP_10010473 | putative ste ste20 ysk protein kinase protein [Eutypa lata UCREL1] | 89.5 | 0.0 |
| DIP_10010474 | putative isocitrate dehydrogenase subunit 1 protein [Eutypa lata UCREL1] | 90.2 | 0.0 |
| DIP_10010475 | hypothetical protein PV09_08855 [Verruconis gallopava] | 55.6 | 3.0E-25 |
| DIP_10010476 | NA | | |
| DIP_10010477 | putative arginine permease protein [Eutypa lata UCREL1] | 66.8 | 0.0 |
| DIP_10010478 | putative fad binding domain protein [Phaeomoniella chlamydospora] | 31.2 | 1.0E-49 |
| DIP_10010479 | NA | | |
| DIP_10010480 | putative ubx domain protein [Eutypa lata UCREL1] | 56.7 | 0.0 |
| DIP_10010481 | hypothetical protein PV09_04661 [Verruconis gallopava] | 56.6 | 5.0E-53 |
| DIP_10010482 | putative proteasome component protein [Eutypa lata UCREL1] | 60.8 | 0.0 |
| DIP_10010483 | hypothetical protein PFICI_00659 [Pestalotiopsis fici W106-1] | 79.8 | 0.0 |
| DIP_10010484 | hypothetical protein GGTG_02052 [Gaeumannomyces graminis var. tritici R3-111a-1] | 50.7 | 2.0E-62 |
| DIP_10010485 | hypothetical protein UCREL1_7284 [Eutypa lata UCREL1] | 46.6 | 7.0E-43 |
| DIP_10010486 | putative nucleoporin nup49 nsp49 protein [Eutypa lata UCREL1] | 63.1 | 1.0E-166 |
| DIP_10010487 | hypothetical protein OIDMADRAFT_29469 [Oidiodendron maius Zn] | 54.9 | 2.0E-86 |
| DIP_10010488 | hypothetical protein A1O5_04748 [Cladophialophora psammophila CBS 110553] | 47.2 | 4.0E-54 |
| DIP_10010489 | hypothetical protein THITE_2115604 [Thielavia terrestris NRRL 8126] | 61 | 5.0E-121 |
| DIP_10010490 | hypothetical protein FGSG_04210 [Fusarium graminearum PH-1] | 31.6 | 7.0E-11 |
| DIP_10010491 | putative xap5 domain-containing protein [Eutypa lata UCREL1] | 53.7 | 1.0E-141 |
| DIP_10010492 | NA | | |
| DIP_10010493 | NA | | |
| DIP_10010494 | hypothetical protein PFICI_12835 [Pestalotiopsis fici W106-1] | 66 | 0.0 |
| DIP_10010495 | putative duf890 domain-containing protein [Eutypa lata UCREL1] | 39.3 | 2.0E-108 |
| DIP_10010496 | hypothetical protein PFICI_14590 [Pestalotiopsis fici W106-1] | 62.1 | 2.0E-100 |
| DIP_10010497 | putative pyridoxine biosynthesis protein [Eutypa lata UCREL1] | 86.9 | 0.0 |
| DIP_10010498 | hypothetical protein PFICI_14574 [Pestalotiopsis fici W106-1] | 76.4 | 0.0 |
| DIP_10010499 | HET-C2 protein [Magnaporthe oryzae 70-15] | 75.5 | 3.0E-116 |
| DIP_10010500 | hypothetical protein UCREL1_4149 [Eutypa lata UCREL1] | 45.4 | 2.0E-64 |
| DIP_10010501 | putative glycogen debranching enzyme protein [Eutypa lata UCREL1] | 76 | 0.0 |
| DIP_10010502 | hypothetical protein V493_08047 [Pseudogymnoascus pannorum VKM F-4281 (FW-2241)] | 51.4 | 4.0E-19 |
| DIP_10010503 | putative glycerol-3-phosphate dehydrogenase protein [Eutypa lata UCREL1] | 75.9 | 0.0 |
| DIP_10010504 | putative gliotoxin biosynthesis protein GliK [Colletotrichum sublineola] | 40.5 | 2.0E-81 |
| DIP_10010505 | NA | | |
| DIP_10010506 | NA | | |
| DIP_10010507 | putative hydantoinase protein [Eutypa lata UCREL1] | 64.4 | 0.0 |
| DIP_10010508 | hypothetical protein ACRE_037320 [Acremonium chrysogenum ATCC 11550] | 38.2 | 1.0E-84 |
| DIP_10010509 | UDP-glucuronosyl/UDP-glucosyltransferase [Metarhizium guizhouense ARSEF 977] | 40.2 | 2.0E-108 |
| DIP_10010510 | putative protein kinase and ribonuclease protein [Eutypa lata UCREL1] | 63.1 | 0.0 |
| DIP_10010511 | NA | | |
| DIP_10010512 | putative delta fatty acid desaturase protein [Eutypa lata UCREL1] | 82 | 0.0 |
| DIP_10010513 | NA | | |
| DIP_10010514 | NA | | |
| DIP_10010515 | hypothetical protein MANI_026369 [Metarhizium anisopliae] | 65.5 | 1.0E-21 |
| DIP_10010516 | hypothetical protein UCREL1_980 [Eutypa lata UCREL1] | 51.2 | 1.0E-28 |
| DIP_10010517 | putative caspase domain-containing protein [Eutypa lata UCREL1] | 75.1 | 0.0 |
| DIP_10010518 | hypothetical protein UCREL1_2322 [Eutypa lata UCREL1] | 42.9 | 2.0E-64 |
| DIP_10010519 | hypothetical protein UCREL1_4887 [Eutypa lata UCREL1] | 65.7 | 0.0 |
| DIP_10010520 | general substrate transporter [Ophiostoma piceae UAMH 11346] | 72.4 | 0.0 |
| DIP_10010521 | putative oxidoreductase-like protein [Acremonium chrysogenum ATCC 11550] | 43 | 4.0E-71 |
| DIP_10010522 | putative poly(3-hydroxybutyrate) depolymerase protein [Eutypa lata UCREL1] | 46.7 | 2.0E-98 |
| DIP_10010523 | hypothetical protein BC1G_00199 [Botrytis cinerea B05.10] | 55.6 | 5.0E-69 |
| DIP_10010524 | hypothetical protein PFICI_01249 [Pestalotiopsis fici W106-1] | 64.3 | 4.0E-61 |
| DIP_10010525 | DnaJ domain-containing protein [Colletotrichum gloeosporioides Cg-14] | 71.7 | 2.0E-118 |
| DIP_10010526 | putative thioredoxin protein [Eutypa lata UCREL1] | 72.9 | 2.0E-168 |
| DIP_10010527 | hypothetical protein CGGC5_9085 [Colletotrichum gloeosporioides Nara gc5] | 74 | 2.0E-121 |
| DIP_10010528 | putative aaa family [Diaporthe ampelina] | 53.3 | 0.0 |
| DIP_10010529 | putative geranylgeranyl pyrophosphate synthetase [Colletotrichum sublineola] | 42.1 | 5.0E-108 |
| DIP_10010530 | putative presequence translocase-associated motor subunit protein [Eutypa lata UCREL1] | 56.7 | 9.0E-86 |
| DIP_10010531 | putative molybdenum cofactor synthesis protein [Eutypa lata UCREL1] | 49.8 | 7.0E-171 |
| DIP_10010532 | hypothetical protein UCRPA7_4529 [Togninia minima UCRPA7] | 50.9 | 1.0E-53 |
| DIP_10010533 | putative ulp1 protease family protein [Eutypa lata UCREL1] | 78.6 | 5.0E-120 |
| DIP_10010534 | putative rho guanyl nucleotide exchange [Diaporthe ampelina] | 61 | 0.0 |
| DIP_10010535 | hypothetical protein PFICI_05074 [Pestalotiopsis fici W106-1] | 64.6 | 3.0E-52 |
| DIP_10010536 | hypothetical protein UCDDS831_g03435 [Diplodia seriata] | 36.8 | 1.0E-11 |
| DIP_10010537 | hypothetical protein UCREL1_5241 [Eutypa lata UCREL1] | 52.5 | 7.0E-30 |
| DIP_10010538 | NA | | |
| DIP_10010539 | monooxygenase, putative [Metarhizium acridum CQMa 102] | 45.9 | 9.0E-119 |
| DIP_10010540 | putative horma domain-containing protein [Eutypa lata UCREL1] | 53.3 | 2.0E-105 |
| DIP_10010541 | putative importin-beta domain-containing protein [Eutypa lata UCREL1] | 62.8 | 0.0 |
| DIP_10010542 | hypothetical protein GLOINDRAFT_233146, partial [Rhizophagus irregularis DAOM 181602] | 52.9 | 8.0E-119 |
| DIP_10010543 | putative clathrin heavy chain protein [Eutypa lata UCREL1] | 73.3 | 0.0 |
| DIP_10010544 | hypothetical protein NEUTE1DRAFT_92235 [Neurospora tetrasperma FGSC 2508] | 83.5 | 6.0E-127 |
| DIP_10010545 | NA | | |
| DIP_10010546 | Pyruvate carboxylase [Pestalotiopsis fici W106-1] | 83.5 | 0.0 |
| DIP_10010547 | Putative UPF0041 domain-containing protein [Torrubiella hemipterigena] | 74.8 | 9.0E-75 |
| DIP_10010548 | putative lem domain-containing protein [Eutypa lata UCREL1] | 78.1 | 3.0E-64 |
| DIP_10010549 | 60S ribosomal protein L11 [Sporothrix schenckii ATCC 58251] | 94.7 | 3.0E-119 |
| DIP_10010550 | NA | | |
| DIP_10010551 | hypothetical protein SNOG_15374 [Phaeosphaeria nodorum SN15] | 36.8 | 1.0E-69 |
| DIP_10010552 | putative uv-damaged dna-binding protein [Eutypa lata UCREL1] | 68.7 | 0.0 |
| DIP_10010553 | NA | | |
| DIP_10010554 | Cytochrome b-c1 complex subunit Rieske [Pestalotiopsis fici W106-1] | 85.5 | 9.0E-149 |
| DIP_10010555 | hypothetical protein PFICI_04897 [Pestalotiopsis fici W106-1] | 57.6 | 1.0E-130 |
| DIP_10010556 | hypothetical protein PFICI_05080 [Pestalotiopsis fici W106-1] | 38.7 | 9.0E-90 |
| DIP_10010557 | putative t-complex protein 1 subunit alpha protein [Eutypa lata UCREL1] | 91.5 | 0.0 |
| DIP_10010558 | prolyl-tRNA synthetase [Colletotrichum graminicola M1.001] | 80.8 | 0.0 |
| DIP_10010559 | putative eukaryotic translation initiation factor 2 subunit gamma protein [Eutypa lata UCREL1] | 92.6 | 0.0 |
| DIP_10010560 | hypothetical protein PFICI_01560 [Pestalotiopsis fici W106-1] | 54.9 | 2.0E-155 |
| DIP_10010561 | hypothetical protein CSUB01_00380 [Colletotrichum sublineola] | 48.8 | 0.0 |
| DIP_10010562 | mitochondrial ATPase inhibitor, putative [Beauveria bassiana ARSEF 2860] | 62 | 7.0E-36 |
| DIP_10010563 | NA | | |
| DIP_10010564 | DNA damage checkpoint protein rad24 [Colletotrichum orbiculare MAFF 240422] | 82.4 | 3.0E-149 |
| DIP_10010565 | putative diphthamide biosynthesis protein 4 protein [Eutypa lata UCREL1] | 43.1 | 4.0E-57 |
| DIP_10010566 | hypothetical protein PFICI_01555 [Pestalotiopsis fici W106-1] | 77.9 | 9.0E-90 |
| DIP_10010567 | putative pre-mrna-splicing atp-dependent rna helicase prp28 protein [Eutypa lata UCREL1] | 71.4 | 0.0 |
| DIP_10010568 | hypothetical protein CLUG_01230 [Clavispora lusitaniae ATCC 42720] | 52.9 | 3.0E-79 |
| DIP_10010569 | putative aminopeptidase 2 protein [Eutypa lata UCREL1] | 87 | 0.0 |
| DIP_10010570 | putative nitrate transporter protein [Eutypa lata UCREL1] | 65 | 0.0 |
| DIP_10010571 | putative gpi anchored serine-threonine rich protein [Eutypa lata UCREL1] | 48.6 | 3.0E-54 |
| DIP_10010572 | putative poly -binding protein [Eutypa lata UCREL1] | 76.7 | 0.0 |
| DIP_10010573 | hypothetical protein TRIVIDRAFT_43213 [Trichoderma virens Gv29-8] | 47.2 | 4.0E-101 |
| DIP_10010574 | hypothetical protein COCMIDRAFT_34178 [Bipolaris oryzae ATCC 44560] | 45.4 | 5.0E-128 |
| DIP_10010575 | hypothetical protein COCMIDRAFT_109392 [Bipolaris oryzae ATCC 44560] | 44.3 | 6.0E-97 |
| DIP_10010576 | dUTP pyrophosphatase [Capronia epimyces CBS 606.96] | 71.7 | 7.0E-85 |
| DIP_10010577 | hypothetical protein PFICI_01767 [Pestalotiopsis fici W106-1] | 61.5 | 2.0E-176 |
| DIP_10010578 | putative proliferating cell nuclear antigen protein [Eutypa lata UCREL1] | 86 | 2.0E-174 |
| DIP_10010579 | putative proliferating cell nuclear antigen protein [Eutypa lata UCREL1] | 70.2 | 0.0 |
| DIP_10010580 | hypothetical protein PFICI_01770 [Pestalotiopsis fici W106-1] | 62.7 | 2.0E-56 |
| DIP_10010581 | hypothetical protein PFICI_01771 [Pestalotiopsis fici W106-1] | 40 | 1.0E-77 |
| DIP_10010582 | putative plasma membrane snare protein [Eutypa lata UCREL1] | 47.9 | 3.0E-111 |
| DIP_10010583 | Acetate non-utilizing protein 9 [Pestalotiopsis fici W106-1] | 73.6 | 5.0E-64 |
| DIP_10010584 | putative helicase associated domain-containing protein [Eutypa lata UCREL1] | 61.1 | 0.0 |
| DIP_10010585 | putative sterol desaturase [Aspergillus ruber CBS 135680] | 77.3 | 0.0 |
| DIP_10010586 | NA | | |
| DIP_10010587 | hypothetical protein HIM_11912 [Hirsutella minnesotensis 3608] | 46.4 | 0.0 |
| DIP_10010588 | hypothetical protein PFICI_01259 [Pestalotiopsis fici W106-1] | 47.7 | 1.0E-172 |
| DIP_10010589 | hypothetical protein MGG_11737 [Magnaporthe oryzae 70-15] | 59.9 | 4.0E-69 |
| DIP_10010590 | hypothetical protein PFICI_01261 [Pestalotiopsis fici W106-1] | 40.4 | 2.0E-93 |
| DIP_10010591 | hypothetical protein PFICI_01263 [Pestalotiopsis fici W106-1] | 53.8 | 0.0 |
| DIP_10010592 | Chitin synthase 2 [Scedosporium apiospermum] | 68.1 | 0.0 |
| DIP_10010593 | NA | | |
| DIP_10010594 | NA | | |
| DIP_10010595 | FAD dependent oxidoreductase-like protein [Myceliophthora thermophila ATCC 42464] | 46.4 | 2.0E-80 |
| DIP_10010596 | NA | | |
| DIP_10010597 | cytochrome P450 monooxygenase-like protein [Myceliophthora thermophila ATCC 42464] | 48.1 | 7.0E-157 |
| DIP_10010598 | hypothetical protein CHGG_09441 [Chaetomium globosum CBS 148.51] | 30.1 | 5.0E-44 |
| DIP_10010599 | hypothetical protein CHGG_09441 [Chaetomium globosum CBS 148.51] | 48.2 | 4.0E-141 |
| DIP_10010600 | NA | | |
| DIP_10010601 | GATA transcription factor AreB [Metarhizium anisopliae] | 44.9 | 4.0E-63 |
| DIP_10010602 | hypothetical protein TRIATDRAFT_300850 [Trichoderma atroviride IMI 206040] | 76.2 | 0.0 |
| DIP_10010603 | putative rrna processing protein [Eutypa lata UCREL1] | 73.7 | 0.0 |
| DIP_10010604 | putative dna-binding protein hgh1 protein [Togninia minima UCRPA7] | 65.4 | 4.0E-171 |
| DIP_10010605 | putative inorganic phosphate transporter pho84 protein [Eutypa lata UCREL1] | 79.7 | 0.0 |
| DIP_10010606 | hypothetical protein THITE_2106359 [Thielavia terrestris NRRL 8126] | 53.7 | 9.0E-144 |
| DIP_10010607 | putative cytochrome b2 protein [Eutypa lata UCREL1] | 83.2 | 0.0 |
| DIP_10010608 | NA | | |
| DIP_10010609 | secretion-related small GTPase [Metarhizium robertsii ARSEF 23] | 66.6 | 2.0E-167 |
| DIP_10010610 | hypothetical protein TRIATDRAFT_136464 [Trichoderma atroviride IMI 206040] | 34.9 | 2.0E-18 |
| DIP_10010611 | hypothetical protein S7711_03584 [Stachybotrys chartarum IBT 7711] | 58.2 | 1.0E-62 |

| | | | |
|---|---|---|---|
| DIP_10010612 | Methylsterol monooxygenase [Pestalotiopsis fici W106-1] | 88.4 | 0.0 |
| DIP_10010613 | NA | | |
| DIP_10010614 | hypothetical protein PFICI_05721 [Pestalotiopsis fici W106-1] | 53.7 | 2.0E-45 |
| DIP_10010615 | helicase mug81 [Pestalotiopsis fici W106-1] | 79.6 | 0.0 |
| DIP_10010616 | Synaptobrevin-like ykt6 [Hirsutella minnesotensis 3608] | 81.7 | 4.0E-118 |
| DIP_10010617 | NA | | |
| DIP_10010618 | putative sister chromatid cohesion protein mis4 protein [Eutypa lata UCREL1] | 63.1 | 0.0 |
| DIP_10010619 | putative transcription factor s-ii protein [Eutypa lata UCREL1] | 78.1 | 8.0E-66 |
| DIP_10010620 | Deoxyhypusine hydroxylase [Pestalotiopsis fici W106-1] | 77.4 | 1.0E-180 |
| DIP_10010621 | putative pyridoxamine 5 -phosphate oxidase protein [Eutypa lata UCREL1] | 63.6 | 1.0E-121 |
| DIP_10010622 | NA | | |
| DIP_10010623 | hypothetical protein PFICI_04450 [Pestalotiopsis fici W106-1] | 58.6 | 0.0 |
| DIP_10010624 | putative f-box domain-containing protein [Eutypa lata UCREL1] | 43.4 | 7.0E-111 |
| DIP_10010625 | hypothetical protein PFICI_04281 [Pestalotiopsis fici W106-1] | 59 | 0.0 |
| DIP_10010626 | hypothetical protein UCREL1_10651 [Eutypa lata UCREL1] | 53.3 | 2.0E-75 |
| DIP_10010627 | hypothetical protein UCREL1_10650 [Eutypa lata UCREL1] | 44.8 | 9.0E-74 |
| DIP_10010628 | NA | | |
| DIP_10010629 | rad55 protein [Colletotrichum orbiculare MAFF 240422] | 50.9 | 7.0E-90 |
| DIP_10010630 | DNA-binding protein creA [Pestalotiopsis fici W106-1] | 69 | 0.0 |
| DIP_10010631 | putative revertant of glycogen synthase kinase mutation protein [Eutypa lata UCREL1] | 60.4 | 0.0 |
| DIP_10010632 | hypothetical protein GLRG_08332 [Colletotrichum graminicola M1.001] | 44.7 | 7.0E-128 |
| DIP_10010633 | NA | | |
| DIP_10010634 | NA | | |
| DIP_10010635 | Cocaine esterase [Fusarium oxysporum f. sp. cubense race 1] | 61.8 | 0.0 |
| DIP_10010636 | putative arf gtpase-activating protein [Eutypa lata UCREL1] | 83.4 | 0.0 |
| DIP_10010637 | Mitochondrial import inner membrane translocase subunit tim-21 [Pestalotiopsis fici W106-1] | 54.2 | 2.0E-78 |
| DIP_10010638 | NA | | |
| DIP_10010639 | NA | | |
| DIP_10010640 | hypothetical protein PFICI_05162 [Pestalotiopsis fici W106-1] | 70.2 | 0.0 |
| DIP_10010641 | putative 2-nitropropane dioxygenase protein [Eutypa lata UCREL1] | 79.6 | 0.0 |
| DIP_10010642 | putative aquaporin protein [Eutypa lata UCREL1] | 67.8 | 1.0E-122 |
| DIP_10010643 | putative short chain dehydrogenase reductase family protein [Togninia minima UCRPA7] | 73.2 | 6.0E-158 |
| DIP_10010644 | NA | | |
| DIP_10010645 | hypothetical protein FVEG_06314 [Fusarium verticillioides 7600] | 43.3 | 3.0E-109 |
| DIP_10010646 | NA | | |
| DIP_10010647 | ion channel, putative [Colletotrichum orbiculare MAFF 240422] | 56.4 | 1.0E-70 |
| DIP_10010648 | NA | | |
| DIP_10010649 | NA | | |
| DIP_10010650 | NA | | |
| DIP_10010651 | NA | | |
| DIP_10010652 | NA | | |
| DIP_10010653 | NA | | |
| DIP_10010654 | NA | | |
| DIP_10010655 | putative prenyltransferase and squalene oxidase protein [Eutypa lata UCREL1] | 47.4 | 4.0E-127 |
| DIP_10010656 | hypothetical protein UCREL1_10507 [Eutypa lata UCREL1] | 31.7 | 9.0E-75 |
| DIP_10010657 | hypothetical protein UCDDA912_g08222 [Diaporthe ampelina] | 54.8 | 7.0E-60 |
| DIP_10010658 | NA | | |
| DIP_10010659 | hypothetical protein UCREL1_11361 [Eutypa lata UCREL1] | 37.9 | 8.0E-17 |
| DIP_10010660 | hypothetical protein MGG_11091 [Magnaporthe oryzae 70-15] | 43.8 | 7.0E-52 |
| DIP_10010661 | putative c2 domain-containing protein [Eutypa lata UCREL1] | 60.9 | 0.0 |
| DIP_10010662 | homocysteine s-methyltransferase [Colletotrichum gloeosporioides Nara gc5] | 50 | 8.0E-101 |
| DIP_10010663 | NA | | |
| DIP_10010664 | putative s-adenosyl-methionine-sterol-c- methyltransferas protein [Eutypa lata UCREL1] | 72.4 | 0.0 |
| DIP_10010665 | NA | | |
| DIP_10010666 | hypothetical protein PFICI_00188 [Pestalotiopsis fici W106-1] | 50 | 2.0E-170 |
| DIP_10010667 | hypothetical protein PFICI_01649 [Pestalotiopsis fici W106-1] | 35.9 | 2.0E-22 |
| DIP_10010668 | hypothetical protein PFICI_01654 [Pestalotiopsis fici W106-1] | 45.6 | 2.0E-36 |
| DIP_10010669 | Cytochrome c oxidase-assembly factor cox-23 [Pestalotiopsis fici W106-1] | 77.6 | 2.0E-36 |
| DIP_10010670 | hypothetical protein CMQ_255 [Grosmannia clavigera kw1407] | 31.6 | 2.0E-53 |
| DIP_10010671 | ATP synthase subunit gamma [Pestalotiopsis fici W106-1] | 78 | 5.0E-167 |
| DIP_10010672 | putative topoisomerase i protein [Eutypa lata UCREL1] | 73.3 | 0.0 |
| DIP_10010673 | putative glycoside hydrolase family 16 protein [Eutypa lata UCREL1] | 65.1 | 5.0E-134 |
| DIP_10010674 | putative dna-directed rna polymerase protein [Eutypa lata UCREL1] | 62.7 | 6.0E-90 |
| DIP_10010675 | hypothetical protein BC1G_12209 [Botrytis cinerea B05.10] | 77.4 | 2.0E-138 |
| DIP_10010676 | putative dnase1 protein [Eutypa lata UCREL1] | 51.7 | 8.0E-47 |
| DIP_10010677 | Peptidase M20 domain-containing protein [Pestalotiopsis fici W106-1] | 57.8 | 6.0E-155 |
| DIP_10010678 | putative methylthioribose-1-phosphate isomerase protein [Eutypa lata UCREL1] | 75.5 | 0.0 |
| DIP_10010679 | putative tmem1 family [Diaporthe ampelina] | 53.6 | 0.0 |
| DIP_10010680 | hypothetical protein UCREL1_5485 [Eutypa lata UCREL1] | 56.1 | 2.0E-68 |
| DIP_10010681 | putative swib mdm2 domain protein [Eutypa lata UCREL1] | 66.3 | 8.0E-110 |
| DIP_10010682 | putative C2H2 finger domain protein [Metarhizium anisopliae] | 55.7 | 0.0 |
| DIP_10010683 | NA | | |
| DIP_10010684 | mitochondrial phosphate carrier protein [Ophiostoma piceae UAMH 11346] | 82.1 | 0.0 |
| DIP_10010685 | hypothetical protein UCREL1_5490 [Eutypa lata UCREL1] | 52.7 | 3.0E-113 |
| DIP_10010686 | putative cytochrome p450 protein [Togninia minima UCRPA7] | 78 | 0.0 |
| DIP_10010687 | hypothetical protein FOC4_g10009697 [Fusarium oxysporum f. sp. cubense race 4] | 55.3 | 2.0E-132 |
| DIP_10010688 | hypothetical protein FG05_03706 [Fusarium graminearum] | 71.8 | 0.0 |
| DIP_10010689 | NA | | |
| DIP_10010690 | hypothetical protein FOTG_14031 [Fusarium oxysporum f. sp. vasinfectum 25433] | 63.4 | 0.0 |
| DIP_10010691 | hypothetical protein FAVG1_09360 [Fusarium avenaceum] | 75.7 | 0.0 |
| DIP_10010692 | trichothecene biosynthesis acetyltransferase, putative [Aspergillus clavatus NRRL 1] | 50.2 | 3.0E-175 |
| DIP_10010693 | NA | | |
| DIP_10010694 | putative atp synthase regulation protein nca2 protein [Eutypa lata UCREL1] | 68.9 | 0.0 |
| DIP_10010695 | putative rab gtpase protein [Eutypa lata UCREL1] | 93.1 | 4.0E-138 |
| DIP_10010696 | hypothetical protein PFICI_05715 [Pestalotiopsis fici W106-1] | 34.3 | 2.0E-42 |
| DIP_10010697 | putative aaa family atpase protein [Eutypa lata UCREL1] | 52.7 | 0.0 |
| DIP_10010698 | putative serine threonine-protein phosphatase pp2a catalytic subunit protein [Togninia minima UCRPA7] | 79 | 1.0E-121 |
| DIP_10010699 | hypothetical protein PV05_07812 [Exophiala xenobiotica] | 43.5 | 1.0E-78 |
| DIP_10010700 | hypothetical protein AURDEDRAFT_164322 [Auricularia delicata TFB-10046 SS5] | 55.4 | 7.0E-24 |

a Data predicted by Sushida et al. (2019). Gene descriptions are derived from non-redundant gene prediction results.

Supplementary Table S4. Predicted gene clusters involved in secondary metabolite biosynthesis in *Hansfordia pulvinata* .

| Region [a] | GeneID | *In vitro Cf/Hp* log2 Fold-change [b] | *In planta Cf/Hp* log2 Fold-change [b] | Length: DNA | Length: No. amino acids | Description | Type [c] |
|---|---|---|---|---|---|---|---|
| 1.5 | DIP_10010298 | 1.11 | | 1521 | 506 | other | fungal-RiPP-like |
| | DIP_10010299 | | -2.60 | 192 | 63 | other | |
| | DIP_10010300 | | 0.84 | 1815 | 604 | biosynthetic-additional | |
| | DIP_10010301 | | 2.51 | 1881 | 626 | other | |
| | DIP_10010302 | | -0.52 | 1884 | 627 | other | |
| | DIP_10010303 | 3.13 | 3.59 | 648 | 215 | other | |
| | DIP_10010304 | 2.64 | 3.54 | 918 | 305 | biosynthetic | |
| | DIP_10010305 | 2.01 | 3.18 | 1302 | 433 | other | |
| | DIP_10010306 | | | 738 | 245 | other | |
| | DIP_10010307 | | 0.59 | 402 | 133 | other | |
| | DIP_10010308 | -0.59 | 0.79 | 465 | 154 | other | |
| | DIP_10010309 | | 1.25 | 513 | 170 | other | |
| | DIP_10010310 | | | 1443 | 480 | other | |
| 23.1 | DIP_10000313 | | 1.50 | 1122 | 373 | other | terpene |
| | DIP_10000314 | | -0.71 | 990 | 329 | biosynthetic | |
| 15.1 | DIP_10001427 | 0.57 | | 2817 | 938 | other | T1PKS |
| | DIP_10001428 | -1.28 | -1.27 | 1590 | 529 | other | |
| | DIP_10001429 | | 0.70 | 207 | 68 | other | |
| | DIP_10001430 | | -1.64 | 1344 | 447 | biosynthetic-additional | |
| | DIP_10001431 | | | 1632 | 543 | biosynthetic-additional | |
| | DIP_10001432 | | | 1539 | 512 | biosynthetic-additional | |
| | DIP_10001433 | | | 774 | 257 | biosynthetic-additional | |
| | DIP_10001434 | | -0.70 | 8424 | 2807 | biosynthetic | |
| | DIP_10001435 | | 2.79 | 1236 | 411 | other | |
| | DIP_10001436 | | 4.57 | 1416 | 471 | biosynthetic-additional | |
| | DIP_10001437 | | | 3150 | 1049 | other | |
| | DIP_10001438 | -0.95 | -2.90 | 921 | 306 | other | |
| 3.1 | DIP_10008267 | | 1.91 | 252 | 83 | other | terpene |
| | DIP_10008268 | | | 1059 | 352 | biosynthetic | |
| | DIP_10008269 | | | 447 | 148 | other | |
| | DIP_10008270 | | | 1002 | 333 | biosynthetic-additional | |
| | DIP_10008271 | | | 2484 | 827 | biosynthetic-additional | |
| 1.4 | DIP_10010170 | 2.19 | | 438 | 145 | other | NRPS-like |
| | DIP_10010171 | | | 234 | 77 | other | |
| | DIP_10010172 | 0.71 | 0.91 | 903 | 300 | other | |
| | DIP_10010173 | | | 120 | 39 | other | |
| | DIP_10010174 | | 0.78 | 675 | 224 | other | |
| | DIP_10010175 | 1.25 | 1.98 | 1389 | 462 | other | |
| | DIP_10010176 | | -0.60 | 1557 | 518 | other | |
| | DIP_10010177 | | -1.15 | 1692 | 563 | other | |
| | DIP_10010178 | 4.95 | 1.06 | 3051 | 1016 | biosynthetic | |
| | DIP_10010179 | 3.60 | | 1164 | 387 | other | |
| | DIP_10010180 | 8.40 | 4.40 | 951 | 316 | other | |
| | DIP_10010181 | 7.68 | | 1386 | 461 | biosynthetic-additional | |
| 1.6 | DIP_10010435 | | | 477 | 158 | other | NRPS-like |
| | DIP_10010436 | 2.66 | 4.48 | 1725 | 574 | other | |
| | DIP_10010437 | | | 1299 | 432 | other | |
| | DIP_10010438 | | 0.61 | 1002 | 333 | other | |
| | DIP_10010439 | | | 3162 | 1053 | biosynthetic | |
| | DIP_10010440 | | -0.63 | 1701 | 566 | other | |
| | DIP_10010441 | | -2.58 | 2250 | 749 | biosynthetic-additional | |
| | DIP_10010442 | 1.82 | 4.11 | 648 | 215 | other | |
| | DIP_10010443 | | | 1296 | 431 | other | |
| 3.2 | DIP_10008464 | | 5.23 | 846 | 281 | biosynthetic-additional | T1PKS |
| | DIP_10008465 | | | 705 | 234 | other | |
| | DIP_10008466 | | | 258 | 85 | other | |
| | DIP_10008467 | | -1.08 | 6423 | 2140 | biosynthetic | |
| | DIP_10008468 | | -4.97 | 1710 | 569 | other | |
| | DIP_10008469 | | -0.82 | 813 | 270 | other | |
| | DIP_10008470 | | | 2655 | 884 | other | |
| 1.3 | DIP_10009629 | | -1.83 | 1467 | 488 | transport | T1PKS |
| | DIP_10009630 | | 2.17 | 1734 | 577 | other | |
| | DIP_10009631 | | 2.95 | 801 | 266 | other | |
| | DIP_10009632 | | | 570 | 189 | other | |
| | DIP_10009633 | | | 1326 | 441 | other | |
| | DIP_10009634 | | 0.59 | 2373 | 790 | other | |
| | DIP_10009635 | -2.97 | -3.16 | 7254 | 2417 | biosynthetic | |
| | DIP_10009636 | | | 1212 | 403 | other | |
| | DIP_10009637 | | | 399 | 132 | other | |
| | DIP_10009638 | | | 1302 | 433 | other | |
| | DIP_10009639 | | | 906 | 301 | other | |
| | DIP_10009640 | | | 2325 | 774 | other | |
| | DIP_10009641 | | -0.62 | 1419 | 472 | other | |
| 1.2 | DIP_10009615 | 2.72 | | 2355 | 784 | other | NRPS |
| | DIP_10009616 | | -1.73 | 942 | 313 | other | |
| | DIP_10009617 | | | 1587 | 528 | other | |
| | DIP_10009618 | | -4.41 | 1173 | 390 | other | |
| | DIP_10009619 | | | 618 | 205 | other | |
| | DIP_10009620 | -1.35 | -2.87 | 5406 | 1801 | biosynthetic | |
| | DIP_10009621 | 6.38 | | 819 | 272 | other | |
| | DIP_10009622 | | | 855 | 284 | other | |
| | DIP_10009623 | | | 990 | 329 | other | |
| | DIP_10009624 | | | 1869 | 622 | biosynthetic-additional | |
| | DIP_10009625 | -3.00 | -1.32 | 1608 | 535 | other | |
| | DIP_10009626 | -1.39 | | 867 | 288 | other | |
| 1.1 | DIP_10009607 | | | 207 | 68 | other | NRPS-like |
| | DIP_10009608 | 1.81 | 2.50 | 675 | 224 | other | |
| | DIP_10009609 | | | 1320 | 439 | other | |
| | DIP_10009610 | 2.20 | -4.25 | 3249 | 1082 | biosynthetic | |
| | DIP_10009611 | | -0.55 | 1575 | 524 | other | |
| | DIP_10009612 | | -0.34 | 2997 | 998 | other | |
| | DIP_10009613 | -1.20 | -1.49 | 3195 | 1064 | other | |
| 5.2 | DIP_10006580 | 4.25 | 3.53 | 2166 | 721 | transport | T1PKS |
| | DIP_10006581 | 1.18 | 0.63 | 3009 | 1002 | biosynthetic-additional | |
| | DIP_10006582 | 5.44 | 5.44 | 972 | 323 | other | |
| | DIP_10006583 | 3.77 | 3.92 | 528 | 175 | other | |
| | DIP_10006584 | 0.78 | | 8463 | 2820 | biosynthetic | |
| | DIP_10006585 | -0.83 | | 2346 | 781 | other | |
| | DIP_10006586 | -1.38 | | 837 | 278 | biosynthetic-additional | |
| | DIP_10006587 | | -1.79 | 1932 | 643 | biosynthetic-additional | |
| | DIP_10006588 | | -3.12 | 1152 | 383 | biosynthetic-additional | |
| | DIP_10006589 | -1.19 | -2.14 | 798 | 265 | other | |
| | DIP_10006590 | | -2.23 | 1383 | 460 | other | |
| | DIP_10006591 | | 0.45 | 1815 | 604 | other | |
| | DIP_10006592 | | | 1098 | 365 | other | |
| 13.2 | DIP_10001961 | | | 687 | 228 | other | fungal-RiPP-like |
| | DIP_10001962 | | | 336 | 111 | other | |
| | DIP_10001963 | | | 450 | 149 | other | |
| | DIP_10001964 | | | 312 | 103 | other | |
| | DIP_10001965 | | | 1089 | 362 | other | |
| | DIP_10001966 | 3.60 | 2.59 | 1752 | 583 | other | |
| | DIP_10001967 | 3.42 | 2.24 | 1326 | 441 | other | |
| | DIP_10001968 | | | 201 | 66 | other | |
| | DIP_10001969 | | | 864 | 287 | biosynthetic | |
| | DIP_10001970 | | | 1713 | 570 | transport | |
| | DIP_10001971 | | | 867 | 288 | other | |
| | DIP_10001972 | 2.17 | -4.54 | 1959 | 652 | other | |
| | DIP_10001973 | | | 138 | 45 | other | |
| | DIP_10001974 | | | 2754 | 917 | other | |
| | DIP_10001975 | -1.29 | -1.33 | 1248 | 415 | other | |
| 13.3 | DIP_10002005 | 1.80 | 1.02 | 1605 | 534 | biosynthetic-additional | terpene |
| | DIP_10002006 | 2.04 | 1.79 | 1716 | 571 | biosynthetic | |
| | DIP_10002007 | | | 1119 | 372 | other | |
| | DIP_10002008 | | -0.65 | 2001 | 666 | other | |
| 5.1 | DIP_10006269 | | 1.44 | 2283 | 760 | other | terpene |
| | DIP_10006270 | | | 1683 | 560 | biosynthetic-additional | |
| | DIP_10006271 | | 2.01 | 1143 | 380 | biosynthetic | |
| | DIP_10006272 | | -0.94 | 870 | 289 | other | |
| | DIP_10006273 | 0.82 | 1.53 | 588 | 195 | other | |
| 13.1 | DIP_10001953 | 1.81 | 1.03 | 1209 | 402 | other | terpene |
| | DIP_10001954 | 2.58 | 1.43 | 1173 | 390 | biosynthetic (*HpDPH1* ) | |
| | DIP_10001955 | 2.27 | 1.22 | 969 | 322 | osynthetic-additional (*HpDPH*. | |
| | DIP_10001956 | 1.96 | | 1488 | 495 | osynthetic-additional (*HpDPH*. | |
| 9.1 | DIP_10003972 | | | 1305 | 434 | biosynthetic-additional | terpene |
| | DIP_10003973 | | | 1557 | 518 | other | |
| | DIP_10003974 | | | 1038 | 345 | biosynthetic-additional | |
| | DIP_10003975 | | 1.28 | 1428 | 475 | biosynthetic | |
| | DIP_10003976 | | -1.35 | 939 | 312 | other | |
| | DIP_10003977 | | -2.41 | 1485 | 494 | transport | |
| | DIP_10003978 | | -2.57 | 729 | 242 | biosynthetic-additional | |
| 11.1 | DIP_10002579 | | 0.73 | 1368 | 455 | other | T1PKS |
| | DIP_10002580 | | | 591 | 196 | other | |
| | DIP_10002581 | | 1.14 | 579 | 192 | other | |
| | DIP_10002582 | | | 1182 | 393 | other | |
| | DIP_10002583 | 1.35 | 2.79 | 1533 | 510 | other | |
| | DIP_10002584 | 3.20 | | 3627 | 1208 | biosynthetic-additional | |
| | DIP_10002585 | | | 2559 | 852 | biosynthetic | |
| | DIP_10002586 | | | 504 | 167 | other | |
| | DIP_10002587 | | | 1080 | 359 | other | |
| | DIP_10002588 | | | 597 | 198 | other | |
| | DIP_10002589 | | | 360 | 119 | other | |
| 27.1 | DIP_10000196 | | 6.89 | 180 | 59 | other | isocyanide |
| | DIP_10000197 | 10.86 | 13.64 | 390 | 129 | other | |
| | DIP_10000198 | | | 1221 | 406 | other | |
| | DIP_10000199 | | | 1566 | 521 | transport | |
| | DIP_10000200 | | 6.29 | 492 | 163 | other | |
| | DIP_10000201 | 5.69 | 10.98 | 324 | 107 | biosynthetic-additional | |
| | DIP_10000202 | | 8.25 | 144 | 47 | other | |
| | DIP_10000203 | | 8.83 | 1962 | 653 | biosynthetic | |
| | DIP_10000204 | | 10.97 | 1365 | 454 | biosynthetic-additional | |
| | DIP_10000205 | | 7.34 | 1548 | 515 | transport | |
| | DIP_10000206 | | 4.00 | 861 | 286 | other | |
| | DIP_10000207 | | 6.51 | 1617 | 538 | other | |
| | DIP_10000208 | | | 912 | 303 | other | |
| | DIP_10000209 | | | 1158 | 385 | other | |
| | DIP_10000210 | 2.03 | 4.27 | 330 | 109 | other | |
| 11.2 | DIP_10002604 | | 0.63 | 1995 | 664 | other | T1PKS |
| | DIP_10002605 | | -1.96 | 1458 | 485 | other | |
| | DIP_10002606 | | -3.10 | 2484 | 827 | other | |
| | DIP_10002607 | 2.67 | 0.60 | 8202 | 2733 | biosynthetic | |
| | DIP_10002608 | 3.28 | 1.53 | 1842 | 613 | biosynthetic-additional | |
| | DIP_10002609 | | | 1167 | 388 | other | |
| | DIP_10002610 | | -0.54 | 2460 | 819 | other | |
| | DIP_10002611 | | 1.34 | 1548 | 515 | other | |
| 2.2 | DIP_10009157 | | 1.62 | 1875 | 624 | transport | NRPS-like |
| | DIP_10009158 | | | 429 | 142 | other | |
| | DIP_10009159 | | | 330 | 109 | other | |
| | DIP_10009160 | | 1.16 | 1188 | 395 | other | |
| | DIP_10009161 | | | 1059 | 352 | other | |
| | DIP_10009162 | | 0.89 | 4641 | 1546 | biosynthetic | |
| | DIP_10009163 | | | 201 | 66 | other | |
| | DIP_10009164 | -2.02 | -6.78 | 381 | 126 | other | |
| 2.3 | DIP_10009226 | | | 2382 | 793 | other | T1PKS |
| | DIP_10009227 | | | 1317 | 438 | other | |
| | DIP_10009228 | | | 726 | 241 | other | |
| | DIP_10009229 | | | 1050 | 349 | biosynthetic-additional | |
| | DIP_10009230 | | -1.86 | 963 | 320 | other | |
| | DIP_10009231 | | -0.90 | 456 | 151 | other | |
| | DIP_10009232 | | -0.94 | 1458 | 485 | other | |
| | DIP_10009233 | | -2.67 | 552 | 183 | other | |
| | DIP_10009234 | | -3.12 | 7317 | 2438 | biosynthetic | |
| | DIP_10009235 | | -2.07 | 1839 | 612 | biosynthetic-additional | |
| | DIP_10009236 | 0.62 | | 1446 | 481 | other | |
| | DIP_10009237 | | | 603 | 200 | other | |
| | DIP_10009238 | 6.14 | 10.38 | 1209 | 402 | other | |
| 2.1 | DIP_10008633 | | | 2178 | 725 | other | NRPS |
| | DIP_10008634 | | 0.99 | 2370 | 789 | other | |
| | DIP_10008635 | | | 960 | 319 | other | |
| | DIP_10008636 | | -2.89 | 1905 | 634 | other | |
| | DIP_10008637 | | 5.61 | 288 | 95 | other | |
| | DIP_10008638 | | -3.00 | 18618 | 6205 | biosynthetic | |
| | DIP_10008639 | 6.24 | 6.85 | 972 | 323 | other | |
| | DIP_10008640 | | 4.93 | 990 | 329 | other | |
| | DIP_10008641 | | | 4671 | 1556 | transport | |
| | DIP_10008642 | | | 924 | 307 | other | |
| 2.4 | DIP_10009258 | -0.62 | | 750 | 249 | other | fungal-RiPP-like |
| | DIP_10009259 | 3.64 | 0.94 | 1740 | 579 | other | |
| | DIP_10009260 | 0.56 | -1.70 | 1119 | 372 | other | |
| | DIP_10009261 | 1.38 | 3.19 | 507 | 168 | other | |
| | DIP_10009262 | | -1.26 | 906 | 301 | other | |
| | DIP_10009263 | 4.34 | 8.05 | 342 | 113 | other | |
| | DIP_10009264 | | | 885 | 294 | other | |
| | DIP_10009265 | | | 1185 | 394 | other | |
| | DIP_10009266 | 0.77 | 1.02 | 1737 | 578 | other | |
| | DIP_10009267 | | | 891 | 296 | biosynthetic | |
| | DIP_10009268 | | 2.25 | 1044 | 347 | other | |
| | DIP_10009269 | | 0.83 | 381 | 126 | other | |
| | DIP_10009270 | -0.87 | 0.52 | 432 | 143 | other | |
| | DIP_10009271 | -0.86 | | 990 | 329 | other | |
| | DIP_10009272 | | | 771 | 256 | other | |
| | DIP_10009273 | | | 1524 | 507 | other | |
| | DIP_10009274 | -1.00 | | 417 | 138 | other | |
| | DIP_10009275 | | 1.16 | 606 | 201 | other | |
| | DIP_10009276 | | | 1260 | 419 | other | |
| | DIP_10009277 | | | 1479 | 492 | other | |
| 16.1 | DIP_10000992 | | -5.39 | 1668 | 555 | transport | NRPS |
| | DIP_10000993 | | 0.39 | 579 | 192 | other | |
| | DIP_10000994 | | | 363 | 120 | other | |
| | DIP_10000995 | | 2.95 | 1044 | 347 | biosynthetic-additional | |
| | DIP_10000996 | 2.75 | 4.70 | 447 | 148 | other | |
| | DIP_10000997 | | -0.75 | 597 | 198 | other | |
| | DIP_10000998 | | | 1797 | 598 | other | |
| | DIP_10000999 | | | 321 | 106 | other | |
| | DIP_10001000 | -3.10 | -4.15 | 8247 | 2748 | biosynthetic | |
| | DIP_10001001 | | | 2415 | 804 | biosynthetic-additional | |
| | DIP_10001002 | | | 243 | 80 | other | |
| | DIP_10001003 | | 1.12 | 1617 | 538 | biosynthetic-additional | |
| 2.5 | DIP_10009427 | | -0.73 | 2670 | 889 | other | NRPS-like |
| | DIP_10009428 | | | 2136 | 711 | biosynthetic-additional | |
| | DIP_10009429 | | -1.89 | 1536 | 511 | transport | |
| | DIP_10009430 | | 0.98 | 975 | 324 | biosynthetic-additional | |
| | DIP_10009431 | | | 3120 | 1039 | biosynthetic | |
| | DIP_10009432 | -1.10 | | 849 | 282 | biosynthetic-additional | |
| | DIP_10009433 | | | 1896 | 631 | other | |
| | DIP_10009434 | | -0.67 | 2562 | 853 | other | |
| | DIP_10009435 | | | 414 | 137 | other | |
| | DIP_10009436 | | | 3306 | 1101 | other | |
| 24.1 | DIP_10000293 | | 1.52 | 1143 | 380 | other | T1PKS |
| | DIP_10000294 | 1.83 | 3.30 | 1902 | 633 | other | |
| | DIP_10000295 | | | 3828 | 1275 | biosynthetic | |
| | DIP_10000296 | | | 150 | 49 | other | |
| 12.1 | DIP_10002251 | 1.72 | 2.24 | 2139 | 712 | other | NRPS-like |
| | DIP_10002252 | | -1.42 | 5169 | 1722 | other | |
| | DIP_10002253 | | | 1404 | 467 | biosynthetic-additional | |
| | DIP_10002254 | | | 3591 | 1196 | biosynthetic | |
| | DIP_10002255 | | | 1374 | 457 | biosynthetic-additional | |
| | DIP_10002256 | | | 1875 | 624 | biosynthetic-additional | |
| | DIP_10002257 | | | 1905 | 634 | transport | |
| | DIP_10002258 | | -1.49 | 1725 | 574 | biosynthetic-additional | |
| | DIP_10002259 | | 3.71 | 1197 | 398 | other | |
| 4.1 | DIP_10007241 | | | 144 | 47 | other | NRPS-like |
| | DIP_10007242 | | | 1236 | 411 | other | |
| | DIP_10007243 | | 2.10 | 2271 | 756 | other | |
| | DIP_10007244 | -0.85 | -0.80 | 450 | 149 | other | |
| | DIP_10007245 | 0.98 | 1.60 | 1635 | 544 | biosynthetic-additional | |
| | DIP_10007246 | | | 819 | 272 | other | |
| | DIP_10007247 | | | 3903 | 1300 | biosynthetic | |
| | DIP_10007248 | | | 1320 | 439 | other | |
| | DIP_10007249 | -0.95 | | 1938 | 645 | biosynthetic-additional | |
| | DIP_10007250 | | -1.61 | 1680 | 559 | other | |
| | DIP_10007251 | | | 2367 | 788 | other | |
| 12.3 | DIP_10002395 | -0.90 | -0.85 | 564 | 187 | other | NRPS |
| | DIP_10002396 | -0.77 | -1.38 | 696 | 231 | other | |
| | DIP_10002397 | 2.67 | | 1659 | 552 | biosynthetic-additional | |
| | DIP_10002398 | 3.61 | -5.97 | 1002 | 333 | other | |
| | DIP_10002399 | 1.68 | 1.21 | 2169 | 722 | biosynthetic-additional | |
| | DIP_10002400 | | | 8469 | 2822 | biosynthetic | |
| | DIP_10002401 | | | 1893 | 630 | other | |
| | DIP_10002402 | | | 1383 | 460 | other | |
| | DIP_10002403 | | | 2367 | 788 | other | |
| 12.2 | DIP_10002373 | 3.27 | 1.97 | 1287 | 428 | other | NRPS |
| | DIP_10002374 | 1.08 | | 1098 | 365 | other | |
| | DIP_10002375 | -1.23 | -2.00 | 930 | 309 | other | |
| | DIP_10002376 | -1.54 | -2.67 | 1011 | 336 | biosynthetic-additional | |
| | DIP_10002377 | | 0.66 | 1929 | 642 | other | |
| | DIP_10002378 | | 0.92 | 2448 | 815 | other | |
| | DIP_10002379 | -1.85 | -1.67 | 3234 | 1077 | biosynthetic | |
| | DIP_10002380 | -0.73 | | 1056 | 351 | other | |
| | DIP_10002381 | | 0.40 | 1293 | 430 | other | |
| | DIP_10002382 | | -1.08 | 1101 | 366 | biosynthetic-additional | |
| | DIP_10002383 | | | 4584 | 1527 | other | |
| | DIP_10002384 | | | 741 | 246 | other | |
| | DIP_10002385 | | | 900 | 299 | other | |
| 4.2 | DIP_10007643 | | 1.28 | 912 | 303 | other | T1PKS |
| | DIP_10007644 | 0.95 | 1.36 | 1932 | 643 | other | |
| | DIP_10007645 | | -1.66 | 537 | 178 | other | |
| | DIP_10007646 | | | 834 | 277 | other | |
| | DIP_10007647 | 2.67 | 2.45 | 315 | 104 | other | |
| | DIP_10007648 | | | 7878 | 2625 | biosynthetic | |
| | DIP_10007649 | -2.95 | -2.77 | 915 | 304 | other | |
| | DIP_10007650 | -3.25 | -6.57 | 1302 | 433 | other | |
| | DIP_10007651 | -1.96 | -3.02 | 996 | 331 | other | |
| | DIP_10007652 | | | 291 | 96 | other | |
| | DIP_10007653 | | | 993 | 330 | other | |
| | DIP_10007654 | | | 474 | 157 | other | |
| 10.2 | DIP_10003450 | 3.48 | | 1257 | 418 | biosynthetic-additional | T1PKS |
| | DIP_10003451 | 7.30 | 5.54 | 1176 | 391 | other | |
| | DIP_10003452 | 2.58 | | 8346 | 2781 | biosynthetic | |
| | DIP_10003453 | | | 4347 | 1448 | other | |
| | DIP_10003454 | | | 195 | 64 | other | |
| | DIP_10003455 | | | 696 | 231 | other | |
| | DIP_10003456 | | 2.78 | 2034 | 677 | other | |
| | DIP_10003457 | 1.96 | 1.83 | 1353 | 450 | other | |
| 10.1 | DIP_10003179 | 5.46 | 3.30 | 1095 | 364 | biosynthetic | terpene |
| | DIP_10003180 | 1.42 | | 846 | 281 | other | |

a Location of secondary metabolite biosynthetic gene clusters in the *H. pulvinata* genome (GCA_006538405.1) predicted by AntiSMASH

b Log2 Fold-change values (log2FC ≥ 1, adjusted *P*-value < 0.05) for genes showing significant expression changes. Blank cells indicate genes with padj ≥ 0.05. All DEG datasets are available in Figshare (10.6084/m9.figshare.29491124)

c Characteristics of secondary metabolite biosynthetic gene clusters predicted by AntiSMASH.

Supplementary Table S5. Small secreted protein-coding genes (≤ 300 aa) in *Hansfordia pulvinata* .

| GeneID | *In vitro Cf/Hp* log2 Fold-change [a] | *In planta Cf/Hp* log2 Fold-change [a] | Number of amino acids | Description (NR) | Description (InterPro) | EffectorP | NetGPI | LOCALIZER (Chloroplast) | LOCALIZER (Mitochondria) | LOCALIZER (Nucleus) |
|---|---|---|---|---|---|---|---|---|---|---|
| DIP_10000066 | 4.11 | 6.33 | 72 | NA | NA | Apoplastic effector | Not GPI-Anchored | - | - | - |
| DIP_10000140 | 5.00 | 7.41 | 141 | NA | NA | Cytoplasmic effector | Not GPI-Anchored | - | - | - |
| DIP_10000197 | 10.86 | 13.64 | 129 | NA | NA | Cytoplasmic/apoplastic effector | Not GPI-Anchored | - | - | - |
| DIP_10000210 | 2.03 | 4.27 | 109 | NA | NA | Apoplastic/cytoplasmic effector | Not GPI-Anchored | - | - | - |
| DIP_10000258 | 6.87 | 11.17 | 180 | NA | NA | Apoplastic effector | Not GPI-Anchored | - | - | - |
| DIP_10000263 | 2.16 | 5.40 | 124 | hypothetical protein VHEMI04368 | NA | Cytoplasmic/apoplastic effector | Not GPI-Anchored | - | - | - |
| DIP_10000278 | 4.72 | 4.20 | 216 | hypothetical protein UCDDA912_g02783 | NA | Non-effector | Not GPI-Anchored | - | - | - |
| DIP_10000320 | 6.32 | 8.41 | 169 | NA | NA | Cytoplasmic effector | Not GPI-Anchored | - | - | - |
| DIP_10000414 | 1.74 | 4.02 | 231 | conserved hypothetical protein | IPR023346; Lysozyme-like domain | Apoplastic effector | Not GPI-Anchored | - | - | - |
| DIP_10000427 | 1.77 | 2.62 | 224 | NA | NA | Cytoplasmic effector | Not GPI-Anchored | - | - | Y (KKAITLTSPKGIRRLRP) |
| DIP_10000454 | 2.24 | 2.03 | 166 | putative immunoreactive protein | IPR008427; Extracellular membrane protein, CFEM domain IPR014005: Extracellular membrane protein | Non-effector | GPI-Anchored | - | - | - |
| DIP_10000474 | 6.14 | 8.97 | 148 | hypothetical protein FOPG_18856 | NA | Apoplastic effector | Not GPI-Anchored | Y (0.996 \| 12-44) | - | - |
| DIP_10000542 | 2.80 | 4.34 | 149 | NA | NA | Apoplastic effector | Not GPI-Anchored | - | - | - |
| DIP_10000703 | 4.80 | 6.25 | 206 | NA | NA | Non-effector | Not GPI-Anchored | - | - | - |
| DIP_10000714 | 6.62 | 7.45 | 247 | NA | NA | Cytoplasmic effector | Not GPI-Anchored | - | - | Y (FKKREDKNRRH) |
| DIP_10000750 | 3.07 | 4.01 | 261 | NA | NA | Cytoplasmic effector | Not GPI-Anchored | - | - | Y (RRTRRRAASKGKPEADTA |
| DIP_10000754 | 2.95 | 3.19 | 285 | putative glycoside hydrolase family 16 protein | IPR000757; Glycoside hydrolase, family 16 IPR013320; Concanavalin A-like lectin/glucanase | Non-effector | Not GPI-Anchored | - | - | - |
| DIP_10000796 | 4.42 | 6.48 | 293 | hypothetical protein UV8b_3126 | IPR008427; Extracellular membrane protein, CFEM domain | Non-effector | Not GPI-Anchored | - | - | Y (RHKKK,RRRHK) |
| DIP_10000797 | 6.06 | 9.81 | 125 | NA | NA | Cytoplasmic effector | Not GPI-Anchored | - | - | Y (SRRG) |
| DIP_10000868 | 3.51 | 3.41 | 204 | NA | NA | Cytoplasmic effector | Not GPI-Anchored | - | - | Y (QKRKGGGGRGGGSSGGK |
| DIP_10000996 | 2.75 | 4.70 | 148 | hypothetical protein PFICI_06810 | NA | Non-effector | Not GPI-Anchored | - | - | - |
| DIP_10001067 | 6.83 | 8.81 | 300 | NA | NA | Cytoplasmic effector | Not GPI-Anchored | - | - | - |
| DIP_10001157 | 1.66 | 4.65 | 262 | NA | NA | Cytoplasmic effector | Not GPI-Anchored | - | - | - |
| DIP_10001187 | 3.57 | 4.95 | 71 | NA | NA | Apoplastic effector | Not GPI-Anchored | - | - | - |
| DIP_10001305 | 1.51 | 4.56 | 184 | putative dnase1 protein | NA | Apoplastic effector | Not GPI-Anchored | - | - | - |
| DIP_10001364 | 1.54 | 2.48 | 139 | NA | NA | Non-effector | Not GPI-Anchored | - | - | Y (AGRRKKVGGGKAR) |
| DIP_10001482 | 2.79 | 4.44 | 217 | putative secreted protein | IPR011089; Domain of unknown function DUF1524 | Non-effector | Not GPI-Anchored | - | - | - |
| DIP_10001573 | 1.60 | 1.17 | 256 | NA | NA | Non-effector | Not GPI-Anchored | - | - | - |
| DIP_10001674 | 5.38 | 7.36 | 194 | NA | NA | Cytoplasmic effector | Not GPI-Anchored | - | - | - |
| DIP_10001679 | 4.57 | 6.03 | 113 | NA | NA | Cytoplasmic effector | Not GPI-Anchored | - | - | - |
| DIP_10001681 | 5.29 | 5.94 | 253 | Chloroperoxidase, partial | IPR000028; Chloroperoxidase | Non-effector | Not GPI-Anchored | - | - | Y (RPRR) |
| DIP_10001933 | 2.74 | 4.94 | 121 | hypothetical protein CSUB01_11738 | NA | Non-effector | GPI-Anchored | - | - | - |
| DIP_10002002 | 3.28 | 3.48 | 198 | NA | NA | Non-effector | Not GPI-Anchored | Y (0.954 \| 38-58) | - | - |
| DIP_10002023 | 6.27 | 7.15 | 299 | NA | NA | Cytoplasmic effector | Not GPI-Anchored | - | - | - |
| DIP_10002040 | 3.36 | 5.43 | 173 | NA | NA | Non-effector | Not GPI-Anchored | - | - | - |
| DIP_10002049 | 3.78 | 3.87 | 193 | hypothetical protein UCRPA7_2912 | IPR032382; Alternaria alternata allergen 1 | Apoplastic effector | Not GPI-Anchored | - | - | - |
| DIP_10002057 | 3.24 | 4.41 | 238 | hypothetical protein PFICI_06676 | NA | Non-effector | GPI-Anchored | - | - | - |
| DIP_10002060 | 3.70 | 4.85 | 256 | Putative Ribonuclease Trv | IPR001568; Ribonuclease T2-like IPR018188; Ribonuclease T2, active site | Non-effector | Not GPI-Anchored | - | - | - |
| DIP_10002224 | 4.88 | 5.94 | 174 | hypothetical protein PFICI_09070 | NA | Non-effector | Not GPI-Anchored | - | - | - |
| DIP_10002281 | 3.79 | 3.26 | 124 | NA | NA | Non-effector | Not GPI-Anchored | - | - | - |
| DIP_10002344 | 3.41 | 4.19 | 177 | NA | NA | Non-effector | Not GPI-Anchored | - | - | - |
| DIP_10002348 | 2.13 | 4.00 | 146 | NA | NA | Apoplastic effector | Not GPI-Anchored | - | - | - |
| DIP_10002354 | 6.43 | 6.32 | 99 | NA | NA | Cytoplasmic effector | Not GPI-Anchored | - | - | Y (KRKR) |
| DIP_10002417 | 2.76 | 5.01 | 274 | hypothetical protein PFICI_03087 | NA | Non-effector | Not GPI-Anchored | - | - | - |
| DIP_10002448 | 2.59 | 3.68 | 198 | NA | NA | Non-effector | Not GPI-Anchored | - | - | - |
| DIP_10002517 | 3.24 | 5.30 | 241 | NA | NA | Non-effector | Not GPI-Anchored | - | - | - |
| DIP_10002520 | 5.49 | 7.66 | 190 | hypothetical protein PV10_03707 | IPR001283; Cysteine-rich secretory protein, allergen V5/Tpx-1-related IPR014044; CAP domain | Apoplastic effector | Not GPI-Anchored | - | - | - |
| DIP_10002538 | 3.27 | 3.99 | 103 | NA | NA | Cytoplasmic effector | Not GPI-Anchored | - | - | - |
| DIP_10002539 | 2.32 | 4.10 | 69 | hypothetical protein CCM_03306 | NA | Non-effector | Not GPI-Anchored | - | - | - |
| DIP_10002542 | 1.53 | 1.37 | 300 | putative nuclease s1 protein | IPR003154; S1/P1 nuclease IPR008947; Phospholipase C/P1 nuclease domain | Non-effector | Not GPI-Anchored | - | - | - |
| DIP_10002620 | 2.72 | 6.95 | 130 | hypothetical protein PFICI_14406 | NA | Apoplastic/cytoplasmic effector | Not GPI-Anchored | - | - | - |
| DIP_10002660 | 4.45 | 8.02 | 58 | NA | NA | Cytoplasmic/apoplastic effector | Not GPI-Anchored | - | - | - |
| DIP_10002759 | 5.43 | 7.60 | 145 | putative epl1 protein | IPR009009; RlpA-like double-psi beta-barrel domain IPR010829; Cerato-platanin | Non-effector | Not GPI-Anchored | - | - | - |
| DIP_10002799 | 5.10 | 5.58 | 230 | NA | NA | Non-effector | Not GPI-Anchored | - | Y (0.966 \| 85-105) | Y (RRAVAPRQPHAARRHRG |
| DIP_10003023 | 4.06 | 8.06 | 100 | NA | NA | Non-effector | Not GPI-Anchored | - | - | - |
| DIP_10003057 | 2.54 | 3.13 | 123 | NA | NA | Apoplastic/cytoplasmic effector | Not GPI-Anchored | - | - | - |
| DIP_10003132 | 1.88 | 1.52 | 203 | putative transmembrane alpha-helix domain-containing protein | IPR002889; Carbohydrate-binding WSC | Non-effector | Not GPI-Anchored | - | - | Y (RRRR) |
| DIP_10003158 | 1.38 | 2.18 | 287 | NA | NA | Non-effector | Not GPI-Anchored | - | - | - |
| DIP_10003205 | 9.92 | 12.67 | 96 | hydrophobin 3 precursor | NA | Apoplastic effector | GPI-Anchored | - | - | - |
| DIP_10003291 | 1.50 | 3.01 | 262 | hypothetical protein FAVG1_06883 | NA | Non-effector | Not GPI-Anchored | - | - | - |
| DIP_10003304 | 6.87 | 8.90 | 186 | hypothetical protein LOTGIDRAFT_88143, partial | NA | Non-effector | Not GPI-Anchored | Y (0.963 \| 12-39) | Y (0.891 \| 38-59) | - |
| DIP_10003361 | 7.69 | 8.85 | 139 | NA | NA | Cytoplasmic effector | Not GPI-Anchored | - | - | - |
| DIP_10003488 | 7.10 | 9.51 | 194 | NA | NA | Apoplastic effector | Not GPI-Anchored | - | - | - |
| DIP_10003534 | 3.48 | 4.97 | 272 | NA | NA | Cytoplasmic effector | Not GPI-Anchored | - | Y (0.83 \| 43-74) | - |
| DIP_10003542 | 3.73 | 5.95 | 119 | NA | NA | Cytoplasmic effector | Not GPI-Anchored | - | - | - |
| DIP_10003548 | 3.46 | 4.01 | 103 | NA | NA | Non-effector | Not GPI-Anchored | - | - | - |
| DIP_10003614 | 6.53 | 8.72 | 296 | NA | NA | Cytoplasmic effector | Not GPI-Anchored | - | - | - |
| DIP_10003711 | 9.24 | 9.85 | 138 | NA | NA | Cytoplasmic effector | Not GPI-Anchored | - | - | - |
| DIP_10003855 | 3.10 | 2.84 | 280 | NA | NA | Non-effector | Not GPI-Anchored | - | - | - |
| DIP_10004049 | 5.91 | 7.67 | 165 | NA | NA | Cytoplasmic effector | Not GPI-Anchored | - | - | - |
| DIP_10004089 | 7.35 | 10.85 | 178 | hypothetical protein NEUTE1DRAFT_138203 | NA | Apoplastic effector | Not GPI-Anchored | - | - | - |
| DIP_10004128 | 2.25 | 4.52 | 160 | NA | NA | Cytoplasmic effector | Not GPI-Anchored | - | - | - |
| DIP_10004145 | 4.59 | 5.97 | 195 | NA | NA | Non-effector | GPI-Anchored | - | - | - |
| DIP_10004302 | 1.40 | 1.72 | 296 | hypothetical protein UCREL1_8406 | NA | Non-effector | GPI-Anchored | - | - | - |
| DIP_10004390 | 4.95 | 6.57 | 222 | NA | NA | Cytoplasmic effector | Not GPI-Anchored | - | - | - |
| DIP_10004543 | 3.42 | 1.58 | 167 | NA | NA | Cytoplasmic/apoplastic effector | Not GPI-Anchored | - | - | - |
| DIP_10004627 | 5.11 | 3.81 | 287 | putative secreted protein | IPR008701; Necrosis inducing protein | Non-effector | Not GPI-Anchored | - | - | - |
| DIP_10004826 | 1.71 | 2.95 | 270 | NA | NA | Non-effector | GPI-Anchored | - | - | - |
| DIP_10004892 | 3.55 | 5.57 | 127 | NA | NA | Cytoplasmic effector | Not GPI-Anchored | - | - | - |
| DIP_10004895 | 2.90 | 5.99 | 206 | NA | NA | Cytoplasmic effector | Not GPI-Anchored | - | - | - |
| DIP_10004922 | 5.14 | 6.88 | 92 | NA | NA | Apoplastic/cytoplasmic effector | Not GPI-Anchored | - | - | - |
| DIP_10004939 | 1.33 | 1.41 | 267 | putative signal sequence receptor alpha chain protein | IPR005595; Translocon-associated protein (TRAP), alpha subunit | Non-effector | Not GPI-Anchored | - | - | Y (PQAKRTK,AKRVKSGPNK |
| DIP_10004949 | 2.96 | 3.41 | 175 | NA | NA | Cytoplasmic effector | Not GPI-Anchored | - | - | Y (KRGSCDLELEKDDECKKR |
| DIP_10005021 | 1.40 | 3.40 | 292 | putative malate dehydrogenase protein | IPR021851; Protein of unknown function DUF3455 | Non-effector | Not GPI-Anchored | - | - | Y (KTEEG) |
| DIP_10005066 | 1.47 | 2.67 | 182 | NA | NA | Cytoplasmic effector | Not GPI-Anchored | - | - | - |
| DIP_10005070 | 3.64 | 5.00 | 297 | NA | NA | Non-effector | Not GPI-Anchored | - | - | Y (RRRR) |
| DIP_10005083 | 3.14 | 4.67 | 256 | Cutinase-like protein | IPR000675; Cutinase IPR011150; Cutinase, monofunctional IPR029058; Alpha/Beta hydrolase | Apoplastic effector | Not GPI-Anchored | - | - | - |
| DIP_10005086 | 4.96 | 6.52 | 125 | NA | NA | Cytoplasmic effector | Not GPI-Anchored | - | - | - |
| DIP_10005099 | 7.13 | 8.12 | 273 | NA | NA | Cytoplasmic effector | Not GPI-Anchored | - | - | Y (PKPS) |
| DIP_10005122 | 2.55 | 3.86 | 143 | hypothetical protein MGG_02347 | NA | Apoplastic effector | Not GPI-Anchored | - | - | - |
| DIP_10005184 | 2.61 | 3.85 | 259 | fungal chitosanase | IPR009939; Fungal chitosanase | Apoplastic effector | Not GPI-Anchored | - | - | - |
| DIP_10005269 | 5.10 | 4.70 | 262 | hypothetical protein MAJ_03385, partial | NA | Non-effector | GPI-Anchored | - | - | - |
| DIP_10005313 | 1.16 | 1.49 | 161 | NA | NA | Apoplastic effector | Not GPI-Anchored | - | - | - |
| DIP_10005315 | 3.47 | 4.09 | 192 | NA | NA | Cytoplasmic effector | Not GPI-Anchored | - | - | - |
| DIP_10005369 | 4.54 | 5.56 | 137 | NA | NA | Non-effector | Not GPI-Anchored | - | - | - |
| DIP_10005420 | 5.52 | 7.12 | 138 | putative rodlet peptide protein | IPR001338; Hydrophobin | Apoplastic effector | Not GPI-Anchored | - | - | - |
| DIP_10005485 | 4.54 | 5.87 | 185 | hypothetical protein PFICI_12551 | NA | Non-effector | Not GPI-Anchored | - | - | - |
| DIP_10005491 | 1.16 | 2.97 | 164 | NA | NA | Non-effector | Not GPI-Anchored | - | - | - |
| DIP_10005547 | 1.30 | 2.52 | 165 | NA | NA | Non-effector | Not GPI-Anchored | - | - | - |
| DIP_10005634 | 2.57 | 3.67 | 284 | NA | NA | Cytoplasmic effector | Not GPI-Anchored | - | - | Y (KKVDMPAKLSSYLRKEGK |
| DIP_10005738 | 2.08 | 3.62 | 210 | conserved hypothetical protein | NA | Cytoplasmic effector | Not GPI-Anchored | - | - | - |
| DIP_10005867 | 1.99 | 1.61 | 217 | NA | NA | Cytoplasmic effector | Not GPI-Anchored | - | - | - |
| DIP_10005970 | 4.12 | 4.69 | 260 | NA | NA | Cytoplasmic effector | Not GPI-Anchored | - | - | - |
| DIP_10005971 | 1.84 | 2.15 | 184 | NA | NA | Non-effector | Not GPI-Anchored | - | - | Y (RRRRRRR) |
| DIP_10005979 | 2.61 | 3.87 | 292 | putative superoxide dismutase protein | IPR001424; Superoxide dismutase, copper/zinc binding domain IPR024134; Superoxide dismutase | Non-effector | GPI-Anchored | - | - | - |
| DIP_10006058 | 3.17 | 5.09 | 247 | hypothetical protein SBOR_9046 | NA | Non-effector | GPI-Anchored | - | - | Y (PPPATPGRRR) |
| DIP_10006333 | 4.25 | 4.56 | 193 | hypothetical protein CGGC5_3556 | NA | Apoplastic effector | Not GPI-Anchored | - | - | - |
| DIP_10006382 | 1.75 | 1.89 | 288 | hypothetical protein NECHADRAFT_104641 | NA | Non-effector | Not GPI-Anchored | - | - | - |
| DIP_10006427 | 3.82 | 4.66 | 251 | NA | NA | Cytoplasmic effector | Not GPI-Anchored | - | - | Y (PRKK) |
| DIP_10006529 | 6.34 | 10.43 | 140 | putative killer kp4 protein | IPR011329; Killer toxin Kp4/SMK IPR015131; Killer toxin, Kp4 | Apoplastic effector | Not GPI-Anchored | - | - | - |
| DIP_10006530 | 8.23 | 12.58 | 129 | putative killer kp4 protein | IPR011329; Killer toxin Kp4/SMK IPR015131; Killer toxin, Kp4 | Apoplastic/cytoplasmic effector | Not GPI-Anchored | - | - | - |
| DIP_10006548 | 2.18 | 5.51 | 56 | NA | NA | Apoplastic/cytoplasmic effector | Not GPI-Anchored | - | - | - |
| DIP_10006639 | 5.85 | 7.26 | 93 | hypothetical protein SNOG_06459 | IPR010636; Cerato-ulmin hydrophobin family | Apoplastic/cytoplasmic effector | Not GPI-Anchored | - | - | - |
| DIP_10006674 | 3.92 | 6.97 | 245 | endoglucanase | IPR002594; Glycoside hydrolase, family 12 IPR013319; Glycoside hydrolase, family 11/12 | Apoplastic effector | Not GPI-Anchored | - | - | - |
| DIP_10006713 | 3.76 | 5.37 | 246 | hypothetical protein UCREL1_9109 | IPR010502; Carbohydrate-binding domain, family 9 | Non-effector | Not GPI-Anchored | - | - | - |
| DIP_10006869 | 2.61 | 3.88 | 254 | NA | NA | Non-effector | Not GPI-Anchored | - | Y (0.797 \| 60-90) | Y (RRRHAVLPADPALSRALR |
| DIP_10006877 | 5.87 | 8.39 | 210 | hypothetical protein UCREL1_9749 | NA | Cytoplasmic effector | Not GPI-Anchored | - | - | - |
| DIP_10007079 | 1.87 | 3.41 | 280 | hypothetical protein MYCTH_2070906 | IPR002818; DJ-1/PfpI IPR029062; Class I glutamine amidotransferase-like | Non-effector | Not GPI-Anchored | - | - | - |
| DIP_10007156 | 1.54 | 1.65 | 172 | NA | NA | Cytoplasmic effector | Not GPI-Anchored | - | - | - |
| DIP_10007328 | 2.37 | 1.74 | 249 | NA | NA | Cytoplasmic effector | Not GPI-Anchored | - | - | - |
| DIP_10007344 | 2.60 | 2.23 | 234 | NA | NA | Cytoplasmic effector | Not GPI-Anchored | - | - | - |
| DIP_10007425 | 6.26 | 7.41 | 214 | hypothetical protein PFICI_08584 | NA | Cytoplasmic effector | Not GPI-Anchored | - | - | - |
| DIP_10007545 | 1.69 | 1.83 | 169 | putative cdp-alcohol phosphatidyltransferase protein | NA | Apoplastic effector | GPI-Anchored | - | - | - |
| DIP_10007554 | 3.35 | 3.82 | 161 | NA | NA | Cytoplasmic effector | Not GPI-Anchored | - | - | - |
| DIP_10007560 | 8.05 | 10.69 | 163 | hypothetical protein Cob_13062 | NA | Cytoplasmic effector | Not GPI-Anchored | - | - | - |
| DIP_10007674 | 4.01 | 3.79 | 294 | NA | NA | Non-effector | Not GPI-Anchored | - | - | - |
| DIP_10007806 | 3.96 | 5.69 | 162 | putative cfem domain-containing protein | IPR008427; Extracellular membrane protein, CFEM domain IPR014005: Extracellular membrane protein | Non-effector | GPI-Anchored | - | - | - |
| DIP_10007817 | 4.77 | 6.30 | 223 | hypothetical protein SBOR_9046 | NA | Non-effector | GPI-Anchored | - | - | - |
| DIP_10007995 | 4.24 | 7.83 | 270 | hypothetical protein PFICI_09074 | IPR009939; Fungal chitosanase | Apoplastic effector | Not GPI-Anchored | - | - | Y (RRWKLEHCINVGSIKDYR |
| DIP_10008037 | 3.53 | 4.81 | 73 | NA | NA | Cytoplasmic effector | Not GPI-Anchored | - | - | - |
| DIP_10008095 | 2.81 | 5.31 | 142 | putative rodlet peptide protein | IPR001338; Hydrophobin | Apoplastic effector | Not GPI-Anchored | - | - | - |
| DIP_10008173 | 2.21 | 5.53 | 211 | NA | NA | Cytoplasmic effector | Not GPI-Anchored | - | - | - |
| DIP_10008179 | 3.84 | 6.30 | 209 | NA | NA | Cytoplasmic effector | Not GPI-Anchored | - | - | - |
| DIP_10008376 | 1.24 | 2.10 | 124 | PREDICTED: extensin-like | NA | Non-effector | Not GPI-Anchored | - | - | - |
| DIP_10008381 | 7.66 | 10.65 | 122 | NA | NA | Apoplastic effector | Not GPI-Anchored | - | - | - |
| DIP_10008424 | 3.08 | 3.52 | 229 | NA | NA | Cytoplasmic effector | Not GPI-Anchored | - | - | - |
| DIP_10008520 | 3.87 | 3.21 | 207 | NA | NA | Cytoplasmic effector | Not GPI-Anchored | - | - | Y (RKKK) |
| DIP_10008577 | 5.85 | 8.55 | 182 | NA | NA | Cytoplasmic effector | Not GPI-Anchored | - | - | - |
| DIP_10008584 | 5.98 | 8.25 | 279 | hypothetical protein M437DRAFT_64485 | NA | Non-effector | GPI-Anchored | - | - | - |
| DIP_10008613 | 5.09 | 4.56 | 110 | NA | NA | Cytoplasmic effector | Not GPI-Anchored | - | - | - |
| DIP_10008815 | 1.78 | 2.28 | 186 | NA | NA | Non-effector | Not GPI-Anchored | - | - | - |
| DIP_10008816 | 4.87 | 6.39 | 251 | NA | NA | Non-effector | Not GPI-Anchored | - | - | - |
| DIP_10008939 | 2.00 | 1.99 | 186 | NA | NA | Apoplastic effector | Not GPI-Anchored | - | - | - |
| DIP_10009037 | 1.18 | 1.20 | 156 | putative membrane protein | IPR001129; Membrane-associated, eicosanoid/glutathione metabolism (MAPEG) protein | Non-effector | Not GPI-Anchored | - | - | - |
| DIP_10009109 | 2.16 | 2.34 | 169 | hypothetical protein UCREL1_1801 | NA | Non-effector | Not GPI-Anchored | - | - | - |
| DIP_10009155 | 3.00 | 5.40 | 162 | hypothetical protein UCREL1_4769 | NA | Apoplastic effector | Not GPI-Anchored | - | - | - |
| DIP_10009263 | 4.34 | 8.05 | 113 | predicted protein | NA | Cytoplasmic effector | Not GPI-Anchored | - | - | - |
| DIP_10009424 | 5.91 | 7.35 | 259 | NA | NA | Cytoplasmic effector | GPI-Anchored | - | - | Y (RKKR,RKEWEKNDEVDKW |
| DIP_10009502 | 2.15 | 1.74 | 208 | hypothetical protein ACRE_032140 | NA | Non-effector | Not GPI-Anchored | - | - | - |
| DIP_10009900 | 3.96 | 3.69 | 228 | extracellular matrix protein | IPR018466; Ser-Thr-rich glycosyl-phosphatidyl-inositol-anchored membrane family | Non-effector | GPI-Anchored | - | - | - |
| DIP_10010025 | 1.53 | 1.95 | 123 | NA | NA | Non-effector | Not GPI-Anchored | - | - | - |
| DIP_10010270 | 3.92 | 5.62 | 54 | NA | NA | Non-effector | Not GPI-Anchored | - | - | - |
| DIP_10010283 | 3.55 | 2.50 | 270 | putative gnat family protein | IPR000182; GNAT domain IPR016181; Acyl-CoA N-acyltransferase | Non-effector | Not GPI-Anchored | - | Y (0.992 \| 49-70) | Y (RRGAKDLHMTIMIRQPKR) |
| DIP_10010303 | 3.13 | 3.59 | 215 | hypothetical protein NFIA_047850 | NA | Cytoplasmic/apoplastic effector | Not GPI-Anchored | - | - | Y (KKRDAPAEEPKKRDD,KK |
| DIP_10010378 | 1.46 | 1.75 | 235 | hypothetical protein UCREL1_2325 | NA | Non-effector | GPI-Anchored | - | - | - |
| DIP_10010390 | 1.87 | 1.95 | 283 | NA | NA | Non-effector | Not GPI-Anchored | - | - | - |
| DIP_10010397 | 3.42 | 4.79 | 282 | putative metalloprotease 1 protein | IPR008754; Peptidase M43, pregnancy-associated plasma-A IPR024079; Metallopeptidase, catalytic | Non-effector | Not GPI-Anchored | - | - | - |
| DIP_10010442 | 1.82 | 4.11 | 215 | hypothetical protein BAUCODRAFT_37640 | NA | Non-effector | Not GPI-Anchored | - | - | - |
| DIP_10010472 | 3.04 | 5.62 | 213 | NA | NA | Cytoplasmic effector | Not GPI-Anchored | - | - | - |
| DIP_10010500 | 1.17 | 1.67 | 253 | hypothetical protein UCREL1_4149 | NA | Apoplastic effector | Not GPI-Anchored | - | - | - |
| DIP_10010505 | 6.19 | 7.79 | 124 | NA | NA | Apoplastic effector | Not GPI-Anchored | - | - | - |
| DIP_10010536 | 3.56 | 5.84 | 202 | hypothetical protein UCDDS831_g03435 | IPR009009; RlpA-like double-psi beta-barrel domain | Apoplastic effector | Not GPI-Anchored | - | - | - |
| DIP_10010571 | 3.28 | 4.03 | 219 | putative gpi anchored serine-threonine rich protein | IPR018466; Ser-Thr-rich glycosyl-phosphatidyl-inositol-anchored membrane family | Non-effector | GPI-Anchored | - | - | - |
| DIP_10010667 | 3.46 | 9.05 | 158 | hypothetical protein PFICI_01649 | NA | Apoplastic effector | Not GPI-Anchored | - | - | - |
| DIP_10010700 | 7.90 | 12.95 | 100 | hypothetical protein AURDEDRAFT_164322 | NA | Non-effector | Not GPI-Anchored | - | - | - |

a Log2 Fold-change values (log2FC ≥ 1, adjusted *P* -value < 0.05) for genes showing significant expression changes. Blank cells indicate genes with padj ≥ 0.05. All DEG datasets are available in Figshare (10.6084/m9.figshare.29491124)

Supplementary Table S6. Key genes encoding secondary metabolism enzymes in *Cladosporium fulvum* .

| **GeneID** | **Description** | **SM key enzyme type** | ***In planta Cf*** **log2 Fold-change** [a] | ***In vitro Cf/Hp*** **log2 Fold-change** [a] | ***In planta Cf/Hp*** **log2 Fold-change** [a] |
|---|---|---|---|---|---|
| CLAFUR5_00278 | Squalestatin hexaketide synthase | T1PKS | 0.69 | | 1.25 |
| CLAFUR5_00877 | Geranylgeranyl pyrophosphate synthase 1 | Terpene | -0.73 | | |
| CLAFUR5_02085 | putative squalene synthase | Terpene | | | |
| CLAFUR5_02708 | Non-canonical non-ribosomal peptide synthetase FUB8 | NRPS-like | | | |
| CLAFUR5_03847 | Apicidin F synthase (*PKS7* ) | NRPS | -1.85 | 0.00 | -2.66 |
| CLAFUR5_04416 | Nonribosomal peptide synthetase fmqA (*NPS5* ) | NRPS | -0.45 | 0.00 | -1.06 |
| CLAFUR5_04428 | Oxalate--CoA ligase | NRPS-like | -1.07 | 0.00 | |
| CLAFUR5_04894 | Peramine synthetase ppzA (*NPS6* ) | NRPS-like | -1.04 | | -0.92 |
| CLAFUR5_06161 | Reducing polyketide synthase PKS2 (*PKS5* ) | T1PKS | -2.08 | -2.31 | -2.86 |
| CLAFUR5_06232 | Hybrid PKS-NRPS synthetase lepA (*HPS1* ) | PKS-NRPS | 0.55 | | 0.26 |
| CLAFUR5_07436 | Nonribosomal peptide synthetase dtxS1 (*NPS4* ) | NRPS | | | -0.52 |
| CLAFUR5_07484 | Non-canonical non-ribosomal peptide synthetase FUB8 | NRPS-like | -0.74 | 0.00 | -1.63 |
| CLAFUR5_08791 | Nonribosomal peptide synthase atnA (*NPS3* ) | NRPS | -1.36 | | -1.94 |
| CLAFUR5_09052 | Putative acyl-CoA synthetase | NRPS | -0.76 | 0.00 | -1.36 |
| CLAFUR5_09474 | Non-canonical non-ribosomal peptide synthetase FUB8 | NRPS-like | | 2.54 | |
| CLAFUR5_09844 | L-2-aminoadipate reductase large subunit | NRPS-like | -0.79 | 0.00 | |
| CLAFUR5_10460 | Terpene cyclase ATR13 | Terpene | | | |
| CLAFUR5_10739 | Nonribosomal peptide synthase (*NPS2* ) | NRPS | -0.56 | 0.00 | -1.69 |
| CLAFUR5_10765 | 6-hydroxymellein synthase (PKS3) | T1PKS | 2.93 | | 4.12 |
| CLAFUR5_10777 | Nonribisomal peptide synthetase (*NPS10* ) | NRPS | 3.15 | 0.00 | |
| CLAFUR5_10780 | Highly reducing polyketide synthase (*PKS8* ) | T1PKS | 1.38 | 2.50 | |
| CLAFUR5_11006 | Non-canonical non-ribosomal peptide synthetase FUB8 | NRPS | -0.64 | | -0.49 |
| CLAFUR5_11318 | Non-canonical non-ribosomal peptide synthetase FUB8 | NRPS | | | |
| CLAFUR5_11407 | Non-reducing polyketide synthase CTB1 (*PKS7* ) | T1PKS | -0.62 | 0.00 | -0.98 |
| CLAFUR5_12064 | Nonribosomal peptide synthetase 8 | NRPS | | | |
| CLAFUR5_12662 | Acyl-CoA ligase sidI | NRPS | -0.96 | 1.21 | 0.95 |
| CLAFUR5_12787 | Adenylate-forming reductase Nps10 | NRPS-like | -3.74 | | -2.83 |
| CLAFUR5_12824 | Non-reducing polyketide synthase PKS1 (*PKS1* ) | T1PKS | | | -1.14 |
| CLAFUR5_12895 | Nonribosomal peptide synthetase TES (*NPS8* ) | NRPS | 0.36 | | 0.66 |
| CLAFUR5_12905 | Atrochrysone carboxylic acid synthase (*PKS6* ) | T1PKS | -2.71 | -1.72 | -3.81 |
| CLAFUR5_12913 | Nonribosomal peptide synthetase sidC (*NPS9* ) | NRPS | -2.23 | | -1.82 |
| CLAFUR5_13164 | Aspulvinone E synthetase melA | NRPS-like | -4.06 | | -4.37 |
| CLAFUR5_13273 | Prosolanapyrone synthase (*PKS2* ) | T1PKS | 0.61 | 0.00 | 0.49 |
| CLAFUR5_13278 | Nonribosomal peptide synthetase (*NPS1* ) | NRPS | | | -0.55 |
| CLAFUR5_13418 | Nonribosomal peptide synthetase dtxS1 | NRPS | -1.94 | | -1.87 |
| CLAFUR5_13788 | Adenylate-forming reductase Nps10 | NRPS-like | | | 4.89 |
| CLAFUR5_13909 | Non-reducing polyketide synthase PKS19 (*PKS9* ) | T1PKS | -1.21 | -0.66 | -1.81 |
| CLAFUR5_13961 | Non-reducing polyketide synthase pkbA | T1PKS | -1.24 | | -1.52 |
| CLAFUR5_13962 | Adenylate-forming reductase cicB | NRPS-like | -1.48 | 0.00 | -2.91 |
| CLAFUR5_14164 | Norsolorinic acid synthase (*PKSA* ) | T1PKS | -1.41 | | -1.67 |
| CLAFUR5_14251 | Bifunctional lycopene cyclase/phytoene synthase | Terpene | -0.47 | 0.00 | -0.31 |
| CLAFUR5_14350 | Linear gramicidin synthase subunit D | NRPS-like | -1.20 | | -2.19 |

a Log2 Fold-change values (log2FC ≥ 1, adjusted *P* -value < 0.05) for genes showing significant expression changes. Blank cells indicate genes with padj ≥ 0.05. All DEG datasets are available in Figshare (10.6084/m9.figshare.29491124)

Supplementary Table S7. Effector genes identified in *Cladosporium fulvum* .

| GeneID | Description [a] | Residue Count | No. cysteine residues | *In planta Cf* log2 Fold-change [b] | *In vitro Cf/Hp* log2 Fold-change [b] | *In planta Cf/Hp* log2 Fold-change [b] | EffectorP | SignalP | AMAPEC: pLDDT | AMPEC: Probability of antimicrobial activity | AMAPEC prediction | Classfication in Fig. S6 |
|---|---|---|---|---|---|---|---|---|---|---|---|---|
| CLAFUR4_00498 | *Avr5* | 103 | 10 | - | - | - | Cytoplasmic/apoplastic effector | SP | 48.960 | 0.969 | Antimicrobial | - |
| CLAFUR5_00004 | *Ecp2* | 165 | 4 | 7.52 | 5.70 | 7.51 | Apoplastic/cytoplasmic effector | SP | 85.417 | 0.955 | Antimicrobial | b |
| CLAFUR5_00034 | *Ecp4* | 119 | 6 | 5.41 | 5.86 | 5.94 | Apoplastic/cytoplasmic effector | SP | 66.331 | 0.164 | Non-antimicrobial | d |
| CLAFUR5_00065 | *Ecp23* | 91 | 4 | 5.42 | 3.84 | 6.09 | Cytoplasmic effector | SP | 72.218 | 0.790 | Antimicrobial | f |
| CLAFUR5_00090 | *Ecp58-1* | 196 | 4 | 1.17 | 3.95 | 1.26 | Non-effector | SP | 86.749 | 0.566 | Antimicrobial | a |
| CLAFUR5_01461 | *Ecp52* | 170 | 4 | 2.43 | 2.79 | 3.05 | Cytoplasmic/apoplastic effector | SP | 77.793 | 0.876 | Antimicrobial | a |
| CLAFUR5_01523 | *Ecp43-1* | 94 | 10 | 7.71 | 2.27 | 7.40 | Apoplastic/cytoplasmic effector | SP | 40.029 | 0.017 | Non-antimicrobial | d |
| CLAFUR5_01609 | *Ecp28-1* | 101 | 6 | 3.01 | | 4.06 | Cytoplasmic/apoplastic effector | SP | 92.401 | 0.957 | Antimicrobial | d |
| CLAFUR5_01926 | *CE39* | 65 | 8 | 6.17 | 1.68 | 5.83 | Apoplastic/cytoplasmic effector | SP | 48.819 | 0.016 | Non-antimicrobial | d |
| CLAFUR5_01962 | *CE21* | 49 | 8 | 2.43 | | 1.47 | Cytoplasmic/apoplastic effector | SP | 48.563 | 0.001 | Non-antimicrobial | a |
| CLAFUR5_02466 | *CfPhiA-2* | 218 | 6 | 1.21 | 2.84 | 1.78 | Apoplastic/cytoplasmic effector | SP | 83.987 | 0.080 | Non-antimicrobial | f |
| CLAFUR5_02840 | *Ecp20-2* | 149 | 4 | 5.10 | 2.47 | 5.48 | Cytoplasmic/apoplastic effector | SP | 90.632 | 0.976 | Antimicrobial | a |
| CLAFUR5_02977 | *Ecp28-3* | 105 | 8 | 1.11 | 0.00 | 2.29 | Cytoplasmic/apoplastic effector | SP | 93.993 | 0.822 | Antimicrobial | a |
| CLAFUR5_03132 | *Ecp57-1* | 285 | 5 | 1.17 | | 1.61 | Apoplastic effector | SP | 74.721 | 0.988 | Antimicrobial | d |
| CLAFUR5_03423 | *Ecp5* | 115 | 6 | 1.92 | | 1.96 | Apoplastic/cytoplasmic effector | SP | 83.752 | 0.458 | Non-antimicrobial | a |
| CLAFUR5_03793 | *CE75* | 239 | 7 | 2.90 | | 2.61 | Apoplastic effector | SP | 84.782 | 0.488 | Non-antimicrobial | b |
| CLAFUR5_03986 | *Ecp10-3* | 77 | 8 | | | | Cytoplasmic/apoplastic effector | SP | 50.114 | 0.106 | Non-antimicrobial | d |
| CLAFUR5_04117 | *Ecp20-3* | 153 | 4 | 1.93 | 3.69 | 4.75 | Apoplastic/cytoplasmic effector | SP | 89.256 | 0.579 | Antimicrobial | b |
| CLAFUR5_04375 | *CE38* | 162 | 7 | 1.50 | | 1.19 | Cytoplasmic/apoplastic effector | SP | 59.413 | 0.959 | Antimicrobial | a |
| CLAFUR5_04595 | *Ecp33* | 210 | 8 | 2.19 | 2.51 | 3.63 | Cytoplasmic effector | SP | 50.193 | 0.000 | Non-antimicrobial | a |
| CLAFUR5_05064 | *Ecp59-1* | 128 | 2 | 2.08 | 6.60 | 3.29 | Apoplastic effector | SP | 92.528 | 0.399 | Non-antimicrobial | f |
| CLAFUR5_05345 | *Ecp49-1* | 104 | 7 | 3.25 | | 4.55 | Apoplastic/cytoplasmic effector | SP | 50.229 | 0.144 | Non-antimicrobial | b |
| CLAFUR5_05411 | *Ecp14-2* | 165 | 14 | | | | Apoplastic/cytoplasmic effector | SP | 61.839 | 0.397 | Non-antimicrobial | a |
| CLAFUR5_05420 | *Ecp32-2* | 195 | 2 | 0.98 | 0.00 | 1.94 | Apoplastic effector | SP | 92.475 | 0.724 | Antimicrobial | a |
| CLAFUR5_05515 | *Ecp2-3* | 372 | 5 | 5.61 | 3.54 | 5.14 | Apoplastic effector | OTHER | 49.879 | 0.946 | Antimicrobial | b |
| CLAFUR5_05549 | *Ecp53-1* | 176 | 2 | -1.89 | 0.00 | 3.64 | Apoplastic effector | SP | 96.646 | 0.376 | Non-antimicrobial | a |
| CLAFUR5_05638 | *Ecp9-9* | 90 | 6 | 3.30 | 3.42 | 4.57 | Cytoplasmic/apoplastic effector | SP | 93.905 | 0.803 | Antimicrobial | a |
| CLAFUR5_06015 | *Ecp14-1* | 206 | 12 | 4.85 | | 4.80 | Apoplastic effector | SP | 49.061 | 0.966 | Antimicrobial | c |
| CLAFUR5_06022 | *Ecp2-2* | 93 | 2 | | 1.65 | 2.07 | Apoplastic effector | SP | 55.814 | 0.007 | Non-antimicrobial | d |
| CLAFUR5_06030 | *Ecp9-8* | 104 | 6 | 1.12 | 2.42 | 2.16 | Cytoplasmic/apoplastic effector | SP | 79.575 | 0.961 | Antimicrobial | a |
| CLAFUR5_06582 | *Ecp11-1-1* | 165 | 10 | 2.13 | 2.11 | 3.60 | Cytoplasmic/apoplastic effector | SP | 97.154 | 0.840 | Antimicrobial | a |
| CLAFUR5_06583 | *Ecp11-1-2* | 165 | 10 | 2.38 | 2.10 | 3.87 | Cytoplasmic/apoplastic effector | SP | 97.154 | 0.829 | Antimicrobial | f |
| CLAFUR5_06629 | *Ecp47* | 148 | 10 | 2.68 | | 0.43 | Cytoplasmic/apoplastic effector | SP | 94.404 | 0.999 | Antimicrobial | a |
| CLAFUR5_06786 | *Ecp9-2* | 77 | 6 | 5.04 | 2.08 | 5.93 | Cytoplasmic/apoplastic effector | SP | 93.608 | 0.882 | Antimicrobial | b |
| CLAFUR5_06841 | *Ecp9-1* | 88 | 6 | 5.34 | | 5.21 | Cytoplasmic/apoplastic effector | SP | 87.897 | 0.896 | Antimicrobial | b |
| CLAFUR5_07035 | *Ecp50-1* | 182 | 0 | 1.88 | | 1.86 | Cytoplasmic effector | SP | 56.986 | 0.073 | Non-antimicrobial | c |
| CLAFUR5_07043 | *Ecp15* | 131 | 8 | 2.01 | | 1.42 | Cytoplasmic/apoplastic effector | SP | 68.413 | 0.848 | Antimicrobial | d |
| CLAFUR5_07332 | *CE78* | 332 | 3 | 6.54 | 2.01 | 6.25 | Non-effector | SP | 92.859 | 0.517 | Antimicrobial | d |
| CLAFUR5_07496 | *Ecp30* | 141 | 6 | 2.29 | 5.32 | 3.40 | Apoplastic/cytoplasmic effector | SP | 87.961 | 0.829 | Antimicrobial | a |
| CLAFUR5_07513 | *Ecp9-6* | 93 | 6 | | | | Cytoplasmic/apoplastic effector | SP | 83.832 | 0.942 | Antimicrobial | c |
| CLAFUR5_07683 | *Ecp39* | 63 | 8 | | | 1.58 | Apoplastic/cytoplasmic effector | SP | 71.155 | 0.480 | Non-antimicrobial | e |
| CLAFUR5_09439 | *Ecp19-2* | 130 | 8 | 2.53 | 1.08 | 1.66 | Cytoplasmic/apoplastic effector | SP | 84.375 | 0.450 | Non-antimicrobial | f |
| CLAFUR5_09751 | *Ecp10-2* | 67 | 6 | 4.07 | 1.36 | 4.39 | Apoplastic/cytoplasmic effector | SP | 46.770 | 0.196 | Non-antimicrobial | c |
| CLAFUR5_09818 | *Avr9* | 63 | 6 | 6.04 | 1.26 | 3.19 | Apoplastic effector | SP | 71.729 | 0.888 | Antimicrobial | d |
| CLAFUR5_09955 | *Ecp54-1* | 160 | 11 | 1.55 | 0.00 | 1.47 | Apoplastic/cytoplasmic effector | SP | 87.823 | 0.982 | Antimicrobial | d |
| CLAFUR5_10085 | *Ecp6* | 222 | 8 | 4.06 | 1.70 | 3.97 | Apoplastic/cytoplasmic effector | SP | 90.765 | 0.898 | Antimicrobial | a |
| CLAFUR5_10106 | *hcf-4* | 210 | 9 | 8.93 | 1.67 | 7.20 | Cytoplasmic effector | SP | 44.027 | 0.207 | Non-antimicrobial | c |
| CLAFUR5_10112 | *Avr4* | 135 | 8 | 5.57 | 1.46 | 5.55 | Cytoplasmic effector | SP | 79.841 | 0.220 | Non-antimicrobial | a |
| CLAFUR5_10278 | *Avr4E* | 121 | 6 | 5.94 | 0.00 | 7.20 | Apoplastic/cytoplasmic effector | SP | 39.594 | 0.511 | Antimicrobial | c |
| CLAFUR5_10379 | *Ecp44* | 146 | 4 | 4.89 | | 4.85 | Non-effector | SP | 92.070 | 0.382 | Non-antimicrobial | a |
| CLAFUR5_10563 | *CE32* | 141 | 6 | -1.91 | 1.42 | -1.72 | Cytoplasmic/apoplastic effector | SP | 59.742 | 0.036 | Non-antimicrobial | a |
| CLAFUR5_10639 | *Ecp28-2* | 105 | 8 | 2.99 | 1.71 | 3.42 | Cytoplasmic effector | SP | 93.071 | 0.870 | Antimicrobial | e |
| CLAFUR5_10648 | *CE23* | 55 | 8 | 5.83 | 2.36 | 5.76 | Apoplastic/cytoplasmic effector | SP | 52.720 | 0.808 | Antimicrobial | a |
| CLAFUR5_10680 | *Ecp19-1* | 162 | 8 | 5.21 | 6.44 | 5.06 | Cytoplasmic/apoplastic effector | SP | 62.501 | 0.731 | Antimicrobial | a |
| CLAFUR5_11007 | *hcf-5* | 74 | 8 | -5.47 | -4.12 | -5.53 | Apoplastic/cytoplasmic effector | SP | 73.043 | 0.101 | Non-antimicrobial | a |
| CLAFUR5_11054 | *hcf-3* | 85 | 8 | 5.16 | | 3.30 | Apoplastic/cytoplasmic effector | SP | 55.969 | 0.473 | Non-antimicrobial | b |
| CLAFUR5_11110 | *CfPhiA-1* | 195 | 4 | 5.44 | | 6.75 | Apoplastic effector | SP | 91.172 | 0.991 | Antimicrobial | e |
| CLAFUR5_11337 | *Ecp21-1* | 184 | 8 | 6.83 | 4.07 | 6.79 | Apoplastic/cytoplasmic effector | SP | 79.777 | 0.921 | Antimicrobial | a |
| CLAFUR5_11716 | *CE43* | 179 | 6 | 8.38 | | 10.62 | Cytoplasmic effector | SP | 40.421 | 0.032 | Non-antimicrobial | a |
| CLAFUR5_11887 | *Ecp45* | 146 | 4 | 1.14 | 5.30 | 4.61 | Apoplastic effector | SP | 95.161 | 0.965 | Antimicrobial | f |
| CLAFUR5_12097 | *Ecp60-1* | 249 | 6 | 1.16 | 1.17 | 1.65 | Apoplastic effector | SP | 87.733 | 0.999 | Antimicrobial | a |
| CLAFUR5_12163 | *Ecp46* | 164 | 2 | 5.91 | 2.83 | 5.80 | Cytoplasmic effector | SP | 54.715 | 0.006 | Non-antimicrobial | c |
| CLAFUR5_12291 | *CE28* | 161 | 10 | 1.39 | | 0.56 | Apoplastic effector | SP | 65.115 | 0.009 | Non-antimicrobial | b |
| CLAFUR5_12426 | *Ecp7* | 116 | 6 | 7.40 | 4.36 | 7.77 | Cytoplasmic/apoplastic effector | SP | 73.902 | 0.747 | Antimicrobial | d |
| CLAFUR5_12448 | *Avr2* | 78 | 8 | 1.43 | 1.69 | 1.94 | Cytoplasmic/apoplastic effector | SP | 65.725 | 0.927 | Antimicrobial | a |
| CLAFUR5_12564 | *Ecp51* | 238 | 8 | 1.35 | 0.00 | | Apoplastic effector | SP | 67.327 | 0.968 | Antimicrobial | e |
| CLAFUR5_13363 | *CE10* | 95 | 8 | 5.94 | 3.10 | 5.46 | Cytoplasmic/apoplastic effector | SP | 55.009 | 0.873 | Antimicrobial | c |
| CLAFUR5_13431 | *Ecp1* | 96 | 8 | 8.12 | 2.34 | 8.49 | Apoplastic/cytoplasmic effector | SP | 72.009 | 0.957 | Antimicrobial | a |
| CLAFUR5_13715 | *hcf-2* | 96 | 8 | 4.08 | | 2.72 | Apoplastic/cytoplasmic effector | SP | 56.462 | 0.715 | Antimicrobial | b |
| CLAFUR5_14044 | *hcf-6* | 168 | 8 | -1.37 | 5.52 | 2.53 | Cytoplasmic/apoplastic effector | SP | 55.520 | 0.999 | Antimicrobial | a |
| CLAFUR5_14656 | *CE46* | 40 | 7 | | | | Cytoplasmic/apoplastic effector | SP | 74.180 | 0.849 | Antimicrobial | a |
| CLAFUR5_14657 | *Ecp10-1* | 70 | 6 | 5.80 | 3.02 | 6.38 | Cytoplasmic/apoplastic effector | SP | 86.948 | 0.929 | Antimicrobial | a |
| CLAFUR5_14658 | *Ecp9-3* | 102 | 6 | | | | Cytoplasmic/apoplastic effector | SP | 83.885 | 0.112 | Non-antimicrobial | b |
| CLAFUR5_14659 | *Ecp42* | 128 | 14 | 4.20 | | 3.93 | Apoplastic/cytoplasmic effector | SP | 36.768 | 0.007 | Non-antimicrobial | f |
| CLAFUR5_14660 | *Ecp20-1* | 166 | 4 | 7.13 | 5.29 | 6.65 | Apoplastic effector | SP | 83.417 | 0.482 | Non-antimicrobial | c |
| CLAFUR5_14661 | *Ecp37* | 79 | 6 | 2.97 | 2.01 | 1.21 | Cytoplasmic/apoplastic effector | SP | 46.795 | 0.372 | Non-antimicrobial | a |
| CLAFUR5_14662 | *Ecp12* | 133 | 8 | 6.67 | 3.27 | 6.33 | Apoplastic/cytoplasmic effector | SP | 37.394 | 0.786 | Antimicrobial | c |
| CLAFUR5_14663 | *Ecp41* | 133 | 9 | 1.34 | | 1.57 | Apoplastic/cytoplasmic effector | SP | 33.234 | 0.055 | Non-antimicrobial | b |
| CLAFUR5_14664 | *Ecp16* | 101 | 8 | 1.17 | | 1.13 | Cytoplasmic/apoplastic effector | SP | 40.414 | 0.034 | Non-antimicrobial | b |
| CLAFUR5_14665 | *Ecp8* | 105 | 8 | 4.66 | 2.53 | 5.28 | Apoplastic/cytoplasmic effector | SP | 37.861 | 0.066 | Non-antimicrobial | e |
| CLAFUR5_14666 | *Ecp9-7* | 86 | 6 | | | | Cytoplasmic/apoplastic effector | SP | 91.198 | 0.972 | Antimicrobial | e |
| CLAFUR5_14667 | *Ecp36-1* | 73 | 10 | 2.71 | 2.71 | 2.72 | Cytoplasmic/apoplastic effector | SP | 79.325 | 0.995 | Antimicrobial | f |
| CLAFUR5_14668 | *Ecp13* | 73 | 10 | 2.13 | 0.00 | | Cytoplasmic/apoplastic effector | SP | 73.214 | 0.739 | Antimicrobial | a |
| CLAFUR5_14669 | *CE29* | 43 | 9 | 6.85 | 0.00 | 7.08 | Apoplastic/cytoplasmic effector | SP | 60.936 | 0.405 | Non-antimicrobial | d |
| CLAFUR5_14670 | *CE52* | 51 | 7 | 0.56 | 0.00 | 1.04 | Apoplastic/cytoplasmic effector | SP | 64.025 | 0.342 | Non-antimicrobial | a |
| CLAFUR5_14671 | *Ecp29* | 266 | 14 | 3.79 | 3.30 | 4.84 | Apoplastic/cytoplasmic effector | SP | 44.370 | 0.974 | Antimicrobial | c |
| CLAFUR5_14672 | *Ecp18* | 81 | 10 | 4.33 | 1.21 | 4.22 | Apoplastic/cytoplasmic effector | SP | 52.960 | 0.099 | Non-antimicrobial | e |
| CLAFUR5_14673 | *Ecp26* | 65 | 6 | 3.17 | | 4.98 | Cytoplasmic/apoplastic effector | SP | 65.607 | 0.965 | Antimicrobial | a |
| CLAFUR5_14674 | *Ecp55* | 239 | 20 | 2.91 | | 2.62 | Apoplastic/cytoplasmic effector | SP | 64.641 | 0.781 | Antimicrobial | a |
| CLAFUR5_14675 | *CE2* | 42 | 7 | 7.23 | 3.34 | 6.20 | Cytoplasmic/apoplastic effector | SP | 67.943 | 0.016 | Non-antimicrobial | e |
| CLAFUR5_14676 | *Ecp48* | 78 | 8 | | 4.16 | 1.17 | Apoplastic/cytoplasmic effector | SP | 60.515 | 0.210 | Non-antimicrobial | a |
| CLAFUR5_14677 | *Ecp56* | 60 | 6 | 6.95 | | 3.96 | Apoplastic/cytoplasmic effector | SP | 52.050 | 0.015 | Non-antimicrobial | f |
| CLAFUR5_14678 | *Avr9B* | 152 | 8 | | -2.09 | -2.98 | Non-effector | SP | 42.750 | 0.000 | Non-antimicrobial | a |
| CLAFUR5_14679 | *Ecp31* | 93 | 10 | 4.57 | 1.93 | 4.17 | Cytoplasmic/apoplastic effector | SP | 44.871 | 0.578 | Antimicrobial | a |
| CLAFUR5_14680 | *CE17* | 73 | 9 | 7.88 | | 6.89 | Cytoplasmic/apoplastic effector | SP | 40.672 | 0.054 | Non-antimicrobial | e |
| CLAFUR5_14681 | *Ecp25* | 79 | 8 | 5.13 | 4.82 | 5.74 | Cytoplasmic/apoplastic effector | SP | 47.056 | 0.102 | Non-antimicrobial | a |
| CLAFUR5_14682 | *Ecp34* | 92 | 8 | 1.44 | | 1.32 | Cytoplasmic/apoplastic effector | SP | 52.665 | 0.179 | Non-antimicrobial | c |
| CLAFUR5_14683 | *Ecp40* | 92 | 8 | | | | Cytoplasmic/apoplastic effector | SP | 38.990 | 0.006 | Non-antimicrobial | a |
| CLAFUR5_14684 | *Ecp22* | 161 | 8 | 4.80 | 1.49 | 5.31 | Cytoplasmic effector | SP | 56.228 | 0.881 | Antimicrobial | a |
| CLAFUR5_14685 | *Ecp38* | 84 | 10 | 3.46 | | 2.94 | Apoplastic effector | SP | 60.248 | 0.117 | Non-antimicrobial | d |
| CLAFUR5_14686 | *Ecp27* | 67 | 4 | 5.92 | 2.08 | 5.71 | Cytoplasmic/apoplastic effector | SP | 65.402 | 0.863 | Antimicrobial | b |
| CLAFUR5_14687 | *Ecp24-2* | 100 | 4 | 13.04 | | 13.00 | Apoplastic/cytoplasmic effector | SP | 54.807 | 0.598 | Antimicrobial | c |
| CLAFUR5_14688 | *Ecp32-1* | 197 | 4 | 2.86 | | 2.69 | Cytoplasmic effector | SP | 92.528 | 0.641 | Antimicrobial | e |
| CLAFUR5_14689 | *Ecp24-1* | 92 | 4 | 3.48 | 1.63 | 3.35 | Apoplastic effector | SP | 56.443 | 0.363 | Non-antimicrobial | e |
| CLAFUR5_14690 | *Ecp35* | 70 | 8 | 6.47 | 2.50 | 6.56 | Apoplastic/cytoplasmic effector | SP | 41.442 | 0.111 | Non-antimicrobial | a |
| CLAFUR5_20078 | *Ecp17* | 62 | 6 | 4.52 | 0.91 | 4.02 | Cytoplasmic/apoplastic effector | SP | 56.534 | 0.199 | Non-antimicrobial | a |

a For *ECP11-1* , the suffixes "-1" and "-2" were added to distinguish duplicated paralogous copies in the genome. As the race5 genome was used, the expression level of Avr5 was not calculated.

b Log2 Fold-change values (log2FC ≥ 1, adjusted *P* -value < 0.05) for genes showing significant expression changes. Blank cells indicate genes with padj ≥ 0.05. All DEG datasets are available in Figshare (10.6084/m9.figshare.29491124)

Supplementary Table S8. Structural homologs of Ecp2 protein identified from PDB25 using the Dali server.

| No. | PDB ID | Chain | Description | %ID | %Similarity | *Z*-score | RMSD | Lali | Nres |
|---|---|---|---|---|---|---|---|---|---|
| 1 | 8acx | 8acx-B | HCE2 domain-containing protein (ZtKP4) | 25.6 | 36.9 | 13.7 | 2 | 103 | 113 |
| 2 | 1kpt | 1kpt-A | KP4 toxin (UmVKP4) | 4.6 | 5.1 | 6.6 | 2.5 | 85 | 105 |
| 3 | 5v6i | 5v6i-A | TMV resistance protein Y3 | 7.3 | 12.1 | 6.4 | 2.5 | 85 | 112 |
| 4 | 4nds | 4nds-A | Alpha-galactosyl-binding lectin | 12.0 | 20.4 | 5.3 | 3.2 | 81 | 94 |
| 5 | 7jsr | 7jsr-B | NAD-specific glutamate dehydrogenase | 1.7 | 2.9 | 3.3 | 4.7 | 80 | 1564 |
| 6 | 3a2e | 3a2e-A | Ginkbilobin-2 | 12.1 | 24.8 | 3.3 | 3.7 | 71 | 108 |

RMSD, root mean square deviation; %ID, % Identity; Lali, length of aligned residues; Nres, number of residues in query

Supplementary Table S9. Species with homologs of Ecp2 and KP4 proteins.

| Group | Organism | Abbreviation [a] | Database [b] | BLASTp HIT | Taxon |
|---|---|---|---|---|---|
| Ecp2 | *Pseudocercospora musae* CBS 116634 | Psemus1 | Psemus1_ExternalModels_proteins_2021-11-19.fasta | 3 | *Dothideomycetes* |
| | *Mycosphaerella eumusae* CBS 114824 | Myceu1 | Myceu1_ExternalModels_proteins_2021-11-19.fasta | 3 | *Dothideomycetes* |
| | *Coniochaeta ligniaria* CBS 111746 | Conli1 | Conli1_GeneCatalog_proteins_20130808.aa | 2 | *Sordariomycetes* |
| | *Dothistroma septosporum* NZE10 v1.0 | Dotse1 | Dotse1fmaa | 2 | *Dothideomycetes* |
| | *Cercospora berteroae* CBS 538.71 | Cerbe1 | Cerbe1_ExternalModels_proteins_2021-11-19.fasta | 2 | *Dothideomycetes* |
| | *Pseudocercospora ulei* ERN8 v1.0 | Pseule1 | Pseule1_GeneCatalog_proteins_20190530.aa | 2 | *Dothideomycetes* |
| | *Pseudocercospora (Mycosphaerella) fijiensis* v2.0 | Mycfi2G | Mycfi2GCaa | 2 | *Dothideomycetes* |
| | *Coniochaeta ligniaria* NRRL 30616 v1.0 | Conlig1 | Conlig1_GeneCatalog_proteins_20140604.aa | 1 | *Sordariomycetes* |
| | *Boothiella tetraspora* CBS 334.67 v1.0 | Bootet1 | Bootet1_GeneCatalog_proteins_20230314.aa | 1 | *Sordariomycetes* |
| | *Cercospora zeae-maydis* v1.0 | Cerzm1 | Cerzm1_GeneCatalog_proteins_20111029.aa | 1 | *Dothideomycetes* |
| | *Cladosporium fulvum* v1.0 | Clafu1 | Clafu1_GeneCatalog_proteins_20110826.aa | 1 | *Dothideomycetes* |
| | *Aspergillus robustus* CBS 428.77 v1.0 | Asprobu1 | Asprobu1_FilteredModels1_proteins_2022-01-21.fasta | 1 | *Eurotiomycetes* |
| | *Podospora inquinata* IMI 300486 v1.0 | Podinq1 | Podinq1_GeneCatalog_proteins_20230206.aa | 1 | *Sordariomycetes* |
| | *Lasiosphaeria ovina* CBS 958.72 v1.0 | Lasov1 | Lasov1_GeneCatalog_proteins_20171021.aa | 1 | *Sordariomycetes* |
| | *Zasmidium cellare* ATCC 36951 v1.0 | Zasce1 | Zasce1_GeneCatalog_proteins_20120808.aa | 1 | *Dothideomycetes* |
| | *Septoria populicola* v1.0 | Seppo1 | Seppo1_GeneCatalog_proteins_20110720.aa | 1 | *Dothideomycetes* |
| | *Septoria musiva* SO2202 v1.0 | Sepmu1 | Sepmu1_GeneCatalog_proteins_20100915.aa | 1 | *Dothideomycetes* |
| | *Sphaerulina populi* pn01.02 v1.0 | Mycpo1 | Mycpo1_FilteredModels1_proteins_2024-03-26.fasta | 1 | *Dothideomycetes* |
| | *Sordaria brevicollis* FGSC1904 v1.0 | Sorbr1 | Sorbr1_GeneCatalog_proteins_20171129.aa | 1 | *Sordariomycetes* |
| | *Teratosphaeria nubilosa* CBS 116005 v1.0 | Ternu1 | Ternu1_GeneCatalog_proteins_20140925.aa | 1 | *Dothideomycetes* |
| ZtKP4 | *Mycosphaerella graminicola* v2.0 | Mycgr3 | Mgraminicolav2All | 7 | *Dothideomycetes* |
| | *Zymoseptoria tritici* IP0323 v2.0 | Zymtr1 | Zymtr1_all_proteins_2023-04-27.fasta | 4 | *Dothideomycetes* |
| UmVKP4 | *Ustilago maydis* KP4 strain | Ustmay* | KP4T_UMV4 (Q90121) | - | *Ustilaginomycetes* |
| | *Aaosphaeria arxii* CBS 175.79 v1.0 | Aaoar1 | Aaoar1_all_proteins_20140429.aa | 3 | *Dothideomycetes* |
| | *Acidothrix acidophila* CBS 136259 v1.0 | Aciaci1 | Aciaci1_all_proteins_20160228.aa | 8 | *Sordariomycetes* |
| | *Alternaria rosae* MPI-PUGE-AT-0040 v1.0 | Altro1 | Altro1_all_proteins_20180320.aa | 4 | *Dothideomycetes* |
| | *Clitocybe* sp. JGI1119283 v1.0 | Cont1 | Cont1119283_primary_alleles_proteins_20190417.aa.fasta | 1 | *Agaricomycetes* |
| | *Aquilomyces patris* DSE4099 v1.0 | Aqupat1 | Aqupat1_all_proteins_20210226.aa | 17 | *Dothideomycetes* |
| | *Aspergillus campestris* IBT 28561 v1.0 | Aspcam1 | Aspcam1_all_proteins_2022-01-21.fasta | 7 | *Eurotiomycetes* |
| | *Mycena filopes* CBHHK001 v1.0 | Mycfil1 | Mycfil1_primary_alleles_proteins_20171109.aa.fasta | 1 | *Agaricomycetes* |
| | *Aspergillus inflatus* CBS 123889 v1.0 | Aspinf1 | Aspinf1_all_proteins_20220407.aa | 1 | *Eurotiomycetes* |
| | *Aspergillus leporis* CBS 151.66 v1.0 | Asplep1 | Asplep1_all_proteins_20161018.aa | 1 | *Eurotiomycetes* |
| | *Aspergillus restrictus* CBS 118.33 v1.0 | Aspres1 | Aspres1_all_proteins_20160629.aa | 1 | *Eurotiomycetes* |
| | *Mycena leptocephala* CBHHK5/15 v1.0 | Myclep1 | Myclep1_secondary_alleles_proteins_20180625.aa.fasta | 1 | *Agaricomycetes* |
| | *Aspergillus zonatus* v1.0 | Aspzo1 | Aspzo1_all_proteins_20121010.aa | 3 | *Eurotiomycetes* |
| | *Astrocystis sublimbata* CBS 130006 v1.0 | Astsub1 | Astsub1_all_proteins_20190325.aa | 1 | *Sordariomycetes* |
| | *Basidiobolus meristosporus* CBS 931.73 v1.0 | Basme2 | Basme2finSC_all_proteins_20150701.aa | 99 | *Basidiobolomycetes* |
| | *Clitocybe* sp. JGI1119283 v1.0 | Cont1 | Cont1119283_all_proteins_20190417.aa | 31 | *Agaricomycetes* |
| | *Mycena vulgaris* CBHHK164 v1.0 | Mycvul1 | Mycvul1_primary_alleles_proteins_20180504.aa.fasta | 1 | *Agaricomycetes* |
| | *Cochliobolus lunatus* m118 v2.0 | Coclu2 | Coclu2_all_proteins_20120521.aa | 2 | *Dothideomycetes* |
| | *Cordyceps* sp. RAO 2017 | CorRAO | CorRAO_2017_1_all_proteins_2021-11-19.fasta | 1 | *Sordariomycetes* |
| | *Epibryaceae* sp. MNA-CCFEE 6094 v1.0 | Epi6094 | Epi6094_1_primary_alleles_proteins_20221014.aa.fasta | 1 | *Eurotiomycetes* |
| | *Ampelomyces quisqualis* HMLAC05119 v1.0 | Ampqui1 | Ampqui1_all_proteins_20160127.aa | 1 | *Dothideomycetes* |
| | *Fusarium acuminatum* 207A_c7_2 v1.0 | Facumi1 | Facumi1_all_proteins_2023-12-28.fasta | 9 | *Sordariomycetes* |
| | *Fusarium acuminatum* F829 v1.0 | Fusacu1 | Fusacu1_all_proteins_2025-05-09.fasta | 9 | *Sordariomycetes* |
| | *Fusarium avenaceum* MPI-SDFR-AT-0044 v1.0 | Fustri1 | Fustri1_all_proteins_20180510.aa | 19 | *Sordariomycetes* |
| | *Fusarium avenaceum* NRRL 54939 v1.0 | Fusave1 | Fusave1_all_proteins_20221012.aa | 10 | *Sordariomycetes* |
| | *Fusarium camptoceras* NRRL 13381 v1.0 | Fusca1 | Fusca1_all_proteins_2023-10-21.fasta | 1 | *Sordariomycetes* |
| | *Fusarium commune* MPI-SDFR-AT-0072 v1.0 | Fusco1 | Fusco1_all_proteins_20180421.aa | 9 | *Sordariomycetes* |
| | *Fusarium concolor* NRRL 13459 v1.0 | Fuscon1 | Fuscon1_all_proteins_20221201.aa | 11 | *Sordariomycetes* |
| | *Fusarium culmorum* UK99 | Fuscu1 | Fuscu1_all_proteins_2021-11-19.fasta | 4 | *Sordariomycetes* |
| | *Fusarium dimerum* 206A_c13_1 v1.0 | Fdimer1 | Fdimer1_all_proteins_2023-12-28.fasta | 15 | *Sordariomycetes* |
| | *Fusarium equiseti* NRRL 66338 v1.0 | Fusequ1 | Fusequ1_all_proteins_20220912.aa | 25 | *Sordariomycetes* |
| | *Fusarium graminearum* PH-1 v2.0 | Fusgr2 | Fusgr2_1_all_proteins_20220618.aa | 31 | *Sordariomycetes* |
| | *Fusarium graminearum* Z3639 v4.0 | Fusgra4 | Fusgra4_all_proteins_20230124.aa | 25 | *Sordariomycetes* |
| | *Fusarium lateritium* NRRL 13622 v1.0 | Fuslat1 | Fuslat1_all_proteins_20230113.aa | 3 | *Sordariomycetes* |
| | *Friedmanniomyces endolithicus* CCFEE 5311 | Frien1 | Frien1_all_proteins_2021-11-19.fasta | 1 | *Dothideomycetes* |
| | *Fusarium nygamai* NRRL 66327 v1.0 | Fusnyg1 | Fusnyg1_all_proteins_20221012.aa | 4 | *Sordariomycetes* |
| | *Fusarium oxysporum* 105B_c14_1 v1.0 | Fusaoxy1 | Fusaoxys1_all_proteins_2023-10-21.fasta | 2 | *Sordariomycetes* |
| | *Setomelanomma holmii* CBS 110217 v1.0 | Setho1 | Setho1_all_proteins_20150424.aa | 1 | *Dothideomycetes* |
| | *Fusarium oxysporum* f. sp. lycopersici 4287 v3.0 | Fusox3 | Fusox3_1_all_proteins_20230106.aa | 22 | *Sordariomycetes* |
| | *Fusarium oxysporum* Fo47 v2.0 | FusoxFo47_2 | FusoxFo47_2_all_proteins_20201023.aa | 18 | *Sordariomycetes* |
| | *Aspergillus candidus* CBS 102.13 v1.0 | Aspcand1 | Aspcand1_all_proteins_2022-01-21.fasta | 3 | *Eurotiomycetes* |
| | *Fusarium proliferatum* ET1 v1.0 | Fusprol1 | Fusprol1_all_proteins_20230113.aa | 2 | *Sordariomycetes* |
| | *Fusarium proliferatum* NRRL 62905 | Fuspr1 | Fuspr1_all_proteins_2021-11-19.fasta | 1 | *Sordariomycetes* |
| | *Fusarium pseudograminearum* NRRL 62612 v1.0 | Fuspse1 | Fuspse1_all_proteins_20221113.aa | 13 | *Sordariomycetes* |
| | *Fusarium* sp. FIESC 12 MPI-CAGE-AA-0113 v1.0 | Fuseq1 | Fuseq1_all_proteins_20180330.aa | 47 | *Sordariomycetes* |
| | *Fusarium sporotirchoides* 7-200 v1.0 | Fusaspor1 | Fusaspor1_all_proteins_2023-12-27.fasta | 3 | *Sordariomycetes* |
| | *Fusarium sporotrichioides* NRRL 3299 v1.0 | Fusspo1 | Fusspo1_all_proteins_20221012.aa | 8 | *Sordariomycetes* |
| | *Fusarium subglutinans* NRRL 66333 v1.0 | Fussub1 | Fussub1_all_proteins_20221115.aa | 13 | *Sordariomycetes* |
| | *Fusarium sulawesiensis* NRRL 66472 v1.0 | Fussul1 | Fussul1_all_proteins_20221115.aa | 7 | *Sordariomycetes* |
| | *Fusarium tricinctum* MPI-SDFR-AT-0068 v1.0 | Fustr1 | Fustr1_all_proteins_20180511.aa | 22 | *Sordariomycetes* |
| | *Fusarium venenatum* MPI-CAGE-CH-0201 v1.0 | Fusven1 | Fusven1_all_proteins_20180620.aa | 19 | *Sordariomycetes* |
| | *Fusarium verticillioides* 7600 v2 | Fusve2 | Fusve2_all_proteins_2022-01-21.fasta | 6 | *Sordariomycetes* |
| | *Geopyxis carbonaria* CBS 144460 v1.0 | Geocar1 | Geocar1_all_proteins_20180620.aa | 18 | *Pezizomycetes* |
| | *Hirsutella rhossiliensis* 206A_T18_1 v2.0 | Hirrh2 | Hirrh2_all_proteins_2024-02-28.fasta | 41 | *Sordariomycetes* |
| | *Hypomyces perniciosus* HP10 | Hypper1 | Hypper1_GeneCatalog_proteins_20210911.aa | 2 | *Sordariomycetes* |
| | *Hypomyces rosellus* CCMJ2808 | Hypros1 | Hypros1_GeneCatalog_proteins_20210912.aa | 8 | *Sordariomycetes* |
| | *Hypoxylon investiens* CIRM-BRFM 1445 v1.0 | Hypinv1 | Hypinv1_all_proteins_20220606.aa | 3 | *Sordariomycetes* |
| | *Hypoxylon* sp. CO27-5 v1.0 | HypCO275_1 | HypCO275_1_all_proteins_20120126.aa | 6 | *Sordariomycetes* |
| | *Hypoxylon* sp. EC38 v3.0 | HypEC38_3 | HypEC38_3_all_proteins_20130613.aa | 5 | *Sordariomycetes* |
| | *Hypoxylon* sp. FL1857 v1.0 | HyFL1857_1 | HyFL1857_1_all_proteins_20190501.aa | 1 | *Sordariomycetes* |
| | *Hypoxylon* sp. NC0597 v1.0 | HyNC0597_1 | HyNC0597_1_all_proteins_20190326.aa | 3 | *Sordariomycetes* |
| | *Hypoxylon trugodes* CBS 135444 v1.0 | Hyptru1 | Hyptru1_all_proteins_20190611.aa | 9 | *Sordariomycetes* |
| | *Inosperma lanatodiscum* 2018PMI007 v1.0 | Inolan1 | Inolan1_all_proteins_20230211.aa | 23 | *Agaricomycetes* |
| | *Inosperma lanatodiscum* 2018PMI007 v1.0 | Inolan1 | Inolan1_primary_alleles_proteins_20230211.aa.fasta | 2 | *Agaricomycetes* |
| | *Isaria fumosorosea* ARSEF 2679 | Corfu1 | Corfu1_all_proteins_2021-11-19.fasta | 4 | *Sordariomycetes* |
| | *Jumillera viridis* CIRM-BRFM 2385 v1.0 | Jumvir1 | Jumvir1_all_proteins_20220412.aa | 3 | *Sordariomycetes* |
| | *Lecanicillium lecanii* RCEF 1005 | Corco1 | Corco1_all_proteins_2021-11-19.fasta | 1 | *Sordariomycetes* |
| | *Lecanicillium muscarium* UNK v1.0 | Lecmu1 | Lecmu1_all_proteins_2024-05-30.fasta | 15 | *Sordariomycetes* |
| | *Leucangium carthusianum* GMNB180 v1.0 | Leuca1 | Leuca1_all_proteins_20190917.aa | 23 | *Pezizomycetes* |
| | *Lizonia empirigonia* CBS 542.76 v1.0 | Lizem1 | Lizem1_all_proteins_20150430.aa | 2 | *Dothideomycetes* |
| | *Marasmius fiardii* PR-910 v1.0 | Marfi1 | Marfi1_all_proteins_20141220.aa | 7 | *Agaricomycetes* |
| | *Metacordyceps chlamydosporia* 111A-C6_1 v1.0 | Mchlamy1 | Mchlamy1_all_proteins_2023-12-28.fasta | 63 | *Sordariomycetes* |
| | *Metacordyceps chlamydosporia* A407-32 v1.0 | Metchl1 | Metchl1_all_proteins_2023-09-14.fasta | 56 | *Sordariomycetes* |
| | *Metarhizium album* ARSEF 1941 | Metal1 | Metal1_all_proteins_2021-11-19.fasta | 3 | *Sordariomycetes* |
| | *Metarhizium anisopliae* ARSEF 549 | Metani1 | Metani1_all_proteins_2021-11-19.fasta | 3 | *Sordariomycetes* |
| | *Metarhizium brunneum* ARSEF3297 | Metbr1 | Metbr1_all_proteins_2021-11-19.fasta | 4 | *Sordariomycetes* |
| | *Metarhizium guizhouense* ARSEF977 | Metgui1 | Metgui1_all_proteins_2021-11-19.fasta | 3 | *Sordariomycetes* |
| | *Metarhizium majus* ARSEF 297 | Metma1 | Metma1_all_proteins_2021-11-19.fasta | 4 | *Sordariomycetes* |
| | *Metarhizium rileyi* RCEF4871 | Metri1 | Metri1_all_proteins_2021-11-19.fasta | 3 | *Sordariomycetes* |
| | *Microthyrium microscopicum* CBS 115976 v1.0 | Micmi1 | Micmi1_all_proteins_20150625.aa | 9 | *Dothideomycetes* |
| | *Mortierella* sp. AD031 | MospAD031_1 | MospAD031_1_all_proteins_2021-11-19.fasta | 1 | *Mortierellomycetes* |
| | *Aspergillus triticus* CBS266.81 v1.0 | Asptr1 | Asptr1_all_proteins_20160629.aa | 1 | *Eurotiomycetes* |
| | *Mortierella* sp. NVP41 | MospNVP41_1 | MospNVP41_1_all_proteins_2021-11-19.fasta | 1 | *Mortierellomycetes* |
| | *Mycena alexandri* CBHHK200 v1.0 | Mycale1 | Mycale1_all_proteins_20171027.aa | 16 | *Agaricomycetes* |
| | *Mycena amicta* BAPX v1.0 | Mycami1 | Mycami1_all_proteins_20171027.aa | 21 | *Agaricomycetes* |
| | *Mycena epipterygia* CBHHK145m v1.0 | Mycepi1 | Mycepi1_all_proteins_20171002.aa | 9 | *Agaricomycetes* |
| | *Mycena filopes* CBHHK001 v1.0 | Mycfil1 | Mycfil1_all_proteins_20171109.aa | 44 | *Agaricomycetes* |
| | *Penicillium discolor* ITEM 17420 v1.0 | Pdis17420_1 | Pdis17420_1_all_proteins_20220801.aa | 1 | *Eurotiomycetes* |
| | *Mycena galericulata* CBHHK162m v1.0 | Mycgale1 | Mycgale1_all_proteins_20180506.aa | 29 | *Agaricomycetes* |
| | *Mycena latifolia* 10383 v1.0 | Myclat1 | Myclat1_all_proteins_20190110.aa | 21 | *Agaricomycetes* |
| | *Mycena leptocephala* CBHHK5/15 v1.0 | Myclep1 | Myclep1_all_proteins_20180625.aa | 30 | *Agaricomycetes* |
| | *Penicillium swiecickii* 182 6C1 v1.0 | Penswi1 | Penswi1_all_proteins_20171024.aa | 1 | *Eurotiomycetes* |
| | *Mycena leptocephala* CBHHK5/15 v1.0 | Myclep1 | Myclep1_primary_alleles_proteins_20180625.aa.fasta | 1 | *Agaricomycetes* |
| | *Mycena maculata* CBHHK188m v1.0 | Mycmac1 | Mycmac1_all_proteins_20180503.aa | 18 | *Agaricomycetes* |
| | *Mycena metata* CBHHK182m v1.0 | Mycmet1 | Mycmet1_all_proteins_20180505.aa | 11 | *Agaricomycetes* |
| | *Mycena polygramma* CBHHK137 v1.0 | Mycpol1 | Mycpol1_all_proteins_20180505.aa | 10 | *Agaricomycetes* |
| | *Mycena rebaudengoi* CBHHK068 v1.0 | Mycreb1 | Mycreb1_all_proteins_20171023.aa | 14 | *Agaricomycetes* |
| | *Mycena sanguinolenta* CBHHK176m v1.0 | Mycsan1 | Mycsan1_all_proteins_20171021.aa | 13 | *Agaricomycetes* |
| | *Mycena vitilis* CBHHK169m v1.0 | Mycvit1 | Mycvit1_all_proteins_20171013.aa | 11 | *Agaricomycetes* |
| | *Mycena vulgaris* CBHHK164 v1.0 | Mycvul1 | Mycvul1_all_proteins_20180504.aa | 17 | *Agaricomycetes* |
| | *Pseudogymnoascus* sp. VKM F-103 | PseVKM1 | PseVKM103_1_all_proteins_2021-11-19.fasta | 1 | *Leotiomycetes* |
| | *Ophiocordyceps australis* Map64 | Ophau1 | Ophau1_all_proteins_2021-11-19.fasta | 3 | *Sordariomycetes* |
| | *Paraphoma fimeti* UNK v1.0 | Parfim1 | Parfim1_all_proteins_20220909.aa | 6 | *Dothideomycetes* |
| | *Penicillium crustosum* ITEM 16721 v1.0 | Pcru16721_1 | Pcru16721_1_all_proteins_20220603.aa | 2 | *Eurotiomycetes* |
| | *Penicillium crustosum* ITEM 16753 v1.0 | Pcru16753_1 | Pcru16753_1_all_proteins_20220601.aa | 2 | *Eurotiomycetes* |
| | *Pseudogymnoascus* sp. VKM F-4520 | PseVKM4520_1 | PseVKM4520_1_all_proteins_2021-11-19.fasta | 1 | *Leotiomycetes* |
| | *Penicillium solitum* IBT 30036 v1.0 | Pensoli1 | Pensoli1_all_proteins_20171003.aa | 2 | *Eurotiomycetes* |
| | *Penicillium* sp. DTU1 v1.0 | Aspcris1 | Aspcris1_all_proteins_20190828.aa | 6 | *Eurotiomycetes* |
| | *Pseudogymnoascus verrucosus* UAMH 10579 | Psever1 | Psever1_all_proteins_2021-11-19.fasta | 1 | *Leotiomycetes* |
| | *Phaeosphaeria poagena* MPI-PUGE-AT-0046c v1.0 | Phapo1 | Phapo1_all_proteins_20180321.aa | 6 | *Dothideomycetes* |
| | *Phoma tracheiphila* IPT5 v1.0 | Photr1 | Photr1_all_proteins_20140424.aa | 6 | *Dothideomycetes* |
| | *Pleosporaceae* sp. PMI_138 v1.0 | PlePMI1 | PlePMI138_1_all_proteins_20190917.aa | 6 | *Dothideomycetes* |
| | *Pochonia chlamydosporia* 170 | Pocchl1 | Pocchl1_all_proteins_2021-11-19.fasta | 4 | *Sordariomycetes* |
| | *Mortierella* sp. GBA35 | MospGBA35_1 | MospGBA35_1_all_proteins_2021-11-19.fasta | 1 | *Mortierellomycetes* |
| | *Physcomitrium patens* | Pp3c23* | PpKP4-2 (Pp3c23_17230) | - | *Plantae* |
| | *Purpureocillium lilacinum* D216_34 v1.0 | Purlil1 | Purlil1_all_proteins_2023-09-14.fasta | 24 | *Sordariomycetes* |
| | *Purpureocillium lilacinum* PLFJ-1 | Purli1 | Purli1_all_proteins_2021-11-19.fasta | 2 | *Sordariomycetes* |
| | *Purpureocillium* sp. UdeA0106 v1.0 | Pursp1 | Pursp1_all_proteins_20160908.aa | 38 | *Sordariomycetes* |
| | *Rasamsonia byssochlamydoides* NRRL 3658 | Rasby1 | Rasby1_GeneCatalog_proteins_20220708.aa | 1 | *Eurotiomycetes* |
| | *Fusarium mangiferae* MRC7560 | Fusma1 | Fusma1_all_proteins_2021-11-19.fasta | 1 | *Sordariomycetes* |
| | *Setophoma terrestris* 302B_T16_1 v1.0 | Setter1 | Setter1_all_proteins_2023-09-14.fasta | 14 | *Dothideomycetes* |
| | *Setophoma terrestris* DSE6109 v1.0 | Sette1 | Sette1_all_proteins_20221202.aa | 7 | *Dothideomycetes* |
| | *Simplicillium lamellicola* cont 1231369 v1.0 | Simlam1 | Simlam1_all_proteins_20220715.aa | 6 | *Sordariomycetes* |
| | *Stagonospora* sp. SRC1lsM3a v1.0 | Stasp1 | Stasp1_all_proteins_20141004.aa | 3 | *Dothideomycetes* |
| | *Thozetella* sp. PMI_491 v2.0 | ThoPMI491_1 | ThoPMI491_1_all_proteins_20140726.aa | 2 | *Sordariomycetes* |
| | *Thyridium* sp. CBS 149464 v1.0 | Phiasp1 | Phiasp1_all_proteins_20220529.aa | 3 | *Sordariomycetes* |
| | *Tolypocladium capitatum* CBS 113982 | Tolca1 | Tolca1_all_proteins_2021-11-19.fasta | 1 | *Sordariomycetes* |
| | *Torrubiella hemipterigena* BCC1449 | Torhe1 | Torhe1_all_proteins_2021-11-19.fasta | 4 | *Sordariomycetes* |
| | *Trematosphaeria pertusa* CBS 122368 v1.0 | Trepe1 | Trepe1_all_proteins_20140415.aa | 3 | *Dothideomycetes* |
| | *Trichoderma aethiopicum* CBS130628 v1.0 | Triaet1 | Triaet1_all_proteins_20210911.aa | 2 | *Sordariomycetes* |
| | *Trichoderma afarasin* CBS130755 v1.0 | Triafa1 | Triafa1_all_proteins_20210911.aa | 6 | *Sordariomycetes* |
| | *Trichoderma aggressivum* SZMC 3109 v1.0 | Triag1 | Triag1_all_proteins_20191108.aa | 2 | *Sordariomycetes* |
| | *Trichoderma alni* SZMC 28000 v1.0 | Trialn1 | Trialn1_all_proteins_20191216.aa | 3 | *Sordariomycetes* |
| | *Trichoderma amazonicum* CBS 121216 v1.0 | Tripleu1 | Tripleu1_all_proteins_20191216.aa | 3 | *Sordariomycetes* |
| | *Trichoderma amazonicum* SZMC 28009 | Triam1 | Triam1_all_proteins_20191216.aa | 1 | *Sordariomycetes* |
| | *Trichoderma barbatum* CBS125733 v1.0 | Tribar1 | Tribar1_all_proteins_20210912.aa | 2 | *Sordariomycetes* |
| | *Trichoderma brevicompactum* SZMC 28004 v1.0 | Tribrev1 | Tribrev1_all_proteins_20191104.aa | 2 | *Sordariomycetes* |
| | *Trichoderma caesareum* CBS 124369 v1.0 | Tricae1 | Tricae1_all_proteins_20210909.aa | 4 | *Sordariomycetes* |
| | *Trichoderma camerunense* CBS 138272 v1.0 | Tricam1 | Tricam1_all_proteins_20210911.aa | 5 | *Sordariomycetes* |
| | *Trichoderma capillare* CBS130629 v1.0 | Tricap1 | Tricap1_all_proteins_20210904.aa | 3 | *Sordariomycetes* |
| | *Trichoderma ceramicum* CBS114576 v1.0 | Tricer1 | Tricer1_all_proteins_20210906.aa | 3 | *Sordariomycetes* |
| | *Trichoderma cerinum* SZMC 28002 v1.0 | Trice1 | Trice1_all_proteins_20191104.aa | 3 | *Sordariomycetes* |
| | *Trichoderma citrinoviride* TUCIM 6016 v4.0 | Trici4 | Trici4_all_proteins_20131015.aa | 11 | *Sordariomycetes* |
| | *Trichoderma compactum* CBS121218 v1.0 | Tricom1 | Tricom1_all_proteins_20210911.aa | 3 | *Sordariomycetes* |
| | *Trichoderma crassum* CBS 336.93 v1.0 | Tricr1 | Tricr1_all_proteins_20210820.aa | 4 | *Sordariomycetes* |
| | *Trichoderma effusum* DAOM 230007 v1.0 | Trieff1 | Trieff1_all_proteins_20191108.aa | 2 | *Sordariomycetes* |
| | *Trichoderma endophyticum* SZMC 28010 v1.0 | Trien1 | Trien1_all_proteins_20191105.aa | 4 | *Sordariomycetes* |
| | *Trichoderma europaeum* CBS 121276 v1.0 | Trieur1 | Trieur1_all_proteins_20210910.aa | 2 | *Sordariomycetes* |
| | *Trichoderma fertile* CBS 339.93 v1.0 | Trifer1 | Trifer1_all_proteins_20210819.aa | 5 | *Sordariomycetes* |
| | *Trichoderma flagellatum* SZMC 28003 v1.0 | Trifla1 | Trifla1_all_proteins_20191109.aa | 4 | *Sordariomycetes* |
| | *Trichoderma ghanense* CBS 259.85 v1.0 | Trigha1 | Trigha1_all_proteins_20210905.aa | 3 | *Sordariomycetes* |
| | *Trichoderma gillesii* CBS130435 v1.0 | Trigil1 | Trigil1_all_proteins_20210909.aa | 2 | *Sordariomycetes* |
| | *Trichoderma gracile* CBS130714 v1.0 | Trigra1 | Trigra1_all_proteins_20210911.aa | 4 | *Sordariomycetes* |
| | *Trichoderma harzianum* CBS 226.95 v1.0 | Triha1 | Triha1_all_proteins_20120308.aa | 19 | *Sordariomycetes* |
| | *Trichoderma harzianum* M10 v1.0 | TriharM10_1 | TriharM10_1_all_proteins_20181017.aa | 29 | *Sordariomycetes* |
| | *Fusarium oxysporum* f. sp. lycopersici 4287 v2 | Fusox2 | Fusox2_all_proteins_2022-01-21.fasta | 1 | *Sordariomycetes* |
| | *Trichoderma harzianum* TR274 v1.0 | Trihar1 | Trihar1_all_proteins_2022-01-21.fasta | 23 | *Sordariomycetes* |
| | *Trichoderma helicolixii* CBS 133499 v1.0 | Triheli | Triheli1_all_proteins_20210820.aa | 6 | *Sordariomycetes* |
| | *Trichoderma inhamatum* CBS 273.78 v1.0 | Triinh1 | Triinh1_all_proteins_20210812.aa | 4 | *Sordariomycetes* |
| | *Trichoderma ivoriense* CBS 125734 v1.0 | Triiv1 | Triiv1_all_proteins_20210820.aa | 3 | *Sordariomycetes* |
| | *Trichoderma konilangbra* CBS 100808 v1.0 | Trikoni1 | Trikoni1_all_proteins_20210909.aa | 3 | *Sordariomycetes* |
| | *Trichoderma lanuginosum* CBS 125718 v1.0 | Trila1 | Trila1_all_proteins_20210819.aa | 4 | *Sordariomycetes* |
| | *Trichoderma longibrachiatum* ATCC 18648 v3.0 | Trilo3 | Trilo3_all_proteins_20130918.aa | 20 | *Sordariomycetes* |
| | *Trichoderma longibrachiatum* CBS 115340 v1.0 | Trior1 | Trior1_all_proteins_20191218.aa | 3 | *Sordariomycetes* |
| | *Trichoderma longibrachiatum* MK1 v1.0 | Trilon1 | Trilon1_all_proteins_20181001.aa | 20 | *Sordariomycetes* |
| | *Trichoderma novae-zelandiae* CBS 472.97 v1.0 | Trinov1 | Trinov1_all_proteins_20210610.aa | 3 | *Sordariomycetes* |
| | *Trichoderma parepimyces* CBS 122769 v1.0 | Tripare1 | Tripare1_all_proteins_20191104.aa | 1 | *Sordariomycetes* |
| | *Fusarium proliferatum* ET1 | Fuspro1 | Fuspro1_all_proteins_2021-11-19.fasta | 1 | *Sordariomycetes* |
| | *Trichoderma reesei* CBS999.97 MAT1-2 | TrireMA12_1 | TrireMAT12_1_all_proteins_2021-11-19.fasta | 1 | *Sordariomycetes* |
| | *Trichoderma rossicum* M135 v1.0 | Triros1 | Triros1_all_proteins_20220328.aa | 6 | *Sordariomycetes* |
| | *Trichoderma sinense* DAOM 230004 v1.0 | Trisin1 | Trisin1_all_proteins_20191108.aa | 3 | *Sordariomycetes* |
| | *Trichoderma* sp. aff breve SZMC 28012 v1.0 | Trsp14807_1 | Trsp14807_all_proteins_20210609.aa | 4 | *Sordariomycetes* |
| | *Trichoderma* sp. aff compactum SZMC 28013 v1.0 | Tri5640_1 | Tri5640_1_all_proteins_20191220.aa | 7 | *Sordariomycetes* |
| | *Trichoderma* sp. aff lixii SZMC 28015 v1.0 | Trich5757_1 | Trich5757_1_all_proteins_20210609.aa | 5 | *Sordariomycetes* |
| | *Trichoderma* sp. aff pleuroticola CBS 121217 v1.0 | Trichpleur1 | Trichpleur1_all_proteins_20210609.aa | 2 | *Sordariomycetes* |
| | *Trichoderma* sp. aff zeloharzianum SZMC 28011 v1.0 | TriTUC5505_1 | TriTUC5505_1_all_proteins_20191216.aa | 6 | *Sordariomycetes* |
| | *Trichoderma* sp. AM6 v1.0 | TrispAM6_1 | TrispAM6_1_all_proteins_2023-10-19.fasta | 6 | *Sordariomycetes* |
| | *Trichoderma spirale* SZMC 28008 v1.0 | Trispi1 | Trispi1_2_all_proteins_20200920.aa | 3 | *Sordariomycetes* |
| | *Trichoderma strictipile* CBS 347.93 v1.0 | Trist1 | Trist1_all_proteins_20191216.aa | 2 | *Sordariomycetes* |
| | *Trichoderma velutinum* CBS 230012 v1.0 | Trivel1 | Trivel1_all_proteins_20191216.aa | 9 | *Sordariomycetes* |
| | *Trichoderma velutinum* SZMC 28001 v1.0 | Triveli | Triveli1_all_proteins_20191216.aa | 4 | *Sordariomycetes* |
| | *Trichoderma harzianum* T22 v1.0 | TriharT | TriharT22_1_all_proteins_20181017.aa | 37 | *Sordariomycetes* |
| | *Trinosporium guianense* CBS132537 v1.0 | Trigu1 | Trigu1_all_proteins_20141217.aa | 8 | *Xylonomycetes* |
| | *Usnea florida* ATCC18376 v1.0 | Usnflo1 | Usnflo1_all_proteins_20160419.aa | 14 | *Lecanoromycetes* |
| | *Whalleya microplaca* YMJ1829 v1.0 | Whamic1 | Whamic1_all_proteins_20190712.aa | 2 | *Sordariomycetes* |
| | *Xylariaceae* sp. FL0255 v1.0 | XylFL0255 | XylFL0255_all_proteins_20190320.aa | 6 | *Sordariomycetes* |
| | *Xylona heveae* TC161 v1.0 | Xylhe1 | Xylhe1_all_proteins_20130827.aa | 2 | *Xylonomycetes* |
| | *Trichoderma reesei* CBS999.97 MAT1-1 | TrireMAT11_1 | TrireMAT11_1_all_proteins_2021-11-19.fasta | 1 | *Sordariomycetes* |
| | *Trichoderma virens* Gv29-8 v2.0 | Trivi2 | Trivi2allaa | 11 | *Sordariomycetes* |

a Abbreviations for species not registered in JGI are indicated by an asterisk.
b Amino acid sequences for KP4T_UMV4 (Q90121) and PpKP4-2 (Pp3c23_17230) were obtained from NCBI (https://www.ncbi.nlm.nih.gov/) and Phytozome v13 (https://phytozome-next.jgi.doe.gov), respectively, and deposited in Figshare.

Supplementary Table S10. Log2 fold change of Defensin -like genes in tomato plants.

| **GeneID** | ***In planta Cf*** **log2 Fold-change** [a] | ***In planta Cf/Hp*** **log2 Fold-change** [a] | **Description** |
|---|---|---|---|
| Solyc01g011435.1 | | | Defensin-like protein 1 (AHRD V3.3 *** A0A1U8FEJ7_CAPAN),Pfam:PF00304 |
| Solyc01g011495.1 | | | Defensin-like protein 1 (AHRD V3.3 *** A0A1U8FEJ7_CAPAN),Pfam:PF00304 |
| Solyc02g078430.3 | | | Defensin-like protein (AHRD V3.3 *-* AT4G21720.3) |
| Solyc04g008470.3 | | | Defensin SD2 (AHRD V3.3 *** A0A2G2ZH52_CAPAN),Pfam:PF00304 |
| Solyc04g072470.4 | | | Defensin Tk-AMP-D6-like (AHRD V3.3 *** A0A1U8GDD3_CAPAN) |
| Solyc05g006813.1 | | | Defensin -like protein 295 (AHRD V3.3 -** XP_015571993.1) |
| Solyc05g012172.1 | | | Defensin -like protein 182 (AHRD V3.3 -** XP_011080983.1) |
| Solyc07g005630.3 | -2.91 | | Defensin -like protein 1 (AHRD V3.3 -** XP_020080865.1),Pfam:PF00304 |
| Solyc07g005655.1 | | | Defensin -like protein 182 (AHRD V3.3 -** XP_011080983.1) |
| Solyc07g006380.3 | | | Defensin-like protein (AHRD V3.3 *** A0A1U8HFN5_CAPAN) |
| Solyc07g007710.4 | | | Defensin protein (AHRD V3.3 *** B1N680_SOLLC),Pfam:PF00304 |
| Solyc07g007730.4 | | | Defensin protein (AHRD V3.3 *** B1N679_SOLLC) |
| Solyc07g007735.1 | | | Defensin protein (AHRD V3.3 *** B1N679_SOLLC),Pfam:PF00304 |
| Solyc07g007740.1 | 3.89 | 5.04 | Defensin -like protein (AHRD V3.3 *** A0A1S3ZY62_TOBAC),Pfam:PF00304 |
| Solyc07g007750.3 | | | Defensin protein (AHRD V3.3 *** B1N678_SOLLC),Pfam:PF00304 |
| Solyc07g007755.1 | -0.90 | -5.15 | Defensin protein (AHRD V3.3 *** B1N681_SOLPI) |
| Solyc07g008373.1 | | | Defensin-like (DEFL) family protein (AHRD V3.3 *** AT4G14276.1) |
| Solyc07g008375.1 | | | Defensin-like (DEFL) family protein (AHRD V3.3 *** AT5G52605.1) |
| Solyc07g008377.1 | | | Defensin-like (DEFL) family protein (AHRD V3.3 *** AT4G14276.1) |
| Solyc07g008980.3 | | | Defensin -like protein 19 (AHRD V3.3 *** A0A2G2ZPK5_CAPAN) |
| Solyc07g009020.2 | | | Defensin -like protein 19 (AHRD V3.3 *** A0A1S4AH79_TOBAC) |
| Solyc07g009030.3 | | 7.53 | Defensin -like protein 19 (AHRD V3.3 *** A0A1S4AH79_TOBAC) |
| Solyc07g009040.3 | | 8.40 | Defensin -like protein 19 (AHRD V3.3 *** A0A1S4AH79_TOBAC) |
| Solyc07g009050.3 | | 7.63 | Defensin -like protein 19 (AHRD V3.3 *** A0A1S4AH79_TOBAC) |
| Solyc07g009060.4 | | 5.33 | Defensin -like protein 19 (AHRD V3.3 *** A0A1S4AH79_TOBAC) |
| Solyc07g009070.4 | | 10.18 | Defensin -like protein 19 (AHRD V3.3 *** A0A1S4AH79_TOBAC) |
| Solyc07g009080.4 | | 8.30 | Defensin -like protein 19 (AHRD V3.3 *** A0A2G2ZPK5_CAPAN) |
| Solyc07g009090.4 | | | Defensin -like protein 19 (AHRD V3.3 *** A0A1S4AH79_TOBAC) |
| Solyc07g009100.3 | | 8.53 | Defensin -like protein 19 (AHRD V3.3 *** A0A1S4AH79_TOBAC) |
| Solyc07g009230.3 | | 3.52 | Defensin-like protein 1 (AHRD V3.3 *** A0A2G2ZS54_CAPAN) |
| Solyc07g009260.3 | -7.96 | | Defensin-like protein 1 (AHRD V3.3 *** A0A2G3AQC7_CAPCH),Pfam:PF00304 |
| Solyc07g009265.1 | | | Defensin-like protein 1 (AHRD V3.3 -** A0A2G3AQC7_CAPCH),Pfam:PF00304 |
| Solyc07g009290.3 | | | Defensin -like protein 1 (AHRD V3.3 *** A0A1U7W6X0_NICSY) |
| Solyc07g016095.1 | | | Defensin-like protein (AHRD V3.3 *** A0A1U8HFN5_CAPAN) |
| Solyc07g016113.1 | | | Defensin-like protein (AHRD V3.3 *** A0A1U8HFN5_CAPAN) |
| Solyc07g016120.3 | | | Defensin-like protein (AHRD V3.3 *** A0A1U8HFN5_CAPAN) |
| Solyc07g017570.2 | | | Defensin -like protein 1 (AHRD V3.3 -** A0A1S3YK59_TOBAC) |
| Solyc07g018145.1 | | | Defensin-like protein 22 (AHRD V3.3 -** A0A2I0VSB5_9ASPA) |
| Solyc07g150108.1 | | | Defensin-like protein (AHRD V3.3 *** A0A1U8HFN5_CAPAN) |
| Solyc07g150139.1 | | | Defensin -like protein 182 (AHRD V3.3 -** XP_011080983.1) |
| Solyc08g060973.1 | | | Defensin -like protein 278 (AHRD V3.3 -** XP_022547472.1) |
| Solyc08g060977.1 | | | Defensin -like protein 278 (AHRD V3.3 -** XP_022547472.1) |
| Solyc09g009725.1 | | | Defensin -like protein 308 (AHRD V3.3 -** XP_006398382.1) |
| Solyc09g074440.3 | | | Defensin-like protein (AHRD V3.3 --* A0A2P4MS75_QUESU),Pfam:PF00304 |
| Solyc10g009040.4 | 1.10 | | Defensin -like protein (AHRD V3.3 *-* AT4G21720.3) |
| Solyc11g006260.2 | | | Defensin-like protein (AHRD V3.3 *** A0A314KXD1_NICAT),Pfam:PF00304 |
| Solyc11g006950.3 | | | Defensin-like protein (AHRD V3.3 *** A0A314LDH2_NICAT),Pfam:PF00304 |
| Solyc11g028040.2 | | | Defensin-like protein (AHRD V3.3 *** A0A1U8HFN5_CAPAN) |
| Solyc11g028060.1 | | | Defensin-like protein (AHRD V3.3 *-* A0A1U8HFN5_CAPAN) |
| Solyc11g028070.2 | | | Defensin-like protein (AHRD V3.3 *** A0A1U8HFN5_CAPAN) |

a Log2 Fold-change values (log2FC ≥ 1, adjusted $P$-value < 0.05) for genes showing significant expression changes. Blank cells indicate genes with padj ≥ 0.05. All DEG datasets are available in Figshare (10.6084/m9.figshare.29491124)
